# Characterization of the Y Chromosome in Newfoundland and Labrador: Evidence of a Founder Effect

**DOI:** 10.1101/2022.08.01.502327

**Authors:** Heather Zurel, Claude Bhérer, Ryan Batten, Margaret E. MacMillan, Sedat Demiriz, Sadra Mirhendi, Edmund Gilbert, Gianpiero L. Cavalleri, Richard A. Leach, Roderick E. M. Scott, Gerald Mugford, Ranjit Randhawa, Alison L. Symington, J. Claiborne Stephens, Michael S. Phillips

**Author notes:** **Corresponding author emails:** Michael S. Phillips.

## Abstract

The population of Newfoundland and Labrador (NL) is largely derived from settlers who migrated primarily from England and Ireland in the 1700-1800s. Previously described as an isolated founder population, based on historical and demographic studies, data on the genetic ancestry of this population remains fragmentary. Here we describe the largest investigation of patrilineal ancestry in NL. To determine the paternal genetic structure of the population, 1,110 Y chromosomes from an NL based cohort were analyzed using 5,761 Y-specific markers. We identified 160 distinct paternal haplotypes, the majority of which (71.4%) belong to the R1b haplogroup. When NL is compared with global reference populations, the haplotype composition and frequencies of the NL paternal lineages primarily resemble the English and Irish ancestral source populations. There is also evidence for genetic contributions from Basque, French, Portuguese, and Spanish fishermen and early settlers that frequented NL. The population structure shows geographical and religious clustering that can be associated with the settlement of ancestral source populations from England and Ireland. For example, the R1b-M222 haplotype, seen in people of Irish descent, is found clustered in the Irish-settled Southeast region of NL. The clustering and expansion of Y haplotypes in conjunction with the geographical and religious clusters illustrate that limited subsequent in-migration, geographic isolation and societal factors have contributed to the genetic substructure of the NL population and its designation as a founder population.

## Introduction

The Canadian province of Newfoundland and Labrador (NL) is home to a unique population that traces its origins to the migration of European communities roughly 300 years ago. The current population is thought to be derived from approximately 25,000 immigrants in the 1700’s and 1800’s who settled in remote coastal communities.^1,2^ These outports were largely isolated from each other, with little settlement in the interior of the island. Communities grew through large families but remained isolated until the 1950s with the advent of paved roads.^1^ The population has continued to expand to its current size of 520,000 and, with the decline of the seafaring economy, is shifting from rural to urban centers.^3^

The main European ancestral source populations that settled NL were from communities around County Waterford and adjacent counties in Ireland and from the counties of Cornwall and Devon as well as fishing ports in Southern England.^1,4^ Following immigration to Newfoundland, English Protestants and Irish Catholics are thought to have remained separated by attending different schools, and rarely inter-married, further isolating these communities.^5,6,7^ Additional European influences that are also thought to have contributed to the genetic landscape of NL are the Portuguese,^8,9^ French^10^ and Highland Scottish.^1^ Norse settlers were present in NL for > 100 years in around 1000 A.D,^11,12,13^ although it appears that they never settled permanently. Also present in NL before, during and after the time of European settlement were Indigenous peoples.^1,7,14^ Since the 1900’s, immigration to Newfoundland has been limited, and the genetic diversity in the province largely traces back to the original European settlers.^15^

Detailed studies of Y chromosome haplotypes have revealed male migration patterns throughout history and led to an understanding of the origins of current human populations.^16,17,18,19,20^ These studies contributed to the development of a standardized Y-DNA phylogenetic tree maintained by the International Society of Genetic Genealogy (ISOGG).^21^ European Y chromosomes are primarily comprised of the haplogroups E, G, I, J, N and R, with the R haplogroup comprising the majority of the Y chromosomes.^22,23,24,25^ While many previous studies are limited by short tandem repeats (STR’s) and/or low resolution single nucleotide polymorphisms (SNPs) panels, ^22,24,25,26,27,28^ they provide information on the composition and frequency of major haplogroups in Europeans.

Supported by studies on the genetic structure of the population^7^ and the presence of numerous rare monogenic disorders,^29^ the population of NL has been described as a founder population. However, information about the haplotypic composition, frequency of Y chromosome variation and ancestral origins across NL is limited. To address these questions, the Y chromosomes of 1,110 individuals from the Newfoundland and Labrador Genome Project (NLGP) cohort^30^ were analyzed in order to: 1) determine the composition and frequency of haplogroups in the paternal lineages; 2) elucidate the population structure of the Y chromosome; 3) understand how the NL population compares with the European ancestral source populations, and 4) identify evidence of founder effects based on haplotypic expansion and regional clustering.

## Materials & Methods

### Newfoundland and Labrador Cohort

Data of the initial 2,500 participants from the NLGP study, a general population cohort from NL, was used for this analysis.^30^ As part of the participants’ self-reported data, we collected information on their religion and birthplace of their ancestors. Each participant provided a saliva sample using the DNA Genotek Oragene OG-600 collection kit (DNA Genotek, Ottawa, Canada). DNA extracted from these samples was genotyped using the Illumina Global Diversity Array (GDA; Illumina, San Diego, CA). Variant calling and quality control (QC) analysis of the genotyping data set was performed using Illumina’s Array Analysis Platform (IAAP) Command Line Interface (CLI) and GTCtoVCF pipeline (github.com/Illumina/GTCtoVCF) (Illumina, San Diego, CA). Out of the 2.1 M variants on the Illumina GDA SNP array, 5,761 SNPs on the male specific portion of the Y chromosome were selected for analysis. QC analysis of the Y chromosome samples determined that 1,110 participants (designated NLGP_1,110_ cohort) had fewer than 200 missing Y chromosome calls (call rate > 96.5%).

### Phylogenetic Reconstruction

The phylogenetic tree was constructed using two different methods: 1) the yHaplo software package,^31^ and, 2) a manual method using maximum parsimony (Supplemental Materials). Although there was concordance between the methods, the manual maximum parsimony approach gave greater haplotype resolution as it enabled the incorporation of SNPs with missing data, singleton SNPs (i.e. variation only observed in a single participant), SNPs without ISOGG designations, and the resolution of phylogenetically inconsistent SNPs (Figure S1, Table S1). Of the 5,761 SNPs, 2,114 were phylogenetically informative (Supplemental Material). The 160 paternal haplotypes, their frequencies, and relationships to each other were used for subsequent analyses. Tag SNPs, associated with specific ISOGG long-form haplotypes, are reported whenever possible to facilitate comparison with the literature.

### Identification of Descendants of NL Founders

A combination of self-reported ethnicity, principal component analysis (PCA) of autosomes, and self-reported birthplaces of paternal ancestors were used to identify individuals whose ancestors descended from early European settlers. Within the NLGP_1,110_ cohort, only 4 participants reported having Indigenous ancestry while 24 participants reported having a mixture of European and Indigenous ancestries (2.6%). Given the limited number of participants with various levels of Indigenous ancestry and the lack of an appropriate reference panel for Indigenous peoples in Eastern North America, we did not investigate the contributions of Y-DNA from Indigenous Peoples in this study. Any participants who were recent immigrants or who reported that their paternal ancestors (up to great-grandfathers) were not from NL were excluded from this analysis. To assess continental ancestry, genotyping data from the autosomes of the NLGP participants was merged with autosome data from the 1000 Genomes project (1KGP3) before running a principal component analysis (PCA) using PLINK 2.0.^32^ Continental ancestry was assigned using the first 5 principal components (PCs) (Figure S2).

To compare NL Y haplotypes with potential ancestral source populations, Y chromosome data from the Irish DNA Atlas^33^ and the People of the British Isles (PoBI)^34^ were analyzed. Of the 812 SNPs overlapped between the NLGP_1,110_ cohort and the PoBI and Irish DNA Atlas data sets, 516 were monomorphic. The remaining 296 SNPs were used to infer major haplogroup frequencies for all 856 Y chromosomes in these data sets. For comparisons with other world populations, the gnomAD allele frequency database^35^ was queried. Since rare variants in European populations are more likely to be population-specific, the 2,114 phylogenetically informative Y-DNA SNPs were inspected for their presence in 7 gnomAD European populations (Basque, Finnish in Finland (FIN), French, British in England and Scotland (GBR), Iberian population in Spain (IBS), Italian, and Toscani in Italia (TSI)). From these, 60 variants were observed in one or two of these populations. Analysis of these 60 variants was extended to all gnomAD populations to assess whether they were informative about potential population ancestry.

### Characterization of the NL Y chromosome population structure

Kinship coefficients were estimated using the KING relationship inference software^36^ implemented in Plink2.^32^ First degree relatives (0.177 < kinship < 0.354) were removed from population analyses. The geographical distribution of the haplotypes was mapped using the birthplace of their most distant paternal ancestor. Regions were assigned based on historical records of settlements and societal and geographic constraints. To evaluate the regional similarities and differences across NL, the province was divided into 5 large regions along the North/South axis and East/West at the point of the Avalon Peninsula isthmus. The St. John’s metropolitan area was designated as a distinct region (Figure 1). These regions were further subdivided into 15 subregions based on major geographical features. The Labrador region, with only 4 participants, was not included in clustering analyses to avoid bias from low numbers. The remaining data set consisted of 831 individuals and 133 haplotypes (designated NL_831_ cohort). Haplotype frequencies were calculated based on geography and religion. Religious affiliation was grouped into 4 categories: Catholic, Protestant, No Religion and Other which includes all other religious/spiritual designations. Notre Dame Bay West, the Northern Peninsula and the West Coast subregions (Figure 1) had less than 25 participants which limited interpretation of this data.

**Figure 1:**
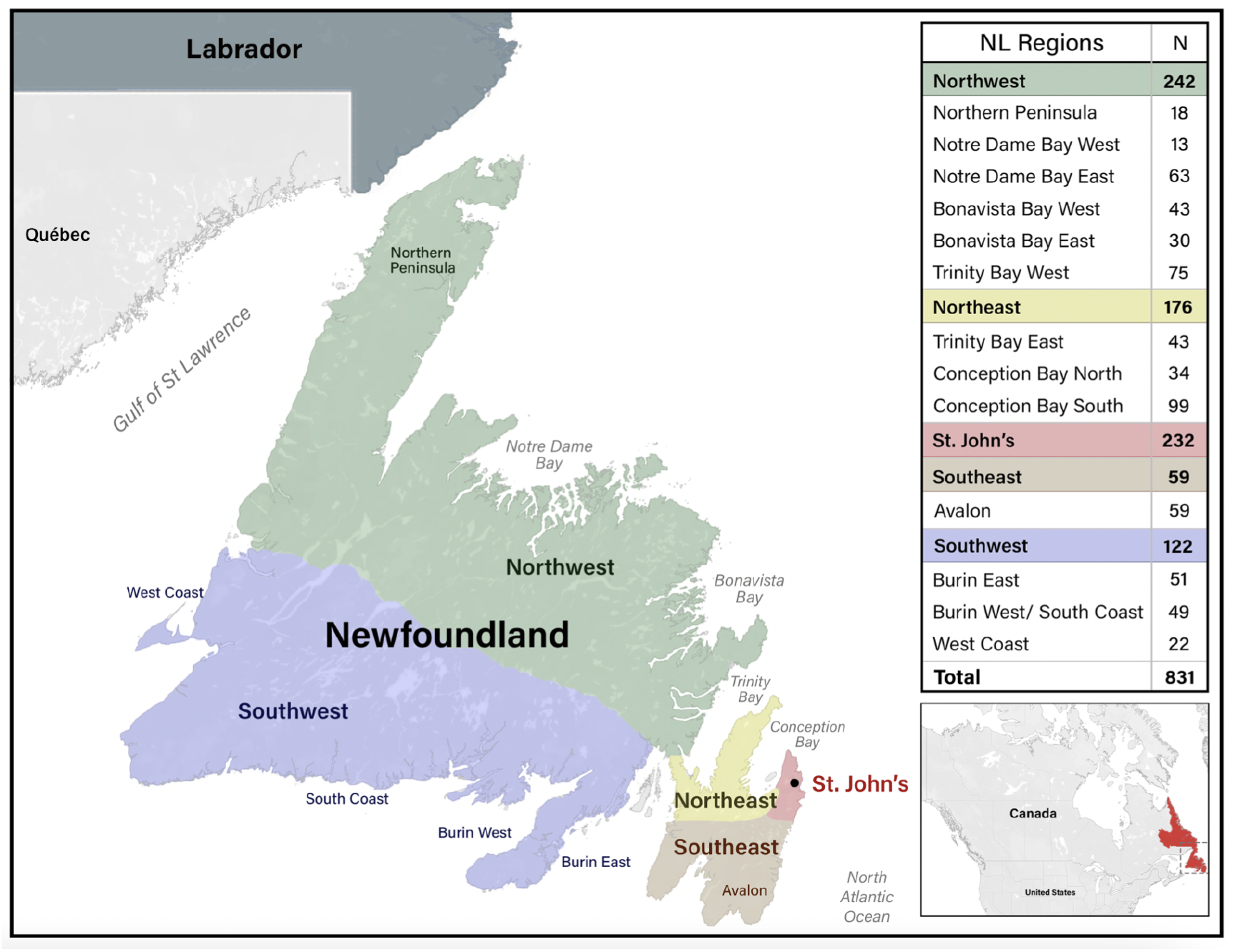
The division of Newfoundland and Labrador into geographical regions and subregions that were used for the analysis of regional Y haplogroup and haplotype frequencies and composition. First degree relatives, recent immigrants and those missing information on the geographic location of paternal ancestors were not included.

## Statistical Methods

### Haplotype Diversity and F_ST_

Haplotype diversity (H), which represents the probability that two randomly sampled haplotypes from a population are different, was calculated for each subregion, as well as the NL_831_ cohort as a whole (n = 831, 133 haplotypes), using the following equation:

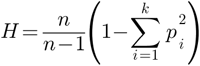

where *n* is the sample size, *k* is the number of distinct haplotypes and *pi* is the frequency of each haplotype.^37^ All distinct haplotypes were used to estimate H for the 5 major regions and 14 remaining subregions. The haplotype diversity term, H, was used to estimate FST of each subregion compared to the NL sample, and pairwise FST was calculated between each pair of regions as described by Slatkin^38^ for haploid genotypes. The pairwise FST values were visualized using multidimensional scaling (MDS) analysis using the cmdscale function in R (v4.1.0).^39^

### Statistical Comparisons

To assess stratification of paternal lineages among the 14 subregions, a PCA was performed based on the variance-covariance matrix of haplotype frequency distribution using the PCAtools R package.^40^ To determine the percentage of variance between populations and groupings, Analysis of MOlecular VAriance (AMOVA) was employed.^41^ The percentage of variation and associated p-values were reported between populations, within populations between subregions and within subregions. AMOVA analyses were conducted using R version 4.1.0 and packages: *ade4* package^42^ obtaining simulated p-values (based on 1000 Monte Carlo simulations). For pairwise comparisons of haplotype composition between regions and subregions, Fisher’s exact test with a simulated p-value (using 1,000 Monte Carlo simulations)^43^ was used with a Benjamini-Hochberg correction.^44^ R version 4.0.3 was used with the *stats* package^39^ to calculate p-values, and results were visualized using the *ggplot2* package.^45^

## Results

### Y Chromosome Structure of the NL population

To construct the NL phylogenetic tree, we used 2,114 phylogenetically informative SNPs in conjunction with the long-form haplotype ISOGG nomenclature to assign 1,110 NL participants to 160 specific haplotypes (Figures 2 & S3, Tables 1, S2 & S3). Seventeen major internal branch points and 7 terminal haplotypes were supported by 20 or more phylogenetically informative SNPs providing confidence in the assembly of the NL Y-DNA tree assembly (Figure S4). The majority of the Y chromosomes in the NLGP_1,110_ cohort occur in the R haplogroup (74.2%, Table 1), predominantly within the R1b haplogroup (71.4% Figure 2, purple). The R1b-S116 haplogroup (R1b1a1b1a1a2) (light purple), comprises 46 distinct haplotypes in the NLGP_1,110_ cohort (43.2%), including R1b-M222 (R1b1a1b1a1a2c1a1a1a1a1), which occurs in 3.1% of the NL Y chromosomes (Table 2). Also present are subclades of major haplogroups I2a, I1a, E1b, R1a, G2a, J2b, J2a in decreasing order of occurrence. The following 7 haplogroups, E1a, H1a, J1a, T1a, O1a, O1b, and Q2a, were detected in single participants, mostly in people who self-reported being born outside of NL.

**Table 1:** List of Major Haplogroups/Haplotypes present in the NLGP_1,110_ cohort.

| ISOGG Subclade<br>or Haplotype | Tag<br>SNP | Haplotype<br>Count | Chromosome<br>Count |
| --- | --- | --- | --- |
| E1a2a1 | CTS10935 | 1 | 1 |
| E1b1b1a1a1 | V12 | 1 | 2 |
| E1b1b1a1b1 | L168 | 3 | 23 |
| E1b1b1a1b2 | L677 | 2 | 5 |
| E1b1b1b | Z827 | 3 | 6 |
| G2a1a1a1a1 | Z6638 | 1 | 1 |
| G2a2a1a2a1a | L166 | 1 | 6 |
| G2a2b1 | M406 | 1 | 1 |
| G2a2b2a1 | L140 | 6 | 11 |
| G2a2b2b1a1 | F872 | 1 | 2 |
| H1a1a4b2 | M2972 | 1 | 1 |
| I1a~ | CTS9857 | 1 | 12 |
| I1a1 | CTS6364 | 4 | 15 |
| I1a2a1a1 | S337 | 3 | 22 |
| I1a2b | S296.1 | 2 | 18 |
| I1a3~ | Z63 | 4 | 29 |
| I2a1a1 | CTS595 | 4 | 18 |
| I2a1a2 | M423 | 2 | 9 |
| I2a1b1 | M223 | 15 | 64 |
| I2a1b2a | L38 | 2 | 10 |
| I2a2 | L596 | 1 | 4 |
| J1a | CTS5368 | 1 | 1 |
| J2a1a | F4326 | 4 | 9 |
| J2b2a | M241 | 2 | 13 |
| O1a1a1 | F446 | 1 | 1 |
| O1b1a1a1a1a1 | M111 | 1 | 1 |
| Q2a1a1a1 | FGC1897 | 1 | 1 |
| R1a1a1 | M417 | 13 | 31 |
| R1b1a1b1 | L23 | 76 | 791 |
| R1b1a1b2a | GG480 | 1 | 1 |
| T1a1a1b2b2b1a1a | CTS6507 | 1 | 1 |
| **Total** |  | **160** | **1,110** |

**Table 2:** Terminal haplotypes and their relative frequencies in the NLGP_1,110_ cohort with a chromosome count of 30 or more.

| ISOGG Haplotype | Tag SNP | Chromosome Count | Frequency in NL (N = 1,110) |
| --- | --- | --- | --- |
| R1b1a1b1a1a | L151 | 65 | 5.9 |
| R1b1a1b1a1a1c1 | S264 | 30 | 2.7 |
| R1b1a1b1a1a1c2b2a1b1a | Z8 | 35 | 3.2 |
| R1b1a1b1a1a2c1a | DF13 | 112 | 10.1 |
| R1b1a1b1a1a2c1a1a1a1a1 | M222 | 34 | 3.1 |
| R1b1a1b1a1a2c1a3a2 | CTS4466 | 33 | 3.0 |
| R1b1a1b1a1a2c1a4a | Z255 | 32 | 2.9 |

**Figure 2.**
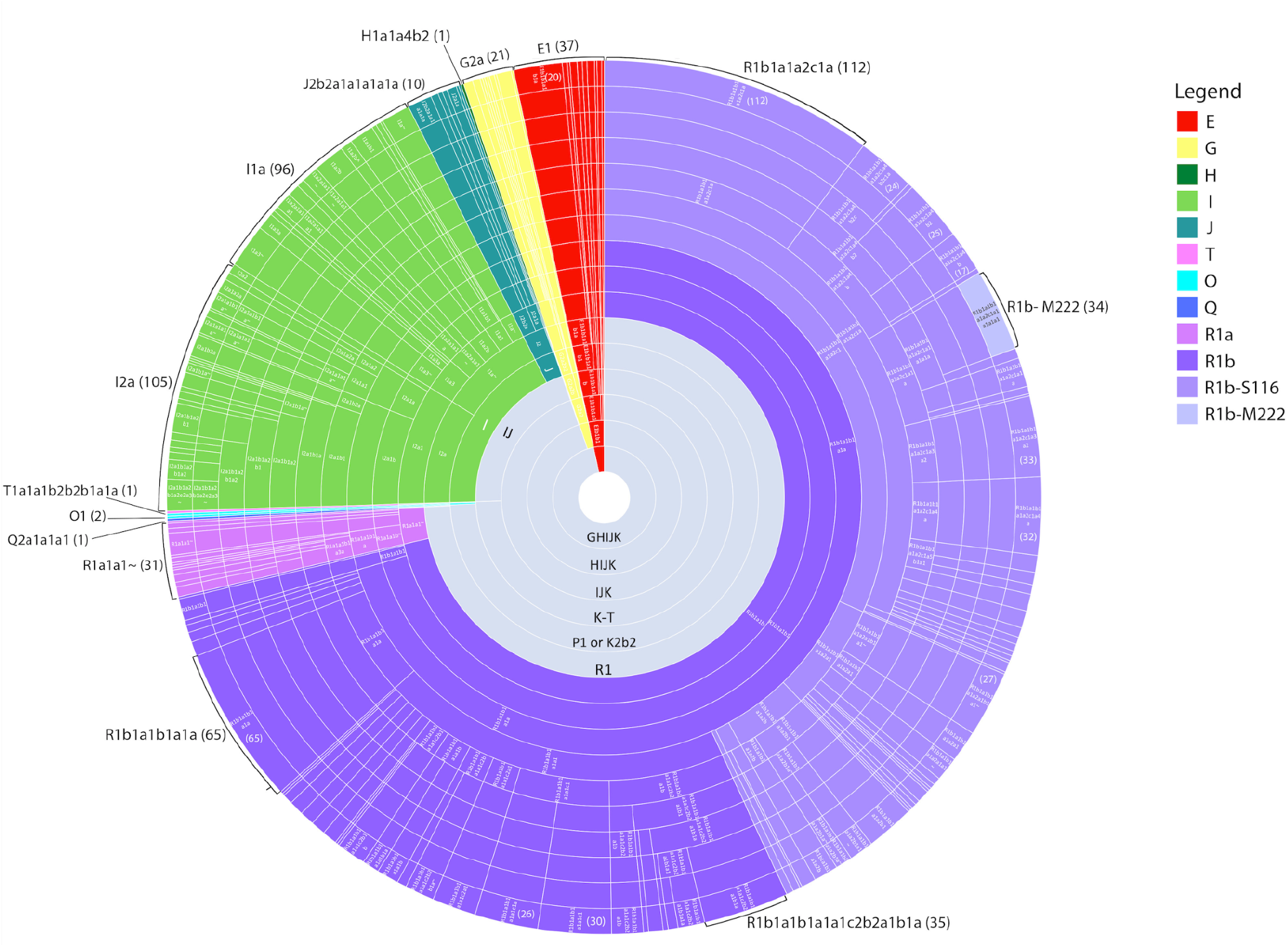
Radial diagram illustrating the proportion of individuals for each haplogroup that are present in the NLGP_1,110_ cohort. The gray inner circles represent mega haplogroups (e.g. K-T, GHIJK) in the phylogenetic tree. Each of the major haplogroups is indicated by a unique color. Each distinct haplotype or haplogroup is separated by a straight white line. Long-form ISOGG nomenclature is provided where possible and can be seen in greater resolution in Figure S3. Each segment of each ring of the radial diagram is proportional to the number of Y chromosomes included in that haplogroup, and segments on the outer ring are proportional to the number of participants belonging to each terminal haplotype. The numbers of participants for select terminal haplotypes are indicated by the brackets on the outer ring.

Review of the NLGP_1,110_ cohort identified 31 terminal haplotypes present in 10 or more individuals (Table S5), and more specifically, 7 terminal haplotypes that were present in 30 or more individuals (Table 2). The largest group was the R1b-DF13 terminal haplotype (R1b1a1b1a1a2c1a) which occurred in 112 individuals.

### Y Chromosome Structure of the NL population

To understand the population structure of NL, we analyzed the haplogroup frequencies by geographical region of the 831 descendants of European founders (NL_831_) (Table 3). Regional differences in R1b haplogroups are observed across NL. The R1b-S116 haplogroup represents greater than 42% of the Y chromosomes except in the Northwest region (28.9%). The frequency of the R1b-M222 haplotype in comparison, is highest in the Southeast region.

**Table 3:** Major haplogroup frequencies detected in the NL_831_ cohort by geographical region. Each NL individual was assigned to a geographical region based on the self-reported birthplace of their most distant paternal ancestor.

| NL Region | Major Haplogroup Frequencies |  |  |  |  |  |  |  |  |  |  | Number of individuals |
| --- | --- | --- | --- | --- | --- | --- | --- | --- | --- | --- | --- | --- |
|  | E1b-M215 | G2a-L31 | I1a-M253 | I2-M438 | J2a-M410 | J2b-M12 | R1a-M198 | R1b-M269 | R1b-S116 | R1b-M222 | T1a-M70 |  |
| Southeast | 5.1 | 1.7 | 10.2 | 3.4 | 0.0 | 0.0 | 0.0 | 23.7 | 47.5 | 8.5 | 0.0 | 59 |
| Northeast | 1.7 | 1.7 | 6.8 | 8.0 | 0.6 | 1.1 | 2.8 | 30.1 | 43.8 | 3.4 | 0.0 | 176 |
| St. John's | 2.6 | 1.3 | 6.9 | 7.3 | 0.0 | 1.3 | 2.6 | 28.9 | 47.0 | 2.2 | 0.0 | 232 |
| Northwest | 2.1 | 1.2 | 10.3 | 15.3 | 1.2 | 0.4 | 2.9 | 34.7 | 28.9 | 2.5 | 0.4 | 242 |
| Southwest | 4.1 | 2.5 | 8.2 | 9.8 | 0.8 | 1.6 | 2.5 | 27.9 | 42.6 | 0.0 | 0.0 | 122 |
| NL Cohort | 2.6 | 1.6 | 8.3 | 9.9 | 0.6 | 1.0 | 2.5 | 30.3 | 40.4 | 2.6 | 0.1 | 831 |

Haplotype diversity based on the frequencies of 133 haplotypes was used to calculate pairwise FST between the 5 major regions. The MDS plot (Figure S5) demonstrates that the major difference among populations (99.4% of the total variance) corresponds to an East-West axis of variation. Results from the AMOVA showed that most of the variation can be explained by the haplotype distribution within subregions (99.3%; p = 0.001; described as “Within populations” in Table S6). A comparison of the Avalon subregion in the East with the Northwest subregions shows significant differences in haplotype composition by Fisher’s exact test (p = 0.02 to 0.001) (Table S7). The East-West geographical haplotype distribution is further supported by PC analysis as represented in the scree plots of the top 5 components (Figure S6).

The coastal communities show distinct patterns in haplogroup frequency and religious affiliation (Figure 3A & 3B). The St. John’s metropolitan area has experienced immigration from many of the coastal communities. As expected, most haplotypes observed in the other regions are present in St. John’s (Figure 3A and S7). In the Northeast region several haplogroup frequencies differ from those observed in the overall NL_831_ cohort suggesting that these subregions might have been settled by immigrants originating from different European regions (Figure 3A). For example, adjacent subregions on the same Peninsula in the Northeast show differential frequencies of I2-M438 ranging from 2.3% in Trinity Bay East to 11.8% in Conception Bay North (Figure 3A). Similarly, in the adjacent Northwest region, Notre Dame Bay East, Bonavista Bay West and Bonavista Bay East subregions show significant differences in haplotype composition when compared with subregions in both the Northeast and Southeast (p = 0.02 - 0.001 by Fisher’s exact test) (Figure 3A, Table S7), further reinforcing the East-West geographical distribution of haplotypes.

**Figure 3:**
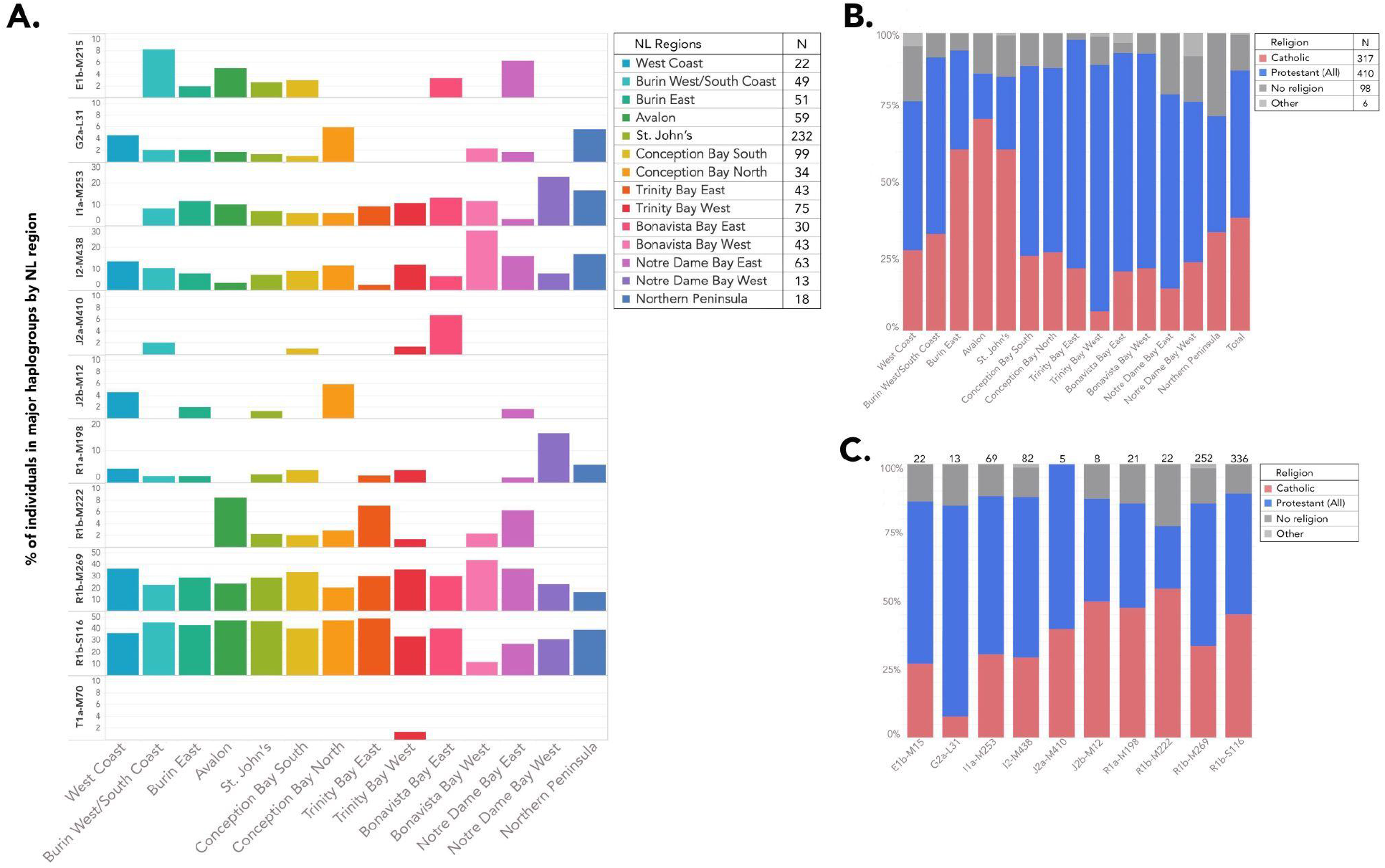
Distribution of major haplogroups by religious affiliation and geographical location. Each NL individual was assigned to a geographical region based on the self-reported birthplace of their most distant paternal ancestor and religion was self-reported. **A:** The frequencies and distribution of the major haplogroups represented as a percentage of the total number of individuals in that region. **B**: The frequencies and distribution of self-reported religious affiliation as a percentage of the total number of individuals in a given region. **C**: The frequencies of self-reported religion represented as a percentage of each major haplogroup. Only one person was reported to have a haplotype of T1a-M70 and therefore is not represented on Figure 3C.

Religion displays some distinctive distribution and frequency patterns across NL as previously described.^1.5.6^ Participants in the Southeast region are predominantly Catholic (>70%) while the Protestant religion predominates in the North (∼ 70%) (Figure 3B). In the NL_831_ cohort, in some regions, some haplotypes appear to be associated with a specific religious affiliation (Figure 3C). For example, the elevated presence of I2-M438 and I1a-M253 haplogroups in the Northwest appears to be mainly associated with Protestant communities (Figure 3C). Similarly, the R1b-M222 haplotype, associated with Irish ancestry,^28^ is observed mainly in Catholic communities (Figure 3C) and is primarily seen in the Avalon Peninsula. As Burin East is the closest subregion of the three to the Avalon subregion and closely resembles the Avalon subregion in terms of religious affiliation, it is noteworthy that R1b-M22, haplotype associated with catholic communities, is absent in this region (Figure 3C).

A PC analysis based on 133 terminal haplotypes was used to visualize the structure of the paternal lineages in the 14 subregions of NL. The first 5 PCs explain >70% of the variation in haplotype frequencies by subregion (Figure S6). A biplot of the first 2 PCs (Figure 4) identifies which haplotypes are the major contributors to the first and second dimensions of PC variation, and shows differentiation between the subregions in the Eastern and Western regions of NL. The R1b-Z255 haplotype, which is mainly observed in Catholics (81%) in the Southeast region is the major contributor to the clustering of the populations in Eastern NL and shows a similar distribution to the R1b-M222 haplotype. R1b-L151 and R1b-Z12 haplotypes which occur mainly in Protestant participants, located in the North Central and West coast regions of NL, appear to be the major haplotypes that are contributing to the clustering of these populations.

**Figure 4:**
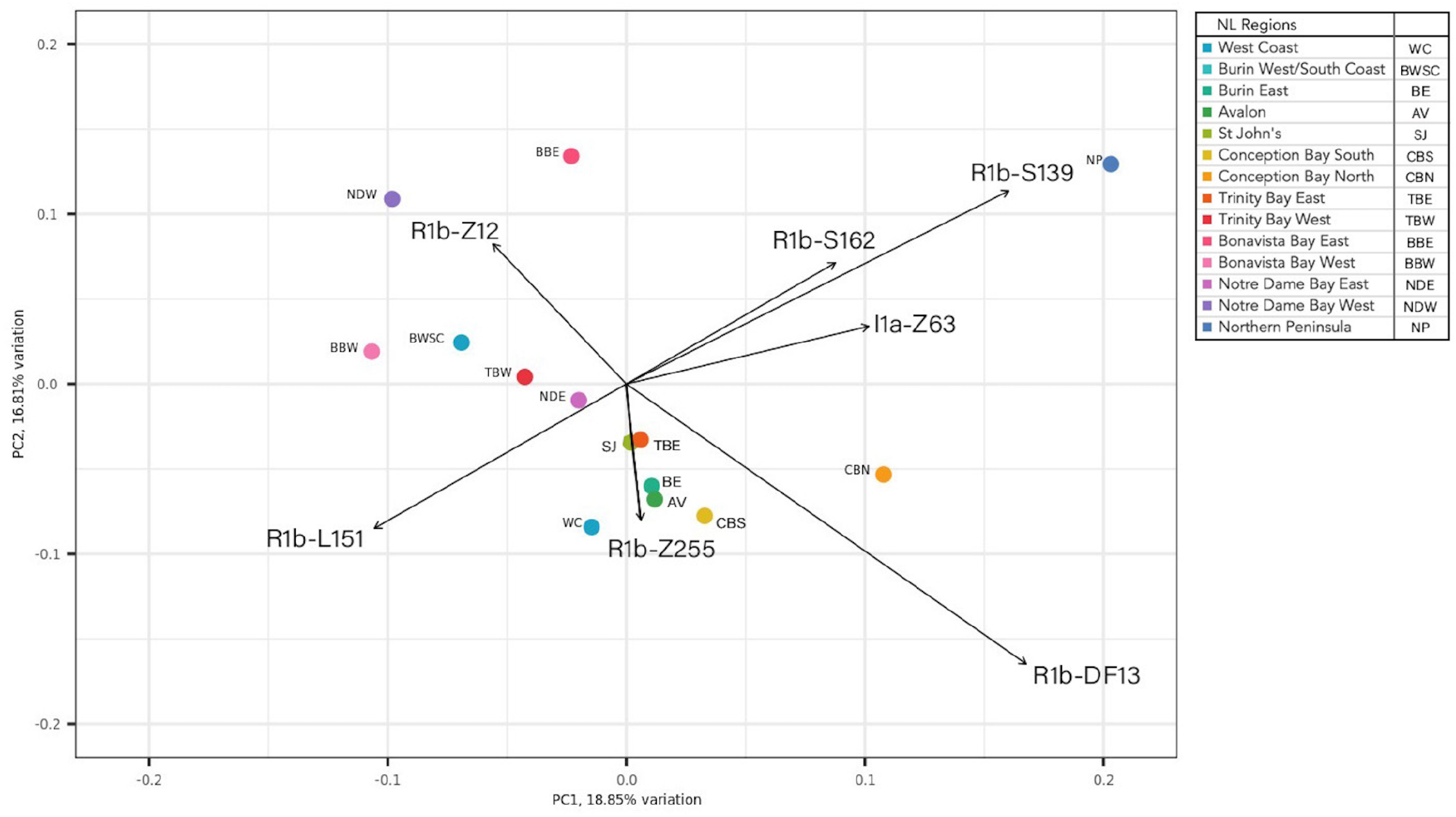
Principal Component (PC) Analysis biplot of the haplotype frequencies in the 14 NL subregions indicating which haplotypes were the major contributors to the top 2 PC axes. Arrows represent the loadings for the 7 haplotypes whose differences in frequencies are the largest contributors to the value of the first two PCs.

### Comparison to Ancestral Source Populations

To infer the origins of the NL paternal lineages, we compared the major Y chromosome haplotypic frequencies found in NL to those of Britain, Ireland and other European source populations using the Irish DNA Atlas,^33^ PoBI,^34^ and the gnomAD allele frequency database^35^ (Table 4). The majority of Y chromosomes within the PoBI and Irish DNA Atlas data sets belong to subclades of the R1b (R1b-M343) haplogroup (Table 4). In addition, analysis of the gnomAD data in combination with data from the PoBI and the Irish DNA Atlas showed evidence of specific haplotypes that could act as markers for the ancestral populations. For example, 3 R1b haplotypes, R1b-U198 (R1b1a1b1a1a1c2a1), R1b-L46 (R1b1a1b1a1a1c2b1a1a), and R1b-Z8 (R1b1a1b1a1a1c2b2a1b1a), seen at relatively high frequency in the NL population, are observed almost exclusively in England in the PoBI and Irish Atlas data sets and likely correspond to English paternal lines. In comparison, R1b-M222 (R1b1a1b1a1a2c1a1a1a1a1) and R1b-Z255 are seen primarily in the Catholic dominated areas of NL, and are almost exclusively seen in Irish populations. The majority of other Y haplotypes seen in the British and Irish populations (subclades of E1b, I1, I2, J2a, J2b, and R1a) were also seen in the NL_831_ cohort, although at different frequencies. For example, R1b-M222 is seen at a frequency of 23.9% in the Irish DNA Atlas data set but only at a frequency of 2.6% in the NL_831_ cohort (Table 4).

**Table 4:**
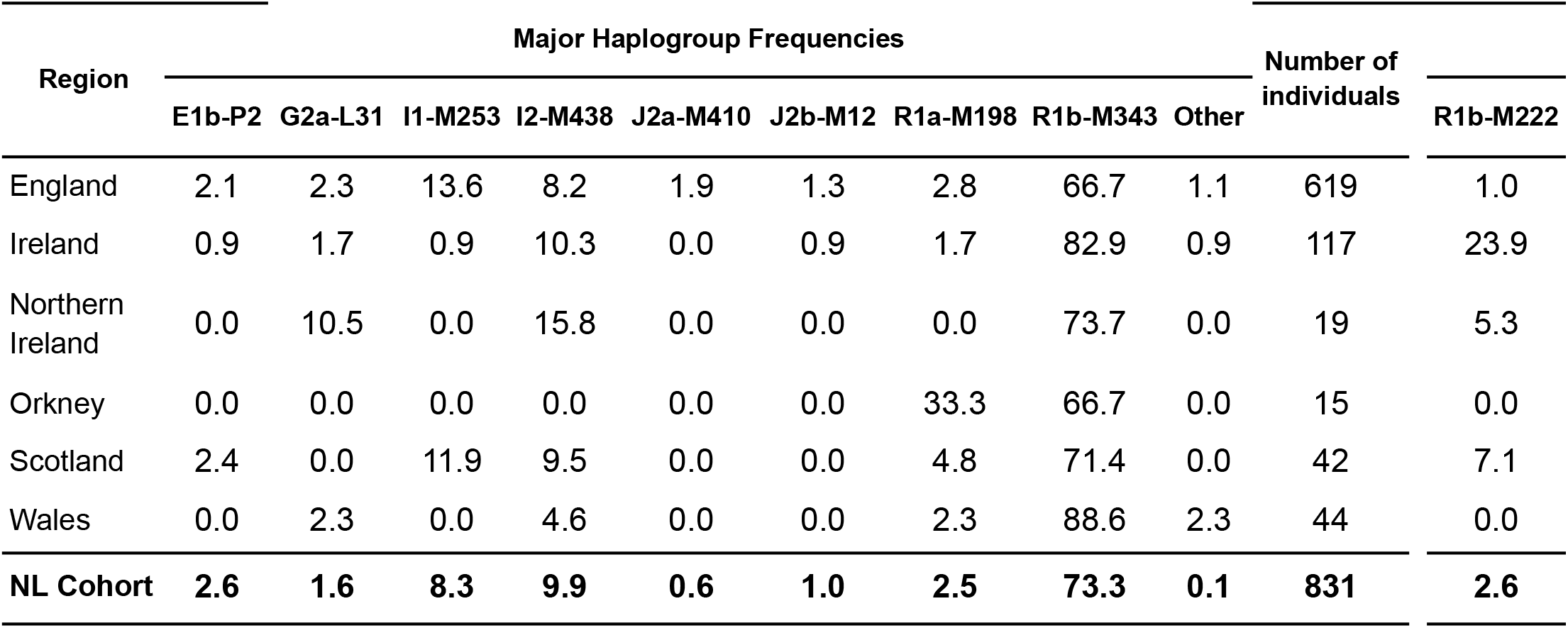
Major haplogroup frequencies detected in the People of the British Isles and Irish DNA Atlas data sets sorted by geographical region. The frequencies of the major haplogroups were calculated and values are represented as a percentage of the total number of individuals in that region. The “Other” column represents haplotypes (F, G, N, Q and T) that were only seen once or twice in the regions (primarily the English population). The R1b-M222 haplotype, included as part of the R1b-M343 haplogroup, is also shown separately as it represents 23.9% of the total Irish Y chromosomes.

| Region | Major Haplogroup Frequencies |  |  |  |  |  |  |  |  | Number of individuals | R1b-M222 |
| --- | --- | --- | --- | --- | --- | --- | --- | --- | --- | --- | --- |
|  | E1b-P2 | G2a-L31 | I1-M253 | I2-M438 | J2a-M410 | J2b-M12 | R1a-M198 | R1b-M343 | Other |  |  |
| England | 2.1 | 2.3 | 13.6 | 8.2 | 1.9 | 1.3 | 2.8 | 66.7 | 1.1 | 619 | 1.0 |
| Ireland | 0.9 | 1.7 | 0.9 | 10.3 | 0.0 | 0.9 | 1.7 | 82.9 | 0.9 | 117 | 23.9 |
| Northern Ireland | 0.0 | 10.5 | 0.0 | 15.8 | 0.0 | 0.0 | 0.0 | 73.7 | 0.0 | 19 | 5.3 |
| Orkney | 0.0 | 0.0 | 0.0 | 0.0 | 0.0 | 0.0 | 33.3 | 66.7 | 0.0 | 15 | 0.0 |
| Scotland | 2.4 | 0.0 | 11.9 | 9.5 | 0.0 | 0.0 | 4.8 | 71.4 | 0.0 | 42 | 7.1 |
| Wales | 0.0 | 2.3 | 0.0 | 4.6 | 0.0 | 0.0 | 2.3 | 88.6 | 2.3 | 44 | 0.0 |
| **NL Cohort** | **2.6** | **1.6** | **8.3** | **9.9** | **0.6** | **1.0** | **2.5** | **73.3** | **0.1** | **831** | **2.6** |

In addition to the English and Irish ancestral populations, several other European populations are known to have fished off the coast of NL.^1^ Under the premise that low-frequency variants are more likely to be population-specific, we looked for rare variants in gnomAD to identify potential source populations for Newfoundland’s founders. We identified 26 distinct haplotypes that were present in one or two of 7 gnomAD European source populations (Basque, Finnish in Finland (FIN), French, British in England and Scotland (GBR), Iberian population in Spain (IBS), Italian, and Toscani in Italia (TSI)). Further analysis of these variants was expanded to all gnomAD populations. Multiple subclades of E1b, and I2a, observed 12 to 64 times in the NLGP_1,110_ cohort, were mainly found in North African and Middle Eastern populations. These haplotypes were associated, at low frequency, with the Southern European populations of France, Iberia, Basque, and Italy. Given that the Y chromosome samples with population designations within gnomAD are limited, further study is required to validate these observations. Analysis of the gnomAD, PoBI and Irish DNA Atlas data revealed several examples of haplotypes that appear to have expanded over time in the NLGP_1,110_ cohort. For example, the R1b-L46 (R1b1a1b1a1a1c2b1a1a) haplotype which is seen in only 2 samples in the English data (PoBI), and not seen in the Irish data, appears in 14 NL participants. Likewise, R1b-Z8 (R1b1a1b1a1a1c2b2a1b1a) is seen in 4 samples in the English data (2 samples in PoBI and 2 samples in gnomAD), but seen in 52 NLGP_1,110_ cohort participants. This observation is suggestive of both possible oversampling of specific haplotypes from England in the settlers who came to NL and possible evidence of local expansion.

## Discussion

This analysis represents the most detailed study of patrilineal ancestry in NL reported to date. In order to characterize the paternal lineages within NL, a high-resolution Y-DNA tree was generated using 2,114 phylogenetically informative markers. Given this level of resolution, this study represents the most detailed study of patrilineal ancestry reported to date for NL. As discussed in the methods, we did not investigate the contributions of Y-DNA from Indigenous Peoples. To address the ancestral contributions of the Indigenous Peoples to the NL Y-DNA tree, a dedicated study of Indigenous Peoples, with and informed by these communities, would be warranted.

The majority of the Y-DNA haplogroups that were identified in the NLGP Y chromosomes appear to be of European origin and reside within the R1b haplogroup (71.4%). The frequency of R1b in the NL cohort is comparable to the English and Irish frequencies observed within the PoBI data, supporting the historical records that immigrants from both these populations settled in NL.

The remaining Y chromosomes in NL, primarily haplogroups I2a1 (9.9%), I1 (8.3%), E1b (2.6%), R1a (2.5%), and J (1.6%), are consistent with haplogroups that are seen in other Western European populations.^23^ Many of these haplogroups have origins in specific European regions, for example R1a and its subclades are commonly observed in Scandinavian populations.^26,49^ It is thought that much of the R1a haplogroup in England and Ireland is associated with Viking settlement.^50^ As the presence of the R1a haplogroup in NL appears to reflect the frequencies seen in these data sets (Table 4), it most likely originated with the English and Irish settlers. The most prevalent of the I haplogroups in the NL_831_ cohort was I2-M438 which comprises I2a, I2b and their respective subclades. Unlike I1, the I2a haplogroup and its subclades are much less frequent in Scandinavia but are reported to comprise 10% of Irish and 6% of Basque Y haplogroups.^24,25,26,28^ While the presence of I2a, which is clustered in the Northwest region of the province (Figure 3, Table 3), is consistent with English-settled communities in NL, it also could be indicative of the presence of Iberian/Basque Y-DNA that originated from Portuguese and Spanish fishermen.^8,9^ Similarly, the J haplogroup, comprising ∼10% of the current Portuguese population^48^ may also have originated in NL with the presence of Portuguese ancestors.

Clustering patterns of haplotypes in specific communities and subregions in the NL_831_ cohort appear to be associated with clustering of self-reported religious affiliation (Figure 3B). The clustering patterns of religion align with historical records of settlements in these regions, primarily Irish Catholics in the Southeast (>70% Catholic) and English Protestants in the Northwest region (70%) (Figure 3B).^1,4,6^ Although religious affiliation can change, our data suggests that self-reported religion in the NL population can be viewed as a surrogate marker for both religion and geographic origin of the participant’s paternal lineage in NL.^4^ The R1b-M222 haplotype, a known Irish haplotype,^22,25,28^ and R1b-Z255, speculated to be of Irish origin,^51^ show localized clustering to known Irish Catholic communities, specifically in the Avalon subregion (Figure 4). Given that early migration of Irish Catholics to NL is well documented, it is likely that these settlers are the primary source of these haplotypes.^1,6^ Although the R1b-M222 haplotype accounts for approximately 25% of Y chromosomes in Ireland,^25^ it is only seen at a frequency of 3% in the NL cohort. This difference is likely because the R1b-M222 haplotype is primarily seen in Northwest Ireland^25^ whereas the historical records suggest that NL was primarily settled by immigrants from Southeast Ireland.^1,25^

NL regional haplotypes exhibit differences along an East to West axis (p = 0.004; Table S6, Figures 3, S5 & S6) and appear to be driven by the ancestral origins of the population with Irish Catholics in the South and East and English Protestants in the North and West. This observation supports the hypothesis that communities were established by settlers who originated from certain communities or specific parishes in Ireland and England and stayed isolated over time. Regions that are directly adjacent to each other, for example Bonavista Bay East and West, only separated by ∼60 km of water, show significant differences in haplogroup composition, supporting the historical records of isolation of coastal communities.^4,5,6^ As expected, the St. John’s Metropolitan region, which has experienced recent immigration from many coastal communities, does not show the same patterns of geographical clustering. The data also indicate that there are Y chromosome contributions from additional European populations such as the Basque, Portuguese, Italian and French. All these observations support the hypothesis that paternal Y haplogroups arrived from distinct European ancestral communities to specific regions within NL.

The unique characteristics of the Y haplotype population structure in NL are indicative of a founder effect. These communities increased over the last 300 years from 25K people to >520K people.^1,2,3^ Evidence of isolation and expansion can be seen by the geographical clustering patterns, and the expansion of certain haplotypes in the NL population (Table 2, Table S4). In fact, 64% of the Y chromosomes in the NLGP_1,110_ cohort show possible evidence of expansion over time as these haplotypes occur in 10 or more people (31 haplotypes in 709 people) (Table 2, Table S4). The expanded haplogroups of R1b-L151 and R1b-Z255 show evidence of regional clustering and expansion as the major haplogroups that differentiate subregions in the East (R1b-Z255) from subregions in the Northwest (R1b-L151) (Figure 4). These observations illustrate that specific ancestral source populations from Europe settled NL, expanded over time, and contributed to the unique clustering patterns seen today.

In summary, NL is an excellent example of a population exhibiting founder effects resulting from limited genetic input followed by generations of geographical and societal isolation which led to regional expansion of specific haplotypes. This data provides a better understanding of the NL genetic population structure which can inform both ancestral history and population structure.

## Acknowledgements

The authors would like to thank all the participants who consented to participate in the Newfoundland and Labrador Genome Project for enabling this research.

This study makes use of data generated by the Irish DNA Atlas Study. A full list of the investigators who contributed to the generation of the data is available from the relevant Irish DNA Atlas papers. The work was in part funded by Science Foundation Ireland Grants 16/RC/3948 and (13/CDA/2223).

This study makes use of data generated by the PoBI project. A full list of the investigators who contributed to the generation of the data is available from the relevant PoBI papers. Part of the funding for the project was provided by the Wellcome Trust under award 088262/Z/09/Z.

## Data Availability

The genotype and sample meta-data from the Newfoundland and Labrador Genome Project (NLGP) are not publicly available due to participant recruitment conditions and consent agreements that protect the privacy of NLGP participants. Reasonable requests for access to the genotyping data should be made to Sequence Bioinformatics. Researchers interested in accessing the NLGP data are encouraged to contact Sequence Bioinformatics.

## Author Contribution Statements

The authors confirm contribution to the paper as follows: study conception and design: H.Z., J.C.S., A.L.S., R.A.L., and M.S.P.; data collection: H.Z., J.C.S., S.M., R.R., R.A.L., and M.S.P.; analysis and interpretation of results: H.Z., J.C.S., C.B., R.B., M.E.M, S.D., S.M., E.G., G.L.C., R.A.L., G.M., R.R., A.L.S., and M.S.P.; draft manuscript preparation: H.Z., J.C.S., C.B., R.B., M.E.M, S.D., S.M., E.G., G.L.C., R.A.L., R.E.M.S., G.M., R.R., A.L.S., and M.S.P.. All authors reviewed the results and approved the final version of the manuscript.

## Ethical Approval

The NL cohort consists of participants recruited with informed consent under a study protocol approved by the Newfoundland and Labrador Health Research Ethics Board (Reference # 2018.243).

## Competing Interests

H.Z., M.E.M, S.D., S.M., R.A.L., G.M., R.R., and M.S.P. are full time employees and shareholders of Sequence BioInformatics, Inc.

R.B., A.L.S. and J.C.S. were paid scientific consultants employed by Sequence BioInformatics, Inc. at the time of this research.

C.B., E.G., and G.L.C. declare no competing interests.

