## Supplemental Information for "Characterization of the Y Chromosome in Newfoundland and Labrador: Evidence of a Founder Effect"

#### Corresponding author emails:

**This PDF file includes:**

Supplementary Methods: Development of the NL Phylogenetic Tree

Figures S1 to S7

Descriptions for Tables S1 to S4 (attached pdf files)
- titles, column headings and descriptions of possible values

Table S5 to S7

### Supplemental Information

#### Supplemental Methods

#### Building the phylogenetic tree

After performing the QC analysis on the genotypes of the NLGP Y chromosomes, as outlined in the methods section of the paper, we identified 5,761 SNPs that could be used to build a NL Y-DNA phylogenetic tree. For this, we employed a multi-stage analysis based on maximum parsimony (described briefly in methods and in more detail below). An overview of the process is shown in Figure S1.

We initially used the yHaplo software to assign haplotypes and construct our Y-DNA tree (https://github.com/23andMe/yhaplo) (Poznik, 2016) from our SNP genotyping dataset. While the older branches of the Y tree were confirmed, the number of phylogenetically informative SNPs available for our study suggested that building the tree manually using maximum parsimony would provide a greater level of detail than could be obtained through yHaplo.

##### Stage I - Identification of clades of the Y-DNA Tree using the highest quality and most complete SNP data subset.

The first stage of the analysis involved identifying all the SNPs that had 1) no missing data and 2) minor allele counts (MAC) of 4 or greater, (1,094 SNPs, Table S1). This subset of SNPs represented the most pristine part of our genotyping dataset. Using this dataset, all cohort participants were then grouped into common SNP genotype clades that contained the same SNP calls. Visual inspection of the clade groupings was performed to identify major binary splits, or “partitions”. Several major partitions were consistently identified. We chose the “M9” branch as a reference point for all clades deduced from our data. Mega haplogroup K-M9 is an ancient clade from which the major haplogroups L, T, N, O, M, P, Q, R, and S are derived (Figure S4).

Stage I analysis was based on the assumptions that most of the 1094 SNPs identified above would 1) create a single binary partition or branch point for our Y chromosomes and 2) be 100% phylogenetically consistent with previous partitions. We observed that 1,072/1,094 of Stage I SNPs fell into this category, creating 100 distinct clades without any suggestion of genotyping error. The 22 remaining Stage I SNPs were identified as phylogenetically inconsistent, meaning that their genotype information conflicted with the phylogenetic inferences drawn from multiple phylogenetically consistent SNPs. Phylogenetic inconsistency can be explained by two processes, genotyping error or genuine homoplasy (the occurrence of parallel or reverse mutations). The investigation and resolution of these inconsistencies is discussed in Stage V below, Table S3. Information on each SNP and the corresponding clade/haplotype is provided in Table S1.

##### Singleton clades

Of the 100 identified clades in Stage I, there were 11 singleton clades. Singleton clades occur when a clade of "*k*" Y chromosomes is further subdivided by a SNP with MAC = *k-1*, splitting one Y chromosome off from the rest. Similarly, clades of 2 or 3 Y chromosomes were also observed in Stage I. Moreover, singleton clades were closely scrutinized and validated during Stage V when all phylogenetically inconsistent SNPs were investigated and resolved (see above and Stage V below). For the purposes of this study, we provisionally recognized these singleton clades until additional information (e.g., ISOGG labeling) could be evaluated. Once validated, the singleton clade is recognized as a singleton haplotype.

##### Stage II - Identification of less frequent clades of the Y-DNA Tree using SNPs with no missing data.

An additional 390/5,761 SNPs were identified as having 1) no missing data and 2) a MAC of 3 or less. Overall, these SNPs helped to further confirm clades identified in Stage I and created refinements that were all phylogenetically consistent with previously established clades. Only 3 of these SNPs were phylogenetically inconsistent, and were set aside for resolution in Stage V. Importantly, many of the 11 singleton clades from Stage I were supported by SNPs with MAC = 1, i.e. multiple singleton SNPs helped to confirm these clades. As SNPs with MAC = 1 are not traditionally used in a typical cladistic analysis, we referred to our analysis method as an "augmented" cladistic analysis. We accepted the singleton haplotypes when they met either of the following criteria:

- C1) When multiple SNPs supported the singleton clade indirectly, as in the discussion below Stage I above, and/or directly as in Stages II, III, or V below;
- C2) When a singleton SNP corresponded to a previously defined ISOGG haplogroup AND the ISOGG haplogroup assignment was consistent with the placement of this singleton haplotype onto our phylogenetic tree.

##### Stage III - SNPs with missing data.

After completion of Stages I and II, 634/5,761 SNPs were identified as having distinct variation in addition to varying degrees of missing data (median 2 missing genotypes). These previous Stages suggested that rates of phylogenetic inconsistency due to genotyping error and/or homoplasy were relatively low (1.7%, 25/1,484 SNPs). As most of the branches in our Y chromosome tree had already been developed in Stages I and II, we were able to place 602/634 SNPs from Stage III onto the existing tree as the missing data was often irrelevant to clade assignment. For example, if the associated clade suggested by the SNP data was deep within M9+ chromosomes, and the missing data was only among M9- chromosomes, there was no difficulty evaluating whether the SNP reinforced an existing, or even defined a new, M9+ clade. In several cases entirely new branches were identified, and these were nonetheless phylogenetically consistent with existing clades.

As expected, Stage III SNPs had a higher frequency of phylogenetic inconsistency (5%, 31/634) than SNPs identified in the previous two Stages of analysis. Additionally, 109/634 (17%) of the SNPs required further review to place on the NL Y-DNA tree as missing data confounded clade assignment. However, even with missing data, most of the missing genotypes could be inferred (imputed) by reliance on downstream SNPs that further refined the haplogroup assignment and many of these SNPs also had ISOGG labels. In addition to providing support for many of the provisional singleton haplotypes, the ISOGG nomenclature often enabled clade imputation in cases of missing data.

##### Stage IV - Monomorphic SNPs

For our Stage IV analysis, 3,643/5,761 SNPs were identified as monomorphic and several also had varying degrees of missing data. Monomorphic SNPs with ISOGG labels (see below) proved to be very valuable for identifying Y chromosome lineages that were absent in our NL cohort (Table S1). Occasionally, a monomorphic SNP had an ISOGG label for a haplogroup that was present in our dataset and these are flagged in Table S1.

##### Stage V - Incorporation of ISOGG Long-Form Nomenclature, Haplotype Assignment, and Reconciliation of Phylogenetically Inconsistent SNPs

In Stage V, the SNPs associated with each clade were manually reviewed and long-form haplogroups were unambiguously assigned to all clades and branch points up to the most highly-defined subclade possible (see detailed phylogenetic tree in Figure S3). When a SNP marker identified a new haplotype that was not reflected in the current ISOGG nomenclature, the new haplotypic extension was annotated with a caret ( ^ ). In several cases, such haplotype extensions were clustered in the same haplogroup. To address this, we annotated the ISOGG long-form root haplogroup with ^, ^^, ^^^, accordingly.

As shown in Table S1, 995 SNPs in Stage I, 302 in Stage II, 515 in Stage III, and 1,988 in Stage IV had an ISOGG designation. Notably, 300 of the 2,114 phylogenetically informative SNPs in Stages I - III have not yet been given an ISOGG label.

At a phylogenetic level, the ISOGG standardized nomenclature proved to be enormously useful in providing additional support for:

1. The confirmation of all identified clades including the singleton haplotypes,
2. The ordering of the various multifurcations arising from maximum parsimony analysis,
3. The confirmation of clades that are absent in the NL cohort, and
4. Resolution of the 56 phylogenetically inconsistent SNPs (see below).

In Stages I-III, 56 SNPs were identified as phylogenetically inconsistent with the NL Y-DNA tree, and were reviewed to assess genotyping error and/or homoplasy (see section on Genotyping Error and Homoplasy below and Figure S1). The subsequent analysis of the 56 SNPs resulted in 53 SNPs being recovered and 3 SNPs that could not be reconciled. The 3 SNPs that could not be reconciled were removed from further analysis. Therefore, in total, 2,114 Y chromosome SNPs were found to be phylogenetically informative and were used for the construction of the phylogenetic tree for the NLGP Y chromosomes

The agreement between the NL Y-DNA tree with the consensus ISOGG phylogenetic tree and long-form labels was outstanding. After review, only 29/3,800 (0.76%) of the ISOGG-labeled SNPs showed any type of discrepancy between our branch placement and the ISOGG labels. Of these 29 SNPs, 9 were phylogenetically informative and consistent with a single clade, 11 were phylogenetically inconsistent with clades emerging from each SNP's resolution either rejected as errors or discrepant with the ISOGG label when accepted, and 9 SNPs were singletons rejected as likely genotyping error (NU in Tables S1 and S3). An additional 22 SNPs were monomorphic in our dataset but whose ISOGG labels nonetheless corresponded to haplogroups present in NL. Upon inspection of each of these, 18 SNPs were missing data for the critical individuals. Since the remaining 4 monomorphic SNPs had discrepancies between our observed clades and their ISOGG labels, we observe a total of 33/3,800 SNPs (0.87%) in which the ISOGG labels were discrepant from our clade assignments.

#### Genotyping Error and Homoplasy

During the process of developing the NL phylogenetic tree (Figure S1**)**, 56 SNPs were identified as phylogenetically inconsistent (Table S3). Phylogenetic inconsistency can be explained by two processes, genotyping error or genuine homoplasy (the occurrence of parallel or reverse mutations). Our resolution of such SNPs assumes that there is an underlying phylogenetic signal, but that this signal has been confounded by secondary changes (homoplasy, H) or by genotyping error (E), or even by both (H+E). These sources of inconsistency can be differentiated, to a degree, by assuming that most genotyping error is random, and that random error is likely to generate a singleton call, that is, differentiating one Y chromosome from the others in an otherwise homogeneous clade. Since our genotypes are strictly binary, an erroneous singleton call amounts to either the generation of a singleton clade (see above), or, more often, to that Y chromosome being assigned to a haplogroup well outside of its consensus placement. Utilizing these principles, we were able to resolve 53/56 phylogenetically inconsistent SNPs identified in our maximum parsimony analysis into 24 that can be attributed to genotyping error (E), 16 that can be explained as homoplasy (H), and 13 that could be explained by homoplasy plus genotyping error (H+E). As mentioned above, the remaining 3 inconsistent SNPs that could not be resolved were not used.

From this process of resolution of the 53 phylogenetically inconsistent SNPs, 13 apparent genotyping errors were identified in our dataset. Another 37 singleton calls resulting from this resolution were rejected as probable genotyping errors, since there was no additional support as singleton clades (see C1 and C2 in Stage II). Similarly, one singleton clade from Stage I and 18 singleton SNPs from Stage II had no additional support and were rejected as potential genotyping errors. Adding to these 69 inferred errors the 14 heterozygous genotypes observed (also assumed to be erroneous), we estimate a genotyping error rate of 0.0035% (83/2,341,277). This illustrates the high quality of our genotyping data.

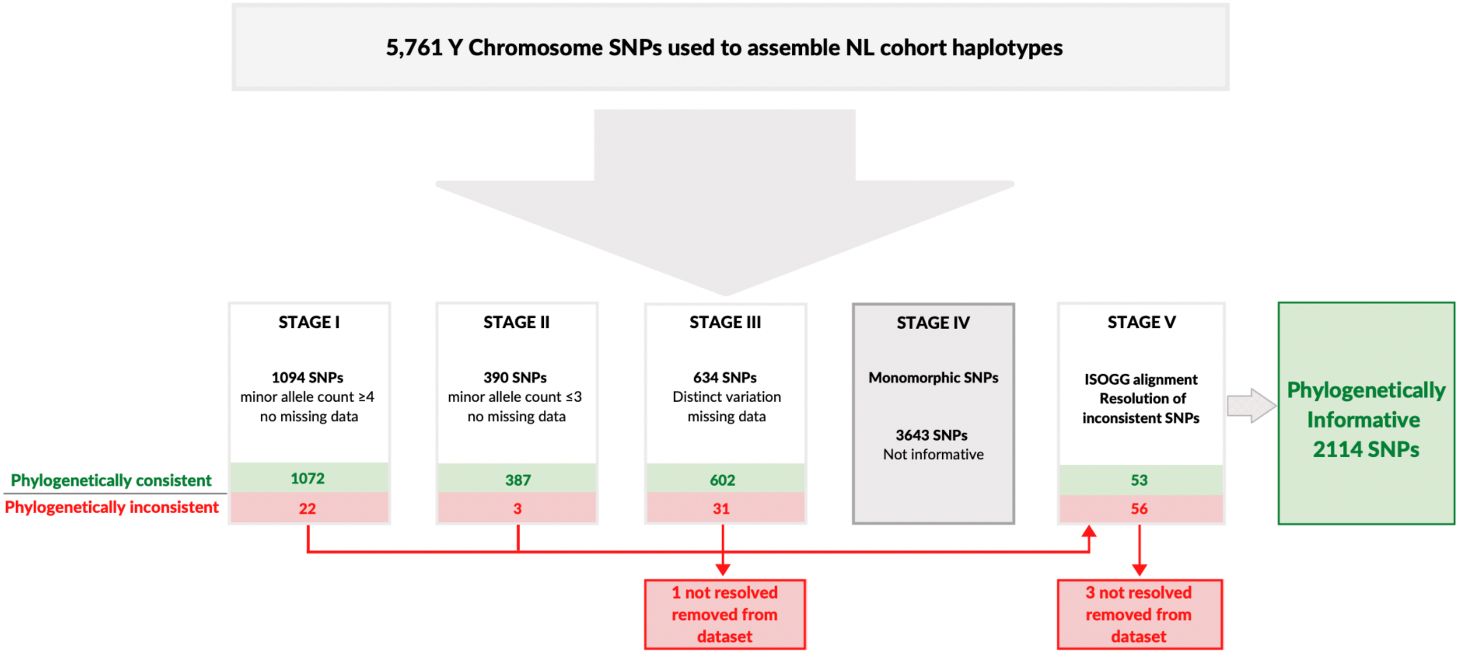

**Figure S1. Analysis Method used for identifying phylogenetically informative Y-DNA SNPs in the NL cohort.** This analysis uses a multi stage maximum parsimony approach to identify both phylogenetically informative and inconsistent markers. In stage V, all SNPs were aligned and compared with known ISOGG haplotypes to finalize the phylogenetic tree and resolve inconsistencies.

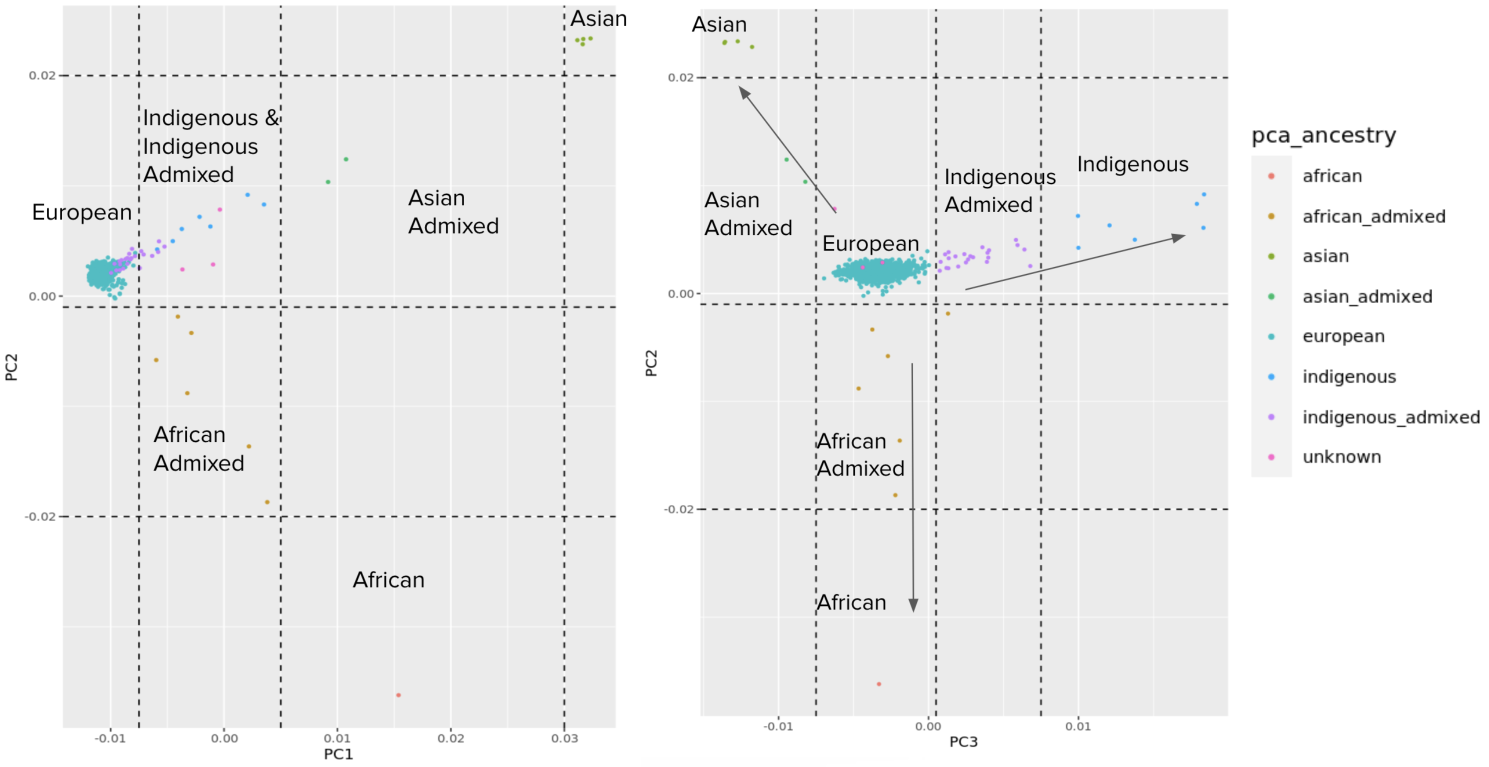

**Figure S2. Principal Component (PC) Analysis of the NL Cohort (males and females).** a) PC1 vs PC2, b) PC3 vs PC2. NLGP samples were merged with 1000 Genomes samples, followed by the PC analysis using Plink2. The first 3 PCs are plotted only for NLGP samples, and coloured by PC-derived continental ancestry. The PC analyses were used to help identify individuals who are descendents of the original settlers to NL. Geographic origins of outliers is based on self-reported ethnicity (i.e. PC1 and PC2 outliers have self-reported African and Asian ancestry or admixture).

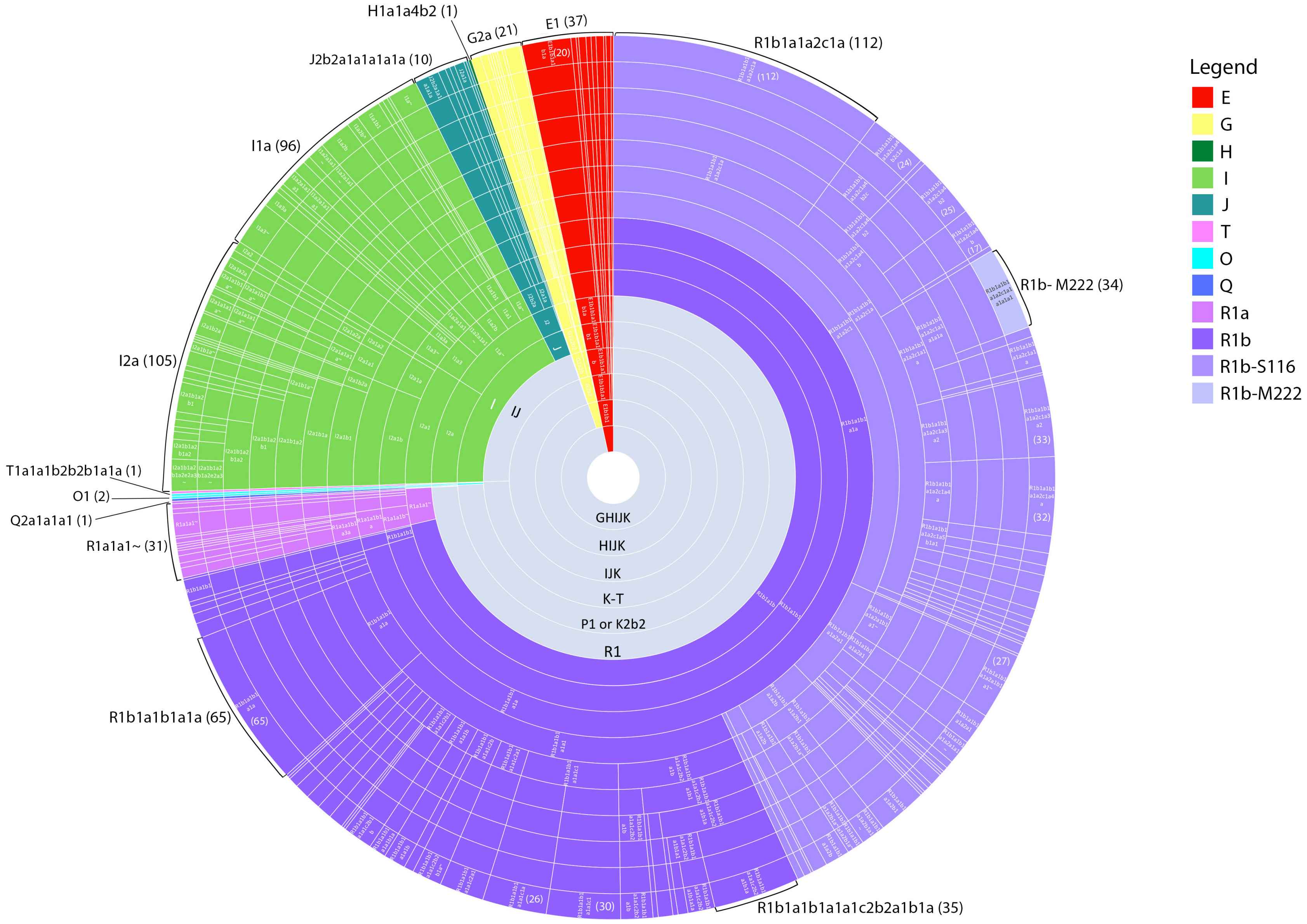

**Figure S3. Radial diagram illustrating the proportion of individuals for each haplogroup that are present in the NLGP$_{1,110}$ cohort.** The grey inner circles represent mega haplogroups (e.g. K-T, GHIJK) in the phylogenetic tree. Each of the major haplogroups are indicated by a unique color. Each distinct haplotype or haplogroup is separated by a straight white line. Long-form ISOGG nomenclature is provided where possible and can be seen in greater resolution in Figure S3. Each segment of each ring of the radial diagram is proportional to the number of Y chromosomes included in that haplogroup, and segments on the outer ring are proportional to the number of participants belonging to each terminal haplotype. The numbers of participants for select terminal haplotypes are indicated by the brackets on the outer ring.

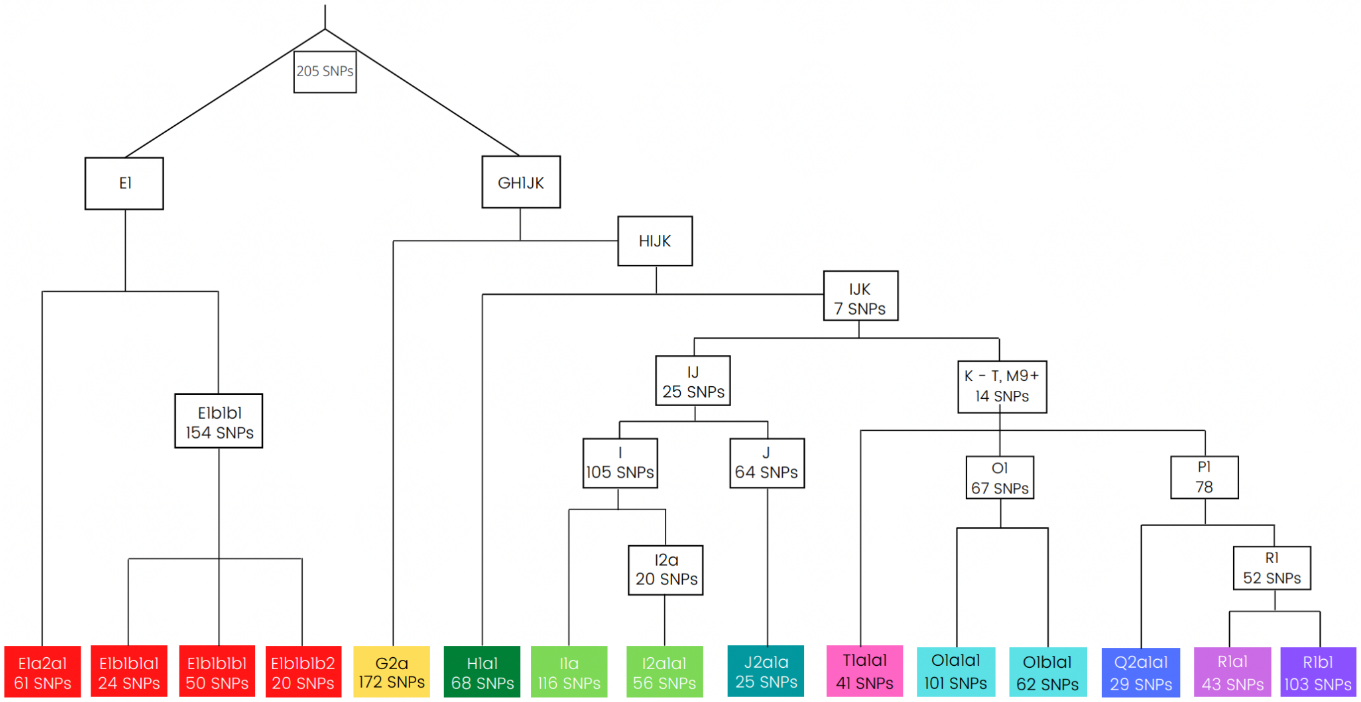

**Figure S4. Schematic diagram illustrating the 17 major internal branch points and 7 terminal haplotypes that are supported by 20 or more phylogenetically informative SNPs.** This figure illustrates the breadth of SNP content that was available to develop the NL cohort Y chromosome phylogenetic tree for the 1,110 participants.

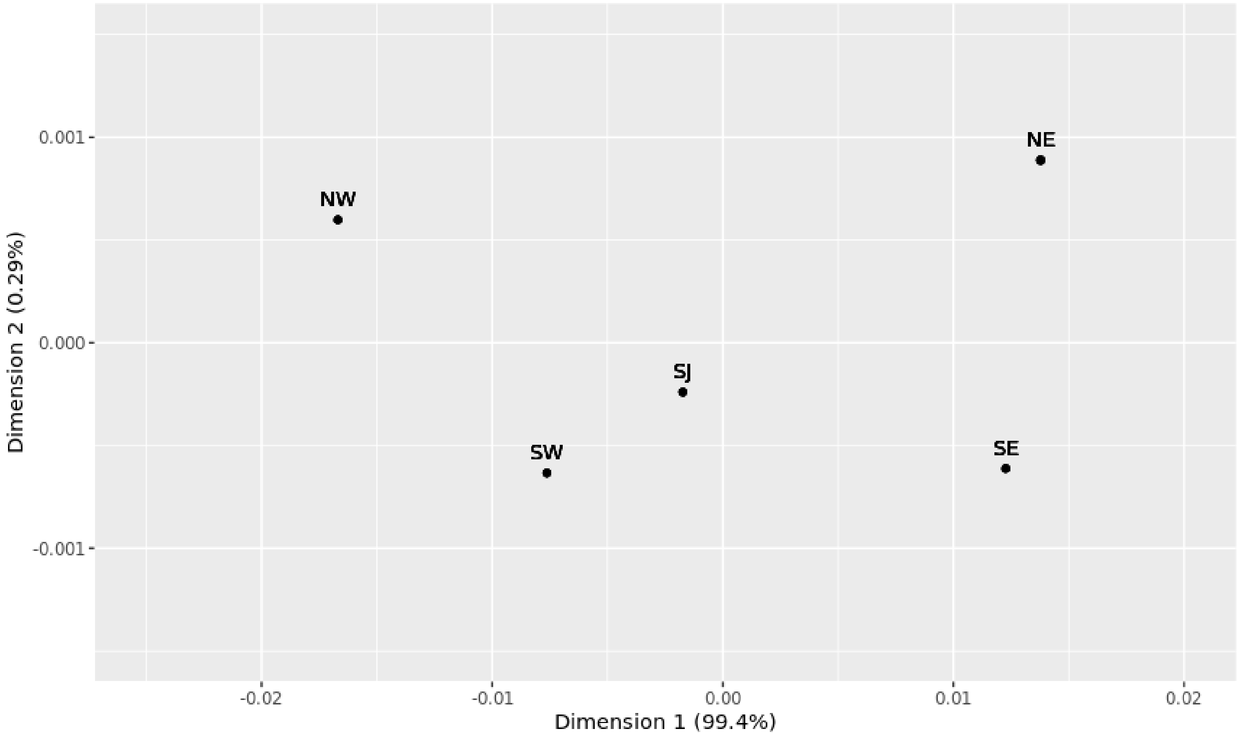

**Figure S5. MDS visualization of pairwise $F_{ST}$ values for 5 major regions in NL.** The plot was generated for the 5 major regions (Southeast (SE), Southwest (SW), St John's (SJ), Northwest (NW) and Northeast (NE)) using $F_{ST}$ values calculated with the 133 haplotypes of the 831 participants that were identified as likely descendants of NL settlers. Values of Dimension 1 were multiplied by -1 so Western regions appear on the left of the plot.

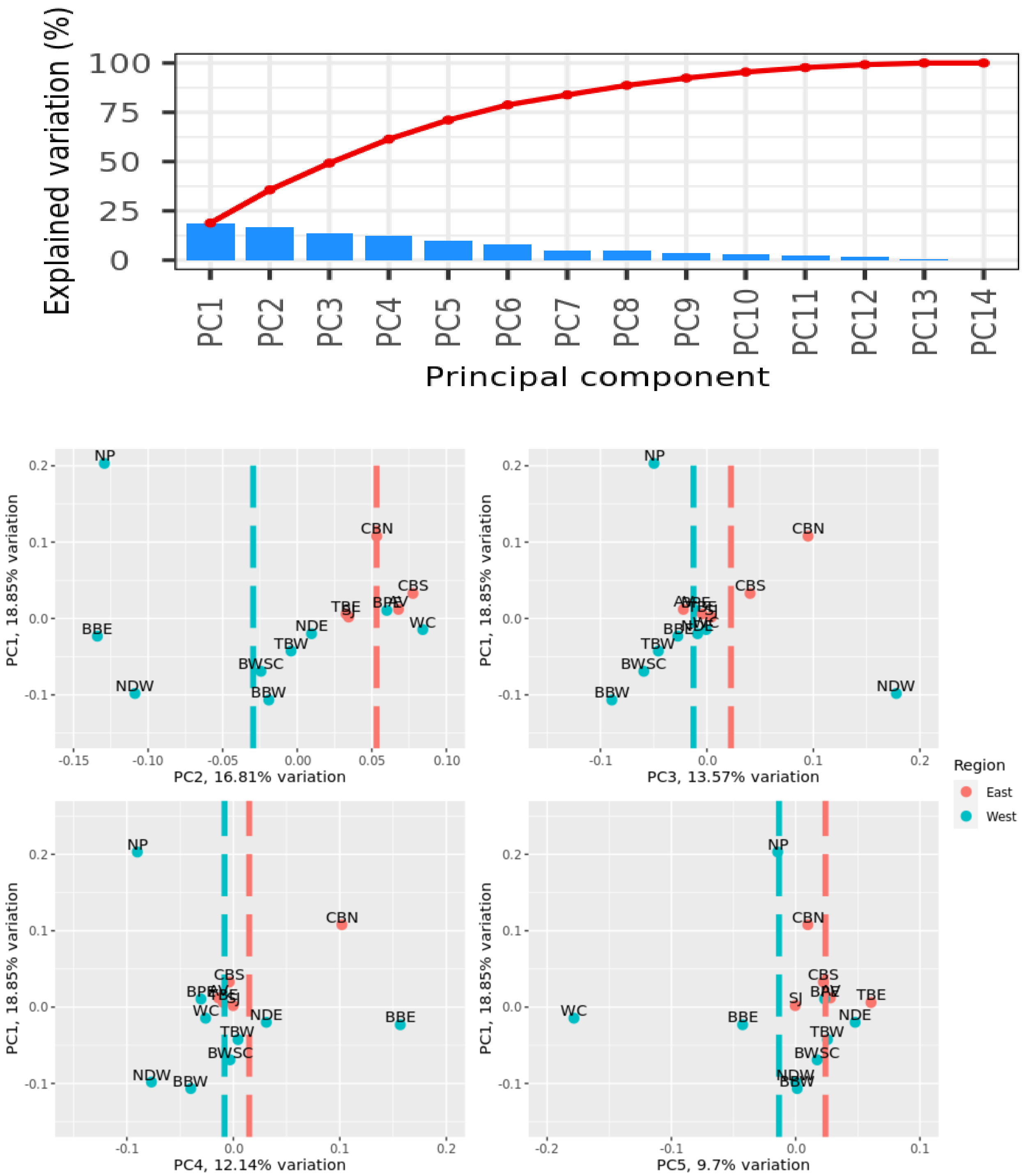

**Figure S6. Principal Components (PC) Analysis of Y chromosome haplotypes in the NL cohort (n = 831).** Top: Scree plot showing proportion of variation explained by each PC. Bottom: PC1 vs PC2-PC5, coloured by major region in NL. Vertical lines indicate the mean value of that PC for Eastern vs Western subregions. Note: values of PC2 in the middle-left plot are multiplied by -1 so the western subregions always appear on the left.

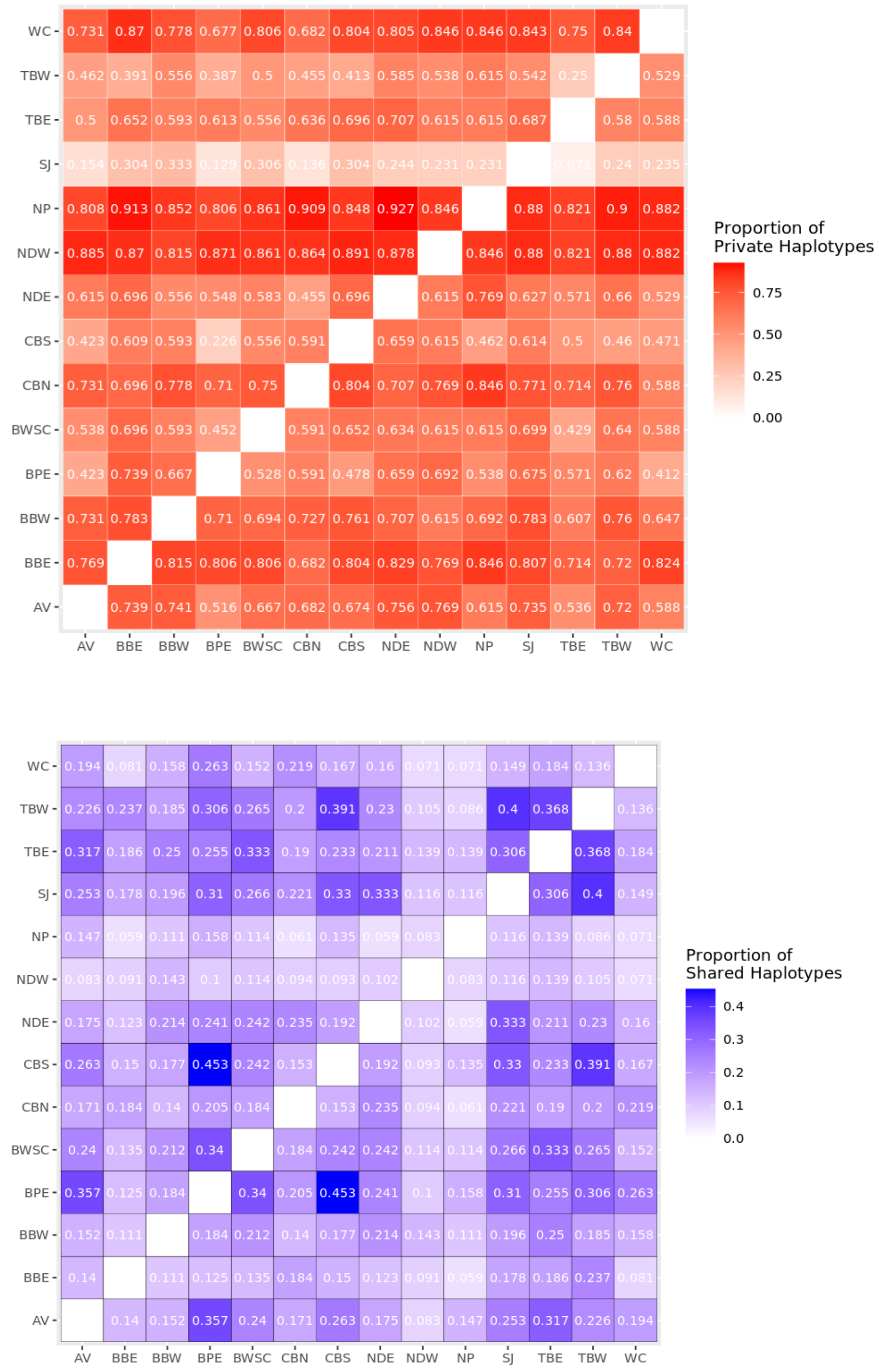

**Figure S7. Heatmaps of the proportion of private and shared haplotypes in 14 subregions in NL.** Private haplotypes are those that are seen in one region, but not in the other region they are being compared to. Shared haplotypes are those that are present in both subregions. As this is a pairwise comparison between two specific subregions, private haplotypes reported here could be present in the remainder of the cohort.

#### Supplemental Tables

*Tables S1-S4 provided in Excel format. Tables S5-S7 are below*

**Table S1. Complete SNP Table.** Results of analysis and phylogenetic reconstruction for all 5761 SNPs. Column headings and a description of possible values are as follows:

- List -- Numerical, ordered by Build 37 Position.
- Stage -- I through V
- M9+/M9- (0, 1,) -- these seven columns are the summary genotype data (counts) for each SNP
- M9? -- M9- are clades splitting off prior to M9; M9+ are those after M9; M9-/M9+ are the M9 split itself; Mono are monomorphic SNPs; phylogenetically inconsistent (PI) SNPs with M9- and M9+ variability are labeled BOTH.
- Clades -- Simple numerical designation of terminal branches or clades, essentially following alphabetical order given by inferred ISOGG Subgroup name. 1 -- 66 for M9- Clades, 1 -- 94 for M9+ Clades. NU: not used, for SNPs in which a single genotyping error was deemed more likely than the clade suggested.
- Prob. -- Problematic SNPs. PI -- phylogenetically inconsistent (56 SNPs); S -- singleton SNPs without additional support, inferred to be genotyping errors; 1H = one heterozygous genotype (13 SNPs), or 2H = heterozygous genotypes (1 SNP), converted to missing data; MD = missing relevant data
- Build 37 Position -- base pair position of SNP according to GRCh37
- Name -- ISOGG tag SNP name
- ISOGG Subgroup Name -- ISOGG longform haplotype name
- Amb. -- ISOGG label ambiguity: A = multiple distinct ISOGG labels assigned; W = wrong for our data

**Table S2. Haplotype Assignments in the NL Cohort.** All clades and their corresponding ISOGG longform haplotype assignments identified in the NL cohort are reported, with counts in the entire cohort (n = 1,110) and for descendants of likely NL founders (n = 831).

**Table S3. Phylogenetically Inconsistent SNPs.** Summary data on phylogenetically inconsistent SNPs and their resolution. Column headings and a description of possible values are as follows:

- List -- Numerical, ordered by GRCh37 Position.
- Stage -- I through V
- M9+/M9- (0, 1, .) -- these seven columns are the genotype data for each SNP
- M9? -- as in Supplementary Table 1, M9- are clades splitting off prior to M9; M9+ are those after M9. BOTH, in the first line summary for each SNP, indicates that variation on both sides of M9 was evaluated for potential clades.
- Clades -- as in Supplementary Table 1, simple numerical designation of terminal branches or clades, essentially following alphabetical order given by inferred ISOGG Subgroup name. 1 -- 66 for M9- Clades, 1 -- 94 for M9+ Clades.

- Res. -- PI SNP Resolution: PI (phylogenetically inconsistent) in each SNP's summary line, followed by one or more of: H (homoplasy), 1E or 2E (1 or 2 genotyping errors predicted), S (flag for singleton calls), MD = missing relevant data, BAD (as above in Clades column).
- Build 37 Position -- base pair position of SNP according to GRCh37 (suppressed in extensions).
- Name -- SNP's identifier (suppressed in extensions).
- ISOGG Subgroup Name -- Subgroup Name assigned each SNP, when available
- Amb. -- ISOGG label ambiguity: A for multiple distinct ISOGG labels available, W for wrong for our data (see Supplementary Table 3).
- List -- Numerical, ordered by GRCh37 Position. Should be identical to the first column.

**Table S4. Reconstructed phylogenetic tree of NLGP participants.** All clades and branch points are indicated, as well as the number of participants in each clade and the ISOGG longform haplogroup associated with each clade.

**Table S5. Haplotypes present in more than 10 participants in the NL cohort (n = 1,110).**

Terminal haplotypes with chromosome count > 10

| M9- | | | M9+ | | |
|---|---|---|---|---|---|
| **Clade** | **ISOGG** | **Count** | **Clade** | **ISOGG** | **Count** |
| 4 | E1b1b1a1b1a | 20 | 18 | R1b1a1b1a1a | 65 |
| 22 | I1a~ | 12 | 28 | R1b1a1b1a1a1c1 | 30 |
| 24 | I1a1b1 | 10 | 29 | R1b1a1b1a1a1c1a | 26 |
| 31 | I1a2b | 13 | 30 | R1b1a1b1a1a1c2a1 | 18 |
| 33 | I1a3~ | 18 | 33 | R1b1a1b1a1a1c2b1b | 12 |
| 49 | I2a1b1a2b1 | 12 | 37 | R1b1a1b1a1a1c2b2a1b | 13 |
| 56 | I2a1b1a2b1a2e2a3~ | 13 | 39 | R1b1a1b1a1a1c2b2a1b1a | 35 |
| 66 | J2b2a1a1a1a1a | 10 | 42 | R1b1a1b1a1a1c2b2a1b1a1a | 11 |
| | | | 45 | R1b1a1b1a1a1c2b2b1a~ | 12 |
| | | | 47 | R1b1a1b1a1a2a1 | 13 |
| | | | 50 | R1b1a1b1a1a2a1a1~ | 10 |
| | | | 58 | R1b1a1b1a1a2a1b1a1~ | 27 |
| | | | 60 | R1b1a1b1a1a2b | 10 |
| | | | 61 | R1b1a1b1a1a2b1 | 20 |
| | | | 63 | R1b1a1b1a1a2b1a1~ | 10 |
| | | | 66 | R1b1a1b1a1a2c1a | 112 |
| | | | 70 | R1b1a1b1a1a2c1a1a | 12 |
| | | | 74 | R1b1a1b1a1a2c1a1a1a1a1 | 34 |
| | | | 78 | R1b1a1b1a1a2c1a3a2 | 33 |
| | | | 79 | R1b1a1b1a1a2c1a4a | 32 |
| | | | 80 | R1b1a1b1a1a2c1a4b | 17 |
| | | | 82 | R1b1a1b1a1a2c1a4b2 | 25 |
| | | | 84 | R1b1a1b1a1a2c1a4b2c1a | 24 |

**Table S6. AMOVA calculated with an equidistant matrix of 133 terminal haplotype frequencies in 14 subregions in NL.** Proportion of variance explained by populations, subregions, and within subregions is reported, as well as p-values based on 999 simulations (n = 831). The Eastern region contains all subregions in the Northeast, southeast, and St. John's regions and the Western regions include all subregions within the Northwest and Southwest regions. Subpopulations/subregions within these groupings are defined in Figure 1.

| Regional Groupings | % total variance | p-value |
|---|---|---|
| Regional analysis (East & West) | | |
| Within populations | 99.3 | 0.001 |
| Among populations within groups | 0.40 | 0.004 |
| Among groups | 0.26 | 0.019 |
| Regional analysis (NE, SE, SJ, NW, SW) | | |
| Within populations | 99.43 | 0.002 |
| Among populations within groups | 0.44 | 0.016 |
| Among groups | 0.13 | 0.137 |
| Subregional analysis (14 subregions) | | |
| Within populations | 99.45 | 0.005 |
| Among populations | 0.55 | 0.005 |

**Table S7. Fisher's exact test with 1000 simulations (n = 831).**

| NL Region | | Northern Peninsula | Notre Dame Bay West | Notre Dame Bay East | Bonavista Bay West | Bonavista Bay East | Trinity Bay West | Trinity Bay East | Conception Bay South | Conception Bay North | St. John's | Avalon | Burin East | Burin West South Coast |
|---|---|---|---|---|---|---|---|---|---|---|---|---|---|---|
| | **N** | 18 | 13 | 63 | 43 | 30 | 75 | 43 | 99 | 34 | 232 | 59 | 51 | 49 |
| West Coast | 22 | 0.083 | 0.441 | 0.209 | 0.045 | 0.024 | 0.515 | 0.321 | 0.129 | 0.199 | 0.698 | 0.049 | 0.558 | 0.317 |
| Burin West South Coast | 49 | 0.086 | 0.500 | 0.354 | 0.109 | 0.100 | 0.852 | 0.726 | 0.001 | 0.011 | 0.033 | 0.013 | 0.661 | |
| Burin East | 51 | 0.153 | 0.140 | 0.129 | 0.043 | 0.001 | 0.987 | 0.584 | 0.902 | 0.091 | 0.722 | 0.161 | | |
| Avalon | 59 | 0.009 | 0.005 | 0.010 | 0.001 | 0.001 | 0.018 | 0.103 | 0.012 | 0.002 | 0.933 | | | |
| St. John's | 232 | 0.099 | 0.084 | 0.013 | 0.002 | 0.001 | 0.075 | 0.965 | 0.128 | 0.471 | | | | |
| Conception Bay North | 34 | 0.049 | 0.096 | 0.423 | 0.002 | 0.051 | 0.167 | 0.138 | 0.049 | | | | | |
| Conception Bay South | 99 | 0.008 | 0.041 | 0.002 | 0.001 | 0.001 | 0.055 | 0.037 | | | | | | |
| Trinity Bay East | 43 | 0.144 | 0.350 | 0.705 | 0.070 | 0.009 | 0.996 | | | | | | | |
| Trinity Bay West | 75 | 0.147 | 0.363 | 0.312 | 0.458 | 0.123 | | | | | | | | |
| Bonavista Bay East | 30 | 0.045 | 0.333 | 0.006 | 0.001 | | | | | | | | | |
| Bonavista Bay West | 43 | 0.002 | 0.108 | 0.029 | | | | | | | | | | |
| Notre Dame Bay East | 63 | 0.028 | 0.271 | | | | | | | | | | | |
| Notre Dame Bay West | 13 | 0.239 | | | | | | | | | | | | |

Using Benjamini-Hochberg correction, anything less than 0.027 is considered statistically significant
