## Supplementary material for "Characterization of the Y Chromosome in Newfoundland and Labrador: Evidence of a Founder Effect": Table S1

| List | Stage | M9+ | | | M9- | | | | | | Build 37 | | ISOGG | | | |
|---|---|---|---|---|---|---|---|---|---|---|---|---|---|---|---|---|
| List | Stage | 0 | 1 | . | 0 | 1 | . | M9? | Clade(s) | Prob. | Position | Name | Subgroup Name | Amb. | List | |
| 2274 | IV | 827 | 0 | 0 | 283 | 0 | 0 | Mono | Mono | | 14814060 | M171 | A (Notes) | | 2274 | |
| 4788 | IV | 827 | 0 | 0 | 283 | 0 | 0 | Mono | Mono | | 21868776 | M59 | A (Notes) | | 4788 | |
| 164 | IV | 827 | 0 | 0 | 282 | 0 | 1 | Mono | Mono | | 2872583 | V190 | A0 | | 164 | NP |
| 530 | IV | 827 | 0 | 0 | 283 | 0 | 0 | Mono | Mono | | 6788191 | V148 | A0 | | 530 | |
| 551 | IV | 826 | 0 | 1 | 283 | 0 | 0 | Mono | Mono | | 6843151 | V149 | A0 | | 551 | |
| 556 | IV | 827 | 0 | 0 | 283 | 0 | 0 | Mono | Mono | | 6846615 | V176 | A0 | | 556 | |
| 574 | IV | 827 | 0 | 0 | 283 | 0 | 0 | Mono | Mono | | 6863229 | V177 | A0 | | 574 | |
| 910 | IV | 827 | 0 | 0 | 283 | 0 | 0 | Mono | Mono | | 7592457 | V223 | A0 | | 910 | |
| 914 | IV | 827 | 0 | 0 | 283 | 0 | 0 | Mono | Mono | | 7596315 | V225 | A0 | | 914 | |
| 926 | IV | 827 | 0 | 0 | 283 | 0 | 0 | Mono | Mono | | 7606633 | V229 | A0 | | 926 | |
| 950 | IV | 826 | 0 | 1 | 282 | 0 | 1 | Mono | Mono | | 7654353 | V239 | A0 | | 950 | |
| 2354 | IV | 827 | 0 | 0 | 283 | 0 | 0 | Mono | Mono | | 14905147 | L995 | A0 | | 2354 | |
| 2635 | IV | 827 | 0 | 0 | 283 | 0 | 0 | Mono | Mono | | 15436806 | L998 | A0 | | 2635 | |
| 5335 | IV | 827 | 0 | 0 | 283 | 0 | 0 | Mono | Mono | | 23117567 | V196 | A0 | | 5335 | |
| 557 | IV | 819 | 0 | 8 | 281 | 0 | 2 | Mono | Mono | | 6847637 | L1098 | A0-T | | 557 | |
| 1934 | IV | 826 | 0 | 1 | 283 | 0 | 0 | Mono | Mono | | 13888035 | L1116 | A0-T | | 1934 | |
| 588 | IV | 827 | 0 | 0 | 283 | 0 | 0 | Mono | Mono | | 6877771 | L1284 | A00 | | 588 | |
| 1600 | IV | 827 | 0 | 0 | 283 | 0 | 0 | Mono | Mono | | 8852748 | AF4 | A00 | | 1600 | |
| 2017 | IV | 827 | 0 | 0 | 283 | 0 | 0 | Mono | Mono | | 14126859 | AF8 | A00 | | 2017 | |
| 2938 | IV | 814 | 0 | 13 | 275 | 0 | 8 | Mono | Mono | | 16193008 | AF9 | A00 | | 2938 | |
| 2457 | IV | 827 | 0 | 0 | 283 | 0 | 0 | Mono | Mono | | 15024040 | L996 | A0a | | 2457 | |
| 4451 | IV | 827 | 0 | 0 | 283 | 0 | 0 | Mono | Mono | | 21288728 | L1011 | A0a | | 4451 | |
| 4821 | IV | 821 | 0 | 6 | 279 | 0 | 4 | Mono | Mono | | 21889673 | L1015 | A0a | | 4821 | |
| 648 | IV | 813 | 0 | 14 | 281 | 0 | 2 | Mono | Mono | | 6932829 | V159 | A0a1 | | 648 | |
| 2015 | IV | 827 | 0 | 0 | 283 | 0 | 0 | Mono | Mono | | 14125645 | L1075 | A0a1 | | 2015 | |
| 564 | IV | 826 | 0 | 1 | 283 | 0 | 0 | Mono | Mono | | 6851720 | V151 | A0a1a | | 564 | |
| 2075 | IV | 827 | 0 | 0 | 283 | 0 | 0 | Mono | Mono | | 14249732 | V181 | A0a1a | | 2075 | |
| 653 | IV | 827 | 0 | 0 | 283 | 0 | 0 | Mono | Mono | | 6933056 | L1289 | A0a1b | | 653 | |
| 2350 | IV | 827 | 0 | 0 | 283 | 0 | 0 | Mono | Mono | | 14902417 | L994 | A0a2 | | 2350 | |
| 3453 | IV | 827 | 0 | 0 | 282 | 0 | 1 | Mono | Mono | | 17419925 | L1007 | A0a2 | | 3453 | |
| 601 | IV | 827 | 0 | 0 | 283 | 0 | 0 | Mono | Mono | | 6894291 | L1038 | A0b | | 601 | |
| 848 | IV | 827 | 0 | 0 | 283 | 0 | 0 | Mono | Mono | | 7400052 | L1039 | A0b | | 848 | |
| 913 | IV | 827 | 0 | 0 | 283 | 0 | 0 | Mono | Mono | | 7596228 | L1040 | A0b | | 913 | |
| 46 | IV | 827 | 0 | 0 | 283 | 0 | 0 | Mono | Mono | | 2710154 | P305 | A1 | | 46 | |
| 955 | IV | 827 | 0 | 0 | 283 | 0 | 0 | Mono | Mono | | 7658712 | V241 | A1 | | 955 | |
| 1162 | IV | 825 | 0 | 2 | 282 | 0 | 1 | Mono | Mono | | 8087059 | L986 | A1 | | 1162 | |
| 2930 | IV | 827 | 0 | 0 | 283 | 0 | 0 | Mono | Mono | | 16181147 | V161.2 | A1 | A | 2930 | |
| 3175 | IV | 827 | 0 | 0 | 283 | 0 | 0 | Mono | Mono | | 16773792 | L1004 | A1 | | 3175 | |
| 3553 | IV | 827 | 0 | 0 | 283 | 0 | 0 | Mono | Mono | | 17645335 | V250 | A1 | | 3553 | |
| 3713 | IV | 827 | 0 | 0 | 282 | 0 | 1 | Mono | Mono | | 17947672 | V168 | A1 | | 3713 | |
| 161 | IV | 825 | 0 | 2 | 283 | 0 | 0 | Mono | Mono | | 2868499 | V215 | A1a | | 161 | |
| 165 | IV | 827 | 0 | 0 | 283 | 0 | 0 | Mono | Mono | | 2872610 | V191 | A1a | | 165 | |
| 598 | IV | 827 | 0 | 0 | 283 | 0 | 0 | Mono | Mono | | 6890869 | V14 | A1a | | 598 | |
| 603 | IV | 827 | 0 | 0 | 283 | 0 | 0 | Mono | Mono | | 6894625 | V25 | A1a | | 603 | |
| 604 | IV | 827 | 0 | 0 | 283 | 0 | 0 | Mono | Mono | | 6894819 | V26 | A1a | | 604 | |
| 610 | IV | 827 | 0 | 0 | 283 | 0 | 0 | Mono | Mono | | 6905202 | V28 | A1a | | 610 | |
| 647 | IV | 826 | 0 | 1 | 283 | 0 | 0 | Mono | Mono | | 6932824 | V30 | A1a | A | 647 | |
| 2939 | IV | 827 | 0 | 0 | 283 | 0 | 0 | Mono | Mono | | 16193088 | V57 | A1a | | 2939 | |
| 4679 | IV | 827 | 0 | 0 | 283 | 0 | 0 | Mono | Mono | | 21739754 | M31 | A1a | | 4679 | |
| 160 | IV | 827 | 0 | 0 | 282 | 0 | 1 | Mono | Mono | | 2868496 | V214 | A1a~ | | 160 | |
| 908 | IV | 825 | 0 | 2 | 283 | 0 | 0 | Mono | Mono | | 7589303 | V221 | A1b | | 908 | |
| 1281 | IV | 824 | 0 | 3 | 238 | 0 | 45 | Mono | Mono | | 8349696 | V1820 | A1b | | 1281 | |
| 2250 | IV | 826 | 0 | 1 | 283 | 0 | 0 | Mono | Mono | | 14774366 | Z17894 | A1b | | 2250 | |
| 3817 | IV | 827 | 0 | 0 | 283 | 0 | 0 | Mono | Mono | | 18180294 | L1009 | A1b | | 3817 | |
| 3961 | IV | 827 | 0 | 0 | 283 | 0 | 0 | Mono | Mono | | 18759708 | L1053 | A1b | | 3961 | |
| 4268 | IV | 818 | 0 | 9 | 281 | 0 | 2 | Mono | Mono | | 19384407 | V4072.1 | A1b | | 4268 | |
| 4270 | IV | 827 | 0 | 0 | 283 | 0 | 0 | Mono | Mono | | 19390790 | V4076 | A1b | | 4270 | |
| 553 | IV | 827 | 0 | 0 | 283 | 0 | 0 | Mono | Mono | | 6845936 | V50 | A1b1a | | 553 | |
| 911 | IV | 826 | 0 | 1 | 283 | 0 | 0 | Mono | Mono | | 7593921 | V224 | A1b1a | | 911 | |
| 3109 | IV | 827 | 0 | 0 | 283 | 0 | 0 | Mono | Mono | | 16582907 | L602 | A1b1a | | 3109 | |
| 676 | IV | 821 | 0 | 6 | 283 | 0 | 0 | Mono | Mono | | 6954470 | Page71 | A1b1a1 | | 676 | |
| 2376 | IV | 827 | 0 | 0 | 283 | 0 | 0 | Mono | Mono | | 14930151 | Page87 | A1b1a1 | | 2376 | |
| 2735 | IV | 827 | 0 | 0 | 283 | 0 | 0 | Mono | Mono | | 15587462 | M206 | A1b1a1 | | 2735 | |
| 4581 | IV | 826 | 0 | 1 | 283 | 0 | 0 | Mono | Mono | | 21610585 | PN3 | A1b1a1 | | 4581 | |
| 4654 | IV | 827 | 0 | 0 | 283 | 0 | 0 | Mono | Mono | | 21722268 | M14 | A1b1a1 | | 4654 | |
| 4672 | IV | 827 | 0 | 0 | 283 | 0 | 0 | Mono | Mono | | 21736239 | M23 | A1b1a1 | | 4672 | |
| 4762 | IV | 827 | 0 | 0 | 283 | 0 | 0 | Mono | Mono | | 21844357 | P5 | A1b1a1 | | 4762 | |
| 4837 | IV | 827 | 0 | 0 | 283 | 0 | 0 | Mono | Mono | | 21894447 | M71 | A1b1a1 | | 4837 | |
| 5140 | IV | 826 | 0 | 1 | 283 | 0 | 0 | Mono | Mono | | 22741728 | P247 | A1b1a1 | | 5140 | |
| 5145 | IV | 827 | 0 | 0 | 283 | 0 | 0 | Mono | Mono | | 22741821 | P248 | A1b1a1 | | 5145 | |
| 5494 | IV | 826 | 0 | 1 | 283 | 0 | 0 | Mono | Mono | | 23497256 | MEH1 | A1b1a1 | | 5494 | |
| 5496 | IV | 827 | 0 | 0 | 283 | 0 | 0 | Mono | Mono | | 23497714 | P4 | A1b1a1 | | 5496 | |
| 2479 | IV | 825 | 0 | 2 | 283 | 0 | 0 | Mono | Mono | | 15030682 | M196 | A1b1a1a | | 2479 | |
| 3878 | IV | 827 | 0 | 0 | 283 | 0 | 0 | Mono | Mono | | 18571026 | M6 | A1b1a1a | | 3878 | |
| 4787 | IV | 827 | 0 | 0 | 283 | 0 | 0 | Mono | Mono | | 21868726 | Page41 | A1b1a1a | | 4787 | |
| 4813 | IV | 827 | 0 | 0 | 283 | 0 | 0 | Mono | Mono | | 21881115 | P28 | A1b1a1a1 | | 4813 | |
| 2697 | IV | 827 | 0 | 0 | 283 | 0 | 0 | Mono | Mono | | 15526695 | M212 | A1b1a1a2a | | 2697 | |
| 4720 | IV | 827 | 0 | 0 | 283 | 0 | 0 | Mono | Mono | | 21765763 | M114 | A1b1a1a2a | | 4720 | |
| 635 | IV | 825 | 0 | 2 | 283 | 0 | 0 | Mono | Mono | | 6932148 | P262 | A1b1a1a2a1a | | 635 | |
| 589 | IV | 827 | 0 | 0 | 283 | 0 | 0 | Mono | Mono | | 6879761 | V79 | A1b1a1a2a1a~ | | 589 | |
| 923 | IV | 827 | 0 | 0 | 283 | 0 | 0 | Mono | Mono | | 7604310 | V228 | A1b1a1a2a1a~ | | 923 | |
| 956 | IV | 827 | 0 | 0 | 283 | 0 | 0 | Mono | Mono | | 7659715 | V242 | A1b1a1a2a1a~ | | 956 | |
| 2931 | IV | 826 | 0 | 1 | 283 | 0 | 0 | Mono | Mono | | 16181491 | V70 | A1b1a1a2a1a~ | | 2931 | |
| 3372 | IV | 814 | 0 | 13 | 280 | 0 | 3 | Mono | Mono | | 17202140 | V61 | A1b1a1a2a1a~ | | 3372 | |
| 3373 | IV | 822 | 0 | 5 | 283 | 0 | 0 | Mono | Mono | | 17202140 | V61 | A1b1a1a2a1a~ | | 3373 | |
| 48 | III | 826 | 0 | 1 | 277 | 6 | 0 | M9- | 12 | | 2710309 | Page52.3 | A1b1a1a2b~ | A,W | 48 | |
| 49 | IV | 827 | 0 | 0 | 283 | 0 | 0 | Mono | Mono | | 2710494 | L949 | A1b1a1a2b~ | | 49 | |
| 642 | IV | 827 | 0 | 0 | 283 | 0 | 0 | Mono | Mono | | 6932648 | L951 | A1b1a1a2b~ | | 642 | |
| 733 | IV | 827 | 0 | 0 | 283 | 0 | 0 | Mono | Mono | | 7076801 | L952 | A1b1a1a2b~ | | 733 | |
| 4129 | IV | 827 | 0 | 0 | 283 | 0 | 0 | Mono | Mono | | 19105935 | L965 | A1b1a1a2b~ | | 4129 | |
| 5096 | IV | 827 | 0 | 0 | 283 | 0 | 0 | Mono | Mono | | 22669438 | L973 | A1b1a1a2b~ | | 5096 | |
| 5286 | IV | 824 | 0 | 3 | 282 | 0 | 1 | Mono | Mono | | 23010731 | L975 | A1b1a1a2b~ | | 5286 | |
| 5415 | IV | 826 | 0 | 1 | 283 | 0 | 0 | Mono | Mono | | 23293047 | L976 | A1b1a1a2b~ | | 5415 | |
| 3283 | IV | 827 | 0 | 0 | 283 | 0 | 0 | Mono | Mono | | 16942397 | Page110 | A1b1a1a3~ | | 3283 | |
| 4681 | IV | 826 | 0 | 1 | 283 | 0 | 0 | Mono | Mono | | 21740436 | M32 | A1b1b | | 4681 | |
| 4661 | IV | 826 | 0 | 1 | 283 | 0 | 0 | Mono | Mono | | 21729839 | M28 | A1b1b1 | | 4661 | |

| | | M9+ | | | M9- | | | | | | Build 37 | | ISOGG | | |
|---|---|---|---|---|---|---|---|---|---|---|---|---|---|---|---|
| List | Stage | 0 | 1 | . | 0 | 1 | . | M9? | Clade(s) | Prob. | Position | Name | Subgroup Name | Amb. | List |
| 614 | IV | 827 | 0 | 0 | 283 | 0 | 0 | Mono | Mono | | 6907838 | V160 | A1b1b2 | | 614 |
| 1371 | IV | 827 | 0 | 0 | 283 | 0 | 0 | Mono | Mono | | 8467082 | PF1372 | A1b1b2 | | 1371 |
| 2407 | IV | 826 | 0 | 1 | 283 | 0 | 0 | Mono | Mono | | 14968527 | PF1373 | A1b1b2 | | 2407 |
| 2637 | IV | 827 | 0 | 0 | 283 | 0 | 0 | Mono | Mono | | 15437050 | M220 | A1b1b2 | | 2637 |
| 3538 | IV | 827 | 0 | 0 | 283 | 0 | 0 | Mono | Mono | | 17596003 | L427 | A1b1b2 | | 3538 |
| 3539 | IV | 827 | 0 | 0 | 283 | 0 | 0 | Mono | Mono | | 17596023 | L430 | A1b1b2 | | 3539 |
| 4883 | IV | 827 | 0 | 0 | 283 | 0 | 0 | Mono | Mono | | 21925500 | M144 | A1b1b2 | | 4883 |
| 5162 | IV | 827 | 0 | 0 | 283 | 0 | 0 | Mono | Mono | | 22750174 | Page17 | A1b1b2 | | 5162 |
| 2405 | IV | 827 | 0 | 0 | 283 | 0 | 0 | Mono | Mono | | 14968331 | Page89 | A1b1b2a | | 2405 |
| 2692 | IV | 827 | 0 | 0 | 283 | 0 | 0 | Mono | Mono | | 15515812 | V3004 | A1b1b2a | | 2692 |
| 4584 | IV | 827 | 0 | 0 | 283 | 0 | 0 | Mono | Mono | | 21612078 | P71 | A1b1b2a | | 4584 |
| 4789 | IV | 827 | 0 | 0 | 283 | 0 | 0 | Mono | Mono | | 21868863 | Page42 | A1b1b2a | | 4789 |
| 1369 | IV | 827 | 0 | 0 | 283 | 0 | 0 | Mono | Mono | | 8466988 | P291 | A1b1b2a1 | | 1369 |
| 2476 | IV | 827 | 0 | 0 | 283 | 0 | 0 | Mono | Mono | | 15029492 | M202 | A1b1b2b | | 2476 |
| 4653 | IV | 827 | 0 | 0 | 283 | 0 | 0 | Mono | Mono | | 21722098 | PF1374 | A1b1b2b | | 4653 |
| 4707 | IV | 827 | 0 | 0 | 283 | 0 | 0 | Mono | Mono | | 21763273 | M127 | A1b1b2b | | 4707 |
| 4796 | IV | 827 | 0 | 0 | 283 | 0 | 0 | Mono | Mono | | 21870638 | PF1079 | A1b1b2b | | 4796 |
| 600 | IV | 826 | 0 | 1 | 283 | 0 | 0 | Mono | Mono | | 6893093 | V10 | A1b1b2b~ | | 600 |
| 2873 | IV | 827 | 0 | 0 | 283 | 0 | 0 | Mono | Mono | | 15879104 | L420 | A1b1b2b~ | | 2873 |
| 4117 | IV | 827 | 0 | 0 | 283 | 0 | 0 | Mono | Mono | | 19077672 | PF962 | A1b1b2b~ | | 4117 |
| 4975 | IV | 822 | 0 | 5 | 280 | 0 | 3 | Mono | Mono | | 22157292 | L439 | A1b1b2b~ | | 4975 |
| 5243 | IV | 827 | 0 | 0 | 283 | 0 | 0 | Mono | Mono | | 22922129 | Page53 | A1b1b2b~ | | 5243 |
| 5634 | IV | 770 | 0 | 57 | 275 | 0 | 8 | Mono | Mono | | 23989836 | PF1287 | A1b1b2b~ | | 5634 |
| 4710 | IV | 827 | 0 | 0 | 283 | 0 | 0 | Mono | Mono | | 21763965 | M118 | A1b1b2b1 | | 4710 |
| 524 | IV | 827 | 0 | 0 | 283 | 0 | 0 | Mono | Mono | | 6777077 | V1 | A1b1b2b2~ | | 524 |
| 586 | IV | 827 | 0 | 0 | 283 | 0 | 0 | Mono | Mono | | 6876484 | V156 | A1b1b2b2~ | | 586 |
| 592 | IV | 827 | 0 | 0 | 283 | 0 | 0 | Mono | Mono | | 6885631 | V155 | A1b1b2b2~ | | 592 |
| 3473 | IV | 822 | 0 | 5 | 283 | 0 | 0 | Mono | Mono | | 17466305 | V51 | A1b1b2b2~ | | 3473 |
| 3554 | IV | 826 | 0 | 1 | 283 | 0 | 0 | Mono | Mono | | 17645441 | V67 | A1c~ | | 3554 |
| 536 | IV | 827 | 0 | 0 | 283 | 0 | 0 | Mono | Mono | | 6804479 | M8680 | B~ | | 536 |
| 4736 | IV | 827 | 0 | 0 | 283 | 0 | 0 | Mono | Mono | | 21779251 | M146 | B1a | | 4736 |
| 2312 | IV | 825 | 0 | 2 | 282 | 0 | 1 | Mono | Mono | | 14869076 | M182 | B2 | | 2312 |
| 2449 | IV | 827 | 0 | 0 | 283 | 0 | 0 | Mono | Mono | | 15020578 | P85 | B2 | | 2449 |
| 467 | IV | 827 | 0 | 0 | 283 | 0 | 0 | Mono | Mono | | 6659577 | V217 | B2~ | | 467 |
| 595 | IV | 827 | 0 | 0 | 283 | 0 | 0 | Mono | Mono | | 6887162 | V85 | B2~ | | 595 |
| 943 | IV | 827 | 0 | 0 | 283 | 0 | 0 | Mono | Mono | | 7644670 | V234 | B2~ | | 943 |
| 2502 | IV | 827 | 0 | 0 | 283 | 0 | 0 | Mono | Mono | | 15064733 | M8760 | B2~ | | 2502 |
| 4793 | IV | 827 | 0 | 0 | 283 | 0 | 0 | Mono | Mono | | 21869519 | M150 | B2a | | 4793 |
| 571 | IV | 824 | 0 | 3 | 282 | 0 | 1 | Mono | Mono | | 6859558 | V78 | B2a1~ | | 571 |
| 2638 | IV | 826 | 0 | 1 | 283 | 0 | 0 | Mono | Mono | | 15437063 | M218 | B2a1a1 | | 2638 |
| 4728 | IV | 827 | 0 | 0 | 283 | 0 | 0 | Mono | Mono | | 21774180 | M109 | B2a1a1a1 | | 4728 |
| 4783 | IV | 827 | 0 | 0 | 283 | 0 | 0 | Mono | Mono | | 21868068 | Page60 | B2a1a1a1 | | 4783 |
| 918 | IV | 826 | 0 | 1 | 283 | 0 | 0 | Mono | Mono | | 7599027 | V227 | B2a1a1a1~ | | 918 |
| 4593 | IV | 827 | 0 | 0 | 283 | 0 | 0 | Mono | Mono | | 21621191 | Page18 | B2a1a1a1~ | | 4593 |
| 5337 | IV | 827 | 0 | 0 | 283 | 0 | 0 | Mono | Mono | | 23120337 | V197 | B2a1a1a1~ | A | 5337 |
| 582 | IV | 827 | 0 | 0 | 283 | 0 | 0 | Mono | Mono | | 6870606 | Z22737 | B2a1a1a1a2 | | 582 |
| 2317 | IV | 827 | 0 | 0 | 283 | 0 | 0 | Mono | Mono | | 14870505 | V2647 | B2a1a1a1a2 | | 2317 |
| 4890 | IV | 827 | 0 | 0 | 283 | 0 | 0 | Mono | Mono | | 21932926 | G1 | B2a1a1a2 | | 4890 |
| 904 | IV | 827 | 0 | 0 | 283 | 0 | 0 | Mono | Mono | | 7583271 | M43 | B2a1a2a | | 904 |
| 4723 | IV | 826 | 0 | 1 | 283 | 0 | 0 | Mono | Mono | | 21767959 | M112 | B2b | | 4723 |
| 681 | IV | 810 | 0 | 17 | 280 | 0 | 3 | Mono | Mono | | 6958165 | Page72 | B2b~ | | 681 |
| 5493 | IV | 826 | 0 | 1 | 282 | 0 | 1 | Mono | Mono | | 23497067 | M8659 | B2b1 | | 5493 |
| 2215 | IV | 827 | 0 | 0 | 283 | 0 | 0 | Mono | Mono | | 14654615 | V2539 | B2b1a1~ | | 2215 |
| 3209 | IV | 827 | 0 | 0 | 283 | 0 | 0 | Mono | Mono | | 16834833 | Page48 | B2b1a1~ | | 3209 |
| 2272 | IV | 827 | 0 | 0 | 283 | 0 | 0 | Mono | Mono | | 14813717 | M169 | B2b1a1a~ | | 2272 |
| 4722 | IV | 827 | 0 | 0 | 283 | 0 | 0 | Mono | Mono | | 21766310 | M115 | B2b1a1a~ | | 4722 |
| 4680 | IV | 827 | 0 | 0 | 283 | 0 | 0 | Mono | Mono | | 21739815 | M30 | B2b1a1b | | 4680 |
| 4891 | IV | 827 | 0 | 0 | 283 | 0 | 0 | Mono | Mono | | 21932935 | M108.2 | B2b1a1b1 | A | 4891 |
| 2710 | IV | 827 | 0 | 0 | 283 | 0 | 0 | Mono | Mono | | 15544614 | M211 | B2b1a1c1 | | 2710 |
| 2369 | IV | 822 | 0 | 5 | 283 | 0 | 0 | Mono | Mono | | 14924617 | Page86 | B2b1a1d~ | | 2369 |
| 1011 | IV | 827 | 0 | 0 | 283 | 0 | 0 | Mono | Mono | | 7743388 | M7601.2 | B2b1a2~ | A | 1011 |
| 4864 | IV | 827 | 0 | 0 | 283 | 0 | 0 | Mono | Mono | | 21906265 | M65.2 | B2b1b2~ | | 4864 |
| 28 | IV | 826 | 0 | 1 | 283 | 0 | 0 | Mono | Mono | | 2669416 | M8945 | BT | | 28 |
| 52 | IV | 827 | 0 | 0 | 283 | 0 | 0 | Mono | Mono | | 2712956 | M8947 | BT | | 52 |
| 91 | IV | 827 | 0 | 0 | 283 | 0 | 0 | Mono | Mono | | 2756519 | V187 | BT | | 91 |
| 147 | IV | 827 | 0 | 0 | 283 | 0 | 0 | Mono | Mono | | 2842830 | M8951 | BT | | 147 |
| 162 | IV | 827 | 0 | 0 | 282 | 0 | 1 | Mono | Mono | | 2868656 | V216 | BT | | 162 |
| 184 | IV | 826 | 0 | 1 | 283 | 0 | 0 | Mono | Mono | | 2889399 | M8954 | BT | | 184 |
| 485 | IV | 827 | 0 | 0 | 283 | 0 | 0 | Mono | Mono | | 6691986 | M8959 | BT | | 485 |
| 500 | IV | 827 | 0 | 0 | 283 | 0 | 0 | Mono | Mono | | 6719048 | M8961 | BT | | 500 |
| 527 | IV | 827 | 0 | 0 | 283 | 0 | 0 | Mono | Mono | | 6779907 | M8967 | BT | | 527 |
| 547 | IV | 827 | 0 | 0 | 283 | 0 | 0 | Mono | Mono | | 6835056 | M8968 | BT | | 547 |
| 572 | IV | 827 | 0 | 0 | 283 | 0 | 0 | Mono | Mono | | 6859691 | V21 | BT | | 572 |
| 578 | IV | 827 | 0 | 0 | 283 | 0 | 0 | Mono | Mono | | 6865490 | M8970 | BT | | 578 |
| 615 | IV | 827 | 0 | 0 | 283 | 0 | 0 | Mono | Mono | | 6908261 | PF1408 | BT | | 615 |
| 649 | IV | 827 | 0 | 0 | 283 | 0 | 0 | Mono | Mono | | 6932831 | V31 | BT | | 649 |
| 655 | IV | 827 | 0 | 0 | 282 | 0 | 1 | Mono | Mono | | 6936412 | M8971 | BT | | 655 |
| 700 | IV | 827 | 0 | 0 | 283 | 0 | 0 | Mono | Mono | | 6993202 | M8972 | BT | | 700 |
| 711 | IV | 827 | 0 | 0 | 283 | 0 | 0 | Mono | Mono | | 7037901 | M8976 | BT | | 711 |
| 779 | IV | 827 | 0 | 0 | 283 | 0 | 0 | Mono | Mono | | 7171532 | M8977 | BT | | 779 |
| 815 | IV | 827 | 0 | 0 | 283 | 0 | 0 | Mono | Mono | | 7269376 | M8980 | BT | | 815 |
| 824 | IV | 827 | 0 | 0 | 283 | 0 | 0 | Mono | Mono | | 7302697 | M8983 | BT | | 824 |
| 835 | IV | 827 | 0 | 0 | 283 | 0 | 0 | Mono | Mono | | 7324963 | M8986 | BT | | 835 |
| 870 | IV | 820 | 0 | 7 | 281 | 0 | 2 | Mono | Mono | | 7531097 | M8988 | BT | | 870 |
| 892 | IV | 827 | 0 | 0 | 283 | 0 | 0 | Mono | Mono | | 7563503 | Z40386 | BT | | 892 |
| 959 | IV | 827 | 0 | 0 | 283 | 0 | 0 | Mono | Mono | | 7661695 | M8994 | BT | | 959 |
| 1001 | IV | 826 | 0 | 1 | 283 | 0 | 0 | Mono | Mono | | 7735760 | M8997 | BT | | 1001 |
| 1022 | IV | 827 | 0 | 0 | 283 | 0 | 0 | Mono | Mono | | 7763132 | M8999 | BT | | 1022 |
| 1048 | IV | 827 | 0 | 0 | 283 | 0 | 0 | Mono | Mono | | 7815370 | M9000 | BT | | 1048 |
| 1055 | IV | 827 | 0 | 0 | 283 | 0 | 0 | Mono | Mono | | 7838397 | M9001 | BT | | 1055 |
| 1078 | IV | 827 | 0 | 0 | 283 | 0 | 0 | Mono | Mono | | 7890655 | M9003 | BT | | 1078 |
| 1084 | IV | 826 | 0 | 1 | 283 | 0 | 0 | Mono | Mono | | 7904101 | M9005 | BT | | 1084 |
| 1088 | IV | 827 | 0 | 0 | 283 | 0 | 0 | Mono | Mono | | 7907933 | M9006 | BT | | 1088 |
| 1106 | IV | 826 | 0 | 1 | 283 | 0 | 0 | Mono | Mono | | 7944491 | M9008 | BT | | 1106 |
| 1127 | IV | 823 | 0 | 4 | 283 | 0 | 0 | Mono | Mono | | 8009052 | M9009 | BT | | 1127 |
| 1143 | IV | 827 | 0 | 0 | 283 | 0 | 0 | Mono | Mono | | 8042040 | Z40387 | BT | | 1143 |

| | | | M9+ | | | M9- | | | | | Build 37 | | ISOGG | | |
|---|---|---|---|---|---|---|---|---|---|---|---|---|---|---|---|
| List | Stage | 0 | 1 | . | 0 | 1 | . | M9? | Clade(s) | Prob. | Position | Name | Subgroup Name | Amb. | List |
| 1178 | IV | 827 | 0 | 0 | 282 | 0 | 1 | Mono | Mono | | 8130509 | M9015 | BT | | 1178 |
| 1194 | IV | 827 | 0 | 0 | 283 | 0 | 0 | Mono | Mono | | 8165769 | M9016 | BT | | 1194 |
| 1195 | IV | 827 | 0 | 0 | 283 | 0 | 0 | Mono | Mono | | 8180027 | Z40388 | BT | | 1195 |
| 1215 | IV | 826 | 0 | 1 | 282 | 0 | 1 | Mono | Mono | | 8226191 | M9017 | BT | | 1215 |
| 1272 | IV | 827 | 0 | 0 | 283 | 0 | 0 | Mono | Mono | | 8330559 | M9019 | BT | | 1272 |
| 1277 | IV | 827 | 0 | 0 | 283 | 0 | 0 | Mono | Mono | | 8337827 | M9020 | BT | | 1277 |
| 1348 | IV | 827 | 0 | 0 | 283 | 0 | 0 | Mono | Mono | | 8446082 | M9025 | BT | | 1348 |
| 1421 | IV | 827 | 0 | 0 | 283 | 0 | 0 | Mono | Mono | | 8511682 | M9026 | BT | | 1421 |
| 1436 | IV | 827 | 0 | 0 | 283 | 0 | 0 | Mono | Mono | | 8540845 | M9028 | BT | | 1436 |
| 1477 | IV | 827 | 0 | 0 | 283 | 0 | 0 | Mono | Mono | | 8594070 | Z40389 | BT | | 1477 |
| 1480 | IV | 827 | 0 | 0 | 283 | 0 | 0 | Mono | Mono | | 8596285 | M9030 | BT | | 1480 |
| 1493 | IV | 827 | 0 | 0 | 283 | 0 | 0 | Mono | Mono | | 8613037 | PF302 | BT | | 1493 |
| 1524 | IV | 826 | 0 | 1 | 283 | 0 | 0 | Mono | Mono | | 8662142 | M9034 | BT | | 1524 |
| 1561 | IV | 826 | 0 | 1 | 282 | 0 | 1 | Mono | Mono | | 8729512 | M9038 | BT | | 1561 |
| 1610 | IV | 827 | 0 | 0 | 283 | 0 | 0 | Mono | Mono | | 8881498 | M9042 | BT | | 1610 |
| 1642 | IV | 827 | 0 | 0 | 283 | 0 | 0 | Mono | Mono | | 9066684 | M9045 | BT | | 1642 |
| 1651 | IV | 827 | 0 | 0 | 283 | 0 | 0 | Mono | Mono | | 9106630 | M9046 | BT | | 1651 |
| 1740 | IV | 827 | 0 | 0 | 283 | 0 | 0 | Mono | Mono | | 9827891 | M9054 | BT | | 1740 |
| 1747 | IV | 827 | 0 | 0 | 283 | 0 | 0 | Mono | Mono | | 9840307 | M9056 | BT | | 1747 |
| 1749 | IV | 827 | 0 | 0 | 283 | 0 | 0 | Mono | Mono | | 9842503 | M9057 | BT | | 1749 |
| 1756 | IV | 824 | 0 | 3 | 283 | 0 | 0 | Mono | Mono | | 9850716 | M9064 | BT | | 1756 |
| 1757 | IV | 823 | 0 | 4 | 283 | 0 | 0 | Mono | Mono | | 9850725 | M9065 | BT | | 1757 |
| 1823 | IV | 827 | 0 | 0 | 282 | 0 | 1 | Mono | Mono | | 10057648 | Z40375 | BT | | 1823 |
| 1825 | IV | 825 | 0 | 2 | 281 | 0 | 2 | Mono | Mono | | 10068588 | Z40376 | BT | | 1825 |
| 1869 | IV | 826 | 0 | 1 | 281 | 0 | 2 | Mono | Mono | | 13668726 | Z40380 | BT | | 1869 |
| 1914 | IV | 815 | 0 | 12 | 280 | 0 | 3 | Mono | Mono | | 13823056 | Z40401 | BT | | 1914 |
| 1956 | IV | 827 | 0 | 0 | 283 | 0 | 0 | Mono | Mono | | 14013228 | M9075 | BT | | 1956 |
| 1958 | IV | 826 | 0 | 1 | 283 | 0 | 0 | Mono | Mono | | 14018605 | M9076 | BT | | 1958 |
| 1986 | II | 825 | 2 | 0 | 283 | 0 | 0 | M9+ | 5 | | 14069414 | M9079 | BT | W | 1986 |
| 1991 | IV | 827 | 0 | 0 | 283 | 0 | 0 | Mono | Mono | | 14075450 | M9080 | BT | | 1991 |
| 2003 | IV | 827 | 0 | 0 | 283 | 0 | 0 | Mono | Mono | | 14096790 | M9081 | BT | | 2003 |
| 2008 | IV | 827 | 0 | 0 | 283 | 0 | 0 | Mono | Mono | | 14108214 | M9083 | BT | | 2008 |
| 2027 | IV | 827 | 0 | 0 | 283 | 0 | 0 | Mono | Mono | | 14150833 | M9089 | BT | | 2027 |
| 2116 | IV | 827 | 0 | 0 | 283 | 0 | 0 | Mono | Mono | | 14344506 | M9094 | BT | | 2116 |
| 2127 | IV | 826 | 0 | 1 | 283 | 0 | 0 | Mono | Mono | | 14386666 | M9097 | BT | | 2127 |
| 2148 | IV | 827 | 0 | 0 | 283 | 0 | 0 | Mono | Mono | | 14461702 | M9099 | BT | | 2148 |
| 2230 | IV | 827 | 0 | 0 | 283 | 0 | 0 | Mono | Mono | | 14720802 | M9110 | BT | | 2230 |
| 2239 | IV | 827 | 0 | 0 | 283 | 0 | 0 | Mono | Mono | | 14762104 | M9114 | BT | | 2239 |
| 2255 | IV | 822 | 0 | 5 | 283 | 0 | 0 | Mono | Mono | | 14778098 | M9115 | BT | | 2255 |
| 2260 | IV | 815 | 0 | 12 | 278 | 0 | 5 | Mono | Mono | | 14786996 | M9116 | BT | | 2260 |
| 2293 | IV | 827 | 0 | 0 | 283 | 0 | 0 | Mono | Mono | | 14839376 | M9117 | BT | | 2293 |
| 2324 | IV | 827 | 0 | 0 | 283 | 0 | 0 | Mono | Mono | | 14885368 | Page81 | BT | | 2324 |
| 2417 | IV | 826 | 0 | 1 | 283 | 0 | 0 | Mono | Mono | | 14980170 | M9121 | BT | | 2417 |
| 2454 | IV | 827 | 0 | 0 | 283 | 0 | 0 | Mono | Mono | | 15022370 | M9122 | BT | | 2454 |
| 2501 | IV | 827 | 0 | 0 | 283 | 0 | 0 | Mono | Mono | | 15063698 | M9124 | BT | | 2501 |
| 2503 | IV | 827 | 0 | 0 | 283 | 0 | 0 | Mono | Mono | | 15068525 | M9125 | BT | | 2503 |
| 2515 | IV | 827 | 0 | 0 | 283 | 0 | 0 | Mono | Mono | | 15095089 | M9126 | BT | | 2515 |
| 2525 | IV | 827 | 0 | 0 | 283 | 0 | 0 | Mono | Mono | | 15112480 | M9128 | BT | | 2525 |
| 2568 | IV | 827 | 0 | 0 | 283 | 0 | 0 | Mono | Mono | | 15256839 | M9133 | BT | | 2568 |
| 2569 | IV | 827 | 0 | 0 | 283 | 0 | 0 | Mono | Mono | | 15261255 | Z40392 | BT | | 2569 |
| 2607 | IV | 827 | 0 | 0 | 283 | 0 | 0 | Mono | Mono | | 15381787 | M9135 | BT | | 2607 |
| 2608 | IV | 827 | 0 | 0 | 283 | 0 | 0 | Mono | Mono | | 15382324 | M9136 | BT | | 2608 |
| 2690 | IV | 827 | 0 | 0 | 283 | 0 | 0 | Mono | Mono | | 15514552 | M9140 | BT | | 2690 |
| 2718 | IV | 827 | 0 | 0 | 283 | 0 | 0 | Mono | Mono | | 15558761 | M9142 | BT | | 2718 |
| 2724 | IV | 827 | 0 | 0 | 283 | 0 | 0 | Mono | Mono | | 15569966 | M9143 | BT | | 2724 |
| 2754 | IV | 820 | 0 | 7 | 280 | 0 | 3 | Mono | Mono | | 15593938 | M9145 | BT | | 2754 |
| 2815 | IV | 827 | 0 | 0 | 282 | 0 | 1 | Mono | Mono | | 15731578 | M9148 | BT | | 2815 |
| 2881 | IV | 827 | 0 | 0 | 283 | 0 | 0 | Mono | Mono | | 15905094 | M9152 | BT | | 2881 |
| 2907 | IV | 794 | 0 | 33 | 272 | 0 | 11 | Mono | Mono | | 15995289 | M9157 | BT | | 2907 |
| 2999 | IV | 826 | 0 | 1 | 283 | 0 | 0 | Mono | Mono | | 16332337 | M9166 | BT | | 2999 |
| 3094 | IV | 827 | 0 | 0 | 283 | 0 | 0 | Mono | Mono | | 16542464 | S1572 | BT | | 3094 |
| 3121 | IV | 825 | 0 | 2 | 283 | 0 | 0 | Mono | Mono | | 16608008 | M9176 | BT | | 3121 |
| 3142 | IV | 827 | 0 | 0 | 283 | 0 | 0 | Mono | Mono | | 16682167 | Y8489 | BT | | 3142 |
| 3148 | IV | 826 | 0 | 1 | 283 | 0 | 0 | Mono | Mono | | 16694398 | M9180 | BT | | 3148 |
| 3170 | IV | 827 | 0 | 0 | 283 | 0 | 0 | Mono | Mono | | 16766467 | M9182 | BT | | 3170 |
| 3192 | IV | 827 | 0 | 0 | 283 | 0 | 0 | Mono | Mono | | 16808769 | Z40393 | BT | | 3192 |
| 3267 | IV | 827 | 0 | 0 | 283 | 0 | 0 | Mono | Mono | | 16913677 | M9187 | BT | | 3267 |
| 3285 | IV | 800 | 0 | 27 | 276 | 0 | 7 | Mono | Mono | | 16946901 | M9188 | BT | | 3285 |
| 3296 | IV | 827 | 0 | 0 | 283 | 0 | 0 | Mono | Mono | | 16959883 | M9189 | BT | | 3296 |
| 3318 | IV | 827 | 0 | 0 | 283 | 0 | 0 | Mono | Mono | | 17037979 | V59 | BT | | 3318 |
| 3322 | IV | 827 | 0 | 0 | 283 | 0 | 0 | Mono | Mono | | 17041558 | M9193 | BT | | 3322 |
| 3368 | IV | 827 | 0 | 0 | 283 | 0 | 0 | Mono | Mono | | 17192307 | M9199 | BT | | 3368 |
| 3405 | IV | 826 | 0 | 1 | 283 | 0 | 0 | Mono | Mono | | 17298439 | M9209 | BT | | 3405 |
| 3428 | IV | 827 | 0 | 0 | 283 | 0 | 0 | Mono | Mono | | 17353147 | M9212 | BT | | 3428 |
| 3437 | IV | 827 | 0 | 0 | 283 | 0 | 0 | Mono | Mono | | 17388578 | M9214 | BT | | 3437 |
| 3442 | IV | 827 | 0 | 0 | 283 | 0 | 0 | Mono | Mono | | 17393971 | M9215 | BT | | 3442 |
| 3485 | IV | 827 | 0 | 0 | 283 | 0 | 0 | Mono | Mono | | 17504731 | M9218 | BT | | 3485 |
| 3492 | IV | 827 | 0 | 0 | 283 | 0 | 0 | Mono | Mono | | 17510373 | L962 | BT | | 3492 |
| 3494 | IV | 827 | 0 | 0 | 283 | 0 | 0 | Mono | Mono | | 17511829 | M9219 | BT | | 3494 |
| 3524 | IV | 813 | 0 | 14 | 280 | 0 | 3 | Mono | Mono | | 17571812 | M9220 | BT | | 3524 |
| 3550 | IV | 827 | 0 | 0 | 283 | 0 | 0 | Mono | Mono | | 17623760 | M9223 | BT | | 3550 |
| 3566 | IV | 826 | 0 | 1 | 283 | 0 | 0 | Mono | Mono | | 17670179 | M9226 | BT | | 3566 |
| 3567 | IV | 827 | 0 | 0 | 283 | 0 | 0 | Mono | Mono | | 17670193 | M9227 | BT | | 3567 |
| 3571 | IV | 825 | 0 | 2 | 282 | 0 | 1 | Mono | Mono | | 17674985 | M9228 | BT | | 3571 |
| 3599 | IV | 827 | 0 | 0 | 283 | 0 | 0 | Mono | Mono | | 17753892 | M9231 | BT | | 3599 |
| 3648 | IV | 827 | 0 | 0 | 282 | 0 | 1 | Mono | Mono | | 17840394 | M9235 | BT | | 3648 |
| 3675 | IV | 827 | 0 | 0 | 283 | 0 | 0 | Mono | Mono | | 17886479 | M9237 | BT | | 3675 |
| 3682 | IV | 827 | 0 | 0 | 283 | 0 | 0 | Mono | Mono | | 17903900 | M9238 | BT | | 3682 |
| 3686 | IV | 827 | 0 | 0 | 283 | 0 | 0 | Mono | Mono | | 17909260 | M9239 | BT | | 3686 |
| 3707 | IV | 827 | 0 | 0 | 283 | 0 | 0 | Mono | Mono | | 17940510 | M9240 | BT | | 3707 |
| 3718 | IV | 825 | 0 | 2 | 283 | 0 | 0 | Mono | Mono | | 17966904 | M9242 | BT | | 3718 |
| 3757 | IV | 827 | 0 | 0 | 283 | 0 | 0 | Mono | Mono | | 18053867 | M9244 | BT | | 3757 |
| 3764 | IV | 827 | 0 | 0 | 283 | 0 | 0 | Mono | Mono | | 18067088 | M9245 | BT | | 3764 |
| 3767 | IV | 827 | 0 | 0 | 283 | 0 | 0 | Mono | Mono | | 18069115 | M9246 | BT | | 3767 |
| 3806 | IV | 827 | 0 | 0 | 283 | 0 | 0 | Mono | Mono | | 18162272 | M9252 | BT | | 3806 |
| 3816 | IV | 827 | 0 | 0 | 282 | 0 | 1 | Mono | Mono | | 18179224 | M9253 | BT | | 3816 |

| | | | M9+ | | | M9- | | | | | Build 37 | | ISOGG | | |
|---|---|---|---|---|---|---|---|---|---|---|---|---|---|---|---|
| List | Stage | 0 | 1 | . | 0 | 1 | . | M9? | Clade(s) | Prob. | Position | Name | Subgroup Name | Amb. | List |
| 3834 | IV | 827 | 0 | 0 | 283 | 0 | 0 | Mono | Mono | | 18242832 | M9254 | BT | | 3834 |
| 3889 | IV | 827 | 0 | 0 | 283 | 0 | 0 | Mono | Mono | | 18585407 | M9255 | BT | | 3889 |
| 3892 | IV | 827 | 0 | 0 | 283 | 0 | 0 | Mono | Mono | | 18589521 | M9257 | BT | | 3892 |
| 3914 | IV | 827 | 0 | 0 | 283 | 0 | 0 | Mono | Mono | | 18633816 | M9261 | BT | | 3914 |
| 3920 | IV | 827 | 0 | 0 | 283 | 0 | 0 | Mono | Mono | | 18643546 | M9262 | BT | | 3920 |
| 4004 | IV | 827 | 0 | 0 | 283 | 0 | 0 | Mono | Mono | | 18837871 | M9265 | BT | | 4004 |
| 4043 | IV | 827 | 0 | 0 | 283 | 0 | 0 | Mono | Mono | | 18917853 | M9267 | BT | | 4043 |
| 4062 | IV | 827 | 0 | 0 | 283 | 0 | 0 | Mono | Mono | | 18958429 | M9269 | BT | | 4062 |
| 4133 | IV | 827 | 0 | 0 | 282 | 0 | 1 | Mono | Mono | | 19118181 | M9277 | BT | | 4133 |
| 4144 | IV | 826 | 0 | 1 | 283 | 0 | 0 | Mono | Mono | | 19140679 | PF969 | BT | | 4144 |
| 4149 | IV | 827 | 0 | 0 | 283 | 0 | 0 | Mono | Mono | | 19149592 | M9280 | BT | | 4149 |
| 4182 | IV | 826 | 0 | 1 | 283 | 0 | 0 | Mono | Mono | | 19207843 | M9282 | BT | | 4182 |
| 4188 | IV | 827 | 0 | 0 | 283 | 0 | 0 | Mono | Mono | | 19218668 | M9283 | BT | | 4188 |
| 4195 | IV | 825 | 0 | 2 | 283 | 0 | 0 | Mono | Mono | | 19233322 | M9284 | BT | | 4195 |
| 4201 | IV | 827 | 0 | 0 | 283 | 0 | 0 | Mono | Mono | | 19235631 | M9285 | BT | | 4201 |
| 4209 | IV | 826 | 0 | 1 | 283 | 0 | 0 | Mono | Mono | | 19249565 | M9286 | BT | | 4209 |
| 4236 | IV | 826 | 0 | 1 | 283 | 0 | 0 | Mono | Mono | | 19306455 | V3998 | BT | | 4236 |
| 4241 | IV | 827 | 0 | 0 | 283 | 0 | 0 | Mono | Mono | | 19317082 | M9289 | BT | | 4241 |
| 4247 | IV | 826 | 0 | 1 | 281 | 0 | 2 | Mono | Mono | | 19337259 | M9291 | BT | | 4247 |
| 4271 | IV | 827 | 0 | 0 | 283 | 0 | 0 | Mono | Mono | | 19391922 | M9292 | BT | | 4271 |
| 4278 | IV | 826 | 0 | 1 | 283 | 0 | 0 | Mono | Mono | | 19416000 | M9293 | BT | | 4278 |
| 4287 | IV | 825 | 0 | 2 | 281 | 0 | 2 | Mono | Mono | | 19445252 | M9295 | BT | | 4287 |
| 4316 | IV | 812 | 0 | 15 | 279 | 0 | 4 | Mono | Mono | | 19519830 | Z12903 | BT | | 4316 |
| 4363 | IV | 827 | 0 | 0 | 283 | 0 | 0 | Mono | Mono | | 21069381 | M9300 | BT | | 4363 |
| 4387 | IV | 827 | 0 | 0 | 283 | 0 | 0 | Mono | Mono | | 21122071 | M9303 | BT | | 4387 |
| 4407 | IV | 825 | 0 | 2 | 282 | 0 | 1 | Mono | Mono | | 21183571 | PF1021 | BT | | 4407 |
| 4425 | IV | 826 | 0 | 1 | 283 | 0 | 0 | Mono | Mono | | 21247623 | M9310 | BT | | 4425 |
| 4426 | IV | 827 | 0 | 0 | 283 | 0 | 0 | Mono | Mono | | 21248962 | Z40394 | BT | | 4426 |
| 4431 | IV | 827 | 0 | 0 | 283 | 0 | 0 | Mono | Mono | | 21253574 | L438 | BT | | 4431 |
| 4437 | IV | 827 | 0 | 0 | 283 | 0 | 0 | Mono | Mono | | 21259569 | M9311 | BT | | 4437 |
| 4443 | IV | 821 | 0 | 6 | 282 | 0 | 1 | Mono | Mono | | 21268114 | M9312 | BT | | 4443 |
| 4452 | IV | 826 | 0 | 1 | 283 | 0 | 0 | Mono | Mono | | 21289152 | M9315 | BT | | 4452 |
| 4453 | IV | 826 | 0 | 1 | 283 | 0 | 0 | Mono | Mono | | 21292327 | M9316 | BT | | 4453 |
| 4464 | IV | 827 | 0 | 0 | 283 | 0 | 0 | Mono | Mono | | 21321168 | M9318 | BT | | 4464 |
| 4490 | IV | 827 | 0 | 0 | 283 | 0 | 0 | Mono | Mono | | 21378573 | Z40395 | BT | | 4490 |
| 4495 | IV | 827 | 0 | 0 | 283 | 0 | 0 | Mono | Mono | | 21391240 | M9321 | BT | | 4495 |
| 4533 | IV | 826 | 0 | 1 | 282 | 0 | 1 | Mono | Mono | | 21486590 | M9325 | BT | | 4533 |
| 4534 | IV | 827 | 0 | 0 | 283 | 0 | 0 | Mono | Mono | | 21490191 | M9326 | BT | | 4534 |
| 4537 | IV | 826 | 0 | 1 | 283 | 0 | 0 | Mono | Mono | | 21492793 | M9327 | BT | | 4537 |
| 4550 | IV | 827 | 0 | 0 | 282 | 0 | 1 | Mono | Mono | | 21518721 | M9328 | BT | | 4550 |
| 4565 | IV | 827 | 0 | 0 | 282 | 0 | 1 | Mono | Mono | | 21567686 | M9331 | BT | | 4565 |
| 4599 | IV | 827 | 0 | 0 | 283 | 0 | 0 | Mono | Mono | | 21627877 | M9334 | BT | | 4599 |
| 4605 | IV | 827 | 0 | 0 | 283 | 0 | 0 | Mono | Mono | | 21637759 | M9335 | BT | | 4605 |
| 4615 | IV | 827 | 0 | 0 | 283 | 0 | 0 | Mono | Mono | | 21646196 | Page26 | BT | | 4615 |
| 4630 | IV | 827 | 0 | 0 | 282 | 0 | 1 | Mono | Mono | | 21674225 | L969 | BT | | 4630 |
| 4750 | IV | 827 | 0 | 0 | 283 | 0 | 0 | Mono | Mono | | 21815245 | M9341 | BT | | 4750 |
| 4777 | IV | 827 | 0 | 0 | 283 | 0 | 0 | Mono | Mono | | 21866840 | M42 | BT | | 4777 |
| 4897 | IV | 827 | 0 | 0 | 283 | 0 | 0 | Mono | Mono | | 21938158 | M94 | BT | | 4897 |
| 4931 | IV | 827 | 0 | 0 | 283 | 0 | 0 | Mono | Mono | | 22025458 | M9346 | BT | | 4931 |
| 4938 | IV | 827 | 0 | 0 | 283 | 0 | 0 | Mono | Mono | | 22062727 | M9347 | BT | | 4938 |
| 4945 | IV | 827 | 0 | 0 | 283 | 0 | 0 | Mono | Mono | | 22079278 | M9348 | BT | | 4945 |
| 4969 | IV | 827 | 0 | 0 | 283 | 0 | 0 | Mono | Mono | | 22143034 | M9349 | BT | | 4969 |
| 4989 | IV | 827 | 0 | 0 | 283 | 0 | 0 | Mono | Mono | | 22178817 | L440 | BT | | 4989 |
| 4991 | IV | 823 | 0 | 4 | 280 | 0 | 3 | Mono | Mono | | 22180636 | M9352 | BT | | 4991 |
| 5086 | IV | 827 | 0 | 0 | 283 | 0 | 0 | Mono | Mono | | 22625732 | M9356 | BT | | 5086 |
| 5119 | IV | 827 | 0 | 0 | 283 | 0 | 0 | Mono | Mono | | 22719912 | M9357 | BT | | 5119 |
| 5125 | IV | 827 | 0 | 0 | 283 | 0 | 0 | Mono | Mono | | 22727889 | M9359 | BT | | 5125 |
| 5128 | IV | 827 | 0 | 0 | 283 | 0 | 0 | Mono | Mono | | 22730555 | M9360 | BT | | 5128 |
| 5129 | IV | 827 | 0 | 0 | 283 | 0 | 0 | Mono | Mono | | 22730777 | M9361 | BT | | 5129 |
| 5159 | IV | 821 | 0 | 6 | 281 | 0 | 2 | Mono | Mono | | 22748506 | M299 | BT | | 5159 |
| 5189 | IV | 827 | 0 | 0 | 283 | 0 | 0 | Mono | Mono | | 22798764 | M9362 | BT | | 5189 |
| 5194 | IV | 827 | 0 | 0 | 283 | 0 | 0 | Mono | Mono | | 22817840 | M9365 | BT | | 5194 |
| 5216 | IV | 827 | 0 | 0 | 266 | 0 | 17 | Mono | Mono | | 22866729 | M9366 | BT | | 5216 |
| 5219 | IV | 827 | 0 | 0 | 283 | 0 | 0 | Mono | Mono | | 22879049 | M9367 | BT | | 5219 |
| 5251 | IV | 827 | 0 | 0 | 283 | 0 | 0 | Mono | Mono | | 22931502 | M9370 | BT | | 5251 |
| 5282 | IV | 821 | 0 | 6 | 281 | 0 | 2 | Mono | Mono | | 23005081 | M9372 | BT | | 5282 |
| 5314 | IV | 826 | 0 | 1 | 283 | 0 | 0 | Mono | Mono | | 23070865 | M9376 | BT | | 5314 |
| 5331 | IV | 825 | 0 | 2 | 283 | 0 | 0 | Mono | Mono | | 23113295 | M9378 | BT | | 5331 |
| 5334 | IV | 827 | 0 | 0 | 283 | 0 | 0 | Mono | Mono | | 23117154 | PF1243 | BT | | 5334 |
| 5371 | IV | 827 | 0 | 0 | 283 | 0 | 0 | Mono | Mono | | 23199069 | Z40409 | BT | | 5371 |
| 5385 | IV | 827 | 0 | 0 | 283 | 0 | 0 | Mono | Mono | | 23244049 | M9379 | BT | | 5385 |
| 5393 | IV | 826 | 0 | 1 | 283 | 0 | 0 | Mono | Mono | | 23256074 | M9380 | BT | | 5393 |
| 5514 | IV | 827 | 0 | 0 | 283 | 0 | 0 | Mono | Mono | | 23558260 | M9393 | BT | | 5514 |
| 5516 | IV | 826 | 0 | 1 | 283 | 0 | 0 | Mono | Mono | | 23565114 | M9394 | BT | | 5516 |
| 5541 | IV | 0 | 803 | 24 | 0 | 275 | 8 | Mono | Mono | | 23652150 | Z40413 | BT | | 5541 |
| 5554 | IV | 827 | 0 | 0 | 283 | 0 | 0 | Mono | Mono | | 23757465 | M9397 | BT | | 5554 |
| 5651 | IV | 825 | 0 | 2 | 283 | 0 | 0 | Mono | Mono | | 24377975 | M9406 | BT | | 5651 |
| 77 | IV | 827 | 0 | 0 | 283 | 0 | 0 | Mono | Mono | | 2734854 | RPS4Y711 | C | | 77 |
| 111 | IV | 778 | 0 | 49 | 270 | 0 | 13 | Mono | Mono | | 2803297 | Y1763 | C | | 111 |
| 773 | IV | 827 | 0 | 0 | 283 | 0 | 0 | Mono | Mono | | 7159950 | F917 | C | | 773 |
| 1188 | IV | 827 | 0 | 0 | 283 | 0 | 0 | Mono | Mono | | 8145871 | F1118 | C | | 1188 |
| 1355 | IV | 825 | 0 | 2 | 283 | 0 | 0 | Mono | Mono | | 8454895 | V2015 | C | | 1355 |
| 1390 | IV | 827 | 0 | 0 | 283 | 0 | 0 | Mono | Mono | | 8482631 | V2099 | C | | 1390 |
| 1433 | IV | 826 | 0 | 1 | 283 | 0 | 0 | Mono | Mono | | 8537273 | V2200 | C | | 1433 |
| 1450 | IV | 827 | 0 | 0 | 283 | 0 | 0 | Mono | Mono | | 8560645 | V2258 | C | | 1450 |
| 1512 | IV | 825 | 0 | 2 | 281 | 0 | 2 | Mono | Mono | | 8640245 | F1367 | C | | 1512 |
| 1537 | IV | 827 | 0 | 0 | 283 | 0 | 0 | Mono | Mono | | 8685038 | P325 | C | | 1537 |
| 1656 | IV | 827 | 0 | 0 | 283 | 0 | 0 | Mono | Mono | | 9118085 | F1490 | C | | 1656 |
| 2053 | IV | 827 | 0 | 0 | 283 | 0 | 0 | Mono | Mono | | 14206892 | F1727 | C | | 2053 |
| 2078 | IV | 795 | 0 | 32 | 274 | 0 | 9 | Mono | Mono | | 14263271 | F1743 | C | | 2078 |
| 2196 | IV | 825 | 0 | 2 | 282 | 0 | 1 | Mono | Mono | | 14603298 | F1804 | C | | 2196 |
| 2394 | IV | 824 | 0 | 3 | 283 | 0 | 0 | Mono | Mono | | 14954047 | F1871 | C | | 2394 |
| 2500 | IV | 824 | 0 | 3 | 281 | 0 | 2 | Mono | Mono | | 15060025 | V2801 | C | | 2500 |
| 2520 | IV | 827 | 0 | 0 | 283 | 0 | 0 | Mono | Mono | | 15097043 | V2823 | C | | 2520 |
| 2642 | IV | 826 | 0 | 1 | 283 | 0 | 0 | Mono | Mono | | 15437564 | M216 | C | | 2642 |
| 2912 | IV | 555 | 0 | 272 | 196 | 0 | 87 | Mono | Mono | | 16035637 | F2067 | C | | 2912 |

| | | | M9+ | | | M9- | | | | | Build 37 | | ISOGG | | |
|---|---|---|---|---|---|---|---|---|---|---|---|---|---|---|---|
| List | Stage | 0 | 1 | . | 0 | 1 | . | M9? | Clade(s) | Prob. | Position | Name | Subgroup Name | Amb. | List |
| 3114 | IV | 827 | 0 | 0 | 283 | 0 | 0 | Mono | Mono | | 16595733 | F2211 | C | | 3114 |
| 3172 | IV | 825 | 0 | 2 | 283 | 0 | 0 | Mono | Mono | | 16768835 | F2258 | C | | 3172 |
| 3239 | IV | 827 | 0 | 0 | 283 | 0 | 0 | Mono | Mono | | 16871809 | F2305 | C | | 3239 |
| 3398 | IV | 827 | 0 | 0 | 283 | 0 | 0 | Mono | Mono | | 17286006 | P324 | C | | 3398 |
| 3419 | IV | 827 | 0 | 0 | 283 | 0 | 0 | Mono | Mono | | 17328425 | F2446 | C | | 3419 |
| 3642 | IV | 827 | 0 | 0 | 283 | 0 | 0 | Mono | Mono | | 17820514 | F2606 | C | | 3642 |
| 3646 | IV | 826 | 0 | 1 | 282 | 0 | 1 | Mono | Mono | | 17833394 | F2607 | C | | 3646 |
| 3712 | IV | 827 | 0 | 0 | 283 | 0 | 0 | Mono | Mono | | 17947542 | V77 | C | | 3712 |
| 3716 | IV | 827 | 0 | 0 | 283 | 0 | 0 | Mono | Mono | | 17957903 | F3712 | C | | 3716 |
| 3741 | IV | 827 | 0 | 0 | 283 | 0 | 0 | Mono | Mono | | 18030738 | F2678 | C | | 3741 |
| 3880 | IV | 827 | 0 | 0 | 283 | 0 | 0 | Mono | Mono | | 18572332 | F2774 | C | | 3880 |
| 3902 | IV | 825 | 0 | 2 | 282 | 0 | 1 | Mono | Mono | | 18613753 | V3332 | C | | 3902 |
| 3979 | IV | 822 | 0 | 5 | 280 | 0 | 3 | Mono | Mono | | 18787125 | V3497 | C | | 3979 |
| 4001 | IV | 827 | 0 | 0 | 283 | 0 | 0 | Mono | Mono | | 18832816 | V3533 | C | | 4001 |
| 4009 | IV | 827 | 0 | 0 | 283 | 0 | 0 | Mono | Mono | | 18843140 | V3554 | C | | 4009 |
| 4026 | IV | 821 | 0 | 6 | 280 | 0 | 3 | Mono | Mono | | 18890063 | V3578 | C | | 4026 |
| 4042 | IV | 827 | 0 | 0 | 282 | 0 | 1 | Mono | Mono | | 18917306 | V3600 | C | | 4042 |
| 4071 | II | 827 | 0 | 0 | 281 | 2 | 0 | M9- | 7 | | 18964479 | V3643 | C | w | 4071 |
| 5002 | IV | 827 | 0 | 0 | 283 | 0 | 0 | Mono | Mono | | 22208408 | F3311 | C | | 5002 |
| 5253 | IV | 826 | 0 | 1 | 282 | 0 | 1 | Mono | Mono | | 22937380 | F3719 | C | | 5253 |
| 5287 | IV | 825 | 0 | 2 | 283 | 0 | 0 | Mono | Mono | | 23020085 | F3388 | C | | 5287 |
| 5295 | IV | 827 | 0 | 0 | 283 | 0 | 0 | Mono | Mono | | 23031841 | F3395 | C | | 5295 |
| 5312 | IV | 827 | 0 | 0 | 282 | 0 | 1 | Mono | Mono | | 23067770 | F3400 | C | | 5312 |
| 5095 | IV | 827 | 0 | 0 | 283 | 0 | 0 | Mono | Mono | | 22669220 | PK2 | C(Notes) | | 5095 |
| 5292 | IV | 827 | 0 | 0 | 283 | 0 | 0 | Mono | Mono | | 23023974 | Z1426 | C1 | | 5292 |
| 5234 | IV | 827 | 0 | 0 | 283 | 0 | 0 | Mono | Mono | | 22914979 | CTS11043 | C1a | | 5234 |
| 821 | IV | 827 | 0 | 0 | 283 | 0 | 0 | Mono | Mono | | 7291534 | M8 | C1a1 | | 821 |
| 2316 | IV | 825 | 0 | 2 | 283 | 0 | 0 | Mono | Mono | | 14870234 | V2645 | C1a1~ | | 2316 |
| 2439 | IV | 826 | 0 | 1 | 283 | 0 | 0 | Mono | Mono | | 15017505 | M240 | C1a1~ | | 2439 |
| 4774 | IV | 827 | 0 | 0 | 281 | 0 | 2 | Mono | Mono | | 21866491 | M105 | C1a1~ | | 4774 |
| 5527 | IV | 827 | 0 | 0 | 283 | 0 | 0 | Mono | Mono | | 23604196 | Z7988 | C1a1a1b | | 5527 |
| 107 | IV | 827 | 0 | 0 | 283 | 0 | 0 | Mono | Mono | | 2795910 | Z28000 | C1a2 | | 107 |
| 116 | IV | 827 | 0 | 0 | 283 | 0 | 0 | Mono | Mono | | 2807426 | Z28001 | C1a2 | | 116 |
| 554 | IV | 827 | 0 | 0 | 283 | 0 | 0 | Mono | Mono | | 6845955 | V20 | C1a2 | | 554 |
| 617 | IV | 825 | 0 | 2 | 283 | 0 | 0 | Mono | Mono | | 6909957 | V86 | C1a2 | | 617 |
| 944 | IV | 827 | 0 | 0 | 283 | 0 | 0 | Mono | Mono | | 7647152 | Z28783 | C1a2 | | 944 |
| 964 | IV | 827 | 0 | 0 | 283 | 0 | 0 | Mono | Mono | | 7676565 | Z28784 | C1a2 | | 964 |
| 1044 | IV | 827 | 0 | 0 | 283 | 0 | 0 | Mono | Mono | | 7811623 | Z28789 | C1a2 | | 1044 |
| 1103 | IV | 821 | 0 | 6 | 283 | 0 | 0 | Mono | Mono | | 7938613 | Z28792 | C1a2 | | 1103 |
| 1110 | IV | 827 | 0 | 0 | 283 | 0 | 0 | Mono | Mono | | 7953413 | Z29320 | C1a2 | | 1110 |
| 1362 | IV | 827 | 0 | 0 | 283 | 0 | 0 | Mono | Mono | | 8462000 | Z28805 | C1a2 | | 1362 |
| 1409 | IV | 827 | 0 | 0 | 283 | 0 | 0 | Mono | Mono | | 8498791 | Z28808 | C1a2 | | 1409 |
| 1454 | IV | 826 | 0 | 1 | 283 | 0 | 0 | Mono | Mono | | 8564465 | Z28811 | C1a2 | | 1454 |
| 1520 | IV | 826 | 0 | 1 | 283 | 0 | 0 | Mono | Mono | | 8646282 | Z28814 | C1a2 | | 1520 |
| 1521 | IV | 797 | 0 | 30 | 276 | 0 | 7 | Mono | Mono | | 8647711 | Z28815 | C1a2 | | 1521 |
| 2251 | IV | 827 | 0 | 0 | 283 | 0 | 0 | Mono | Mono | | 14774539 | Z29322 | C1a2 | | 2251 |
| 2282 | IV | 827 | 0 | 0 | 283 | 0 | 0 | Mono | Mono | | 14820317 | Z29323 | C1a2 | | 2282 |
| 2859 | IV | 827 | 0 | 0 | 283 | 0 | 0 | Mono | Mono | | 15831202 | Z28863 | C1a2 | | 2859 |
| 2865 | IV | 305 | 0 | 522 | 90 | 0 | 193 | Mono | Mono | | 15852360 | Z28864 | C1a2 | | 2865 |
| 2887 | IV | 827 | 0 | 0 | 283 | 0 | 0 | Mono | Mono | | 15933881 | Z28867 | C1a2 | | 2887 |
| 3325 | IV | 827 | 0 | 0 | 283 | 0 | 0 | Mono | Mono | | 17048894 | M1009 | C1a2 | | 3325 |
| 3883 | IV | 827 | 0 | 0 | 283 | 0 | 0 | Mono | Mono | | 18576889 | Z28932 | C1a2 | | 3883 |
| 3934 | IV | 827 | 0 | 0 | 283 | 0 | 0 | Mono | Mono | | 18679308 | Z28938 | C1a2 | | 3934 |
| 4088 | IV | 826 | 0 | 1 | 283 | 0 | 0 | Mono | Mono | | 19024253 | Z28946 | C1a2 | | 4088 |
| 4092 | IV | 825 | 0 | 2 | 280 | 0 | 3 | Mono | Mono | | 19035347 | Z28948 | C1a2 | | 4092 |
| 4130 | IV | 827 | 0 | 0 | 283 | 0 | 0 | Mono | Mono | | 19108008 | Z28951 | C1a2 | | 4130 |
| 4301 | IV | 827 | 0 | 0 | 283 | 0 | 0 | Mono | Mono | | 19490323 | Z29324 | C1a2 | | 4301 |
| 1347 | IV | 827 | 0 | 0 | 283 | 0 | 0 | Mono | Mono | | 8445233 | Z28803 | C1a2a | | 1347 |
| 1453 | IV | 827 | 0 | 0 | 283 | 0 | 0 | Mono | Mono | | 8563946 | Z28810 | C1a2a | | 1453 |
| 2553 | IV | 825 | 0 | 2 | 283 | 0 | 0 | Mono | Mono | | 15222486 | Z28857 | C1a2a | | 2553 |
| 3879 | IV | 827 | 0 | 0 | 283 | 0 | 0 | Mono | Mono | | 18572315 | V3287 | C1a2a | | 3879 |
| 4037 | IV | 824 | 0 | 3 | 283 | 0 | 0 | Mono | Mono | | 18908253 | Z28945 | C1a2a | | 4037 |
| 56 | IV | 827 | 0 | 0 | 283 | 0 | 0 | Mono | Mono | | 2717985 | Z28767 | C1a2a1 | | 56 |
| 879 | IV | 827 | 0 | 0 | 283 | 0 | 0 | Mono | Mono | | 7543039 | Z28781 | C1a2a1 | | 879 |
| 909 | IV | 827 | 0 | 0 | 283 | 0 | 0 | Mono | Mono | | 7589937 | V222 | C1a2a1 | | 909 |
| 1392 | IV | 827 | 0 | 0 | 283 | 0 | 0 | Mono | Mono | | 8482793 | Z28807 | C1a2a1 | | 1392 |
| 1464 | IV | 827 | 0 | 0 | 283 | 0 | 0 | Mono | Mono | | 8578064 | Z28812 | C1a2a1 | | 1464 |
| 3896 | IV | 827 | 0 | 0 | 283 | 0 | 0 | Mono | Mono | | 18600043 | Z28934 | C1a2a1 | | 3896 |
| 4017 | IV | 827 | 0 | 0 | 281 | 0 | 2 | Mono | Mono | | 18868114 | Z28935 | C1a2a1 | | 4017 |
| 4190 | IV | 827 | 0 | 0 | 283 | 0 | 0 | Mono | Mono | | 19219752 | Z28955 | C1a2a1 | | 4190 |
| 4192 | IV | 827 | 0 | 0 | 283 | 0 | 0 | Mono | Mono | | 19224173 | Z28956 | C1a2a1 | | 4192 |
| 1517 | IV | 824 | 0 | 3 | 283 | 0 | 0 | Mono | Mono | | 8643365 | F1370 | C1b | | 1517 |
| 4752 | IV | 826 | 0 | 1 | 283 | 0 | 0 | Mono | Mono | | 21815765 | Z16480 | C1b | | 4752 |
| 183 | IV | 826 | 0 | 1 | 283 | 0 | 0 | Mono | Mono | | 2889366 | K43 | C1b1a1a1 | | 183 |
| 2302 | IV | 827 | 0 | 0 | 283 | 0 | 0 | Mono | Mono | | 14850853 | P92 | C1b1a1a1 | | 2302 |
| 103 | IV | 827 | 0 | 0 | 283 | 0 | 0 | Mono | Mono | | 2784458 | K41 | C1b1a1a1a | | 103 |
| 817 | IV | 827 | 0 | 0 | 283 | 0 | 0 | Mono | Mono | | 7279954 | K107 | C1b1a1a1a1a | | 817 |
| 1290 | IV | 827 | 0 | 0 | 283 | 0 | 0 | Mono | Mono | | 8367491 | K131 | C1b1a1a1a1a | | 1290 |
| 106 | IV | 827 | 0 | 0 | 283 | 0 | 0 | Mono | Mono | | 2794557 | K42 | C1b1a1a1a1a1 | | 106 |
| 5568 | IV | 824 | 0 | 3 | 282 | 0 | 1 | Mono | Mono | | 23792572 | Z12518 | C1b1a1a1a1a1a | | 5568 |
| 185 | IV | 825 | 0 | 2 | 283 | 0 | 0 | Mono | Mono | | 2890490 | Z5900 | C1b1a1a2a1 | | 185 |
| 80 | IV | 827 | 0 | 0 | 283 | 0 | 0 | Mono | Mono | | 2737462 | F725 | C1b1a2b | | 80 |
| 125 | IV | 824 | 0 | 3 | 283 | 0 | 0 | Mono | Mono | | 2816523 | F736 | C1b1a2b | | 125 |
| 476 | IV | 827 | 0 | 0 | 283 | 0 | 0 | Mono | Mono | | 6668360 | F796 | C1b1a2b | | 476 |
| 515 | IV | 825 | 0 | 2 | 282 | 0 | 1 | Mono | Mono | | 6741680 | F821 | C1b1a2b | | 515 |
| 650 | IV | 827 | 0 | 0 | 283 | 0 | 0 | Mono | Mono | | 6932877 | F873 | C1b1a2b | | 650 |
| 651 | IV | 827 | 0 | 0 | 283 | 0 | 0 | Mono | Mono | | 6932987 | F4067 | C1b1a2b | | 651 |
| 668 | IV | 827 | 0 | 0 | 283 | 0 | 0 | Mono | Mono | | 6951737 | F884 | C1b1a2b | | 668 |
| 882 | IV | 827 | 0 | 0 | 282 | 0 | 1 | Mono | Mono | | 7545688 | F988 | C1b1a2b | | 882 |
| 941 | IV | 826 | 0 | 1 | 283 | 0 | 0 | Mono | Mono | | 7643777 | F1033 | C1b1a2b | | 941 |
| 1125 | IV | 823 | 0 | 4 | 282 | 0 | 1 | Mono | Mono | | 7997470 | F3859 | C1b1a2b | | 1125 |
| 1170 | IV | 823 | 0 | 4 | 283 | 0 | 0 | Mono | Mono | | 8117297 | F3860 | C1b1a2b | | 1170 |
| 1176 | IV | 827 | 0 | 0 | 283 | 0 | 0 | Mono | Mono | | 8128296 | F1109 | C1b1a2b | | 1176 |
| 1299 | IV | 827 | 0 | 0 | 283 | 0 | 0 | Mono | Mono | | 8381922 | F3863 | C1b1a2b | | 1299 |
| 1359 | IV | 827 | 0 | 0 | 283 | 0 | 0 | Mono | Mono | | 8460176 | F1220 | C1b1a2b | | 1359 |
| 1408 | IV | 827 | 0 | 0 | 283 | 0 | 0 | Mono | Mono | | 8498515 | F1259 | C1b1a2b | | 1408 |

| | | | M9+ | | | M9- | | | | | Build 37 | | ISOGG | | |
|---|---|---|---|---|---|---|---|---|---|---|---|---|---|---|---|
| List | Stage | 0 | 1 | . | 0 | 1 | . | M9? | Clade(s) | Prob. | Position | Name | Subgroup Name | Amb. | List |
| 1472 | IV | 826 | 0 | 1 | 283 | 0 | 0 | Mono | Mono | | 8589831 | F1331 | C1b1a2b | | 1472 |
| 1494 | IV | 827 | 0 | 0 | 283 | 0 | 0 | Mono | Mono | | 8613151 | F4116 | C1b1a2b | | 1494 |
| 1603 | IV | 825 | 0 | 2 | 283 | 0 | 0 | Mono | Mono | | 8862487 | F3882 | C1b1a2b | | 1603 |
| 1984 | IV | 827 | 0 | 0 | 283 | 0 | 0 | Mono | Mono | | 14065152 | F1686 | C1b1a2b | | 1984 |
| 2083 | IV | 827 | 0 | 0 | 283 | 0 | 0 | Mono | Mono | | 14275268 | F3915 | C1b1a2b | | 2083 |
| 2130 | IV | 815 | 0 | 12 | 281 | 0 | 2 | Mono | Mono | | 14397271 | F1782 | C1b1a2b | | 2130 |
| 2226 | IV | 827 | 0 | 0 | 283 | 0 | 0 | Mono | Mono | | 14697887 | F3922 | C1b1a2b | | 2226 |
| 2551 | IV | 825 | 0 | 2 | 283 | 0 | 0 | Mono | Mono | | 15219892 | F1914 | C1b1a2b | | 2551 |
| 2826 | IV | 826 | 0 | 1 | 281 | 0 | 2 | Mono | Mono | | 15764766 | F1990 | C1b1a2b | | 2826 |
| 2864 | IV | 827 | 0 | 0 | 283 | 0 | 0 | Mono | Mono | | 15847267 | F2010 | C1b1a2b | | 2864 |
| 2877 | IV | 827 | 0 | 0 | 283 | 0 | 0 | Mono | Mono | | 15899755 | F2031 | C1b1a2b | | 2877 |
| 2944 | II | 826 | 1 | 0 | 283 | 0 | 0 | M9+ | NU | | 16201991 | F2085 | C1b1a2b | w | 2944 |
| 2957 | IV | 825 | 0 | 2 | 283 | 0 | 0 | Mono | Mono | | 16224533 | F2098 | C1b1a2b | | 2957 |
| 3017 | IV | 827 | 0 | 0 | 283 | 0 | 0 | Mono | Mono | | 16368478 | F2143 | C1b1a2b | | 3017 |
| 3079 | IV | 821 | 0 | 6 | 282 | 0 | 1 | Mono | Mono | | 16505425 | F2181 | C1b1a2b | | 3079 |
| 3080 | IV | 827 | 0 | 0 | 282 | 0 | 1 | Mono | Mono | | 16508917 | F2182 | C1b1a2b | | 3080 |
| 3159 | IV | 827 | 0 | 0 | 283 | 0 | 0 | Mono | Mono | | 16736046 | F2248 | C1b1a2b | | 3159 |
| 3225 | IV | 826 | 0 | 1 | 283 | 0 | 0 | Mono | Mono | | 16854044 | F2298 | C1b1a2b | | 3225 |
| 3287 | IV | 827 | 0 | 0 | 283 | 0 | 0 | Mono | Mono | | 16947886 | F2333 | C1b1a2b | | 3287 |
| 3290 | IV | 827 | 0 | 0 | 283 | 0 | 0 | Mono | Mono | | 16951198 | F2335 | C1b1a2b | | 3290 |
| 3338 | IV | 827 | 0 | 0 | 283 | 0 | 0 | Mono | Mono | | 17110718 | F2381 | C1b1a2b | | 3338 |
| 3456 | IV | 827 | 0 | 0 | 283 | 0 | 0 | Mono | Mono | | 17421589 | F2476 | C1b1a2b | | 3456 |
| 3558 | IV | 827 | 0 | 0 | 283 | 0 | 0 | Mono | Mono | | 17655987 | F2545 | C1b1a2b | | 3558 |
| 3559 | IV | 827 | 0 | 0 | 283 | 0 | 0 | Mono | Mono | | 17656418 | F3966 | C1b1a2b | | 3559 |
| 3649 | IV | 827 | 0 | 0 | 283 | 0 | 0 | Mono | Mono | | 17840955 | F2611 | C1b1a2b | | 3649 |
| 3702 | IV | 813 | 0 | 14 | 279 | 0 | 4 | Mono | Mono | | 17934375 | F2641 | C1b1a2b | | 3702 |
| 3941 | IV | 826 | 0 | 1 | 283 | 0 | 0 | Mono | Mono | | 18710779 | F4265 | C1b1a2b | | 3941 |
| 3955 | IV | 814 | 0 | 13 | 280 | 0 | 3 | Mono | Mono | | 18744132 | F2832 | C1b1a2b | | 3955 |
| 3980 | IV | 827 | 0 | 0 | 283 | 0 | 0 | Mono | Mono | | 18787373 | F2848 | C1b1a2b | | 3980 |
| 4012 | IV | 821 | 0 | 6 | 282 | 0 | 1 | Mono | Mono | | 18852625 | F2873 | C1b1a2b | | 4012 |
| 4079 | IV | 827 | 0 | 0 | 283 | 0 | 0 | Mono | Mono | | 18999411 | F2915 | C1b1a2b | | 4079 |
| 4140 | IV | 827 | 0 | 0 | 283 | 0 | 0 | Mono | Mono | | 19135700 | F2946 | C1b1a2b | | 4140 |
| 4158 | IV | 827 | 0 | 0 | 283 | 0 | 0 | Mono | Mono | | 19161668 | F2959 | C1b1a2b | | 4158 |
| 4230 | IV | 826 | 0 | 1 | 283 | 0 | 0 | Mono | Mono | | 19285282 | F3007 | C1b1a2b | | 4230 |
| 4356 | IV | 826 | 0 | 1 | 283 | 0 | 0 | Mono | Mono | | 21052319 | F3107 | C1b1a2b | | 4356 |
| 4369 | IV | 827 | 0 | 0 | 283 | 0 | 0 | Mono | Mono | | 21083886 | F3114 | C1b1a2b | | 4369 |
| 4376 | IV | 827 | 0 | 0 | 283 | 0 | 0 | Mono | Mono | | 21097978 | F3120 | C1b1a2b | | 4376 |
| 4396 | IV | 827 | 0 | 0 | 283 | 0 | 0 | Mono | Mono | | 21148137 | F3137 | C1b1a2b | | 4396 |
| 4601 | IV | 826 | 0 | 1 | 283 | 0 | 0 | Mono | Mono | | 21628445 | F3219 | C1b1a2b | | 4601 |
| 4662 | IV | 827 | 0 | 0 | 283 | 0 | 0 | Mono | Mono | | 21730167 | F3244 | C1b1a2b | | 4662 |
| 5254 | IV | 826 | 0 | 1 | 283 | 0 | 0 | Mono | Mono | | 22940275 | F3378.2 | C1b1a2b | | 5254 |
| 5255 | IV | 825 | 0 | 2 | 283 | 0 | 0 | Mono | Mono | | 22942170 | F3381 | C1b1a2b | | 5255 |
| 5259 | IV | 827 | 0 | 0 | 283 | 0 | 0 | Mono | Mono | | 22946162 | F4020 | C1b1a2b | | 5259 |
| 1320 | IV | 826 | 0 | 1 | 283 | 0 | 0 | Mono | Mono | | 8418520 | B465 | C1b1a2b1a | | 1320 |
| 654 | IV | 827 | 0 | 0 | 283 | 0 | 0 | Mono | Mono | | 6933080 | SK1010 | C1b2 | | 654 |
| 1437 | IV | 827 | 0 | 0 | 283 | 0 | 0 | Mono | Mono | | 8541673 | Z31886 | C1b2 | | 1437 |
| 2216 | IV | 824 | 0 | 3 | 283 | 0 | 0 | Mono | Mono | | 14655853 | SK1013 | C1b2 | | 2216 |
| 2997 | IV | 827 | 0 | 0 | 283 | 0 | 0 | Mono | Mono | | 16330869 | Z31878 | C1b2 | | 2997 |
| 4039 | IV | 827 | 0 | 0 | 283 | 0 | 0 | Mono | Mono | | 18913123 | PH4087 | C1b2 | | 4039 |
| 4186 | IV | 827 | 0 | 0 | 282 | 0 | 1 | Mono | Mono | | 19215759 | PH4346 | C1b2 | | 4186 |
| 5278 | IV | 827 | 0 | 0 | 283 | 0 | 0 | Mono | Mono | | 22995305 | Z31881 | C1b2 | | 5278 |
| 5323 | IV | 826 | 0 | 1 | 283 | 0 | 0 | Mono | Mono | | 23081529 | PH5328 | C1b2 | | 5323 |
| 5727 | IV | 827 | 0 | 0 | 283 | 0 | 0 | Mono | Mono | | 28767538 | Z31884 | C1b2 | | 5727 |
| 4687 | IV | 827 | 0 | 0 | 283 | 0 | 0 | Mono | Mono | | 21742158 | M38 | C1b2a | | 4687 |
| 2729 | IV | 827 | 0 | 0 | 283 | 0 | 0 | Mono | Mono | | 15576203 | M208 | C1b2a1 | | 2729 |
| 172 | IV | 827 | 0 | 0 | 283 | 0 | 0 | Mono | Mono | | 2877479 | M347 | C1b2b | | 172 |
| 2727 | IV | 826 | 0 | 1 | 283 | 0 | 0 | Mono | Mono | | 15575780 | M210 | C1b2b1 | | 2727 |
| 2162 | IV | 827 | 0 | 0 | 283 | 0 | 0 | Mono | Mono | | 14488952 | P55 | C1b3~ | | 2162 |
| 2640 | IV | 827 | 0 | 0 | 283 | 0 | 0 | Mono | Mono | | 15437333 | M217 | C2 | | 2640 |
| 4086 | IV | 827 | 0 | 0 | 283 | 0 | 0 | Mono | Mono | | 19020366 | Z1453 | C2 | | 4086 |
| 5117 | IV | 827 | 0 | 0 | 282 | 0 | 1 | Mono | Mono | | 22711592 | L1373 | C2a | | 5117 |
| 2675 | I | 796 | 31 | 0 | 272 | 11 | 0 | BOTH | | PI | 15491978 | FGC22105 | C2a1a1a1a1a1~ | w | 2675 |
| 3156 | III | 825 | 0 | 2 | 245 | 29 | 9 | M9- | 1 – 10 | MD | 16723271 | FT59 | C2a1a1b1a2b~ | w | 3156 |
| 4689 | IV | 827 | 0 | 0 | 283 | 0 | 0 | Mono | Mono | | 21749881 | M48 | C2a1a2 | | 4689 |
| 4702 | IV | 827 | 0 | 0 | 283 | 0 | 0 | Mono | Mono | | 21759138 | M77 | C2a1a2a | | 4702 |
| 4862 | IV | 827 | 0 | 0 | 283 | 0 | 0 | Mono | Mono | | 21905917 | M86 | C2a1a2a | | 4862 |
| 867 | IV | 826 | 0 | 1 | 283 | 0 | 0 | Mono | Mono | | 7527930 | IMS-JST002613- | C2b1 | | 867 |
| 3651 | IV | 827 | 0 | 0 | 283 | 0 | 0 | Mono | Mono | | 17844001 | Z1338 | C2b1 | | 3651 |
| 4705 | IV | 827 | 0 | 0 | 283 | 0 | 0 | Mono | Mono | | 21761737 | Page12 | C2b1 | | 4705 |
| 1038 | IV | 827 | 0 | 0 | 283 | 0 | 0 | Mono | Mono | | 7801788 | Z1300 | C2b1a | | 1038 |
| 2589 | IV | 827 | 0 | 0 | 283 | 0 | 0 | Mono | Mono | | 15339328 | Z1301 | C2b1a | | 2589 |
| 2811 | IV | 827 | 0 | 0 | 279 | 0 | 4 | Mono | Mono | | 15724326 | CTS4590 | C2b1a | | 2811 |
| 4183 | IV | 827 | 0 | 0 | 283 | 0 | 0 | Mono | Mono | | 19207887 | F2982 | C2b1a | | 4183 |
| 87 | IV | 827 | 0 | 0 | 283 | 0 | 0 | Mono | Mono | | 2750408 | M407 | C2b1a1a1a | | 87 |
| 1581 | IV | 827 | 0 | 0 | 283 | 0 | 0 | Mono | Mono | | 8804321 | F3880 | C2b1a2 | | 1581 |
| 1458 | IV | 827 | 0 | 0 | 283 | 0 | 0 | Mono | Mono | | 8571457 | F1319 | C2b1a2a | | 1458 |
| 2974 | IV | 824 | 0 | 3 | 280 | 0 | 3 | Mono | Mono | | 16262392 | F3777 | C2b1a2a1 | | 2974 |
| 3929 | IV | 826 | 0 | 1 | 283 | 0 | 0 | Mono | Mono | | 18666927 | Z2201.2 | C2b1a2a1 | | 3929 |
| 585 | IV | 827 | 0 | 0 | 283 | 0 | 0 | Mono | Mono | | 6875257 | F845 | C2b1b | | 585 |
| 3157 | IV | 827 | 0 | 0 | 283 | 0 | 0 | Mono | Mono | | 16734196 | K548 | C2b1b1 | | 3157 |
| 5206 | IV | 827 | 0 | 0 | 283 | 0 | 0 | Mono | Mono | | 22836292 | CTS10923 | C2b1b1 | | 5206 |
| 2165 | IV | 827 | 0 | 0 | 283 | 0 | 0 | Mono | Mono | | 14495229 | Z12284^ | C2b1b1a1 | | 2165 |
| 4855 | IV | 827 | 0 | 0 | 283 | 0 | 0 | Mono | Mono | | 21902505 | M93 | C2b1b1b4~ | | 4855 |
| 2043 | I | 827 | 0 | 0 | 246 | 37 | 0 | M9- | 1 – 10 | | 14197867 | PF2587 | CF | | 2043 |
| 2544 | I | 827 | 0 | 0 | 246 | 37 | 0 | M9- | 1 – 10 | | 15203676 | PF2668 | CF | | 2544 |
| 3233 | III | 826 | 0 | 1 | 246 | 37 | 0 | M9- | 1 – 10 | | 16863259 | PF2697 | CF | | 3233 |
| 74 | IV | 825 | 0 | 2 | 283 | 0 | 0 | Mono | Mono | | 2733618 | M8948 | CT | | 74 |
| 83 | IV | 826 | 0 | 1 | 283 | 0 | 0 | Mono | Mono | | 2744386 | M5576 | CT | | 83 |
| 517 | IV | 826 | 0 | 1 | 283 | 0 | 0 | Mono | Mono | | 6744622 | M5583 | CT | | 517 |
| 545 | IV | 827 | 0 | 0 | 283 | 0 | 0 | Mono | Mono | | 6831340 | M5584 | CT | | 545 |
| 577 | IV | 827 | 0 | 0 | 283 | 0 | 0 | Mono | Mono | | 6864267 | V9 | CT | | 577 |
| 698 | IV | 827 | 0 | 0 | 283 | 0 | 0 | Mono | Mono | | 6991000 | PF210 | CT | | 698 |
| 712 | IV | 827 | 0 | 0 | 283 | 0 | 0 | Mono | Mono | | 7041647 | M5590 | CT | | 712 |
| 768 | IV | 827 | 0 | 0 | 282 | 0 | 1 | Mono | Mono | | 7153375 | PF223 | CT | | 768 |
| 840 | IV | 825 | 0 | 2 | 282 | 0 | 1 | Mono | Mono | | 7358315 | PF234 | CT | | 840 |
| 857 | IV | 827 | 0 | 0 | 283 | 0 | 0 | Mono | Mono | | 7416169 | M5600 | CT | | 857 |

| | | M9+ | | | M9- | | | | | | Build 37 | | ISOGG | | |
|---|---|---|---|---|---|---|---|---|---|---|---|---|---|---|---|
| List | Stage | 0 | 1 | . | 0 | 1 | . | M9? | Clade(s) | Prob. | Position | Name | Subgroup Name | Amb. | List |
| 871 | IV | 824 | 0 | 3 | 283 | 0 | 0 | Mono | Mono | | 7533511 | M5601 | CT | | 871 |
| 916 | IV | 827 | 0 | 0 | 283 | 0 | 0 | Mono | Mono | | 7597185 | V226 | CT | | 916 |
| 984 | IV | 827 | 0 | 0 | 282 | 0 | 1 | Mono | Mono | | 7707484 | M5607 | CT | | 984 |
| 995 | IV | 827 | 0 | 0 | 283 | 0 | 0 | Mono | Mono | | 7729324 | PF258 | CT | | 995 |
| 1007 | IV | 827 | 0 | 0 | 283 | 0 | 0 | Mono | Mono | | 7738840 | M5609 | CT | | 1007 |
| 1008 | IV | 827 | 0 | 0 | 283 | 0 | 0 | Mono | Mono | | 7740081 | M5610 | CT | | 1008 |
| 1028 | IV | 826 | 0 | 1 | 282 | 0 | 1 | Mono | Mono | | 7778691 | PF263 | CT | | 1028 |
| 1036 | IV | 827 | 0 | 0 | 283 | 0 | 0 | Mono | Mono | | 7796647 | M5613 | CT | | 1036 |
| 1076 | IV | 826 | 0 | 1 | 283 | 0 | 0 | Mono | Mono | | 7887815 | M5615 | CT | | 1076 |
| 1098 | IV | 827 | 0 | 0 | 283 | 0 | 0 | Mono | Mono | | 7921525 | PF272 | CT | | 1098 |
| 1117 | IV | 827 | 0 | 0 | 283 | 0 | 0 | Mono | Mono | | 7970612 | PF278 | CT | | 1117 |
| 1185 | IV | 779 | 0 | 48 | 274 | 0 | 9 | Mono | Mono | | 8138589 | M5625 | CT | | 1185 |
| 1196 | IV | 827 | 0 | 0 | 283 | 0 | 0 | Mono | Mono | | 8183439 | M5627 | CT | | 1196 |
| 1243 | IV | 827 | 0 | 0 | 283 | 0 | 0 | Mono | Mono | | 8262951 | M5629 | CT | | 1243 |
| 1309 | IV | 827 | 0 | 0 | 283 | 0 | 0 | Mono | Mono | | 8396636 | PF292 | CT | | 1309 |
| 1428 | IV | 827 | 0 | 0 | 283 | 0 | 0 | Mono | Mono | | 8526565 | M5632 | CT | | 1428 |
| 1439 | IV | 827 | 0 | 0 | 283 | 0 | 0 | Mono | Mono | | 8543804 | M5633 | CT | | 1439 |
| 1510 | IV | 827 | 0 | 0 | 283 | 0 | 0 | Mono | Mono | | 8635526 | M5636 | CT | | 1510 |
| 1713 | IV | 825 | 0 | 2 | 282 | 0 | 1 | Mono | Mono | | 9758726 | PF342 | CT | | 1713 |
| 1821 | IV | 827 | 0 | 0 | 283 | 0 | 0 | Mono | Mono | | 10045515 | Z17707 | CT | | 1821 |
| 1985 | IV | 827 | 0 | 0 | 283 | 0 | 0 | Mono | Mono | | 14065693 | M5649 | CT | | 1985 |
| 2013 | IV | 820 | 0 | 7 | 282 | 0 | 1 | Mono | Mono | | 14124435 | M5650 | CT | | 2013 |
| 2024 | IV | 827 | 0 | 0 | 283 | 0 | 0 | Mono | Mono | | 14149010 | M5651 | CT | | 2024 |
| 2025 | IV | 827 | 0 | 0 | 283 | 0 | 0 | Mono | Mono | | 14149520 | M5652 | CT | | 2025 |
| 2034 | IV | 827 | 0 | 0 | 283 | 0 | 0 | Mono | Mono | | 14172842 | M5653 | CT | | 2034 |
| 2092 | IV | 826 | 0 | 1 | 283 | 0 | 0 | Mono | Mono | | 14289530 | M5657 | CT | | 2092 |
| 2143 | IV | 827 | 0 | 0 | 283 | 0 | 0 | Mono | Mono | | 14439817 | CTS2711 | CT | | 2143 |
| 2155 | IV | 827 | 0 | 0 | 283 | 0 | 0 | Mono | Mono | | 14472971 | Y1526 | CT | | 2155 |
| 2221 | IV | 826 | 0 | 1 | 283 | 0 | 0 | Mono | Mono | | 14671991 | M5664 | CT | | 2221 |
| 2273 | IV | 827 | 0 | 0 | 283 | 0 | 0 | Mono | Mono | | 14813991 | PF1416 | CT | | 2273 |
| 2361 | IV | 825 | 0 | 2 | 283 | 0 | 0 | Mono | Mono | | 14917660 | M5669 | CT | | 2361 |
| 2382 | IV | 820 | 0 | 7 | 280 | 0 | 3 | Mono | Mono | | 14943290 | M5670 | CT | | 2382 |
| 2521 | IV | 827 | 0 | 0 | 283 | 0 | 0 | Mono | Mono | | 15097073 | M5671 | CT | | 2521 |
| 2594 | IV | 827 | 0 | 0 | 283 | 0 | 0 | Mono | Mono | | 15344987 | PF719 | CT | | 2594 |
| 2763 | IV | 827 | 0 | 0 | 283 | 0 | 0 | Mono | Mono | | 15614295 | M5679 | CT | | 2763 |
| 2767 | IV | 827 | 0 | 0 | 283 | 0 | 0 | Mono | Mono | | 15615637 | M5680 | CT | | 2767 |
| 2786 | IV | 826 | 0 | 1 | 283 | 0 | 0 | Mono | Mono | | 15662994 | M5682 | CT | | 2786 |
| 2838 | IV | 827 | 0 | 0 | 283 | 0 | 0 | Mono | Mono | | 15792531 | M5684 | CT | | 2838 |
| 2926 | IV | 827 | 0 | 0 | 283 | 0 | 0 | Mono | Mono | | 16131945 | Y1531 | CT | | 2926 |
| 2950 | IV | 811 | 0 | 16 | 279 | 0 | 4 | Mono | Mono | | 16203547 | M5690 | CT | | 2950 |
| 2973 | IV | 823 | 0 | 4 | 282 | 0 | 1 | Mono | Mono | | 16262390 | PF779 | CT | | 2973 |
| 2995 | IV | 827 | 0 | 0 | 283 | 0 | 0 | Mono | Mono | | 16325663 | M5692 | CT | | 2995 |
| 3007 | IV | 827 | 0 | 0 | 283 | 0 | 0 | Mono | Mono | | 16346371 | M5694 | CT | | 3007 |
| 3061 | IV | 827 | 0 | 0 | 283 | 0 | 0 | Mono | Mono | | 16465771 | M5697 | CT | | 3061 |
| 3173 | IV | 826 | 0 | 1 | 283 | 0 | 0 | Mono | Mono | | 16770207 | M5700 | CT | | 3173 |
| 3204 | IV | 827 | 0 | 0 | 283 | 0 | 0 | Mono | Mono | | 16822011 | M5705 | CT | | 3204 |
| 3219 | IV | 827 | 0 | 0 | 283 | 0 | 0 | Mono | Mono | | 16844116 | PF815 | CT | | 3219 |
| 3234 | IV | 826 | 0 | 1 | 283 | 0 | 0 | Mono | Mono | | 16865757 | M5707 | CT | | 3234 |
| 3241 | IV | 826 | 0 | 1 | 283 | 0 | 0 | Mono | Mono | | 16877090 | M5708 | CT | | 3241 |
| 3250 | IV | 827 | 0 | 0 | 283 | 0 | 0 | Mono | Mono | | 16887784 | M5709 | CT | | 3250 |
| 3335 | IV | 826 | 0 | 1 | 283 | 0 | 0 | Mono | Mono | | 17104433 | M5712 | CT | | 3335 |
| 3340 | IV | 827 | 0 | 0 | 283 | 0 | 0 | Mono | Mono | | 17114434 | M5713 | CT | | 3340 |
| 3359 | IV | 827 | 0 | 0 | 283 | 0 | 0 | Mono | Mono | | 17174326 | M5714 | CT | | 3359 |
| 3409 | IV | 827 | 0 | 0 | 283 | 0 | 0 | Mono | Mono | | 17301733 | PF840 | CT | | 3409 |
| 3498 | IV | 827 | 0 | 0 | 283 | 0 | 0 | Mono | Mono | | 17515027 | V52 | CT | | 3498 |
| 3501 | IV | 824 | 0 | 3 | 282 | 0 | 1 | Mono | Mono | | 17517587 | M5722 | CT | | 3501 |
| 3565 | IV | 827 | 0 | 0 | 283 | 0 | 0 | Mono | Mono | | 17670046 | M5726 | CT | | 3565 |
| 3593 | IV | 827 | 0 | 0 | 283 | 0 | 0 | Mono | Mono | | 17742131 | M5727 | CT | | 3593 |
| 3598 | IV | 826 | 0 | 1 | 283 | 0 | 0 | Mono | Mono | | 17751557 | M5729 | CT | | 3598 |
| 3680 | IV | 827 | 0 | 0 | 283 | 0 | 0 | Mono | Mono | | 17894575 | M5736 | CT | | 3680 |
| 3681 | IV | 827 | 0 | 0 | 283 | 0 | 0 | Mono | Mono | | 17897543 | PF892 | CT | | 3681 |
| 3687 | IV | 827 | 0 | 0 | 283 | 0 | 0 | Mono | Mono | | 17909505 | M5738 | CT | | 3687 |
| 3722 | IV | 827 | 0 | 0 | 283 | 0 | 0 | Mono | Mono | | 17975155 | Y1537 | CT | | 3722 |
| 3774 | IV | 826 | 0 | 1 | 283 | 0 | 0 | Mono | Mono | | 18077583 | M5743 | CT | | 3774 |
| 3815 | IV | 827 | 0 | 0 | 283 | 0 | 0 | Mono | Mono | | 18179043 | M5747 | CT | | 3815 |
| 3893 | IV | 827 | 0 | 0 | 283 | 0 | 0 | Mono | Mono | | 18592845 | M5748 | CT | | 3893 |
| 3906 | IV | 827 | 0 | 0 | 283 | 0 | 0 | Mono | Mono | | 18616467 | M5750 | CT | | 3906 |
| 3943 | IV | 825 | 0 | 2 | 283 | 0 | 0 | Mono | Mono | | 18715075 | PF937 | CT | | 3943 |
| 4053 | IV | 827 | 0 | 0 | 283 | 0 | 0 | Mono | Mono | | 18940202 | M5754 | CT | | 4053 |
| 4057 | IV | 827 | 0 | 0 | 283 | 0 | 0 | Mono | Mono | | 18948988 | PF951 | CT | | 4057 |
| 4066 | IV | 820 | 0 | 7 | 278 | 0 | 5 | Mono | Mono | | 18961874 | M5758 | CT | | 4066 |
| 4095 | IV | 827 | 0 | 0 | 283 | 0 | 0 | Mono | Mono | | 19039750 | M5762 | CT | | 4095 |
| 4108 | IV | 825 | 0 | 2 | 282 | 0 | 1 | Mono | Mono | | 19059200 | M5763 | CT | | 4108 |
| 4124 | IV | 827 | 0 | 0 | 283 | 0 | 0 | Mono | Mono | | 19090637 | M5764 | CT | | 4124 |
| 4162 | IV | 826 | 0 | 1 | 283 | 0 | 0 | Mono | Mono | | 19167672 | CTS9948 | CT | | 4162 |
| 4163 | IV | 827 | 0 | 0 | 283 | 0 | 0 | Mono | Mono | | 19169483 | PF970 | CT | | 4163 |
| 4184 | IV | 826 | 0 | 1 | 283 | 0 | 0 | Mono | Mono | | 19212465 | M5766 | CT | | 4184 |
| 4281 | IV | 827 | 0 | 0 | 283 | 0 | 0 | Mono | Mono | | 19421388 | M5770 | CT | | 4281 |
| 4373 | IV | 827 | 0 | 0 | 283 | 0 | 0 | Mono | Mono | | 21095622 | PF1016 | CT | | 4373 |
| 4428 | IV | 825 | 0 | 2 | 282 | 0 | 1 | Mono | Mono | | 21250559 | M5775 | CT | | 4428 |
| 4436 | IV | 827 | 0 | 0 | 283 | 0 | 0 | Mono | Mono | | 21258894 | PF1029 | CT | | 4436 |
| 4442 | IV | 827 | 0 | 0 | 283 | 0 | 0 | Mono | Mono | | 21267676 | M5777 | CT | | 4442 |
| 4474 | IV | 827 | 0 | 0 | 283 | 0 | 0 | Mono | Mono | | 21353965 | PF1040 | CT | | 4474 |
| 4510 | IV | 827 | 0 | 0 | 283 | 0 | 0 | Mono | Mono | | 21413635 | PF1046 | CT | | 4510 |
| 4514 | IV | 827 | 0 | 0 | 283 | 0 | 0 | Mono | Mono | | 21429988 | M5783 | CT | | 4514 |
| 4619 | IV | 827 | 0 | 0 | 283 | 0 | 0 | Mono | Mono | | 21650381 | PF1061 | CT | | 4619 |
| 4632 | IV | 827 | 0 | 0 | 283 | 0 | 0 | Mono | Mono | | 21674378 | PF1065 | CT | | 4632 |
| 4634 | IV | 827 | 0 | 0 | 283 | 0 | 0 | Mono | Mono | | 21683734 | M5788 | CT | | 4634 |
| 4940 | IV | 827 | 0 | 0 | 283 | 0 | 0 | Mono | Mono | | 22067044 | PF1092 | CT | | 4940 |
| 4965 | IV | 826 | 0 | 1 | 283 | 0 | 0 | Mono | Mono | | 22136242 | PF1097 | CT | | 4965 |
| 5059 | IV | 827 | 0 | 0 | 283 | 0 | 0 | Mono | Mono | | 22571947 | PF1203 | CT | | 5059 |
| 5151 | IV | 827 | 0 | 0 | 283 | 0 | 0 | Mono | Mono | | 22744945 | M294 | CT | | 5151 |
| 5210 | IV | 825 | 0 | 2 | 283 | 0 | 0 | Mono | Mono | | 22850240 | M5804 | CT | | 5210 |
| 5325 | IV | 827 | 0 | 0 | 283 | 0 | 0 | Mono | Mono | | 23081753 | M5808 | CT | | 5325 |
| 5329 | IV | 825 | 0 | 2 | 282 | 0 | 1 | Mono | Mono | | 23105586 | M5812 | CT | | 5329 |
| 5374 | IV | 825 | 0 | 2 | 283 | 0 | 0 | Mono | Mono | | 23234852 | Y1571 | CT | | 5374 |

| | | M9+ | | | M9- | | | | | | Build 37 | | ISOGG | | |
|---|---|---|---|---|---|---|---|---|---|---|---|---|---|---|---|
| List | Stage | 0 | 1 | . | 0 | 1 | . | M9? | Clade(s) | Prob. | Position | Name | Subgroup Name | Amb. | List |
| 5416 | IV | 827 | 0 | 0 | 283 | 0 | 0 | Mono | Mono | | 23298645 | M5816 | CT | | 5416 |
| 5417 | IV | 826 | 0 | 1 | 283 | 0 | 0 | Mono | Mono | | 23302389 | L977 | CT | | 5417 |
| 5476 | IV | 825 | 0 | 2 | 283 | 0 | 0 | Mono | Mono | | 23466468 | M5819 | CT | | 5476 |
| 5511 | IV | 826 | 0 | 1 | 283 | 0 | 0 | Mono | Mono | | 23555082 | M5821 | CT | | 5511 |
| 5643 | IV | 825 | 0 | 2 | 283 | 0 | 0 | Mono | Mono | | 24358251 | Y1574 | CT | | 5643 |
| 5674 | IV | 825 | 0 | 2 | 283 | 0 | 0 | Mono | Mono | | 24443898 | M5825 | CT | | 5674 |
| 5709 | IV | 827 | 0 | 0 | 281 | 0 | 2 | Mono | Mono | | 28590906 | Y1578 | CT | | 5709 |
| 4849 | IV | 827 | 0 | 0 | 283 | 0 | 0 | Mono | Mono | | 21899888 | N2 | D (Notes) | | 4849 |
| 66 | IV | 827 | 0 | 0 | 283 | 0 | 0 | Mono | Mono | | 2726402 | CTS94 | D1 | | 66 |
| 135 | IV | 827 | 0 | 0 | 283 | 0 | 0 | Mono | Mono | | 2828425 | IMS-JST021355 | D1 | | 135 |
| 463 | IV | 827 | 0 | 0 | 283 | 0 | 0 | Mono | Mono | | 6630211 | F3835 | D1 | | 463 |
| 750 | IV | 827 | 0 | 0 | 283 | 0 | 0 | Mono | Mono | | 7129691 | Z1588 | D1 | | 750 |
| 864 | IV | 827 | 0 | 0 | 283 | 0 | 0 | Mono | Mono | | 7520390 | F975 | D1 | | 864 |
| 942 | IV | 826 | 0 | 1 | 283 | 0 | 0 | Mono | Mono | | 7643998 | V1320 | D1 | | 942 |
| 1031 | IV | 827 | 0 | 0 | 283 | 0 | 0 | Mono | Mono | | 7792496 | V1437 | D1 | | 1031 |
| 1251 | IV | 827 | 0 | 0 | 283 | 0 | 0 | Mono | Mono | | 8280433 | Z1591 | D1 | | 1251 |
| 1414 | IV | 827 | 0 | 0 | 283 | 0 | 0 | Mono | Mono | | 8503447 | V2153 | D1 | | 1414 |
| 1449 | IV | 827 | 0 | 0 | 283 | 0 | 0 | Mono | Mono | | 8559992 | V2256 | D1 | | 1449 |
| 1604 | IV | 826 | 0 | 1 | 282 | 0 | 1 | Mono | Mono | | 8869318 | F1439 | D1 | | 1604 |
| 1741 | IV | 827 | 0 | 0 | 283 | 0 | 0 | Mono | Mono | | 9828418 | F1612 | D1 | | 1741 |
| 1772 | IV | 823 | 0 | 4 | 282 | 0 | 1 | Mono | Mono | | 9901153 | F1642 | D1 | | 1772 |
| 2201 | IV | 808 | 0 | 19 | 281 | 0 | 2 | Mono | Mono | | 14609947 | F3920 | D1 | | 2201 |
| 2566 | IV | 821 | 0 | 6 | 283 | 0 | 0 | Mono | Mono | | 15253322 | V2849 | D1 | | 2566 |
| 2585 | IV | 826 | 0 | 1 | 283 | 0 | 0 | Mono | Mono | | 15315059 | V2885 | D1 | | 2585 |
| 2904 | IV | 827 | 0 | 0 | 283 | 0 | 0 | Mono | Mono | | 15985397 | F2062 | D1 | | 2904 |
| 3018 | IV | 827 | 0 | 0 | 283 | 0 | 0 | Mono | Mono | | 16370123 | Z1608 | D1 | | 3018 |
| 3032 | IV | 827 | 0 | 0 | 283 | 0 | 0 | Mono | Mono | | 16403895 | F2157 | D1 | | 3032 |
| 3040 | IV | 827 | 0 | 0 | 283 | 0 | 0 | Mono | Mono | | 16415936 | Z1610 | D1 | | 3040 |
| 3258 | IV | 827 | 0 | 0 | 283 | 0 | 0 | Mono | Mono | | 16902070 | F2318 | D1 | | 3258 |
| 3268 | IV | 827 | 0 | 0 | 283 | 0 | 0 | Mono | Mono | | 16915498 | F2325 | D1 | | 3268 |
| 3429 | IV | 827 | 0 | 0 | 283 | 0 | 0 | Mono | Mono | | 17358069 | F2455 | D1 | | 3429 |
| 3536 | IV | 827 | 0 | 0 | 283 | 0 | 0 | Mono | Mono | | 17595451 | F3709 | D1 | | 3536 |
| 3725 | IV | 827 | 0 | 0 | 283 | 0 | 0 | Mono | Mono | | 17980858 | Z1612 | D1 | | 3725 |
| 3763 | IV | 827 | 0 | 0 | 283 | 0 | 0 | Mono | Mono | | 18066200 | F3976 | D1 | | 3763 |
| 3863 | IV | 827 | 0 | 0 | 283 | 0 | 0 | Mono | Mono | | 18538031 | F2765 | D1 | | 3863 |
| 4087 | IV | 827 | 0 | 0 | 283 | 0 | 0 | Mono | Mono | | 19020916 | V3694 | D1 | | 4087 |
| 4155 | IV | 826 | 0 | 1 | 283 | 0 | 0 | Mono | Mono | | 19156924 | F2956 | D1 | | 4155 |
| 4972 | IV | 827 | 0 | 0 | 283 | 0 | 0 | Mono | Mono | | 22148722 | F3289 | D1 | | 4972 |
| 5067 | IV | 820 | 0 | 7 | 279 | 0 | 4 | Mono | Mono | | 22584021 | F3321 | D1 | | 5067 |
| 5085 | IV | 827 | 0 | 0 | 283 | 0 | 0 | Mono | Mono | | 22625398 | Z1620 | D1 | | 5085 |
| 5239 | IV | 827 | 0 | 0 | 283 | 0 | 0 | Mono | Mono | | 22919926 | F3370 | D1 | | 5239 |
| 5550 | IV | 826 | 0 | 1 | 282 | 0 | 1 | Mono | Mono | | 23749005 | F3519 | D1 | | 5550 |
| 5655 | IV | 825 | 0 | 2 | 283 | 0 | 0 | Mono | Mono | | 24390651 | F3615 | D1 | | 5655 |
| 5688 | IV | 821 | 0 | 6 | 282 | 0 | 1 | Mono | Mono | | 24495962 | F3666 | D1 | | 5688 |
| 1992 | IV | 827 | 0 | 0 | 283 | 0 | 0 | Mono | Mono | | 14076879 | F1690 | D1~ | | 1992 |
| 3317 | IV | 827 | 0 | 0 | 283 | 0 | 0 | Mono | Mono | | 17033780 | F3704 | D1~ | | 3317 |
| 3208 | IV | 827 | 0 | 0 | 283 | 0 | 0 | Mono | Mono | | 16832763 | F2285 | D1a | A | 3208 |
| 5365 | IV | 827 | 0 | 0 | 283 | 0 | 0 | Mono | Mono | | 23165423 | CTS11577 | D1a | | 5365 |
| 3147 | IV | 827 | 0 | 0 | 282 | 0 | 1 | Mono | Mono | | 16693532 | F2238 | D1a1a1 | | 3147 |
| 1926 | IV | 827 | 0 | 0 | 283 | 0 | 0 | Mono | Mono | | 13864428 | Z26282.2 | D1a1a1a1a1b2b~ | | 1926 |
| 2171 | IV | 825 | 0 | 2 | 283 | 0 | 0 | Mono | Mono | | 14497168 | P47 | D1a1b1~ | | 2171 |
| 50 | IV | 827 | 0 | 0 | 283 | 0 | 0 | Mono | Mono | | 2711801 | Y14732 | D1a1b1a | | 50 |
| 1575 | IV | 827 | 0 | 0 | 283 | 0 | 0 | Mono | Mono | | 8784179 | F1412.2 | D1a1b1a | A | 1575 |
| 2007 | IV | 826 | 0 | 1 | 283 | 0 | 0 | Mono | Mono | | 14106253 | Y14779 | D1a1b1a | | 2007 |
| 2292 | IV | 827 | 0 | 0 | 283 | 0 | 0 | Mono | Mono | | 14838700 | Page31 | D1a2a | | 2292 |
| 3781 | IV | 827 | 0 | 0 | 283 | 0 | 0 | Mono | Mono | | 18097249 | P190 | D1a2a | | 3781 |
| 4800 | IV | 827 | 0 | 0 | 283 | 0 | 0 | Mono | Mono | | 21872738 | M55 | D1a2a | | 4800 |
| 4719 | IV | 825 | 0 | 2 | 282 | 0 | 1 | Mono | Mono | | 21765281 | M116.2 | D1a2a1 | A | 4719 |
| 4888 | IV | 827 | 0 | 0 | 283 | 0 | 0 | Mono | Mono | | 21930287 | M125 | D1a2a1a | | 4888 |
| 72 | IV | 827 | 0 | 0 | 283 | 0 | 0 | Mono | Mono | | 2731887 | IMS-JST055457 | D1a2a1a2 | | 72 |
| 5683 | IV | 825 | 0 | 2 | 283 | 0 | 0 | Mono | Mono | | 24464597 | IMS-JST022457 | D1a2a1a2 | | 5683 |
| 1391 | IV | 823 | 0 | 4 | 283 | 0 | 0 | Mono | Mono | | 8482714 | Z3611 | D1a2a1a2b1 | A | 1391 |
| 2022 | IV | 827 | 0 | 0 | 283 | 0 | 0 | Mono | Mono | | 14141411 | Z1498 | D1a2a1a2b1 | A | 2022 |
| 2330 | IV | 827 | 0 | 0 | 283 | 0 | 0 | Mono | Mono | | 14889059 | CTS3397 | D1a2a1a2b1 | A | 2330 |
| 4754 | IV | 827 | 0 | 0 | 283 | 0 | 0 | Mono | Mono | | 21817062 | Z3642 | D1a2a1a2b1 | A | 4754 |
| 952 | IV | 827 | 0 | 0 | 283 | 0 | 0 | Mono | Mono | | 7656808 | Z3605 | D1a2a1a2b1a | A | 952 |
| 1023 | IV | 826 | 0 | 1 | 283 | 0 | 0 | Mono | Mono | | 7765271 | Z3607 | D1a2a1a2b1a | A | 1023 |
| 1261 | IV | 827 | 0 | 0 | 283 | 0 | 0 | Mono | Mono | | 8292501 | Z1501 | D1a2a1a2b1a | A | 1261 |
| 1535 | IV | 587 | 0 | 240 | 202 | 0 | 81 | Mono | Mono | | 8681455 | Z3617 | D1a2a1a2b1a | A | 1535 |
| 3663 | IV | 827 | 0 | 0 | 283 | 0 | 0 | Mono | Mono | | 17864712 | CTS8181 | D1a2a1a2b1a | A | 3663 |
| 3967 | IV | 827 | 0 | 0 | 283 | 0 | 0 | Mono | Mono | | 18770421 | Z1502 | D1a2a1a2b1a | A | 3967 |
| 4309 | IV | 827 | 0 | 0 | 283 | 0 | 0 | Mono | Mono | | 19502802 | Z1503 | D1a2a1a2b1a | A | 4309 |
| 5082 | IV | 827 | 0 | 0 | 283 | 0 | 0 | Mono | Mono | | 22617318 | Z3651 | D1a2a1a2b1a | A | 5082 |
| 4874 | IV | 826 | 0 | 1 | 281 | 0 | 2 | Mono | Mono | | 21916474 | Z1504 | D1a2a1a2b1a1 | A | 4874 |
| 2629 | IV | 827 | 0 | 0 | 283 | 0 | 0 | Mono | Mono | | 15426000 | P120 | D1a2a1b~ | | 2629 |
| 537 | IV | 827 | 0 | 0 | 283 | 0 | 0 | Mono | Mono | | 6804892 | Z1569 | D1a2a1c1 | | 537 |
| 737 | IV | 827 | 0 | 0 | 283 | 0 | 0 | Mono | Mono | | 7079046 | Z1570 | D1a2a1c1 | | 737 |
| 4842 | IV | 827 | 0 | 0 | 283 | 0 | 0 | Mono | Mono | | 21897321 | Z14878 | D1a2a1c1 | | 4842 |
| 820 | IV | 826 | 0 | 1 | 283 | 0 | 0 | Mono | Mono | | 7283222 | Z1533 | D1a2a1c1a | | 820 |
| 5684 | IV | 820 | 0 | 7 | 283 | 0 | 0 | Mono | Mono | | 24471706 | IMS-JST022456 | D1a2a1c1a | | 5684 |
| 126 | IV | 827 | 0 | 0 | 283 | 0 | 0 | Mono | Mono | | 2818823 | Z1527 | D1a2a1c1a1 | | 126 |
| 186 | IV | 826 | 0 | 1 | 283 | 0 | 0 | Mono | Mono | | 2890647 | Z1528 | D1a2a1c1a1 | A | 186 |
| 4827 | IV | 827 | 0 | 0 | 283 | 0 | 0 | Mono | Mono | | 21892634 | M151 | D1a2a1c1a1c~ | | 4827 |
| 5221 | IV | 827 | 0 | 0 | 283 | 0 | 0 | Mono | Mono | | 22882865 | M6525 | D1a2a1c1b | | 5221 |
| 88 | IV | 826 | 0 | 1 | 283 | 0 | 0 | Mono | Mono | | 2750950 | CTS131 | D1a2a2 | | 88 |
| 2742 | III | 826 | 0 | 1 | 281 | 2 | 0 | M9- | 34 | | 15591445 | M228.2 | D1a2a2a2b | A,W | 2742 |
| 5560 | IV | 827 | 0 | 0 | 283 | 0 | 0 | Mono | Mono | | 23769206 | V8035^^ | D1a2b1 | A | 5560 |
| 133 | IV | 827 | 0 | 0 | 283 | 0 | 0 | Mono | Mono | | 2828140 | L1378 | D1b | | 133 |
| 2744 | IV | 827 | 0 | 0 | 283 | 0 | 0 | Mono | Mono | | 15591447 | M226.2 | D1b | A | 2744 |
| 71 | I | 827 | 0 | 0 | 246 | 37 | 0 | M9- | 1 – 10 | | 2728927 | PF1447 | DE | | 71 |
| 831 | III | 827 | 0 | 0 | 245 | 37 | 1 | M9- | 1 – 10 | | 7320623 | PF1423 | DE | | 831 |
| 876 | I | 827 | 0 | 0 | 246 | 37 | 0 | M9- | 1 – 10 | | 7540450 | M5402 | DE | | 876 |
| 1156 | I | 827 | 0 | 0 | 246 | 37 | 0 | M9- | 1 – 10 | | 8072664 | PF1425 | DE | | 1156 |
| 1758 | I | 827 | 0 | 0 | 246 | 37 | 0 | M9- | 1 – 10 | | 9851457 | PF1430 | DE | | 1758 |
| 2296 | I | 827 | 0 | 0 | 246 | 37 | 0 | M9- | 1 – 10 | | 14842223 | PF1432 | DE | | 2296 |
| 2709 | III | 826 | 0 | 1 | 246 | 37 | 0 | M9- | 1 – 10 | | 15543875 | PF1433 | DE | | 2709 |

| | | | M9+ | | | M9- | | | | | Build 37 | | ISOGG | | |
|---|---|---|---|---|---|---|---|---|---|---|---|---|---|---|---|
| List | Stage | 0 | 1 | . | 0 | 1 | . | M9? | Clade(s) | Prob. | Position | Name | Subgroup Name | Amb. | List |
| 2740 | I | 827 | 0 | 0 | 246 | 37 | 0 | M9- | 1 – 10 | | 15590674 | PF1434 | DE | | 2740 |
| 2748 | I | 827 | 0 | 0 | 246 | 37 | 0 | M9- | 1 – 10 | | 15591537 | PF1435 | DE | | 2748 |
| 3873 | I | 827 | 0 | 0 | 246 | 37 | 0 | M9- | 1 – 10 | | 18561042 | P153 | DE | | 3873 |
| 4036 | III | 824 | 0 | 3 | 246 | 37 | 0 | M9- | 1 – 10 | | 18907927 | PF1770 | DE | | 4036 |
| 4248 | I | 827 | 0 | 0 | 246 | 37 | 0 | M9- | 1 – 10 | | 19338287 | M5518 | DE | | 4248 |
| 4257 | I | 827 | 0 | 0 | 246 | 37 | 0 | M9- | 1 – 10 | | 19370916 | PF1439 | DE | | 4257 |
| 4421 | I | 827 | 0 | 0 | 246 | 37 | 0 | M9- | 1 – 10 | | 21227423 | PF1442 | DE | | 4421 |
| 4457 | I | 827 | 0 | 0 | 246 | 37 | 0 | M9- | 1 – 10 | | 21306315 | PF1797.2 | DE | A | 4457 |
| 4646 | I | 827 | 0 | 0 | 246 | 37 | 0 | M9- | 1 – 10 | | 21717208 | PF1444 | DE | | 4646 |
| 5122 | III | 826 | 0 | 1 | 246 | 36 | 1 | M9- | 1 – 10 | MD | 22721522 | PF1876 | DE | | 5122 |
| 5656 | III | 823 | 0 | 4 | 243 | 36 | 4 | M9- | 1 – 10 | MD | 24392653 | M5568 | DE | | 5656 |
| 519 | I | 827 | 0 | 0 | 246 | 37 | 0 | M9- | 1 – 10 | | 6750775 | PF1504 | E | | 519 |
| 573 | I | 827 | 0 | 0 | 246 | 37 | 0 | M9- | 1 – 10 | | 6861075 | PF1507 | E | | 573 |
| 688 | I | 827 | 0 | 0 | 246 | 37 | 0 | M9- | 1 – 10 | | 6965215 | PF1510 | E | | 688 |
| 702 | I | 827 | 0 | 0 | 246 | 37 | 0 | M9- | 1 – 10 | | 6995523 | P173 | E | | 702 |
| 742 | I | 827 | 0 | 0 | 246 | 37 | 0 | M9- | 1 – 10 | | 7098985 | M5389 | E | | 742 |
| 921 | I | 827 | 0 | 0 | 246 | 37 | 0 | M9- | 1 – 10 | | 7602644 | V1254 | E | | 921 |
| 1094 | I | 827 | 0 | 0 | 246 | 37 | 0 | M9- | 1 – 10 | | 7913358 | PF1539 | E | | 1094 |
| 1197 | I | 827 | 0 | 0 | 246 | 37 | 0 | M9- | 1 – 10 | | 8188581 | V1692 | E | | 1197 |
| 1376 | I | 827 | 0 | 0 | 246 | 37 | 0 | M9- | 1 – 10 | | 8469322 | PF1552 | E | | 1376 |
| 1398 | I | 827 | 0 | 0 | 246 | 37 | 0 | M9- | 1 – 10 | | 8485542 | PF1553.2 | E | A | 1398 |
| 1430 | I | 827 | 0 | 0 | 246 | 37 | 0 | M9- | 1 – 10 | | 8532844 | V2187 | E | | 1430 |
| 1492 | I | 827 | 0 | 0 | 246 | 37 | 0 | M9- | 1 – 10 | | 8612630 | V2350 | E | | 1492 |
| 1550 | I | 827 | 0 | 0 | 246 | 37 | 0 | M9- | 1 – 10 | | 8703052 | V2471 | E | | 1550 |
| 1587 | I | 827 | 0 | 0 | 246 | 37 | 0 | M9- | 1 – 10 | | 8810775 | PF1564 | E | | 1587 |
| 1623 | I | 827 | 0 | 0 | 246 | 37 | 0 | M9- | 1 – 10 | | 8908270 | PF1567 | E | | 1623 |
| 1683 | III | 824 | 0 | 3 | 246 | 36 | 1 | M9- | 1 – 10 | MD | 9394763 | PF1576 | E | | 1683 |
| 1690 | III | 825 | 0 | 2 | 244 | 37 | 2 | M9- | 1 – 10 | 1H | 9443407 | PF1583 | E | | 1690 |
| 1864 | III | 827 | 0 | 0 | 244 | 36 | 3 | M9- | 1 – 10 | MD | 13666276 | Z15671 | E | | 1864 |
| 1876 | III | 826 | 0 | 1 | 244 | 37 | 2 | M9- | 1 – 10 | | 13679813 | Z15672 | E | | 1876 |
| 1905 | III | 825 | 0 | 2 | 246 | 37 | 0 | M9- | 1 – 10 | | 13802034 | PF1618 | E | | 1905 |
| 1916 | I | 827 | 0 | 0 | 246 | 37 | 0 | M9- | 1 – 10 | | 13828327 | Z15673 | E | | 1916 |
| 2101 | I | 827 | 0 | 0 | 246 | 37 | 0 | M9- | 1 – 10 | | 14329233 | PF1636 | E | | 2101 |
| 2172 | I | 827 | 0 | 0 | 246 | 37 | 0 | M9- | 1 – 10 | | 14497207 | PF1643 | E | | 2172 |
| 2218 | I | 827 | 0 | 0 | 246 | 37 | 0 | M9- | 1 – 10 | | 14664631 | PF1646 | E | | 2218 |
| 2267 | I | 827 | 0 | 0 | 246 | 37 | 0 | M9- | 1 – 10 | | 14804077 | P175 | E | | 2267 |
| 2452 | III | 825 | 0 | 2 | 246 | 37 | 0 | M9- | 1 – 10 | | 15021522 | PF1656 | E | | 2452 |
| 2768 | I | 827 | 0 | 0 | 246 | 37 | 0 | M9- | 1 – 10 | | 15617734 | PF1669 | E | | 2768 |
| 2827 | I | 827 | 0 | 0 | 246 | 37 | 0 | M9- | 1 – 10 | | 15768559 | PF1671 | E | | 2827 |
| 2845 | I | 827 | 0 | 0 | 246 | 37 | 0 | M9- | 1 – 10 | | 15809326 | PF1674 | E | | 2845 |
| 2889 | I | 827 | 0 | 0 | 246 | 37 | 0 | M9- | 1 – 10 | | 15935524 | PF1678 | E | | 2889 |
| 2892 | I | 827 | 0 | 0 | 246 | 37 | 0 | M9- | 1 – 10 | | 15945309 | M5468 | E | | 2892 |
| 2947 | I | 827 | 0 | 0 | 246 | 37 | 0 | M9- | 1 – 10 | | 16203354 | PF1683 | E | | 2947 |
| 3003 | III | 827 | 0 | 0 | 245 | 37 | 1 | M9- | 1 – 10 | | 16338537 | PF1687 | E | | 3003 |
| 3129 | I | 827 | 0 | 0 | 246 | 37 | 0 | M9- | 1 – 10 | | 16636022 | PF1697 | E | | 3129 |
| 3221 | I | 827 | 0 | 0 | 246 | 37 | 0 | M9- | 1 – 10 | | 16846439 | PF1704 | E | | 3221 |
| 3297 | I | 827 | 0 | 0 | 246 | 37 | 0 | M9- | 1 – 10 | | 16960786 | PF1710 | E | | 3297 |
| 3332 | I | 827 | 0 | 0 | 246 | 37 | 0 | M9- | 1 – 10 | | 17092499 | PF1713 | E | | 3332 |
| 3454 | I | 827 | 0 | 0 | 246 | 37 | 0 | M9- | 1 – 10 | | 17420017 | PF1725 | E | | 3454 |
| 3576 | III | 825 | 0 | 2 | 246 | 37 | 0 | M9- | 1 – 10 | | 17686886 | PF1732 | E | | 3576 |
| 3635 | III | 826 | 0 | 1 | 246 | 37 | 0 | M9- | 1 – 10 | | 17814087 | PF1738 | E | | 3635 |
| 3666 | III | 823 | 0 | 4 | 246 | 37 | 0 | M9- | 1 – 10 | | 17867791 | M5500 | E | | 3666 |
| 3684 | I | 827 | 0 | 0 | 246 | 37 | 0 | M9- | 1 – 10 | | 17907131 | PF1743 | E | | 3684 |
| 3819 | I | 827 | 0 | 0 | 246 | 37 | 0 | M9- | 1 – 10 | | 18181803 | PF1752 | E | | 3819 |
| 4030 | III | 802 | 0 | 25 | 239 | 36 | 8 | M9- | 1 – 10 | MD | 18894151 | PF1769 | E | | 4030 |
| 4306 | I | 827 | 0 | 0 | 246 | 37 | 0 | M9- | 1 – 10 | | 19500107 | PF1788 | E | | 4306 |
| 4409 | I | 827 | 0 | 0 | 246 | 37 | 0 | M9- | 1 – 10 | | 21192906 | PF1795 | E | | 4409 |
| 4434 | I | 827 | 0 | 0 | 246 | 37 | 0 | M9- | 1 – 10 | | 21256219 | PF1796 | E | | 4434 |
| 4462 | I | 827 | 0 | 0 | 246 | 37 | 0 | M9- | 1 – 10 | | 21314704 | PF1798 | E | | 4462 |
| 4492 | I | 827 | 0 | 0 | 246 | 37 | 0 | M9- | 1 – 10 | | 21385724 | PF1803 | E | | 4492 |
| 4564 | I | 827 | 0 | 0 | 246 | 37 | 0 | M9- | 1 – 10 | | 21565962 | M5539 | E | | 4564 |
| 4571 | III | 760 | 0 | 67 | 230 | 37 | 16 | M9- | 1 – 10 | | 21576051 | PF1816 | E | | 4571 |
| 4590 | I | 827 | 0 | 0 | 246 | 37 | 0 | M9- | 1 – 10 | | 21618583 | P168 | E | | 4590 |
| 4620 | I | 827 | 0 | 0 | 246 | 37 | 0 | M9- | 1 – 10 | | 21652284 | PF1819 | E | | 4620 |
| 4688 | I | 827 | 0 | 0 | 246 | 37 | 0 | M9- | 1 – 10 | | 21747107 | PF1822 | E | | 4688 |
| 4734 | I | 827 | 0 | 0 | 246 | 37 | 0 | M9- | 1 – 10 | | 21778998 | PF1823 | E | | 4734 |
| 4977 | I | 827 | 0 | 0 | 246 | 37 | 0 | M9- | 1 – 10 | | 22158261 | PF1834 | E | | 4977 |
| 5042 | I | 827 | 0 | 0 | 246 | 37 | 0 | M9- | 1 – 10 | | 22538799 | PF1868 | E | | 5042 |
| 5091 | III | 827 | 0 | 0 | 245 | 37 | 1 | M9- | 1 – 10 | | 22646728 | PF1873.2 | E | A | 5091 |
| 5108 | I | 827 | 0 | 0 | 246 | 37 | 0 | M9- | 1 – 10 | | 22688731 | PF1874 | E | | 5108 |
| 5202 | III | 826 | 0 | 1 | 246 | 37 | 0 | M9- | 1 – 10 | | 22823374 | M5552 | E | | 5202 |
| 5236 | III | 823 | 0 | 4 | 246 | 37 | 0 | M9- | 1 – 10 | | 22918577 | PF1878 | E | | 5236 |
| 5306 | III | 825 | 0 | 2 | 246 | 37 | 0 | M9- | 1 – 10 | | 23059591 | PF1888 | E | | 5306 |
| 5369 | III | 822 | 0 | 5 | 246 | 37 | 0 | M9- | 1 – 10 | | 23185647 | PF1893 | E | | 5369 |
| 5388 | III | 801 | 0 | 26 | 237 | 37 | 9 | M9- | 1 – 10 | | 23249378 | YSC0000032 | E | | 5388 |
| 5432 | I | 827 | 0 | 0 | 246 | 37 | 0 | M9- | 1 – 10 | | 23353901 | PF1901 | E | | 5432 |
| 5468 | I | 827 | 0 | 0 | 246 | 37 | 0 | M9- | 1 – 10 | | 23443971 | PF1903 | E | | 5468 |
| 5475 | I | 827 | 0 | 0 | 246 | 37 | 0 | M9- | 1 – 10 | | 23463283 | PF1904 | E | | 5475 |
| 5503 | III | 826 | 0 | 1 | 246 | 37 | 0 | M9- | 1 – 10 | | 23538555 | PF1906 | E | | 5503 |
| 5641 | III | 827 | 0 | 0 | 245 | 37 | 1 | M9- | 1 – 10 | | 24356128 | PF1917 | E | | 5641 |
| 5660 | III | 826 | 0 | 1 | 246 | 37 | 0 | M9- | 1 – 10 | | 24399592 | PF1919 | E | | 5660 |
| 5672 | III | 825 | 0 | 2 | 248 | 35 | 0 | M9- | | PI | 24437979 | S7474 | E | | 5672 |
| 605 | IV | 827 | 0 | 0 | 283 | 0 | 0 | Mono | Mono | | 6896051 | V27 | E (Notes) | | 605 |
| 2438 | IV | 827 | 0 | 0 | 283 | 0 | 0 | Mono | Mono | | 15016536 | M195 | E (Notes) | | 2438 |
| 4647 | IV | 827 | 0 | 0 | 283 | 0 | 0 | Mono | Mono | | 21717227 | M156 | E (Notes) | | 4647 |
| 4674 | IV | 827 | 0 | 0 | 283 | 0 | 0 | Mono | Mono | | 21736331 | M155 | E (Notes) | | 4674 |
| 4814 | IV | 827 | 0 | 0 | 282 | 0 | 1 | Mono | Mono | | 21881573 | M66 | E (Notes) | | 4814 |
| 4839 | IV | 827 | 0 | 0 | 283 | 0 | 0 | Mono | Mono | | 21896200 | V19 | E (Notes) | | 4839 |
| 4841 | IV | 825 | 0 | 2 | 282 | 0 | 1 | Mono | Mono | | 21896248 | M149 | E (Notes) | | 4841 |
| 4889 | IV | 827 | 0 | 0 | 283 | 0 | 0 | Mono | Mono | | 21931638 | M107 | E (Notes) | | 4889 |
| 5633 | IV | 827 | 0 | 0 | 281 | 0 | 2 | Mono | Mono | | 23989677 | P268 | E (Notes) | | 5633 |
| 1292 | I | 827 | 0 | 0 | 246 | 37 | 0 | M9- | 1 – 10 | | 8375407 | V1844 | E1 | | 1292 |
| 1519 | I | 827 | 0 | 0 | 246 | 37 | 0 | M9- | 1 – 10 | | 8644218 | PF1559 | E1 | | 1519 |
| 3735 | III | 826 | 0 | 1 | 246 | 37 | 0 | M9- | 1 – 10 | | 18023975 | PF1745 | E1 | | 3735 |
| 3926 | I | 827 | 0 | 0 | 246 | 37 | 0 | M9- | 1 – 10 | | 18662674 | PF1764 | E1 | | 3926 |
| 4107 | I | 827 | 0 | 0 | 246 | 37 | 0 | M9- | 1 – 10 | | 19058376 | PF1776 | E1 | | 4107 |

| | | M9+ | | | M9- | | | | | | Build 37 | | ISOGG | | |
|---|---|---|---|---|---|---|---|---|---|---|---|---|---|---|---|
| List | Stage | 0 | 1 | . | 0 | 1 | . | M9? | Clade(s) | Prob. | Position | Name | Subgroup Name | Amb. | List |
| 4368 | I | 827 | 0 | 0 | 246 | 37 | 0 | M9- | 1 – 10 | | 21083420 | PF1938 | E1 | | 4368 |
| 4521 | I | 827 | 0 | 0 | 246 | 37 | 0 | M9- | 1 – 10 | | 21453551 | PF1809 | E1 | | 4521 |
| 143 | IV | 827 | 0 | 0 | 282 | 0 | 1 | Mono | Mono | MD | 2839051 | V1118 | E1a | | 143 |
| 488 | II | 827 | 0 | 0 | 282 | 1 | 0 | M9- | 1 | | 6697474 | CTS340 | E1a | | 488 |
| 611 | II | 827 | 0 | 0 | 282 | 1 | 0 | M9- | 1 | | 6905504 | L481 | E1a | | 611 |
| 729 | II | 827 | 0 | 0 | 282 | 1 | 0 | M9- | 1 | | 7074048 | CTS895 | E1a | | 729 |
| 930 | III | 826 | 0 | 1 | 282 | 1 | 0 | M9- | 1 | | 7618399 | Z15456 | E1a | | 930 |
| 954 | II | 827 | 0 | 0 | 282 | 1 | 0 | M9- | 1 | | 7658259 | Z904 | E1a | | 954 |
| 978 | II | 827 | 0 | 0 | 282 | 1 | 0 | M9- | 1 | | 7693248 | Z15459 | E1a | | 978 |
| 1128 | II | 827 | 0 | 0 | 282 | 1 | 0 | M9- | 1 | | 8009314 | Z15461 | E1a | | 1128 |
| 1190 | III | 826 | 0 | 1 | 282 | 1 | 0 | M9- | 1 | | 8148970 | Z15462 | E1a | | 1190 |
| 1201 | III | 823 | 0 | 4 | 281 | 1 | 1 | M9- | 1 | | 8193988 | Z15464 | E1a | | 1201 |
| 1217 | II | 827 | 0 | 0 | 282 | 1 | 0 | M9- | 1 | | 8229982 | Z17396 | E1a | | 1217 |
| 1230 | II | 827 | 0 | 0 | 282 | 1 | 0 | M9- | 1 | | 8243264 | Z907 | E1a | | 1230 |
| 1351 | II | 827 | 0 | 0 | 282 | 1 | 0 | M9- | 1 | | 8450180 | Z910 | E1a | | 1351 |
| 1374 | III | 819 | 0 | 8 | 281 | 1 | 1 | M9- | 1 | | 8467563 | Z15467 | E1a | | 1374 |
| 1383 | II | 827 | 0 | 0 | 282 | 1 | 0 | M9- | 1 | | 8478528 | Z911 | E1a | A | 1383 |
| 1395 | II | 827 | 0 | 0 | 282 | 1 | 0 | M9- | 1 | | 8484355 | Z15468 | E1a | | 1395 |
| 1402 | II | 827 | 0 | 0 | 282 | 1 | 0 | M9- | 1 | | 8489812 | Z912 | E1a | | 1402 |
| 1475 | II | 827 | 0 | 0 | 282 | 1 | 0 | M9- | 1 | | 8590777 | Z916 | E1a | | 1475 |
| 1929 | III | 826 | 0 | 1 | 281 | 1 | 1 | M9- | 1 | | 13869975 | Z15496 | E1a | | 1929 |
| 2205 | II | 827 | 0 | 0 | 282 | 1 | 0 | M9- | 1 | | 14630009 | Z930 | E1a | | 2205 |
| 2283 | II | 827 | 0 | 0 | 282 | 1 | 0 | M9- | 1 | | 14822216 | V2623 | E1a | | 2283 |
| 2419 | II | 827 | 0 | 0 | 282 | 1 | 0 | M9- | 1 | | 14981420 | V2736 | E1a | | 2419 |
| 2492 | III | 825 | 0 | 2 | 282 | 1 | 0 | M9- | 1 | | 15041425 | V2785 | E1a | | 2492 |
| 2783 | III | 826 | 0 | 1 | 277 | 1 | 5 | M9- | 1 | | 15660662 | V3085 | E1a | | 2783 |
| 2932 | II | 827 | 0 | 0 | 282 | 1 | 0 | M9- | 1 | | 16183346 | CTS5283 | E1a | | 2932 |
| 2994 | II | 827 | 0 | 0 | 282 | 1 | 0 | M9- | 1 | | 16324781 | Z937 | E1a | | 2994 |
| 3298 | II | 827 | 0 | 0 | 282 | 1 | 0 | M9- | 1 | | 16962160 | CTS6516 | E1a | | 3298 |
| 3665 | II | 827 | 0 | 0 | 282 | 1 | 0 | M9- | 1 | | 17866126 | CTS8185 | E1a | | 3665 |
| 3923 | III | 827 | 0 | 0 | 281 | 1 | 1 | M9- | 1 | | 18655034 | V3362 | E1a | | 3923 |
| 3936 | II | 827 | 0 | 0 | 282 | 1 | 0 | M9- | 1 | | 18688987 | V3403 | E1a | | 3936 |
| 3945 | II | 827 | 0 | 0 | 282 | 1 | 0 | M9- | 1 | | 18718543 | CTS9155 | E1a | | 3945 |
| 3952 | II | 827 | 0 | 0 | 282 | 1 | 0 | M9- | 1 | | 18742587 | V3464 | E1a | | 3952 |
| 4152 | II | 827 | 0 | 0 | 282 | 1 | 0 | M9- | 1 | | 19150795 | CTS9927 | E1a | | 4152 |
| 4239 | II | 827 | 0 | 0 | 282 | 1 | 0 | M9- | 1 | | 19316047 | V4005 | E1a | | 4239 |
| 4299 | II | 827 | 0 | 0 | 282 | 1 | 0 | M9- | 1 | | 19486772 | V4170 | E1a | | 4299 |
| 4321 | III | 826 | 0 | 1 | 282 | 1 | 0 | M9- | 1 | | 19543393 | V4219 | E1a | | 4321 |
| 4424 | II | 827 | 0 | 0 | 282 | 1 | 0 | M9- | 1 | | 21237511 | Z944 | E1a | | 4424 |
| 4455 | II | 827 | 0 | 0 | 282 | 1 | 0 | M9- | 1 | | 21298893 | Z17399 | E1a | | 4455 |
| 4549 | III | 826 | 0 | 1 | 282 | 1 | 0 | M9- | 1 | | 21516787 | Z15512 | E1a | | 4549 |
| 4594 | II | 827 | 0 | 0 | 282 | 1 | 0 | M9- | 1 | | 21621477 | Z17400 | E1a | | 4594 |
| 4627 | III | 826 | 0 | 1 | 282 | 1 | 0 | M9- | 1 | | 21667411 | Z948 | E1a | | 4627 |
| 4682 | II | 827 | 0 | 0 | 282 | 1 | 0 | M9- | 1 | | 21740450 | M33 | E1a | | 4682 |
| 5068 | II | 827 | 0 | 0 | 282 | 1 | 0 | M9- | 1 | | 22584859 | Z15542 | E1a | | 5068 |
| 5175 | III | 826 | 0 | 1 | 282 | 1 | 0 | M9- | 1 | | 22758062 | CTS10752 | E1a | | 5175 |
| 5213 | II | 827 | 0 | 0 | 282 | 1 | 0 | M9- | 1 | | 22858980 | CTS10964 | E1a | | 5213 |
| 5297 | II | 827 | 0 | 0 | 282 | 1 | 0 | M9- | 1 | | 23033239 | CTS11248 | E1a | | 5297 |
| 5302 | II | 827 | 0 | 0 | 282 | 1 | 0 | M9- | 1 | | 23046558 | Z960 | E1a | | 5302 |
| 5318 | II | 827 | 0 | 0 | 282 | 1 | 0 | M9- | 1 | | 23073381 | CTS11329 | E1a | | 5318 |
| 587 | IV | 827 | 0 | 0 | 283 | 0 | 0 | Mono | Mono | | 6877229 | L1239 | E1a1 | | 587 |
| 1221 | IV | 827 | 0 | 0 | 283 | 0 | 0 | Mono | Mono | | 8233083 | Z17417 | E1a1 | | 1221 |
| 1252 | IV | 827 | 0 | 0 | 283 | 0 | 0 | Mono | Mono | | 8282207 | Z17418 | E1a1 | | 1252 |
| 1340 | IV | 827 | 0 | 0 | 283 | 0 | 0 | Mono | Mono | | 8436218 | Z17419 | E1a1 | | 1340 |
| 1378 | IV | 827 | 0 | 0 | 283 | 0 | 0 | Mono | Mono | | 8472581 | Z17420 | E1a1 | | 1378 |
| 1410 | IV | 827 | 0 | 0 | 283 | 0 | 0 | Mono | Mono | | 8498833 | Z17421 | E1a1 | | 1410 |
| 1478 | IV | 826 | 0 | 1 | 283 | 0 | 0 | Mono | Mono | | 8594203 | Z17422 | E1a1 | | 1478 |
| 1515 | IV | 827 | 0 | 0 | 283 | 0 | 0 | Mono | Mono | | 8641183 | Z17423 | E1a1 | | 1515 |
| 2441 | IV | 827 | 0 | 0 | 283 | 0 | 0 | Mono | Mono | | 15018495 | L632 | E1a1 | | 2441 |
| 2489 | IV | 827 | 0 | 0 | 283 | 0 | 0 | Mono | Mono | | 15040134 | Z17439 | E1a1 | | 2489 |
| 2688 | IV | 827 | 0 | 0 | 283 | 0 | 0 | Mono | Mono | | 15512685 | Z17443 | E1a1 | | 2688 |
| 2691 | IV | 826 | 0 | 1 | 283 | 0 | 0 | Mono | Mono | | 15514675 | Z17444 | E1a1 | | 2691 |
| 2732 | IV | 821 | 0 | 6 | 281 | 0 | 2 | Mono | Mono | | 15582383 | Z17445 | E1a1 | | 2732 |
| 2765 | IV | 827 | 0 | 0 | 282 | 0 | 1 | Mono | Mono | | 15615452 | L1240 | E1a1 | | 2765 |
| 2841 | IV | 827 | 0 | 0 | 283 | 0 | 0 | Mono | Mono | | 15800133 | L634 | E1a1 | | 2841 |
| 2861 | IV | 827 | 0 | 0 | 283 | 0 | 0 | Mono | Mono | | 15840381 | Z17446 | E1a1 | | 2861 |
| 3890 | IV | 826 | 0 | 1 | 283 | 0 | 0 | Mono | Mono | | 18587512 | Z17466 | E1a1 | | 3890 |
| 4045 | IV | 827 | 0 | 0 | 283 | 0 | 0 | Mono | Mono | | 18919035 | Z17468 | E1a1 | | 4045 |
| 4047 | IV | 827 | 0 | 0 | 283 | 0 | 0 | Mono | Mono | | 18925978 | Z17469 | E1a1 | | 4047 |
| 4225 | IV | 827 | 0 | 0 | 283 | 0 | 0 | Mono | Mono | | 19280369 | Z17472 | E1a1 | | 4225 |
| 4696 | IV | 826 | 0 | 1 | 283 | 0 | 0 | Mono | Mono | | 21752644 | M44 | E1a1 | | 4696 |
| 141 | II | 827 | 0 | 0 | 282 | 1 | 0 | M9- | 1 | | 2831375 | V1113 | E1a2 | | 141 |
| 1235 | II | 827 | 0 | 0 | 282 | 1 | 0 | M9- | 1 | | 8249966 | Z908 | E1a2 | A | 1235 |
| 1287 | III | 812 | 0 | 15 | 279 | 1 | 3 | M9- | 1 | | 8363564 | Z15131 | E1a2 | | 1287 |
| 3932 | II | 827 | 0 | 0 | 282 | 1 | 0 | M9- | 1 | | 18671174 | V3384 | E1a2 | | 3932 |
| 5088 | III | 826 | 0 | 1 | 281 | 1 | 1 | M9- | 1 | | 22634180 | Z956 | E1a2 | | 5088 |
| 5120 | II | 827 | 0 | 0 | 282 | 1 | 0 | M9- | 1 | | 22720028 | Z958 | E1a2 | | 5120 |
| 144 | II | 827 | 0 | 0 | 282 | 1 | 0 | M9- | 1 | | 2840301 | V1119 | E1a2a | | 144 |
| 2795 | III | 826 | 0 | 1 | 282 | 1 | 0 | M9- | 1 | | 15691160 | CTS4519 | E1a2a | | 2795 |
| 3930 | III | 827 | 0 | 0 | 281 | 1 | 1 | M9- | 1 | | 18667241 | V3379 | E1a2a | | 3930 |
| 5041 | II | 827 | 0 | 0 | 282 | 1 | 0 | M9- | 1 | | 22536067 | Z15271 | E1a2a1 | | 5041 |
| 5207 | II | 827 | 0 | 0 | 282 | 1 | 0 | M9- | 1 | | 22843448 | CTS10935 | E1a2a1 | | 5207 |
| 1218 | IV | 827 | 0 | 0 | 283 | 0 | 0 | Mono | Mono | | 8229995 | Z885 | E1a2a1a | | 1218 |
| 2134 | IV | 827 | 0 | 0 | 283 | 0 | 0 | Mono | Mono | | 14405427 | CTS2648 | E1a2a1a | | 2134 |
| 2321 | IV | 827 | 0 | 0 | 283 | 0 | 0 | Mono | Mono | | 14881244 | CTS3380 | E1a2a1a | | 2321 |
| 2652 | IV | 827 | 0 | 0 | 283 | 0 | 0 | Mono | Mono | | 15460054 | CTS4187 | E1a2a1a | A | 2652 |
| 1972 | IV | 827 | 0 | 0 | 283 | 0 | 0 | Mono | Mono | | 14049582 | CTS1753 | E1a2a1a1 | | 1972 |
| 3821 | IV | 827 | 0 | 0 | 283 | 0 | 0 | Mono | Mono | | 18184213 | Z881 | E1a2a1b | | 3821 |
| 594 | IV | 827 | 0 | 0 | 283 | 0 | 0 | Mono | Mono | | 6886942 | CTS602 | E1a2a1b1 | | 594 |
| 2308 | IV | 827 | 0 | 0 | 283 | 0 | 0 | Mono | Mono | | 14859550 | CTS3351 | E1a2a1b1 | | 2308 |
| 2505 | IV | 827 | 0 | 0 | 283 | 0 | 0 | Mono | Mono | | 15071710 | CTS3615 | E1a2a1b1 | | 2505 |
| 2374 | IV | 827 | 0 | 0 | 283 | 0 | 0 | Mono | Mono | | 14928196 | Z15350 | E1a2a1b1a | | 2374 |
| 845 | IV | 827 | 0 | 0 | 283 | 0 | 0 | Mono | Mono | | 7383272 | Z15188^ | E1a2a2 | A | 845 |
| 1666 | IV | 827 | 0 | 0 | 283 | 0 | 0 | Mono | Mono | | 9145267 | Z15088 | E1a2b | | 1666 |
| 108 | IV | 812 | 0 | 15 | 278 | 0 | 5 | Mono | Mono | | 2800805 | Z15068 | E1a2b1 | | 108 |
| 2628 | IV | 827 | 0 | 0 | 283 | 0 | 0 | Mono | Mono | | 15425719 | L94 | E1a2b1 | | 2628 |

| | | M9+ | | | M9- | | | | | | Build 37 | | ISOGG | | |
|---|---|---|---|---|---|---|---|---|---|---|---|---|---|---|---|
| List | Stage | 0 | 1 | . | 0 | 1 | . | M9? | Clade(s) | Prob. | Position | Name | Subgroup Name | Amb. | List |
| 2734 | IV | 826 | 0 | 1 | 283 | 0 | 0 | Mono | Mono | | 15586322 | Z5988 | E1a2b1a1 | | 2734 |
| 2029 | I | 827 | 0 | 0 | 247 | 36 | 0 | M9- | 2 – 10 | | 14159846 | PF1939 | E1b | | 2029 |
| 1978 | I | 827 | 0 | 0 | 247 | 36 | 0 | M9- | 2 – 10 | | 14060308 | P179 | E1b1 | | 1978 |
| 3443 | I | 827 | 0 | 0 | 247 | 36 | 0 | M9- | 2 – 10 | | 17394111 | P181 | E1b1 | | 3443 |
| 3898 | I | 827 | 0 | 0 | 247 | 36 | 0 | M9- | 2 – 10 | | 18601274 | PF1941 | E1b1 | | 3898 |
| 4582 | I | 827 | 0 | 0 | 247 | 36 | 0 | M9- | 2 – 10 | | 21610831 | PN2 | E1b1 | | 4582 |
| 81 | I | 827 | 0 | 0 | 247 | 36 | 0 | M9- | 2 – 10 | | 2739468 | V1048 | E1b1~ | | 81 |
| 486 | III | 827 | 0 | 0 | 247 | 35 | 1 | M9- | 2 – 10 | MD | 6692469 | PF1503 | E1b1~ | | 486 |
| 851 | I | 827 | 0 | 0 | 247 | 36 | 0 | M9- | 2 – 10 | | 7401836 | PF1526 | E1b1~ | | 851 |
| 1003 | I | 827 | 0 | 0 | 247 | 36 | 0 | M9- | 2 – 10 | | 7736744 | PF1535 | E1b1~ | | 1003 |
| 1020 | I | 827 | 0 | 0 | 247 | 36 | 0 | M9- | 2 – 10 | | 7762688 | PF1536 | E1b1~ | | 1020 |
| 1192 | III | 825 | 0 | 2 | 247 | 36 | 0 | M9- | 2 – 10 | | 8155660 | PF1544 | E1b1~ | A | 1192 |
| 1236 | III | 825 | 0 | 2 | 247 | 36 | 0 | M9- | 2 – 10 | | 8254171 | PF1547 | E1b1~ | | 1236 |
| 1266 | I | 827 | 0 | 0 | 247 | 36 | 0 | M9- | 2 – 10 | | 8308572 | V1798 | E1b1~ | | 1266 |
| 1334 | III | 826 | 0 | 1 | 247 | 36 | 0 | M9- | 2 – 10 | | 8430640 | PF1551 | E1b1~ | | 1334 |
| 1506 | I | 827 | 0 | 0 | 247 | 36 | 0 | M9- | 2 – 10 | | 8633213 | PF1558 | E1b1~ | A | 1506 |
| 1553 | III | 826 | 0 | 1 | 247 | 36 | 0 | M9- | 2 – 10 | | 8711273 | V2484^ | E1b1~ | | 1553 |
| 1725 | III | 824 | 0 | 3 | 247 | 35 | 1 | M9- | 2 – 10 | MD | 9785893 | PF1589 | E1b1~ | | 1725 |
| 2018 | I | 827 | 0 | 0 | 247 | 36 | 0 | M9- | 2 – 10 | | 14127014 | PF1625 | E1b1~ | | 2018 |
| 2020 | I | 827 | 0 | 0 | 247 | 36 | 0 | M9- | 2 – 10 | | 14140277 | PF1626 | E1b1~ | | 2020 |
| 2046 | I | 827 | 0 | 0 | 247 | 36 | 0 | M9- | 2 – 10 | | 14199508 | PF1628 | E1b1~ | | 2046 |
| 2139 | III | 823 | 0 | 4 | 245 | 36 | 2 | M9- | 2 – 10 | | 14419185 | PF1639 | E1b1~ | | 2139 |
| 2243 | III | 827 | 0 | 0 | 246 | 36 | 1 | M9- | 2 – 10 | | 14768679 | PF1650 | E1b1~ | | 2243 |
| 2253 | I | 827 | 0 | 0 | 247 | 36 | 0 | M9- | 2 – 10 | | 14777360 | V2598 | E1b1~ | | 2253 |
| 2311 | I | 827 | 0 | 0 | 247 | 36 | 0 | M9- | 2 – 10 | | 14865477 | PF1654 | E1b1~ | | 2311 |
| 2706 | I | 827 | 0 | 0 | 247 | 36 | 0 | M9- | 2 – 10 | | 15541126 | PF1667 | E1b1~ | | 2706 |
| 2961 | III | 826 | 0 | 1 | 247 | 36 | 0 | M9- | 2 – 10 | | 16230262 | PF1684 | E1b1~ | | 2961 |
| 2969 | I | 827 | 0 | 0 | 247 | 36 | 0 | M9- | 2 – 10 | | 16252670 | PF1685 | E1b1~ | | 2969 |
| 3165 | III | 826 | 0 | 1 | 247 | 36 | 0 | M9- | 2 – 10 | | 16751001 | M5477 | E1b1~ | | 3165 |
| 3205 | I | 827 | 0 | 0 | 247 | 36 | 0 | M9- | 2 – 10 | | 16824270 | PF1702^ | E1b1~ | | 3205 |
| 3351 | I | 827 | 0 | 0 | 247 | 36 | 0 | M9- | 2 – 10 | | 17140219 | PF1719 | E1b1~ | | 3351 |
| 3591 | I | 827 | 0 | 0 | 247 | 36 | 0 | M9- | 2 – 10 | | 17737313 | PF1734 | E1b1~ | | 3591 |
| 3717 | I | 827 | 0 | 0 | 247 | 36 | 0 | M9- | 2 – 10 | | 17961512 | PF1744 | E1b1~ | | 3717 |
| 3793 | I | 827 | 0 | 0 | 247 | 36 | 0 | M9- | 2 – 10 | | 18118314 | PF1750 | E1b1~ | | 3793 |
| 3823 | I | 827 | 0 | 0 | 247 | 36 | 0 | M9- | 2 – 10 | | 18206187 | PF1753 | E1b1~ | | 3823 |
| 3865 | I | 827 | 0 | 0 | 247 | 36 | 0 | M9- | 2 – 10 | | 18538297 | PF1756 | E1b1~ | | 3865 |
| 3882 | I | 827 | 0 | 0 | 247 | 36 | 0 | M9- | 2 – 10 | | 18573243 | V3292 | E1b1~ | | 3882 |
| 4469 | I | 827 | 0 | 0 | 247 | 36 | 0 | M9- | 2 – 10 | | 21331627 | PF1799 | E1b1~ | | 4469 |
| 4638 | III | 825 | 0 | 2 | 247 | 36 | 0 | M9- | 2 – 10 | | 21700267 | PF1821 | E1b1~ | | 4638 |
| 5268 | I | 827 | 0 | 0 | 247 | 36 | 0 | M9- | 2 – 10 | | 22968980 | PF1880 | E1b1~ | | 5268 |
| 5430 | I | 827 | 0 | 0 | 247 | 36 | 0 | M9- | 2 – 10 | | 23351411 | PF1900 | E1b1~ | | 5430 |
| 5461 | I | 827 | 0 | 0 | 247 | 36 | 0 | M9- | 2 – 10 | | 23415873 | PF1902 | E1b1~ | | 5461 |
| 5661 | III | 825 | 0 | 2 | 247 | 36 | 0 | M9- | 2 – 10 | | 24401940 | PF1920 | E1b1~ | | 5661 |
| 543 | IV | 827 | 0 | 0 | 283 | 0 | 0 | Mono | Mono | | 6818291 | V38 | E1b1a | | 543 |
| 192 | IV | 827 | 0 | 0 | 283 | 0 | 0 | Mono | Mono | | 2911033 | Z1101 | E1b1a1 | | 192 |
| 1469 | IV | 827 | 0 | 0 | 283 | 0 | 0 | Mono | Mono | | 8587013 | Z1116 | E1b1a1 | | 1469 |
| 3272 | IV | 827 | 0 | 0 | 283 | 0 | 0 | Mono | Mono | | 16927082 | Z1125 | E1b1a1 | | 3272 |
| 3450 | IV | 815 | 0 | 12 | 279 | 0 | 4 | Mono | Mono | | 17415700 | Z1127 | E1b1a1 | | 3450 |
| 4445 | IV | 827 | 0 | 0 | 283 | 0 | 0 | Mono | Mono | | 21282471 | Z1130 | E1b1a1 | | 4445 |
| 4585 | IV | 827 | 0 | 0 | 283 | 0 | 0 | Mono | Mono | | 21612167 | PN1 | E1b1a1 | | 4585 |
| 4617 | IV | 827 | 0 | 0 | 283 | 0 | 0 | Mono | Mono | | 21646710 | Page106 | E1b1a1 | | 4617 |
| 4992 | IV | 827 | 0 | 0 | 283 | 0 | 0 | Mono | Mono | | 22181421 | Z1133 | E1b1a1 | | 4992 |
| 5146 | IV | 827 | 0 | 0 | 283 | 0 | 0 | Mono | Mono | | 22743597 | M291 | E1b1a1 | | 5146 |
| 5183 | IV | 827 | 0 | 0 | 283 | 0 | 0 | Mono | Mono | | 22784352 | Z1135 | E1b1a1 | A | 5183 |
| 113 | IV | 826 | 0 | 1 | 283 | 0 | 0 | Mono | Mono | | 2804671 | V1080 | E1b1a1~ | | 113 |
| 131 | IV | 827 | 0 | 0 | 283 | 0 | 0 | Mono | Mono | | 2826187 | M4714 | E1b1a1~ | | 131 |
| 170 | IV | 827 | 0 | 0 | 283 | 0 | 0 | Mono | Mono | | 2875993 | Z1100 | E1b1a1~ | | 170 |
| 466 | IV | 827 | 0 | 0 | 283 | 0 | 0 | Mono | Mono | | 6647632 | M4719 | E1b1a1~ | | 466 |
| 470 | IV | 826 | 0 | 1 | 282 | 0 | 1 | Mono | Mono | | 6660077 | M4720 | E1b1a1~ | | 470 |
| 609 | IV | 827 | 0 | 0 | 283 | 0 | 0 | Mono | Mono | | 6902475 | M4723 | E1b1a1~ | | 609 |
| 707 | IV | 827 | 0 | 0 | 283 | 0 | 0 | Mono | Mono | | 7028463 | CTS809 | E1b1a1~ | | 707 |
| 850 | IV | 827 | 0 | 0 | 283 | 0 | 0 | Mono | Mono | | 7401585 | Z1107 | E1b1a1~ | | 850 |
| 953 | IV | 827 | 0 | 0 | 283 | 0 | 0 | Mono | Mono | | 7658027 | Z1108 | E1b1a1~ | | 953 |
| 963 | IV | 827 | 0 | 0 | 283 | 0 | 0 | Mono | Mono | | 7672672 | M4734 | E1b1a1~ | | 963 |
| 979 | IV | 826 | 0 | 1 | 281 | 0 | 2 | Mono | Mono | | 7697222 | Z1109 | E1b1a1~ | | 979 |
| 996 | IV | 827 | 0 | 0 | 283 | 0 | 0 | Mono | Mono | | 7729538 | Z1110 | E1b1a1~ | | 996 |
| 1058 | IV | 824 | 0 | 3 | 283 | 0 | 0 | Mono | Mono | | 7843853 | M4738 | E1b1a1~ | | 1058 |
| 1109 | IV | 827 | 0 | 0 | 283 | 0 | 0 | Mono | Mono | | 7951143 | M4741 | E1b1a1~ | | 1109 |
| 1116 | IV | 826 | 0 | 1 | 281 | 0 | 2 | Mono | Mono | | 7965809 | M4742 | E1b1a1~ | | 1116 |
| 1177 | IV | 827 | 0 | 0 | 283 | 0 | 0 | Mono | Mono | | 8129921 | M4744 | E1b1a1~ | | 1177 |
| 1211 | IV | 799 | 0 | 28 | 275 | 0 | 8 | Mono | Mono | | 8219002 | V1725 | E1b1a1~ | | 1211 |
| 1288 | IV | 827 | 0 | 0 | 283 | 0 | 0 | Mono | Mono | | 8363884 | Z1113 | E1b1a1~ | | 1288 |
| 1329 | IV | 827 | 0 | 0 | 283 | 0 | 0 | Mono | Mono | | 8428087 | V1945 | E1b1a1~ | | 1329 |
| 1375 | IV | 827 | 0 | 0 | 283 | 0 | 0 | Mono | Mono | | 8467732 | V2055 | E1b1a1~ | | 1375 |
| 1448 | IV | 825 | 0 | 2 | 283 | 0 | 0 | Mono | Mono | | 8559787 | V2255 | E1b1a1~ | | 1448 |
| 1496 | IV | 827 | 0 | 0 | 283 | 0 | 0 | Mono | Mono | | 8613490 | V2355 | E1b1a1~ | | 1496 |
| 1531 | IV | 827 | 0 | 0 | 283 | 0 | 0 | Mono | Mono | | 8677609 | M4761 | E1b1a1~ | | 1531 |
| 1546 | IV | 827 | 0 | 0 | 283 | 0 | 0 | Mono | Mono | | 8697309 | V2468 | E1b1a1~ | | 1546 |
| 1565 | IV | 827 | 0 | 0 | 283 | 0 | 0 | Mono | Mono | | 8746440 | M4763 | E1b1a1~ | | 1565 |
| 1650 | IV | 827 | 0 | 0 | 283 | 0 | 0 | Mono | Mono | | 9101592 | M4766 | E1b1a1~ | | 1650 |
| 1658 | IV | 827 | 0 | 0 | 283 | 0 | 0 | Mono | Mono | | 9125158 | M4767 | E1b1a1~ | | 1658 |
| 1689 | IV | 827 | 0 | 0 | 283 | 0 | 0 | Mono | Mono | | 9438818 | M4769 | E1b1a1~ | | 1689 |
| 1753 | IV | 826 | 0 | 1 | 283 | 0 | 0 | Mono | Mono | | 9848099 | M4772 | E1b1a1~ | | 1753 |
| 1767 | IV | 827 | 0 | 0 | 283 | 0 | 0 | Mono | Mono | | 9882301 | L488 | E1b1a1~ | | 1767 |
| 1920 | IV | 827 | 0 | 0 | 283 | 0 | 0 | Mono | Mono | | 13849387 | Y1696 | E1b1a1~ | | 1920 |
| 1990 | IV | 827 | 0 | 0 | 283 | 0 | 0 | Mono | Mono | | 14074998 | CTS1804 | E1b1a1~ | | 1990 |
| 2002 | IV | 827 | 0 | 0 | 282 | 0 | 1 | Mono | Mono | | 14096577 | SY81 | E1b1a1~ | | 2002 |
| 2009 | IV | 827 | 0 | 0 | 282 | 0 | 1 | Mono | Mono | | 14108294 | Z1117 | E1b1a1~ | | 2009 |
| 2023 | IV | 827 | 0 | 0 | 283 | 0 | 0 | Mono | Mono | | 14145891 | M4778 | E1b1a1~ | | 2023 |
| 2033 | IV | 826 | 0 | 1 | 280 | 0 | 3 | Mono | Mono | | 14172225 | M4780 | E1b1a1~ | | 2033 |
| 2109 | IV | 802 | 0 | 25 | 275 | 0 | 8 | Mono | Mono | | 14334794 | M4788 | E1b1a1~ | | 2109 |
| 2151 | IV | 827 | 0 | 0 | 283 | 0 | 0 | Mono | Mono | | 14469450 | L610 | E1b1a1~ | | 2151 |
| 2219 | IV | 827 | 0 | 0 | 283 | 0 | 0 | Mono | Mono | | 14666569 | V2550 | E1b1a1~ | | 2219 |
| 2248 | IV | 827 | 0 | 0 | 283 | 0 | 0 | Mono | Mono | | 14773154 | M4797 | E1b1a1~ | | 2248 |
| 2270 | IV | 826 | 0 | 1 | 283 | 0 | 0 | Mono | Mono | | 14811240 | M4798 | E1b1a1~ | | 2270 |
| 2295 | IV | 823 | 0 | 4 | 282 | 0 | 1 | Mono | Mono | | 14841903 | M4799 | E1b1a1~ | | 2295 |

| | | | M9+ | | | | M9- | | | | | Build 37 | | ISOGG | |
|---|---|---|---|---|---|---|---|---|---|---|---|---|---|---|---|
| List | Stage | 0 | 1 | . | 0 | 1 | . | M9? | Clade(s) | Prob. | Position | Name | Subgroup Name | Amb. | List |
| 2543 | IV | 827 | 0 | 0 | 283 | 0 | 0 | Mono | Mono | | 15196452 | M4805 | E1b1a1~ | | 2543 |
| 2584 | IV | 827 | 0 | 0 | 283 | 0 | 0 | Mono | Mono | | 15315047 | V2884 | E1b1a1~ | | 2584 |
| 2590 | IV | 827 | 0 | 0 | 283 | 0 | 0 | Mono | Mono | | 15339461 | L491 | E1b1a1~ | | 2590 |
| 2647 | IV | 825 | 0 | 2 | 283 | 0 | 0 | Mono | Mono | | 15440731 | V2953 | E1b1a1~ | | 2647 |
| 2687 | IV | 827 | 0 | 0 | 283 | 0 | 0 | Mono | Mono | | 15510490 | Z1120 | E1b1a1~ | | 2687 |
| 2760 | IV | 827 | 0 | 0 | 283 | 0 | 0 | Mono | Mono | | 15603077 | V3068 | E1b1a1~ | | 2760 |
| 2770 | IV | 827 | 0 | 0 | 283 | 0 | 0 | Mono | Mono | | 15633954 | M4812 | E1b1a1~ | | 2770 |
| 2772 | IV | 827 | 0 | 0 | 283 | 0 | 0 | Mono | Mono | | 15635573 | CTS4415 | E1b1a1~ | | 2772 |
| 2818 | IV | 827 | 0 | 0 | 283 | 0 | 0 | Mono | Mono | | 15744876 | Z1121 | E1b1a1~ | | 2818 |
| 2880 | IV | 827 | 0 | 0 | 283 | 0 | 0 | Mono | Mono | | 15904338 | V3225 | E1b1a1~ | | 2880 |
| 2977 | IV | 826 | 0 | 1 | 283 | 0 | 0 | Mono | Mono | | 16264477 | M4820 | E1b1a1~ | | 2977 |
| 3009 | IV | 827 | 0 | 0 | 283 | 0 | 0 | Mono | Mono | | 16348910 | CTS5539 | E1b1a1~ | | 3009 |
| 3019 | IV | 827 | 0 | 0 | 283 | 0 | 0 | Mono | Mono | | 16371619 | M4822 | E1b1a1~ | | 3019 |
| 3119 | IV | 826 | 0 | 1 | 282 | 0 | 1 | Mono | Mono | | 16604139 | M4828 | E1b1a1~ | | 3119 |
| 3152 | IV | 827 | 0 | 0 | 283 | 0 | 0 | Mono | Mono | | 16713318 | M4832 | E1b1a1~ | | 3152 |
| 3196 | IV | 827 | 0 | 0 | 283 | 0 | 0 | Mono | Mono | | 16812870 | M4833 | E1b1a1~ | | 3196 |
| 3200 | IV | 827 | 0 | 0 | 283 | 0 | 0 | Mono | Mono | | 16817923 | Z1124 | E1b1a1~ | | 3200 |
| 3269 | IV | 827 | 0 | 0 | 283 | 0 | 0 | Mono | Mono | | 16922783 | M4834 | E1b1a1~ | | 3269 |
| 3271 | IV | 827 | 0 | 0 | 283 | 0 | 0 | Mono | Mono | | 16925100 | CTS6474 | E1b1a1~ | | 3271 |
| 3321 | IV | 827 | 0 | 0 | 283 | 0 | 0 | Mono | Mono | | 17040593 | M4836 | E1b1a1~ | | 3321 |
| 3361 | IV | 827 | 0 | 0 | 283 | 0 | 0 | Mono | Mono | | 17175506 | M4840 | E1b1a1~ | | 3361 |
| 3399 | IV | 827 | 0 | 0 | 283 | 0 | 0 | Mono | Mono | | 17286288 | CTS7095 | E1b1a1~ | | 3399 |
| 3481 | IV | 827 | 0 | 0 | 283 | 0 | 0 | Mono | Mono | | 17488242 | M4847 | E1b1a1~ | | 3481 |
| 3527 | IV | 827 | 0 | 0 | 283 | 0 | 0 | Mono | Mono | | 17583053 | CTS7641 | E1b1a1~ | | 3527 |
| 3551 | IV | 827 | 0 | 0 | 283 | 0 | 0 | Mono | Mono | | 17630793 | M4858 | E1b1a1~ | | 3551 |
| 3602 | IV | 827 | 0 | 0 | 283 | 0 | 0 | Mono | Mono | | 17758085 | M4862 | E1b1a1~ | | 3602 |
| 3645 | IV | 827 | 0 | 0 | 283 | 0 | 0 | Mono | Mono | | 17821258 | CTS8068 | E1b1a1~ | | 3645 |
| 3650 | IV | 827 | 0 | 0 | 283 | 0 | 0 | Mono | Mono | | 17842573 | CTS8123 | E1b1a1~ | | 3650 |
| 3742 | IV | 827 | 0 | 0 | 283 | 0 | 0 | Mono | Mono | | 18032753 | S27799 | E1b1a1~ | | 3742 |
| 3778 | IV | 827 | 0 | 0 | 283 | 0 | 0 | Mono | Mono | | 18083482 | M4867 | E1b1a1~ | | 3778 |
| 3845 | IV | 827 | 0 | 0 | 283 | 0 | 0 | Mono | Mono | | 18390543 | Y1705 | E1b1a1~ | | 3845 |
| 3853 | IV | 827 | 0 | 0 | 283 | 0 | 0 | Mono | Mono | | 18396767 | Y1746 | E1b1a1~ | | 3853 |
| 3860 | IV | 827 | 0 | 0 | 283 | 0 | 0 | Mono | Mono | | 18418203 | L501 | E1b1a1~ | | 3860 |
| 3875 | IV | 827 | 0 | 0 | 283 | 0 | 0 | Mono | Mono | | 18563482 | V3281 | E1b1a1~ | | 3875 |
| 3950 | IV | 827 | 0 | 0 | 283 | 0 | 0 | Mono | Mono | | 18736860 | V3455 | E1b1a1~ | | 3950 |
| 4003 | IV | 827 | 0 | 0 | 283 | 0 | 0 | Mono | Mono | | 18835911 | V3545 | E1b1a1~ | | 4003 |
| 4067 | IV | 827 | 0 | 0 | 283 | 0 | 0 | Mono | Mono | | 18962262 | M4876 | E1b1a1~ | | 4067 |
| 4103 | IV | 827 | 0 | 0 | 282 | 0 | 1 | Mono | Mono | | 19053340 | V3717 | E1b1a1~ | | 4103 |
| 4111 | IV | 827 | 0 | 0 | 283 | 0 | 0 | Mono | Mono | | 19061526 | M4879 | E1b1a1~ | | 4111 |
| 4173 | IV | 822 | 0 | 5 | 283 | 0 | 0 | Mono | Mono | | 19182415 | V3871 | E1b1a1~ | | 4173 |
| 4205 | IV | 805 | 0 | 22 | 278 | 0 | 5 | Mono | Mono | | 19238511 | CTS10066 | E1b1a1~ | | 4205 |
| 4219 | IV | 827 | 0 | 0 | 283 | 0 | 0 | Mono | Mono | | 19272435 | V3958 | E1b1a1~ | | 4219 |
| 4386 | IV | 827 | 0 | 0 | 283 | 0 | 0 | Mono | Mono | | 21121563 | S27751 | E1b1a1~ | | 4386 |
| 4411 | IV | 826 | 0 | 1 | 282 | 0 | 1 | Mono | Mono | | 21200625 | M4892 | E1b1a1~ | | 4411 |
| 4412 | IV | 822 | 0 | 5 | 281 | 0 | 2 | Mono | Mono | | 21201549 | M9307 | E1b1a1~ | | 4412 |
| 4481 | IV | 826 | 0 | 1 | 283 | 0 | 0 | Mono | Mono | | 21362700 | Y1604 | E1b1a1~ | | 4481 |
| 4556 | IV | 825 | 0 | 2 | 282 | 0 | 1 | Mono | Mono | | 21550026 | M4898 | E1b1a1~ | | 4556 |
| 4592 | IV | 826 | 0 | 1 | 283 | 0 | 0 | Mono | Mono | | 21620016 | M4901 | E1b1a1~ | | 4592 |
| 4622 | IV | 827 | 0 | 0 | 283 | 0 | 0 | Mono | Mono | | 21654965 | S27705 | E1b1a1~ | | 4622 |
| 4623 | IV | 827 | 0 | 0 | 283 | 0 | 0 | Mono | Mono | | 21661267 | M4903 | E1b1a1~ | | 4623 |
| 4628 | IV | 827 | 0 | 0 | 283 | 0 | 0 | Mono | Mono | | 21672442 | M4904 | E1b1a1~ | | 4628 |
| 4740 | IV | 827 | 0 | 0 | 283 | 0 | 0 | Mono | Mono | | 21792328 | M4905 | E1b1a1~ | | 4740 |
| 4950 | IV | 826 | 0 | 1 | 283 | 0 | 0 | Mono | Mono | | 22098943 | M4911 | E1b1a1~ | | 4950 |
| 4958 | IV | 827 | 0 | 0 | 282 | 0 | 1 | Mono | Mono | | 22115888 | M4913 | E1b1a1~ | | 4958 |
| 4976 | IV | 827 | 0 | 0 | 283 | 0 | 0 | Mono | Mono | | 22158010 | S27802 | E1b1a1~ | | 4976 |
| 4994 | IV | 824 | 0 | 3 | 281 | 0 | 2 | Mono | Mono | | 22187481 | M4915 | E1b1a1~ | | 4994 |
| 5001 | IV | 827 | 0 | 0 | 283 | 0 | 0 | Mono | Mono | | 22206380 | M4917 | E1b1a1~ | | 5001 |
| 5035 | IV | 827 | 0 | 0 | 283 | 0 | 0 | Mono | Mono | | 22521760 | Y1750 | E1b1a1~ | | 5035 |
| 5099 | IV | 827 | 0 | 0 | 283 | 0 | 0 | Mono | Mono | | 22672031 | CTS10638 | E1b1a1~ | | 5099 |
| 5212 | IV | 827 | 0 | 0 | 283 | 0 | 0 | Mono | Mono | | 22854091 | M4924 | E1b1a1~ | | 5212 |
| 5252 | IV | 824 | 0 | 3 | 281 | 0 | 2 | Mono | Mono | | 22933080 | M4925 | E1b1a1~ | | 5252 |
| 5338 | IV | 827 | 0 | 0 | 283 | 0 | 0 | Mono | Mono | | 23121708 | KHS1479188 | E1b1a1~ | | 5338 |
| 5359 | IV | 827 | 0 | 0 | 283 | 0 | 0 | Mono | Mono | | 23160649 | Z1136 | E1b1a1~ | | 5359 |
| 5366 | IV | 827 | 0 | 0 | 283 | 0 | 0 | Mono | Mono | | 23165895 | CTS11579 | E1b1a1~ | | 5366 |
| 5378 | IV | 827 | 0 | 0 | 283 | 0 | 0 | Mono | Mono | | 23241047 | CTS11729 | E1b1a1~ | | 5378 |
| 5387 | IV | 827 | 0 | 0 | 283 | 0 | 0 | Mono | Mono | | 23248930 | M4930 | E1b1a1~ | | 5387 |
| 5391 | IV | 827 | 0 | 0 | 283 | 0 | 0 | Mono | Mono | | 23254814 | Z1137 | E1b1a1~ | | 5391 |
| 5500 | IV | 816 | 0 | 11 | 277 | 0 | 6 | Mono | Mono | | 23532033 | M4935 | E1b1a1~ | | 5500 |
| 5508 | IV | 823 | 0 | 4 | 283 | 0 | 0 | Mono | Mono | | 23551491 | M4937 | E1b1a1~ | | 5508 |
| 5510 | IV | 827 | 0 | 0 | 283 | 0 | 0 | Mono | Mono | | 23554992 | M4938 | E1b1a1~ | | 5510 |
| 5562 | IV | 827 | 0 | 0 | 283 | 0 | 0 | Mono | Mono | | 23778425 | M4941 | E1b1a1~ | | 5562 |
| 5611 | IV | 827 | 0 | 0 | 283 | 0 | 0 | Mono | Mono | | 23900442 | Y6734 | E1b1a1~ | | 5611 |
| 5652 | IV | 825 | 0 | 2 | 283 | 0 | 0 | Mono | Mono | | 24379367 | S27745 | E1b1a1~ | | 5652 |
| 546 | IV | 822 | 0 | 5 | 282 | 0 | 1 | Mono | Mono | | 6832587 | M4722 | E1b1a1a1 | | 546 |
| 1045 | IV | 827 | 0 | 0 | 283 | 0 | 0 | Mono | Mono | | 7812445 | Z1111 | E1b1a1a1 | | 1045 |
| 1067 | IV | 827 | 0 | 0 | 283 | 0 | 0 | Mono | Mono | | 7855759 | S27755 | E1b1a1a1 | | 1067 |
| 1268 | IV | 827 | 0 | 0 | 283 | 0 | 0 | Mono | Mono | | 8315095 | Z1112 | E1b1a1a1 | | 1268 |
| 1333 | IV | 826 | 0 | 1 | 283 | 0 | 0 | Mono | Mono | | 8429714 | V1952 | E1b1a1a1 | | 1333 |
| 1832 | IV | 827 | 0 | 0 | 281 | 0 | 2 | Mono | Mono | | 13225125 | Y1694 | E1b1a1a1 | | 1832 |
| 2000 | IV | 827 | 0 | 0 | 283 | 0 | 0 | Mono | Mono | | 14092982 | CTS1847 | E1b1a1a1 | | 2000 |
| 2049 | IV | 821 | 0 | 6 | 279 | 0 | 4 | Mono | Mono | | 14200713 | M4781 | E1b1a1a1 | | 2049 |
| 2300 | IV | 827 | 0 | 0 | 283 | 0 | 0 | Mono | Mono | | 14850341 | P88 | E1b1a1a1 | | 2300 |
| 2358 | IV | 827 | 0 | 0 | 283 | 0 | 0 | Mono | Mono | | 14912545 | V2680 | E1b1a1a1 | | 2358 |
| 2518 | IV | 827 | 0 | 0 | 283 | 0 | 0 | Mono | Mono | | 15095580 | M4803 | E1b1a1a1 | | 2518 |
| 2600 | IV | 823 | 0 | 4 | 282 | 0 | 1 | Mono | Mono | | 15360167 | V2906 | E1b1a1a1 | | 2600 |
| 2604 | IV | 827 | 0 | 0 | 283 | 0 | 0 | Mono | Mono | | 15375740 | M4809 | E1b1a1a1 | | 2604 |
| 3128 | IV | 827 | 0 | 0 | 283 | 0 | 0 | Mono | Mono | | 16633829 | M4830 | E1b1a1a1 | | 3128 |
| 3144 | IV | 827 | 0 | 0 | 283 | 0 | 0 | Mono | Mono | | 16691822 | V43 | E1b1a1a1 | | 3144 |
| 3371 | IV | 827 | 0 | 0 | 283 | 0 | 0 | Mono | Mono | | 17201259 | M4841 | E1b1a1a1 | | 3371 |
| 3573 | IV | 827 | 0 | 0 | 283 | 0 | 0 | Mono | Mono | | 17678693 | Page66 | E1b1a1a1 | | 3573 |
| 4202 | IV | 827 | 0 | 0 | 283 | 0 | 0 | Mono | Mono | | 19236447 | P182 | E1b1a1a1 | | 4202 |
| 4635 | IV | 827 | 0 | 0 | 283 | 0 | 0 | Mono | Mono | | 21688300 | Z1132 | E1b1a1a1 | | 4635 |
| 5089 | IV | 827 | 0 | 0 | 282 | 0 | 1 | Mono | Mono | | 22634764 | Y1753 | E1b1a1a1 | | 5089 |
| 5106 | IV | 827 | 0 | 0 | 283 | 0 | 0 | Mono | Mono | | 22684420 | CTS10659 | E1b1a1a1 | | 5106 |
| 5203 | IV | 827 | 0 | 0 | 283 | 0 | 0 | Mono | Mono | | 22833310 | CTS10914 | E1b1a1a1 | | 5203 |
| 5379 | IV | 827 | 0 | 0 | 283 | 0 | 0 | Mono | Mono | | 23241568 | KHS1381212 | E1b1a1a1 | | 5379 |

| | | M9+ | | | M9- | | | | | | Build 37 | | ISOGG | | |
|---|---|---|---|---|---|---|---|---|---|---|---|---|---|---|---|
| List | Stage | 0 | 1 | . | 0 | 1 | . | M9? | Clade(s) | Prob. | Position | Name | Subgroup Name | Amb. | List |
| 5664 | IV | 825 | 0 | 2 | 283 | 0 | 0 | Mono | Mono | | 24407921 | M4948 | E1b1a1a1 | | 5664 |
| 5681 | IV | 825 | 0 | 2 | 283 | 0 | 0 | Mono | Mono | | 24461432 | M4949 | E1b1a1a1 | A | 5681 |
| 755 | IV | 827 | 0 | 0 | 283 | 0 | 0 | Mono | Mono | | 7136219 | M4727 | E1b1a1a1a | | 755 |
| 931 | IV | 825 | 0 | 2 | 283 | 0 | 0 | Mono | Mono | | 7620822 | M4732 | E1b1a1a1a | | 931 |
| 4501 | IV | 827 | 0 | 0 | 283 | 0 | 0 | Mono | Mono | | 21401576 | M4895 | E1b1a1a1a | | 4501 |
| 4616 | IV | 827 | 0 | 0 | 283 | 0 | 0 | Mono | Mono | | 21646675 | Page27 | E1b1a1a1a1a | | 4616 |
| 4673 | IV | 827 | 0 | 0 | 283 | 0 | 0 | Mono | Mono | | 21736304 | M58 | E1b1a1a1a1a | | 4673 |
| 4655 | IV | 827 | 0 | 0 | 283 | 0 | 0 | Mono | Mono | | 21722998 | M10 | E1b1a1a1a1b~ | | 4655 |
| 1601 | IV | 827 | 0 | 0 | 283 | 0 | 0 | Mono | Mono | | 8852858 | L485 | E1b1a1a1a1c | | 1601 |
| 1313 | IV | 827 | 0 | 0 | 282 | 0 | 1 | Mono | Mono | | 8402223 | Z1291 | E1b1a1a1a1c1 | | 1313 |
| 2207 | IV | 827 | 0 | 0 | 283 | 0 | 0 | Mono | Mono | | 14636457 | L514 | E1b1a1a1a1c1 | | 2207 |
| 5501 | IV | 827 | 0 | 0 | 283 | 0 | 0 | Mono | Mono | | 23533199 | M4708 | E1b1a1a1a1c1 | | 5501 |
| 3915 | IV | 819 | 0 | 8 | 280 | 0 | 3 | Mono | Mono | | 18635977 | M4693 | E1b1a1a1a1c1a | | 3915 |
| 4365 | IV | 827 | 0 | 0 | 283 | 0 | 0 | Mono | Mono | | 21071521 | M4696 | E1b1a1a1a1c1a | | 4365 |
| 4547 | IV | 692 | 0 | 135 | 231 | 0 | 52 | Mono | Mono | | 21514118 | M4699 | E1b1a1a1a1c1a | | 4547 |
| 4658 | IV | 827 | 0 | 0 | 283 | 0 | 0 | Mono | Mono | | 21726388 | Z1712.1 | E1b1a1a1a1c1a | | 4658 |
| 4923 | IV | 827 | 0 | 0 | 283 | 0 | 0 | Mono | Mono | | 22009958 | M4700 | E1b1a1a1a1c1a | | 4923 |
| 5033 | IV | 827 | 0 | 0 | 283 | 0 | 0 | Mono | Mono | | 22515871 | Z22334 | E1b1a1a1a1c1a | | 5033 |
| 5360 | IV | 822 | 0 | 5 | 283 | 0 | 0 | Mono | Mono | | 23161339 | CTS11559 | E1b1a1a1a1c1a | | 5360 |
| 736 | IV | 827 | 0 | 0 | 283 | 0 | 0 | Mono | Mono | | 7078953 | M4684 | E1b1a1a1a1c1a1 | | 736 |
| 2446 | IV | 827 | 0 | 0 | 283 | 0 | 0 | Mono | Mono | | 15019613 | P86 | E1b1a1a1a1c1a1 | | 2446 |
| 3560 | IV | 827 | 0 | 0 | 283 | 0 | 0 | Mono | Mono | | 17656496 | Z1293 | E1b1a1a1a1c1a1 | | 3560 |
| 912 | IV | 827 | 0 | 0 | 283 | 0 | 0 | Mono | Mono | | 7595638 | V1245 | E1b1a1a1a1c1a1a | | 912 |
| 2804 | IV | 827 | 0 | 0 | 283 | 0 | 0 | Mono | Mono | | 15708715 | M4675 | E1b1a1a1a1c1a1a | | 2804 |
| 2967 | IV | 827 | 0 | 0 | 283 | 0 | 0 | Mono | Mono | | 16251357 | U174 | E1b1a1a1a1c1a1a | | 2967 |
| 4578 | IV | 827 | 0 | 0 | 283 | 0 | 0 | Mono | Mono | | 21605152 | Z1715 | E1b1a1a1a1c1a1a | | 4578 |
| 740 | IV | 827 | 0 | 0 | 283 | 0 | 0 | Mono | Mono | | 7087671 | M4670 | E1b1a1a1a1c1a1a3 | | 740 |
| 2550 | IV | 826 | 0 | 1 | 283 | 0 | 0 | Mono | Mono | | 15209074 | M4673 | E1b1a1a1a1c1a1a3 | | 2550 |
| 3449 | IV | 827 | 0 | 0 | 283 | 0 | 0 | Mono | Mono | | 17413565 | Z1714 | E1b1a1a1a1c1a1a3 | | 3449 |
| 602 | IV | 827 | 0 | 0 | 283 | 0 | 0 | Mono | Mono | | 6894595 | M4437 | E1b1a1a1a1c1a1a3a1a | | 602 |
| 3935 | IV | 827 | 0 | 0 | 283 | 0 | 0 | Mono | Mono | | 18683145 | M4451 | E1b1a1a1a1c1a1a3a1a1 | | 3935 |
| 510 | IV | 827 | 0 | 0 | 283 | 0 | 0 | Mono | Mono | | 6740171 | P113 | E1b1a1a1a1c1a1a3a1a1a1~ | | 510 |
| 101 | IV | 824 | 0 | 3 | 281 | 0 | 2 | Mono | Mono | | 2772928 | V199 | E1b1a1a1a1c1a1a3m3~ | A | 101 |
| 761 | IV | 827 | 0 | 0 | 283 | 0 | 0 | Mono | Mono | | 7144164 | M4260 | E1b1a1a1a1c1b | | 761 |
| 982 | IV | 827 | 0 | 0 | 283 | 0 | 0 | Mono | Mono | | 7703243 | Z1138 | E1b1a1a1a1c1b | | 982 |
| 1063 | IV | 827 | 0 | 0 | 283 | 0 | 0 | Mono | Mono | | 7851549 | V1473 | E1b1a1a1a1c1b | | 1063 |
| 1238 | IV | 826 | 0 | 1 | 283 | 0 | 0 | Mono | Mono | | 8258764 | V1760 | E1b1a1a1a1c1b | | 1238 |
| 1239 | IV | 827 | 0 | 0 | 283 | 0 | 0 | Mono | Mono | | 8259636 | Z1139 | E1b1a1a1a1c1b | | 1239 |
| 1246 | IV | 827 | 0 | 0 | 282 | 0 | 1 | Mono | Mono | | 8267895 | L515 | E1b1a1a1a1c1b | | 1246 |
| 1280 | IV | 827 | 0 | 0 | 283 | 0 | 0 | Mono | Mono | | 8349250 | Z22224 | E1b1a1a1a1c1b | | 1280 |
| 1744 | IV | 827 | 0 | 0 | 283 | 0 | 0 | Mono | Mono | | 9833459 | M4268 | E1b1a1a1a1c1b | | 1744 |
| 1766 | IV | 807 | 0 | 20 | 274 | 0 | 9 | Mono | Mono | | 9879569 | Z1893 | E1b1a1a1a1c1b | | 1766 |
| 2437 | IV | 827 | 0 | 0 | 283 | 0 | 0 | Mono | Mono | | 15016438 | M263.2 | E1b1a1a1a1c1b | | 2437 |
| 3072 | IV | 827 | 0 | 0 | 283 | 0 | 0 | Mono | Mono | | 16491559 | M4282 | E1b1a1a1a1c1b | | 3072 |
| 4128 | IV | 827 | 0 | 0 | 283 | 0 | 0 | Mono | Mono | | 19105726 | L516 | E1b1a1a1a1c1b | | 4128 |
| 4340 | IV | 827 | 0 | 0 | 283 | 0 | 0 | Mono | Mono | | 20821220 | Z22228 | E1b1a1a1a1c1b | | 4340 |
| 4574 | IV | 827 | 0 | 0 | 283 | 0 | 0 | Mono | Mono | | 21593470 | L517 | E1b1a1a1a1c1b | | 4574 |
| 5512 | IV | 826 | 0 | 1 | 283 | 0 | 0 | Mono | Mono | | 23555194 | M4298 | E1b1a1a1a1c1b | | 5512 |
| 2328 | IV | 827 | 0 | 0 | 283 | 0 | 0 | Mono | Mono | | 14887218 | Z1894 | E1b1a1a1a1c1b1 | | 2328 |
| 3052 | IV | 826 | 0 | 1 | 283 | 0 | 0 | Mono | Mono | | 16450352 | M4281 | E1b1a1a1a1c1b2a | | 3052 |
| 5226 | IV | 827 | 0 | 0 | 283 | 0 | 0 | Mono | Mono | | 22900330 | Z22243 | E1b1a1a1a1c1b2a | | 5226 |
| 5467 | IV | 827 | 0 | 0 | 283 | 0 | 0 | Mono | Mono | | 23443721 | M4297 | E1b1a1a1a1c1b2a | | 5467 |
| 776 | IV | 826 | 0 | 1 | 282 | 0 | 1 | Mono | Mono | | 7168798 | Z15939 | E1b1a1a1a1c2 | | 776 |
| 866 | IV | 826 | 0 | 1 | 283 | 0 | 0 | Mono | Mono | | 7526997 | Z15940 | E1b1a1a1a1c2 | | 866 |
| 1131 | IV | 827 | 0 | 0 | 283 | 0 | 0 | Mono | Mono | | 8019387 | Z15941 | E1b1a1a1a1c2 | | 1131 |
| 1596 | IV | 827 | 0 | 0 | 283 | 0 | 0 | Mono | Mono | | 8841448 | Z15943 | E1b1a1a1a1c2 | | 1596 |
| 2056 | IV | 826 | 0 | 1 | 283 | 0 | 0 | Mono | Mono | | 14210641 | Z15953 | E1b1a1a1a1c2 | | 2056 |
| 3108 | IV | 827 | 0 | 0 | 283 | 0 | 0 | Mono | Mono | | 16575643 | CTS5961 | E1b1a1a1a1c2 | | 3108 |
| 3690 | III | 822 | 1 | 4 | 283 | 0 | 0 | M9+ | 65 | | 17914149 | Z15956 | E1b1a1a1a1c2 | W | 3690 |
| 4135 | IV | 827 | 0 | 0 | 283 | 0 | 0 | Mono | Mono | | 19118921 | CTS9883 | E1b1a1a1a1c2 | | 4135 |
| 4319 | IV | 827 | 0 | 0 | 283 | 0 | 0 | Mono | Mono | | 19538933 | CTS10560 | E1b1a1a1a1c2 | | 4319 |
| 4651 | IV | 827 | 0 | 0 | 283 | 0 | 0 | Mono | Mono | | 21718388 | Z15958 | E1b1a1a1a1c2 | | 4651 |
| 4970 | IV | 812 | 0 | 15 | 279 | 0 | 4 | Mono | Mono | | 22145251 | ZS6144 | E1b1a1a1a1c2 | | 4970 |
| 5211 | IV | 825 | 0 | 2 | 283 | 0 | 0 | Mono | Mono | | 22850513 | Z15962 | E1b1a1a1a1c2 | | 5211 |
| 1719 | IV | 823 | 0 | 4 | 282 | 0 | 1 | Mono | Mono | | 9765455 | Z15947 | E1b1a1a1a1c2~ | | 1719 |
| 2090 | IV | 827 | 0 | 0 | 283 | 0 | 0 | Mono | Mono | | 14287415 | Z16002 | E1b1a1a1a1c2b | | 2090 |
| 5309 | IV | 827 | 0 | 0 | 283 | 0 | 0 | Mono | Mono | | 23065208 | Z16044 | E1b1a1a1a1c2b2 | | 5309 |
| 5509 | IV | 827 | 0 | 0 | 283 | 0 | 0 | Mono | Mono | | 23553620 | Z16045 | E1b1a1a1a1c2b2 | | 5509 |
| 974 | IV | 827 | 0 | 0 | 283 | 0 | 0 | Mono | Mono | | 7689689 | Z16056 | E1b1a1a1a1c2c | | 974 |
| 2258 | IV | 821 | 0 | 6 | 282 | 0 | 1 | Mono | Mono | | 14785945 | CTS3274 | E1b1a1a1a1c2c | | 2258 |
| 3379 | IV | 825 | 0 | 2 | 283 | 0 | 0 | Mono | Mono | | 17217538 | CTS6973 | E1b1a1a1a1c2c1 | | 3379 |
| 5460 | IV | 827 | 0 | 0 | 283 | 0 | 0 | Mono | Mono | | 23414794 | CTS12004 | E1b1a1a1a1c2c1 | | 5460 |
| 180 | IV | 827 | 0 | 0 | 283 | 0 | 0 | Mono | Mono | | 2888392 | CTS308 | E1b1a1a1a1c2c1a | A | 180 |
| 596 | IV | 827 | 0 | 0 | 283 | 0 | 0 | Mono | Mono | | 6888782 | Z6010 | E1b1a1a1a1c2c2 | | 596 |
| 1083 | IV | 827 | 0 | 0 | 283 | 0 | 0 | Mono | Mono | | 7903471 | Z17016 | E1b1a1a1a1c2c2 | | 1083 |
| 114 | IV | 827 | 0 | 0 | 283 | 0 | 0 | Mono | Mono | | 2806457 | Z6015 | E1b1a1a1a1c2c3a1b | | 114 |
| 1508 | IV | 827 | 0 | 0 | 283 | 0 | 0 | Mono | Mono | | 8633967 | Z16104 | E1b1a1a1a1c2c3a1b | | 1508 |
| 5182 | IV | 826 | 0 | 1 | 283 | 0 | 0 | Mono | Mono | | 22784044 | Z16155 | E1b1a1a1a1c2c3a2a | | 5182 |
| 140 | IV | 827 | 0 | 0 | 283 | 0 | 0 | Mono | Mono | | 2831112 | V1112^ | E1b1a1a1a2 | | 140 |
| 1012 | IV | 827 | 0 | 0 | 283 | 0 | 0 | Mono | Mono | | 7747954 | M4231 | E1b1a1a1a2 | | 1012 |
| 1349 | IV | 827 | 0 | 0 | 283 | 0 | 0 | Mono | Mono | | 8446807 | V2003^ | E1b1a1a1a2 | | 1349 |
| 2970 | IV | 827 | 0 | 0 | 283 | 0 | 0 | Mono | Mono | | 16253694 | U175 | E1b1a1a1a2a | | 2970 |
| 731 | IV | 827 | 0 | 0 | 283 | 0 | 0 | Mono | Mono | | 7074820 | M4227 | E1b1a1a1a2a1 | | 731 |
| 972 | IV | 827 | 0 | 0 | 283 | 0 | 0 | Mono | Mono | | 7687135 | M4230 | E1b1a1a1a2a1 | | 972 |
| 1568 | IV | 827 | 0 | 0 | 283 | 0 | 0 | Mono | Mono | | 8765851 | M4236 | E1b1a1a1a2a1 | A | 1568 |
| 2561 | IV | 827 | 0 | 0 | 283 | 0 | 0 | Mono | Mono | | 15239487 | CTS3878 | E1b1a1a1a2a1 | | 2561 |
| 2617 | IV | 827 | 0 | 0 | 283 | 0 | 0 | Mono | Mono | | 15401647 | V2926 | E1b1a1a1a2a1 | | 2617 |
| 2730 | IV | 827 | 0 | 0 | 283 | 0 | 0 | Mono | Mono | | 15579215 | P277 | E1b1a1a1a2a1 | | 2730 |
| 2857 | IV | 825 | 0 | 2 | 281 | 0 | 2 | Mono | Mono | | 15821083 | V3154 | E1b1a1a1a2a1 | | 2857 |
| 2879 | IV | 827 | 0 | 0 | 283 | 0 | 0 | Mono | Mono | | 15903731 | V3224 | E1b1a1a1a2a1 | | 2879 |
| 4554 | IV | 827 | 0 | 0 | 283 | 0 | 0 | Mono | Mono | | 21541514 | M4254 | E1b1a1a1a2a1 | | 4554 |
| 1358 | IV | 824 | 0 | 3 | 283 | 0 | 0 | Mono | Mono | | 8458731 | V2027 | E1b1a1a1a2a1~ | | 1358 |
| 3403 | IV | 827 | 0 | 0 | 283 | 0 | 0 | Mono | Mono | | 17294958 | U209 | E1b1a1a1a2a1~ | A | 3403 |
| 1061 | IV | 825 | 0 | 2 | 283 | 0 | 0 | Mono | Mono | | 7850169 | M4232 | E1b1a1a1a2a1a | | 1061 |
| 1145 | IV | 827 | 0 | 0 | 283 | 0 | 0 | Mono | Mono | | 8048543 | Z1807 | E1b1a1a1a2a1a | | 1145 |
| 5074 | IV | 827 | 0 | 0 | 283 | 0 | 0 | Mono | Mono | | 22596423 | M4255 | E1b1a1a1a2a1a | | 5074 |

| | | M9+ | | | M9- | | | | | | Build 37 | | ISOGG | | |
|---|---|---|---|---|---|---|---|---|---|---|---|---|---|---|---|
| List | Stage | 0 | 1 | . | 0 | 1 | . | M9? | Clade(s) | Prob. | Position | Name | Subgroup Name | Amb. | List |
| 5536 | IV | 801 | 0 | 26 | 263 | 0 | 20 | Mono | Mono | | 23621784 | Z1720 | E1b1a1a1a2a1a3 | | 5536 |
| 901 | IV | 785 | 0 | 42 | 272 | 0 | 11 | Mono | Mono | | 7572456 | M4215 | E1b1a1a1a2a1a3a | | 901 |
| 2519 | IV | 827 | 0 | 0 | 283 | 0 | 0 | Mono | Mono | | 15096961 | Z1776 | E1b1a1a1a2a1a3a | | 2519 |
| 3895 | IV | 827 | 0 | 0 | 283 | 0 | 0 | Mono | Mono | | 18599285 | M4221 | E1b1a1a1a2a1a3a | | 3895 |
| 5300 | IV | 827 | 0 | 0 | 283 | 0 | 0 | Mono | Mono | | 23041577 | M4223 | E1b1a1a1a2a1a3a | | 5300 |
| 5327 | IV | 825 | 0 | 2 | 283 | 0 | 0 | Mono | Mono | | 23090298 | M4224 | E1b1a1a1a2a1a3a | | 5327 |
| 5264 | IV | 827 | 0 | 0 | 283 | 0 | 0 | Mono | Mono | | 22956502 | M4222 | E1b1a1a1a2a1a3a~ | | 5264 |
| 664 | IV | 827 | 0 | 0 | 283 | 0 | 0 | Mono | Mono | | 6942201 | M4102 | E1b1a1a1a2a1a3a1 | | 664 |
| 4704 | IV | 826 | 0 | 1 | 282 | 0 | 1 | Mono | Mono | | 21761058 | M4062 | E1b1a1a1a2a1a3a1 | | 4704 |
| 4198 | IV | 827 | 0 | 0 | 283 | 0 | 0 | Mono | Mono | | 19234486 | M4090 | E1b1a1a1a2a1a3a1~ | | 4198 |
| 3334 | IV | 827 | 0 | 0 | 283 | 0 | 0 | Mono | Mono | | 17102607 | M4041 | E1b1a1a1a2a1a3b | | 3334 |
| 23 | IV | 825 | 0 | 2 | 283 | 0 | 0 | Mono | Mono | | 2662740 | M3988 | E1b1a1a1a2a1a3b1 | | 23 |
| 1130 | IV | 803 | 0 | 24 | 273 | 0 | 10 | Mono | Mono | | 8017283 | M3991 | E1b1a1a1a2a1a3b1 | | 1130 |
| 2106 | IV | 827 | 0 | 0 | 283 | 0 | 0 | Mono | Mono | | 14332342 | Z1717 | E1b1a1a1a2a1a3b1 | | 2106 |
| 4614 | IV | 827 | 0 | 0 | 283 | 0 | 0 | Mono | Mono | | 21646058 | U290 | E1b1a1a1a2a1a3b1a | | 4614 |
| 2339 | IV | 827 | 0 | 0 | 283 | 0 | 0 | Mono | Mono | | 14898117 | L651 | E1b1a1a1a2a1a3b1a19 | | 2339 |
| 5149 | IV | 827 | 0 | 0 | 283 | 0 | 0 | Mono | Mono | | 22744795 | M292 | E1b1a1a1a2a1a3b1a19a~ | | 5149 |
| 70 | IV | 826 | 0 | 1 | 282 | 0 | 1 | Mono | Mono | | 2728042 | Z1725 | E1b1a1a1a2a1a3b1a2 | | 70 |
| 3308 | IV | 827 | 0 | 0 | 283 | 0 | 0 | Mono | Mono | | 17010888 | CTS6613 | E1b1a1a1a2a1a3b1a2~ | | 3308 |
| 4828 | IV | 827 | 0 | 0 | 283 | 0 | 0 | Mono | Mono | | 21892677 | M154 | E1b1a1a1a2a1a3b1b~ | | 4828 |
| 5317 | IV | 826 | 0 | 1 | 283 | 0 | 0 | Mono | Mono | | 23073358 | Z1789 | E1b1a1a1a2a1a3b1d1c | | 5317 |
| 2240 | IV | 827 | 0 | 0 | 283 | 0 | 0 | Mono | Mono | | 14762903 | Z1781 | E1b1a1a1a2a1a3b1d1c1 | | 2240 |
| 3626 | IV | 827 | 0 | 0 | 283 | 0 | 0 | Mono | Mono | | 17800068 | M3850 | E1b1a1a1a2a1a3b1d1c1~ | | 3626 |
| 3627 | IV | 827 | 0 | 0 | 283 | 0 | 0 | Mono | Mono | | 17800185 | M3851 | E1b1a1a1a2a1a3b1d1c1a | | 3627 |
| 4523 | IV | 827 | 0 | 0 | 283 | 0 | 0 | Mono | Mono | | 21454514 | M3852 | E1b1a1a1a2a1a3b1d1c1a | | 4523 |
| 1455 | II | 827 | 0 | 0 | 281 | 2 | 0 | M9- | 5 | | 8567314 | Z12348^ | E1b1a1a1a2a1a3b3 | A,W | 1455 |
| 2082 | IV | 827 | 0 | 0 | 283 | 0 | 0 | Mono | Mono | | 14274995 | CTS2359 | E1b1a1a1a2a1a3b3 | | 2082 |
| 2431 | IV | 822 | 0 | 5 | 281 | 0 | 2 | Mono | Mono | | 15004487 | Z21813 | E1b1a1a1a2a1a3b3 | | 2431 |
| 3001 | IV | 827 | 0 | 0 | 283 | 0 | 0 | Mono | Mono | | 16334728 | Z21814^ | E1b1a1a1a2a1a3b3 | | 3001 |
| 3063 | III | 825 | 1 | 1 | 283 | 0 | 0 | M9+ | NU | | 16472227 | CTS5768 | E1b1a1a1a2a1a3b3 | W | 3063 |
| 3672 | IV | 827 | 0 | 0 | 283 | 0 | 0 | Mono | Mono | | 17881472 | Z21815 | E1b1a1a1a2a1a3b3 | | 3672 |
| 4470 | IV | 826 | 0 | 1 | 283 | 0 | 0 | Mono | Mono | | 21331875 | Z21817^ | E1b1a1a1a2a1a3b3 | | 4470 |
| 5410 | IV | 827 | 0 | 0 | 283 | 0 | 0 | Mono | Mono | | 23273932 | CTS11789^ | E1b1a1a1a2a1a3b3 | | 5410 |
| 5572 | IV | 827 | 0 | 0 | 283 | 0 | 0 | Mono | Mono | | 23800883 | Z21820^ | E1b1a1a1a2a1a3b3 | | 5572 |
| 5455 | IV | 827 | 0 | 0 | 283 | 0 | 0 | Mono | Mono | | 23402870 | Z21819 | E1b1a1a1a2a1a3b3~ | | 5455 |
| 169 | IV | 827 | 0 | 0 | 283 | 0 | 0 | Mono | Mono | | 2875527 | M329 | E1b1a2 | | 169 |
| 132 | III | 801 | 0 | 26 | 243 | 33 | 7 | M9- | 2 – 10 | MD | 2827409 | PF1450 | E1b1b | | 132 |
| 1085 | III | 826 | 0 | 1 | 247 | 36 | 0 | M9- | 2 – 10 | | 7905833 | PF1537 | E1b1b | | 1085 |
| 1086 | I | 827 | 0 | 0 | 247 | 36 | 0 | M9- | 2 – 10 | | 7906010 | PF1538 | E1b1b | | 1086 |
| 1136 | I | 827 | 0 | 0 | 247 | 36 | 0 | M9- | 2 – 10 | | 8028896 | V1575 | E1b1b | | 1136 |
| 1669 | I | 827 | 0 | 0 | 247 | 36 | 0 | M9- | 2 – 10 | | 9160895 | PF1572 | E1b1b | | 1669 |
| 2126 | III | 826 | 0 | 1 | 247 | 36 | 0 | M9- | 2 – 10 | | 14377079 | PF1637 | E1b1b | | 2126 |
| 2443 | I | 827 | 0 | 0 | 247 | 36 | 0 | M9- | 2 – 10 | | 15019092 | PF1943 | E1b1b | | 2443 |
| 2583 | III | 826 | 0 | 1 | 247 | 36 | 0 | M9- | 2 – 10 | | 15313925 | PF1664 | E1b1b | | 2583 |
| 2660 | I | 827 | 0 | 0 | 247 | 36 | 0 | M9- | 2 – 10 | | 15467824 | PF1942 | E1b1b | | 2660 |
| 3006 | I | 827 | 0 | 0 | 247 | 36 | 0 | M9- | 2 – 10 | | 16346117 | PF1689 | E1b1b | | 3006 |
| 3068 | I | 827 | 0 | 0 | 247 | 36 | 0 | M9- | 2 – 10 | | 16484019 | PF1692 | E1b1b | | 3068 |
| 3374 | III | 826 | 0 | 1 | 247 | 36 | 0 | M9- | 2 – 10 | | 17209379 | M5221 | E1b1b | | 3374 |
| 3547 | I | 827 | 0 | 0 | 247 | 36 | 0 | M9- | 2 – 10 | | 17612676 | PF1729 | E1b1b | | 3547 |
| 3751 | I | 827 | 0 | 0 | 247 | 36 | 0 | M9- | 2 – 10 | | 18045601 | PF1746.2 | E1b1b | A | 3751 |
| 3908 | I | 827 | 0 | 0 | 247 | 36 | 0 | M9- | 2 – 10 | | 18617336 | V3341 | E1b1b | | 3908 |
| 3916 | I | 827 | 0 | 0 | 247 | 36 | 0 | M9- | 2 – 10 | | 18637397 | PF1763 | E1b1b | | 3916 |
| 4310 | I | 827 | 0 | 0 | 247 | 36 | 0 | M9- | 2 – 10 | | 19503700 | V4197 | E1b1b | | 4310 |
| 4507 | I | 827 | 0 | 0 | 247 | 36 | 0 | M9- | 2 – 10 | | 21410887 | PF1807 | E1b1b | | 4507 |
| 4608 | I | 827 | 0 | 0 | 247 | 36 | 0 | M9- | 2 – 10 | | 21643140 | PF1818 | E1b1b | | 4608 |
| 4858 | I | 827 | 0 | 0 | 247 | 36 | 0 | M9- | 2 – 10 | | 21903853 | PF1825 | E1b1b | | 4858 |
| 4870 | I | 827 | 0 | 0 | 247 | 36 | 0 | M9- | 2 – 10 | | 21909401 | PF1826 | E1b1b | | 4870 |
| 4913 | I | 827 | 0 | 0 | 247 | 36 | 0 | M9- | 2 – 10 | | 21983698 | PF1831 | E1b1b | | 4913 |
| 5098 | I | 827 | 0 | 0 | 247 | 36 | 0 | M9- | 2 – 10 | | 22671606 | M5327 | E1b1b | | 5098 |
| 5110 | I | 827 | 0 | 0 | 247 | 36 | 0 | M9- | 2 – 10 | | 22700429 | PF1875 | E1b1b | | 5110 |
| 5288 | III | 824 | 0 | 3 | 247 | 36 | 0 | M9- | 2 – 10 | | 23021729 | PF1883 | E1b1b | | 5288 |
| 726 | I | 827 | 0 | 0 | 247 | 36 | 0 | M9- | 2 – 10 | | 7071937 | PF1513 | E1b1b1 | | 726 |
| 842 | I | 827 | 0 | 0 | 247 | 36 | 0 | M9- | 2 – 10 | | 7377029 | PF1525 | E1b1b1 | | 842 |
| 924 | I | 827 | 0 | 0 | 247 | 36 | 0 | M9- | 2 – 10 | | 7605031 | V1257 | E1b1b1 | | 924 |
| 965 | I | 827 | 0 | 0 | 247 | 36 | 0 | M9- | 2 – 10 | | 7678694 | PF1532 | E1b1b1 | | 965 |
| 990 | I | 827 | 0 | 0 | 247 | 36 | 0 | M9- | 2 – 10 | | 7721674 | M5078 | E1b1b1 | | 990 |
| 1158 | I | 827 | 0 | 0 | 247 | 36 | 0 | M9- | 2 – 10 | | 8082372 | V1602 | E1b1b1 | | 1158 |
| 1187 | I | 827 | 0 | 0 | 247 | 36 | 0 | M9- | 2 – 10 | | 8143135 | V1659 | E1b1b1 | | 1187 |
| 1495 | I | 827 | 0 | 0 | 247 | 36 | 0 | M9- | 2 – 10 | | 8613281 | V2353 | E1b1b1 | | 1495 |
| 1609 | III | 824 | 0 | 3 | 247 | 36 | 0 | M9- | 2 – 10 | | 8880108 | PF1566 | E1b1b1 | | 1609 |
| 1682 | III | 826 | 0 | 1 | 247 | 36 | 0 | M9- | 2 – 10 | | 9389773 | PF1575 | E1b1b1 | | 1682 |
| 2062 | I | 827 | 0 | 0 | 247 | 36 | 0 | M9- | 2 – 10 | | 14221285 | PF1629 | E1b1b1 | | 2062 |
| 2099 | III | 826 | 0 | 1 | 247 | 36 | 0 | M9- | 2 – 10 | | 14320934 | PF1635 | E1b1b1 | | 2099 |
| 2129 | I | 827 | 0 | 0 | 247 | 36 | 0 | M9- | 2 – 10 | | 14393170 | PF1638 | E1b1b1 | | 2129 |
| 2421 | I | 827 | 0 | 0 | 247 | 36 | 0 | M9- | 2 – 10 | | 14983106 | V2737 | E1b1b1 | | 2421 |
| 2464 | I | 827 | 0 | 0 | 247 | 36 | 0 | M9- | 2 – 10 | | 15026633 | PF1657 | E1b1b1 | | 2464 |
| 2511 | I | 827 | 0 | 0 | 247 | 36 | 0 | M9- | 2 – 10 | | 15089380 | PF1659 | E1b1b1 | | 2511 |
| 2674 | I | 827 | 0 | 0 | 247 | 36 | 0 | M9- | 2 – 10 | | 15489314 | M5166 | E1b1b1 | | 2674 |
| 2870 | III | 827 | 0 | 0 | 246 | 36 | 1 | M9- | 2 – 10 | | 15869722 | V3194 | E1b1b1 | | 2870 |
| 3193 | III | 827 | 0 | 0 | 246 | 36 | 1 | M9- | 2 – 10 | | 16808859 | PF1701 | E1b1b1 | | 3193 |
| 3345 | III | 827 | 0 | 0 | 247 | 13 | 23 | M9- | 2 – 10 | MD | 17119601 | PF1716 | E1b1b1 | | 3345 |
| 3348 | I | 827 | 0 | 0 | 247 | 36 | 0 | M9- | 2 – 10 | | 17138251 | PF1718 | E1b1b1 | | 3348 |
| 3418 | I | 827 | 0 | 0 | 247 | 36 | 0 | M9- | 2 – 10 | | 17325559 | PF1724.2 | E1b1b1 | A | 3418 |
| 3618 | I | 827 | 0 | 0 | 247 | 36 | 0 | M9- | 2 – 10 | | 17780707 | PF1735 | E1b1b1 | | 3618 |
| 3656 | I | 827 | 0 | 0 | 247 | 36 | 0 | M9- | 2 – 10 | | 17846754 | PF1740 | E1b1b1 | | 3656 |
| 3842 | III | 827 | 0 | 0 | 247 | 35 | 1 | M9- | 2 – 10 | MD | 18384838 | PF1755 | E1b1b1 | | 3842 |
| 3881 | I | 827 | 0 | 0 | 247 | 36 | 0 | M9- | 2 – 10 | | 18573064 | V3289 | E1b1b1 | | 3881 |
| 3998 | I | 827 | 0 | 0 | 247 | 36 | 0 | M9- | 2 – 10 | | 18823372 | PF1768 | E1b1b1 | | 3998 |
| 4164 | I | 827 | 0 | 0 | 247 | 36 | 0 | M9- | 2 – 10 | | 19170454 | V3860 | E1b1b1 | | 4164 |
| 4240 | I | 827 | 0 | 0 | 247 | 36 | 0 | M9- | 2 – 10 | | 19316389 | V4006 | E1b1b1 | | 4240 |
| 4476 | I | 827 | 0 | 0 | 247 | 36 | 0 | M9- | 2 – 10 | | 21358197 | PF1801 | E1b1b1 | | 4476 |
| 4536 | III | 795 | 0 | 32 | 241 | 36 | 6 | M9- | 2 – 10 | | 21491115 | PF1812 | E1b1b1 | | 4536 |
| 4910 | III | 826 | 0 | 1 | 246 | 36 | 1 | M9- | 2 – 10 | | 21977569 | PF1830 | E1b1b1 | | 4910 |
| 4984 | I | 827 | 0 | 0 | 247 | 36 | 0 | M9- | 2 – 10 | | 22168598 | PF1835 | E1b1b1 | | 4984 |
| 4993 | I | 827 | 0 | 0 | 247 | 36 | 0 | M9- | 2 – 10 | | 22181731 | PF1836 | E1b1b1 | | 4993 |
| 5069 | I | 827 | 0 | 0 | 247 | 36 | 0 | M9- | 2 – 10 | | 22588254 | PF1871 | E1b1b1 | | 5069 |

| | | | M9+ | | | M9- | | | | | Build 37 | | ISOGG | | |
|---|---|---|---|---|---|---|---|---|---|---|---|---|---|---|---|
| List | Stage | 0 | 1 | . | 0 | 1 | . | M9? | Clade(s) | Prob. | Position | Name | Subgroup Name | Amb. | List |
| 5533 | I | 827 | 0 | 0 | 247 | 36 | 0 | M9- | 2 – 10 | | 23618826 | PF1909 | E1b1b1 | | 5533 |
| 584 | I | 827 | 0 | 0 | 247 | 36 | 0 | M9- | 2 – 10 | | 6875047 | PF1508 | E1b1b1~ | | 584 |
| 745 | III | 827 | 0 | 0 | 247 | 35 | 1 | M9- | 2 – 10 | MD | 7108366 | PF1515 | E1b1b1~ | | 745 |
| 813 | I | 827 | 0 | 0 | 247 | 36 | 0 | M9- | 2 – 10 | | 7264950 | PF1521 | E1b1b1~ | | 813 |
| 1205 | I | 827 | 0 | 0 | 247 | 36 | 0 | M9- | 2 – 10 | | 8202992 | V1707 | E1b1b1~ | | 1205 |
| 1256 | III | 823 | 0 | 4 | 247 | 36 | 0 | M9- | 2 – 10 | | 8287689 | V1779 | E1b1b1~ | | 1256 |
| 2096 | III | 826 | 0 | 1 | 247 | 36 | 0 | M9- | 2 – 10 | | 14308835 | PF1634 | E1b1b1~ | | 2096 |
| 2222 | III | 827 | 0 | 0 | 246 | 36 | 1 | M9- | 2 – 10 | | 14675847 | PF1647 | E1b1b1~ | | 2222 |
| 2412 | I | 827 | 0 | 0 | 247 | 36 | 0 | M9- | 2 – 10 | | 14972667 | M5149 | E1b1b1~ | | 2412 |
| 2497 | I | 827 | 0 | 0 | 247 | 36 | 0 | M9- | 2 – 10 | | 15051084 | PF1658 | E1b1b1~ | | 2497 |
| 2582 | I | 827 | 0 | 0 | 247 | 36 | 0 | M9- | 2 – 10 | | 15312020 | PF1663 | E1b1b1~ | | 2582 |
| 2667 | I | 827 | 0 | 0 | 247 | 36 | 0 | M9- | 2 – 10 | | 15472586 | V2964 | E1b1b1~ | | 2667 |
| 2774 | III | 825 | 0 | 2 | 246 | 36 | 1 | M9- | 2 – 10 | | 15649242 | V3082 | E1b1b1~ | | 2774 |
| 2808 | III | 827 | 0 | 0 | 247 | 32 | 4 | M9- | 2 – 10 | MD | 15719310 | PF1670 | E1b1b1~ | | 2808 |
| 2829 | I | 827 | 0 | 0 | 247 | 36 | 0 | M9- | 2 – 10 | | 15772820 | PF1672 | E1b1b1~ | | 2829 |
| 3036 | I | 827 | 0 | 0 | 247 | 36 | 0 | M9- | 2 – 10 | | 16405140 | PF1690 | E1b1b1~ | | 3036 |
| 3082 | I | 827 | 0 | 0 | 247 | 36 | 0 | M9- | 2 – 10 | | 16514009 | PF1693 | E1b1b1~ | | 3082 |
| 3120 | I | 827 | 0 | 0 | 247 | 36 | 0 | M9- | 2 – 10 | | 16604397 | M5033 | E1b1b1~ | | 3120 |
| 3353 | I | 827 | 0 | 0 | 247 | 36 | 0 | M9- | 2 – 10 | | 17144649 | PF1721 | E1b1b1~ | | 3353 |
| 3497 | I | 827 | 0 | 0 | 247 | 36 | 0 | M9- | 2 – 10 | | 17514700 | PF1727 | E1b1b1~ | | 3497 |
| 3561 | I | 827 | 0 | 0 | 247 | 36 | 0 | M9- | 2 – 10 | | 17656884 | PF1730 | E1b1b1~ | | 3561 |
| 3624 | I | 827 | 0 | 0 | 247 | 36 | 0 | M9- | 2 – 10 | | 17793914 | PF1736 | E1b1b1~ | | 3624 |
| 3776 | III | 826 | 0 | 1 | 247 | 36 | 0 | M9- | 2 – 10 | | 18081435 | PF1748 | E1b1b1~ | | 3776 |
| 3903 | III | 824 | 0 | 3 | 247 | 36 | 0 | M9- | 2 – 10 | | 18614094 | V3333 | E1b1b1~ | | 3903 |
| 3990 | I | 827 | 0 | 0 | 247 | 36 | 0 | M9- | 2 – 10 | | 18814579 | V3515 | E1b1b1~ | | 3990 |
| 4069 | III | 826 | 0 | 1 | 247 | 35 | 1 | M9- | 2 – 10 | MD | 18963497 | PF1771 | E1b1b1~ | | 4069 |
| 4284 | III | 826 | 0 | 1 | 247 | 36 | 0 | M9- | 2 – 10 | | 19443039 | V4126 | E1b1b1~ | | 4284 |
| 4333 | III | 825 | 0 | 2 | 247 | 36 | 0 | M9- | 2 – 10 | | 20812311 | PF1791 | E1b1b1~ | | 4333 |
| 4463 | I | 827 | 0 | 0 | 247 | 36 | 0 | M9- | 2 – 10 | | 21315746 | S22282 | E1b1b1~ | | 4463 |
| 4494 | III | 810 | 0 | 17 | 246 | 35 | 2 | M9- | 2 – 10 | MD, 1H | 21389667 | PF1804 | E1b1b1~ | | 4494 |
| 5205 | I | 827 | 0 | 0 | 247 | 36 | 0 | M9- | 2 – 10 | | 22835237 | PF1877 | E1b1b1~ | | 5205 |
| 5283 | III | 803 | 0 | 24 | 242 | 36 | 5 | M9- | 2 – 10 | | 23009991 | PF1881 | E1b1b1~ | | 5283 |
| 5301 | III | 801 | 0 | 26 | 240 | 36 | 7 | M9- | 2 – 10 | | 23045843 | PF1887 | E1b1b1~ | | 5301 |
| 5530 | I | 827 | 0 | 0 | 247 | 36 | 0 | M9- | 2 – 10 | | 23614600 | PF1908 | E1b1b1~ | | 5530 |
| 5538 | I | 827 | 0 | 0 | 247 | 36 | 0 | M9- | 2 – 10 | | 23629008 | PF1910 | E1b1b1~ | | 5538 |
| 5563 | I | 827 | 0 | 0 | 247 | 36 | 0 | M9- | 2 – 10 | | 23781504 | PF1912 | E1b1b1~ | | 5563 |
| 5686 | III | 826 | 0 | 1 | 247 | 36 | 0 | M9- | 2 – 10 | | 24492875 | PF1924 | E1b1b1~ | | 5686 |
| 1039 | III | 827 | 0 | 0 | 253 | 26 | 4 | M9- | 2 – 7 | MD | 7804308 | PF2108 | E1b1b1a | | 1039 |
| 1220 | I | 827 | 0 | 0 | 253 | 30 | 0 | M9- | 2 – 7 | | 8232450 | V1739 | E1b1b1a | | 1220 |
| 1286 | I | 827 | 0 | 0 | 253 | 30 | 0 | M9- | 2 – 7 | | 8361073 | PF2115 | E1b1b1a | | 1286 |
| 2070 | III | 819 | 0 | 8 | 251 | 30 | 2 | M9- | 2 – 7 | | 14242950 | PF2127 | E1b1b1a | | 2070 |
| 2669 | III | 826 | 0 | 1 | 253 | 30 | 0 | M9- | 2 – 7 | | 15479586 | Z1209 | E1b1b1a | | 2669 |
| 2695 | I | 827 | 0 | 0 | 253 | 30 | 0 | M9- | 2 – 7 | | 15520497 | V3011 | E1b1b1a | | 2695 |
| 3596 | III | 795 | 0 | 32 | 245 | 30 | 8 | M9- | 2 – 7 | | 17744738 | PF2154 | E1b1b1a | | 3596 |
| 3804 | I | 827 | 0 | 0 | 253 | 30 | 0 | M9- | 2 – 7 | | 18149476 | PF2157 | E1b1b1a | | 3804 |
| 3864 | III | 821 | 0 | 6 | 252 | 30 | 1 | M9- | 2 – 7 | | 18538216 | CTS8899 | E1b1b1a | | 3864 |
| 3963 | III | 826 | 0 | 1 | 253 | 30 | 0 | M9- | 2 – 7 | | 18765146 | PF2162 | E1b1b1a | | 3963 |
| 4044 | III | 812 | 0 | 15 | 248 | 30 | 5 | M9- | 2 – 7 | | 18918010 | V3603 | E1b1b1a | | 4044 |
| 4058 | I | 827 | 0 | 0 | 253 | 30 | 0 | M9- | 2 – 7 | | 18949452 | PF2280 | E1b1b1a | | 4058 |
| 4273 | I | 827 | 0 | 0 | 253 | 30 | 0 | M9- | 2 – 7 | | 19396726 | V4083 | E1b1b1a | | 4273 |
| 4351 | I | 827 | 0 | 0 | 253 | 30 | 0 | M9- | 2 – 7 | | 21036413 | Z1211 | E1b1b1a | | 4351 |
| 4489 | I | 827 | 0 | 0 | 253 | 30 | 0 | M9- | 2 – 7 | | 21369924 | S7478 | E1b1b1a | | 4489 |
| 4741 | I | 827 | 0 | 0 | 253 | 30 | 0 | M9- | 2 – 7 | | 21792508 | PF2182 | E1b1b1a | | 4741 |
| 4767 | I | 827 | 0 | 0 | 253 | 30 | 0 | M9- | 2 – 7 | | 21860060 | Z1214 | E1b1b1a | | 4767 |
| 5225 | I | 827 | 0 | 0 | 253 | 30 | 0 | M9- | 2 – 7 | | 22898839 | PF1955 | E1b1b1a | | 5225 |
| 5361 | I | 827 | 0 | 0 | 253 | 30 | 0 | M9- | 2 – 7 | | 23161406 | PF2200 | E1b1b1a | | 5361 |
| 5622 | III | 826 | 0 | 1 | 253 | 30 | 0 | M9- | 2 – 7 | | 23973415 | PF2203 | E1b1b1a | | 5622 |
| 1394 | I | 827 | 0 | 0 | 253 | 30 | 0 | M9- | 2 – 7 | | 8483852 | V2104 | E1b1b1a1 | | 1394 |
| 4830 | I | 827 | 0 | 0 | 253 | 30 | 0 | M9- | 2 – 7 | | 21893303 | PF2186 | E1b1b1a1 | | 4830 |
| 5200 | III | 824 | 0 | 3 | 281 | 2 | 0 | M9- | 2 | | 22823155 | Z1902 | E1b1b1a1a | | 5200 |
| 2213 | II | 827 | 0 | 0 | 281 | 2 | 0 | M9- | 2 | | 14645286 | Z1219 | E1b1b1a1a1 | | 2213 |
| 2490 | II | 827 | 0 | 0 | 281 | 2 | 0 | M9- | 2 | | 15040454 | Z1220 | E1b1b1a1a1 | | 2490 |
| 3973 | II | 827 | 0 | 0 | 281 | 2 | 0 | M9- | 2 | | 18776848 | Z1225 | E1b1b1a1a1 | | 3973 |
| 4829 | IV | 827 | 0 | 0 | 273 | 0 | 10 | Mono | Mono | | 21893299 | M224 | E1b1b1a1a1a~ | | 4829 |
| 646 | IV | 827 | 0 | 0 | 283 | 0 | 0 | Mono | Mono | | 6932821 | V32 | E1b1b1a1a1b | | 646 |
| 1462 | IV | 827 | 0 | 0 | 283 | 0 | 0 | Mono | Mono | | 8576554 | V2296 | E1b1b1a1a2 | | 1462 |
| 4132 | IV | 826 | 0 | 1 | 283 | 0 | 0 | Mono | Mono | | 19118052 | Z1240 | E1b1b1a1a2 | | 4132 |
| 4285 | IV | 827 | 0 | 0 | 283 | 0 | 0 | Mono | Mono | | 19443230 | V4127 | E1b1b1a1a2 | | 4285 |
| 4865 | IV | 827 | 0 | 0 | 283 | 0 | 0 | Mono | Mono | | 21906413 | PF2287 | E1b1b1a1a2a | A | 4865 |
| 1175 | IV | 827 | 0 | 0 | 283 | 0 | 0 | Mono | Mono | | 8124473 | M5624 | E1b1b1a1a2a1b~ | A | 1175 |
| 2680 | I | 827 | 0 | 0 | 255 | 28 | 0 | M9- | 3 – 7 | | 15502485 | Z1920 | E1b1b1a1b | | 2680 |
| 1242 | I | 827 | 0 | 0 | 260 | 23 | 0 | M9- | 3 – 5 | | 8262442 | V1764 | E1b1b1a1b1 | | 1242 |
| 542 | I | 827 | 0 | 0 | 261 | 22 | 0 | M9- | 4 – 5 | | 6814246 | V36 | E1b1b1a1b1a | | 542 |
| 550 | I | 827 | 0 | 0 | 261 | 22 | 0 | M9- | 4 – 5 | | 6842263 | V13 | E1b1b1a1b1a | | 550 |
| 5148 | III | 827 | 0 | 0 | 261 | 21 | 1 | M9- | 4 – 5 | MD | 22744616 | Page102 | E1b1b1a1b1a | | 5148 |
| 4060 | I | 827 | 0 | 0 | 278 | 5 | 0 | M9- | 6 – 7 | | 18957059 | L677 | E1b1b1a1b2 | | 4060 |
| 4838 | IV | 827 | 0 | 0 | 283 | 0 | 0 | Mono | Mono | | 21896093 | M148 | E1b1b1a1b2a1~ | | 4838 |
| 1549 | IV | 827 | 0 | 0 | 283 | 0 | 0 | Mono | Mono | | 8702003 | F1386 | E1b1b1a1b2a4c1~ | | 1549 |
| 544 | IV | 827 | 0 | 0 | 283 | 0 | 0 | Mono | Mono | | 6822948 | M521 | E1b1b1a1c~ | | 544 |
| 44 | II | 827 | 0 | 0 | 281 | 2 | 0 | M9- | 8 | | 2710013 | PF2300 | E1b1b1b1 | | 44 |
| 832 | II | 827 | 0 | 0 | 281 | 2 | 0 | M9- | 8 | | 7322178 | PF2314 | E1b1b1b1 | | 832 |
| 858 | II | 827 | 0 | 0 | 281 | 2 | 0 | M9- | 8 | | 7432723 | PF2318 | E1b1b1b1 | | 858 |
| 968 | II | 827 | 0 | 0 | 281 | 2 | 0 | M9- | 8 | | 7682321 | Z1162 | E1b1b1b1 | | 968 |
| 1341 | II | 827 | 0 | 0 | 281 | 2 | 0 | M9- | 8 | | 8438286 | Z1166 | E1b1b1b1 | A | 1341 |
| 2182 | II | 827 | 0 | 0 | 281 | 2 | 0 | M9- | 8 | | 14541624 | PF2346 | E1b1b1b1 | | 2182 |
| 2280 | III | 826 | 0 | 1 | 281 | 2 | 0 | M9- | 8 | | 14817699 | PF2347 | E1b1b1b1 | | 2280 |
| 3012 | III | 827 | 0 | 0 | 281 | 1 | 1 | M9- | 8 | MD | 16361269 | Z1183 | E1b1b1b1 | | 3012 |
| 3026 | III | 674 | 0 | 153 | 228 | 2 | 53 | M9- | 8 | | 16390272 | PF2362 | E1b1b1b1 | | 3026 |
| 3256 | II | 827 | 0 | 0 | 281 | 2 | 0 | M9- | 8 | | 16899460 | PF2365 | E1b1b1b1 | | 3256 |
| 3331 | II | 827 | 0 | 0 | 281 | 2 | 0 | M9- | 8 | | 17091865 | Z1187 | E1b1b1b1 | | 3331 |
| 3363 | II | 826 | 1 | 0 | 281 | 2 | 0 | BOTH | | PI | 17190078 | PF2372 | E1b1b1b1 | | 3363 |
| 3377 | II | 827 | 0 | 0 | 281 | 2 | 0 | M9- | 8 | | 17210606 | M5222 | E1b1b1b1 | | 3377 |
| 3568 | II | 827 | 0 | 0 | 281 | 2 | 0 | M9- | 8 | | 17670232 | PF2378 | E1b1b1b1 | | 3568 |
| 3569 | II | 827 | 0 | 0 | 281 | 2 | 0 | M9- | 8 | | 17670258 | Z1193 | E1b1b1b1 | | 3569 |
| 3797 | II | 827 | 0 | 0 | 281 | 2 | 0 | M9- | 8 | | 18128763 | PF2381 | E1b1b1b1 | | 3797 |
| 3858 | II | 827 | 0 | 0 | 281 | 2 | 0 | M9- | 8 | | 18416012 | Z1196 | E1b1b1b1 | | 3858 |

| | | M9+ | | | M9- | | | | | | Build 37 | | ISOGG | | |
|---|---|---|---|---|---|---|---|---|---|---|---|---|---|---|---|
| List | Stage | 0 | 1 | . | 0 | 1 | . | M9? | Clade(s) | Prob. | Position | Name | Subgroup Name | Amb. | List |
| 3861 | II | 827 | 0 | 0 | 281 | 2 | 0 | M9- | 8 | | 18419547 | Z1197 | E1b1b1b1 | | 3861 |
| 4174 | II | 827 | 0 | 0 | 281 | 2 | 0 | M9- | 8 | | 19193159 | V3883 | E1b1b1b1 | | 4174 |
| 4497 | II | 827 | 0 | 0 | 281 | 2 | 0 | M9- | 8 | | 21397600 | PF2394 | E1b1b1b1 | | 4497 |
| 4545 | III | 827 | 0 | 0 | 280 | 2 | 1 | M9- | 8 | | 21508604 | PF2395 | E1b1b1b1 | | 4545 |
| 4746 | II | 827 | 0 | 0 | 281 | 2 | 0 | M9- | 8 | | 21807861 | PF2396 | E1b1b1b1 | | 4746 |
| 4916 | III | 797 | 0 | 30 | 275 | 2 | 6 | M9- | 8 | | 21989906 | PF2398 | E1b1b1b1 | | 4916 |
| 5167 | II | 827 | 0 | 0 | 281 | 2 | 0 | M9- | 8 | | 22751863 | PF2402.2 | E1b1b1b1 | A | 5167 |
| 5372 | II | 827 | 0 | 0 | 281 | 2 | 0 | M9- | 8 | | 23225611 | PF2405 | E1b1b1b1 | | 5372 |
| 5576 | II | 827 | 0 | 0 | 281 | 2 | 0 | M9- | 8 | | 23809883 | PF2410 | E1b1b1b1 | | 5576 |
| 5625 | II | 827 | 0 | 0 | 281 | 2 | 0 | M9- | 8 | | 23976740 | PF2413 | E1b1b1b1 | | 5625 |
| 825 | II | 827 | 0 | 0 | 281 | 2 | 0 | M9- | 8 | | 7303900 | M5064 | E1b1b1b1~ | | 825 |
| 1183 | III | 821 | 0 | 6 | 279 | 2 | 2 | M9- | 8 | | 8134356 | V1649 | E1b1b1b1~ | | 1183 |
| 1606 | II | 827 | 0 | 0 | 281 | 2 | 0 | M9- | 8 | | 8871735 | M5107 | E1b1b1b1~ | | 1606 |
| 2098 | II | 827 | 0 | 0 | 281 | 2 | 0 | M9- | 8 | | 14320818 | PF2344 | E1b1b1b1~ | | 2098 |
| 2509 | II | 827 | 0 | 0 | 281 | 2 | 0 | M9- | 8 | | 15085950 | V2814 | E1b1b1b1~ | | 2509 |
| 2575 | II | 827 | 0 | 0 | 281 | 2 | 0 | M9- | 8 | | 15273119 | Z1178 | E1b1b1b1~ | | 2575 |
| 2650 | III | 825 | 0 | 2 | 281 | 2 | 0 | M9- | 8 | | 15452759 | PF2353 | E1b1b1b1~ | | 2650 |
| 2689 | II | 827 | 0 | 0 | 281 | 2 | 0 | M9- | 8 | | 15512696 | V3001 | E1b1b1b1~ | | 2689 |
| 2874 | II | 827 | 0 | 0 | 281 | 2 | 0 | M9- | 8 | | 15879807 | PF2356 | E1b1b1b1~ | | 2874 |
| 2956 | III | 826 | 0 | 1 | 281 | 2 | 0 | M9- | 8 | | 16223362 | PF2359 | E1b1b1b1~ | | 2956 |
| 3342 | II | 827 | 0 | 0 | 281 | 2 | 0 | M9- | 8 | | 17118046 | PF2370 | E1b1b1b1~ | | 3342 |
| 3381 | II | 827 | 0 | 0 | 281 | 2 | 0 | M9- | 8 | | 17235818 | PF2374 | E1b1b1b1~ | | 3381 |
| 3480 | III | 827 | 0 | 0 | 280 | 2 | 1 | M9- | 8 | 1H | 17486942 | PF2377 | E1b1b1b1~ | | 3480 |
| 4292 | II | 827 | 0 | 0 | 281 | 2 | 0 | M9- | 8 | | 19475462 | Z1198 | E1b1b1b1~ | | 4292 |
| 4359 | II | 827 | 0 | 0 | 281 | 2 | 0 | M9- | 8 | | 21066342 | M5285 | E1b1b1b1~ | | 4359 |
| 5421 | II | 827 | 0 | 0 | 281 | 2 | 0 | M9- | 8 | | 23314799 | Z1202 | E1b1b1b1~ | | 5421 |
| 5435 | II | 827 | 0 | 0 | 281 | 2 | 0 | M9- | 8 | | 23367445 | PF2407 | E1b1b1b1~ | | 5435 |
| 5442 | II | 827 | 0 | 0 | 281 | 2 | 0 | M9- | 8 | | 23385983 | PF2408 | E1b1b1b1~ | | 5442 |
| 5659 | III | 825 | 0 | 2 | 281 | 2 | 0 | M9- | 8 | | 24396804 | PF2414 | E1b1b1b1~ | | 5659 |
| 4825 | IV | 827 | 0 | 0 | 283 | 0 | 0 | Mono | Mono | | 21892572 | PF2554 | E1b1b1b1a | | 4825 |
| 62 | IV | 827 | 0 | 0 | 283 | 0 | 0 | Mono | Mono | | 2723620 | V1030 | E1b1b1b1a1 | | 62 |
| 830 | IV | 809 | 0 | 18 | 278 | 0 | 5 | Mono | Mono | | 7317593 | Z1161 | E1b1b1b1a1 | | 830 |
| 928 | IV | 827 | 0 | 0 | 283 | 0 | 0 | Mono | Mono | | 7612987 | V1272 | E1b1b1b1a1 | | 928 |
| 1014 | IV | 827 | 0 | 0 | 283 | 0 | 0 | Mono | Mono | | 7752540 | PF2486 | E1b1b1b1a1 | | 1014 |
| 1316 | IV | 827 | 0 | 0 | 283 | 0 | 0 | Mono | Mono | | 8415334 | Z1165 | E1b1b1b1a1 | | 1316 |
| 1324 | IV | 827 | 0 | 0 | 283 | 0 | 0 | Mono | Mono | | 8423288 | V1937 | E1b1b1b1a1 | | 1324 |
| 1611 | IV | 826 | 0 | 1 | 283 | 0 | 0 | Mono | Mono | | 8881572 | PF2498 | E1b1b1b1a1 | | 1611 |
| 1620 | IV | 826 | 0 | 1 | 282 | 0 | 1 | Mono | Mono | | 8900662 | PF2500 | E1b1b1b1a1 | | 1620 |
| 1662 | IV | 826 | 0 | 1 | 283 | 0 | 0 | Mono | Mono | | 9135484 | PF2501 | E1b1b1b1a1 | | 1662 |
| 1996 | IV | 827 | 0 | 0 | 283 | 0 | 0 | Mono | Mono | | 14082877 | Z1172 | E1b1b1b1a1 | | 1996 |
| 2307 | IV | 827 | 0 | 0 | 283 | 0 | 0 | Mono | Mono | | 14858228 | PF2510 | E1b1b1b1a1 | | 2307 |
| 2329 | IV | 827 | 0 | 0 | 283 | 0 | 0 | Mono | Mono | | 14888783 | Page33 | E1b1b1b1a1 | | 2329 |
| 2355 | IV | 827 | 0 | 0 | 283 | 0 | 0 | Mono | Mono | | 14906675 | PF2348 | E1b1b1b1a1 | | 2355 |
| 2557 | IV | 827 | 0 | 0 | 283 | 0 | 0 | Mono | Mono | | 15236994 | Z1177 | E1b1b1b1a1 | | 2557 |
| 2681 | IV | 827 | 0 | 0 | 283 | 0 | 0 | Mono | Mono | | 15502732 | Z1179 | E1b1b1b1a1 | | 2681 |
| 3073 | IV | 827 | 0 | 0 | 283 | 0 | 0 | Mono | Mono | | 16491953 | Z1184 | E1b1b1b1a1 | | 3073 |
| 3088 | IV | 827 | 0 | 0 | 283 | 0 | 0 | Mono | Mono | | 16534130 | PF2363 | E1b1b1b1a1 | | 3088 |
| 3765 | IV | 827 | 0 | 0 | 282 | 0 | 1 | Mono | Mono | | 18068338 | Z1195 | E1b1b1b1a1 | | 3765 |
| 4244 | IV | 827 | 0 | 0 | 283 | 0 | 0 | Mono | Mono | | 19323263 | V4016 | E1b1b1b1a1 | | 4244 |
| 4272 | IV | 827 | 0 | 0 | 283 | 0 | 0 | Mono | Mono | | 19396045 | V4081 | E1b1b1b1a1 | | 4272 |
| 4297 | IV | 827 | 0 | 0 | 283 | 0 | 0 | Mono | Mono | | 19482455 | V4166 | E1b1b1b1a1 | | 4297 |
| 4358 | IV | 827 | 0 | 0 | 283 | 0 | 0 | Mono | Mono | | 21064067 | PF2546 | E1b1b1b1a1 | | 4358 |
| 4468 | IV | 827 | 0 | 0 | 283 | 0 | 0 | Mono | Mono | | 21327286 | PF2548 | E1b1b1b1a1 | | 4468 |
| 4739 | IV | 824 | 0 | 3 | 282 | 0 | 1 | Mono | Mono | | 21785697 | PF2551 | E1b1b1b1a1 | | 4739 |
| 4751 | IV | 827 | 0 | 0 | 283 | 0 | 0 | Mono | Mono | | 21815268 | PF2553 | E1b1b1b1a1 | | 4751 |
| 5056 | IV | 827 | 0 | 0 | 283 | 0 | 0 | Mono | Mono | | 22566659 | Z1201 | E1b1b1b1a1 | | 5056 |
| 5275 | IV | 827 | 0 | 0 | 283 | 0 | 0 | Mono | Mono | | 22978109 | PF2560 | E1b1b1b1a1 | | 5275 |
| 641 | IV | 827 | 0 | 0 | 282 | 0 | 1 | Mono | Mono | | 6932547 | PF2482 | E1b1b1b1a1~ | | 641 |
| 1049 | IV | 827 | 0 | 0 | 283 | 0 | 0 | Mono | Mono | | 7830010 | Z1163 | E1b1b1b1a1~ | | 1049 |
| 1423 | IV | 827 | 0 | 0 | 283 | 0 | 0 | Mono | Mono | | 8518796 | Z1167 | E1b1b1b1a1~ | | 1423 |
| 1435 | IV | 827 | 0 | 0 | 283 | 0 | 0 | Mono | Mono | | 8540752 | V2208 | E1b1b1b1a1~ | A | 1435 |
| 1559 | IV | 827 | 0 | 0 | 283 | 0 | 0 | Mono | Mono | | 8726283 | V2505 | E1b1b1b1a1~ | | 1559 |
| 2038 | IV | 819 | 0 | 8 | 282 | 0 | 1 | Mono | Mono | | 14186688 | Z1175 | E1b1b1b1a1~ | | 2038 |
| 2120 | IV | 827 | 0 | 0 | 283 | 0 | 0 | Mono | Mono | | 14365288 | Z1176 | E1b1b1b1a1~ | | 2120 |
| 2526 | IV | 826 | 0 | 1 | 283 | 0 | 0 | Mono | Mono | | 15114328 | PF2512 | E1b1b1b1a1~ | | 2526 |
| 2812 | IV | 825 | 0 | 2 | 279 | 0 | 4 | Mono | Mono | | 15725315 | PF2568 | E1b1b1b1a1~ | | 2812 |
| 2900 | IV | 827 | 0 | 0 | 283 | 0 | 0 | Mono | Mono | | 15978952 | PF2516 | E1b1b1b1a1~ | | 2900 |
| 3016 | IV | 827 | 0 | 0 | 283 | 0 | 0 | Mono | Mono | | 16367635 | PF2518 | E1b1b1b1a1~ | | 3016 |
| 3111 | IV | 823 | 0 | 4 | 283 | 0 | 0 | Mono | Mono | | 16586893 | Z1185 | E1b1b1b1a1~ | | 3111 |
| 3350 | IV | 827 | 0 | 0 | 283 | 0 | 0 | Mono | Mono | | 17139711 | Z1190 | E1b1b1b1a1~ | | 3350 |
| 4235 | IV | 824 | 0 | 3 | 282 | 0 | 1 | Mono | Mono | | 19304315 | PF2543 | E1b1b1b1a1~ | | 4235 |
| 4515 | IV | 827 | 0 | 0 | 283 | 0 | 0 | Mono | Mono | | 21437042 | M5300 | E1b1b1b1a1~ | | 4515 |
| 4738 | IV | 826 | 0 | 1 | 282 | 0 | 1 | Mono | Mono | | 21785013 | PF2550 | E1b1b1b1a1~ | | 4738 |
| 4944 | IV | 827 | 0 | 0 | 283 | 0 | 0 | Mono | Mono | | 22076595 | Z1200 | E1b1b1b1a1~ | | 4944 |
| 5320 | IV | 827 | 0 | 0 | 283 | 0 | 0 | Mono | Mono | | 23074213 | PF2562 | E1b1b1b1a1~ | | 5320 |
| 5341 | IV | 822 | 0 | 5 | 280 | 0 | 3 | Mono | Mono | | 23124284 | PF2564 | E1b1b1b1a1~ | | 5341 |
| 5524 | IV | 827 | 0 | 0 | 282 | 0 | 1 | Mono | Mono | | 23583756 | M5358 | E1b1b1b1a1~ | | 5524 |
| 5696 | IV | 825 | 0 | 2 | 282 | 0 | 1 | Mono | Mono | | 24513116 | Z1203 | E1b1b1b1a1~ | | 5696 |
| 1738 | I | 826 | 1 | 0 | 270 | 13 | 0 | BOTH | | PI | 9823657 | PF3985 | E1b1b1b1b | A | 1738 |
| 674 | I | 827 | 0 | 0 | 279 | 4 | 0 | M9- | 9 – 10 | | 6953603 | Z829 | E1b1b1b2 | | 674 |
| 997 | I | 827 | 0 | 0 | 279 | 4 | 0 | M9- | 9 – 10 | | 7729622 | Z830 | E1b1b1b2 | | 997 |
| 1729 | III | 814 | 0 | 13 | 274 | 4 | 5 | M9- | 9 – 10 | | 9795609 | Z1076 | E1b1b1b2 | | 1729 |
| 2381 | III | 822 | 0 | 5 | 278 | 4 | 1 | M9- | 9 – 10 | | 14941989 | PF1973 | E1b1b1b2 | | 2381 |
| 2989 | I | 827 | 0 | 0 | 279 | 4 | 0 | M9- | 9 – 10 | | 16313785 | Z832 | E1b1b1b2 | | 2989 |
| 2990 | I | 827 | 0 | 0 | 279 | 4 | 0 | M9- | 9 – 10 | | 16314844 | Z833 | E1b1b1b2 | | 2990 |
| 3288 | I | 827 | 0 | 0 | 279 | 4 | 0 | M9- | 9 – 10 | | 16948834 | Z834 | E1b1b1b2 | | 3288 |
| 3329 | III | 827 | 0 | 0 | 279 | 3 | 1 | M9- | 9 – 10 | MD | 17077753 | Z835 | E1b1b1b2 | | 3329 |
| 3664 | I | 827 | 0 | 0 | 279 | 4 | 0 | M9- | 9 – 10 | | 17865060 | CTS8182 | E1b1b1b2 | | 3664 |
| 3822 | III | 807 | 0 | 20 | 273 | 4 | 6 | M9- | 9 – 10 | | 18203565 | PF1989 | E1b1b1b2 | | 3822 |
| 4264 | I | 827 | 0 | 0 | 279 | 4 | 0 | M9- | 9 – 10 | | 19379842 | V4065 | E1b1b1b2 | | 4264 |
| 4742 | III | 827 | 0 | 0 | 278 | 4 | 1 | M9- | 9 – 10 | | 21797330 | PF1997 | E1b1b1b2 | | 4742 |
| 5402 | I | 827 | 0 | 0 | 279 | 4 | 0 | M9- | 9 – 10 | | 23268660 | PF2006 | E1b1b1b2 | | 5402 |
| 4072 | I | 827 | 0 | 0 | 279 | 4 | 0 | M9- | 9 – 10 | | 18976881 | V3649 | E1b1b1b2a1 | | 4072 |
| 4716 | I | 827 | 0 | 0 | 279 | 4 | 0 | M9- | 9 – 10 | | 21764586 | PF2023 | E1b1b1b2a1 | | 4716 |
| 1227 | III | 827 | 0 | 0 | 278 | 4 | 1 | M9- | 9 – 10 | | 8240479 | V1750 | E1b1b1b2a1a~ | | 1227 |
| 1551 | III | 826 | 0 | 1 | 279 | 4 | 0 | M9- | 9 – 10 | | 8704571 | Z1148 | E1b1b1b2a1a~ | | 1551 |

| List | Stage | M9+ | | | M9- | | | M9? | Clade(s) | Prob. | Build 37 | | ISOGG | Amb. | List |
|---|---|---|---|---|---|---|---|---|---|---|---|---|---|---|---|
| | | 0 | 1 | . | 0 | 1 | . | | | | Position | Name | Subgroup Name | | |
| 4683 | I | 827 | 0 | 0 | 279 | 4 | 0 | M9- | 9 – 10 | | 21740716 | PF2022 | E1b1b1b2a1a~ | | 4683 |
| 3275 | II | 827 | 0 | 0 | 280 | 3 | 0 | M9- | 9 | | 16936081 | Page47 | E1b1b1b2a1a1 | | 3275 |
| 1228 | III | 825 | 0 | 2 | 280 | 3 | 0 | M9- | 9 | | 8240582 | L788 | E1b1b1b2a1a1~ | | 1228 |
| 1684 | III | 824 | 1 | 2 | 282 | 0 | 1 | M9+ | NU | 1H | 9398519 | F1539 | E1b1b1b2a1a1a1a1b1~ | A,W | 1684 |
| 3954 | IV | 827 | 0 | 0 | 283 | 0 | 0 | Mono | Mono | | 18743050 | V3465 | E1b1b1b2a1a1a1a1f1b1a | A | 3954 |
| 4811 | IV | 827 | 0 | 0 | 283 | 0 | 0 | Mono | Mono | | 21878825 | M290 | E1b1b1b2a1a2 | | 4811 |
| 3711 | III | 826 | 0 | 1 | 282 | 1 | 0 | M9- | 10 | | 17947414 | L792 | E1b1b1b2a1a4 | | 3711 |
| 5193 | IV | 827 | 0 | 0 | 283 | 0 | 0 | Mono | Mono | | 22817188 | CTS10880 | E1b1b1b2b | | 5193 |
| 5227 | IV | 827 | 0 | 0 | 283 | 0 | 0 | Mono | Mono | | 22905131 | CTS11025 | E1b1b1b2b | | 5227 |
| 5241 | IV | 827 | 0 | 0 | 283 | 0 | 0 | Mono | Mono | | 22920950 | CTS11051 | E1b1b1b2b | | 5241 |
| 5363 | IV | 826 | 0 | 1 | 283 | 0 | 0 | Mono | Mono | | 23164107 | Z1283 | E1b1b1b2b | | 5363 |
| 4583 | IV | 827 | 0 | 0 | 283 | 0 | 0 | Mono | Mono | | 21610853 | P72 | E1b1b1b2b1~ | | 4583 |
| 810 | IV | 827 | 0 | 0 | 283 | 0 | 0 | Mono | Mono | | 7253049 | CTS1177 | E1b1b1b2b2 | | 810 |
| 4197 | IV | 827 | 0 | 0 | 283 | 0 | 0 | Mono | Mono | | 19233913 | V3935 | E1b1b1b2b2 | | 4197 |
| 5208 | IV | 827 | 0 | 0 | 283 | 0 | 0 | Mono | Mono | | 22845992 | CTS10942 | E1b1b1b2b2 | | 5208 |
| 5150 | IV | 827 | 0 | 0 | 283 | 0 | 0 | Mono | Mono | | 22744939 | M293 | E1b1b1b2b2a1 | | 5150 |
| 2077 | IV | 827 | 0 | 0 | 283 | 0 | 0 | Mono | Mono | | 14254220 | CTS2297 | E1b1b1b2b2a1a~ | | 2077 |
| 630 | IV | 827 | 0 | 0 | 283 | 0 | 0 | Mono | Mono | | 6932007 | V6 | E1b1b1c | A | 630 |
| 4714 | IV | 827 | 0 | 0 | 283 | 0 | 0 | Mono | Mono | | 21764500 | M281 | E1b1b2 | | 4714 |
| 30 | IV | 827 | 0 | 0 | 283 | 0 | 0 | Mono | Mono | | 2671852 | CTS20 | E2 | | 30 |
| 41 | IV | 827 | 0 | 0 | 283 | 0 | 0 | Mono | Mono | | 2708952 | V1006 | E2 | | 41 |
| 145 | IV | 827 | 0 | 0 | 283 | 0 | 0 | Mono | Mono | | 2842030 | V1124 | E2 | | 145 |
| 181 | IV | 827 | 0 | 0 | 283 | 0 | 0 | Mono | Mono | | 2888397 | V1152 | E2 | | 181 |
| 946 | IV | 823 | 0 | 4 | 283 | 0 | 0 | Mono | Mono | | 7648175 | Z15718 | E2 | | 946 |
| 1015 | IV | 823 | 0 | 4 | 282 | 0 | 1 | Mono | Mono | | 7753973 | Z999 | E2 | | 1015 |
| 1255 | IV | 827 | 0 | 0 | 283 | 0 | 0 | Mono | Mono | | 8285892 | Z15725 | E2 | | 1255 |
| 1401 | IV | 825 | 0 | 2 | 283 | 0 | 0 | Mono | Mono | | 8489122 | Z15727 | E2 | | 1401 |
| 2320 | IV | 827 | 0 | 0 | 283 | 0 | 0 | Mono | Mono | | 14876810 | Z1009 | E2 | | 2320 |
| 2389 | IV | 827 | 0 | 0 | 283 | 0 | 0 | Mono | Mono | | 14946181 | Z1010 | E2 | | 2389 |
| 2717 | IV | 827 | 0 | 0 | 283 | 0 | 0 | Mono | Mono | | 15555421 | V3029 | E2 | | 2717 |
| 2866 | IV | 826 | 0 | 1 | 283 | 0 | 0 | Mono | Mono | | 15862752 | V3185 | E2 | | 2866 |
| 4022 | IV | 826 | 0 | 1 | 283 | 0 | 0 | Mono | Mono | | 18885293 | V3573 | E2 | | 4022 |
| 4064 | IV | 827 | 0 | 0 | 283 | 0 | 0 | Mono | Mono | | 18960670 | V3638 | E2 | | 4064 |
| 4280 | IV | 827 | 0 | 0 | 283 | 0 | 0 | Mono | Mono | | 19420881 | Z1024 | E2 | | 4280 |
| 4824 | IV | 827 | 0 | 0 | 283 | 0 | 0 | Mono | Mono | | 21890177 | M75 | E2 | | 4824 |
| 5308 | IV | 827 | 0 | 0 | 283 | 0 | 0 | Mono | Mono | | 23063397 | M10999 | E2 | | 5308 |
| 2367 | IV | 826 | 0 | 1 | 283 | 0 | 0 | Mono | Mono | | 14923438 | M237 | E2a~ | | 2367 |
| 68 | IV | 827 | 0 | 0 | 283 | 0 | 0 | Mono | Mono | | 2727675 | V1033 | E2b | | 68 |
| 89 | IV | 827 | 0 | 0 | 283 | 0 | 0 | Mono | Mono | | 2751973 | V1057 | E2b | | 89 |
| 474 | IV | 827 | 0 | 0 | 283 | 0 | 0 | Mono | Mono | | 6667139 | Z15793 | E2b | | 474 |
| 497 | IV | 827 | 0 | 0 | 283 | 0 | 0 | Mono | Mono | | 6716831 | CTS375 | E2b | | 497 |
| 939 | IV | 827 | 0 | 0 | 283 | 0 | 0 | Mono | Mono | | 7638542 | Z15797 | E2b | | 939 |
| 1160 | IV | 827 | 0 | 0 | 283 | 0 | 0 | Mono | Mono | | 8083622 | Z15801 | E2b | | 1160 |
| 1304 | IV | 819 | 0 | 8 | 281 | 0 | 2 | Mono | Mono | | 8390514 | Z15804 | E2b | | 1304 |
| 1476 | IV | 827 | 0 | 0 | 283 | 0 | 0 | Mono | Mono | | 8592165 | Z1002 | E2b | | 1476 |
| 2217 | IV | 827 | 0 | 0 | 283 | 0 | 0 | Mono | Mono | | 14659581 | V2543 | E2b | | 2217 |
| 2289 | IV | 827 | 0 | 0 | 283 | 0 | 0 | Mono | Mono | | 14837777 | V2630 | E2b | | 2289 |
| 2876 | IV | 827 | 0 | 0 | 283 | 0 | 0 | Mono | Mono | | 15899708 | Z1013 | E2b | | 2876 |
| 3907 | IV | 827 | 0 | 0 | 283 | 0 | 0 | Mono | Mono | | 18617044 | Z1021 | E2b | | 3907 |
| 3988 | IV | 827 | 0 | 0 | 283 | 0 | 0 | Mono | Mono | | 18805281 | CTS9295 | E2b | | 3988 |
| 4005 | IV | 826 | 0 | 1 | 282 | 0 | 1 | Mono | Mono | | 18840959 | V3551 | E2b | | 4005 |
| 4032 | IV | 827 | 0 | 0 | 282 | 0 | 1 | Mono | Mono | | 18899203 | V3584 | E2b | | 4032 |
| 4033 | IV | 827 | 0 | 0 | 283 | 0 | 0 | Mono | Mono | | 18903250 | V3586 | E2b | | 4033 |
| 4065 | IV | 826 | 0 | 1 | 282 | 0 | 1 | Mono | Mono | | 18961021 | V3640 | E2b | | 4065 |
| 4249 | IV | 827 | 0 | 0 | 283 | 0 | 0 | Mono | Mono | | 19346290 | Z1023 | E2b | | 4249 |
| 4735 | IV | 826 | 0 | 1 | 283 | 0 | 0 | Mono | Mono | | 21779234 | M98 | E2b | | 4735 |
| 4882 | IV | 827 | 0 | 0 | 283 | 0 | 0 | Mono | Mono | | 21924329 | M90 | E2b1 | | 4882 |
| 4840 | IV | 827 | 0 | 0 | 283 | 0 | 0 | Mono | Mono | | 21896216 | M85 | E2b1a | | 4840 |
| 100 | IV | 826 | 0 | 1 | 283 | 0 | 0 | Mono | Mono | | 2771528 | Z971 | E2b1a1~ | | 100 |
| 523 | IV | 827 | 0 | 0 | 283 | 0 | 0 | Mono | Mono | | 6764869 | CTS462 | E2b1a1~ | | 523 |
| 714 | IV | 827 | 0 | 0 | 283 | 0 | 0 | Mono | Mono | | 7046076 | CTS845 | E2b1a1~ | | 714 |
| 2696 | IV | 826 | 0 | 1 | 283 | 0 | 0 | Mono | Mono | | 15520874 | V3013 | E2b1a1~ | | 2696 |
| 4812 | IV | 827 | 0 | 0 | 283 | 0 | 0 | Mono | Mono | | 21880748 | P45 | E2b1a1a~ | | 4812 |
| 1980 | IV | 827 | 0 | 0 | 283 | 0 | 0 | Mono | Mono | | 14063951 | CTS1779 | E2b2 | | 1980 |
| 2445 | IV | 827 | 0 | 0 | 283 | 0 | 0 | Mono | Mono | | 15019403 | CTS3562 | E2b2 | | 2445 |
| 59 | I | 827 | 0 | 0 | 246 | 37 | 0 | M9- | 1 – 10 | | 2722506 | S8235 | F | | 59 |
| 157 | I | 827 | 0 | 0 | 246 | 37 | 0 | M9- | 1 – 10 | | 2863665 | V205 | F | | 157 |
| 193 | III | 823 | 0 | 4 | 245 | 37 | 1 | M9- | 1 – 10 | | 2912385 | PF2591 | F | | 193 |
| 663 | I | 827 | 0 | 0 | 246 | 37 | 0 | M9- | 1 – 10 | | 6941218 | PF2602 | F | | 663 |
| 862 | I | 827 | 0 | 0 | 246 | 37 | 0 | M9- | 1 – 10 | | 7517055 | PF2608 | F | | 862 |
| 961 | I | 827 | 0 | 0 | 246 | 37 | 0 | M9- | 1 – 10 | | 7671530 | YSC0001295 | F | | 961 |
| 992 | I | 827 | 0 | 0 | 246 | 37 | 0 | M9- | 1 – 10 | | 7726849 | PF2611 | F | | 992 |
| 1179 | III | 825 | 0 | 2 | 246 | 37 | 0 | M9- | 1 – 10 | | 8131538 | M3650 | F | | 1179 |
| 1216 | I | 827 | 0 | 0 | 246 | 37 | 0 | M9- | 1 – 10 | | 8227605 | PF2615 | F | | 1216 |
| 1325 | I | 827 | 0 | 0 | 246 | 37 | 0 | M9- | 1 – 10 | | 8424089 | PF2617 | F | | 1325 |
| 1345 | I | 827 | 0 | 0 | 246 | 37 | 0 | M9- | 1 – 10 | | 8442341 | YSC0001297 | F | | 1345 |
| 1379 | I | 827 | 0 | 0 | 246 | 37 | 0 | M9- | 1 – 10 | | 8474189 | PF2618 | F | | 1379 |
| 1459 | I | 827 | 0 | 0 | 246 | 37 | 0 | M9- | 1 – 10 | | 8572150 | PF2621 | F | | 1459 |
| 1470 | I | 827 | 0 | 0 | 246 | 37 | 0 | M9- | 1 – 10 | | 8589031 | YSC0001299 | F | A | 1470 |
| 1483 | I | 827 | 0 | 0 | 246 | 37 | 0 | M9- | 1 – 10 | | 8602415 | PF2623 | F | | 1483 |
| 1533 | I | 827 | 0 | 0 | 246 | 37 | 0 | M9- | 1 – 10 | | 8680661 | PF2625 | F | | 1533 |
| 2026 | I | 827 | 0 | 0 | 246 | 37 | 0 | M9- | 1 – 10 | | 14149772 | PF2651 | F | | 2026 |
| 2032 | I | 827 | 0 | 0 | 246 | 37 | 0 | M9- | 1 – 10 | | 14171665 | PF2653 | F | | 2032 |
| 2045 | I | 827 | 0 | 0 | 246 | 37 | 0 | M9- | 1 – 10 | | 14199284 | PF2655 | F | | 2045 |
| 2093 | I | 827 | 0 | 0 | 246 | 37 | 0 | M9- | 1 – 10 | | 14294504 | PF2658 | F | | 2093 |
| 2108 | I | 827 | 0 | 0 | 246 | 37 | 0 | M9- | 1 – 10 | | 14334396 | PF2660 | F | | 2108 |
| 2430 | I | 827 | 0 | 0 | 246 | 37 | 0 | M9- | 1 – 10 | | 14996960 | PF2666 | F | | 2430 |
| 2516 | I | 827 | 0 | 0 | 246 | 37 | 0 | M9- | 1 – 10 | | 15095345 | PF2667 | F | | 2516 |
| 2556 | I | 827 | 0 | 0 | 246 | 37 | 0 | M9- | 1 – 10 | | 15234830 | PF2669 | F | | 2556 |
| 2587 | I | 827 | 0 | 0 | 246 | 37 | 0 | M9- | 1 – 10 | | 15323154 | PF2671 | F | | 2587 |
| 2698 | I | 827 | 0 | 0 | 246 | 37 | 0 | M9- | 1 – 10 | | 15526751 | PF2673 | F | | 2698 |
| 2775 | III | 826 | 0 | 1 | 245 | 37 | 1 | M9- | 1 – 10 | | 15651438 | PF2677 | F | | 2775 |
| 2895 | I | 827 | 0 | 0 | 246 | 37 | 0 | M9- | 1 – 10 | | 15955432 | V3268 | F | | 2895 |
| 2928 | I | 827 | 0 | 0 | 246 | 37 | 0 | M9- | 1 – 10 | | 16178042 | PF2685 | F | | 2928 |
| 2963 | I | 827 | 0 | 0 | 246 | 37 | 0 | M9- | 1 – 10 | | 16242316 | PF2686 | F | | 2963 |
| 3015 | III | 825 | 0 | 2 | 246 | 37 | 0 | M9- | 1 – 10 | | 16364286 | PF2688 | F | | 3015 |

| | | | M9+ | | | M9- | | | | | Build 37 | | ISOGG | | |
|---|---|---|---|---|---|---|---|---|---|---|---|---|---|---|---|
| List | Stage | 0 | 1 | . | 0 | 1 | . | M9? | Clade(s) | Prob. | Position | Name | Subgroup Name | Amb. | List |
| 3029 | I | 827 | 0 | 0 | 246 | 37 | 0 | M9- | 1 – 10 | | 16401339 | PF2690 | F | | 3029 |
| 3143 | I | 827 | 0 | 0 | 246 | 37 | 0 | M9- | 1 – 10 | | 16690780 | PF2693 | F | | 3143 |
| 3216 | I | 827 | 0 | 0 | 246 | 37 | 0 | M9- | 1 – 10 | | 16839641 | PF2696 | F | | 3216 |
| 3265 | I | 827 | 0 | 0 | 246 | 37 | 0 | M9- | 1 – 10 | | 16906683 | PF2698 | F | | 3265 |
| 3355 | I | 827 | 0 | 0 | 246 | 37 | 0 | M9- | 1 – 10 | | 17152659 | PF2700 | F | | 3355 |
| 3392 | I | 827 | 0 | 0 | 246 | 37 | 0 | M9- | 1 – 10 | | 17256018 | PF2702 | F | | 3392 |
| 3414 | I | 827 | 0 | 0 | 246 | 37 | 0 | M9- | 1 – 10 | | 17311975 | PF2703 | F | | 3414 |
| 3445 | I | 827 | 0 | 0 | 246 | 37 | 0 | M9- | 1 – 10 | | 17398598 | PF2704 | F | | 3445 |
| 3472 | III | 821 | 0 | 6 | 246 | 37 | 0 | M9- | 1 – 10 | | 17464197 | L132.2 | F | A | 3472 |
| 3482 | I | 827 | 0 | 0 | 246 | 37 | 0 | M9- | 1 – 10 | | 17493513 | PF2706 | F | | 3482 |
| 3606 | I | 827 | 0 | 0 | 246 | 37 | 0 | M9- | 1 – 10 | | 17762668 | YSC0001308 | F | | 3606 |
| 3755 | I | 827 | 0 | 0 | 246 | 37 | 0 | M9- | 1 – 10 | | 18051289 | PF2716 | F | | 3755 |
| 3782 | I | 827 | 0 | 0 | 246 | 37 | 0 | M9- | 1 – 10 | | 18097251 | PF2717 | F | | 3782 |
| 3814 | III | 818 | 0 | 9 | 244 | 37 | 2 | M9- | 1 – 10 | | 18178091 | PF2719 | F | | 3814 |
| 3884 | I | 827 | 0 | 0 | 246 | 37 | 0 | M9- | 1 – 10 | | 18578476 | PF2720 | F | | 3884 |
| 3965 | I | 827 | 0 | 0 | 246 | 37 | 0 | M9- | 1 – 10 | | 18765649 | PF2722 | F | | 3965 |
| 4180 | I | 827 | 0 | 0 | 246 | 37 | 0 | M9- | 1 – 10 | | 19205814 | PF2730 | F | | 4180 |
| 4203 | I | 827 | 0 | 0 | 246 | 37 | 0 | M9- | 1 – 10 | | 19237093 | V3940 | F | | 4203 |
| 4251 | III | 821 | 0 | 6 | 245 | 37 | 1 | M9- | 1 – 10 | | 19349615 | PF2734 | F | | 4251 |
| 4366 | I | 827 | 0 | 0 | 246 | 37 | 0 | M9- | 1 – 10 | | 21080707 | PF2736 | F | | 4366 |
| 4395 | I | 827 | 0 | 0 | 246 | 37 | 0 | M9- | 1 – 10 | | 21147336 | PF2737 | F | | 4395 |
| 4591 | I | 827 | 0 | 0 | 246 | 37 | 0 | M9- | 1 – 10 | | 21618856 | PF2741 | F | | 4591 |
| 4748 | I | 827 | 0 | 0 | 246 | 37 | 0 | M9- | 1 – 10 | | 21811276 | PF2745 | F | | 4748 |
| 4876 | I | 827 | 0 | 0 | 246 | 37 | 0 | M9- | 1 – 10 | | 21917313 | PF2746 | F | | 4876 |
| 4896 | I | 827 | 0 | 0 | 246 | 37 | 0 | M9- | 1 – 10 | | 21937573 | PF2747 | F | | 4896 |
| 4953 | I | 827 | 0 | 0 | 246 | 37 | 0 | M9- | 1 – 10 | | 22108101 | PF2748 | F | | 4953 |
| 5100 | I | 827 | 0 | 0 | 246 | 37 | 0 | M9- | 1 – 10 | | 22673903 | PF2758 | F | | 5100 |
| 5299 | I | 827 | 0 | 0 | 246 | 37 | 0 | M9- | 1 – 10 | | 23040647 | PF2762 | F | | 5299 |
| 5644 | III | 825 | 0 | 2 | 246 | 37 | 0 | M9- | 1 – 10 | | 24359931 | PF2771 | F | | 5644 |
| 5713 | I | 827 | 0 | 0 | 246 | 37 | 0 | M9- | 1 – 10 | | 28650343 | PF2775 | F | | 5713 |
| 2168 | IV | 827 | 0 | 0 | 283 | 0 | 0 | Mono | Mono | | 14496387 | M481 | F3 | | 2168 |
| 2202 | III | 1 | 820 | 6 | 21 | 260 | 2 | BOTH | | PI | 14624254 | PF3499 | Freq. Mut. SNP in R | W | 2202 |
| 31 | I | 827 | 0 | 0 | 262 | 21 | 0 | M9- | 11 – 20 | | 2681740 | PF2780 | G | | 31 |
| 104 | I | 827 | 0 | 0 | 262 | 21 | 0 | M9- | 11 – 20 | | 2785630 | PF2782 | G | | 104 |
| 151 | I | 827 | 0 | 0 | 262 | 21 | 0 | M9- | 11 – 20 | | 2846401 | U17 | G | | 151 |
| 163 | I | 827 | 0 | 0 | 262 | 21 | 0 | M9- | 11 – 20 | | 2871867 | M3445 | G | | 163 |
| 496 | III | 826 | 0 | 1 | 261 | 21 | 1 | M9- | 11 – 20 | | 6716150 | M3448 | G | | 496 |
| 625 | I | 827 | 0 | 0 | 262 | 21 | 0 | M9- | 11 – 20 | | 6931141 | PF2844 | G | | 625 |
| 679 | I | 827 | 0 | 0 | 262 | 21 | 0 | M9- | 11 – 20 | | 6955839 | M3452 | G | | 679 |
| 771 | I | 827 | 0 | 0 | 262 | 21 | 0 | M9- | 11 – 20 | | 7157834 | PF2850 | G | | 771 |
| 874 | III | 827 | 0 | 0 | 262 | 19 | 2 | M9- | 11 – 20 | MD | 7537241 | PF2857 | G | | 874 |
| 875 | III | 826 | 0 | 1 | 262 | 21 | 0 | M9- | 11 – 20 | | 7537950 | PF2858 | G | | 875 |
| 893 | III | 826 | 0 | 1 | 261 | 21 | 1 | M9- | 11 – 20 | | 7565637 | PF2859 | G | | 893 |
| 1056 | I | 827 | 0 | 0 | 262 | 21 | 0 | M9- | 11 – 20 | | 7840218 | PF2866 | G | | 1056 |
| 1080 | I | 827 | 0 | 0 | 262 | 21 | 0 | M9- | 11 – 20 | | 7899682 | PF2867 | G | | 1080 |
| 1100 | I | 827 | 0 | 0 | 262 | 21 | 0 | M9- | 11 – 20 | | 7927218 | PF2868 | G | | 1100 |
| 1101 | I | 827 | 0 | 0 | 262 | 21 | 0 | M9- | 11 – 20 | | 7930724 | PF2869 | G | | 1101 |
| 1123 | I | 827 | 0 | 0 | 262 | 21 | 0 | M9- | 11 – 20 | | 7992031 | PF2872 | G | | 1123 |
| 1212 | I | 827 | 0 | 0 | 262 | 21 | 0 | M9- | 11 – 20 | | 8219021 | PF2876 | G | | 1212 |
| 1219 | I | 827 | 0 | 0 | 262 | 21 | 0 | M9- | 11 – 20 | | 8231862 | PF2877 | G | | 1219 |
| 1229 | I | 827 | 0 | 0 | 262 | 21 | 0 | M9- | 11 – 20 | | 8240725 | L519 | G | | 1229 |
| 1271 | I | 827 | 0 | 0 | 262 | 21 | 0 | M9- | 11 – 20 | | 8327892 | PF2879 | G | | 1271 |
| 1302 | I | 827 | 0 | 0 | 262 | 21 | 0 | M9- | 11 – 20 | | 8387539 | PF2880 | G | | 1302 |
| 1323 | I | 827 | 0 | 0 | 262 | 21 | 0 | M9- | 11 – 20 | | 8422993 | PF2881 | G | | 1323 |
| 1452 | I | 827 | 0 | 0 | 262 | 21 | 0 | M9- | 11 – 20 | | 8563874 | PF2888 | G | | 1452 |
| 1482 | I | 827 | 0 | 0 | 262 | 21 | 0 | M9- | 11 – 20 | | 8600158 | PF2889 | G | | 1482 |
| 1485 | I | 827 | 0 | 0 | 262 | 21 | 0 | M9- | 11 – 20 | | 8602816 | PF2890 | G | | 1485 |
| 1497 | I | 827 | 0 | 0 | 262 | 21 | 0 | M9- | 11 – 20 | | 8614138 | PF3139 | G | | 1497 |
| 1543 | I | 827 | 0 | 0 | 262 | 21 | 0 | M9- | 11 – 20 | | 8691654 | Z3215 | G | | 1543 |
| 1548 | I | 824 | 3 | 0 | 262 | 21 | 0 | BOTH | | PI | 8700380 | PF2892 | G | | 1548 |
| 1692 | I | 827 | 0 | 0 | 262 | 21 | 0 | M9- | 11 – 20 | | 9448354 | PF2899 | G | | 1692 |
| 1707 | III | 827 | 0 | 0 | 262 | 18 | 3 | M9- | 11 – 20 | MD | 9523592 | PF2901 | G | | 1707 |
| 1875 | III | 826 | 0 | 1 | 260 | 21 | 2 | M9- | 11 – 20 | | 13679469 | PF2918 | G | | 1875 |
| 1945 | I | 827 | 0 | 0 | 262 | 21 | 0 | M9- | 11 – 20 | | 13987899 | PF2924 | G | | 1945 |
| 2039 | I | 827 | 0 | 0 | 262 | 21 | 0 | M9- | 11 – 20 | | 14188094 | PF2935 | G | | 2039 |
| 2040 | I | 827 | 0 | 0 | 262 | 21 | 0 | M9- | 11 – 20 | | 14190812 | M3512 | G | | 2040 |
| 2047 | I | 827 | 0 | 0 | 262 | 21 | 0 | M9- | 11 – 20 | | 14199646 | PF3265 | G | | 2047 |
| 2054 | I | 827 | 0 | 0 | 262 | 21 | 0 | M9- | 11 – 20 | | 14207268 | PF2939 | G | | 2054 |
| 2061 | I | 827 | 0 | 0 | 262 | 21 | 0 | M9- | 11 – 20 | | 14220356 | PF2940 | G | | 2061 |
| 2066 | III | 827 | 0 | 0 | 262 | 14 | 7 | M9- | 11 – 20 | MD | 14235140 | PF2941.2 | G | A | 2066 |
| 2071 | III | 814 | 0 | 13 | 259 | 21 | 3 | M9- | 11 – 20 | | 14243137 | M3516 | G | | 2071 |
| 2113 | I | 827 | 0 | 0 | 262 | 21 | 0 | M9- | 11 – 20 | | 14338503 | PF2946 | G | | 2113 |
| 2140 | I | 0 | 827 | 0 | 21 | 262 | 0 | M9- | 11 – 20 | | 14432928 | U6 | G | | 2140 |
| 2150 | I | 0 | 827 | 0 | 21 | 262 | 0 | M9- | 11 – 20 | | 14469411 | PF2951 | G | | 2150 |
| 2191 | I | 0 | 827 | 0 | 21 | 262 | 0 | M9- | 11 – 20 | | 14577177 | U2 | G | | 2191 |
| 2401 | III | 0 | 826 | 1 | 21 | 262 | 0 | M9- | 11 – 20 | | 14958218 | PF3135 | G | | 2401 |
| 2425 | I | 0 | 827 | 0 | 21 | 262 | 0 | M9- | 11 – 20 | | 14989721 | S284 | G | | 2425 |
| 2470 | I | 0 | 827 | 0 | 21 | 262 | 0 | M9- | 11 – 20 | | 15027529 | PF2957 | G | | 2470 |
| 2510 | I | 0 | 827 | 0 | 21 | 262 | 0 | M9- | 11 – 20 | | 15086183 | U7 | G | | 2510 |
| 2545 | I | 0 | 827 | 0 | 21 | 262 | 0 | M9- | 11 – 20 | | 15204708 | L402 | G | | 2545 |
| 2546 | I | 0 | 827 | 0 | 21 | 262 | 0 | M9- | 11 – 20 | | 15204710 | U21 | G | | 2546 |
| 2682 | III | 827 | 0 | 0 | 262 | 11 | 10 | M9- | 11 – 20 | MD | 15504804 | PF2966 | G | | 2682 |
| 2790 | I | 827 | 0 | 0 | 262 | 21 | 0 | M9- | 11 – 20 | | 15667235 | PF2973 | G | | 2790 |
| 2797 | I | 827 | 0 | 0 | 262 | 21 | 0 | M9- | 11 – 20 | | 15693336 | PF2974 | G | | 2797 |
| 2844 | I | 827 | 0 | 0 | 262 | 21 | 0 | M9- | 11 – 20 | | 15802681 | PF2978 | G | | 2844 |
| 2934 | I | 827 | 0 | 0 | 262 | 21 | 0 | M9- | 11 – 20 | | 16185081 | PF3136 | G | | 2934 |
| 2948 | I | 827 | 0 | 0 | 262 | 21 | 0 | M9- | 11 – 20 | | 16203361 | M3537 | G | | 2948 |
| 3038 | I | 827 | 0 | 0 | 262 | 21 | 0 | M9- | 11 – 20 | | 16408569 | M3540 | G | | 3038 |
| 3043 | I | 827 | 0 | 0 | 262 | 21 | 0 | M9- | 11 – 20 | | 16419934 | PF2985 | G | | 3043 |
| 3047 | III | 825 | 0 | 2 | 262 | 21 | 0 | M9- | 11 – 20 | | 16439267 | M3542 | G | | 3047 |
| 3078 | I | 827 | 0 | 0 | 262 | 21 | 0 | M9- | 11 – 20 | | 16499780 | PF2990 | G | | 3078 |
| 3134 | I | 827 | 0 | 0 | 262 | 21 | 0 | M9- | 11 – 20 | | 16651032 | PF2992 | G | | 3134 |
| 3253 | I | 827 | 0 | 0 | 262 | 21 | 0 | M9- | 11 – 20 | | 16896148 | L836 | G | | 3253 |
| 3273 | I | 827 | 0 | 0 | 262 | 21 | 0 | M9- | 11 – 20 | | 16929270 | PF2996 | G | | 3273 |
| 3343 | III | 826 | 0 | 1 | 262 | 21 | 0 | M9- | 11 – 20 | | 17119047 | M3550 | G | | 3343 |
| 3378 | III | 827 | 0 | 0 | 261 | 21 | 1 | M9- | 11 – 20 | | 17210745 | PF3005 | G | | 3378 |

| | | | M9+ | | | M9- | | | | | Build 37 | | ISOGG | | |
|---|---|---|---|---|---|---|---|---|---|---|---|---|---|---|---|
| List | Stage | 0 | 1 | . | 0 | 1 | . | M9? | Clade(s) | Prob. | Position | Name | Subgroup Name | Amb. | List |
| 3396 | I | 827 | 0 | 0 | 262 | 21 | 0 | M9- | 11 – 20 | | 17281783 | PF3006 | G | | 3396 |
| 3441 | I | 827 | 0 | 0 | 262 | 21 | 0 | M9- | 11 – 20 | | 17393643 | M3557 | G | | 3441 |
| 3468 | III | 826 | 0 | 1 | 262 | 21 | 0 | M9- | 11 – 20 | | 17453851 | PF3009 | G | | 3468 |
| 3546 | I | 827 | 0 | 0 | 262 | 21 | 0 | M9- | 11 – 20 | | 17610571 | PF3015 | G | | 3546 |
| 3597 | I | 827 | 0 | 0 | 262 | 21 | 0 | M9- | 11 – 20 | | 17747521 | PF3016 | G | | 3597 |
| 3625 | III | 827 | 0 | 0 | 261 | 21 | 1 | M9- | 11 – 20 | | 17798903 | PF3017 | G | | 3625 |
| 3657 | I | 827 | 0 | 0 | 262 | 21 | 0 | M9- | 11 – 20 | | 17853245 | L837 | G | | 3657 |
| 3768 | I | 827 | 0 | 0 | 262 | 21 | 0 | M9- | 11 – 20 | | 18070349 | M3565 | G | | 3768 |
| 3805 | III | 827 | 0 | 0 | 261 | 21 | 1 | M9- | 11 – 20 | | 18162022 | M3566 | G | | 3805 |
| 3848 | III | 825 | 0 | 2 | 262 | 21 | 0 | M9- | 11 – 20 | | 18393536 | PF3023 | G | | 3848 |
| 3951 | III | 826 | 0 | 1 | 261 | 19 | 3 | M9- | 11 – 20 | MD | 18737609 | PF3025 | G | | 3951 |
| 4061 | I | 827 | 0 | 0 | 262 | 21 | 0 | M9- | 11 – 20 | | 18957208 | PF3027 | G | | 4061 |
| 4075 | III | 826 | 0 | 1 | 262 | 21 | 0 | M9- | 11 – 20 | | 18979775 | M3570 | G | | 4075 |
| 4089 | I | 827 | 0 | 0 | 262 | 21 | 0 | M9- | 11 – 20 | | 19030998 | PF3031 | G | | 4089 |
| 4090 | I | 827 | 0 | 0 | 262 | 21 | 0 | M9- | 11 – 20 | | 19033112 | PF3032 | G | | 4090 |
| 4185 | III | 826 | 0 | 1 | 262 | 21 | 0 | M9- | 11 – 20 | | 19215139 | PF3037 | G | | 4185 |
| 4343 | I | 827 | 0 | 0 | 262 | 21 | 0 | M9- | 11 – 20 | | 20823823 | PF3045 | G | | 4343 |
| 4402 | III | 826 | 0 | 1 | 262 | 21 | 0 | M9- | 11 – 20 | | 21162869 | PF3046 | G | | 4402 |
| 4480 | I | 827 | 0 | 0 | 262 | 21 | 0 | M9- | 11 – 20 | | 21362016 | PF3050 | G | | 4480 |
| 4509 | I | 827 | 0 | 0 | 262 | 21 | 0 | M9- | 11 – 20 | | 21412501 | PF3052 | G | | 4509 |
| 4518 | I | 827 | 0 | 0 | 262 | 21 | 0 | M9- | 11 – 20 | | 21447363 | PF3053 | G | | 4518 |
| 4528 | III | 824 | 0 | 3 | 262 | 21 | 0 | M9- | 11 – 20 | | 21469197 | PF3054 | G | | 4528 |
| 4539 | III | 826 | 0 | 1 | 261 | 21 | 1 | M9- | 11 – 20 | | 21495813 | PF3057 | G | | 4539 |
| 4597 | I | 827 | 0 | 0 | 262 | 21 | 0 | M9- | 11 – 20 | | 21626642 | PF3061 | G | | 4597 |
| 4618 | I | 827 | 0 | 0 | 262 | 21 | 0 | M9- | 11 – 20 | | 21648433 | PF3063 | G | | 4618 |
| 4771 | III | 827 | 0 | 0 | 261 | 21 | 1 | M9- | 11 – 20 | | 21865624 | PF3069 | G | | 4771 |
| 4900 | III | 826 | 0 | 1 | 262 | 21 | 0 | M9- | 11 – 20 | | 21939157 | PF3070 | G | | 4900 |
| 4904 | I | 827 | 0 | 0 | 262 | 21 | 0 | M9- | 11 – 20 | | 21954611 | PF3071 | G | | 4904 |
| 4979 | III | 827 | 0 | 0 | 261 | 21 | 1 | M9- | 11 – 20 | | 22163252 | PF3075 | G | | 4979 |
| 4983 | I | 827 | 0 | 0 | 262 | 21 | 0 | M9- | 11 – 20 | | 22167631 | S7530 | G | | 4983 |
| 5063 | III | 820 | 0 | 7 | 262 | 21 | 0 | M9- | 11 – 20 | | 22579947 | PF3092 | G | | 5063 |
| 5126 | III | 824 | 0 | 3 | 260 | 21 | 2 | M9- | 11 – 20 | | 22729194 | M3611 | G | | 5126 |
| 5130 | I | 827 | 0 | 0 | 262 | 21 | 0 | M9- | 11 – 20 | | 22730922 | M3612 | G | | 5130 |
| 5204 | I | 827 | 0 | 0 | 262 | 21 | 0 | M9- | 11 – 20 | | 22834341 | PF3098 | G | | 5204 |
| 5209 | I | 827 | 0 | 0 | 262 | 21 | 0 | M9- | 11 – 20 | | 22848965 | M3206 | G | | 5209 |
| 5280 | I | 827 | 0 | 0 | 262 | 21 | 0 | M9- | 11 – 20 | | 22997377 | PF3100 | G | | 5280 |
| 5290 | I | 827 | 0 | 0 | 262 | 21 | 0 | M9- | 11 – 20 | | 23023554 | M3618 | G | | 5290 |
| 5305 | I | 827 | 0 | 0 | 262 | 21 | 0 | M9- | 11 – 20 | | 23059496 | PF3104 | G | | 5305 |
| 5319 | I | 827 | 0 | 0 | 262 | 21 | 0 | M9- | 11 – 20 | | 23074190 | M3620 | G | | 5319 |
| 5328 | III | 824 | 0 | 3 | 261 | 21 | 1 | M9- | 11 – 20 | | 23095144 | PF3106 | G | | 5328 |
| 5352 | I | 827 | 0 | 0 | 262 | 21 | 0 | M9- | 11 – 20 | | 23151673 | M3622 | G | | 5352 |
| 5428 | I | 827 | 0 | 0 | 262 | 21 | 0 | M9- | 11 – 20 | | 23346582 | PF3114 | G | | 5428 |
| 5522 | I | 827 | 0 | 0 | 262 | 21 | 0 | M9- | 11 – 20 | | 23578115 | PF3117 | G | | 5522 |
| 5535 | III | 827 | 0 | 0 | 261 | 21 | 1 | M9- | 11 – 20 | | 23621266 | PF3118 | G | | 5535 |
| 5569 | I | 827 | 0 | 0 | 262 | 21 | 0 | M9- | 11 – 20 | | 23793740 | PF3122 | G | | 5569 |
| 5642 | III | 825 | 0 | 2 | 262 | 20 | 1 | M9- | 11 – 20 | MD | 24357567 | PF3123 | G | | 5642 |
| 2135 | IV | 827 | 0 | 0 | 283 | 0 | 0 | Mono | Mono | | 14406527 | Z3308 | G1 | | 2135 |
| 2707 | IV | 827 | 0 | 0 | 283 | 0 | 0 | Mono | Mono | | 15541217 | L833 | G1 | | 2707 |
| 5141 | IV | 827 | 0 | 0 | 283 | 0 | 0 | Mono | Mono | | 22741740 | M285 | G1 | | 5141 |
| 5383 | IV | 827 | 0 | 0 | 283 | 0 | 0 | Mono | Mono | | 23243942 | M342 | G1 | | 5383 |
| 1393 | IV | 827 | 0 | 0 | 283 | 0 | 0 | Mono | Mono | | 8483661 | L1324 | G1a1a1a | | 1393 |
| 2158 | IV | 826 | 0 | 1 | 283 | 0 | 0 | Mono | Mono | | 14481609 | L1325 | G1a1a1a | | 2158 |
| 4166 | IV | 827 | 0 | 0 | 283 | 0 | 0 | Mono | Mono | | 19179085 | L1414 | G1a1a1a | | 4166 |
| 4207 | IV | 827 | 0 | 0 | 283 | 0 | 0 | Mono | Mono | | 19240002 | L1327 | G1a1a1a | | 4207 |
| 616 | IV | 827 | 0 | 0 | 283 | 0 | 0 | Mono | Mono | | 6909339 | L1323 | G1a1a1a2 | | 616 |
| 4023 | IV | 827 | 0 | 0 | 283 | 0 | 0 | Mono | Mono | | 18886769 | F2885 | G1a2 | | 4023 |
| 576 | IV | 827 | 0 | 0 | 283 | 0 | 0 | Mono | Mono | | 6863908 | L830 | G1b | | 576 |
| 2088 | IV | 827 | 0 | 0 | 283 | 0 | 0 | Mono | Mono | | 14286626 | L832 | G1b | | 2088 |
| 3619 | IV | 827 | 0 | 0 | 283 | 0 | 0 | Mono | Mono | | 17782015 | L835 | G1b | | 3619 |
| 900 | III | 827 | 0 | 0 | 262 | 20 | 1 | M9- | 11 – 20 | MD | 7571775 | PF2860 | G2 | | 900 |
| 1120 | I | 827 | 0 | 0 | 262 | 21 | 0 | M9- | 11 – 20 | | 7978725 | PF3138 | G2 | A | 1120 |
| 1328 | I | 827 | 0 | 0 | 262 | 21 | 0 | M9- | 11 – 20 | | 8427005 | PF2883 | G2 | | 1328 |
| 1389 | I | 827 | 0 | 0 | 262 | 21 | 0 | M9- | 11 – 20 | | 8482393 | PF2886 | G2 | | 1389 |
| 1440 | I | 827 | 0 | 0 | 262 | 21 | 0 | M9- | 11 – 20 | | 8545324 | PF2887 | G2 | | 1440 |
| 1542 | I | 827 | 0 | 0 | 262 | 21 | 0 | M9- | 11 – 20 | | 8687693 | PF2891 | G2 | | 1542 |
| 1557 | I | 827 | 0 | 0 | 262 | 21 | 0 | M9- | 11 – 20 | | 8719593 | PF2893 | G2 | | 1557 |
| 1775 | I | 827 | 0 | 0 | 262 | 21 | 0 | M9- | 11 – 20 | | 9907842 | PF2904 | G2 | | 1775 |
| 1829 | III | 827 | 0 | 0 | 260 | 21 | 2 | M9- | 11 – 20 | | 13205148 | PF2909 | G2 | | 1829 |
| 2006 | I | 827 | 0 | 0 | 262 | 21 | 0 | M9- | 11 – 20 | | 14103263 | PF2929 | G2 | | 2006 |
| 2011 | III | 825 | 0 | 2 | 262 | 21 | 0 | M9- | 11 – 20 | | 14116322 | PF2931 | G2 | | 2011 |
| 2700 | I | 827 | 0 | 0 | 262 | 21 | 0 | M9- | 11 – 20 | | 15528792 | PF2968 | G2 | | 2700 |
| 2771 | I | 827 | 0 | 0 | 262 | 21 | 0 | M9- | 11 – 20 | | 15635425 | PF2971 | G2 | | 2771 |
| 3199 | I | 827 | 0 | 0 | 262 | 21 | 0 | M9- | 11 – 20 | | 16817402 | M3545 | G2 | | 3199 |
| 3260 | I | 827 | 0 | 0 | 262 | 21 | 0 | M9- | 11 – 20 | | 16903051 | PF2995 | G2 | | 3260 |
| 3330 | I | 827 | 0 | 0 | 262 | 21 | 0 | M9- | 11 – 20 | | 17088129 | PF2999 | G2 | | 3330 |
| 3360 | III | 576 | 0 | 251 | 179 | 21 | 83 | M9- | 11 – 20 | | 17174741 | PF3002 | G2 | | 3360 |
| 3478 | I | 827 | 0 | 0 | 262 | 21 | 0 | M9- | 11 – 20 | | 17476068 | PF3010 | G2 | | 3478 |
| 4136 | I | 827 | 0 | 0 | 262 | 21 | 0 | M9- | 11 – 20 | | 19119067 | PF3035 | G2 | A | 4136 |
| 4304 | III | 825 | 0 | 2 | 262 | 21 | 0 | M9- | 11 – 20 | | 19493301 | PF3042 | G2 | | 4304 |
| 4394 | I | 827 | 0 | 0 | 262 | 21 | 0 | M9- | 11 – 20 | | 21147058 | Z6103 | G2 | | 4394 |
| 4500 | I | 827 | 0 | 0 | 262 | 21 | 0 | M9- | 11 – 20 | | 21401188 | M3584 | G2 | | 4500 |
| 4604 | I | 827 | 0 | 0 | 262 | 21 | 0 | M9- | 11 – 20 | | 21637589 | M3592 | G2 | | 4604 |
| 4625 | I | 827 | 0 | 0 | 262 | 21 | 0 | M9- | 11 – 20 | | 21663882 | PF3064 | G2 | | 4625 |
| 4942 | I | 827 | 0 | 0 | 262 | 21 | 0 | M9- | 11 – 20 | | 22072097 | PF3140 | G2 | | 4942 |
| 5109 | I | 827 | 0 | 0 | 262 | 21 | 0 | M9- | 11 – 20 | | 22697266 | PF3095 | G2 | | 5109 |
| 5547 | III | 827 | 0 | 0 | 261 | 14 | 8 | M9- | 11 – 20 | MD | 23739606 | PF3119 | G2 | | 5547 |
| 122 | I | 827 | 0 | 0 | 262 | 21 | 0 | M9- | 11 – 20 | | 2812741 | PF2783 | G2a | | 122 |
| 993 | I | 827 | 0 | 0 | 262 | 21 | 0 | M9- | 11 – 20 | | 7727677 | PF2863 | G2a | | 993 |
| 1962 | I | 827 | 0 | 0 | 262 | 21 | 0 | M9- | 11 – 20 | | 14028148 | S149 | G2a | | 1962 |
| 2738 | I | 827 | 0 | 0 | 262 | 21 | 0 | M9- | 11 – 20 | | 15588776 | PF2969 | G2a | | 2738 |
| 2782 | I | 827 | 0 | 0 | 262 | 21 | 0 | M9- | 11 – 20 | | 15660640 | PF2972 | G2a | | 2782 |
| 2972 | III | 826 | 0 | 1 | 262 | 21 | 0 | M9- | 11 – 20 | | 16262350 | M3320 | G2a | | 2972 |
| 3188 | III | 826 | 0 | 1 | 262 | 21 | 0 | M9- | 11 – 20 | | 16802506 | PF2993 | G2a | | 3188 |
| 3232 | III | 826 | 0 | 1 | 262 | 21 | 0 | M9- | 11 – 20 | | 16861108 | PF2994 | G2a | | 3232 |
| 3522 | I | 827 | 0 | 0 | 262 | 21 | 0 | M9- | 11 – 20 | | 17571517 | PF3013 | G2a | | 3522 |
| 4538 | III | 826 | 0 | 1 | 262 | 21 | 0 | M9- | 11 – 20 | | 21493984 | PF3056 | G2a | | 4538 |

| | | | M9+ | | | M9- | | | | | Build 37 | | ISOGG | | |
|---|---|---|---|---|---|---|---|---|---|---|---|---|---|---|---|
| List | Stage | 0 | 1 | . | 0 | 1 | . | M9? | Clade(s) | Prob. | Position | Name | Subgroup Name | Amb. | List |
| 4579 | III | 826 | 0 | 1 | 262 | 21 | 0 | M9- | 11 – 20 | | 21605685 | PF3060 | G2a | | 4579 |
| 4703 | I | 827 | 0 | 0 | 262 | 21 | 0 | M9- | 11 – 20 | | 21760742 | PF3066 | G2a | | 4703 |
| 4954 | I | 827 | 0 | 0 | 262 | 21 | 0 | M9- | 11 – 20 | | 22109159 | PF3073 | G2a | | 4954 |
| 5340 | I | 827 | 0 | 0 | 262 | 21 | 0 | M9- | 11 – 20 | | 23122426 | PF3108 | G2a | | 5340 |
| 5384 | I | 827 | 0 | 0 | 262 | 21 | 0 | M9- | 11 – 20 | | 23244026 | PF3112 | G2a | | 5384 |
| 5478 | I | 827 | 0 | 0 | 262 | 21 | 0 | M9- | 11 – 20 | | 23475051 | PF3116 | G2a | | 5478 |
| 591 | II | 827 | 0 | 0 | 282 | 1 | 0 | M9- | 11 | | 6882236 | Z6638 | G2a1a1a1a1 | | 591 |
| 3891 | IV | 827 | 0 | 0 | 283 | 0 | 0 | Mono | Mono | | 18588967 | Z8011 | G2a1a1a1a1a1a1a1 | | 3891 |
| 2817 | IV | 827 | 0 | 0 | 283 | 0 | 0 | Mono | Mono | | 15738379 | FGC719 | G2a1a1a1a1a1a1a1a2 | | 2817 |
| 2764 | I | 827 | 0 | 0 | 263 | 20 | 0 | M9- | 12 – 20 | | 15615340 | PF2970 | G2a2 | | 2764 |
| 1005 | I | 827 | 0 | 0 | 277 | 6 | 0 | M9- | 12 | | 7738069 | PF3147 | G2a2a | | 1005 |
| 1724 | III | 818 | 0 | 9 | 274 | 6 | 3 | M9- | 12 | | 9785736 | PF3151 | G2a2a | | 1724 |
| 2800 | I | 827 | 0 | 0 | 277 | 6 | 0 | M9- | 12 | | 15702713 | PF3161 | G2a2a | | 2800 |
| 4467 | III | 827 | 0 | 0 | 273 | 6 | 4 | M9- | 12 | | 21327198 | PF3177 | G2a2a1a | | 4467 |
| 5143 | IV | 827 | 0 | 0 | 283 | 0 | 0 | Mono | Mono | | 22741799 | M286 | G2a2a1a1a1 | | 5143 |
| 4613 | I | 827 | 0 | 0 | 277 | 6 | 0 | M9- | 12 | | 21645555 | S285 | G2a2a1a2 | | 4613 |
| 4745 | III | 823 | 0 | 4 | 276 | 2 | 5 | M9- | 12 | MD | 21802302 | Z6489^ | G2a2a1a2a1 | | 4745 |
| 948 | I | 827 | 0 | 0 | 277 | 6 | 0 | M9- | 12 | | 7652689 | FGC5671 | G2a2a1a2a1a | | 948 |
| 1042 | III | 826 | 0 | 1 | 277 | 6 | 0 | M9- | 12 | | 7811508 | Z1370.2 | G2a2a1a2a1a | | 1042 |
| 3847 | I | 827 | 0 | 0 | 277 | 6 | 0 | M9- | 12 | | 18392335 | FGC5721 | G2a2a1a2a1a | | 3847 |
| 5635 | I | 827 | 0 | 0 | 277 | 6 | 0 | M9- | 12 | | 23989884 | L166 | G2a2a1a2a1a | | 5635 |
| 449 | III | 827 | 0 | 0 | 268 | 14 | 1 | M9- | 13 – 20 | | 6425529 | PF3254 | G2a2b | | 449 |
| 462 | III | 827 | 0 | 0 | 268 | 14 | 1 | M9- | 13 – 20 | | 6627544 | PF3255 | G2a2b | | 462 |
| 568 | III | 826 | 0 | 1 | 269 | 14 | 0 | M9- | 13 – 20 | | 6855308 | M3237 | G2a2b | | 568 |
| 1249 | III | 827 | 0 | 0 | 269 | 13 | 1 | M9- | 13 – 20 | MD | 8275094 | PF3261 | G2a2b | | 1249 |
| 2225 | I | 0 | 827 | 0 | 14 | 269 | 0 | M9- | 13 – 20 | | 14692227 | U8 | G2a2b | | 2225 |
| 2762 | I | 827 | 0 | 0 | 269 | 14 | 0 | M9- | 13 – 20 | | 15604899 | S126 | G2a2b | | 2762 |
| 3532 | I | 827 | 0 | 0 | 269 | 14 | 0 | M9- | 13 – 20 | | 17589788 | PF3270 | G2a2b | | 3532 |
| 3960 | I | 827 | 0 | 0 | 269 | 14 | 0 | M9- | 13 – 20 | | 18759690 | M485 | G2a2b | | 3960 |
| 4109 | I | 827 | 0 | 0 | 269 | 14 | 0 | M9- | 13 – 20 | | 19060222 | PF3271 | G2a2b | | 4109 |
| 4179 | I | 827 | 0 | 0 | 269 | 14 | 0 | M9- | 13 – 20 | | 19205722 | PF3272 | G2a2b | | 4179 |
| 4290 | I | 827 | 0 | 0 | 269 | 14 | 0 | M9- | 13 – 20 | | 19461366 | PF3273 | G2a2b | | 4290 |
| 4398 | I | 827 | 0 | 0 | 269 | 14 | 0 | M9- | 13 – 20 | | 21151007 | PF3274 | G2a2b | | 4398 |
| 5048 | I | 827 | 0 | 0 | 269 | 14 | 0 | M9- | 13 – 20 | | 22553146 | PF3278 | G2a2b | | 5048 |
| 5315 | III | 826 | 0 | 1 | 266 | 14 | 3 | M9- | 13 – 20 | | 23071312 | PF3279 | G2a2b | | 5315 |
| 5594 | III | 827 | 0 | 0 | 269 | 12 | 2 | M9- | 13 – 20 | MD | 23871230 | PF3281 | G2a2b | | 5594 |
| 86 | III | 826 | 0 | 1 | 282 | 1 | 0 | M9- | 13 | | 2749995 | PF3285 | G2a2b1 | | 86 |
| 917 | III | 825 | 0 | 2 | 282 | 1 | 0 | M9- | 13 | | 7598950 | PF3287 | G2a2b1 | | 917 |
| 1425 | III | 827 | 0 | 0 | 281 | 1 | 1 | M9- | 13 | | 8519704 | PF3292.3 | G2a2b1 | A | 1425 |
| 1952 | II | 827 | 0 | 0 | 282 | 1 | 0 | M9- | 13 | | 14002366 | PF3297 | G2a2b1 | | 1952 |
| 2821 | II | 827 | 0 | 0 | 282 | 1 | 0 | M9- | 13 | | 15752242 | PF3301 | G2a2b1 | | 2821 |
| 2996 | II | 827 | 0 | 0 | 282 | 1 | 0 | M9- | 13 | | 16327939 | PF3304 | G2a2b1 | | 2996 |
| 4082 | IV | 825 | 0 | 2 | 281 | 0 | 2 | Mono | Mono | MD | 19005164 | PF3312 | G2a2b1 | | 4082 |
| 4122 | III | 827 | 0 | 0 | 281 | 1 | 1 | M9- | 13 | | 19087143 | M461 | G2a2b1 | | 4122 |
| 4314 | III | 825 | 0 | 2 | 282 | 1 | 0 | M9- | 13 | | 19513372 | PF3315 | G2a2b1 | | 4314 |
| 5424 | II | 827 | 0 | 0 | 282 | 1 | 0 | M9- | 13 | | 23327970 | PF3319 | G2a2b1 | | 5424 |
| 4600 | IV | 827 | 0 | 0 | 283 | 0 | 0 | Mono | Mono | | 21628300 | S133 | G2a2b1a1a1 | | 4600 |
| 5235 | IV | 827 | 0 | 0 | 283 | 0 | 0 | Mono | Mono | | 22917995 | U16 | G2a2b1a1a1 | | 5235 |
| 1465 | IV | 827 | 0 | 0 | 283 | 0 | 0 | Mono | Mono | | 8578140 | S27195 | G2a2b1a1a1a1b~ | | 1465 |
| 4611 | IV | 827 | 0 | 0 | 283 | 0 | 0 | Mono | Mono | | 21645133 | Page25 | G2a2b1a1a1a4~ | | 4611 |
| 2522 | IV | 827 | 0 | 0 | 283 | 0 | 0 | Mono | Mono | | 15104273 | M3302 | G2a2b1a2 | | 2522 |
| 5398 | IV | 827 | 0 | 0 | 283 | 0 | 0 | Mono | Mono | | 23265883 | M3422 | G2a2b1a2a | | 5398 |
| 730 | IV | 827 | 0 | 0 | 283 | 0 | 0 | Mono | Mono | | 7074221 | M3240 | G2a2b1a2a1 | | 730 |
| 518 | I | 827 | 0 | 0 | 272 | 11 | 0 | M9- | 14 – 19 | | 6744902 | PF3328 | G2a2b2a | | 518 |
| 673 | III | 823 | 0 | 4 | 271 | 11 | 1 | M9- | 14 – 19 | | 6953417 | CTS688 | G2a2b2a | | 673 |
| 743 | III | 827 | 0 | 0 | 265 | 11 | 7 | M9- | 14 – 19 | | 7100848 | CTS946 | G2a2b2a | | 743 |
| 1382 | I | 827 | 0 | 0 | 272 | 11 | 0 | M9- | 14 – 19 | | 8478026 | PF3329 | G2a2b2a | | 1382 |
| 1541 | III | 809 | 0 | 18 | 266 | 11 | 6 | M9- | 14 – 19 | | 8687649 | PF3330 | G2a2b2a | | 1541 |
| 2778 | I | 827 | 0 | 0 | 272 | 11 | 0 | M9- | 14 – 19 | | 15655268 | PF3335^ | G2a2b2a | | 2778 |
| 4371 | III | 826 | 0 | 1 | 272 | 11 | 0 | M9- | 14 – 19 | | 21089622 | PF3339 | G2a2b2a | | 4371 |
| 4612 | I | 827 | 0 | 0 | 272 | 11 | 0 | M9- | 14 – 19 | | 21645348 | Z765 | G2a2b2a | | 4612 |
| 4711 | I | 827 | 0 | 0 | 272 | 11 | 0 | M9- | 14 – 19 | | 21764212 | Z766 | G2a2b2a | | 4711 |
| 4892 | I | 827 | 0 | 0 | 272 | 11 | 0 | M9- | 14 – 19 | | 21935550 | Z3243 | G2a2b2a | | 4892 |
| 4893 | I | 827 | 0 | 0 | 272 | 11 | 0 | M9- | 14 – 19 | | 21935606 | PF3342 | G2a2b2a | | 4893 |
| 3734 | III | 798 | 0 | 29 | 259 | 10 | 14 | M9- | 14 – 19 | MD | 18023391 | Z764 | G2a2b2a1 | | 3734 |
| 5724 | I | 827 | 0 | 0 | 272 | 11 | 0 | M9- | 14 – 19 | | 28741231 | PF3347 | G2a2b2a1 | | 5724 |
| 5690 | III | 824 | 0 | 3 | 273 | 10 | 0 | M9- | 15 – 19 | | 24498136 | PF3346 | G2a2b2a1a | | 5690 |
| 5540 | I | 827 | 0 | 0 | 273 | 10 | 0 | M9- | 15 – 19 | | 23640536 | PF3345 | G2a2b2a1a1 | | 5540 |
| 735 | III | 826 | 0 | 1 | 282 | 1 | 0 | M9- | 16 | | 7077897 | Z1993 | G2a2b2a1a1a1a | | 735 |
| 2227 | II | 0 | 827 | 0 | 1 | 282 | 0 | M9- | 16 | | 14698928 | U13 | G2a2b2a1a1a1a | | 2227 |
| 2318 | II | 0 | 827 | 0 | 1 | 282 | 0 | M9- | 16 | | 14871976 | M527 | G2a2b2a1a1a1a | | 2318 |
| 508 | II | 827 | 0 | 0 | 282 | 1 | 0 | M9- | 16 | | 6738741 | Z1991 | G2a2b2a1a1a1a1 | | 508 |
| 4139 | II | 827 | 0 | 0 | 282 | 1 | 0 | M9- | 16 | | 19134681 | Z2022 | G2a2b2a1a1a1a1 | | 4139 |
| 1262 | I | 827 | 0 | 0 | 276 | 7 | 0 | M9- | 17 – 18 | | 8302276 | Z728 | G2a2b2a1a1b | | 1262 |
| 2010 | I | 827 | 0 | 0 | 276 | 7 | 0 | M9- | 17 – 18 | | 14116041 | Z738 | G2a2b2a1a1b | | 2010 |
| 3404 | I | 827 | 0 | 0 | 276 | 7 | 0 | M9- | 17 – 18 | | 17295306 | Z748 | G2a2b2a1a1b | | 3404 |
| 3457 | I | 827 | 0 | 0 | 276 | 7 | 0 | M9- | 17 – 18 | | 17423320 | S317 | G2a2b2a1a1b | | 3457 |
| 4099 | I | 827 | 0 | 0 | 276 | 7 | 0 | M9- | 17 – 18 | | 19047630 | Z1815 | G2a2b2a1a1b1 | | 4099 |
| 5321 | III | 827 | 0 | 0 | 277 | 4 | 2 | M9- | | PI | 23080197 | Z759 | G2a2b2a1a1b1a | | 5321 |
| 3705 | IV | 827 | 0 | 0 | 283 | 0 | 0 | Mono | Mono | | 17937365 | S147 | G2a2b2a1a1b1a1a1 | | 3705 |
| 3135 | IV | 827 | 0 | 0 | 283 | 0 | 0 | Mono | Mono | | 16660759 | S146 | G2a2b2a1a1b1a1a1a | | 3135 |
| 3647 | IV | 827 | 0 | 0 | 283 | 0 | 0 | Mono | Mono | | 17833928 | F2610.2 | G2a2b2a1a1b1a1a1a1a4~ | A | 3647 |
| 3427 | IV | 827 | 0 | 0 | 283 | 0 | 0 | Mono | Mono | | 17353144 | YSC0000033 | G2a2b2a1a1b1a1a1a1b | | 3427 |
| 3339 | I | 827 | 0 | 0 | 279 | 4 | 0 | M9- | 18 | | 17111777 | Z726 | G2a2b2a1a1b1a1a2 | | 3339 |
| 2860 | I | 827 | 0 | 0 | 279 | 4 | 0 | M9- | 18 | | 15833180 | CTS4803 | G2a2b2a1a1b1a1a2a | | 2860 |
| 5045 | I | 827 | 0 | 0 | 279 | 4 | 0 | M9- | 18 | | 22539461 | S2808 | G2a2b2a1a1b1a1a2a1 | | 5045 |
| 3503 | IV | 827 | 0 | 0 | 283 | 0 | 0 | Mono | Mono | | 17533255 | M2226.2 | G2a2b2a1a1b1a1a2a1a1b1e | A | 3503 |
| 5526 | IV | 827 | 0 | 0 | 283 | 0 | 0 | Mono | Mono | | 23592051 | F3484.2 | G2a2b2a1a1b1a1a2a2a1 | | 5526 |
| 1570 | IV | 826 | 0 | 1 | 283 | 0 | 0 | Mono | Mono | | 8775178 | P293.2 | G2a2b2a1a1b1a1a2d~ | A | 1570 |
| 2167 | IV | 827 | 0 | 0 | 283 | 0 | 0 | Mono | Mono | | 14496103 | Page11 | G2a2b2a1a1b3~ | | 2167 |
| 490 | II | 827 | 0 | 0 | 282 | 1 | 0 | M9- | 19 | | 6700171 | CTS342 | G2a2b2a1a1c | | 490 |
| 3749 | IV | 827 | 0 | 0 | 283 | 0 | 0 | Mono | Mono | | 18043703 | PF6867 | G2a2b2a1a1c1 | | 3749 |
| 2787 | IV | 827 | 0 | 0 | 283 | 0 | 0 | Mono | Mono | | 15663626 | CTS4472 | G2a2b2a1a1c1a | | 2787 |
| 55 | IV | 827 | 0 | 0 | 283 | 0 | 0 | Mono | Mono | | 2714659 | CTS77 | G2a2b2a1a1c1a1 | | 55 |
| 148 | IV | 827 | 0 | 0 | 282 | 0 | 1 | Mono | Mono | | 2843538 | Z3038 | G2a2b2a1a1c1a1 | | 148 |
| 3116 | IV | 827 | 0 | 0 | 283 | 0 | 0 | Mono | Mono | | 16596946 | Z1903 | G2a2b2a1a1c1a1 | | 3116 |

| | | | M9+ | | | M9- | | | | | Build 37 | | ISOGG | | |
|---|---|---|---|---|---|---|---|---|---|---|---|---|---|---|---|
| List | Stage | 0 | 1 | . | 0 | 1 | . | M9? | Clade(s) | Prob. | Position | Name | Subgroup Name | Amb. | List |
| 2014 | IV | 827 | 0 | 0 | 283 | 0 | 0 | Mono | Mono | | 14125529 | CTS1934 | G2a2b2a1a1c1a1a1 | | 2014 |
| 2359 | IV | 827 | 0 | 0 | 283 | 0 | 0 | Mono | Mono | | 14912952 | CTS3426 | G2a2b2a1a1c1a1a1 | | 2359 |
| 3849 | IV | 827 | 0 | 0 | 283 | 0 | 0 | Mono | Mono | | 18393688 | L640 | G2a2b2a1a1c1a1a1 | | 3849 |
| 3344 | IV | 827 | 0 | 0 | 283 | 0 | 0 | Mono | Mono | | 17119057 | M4839.2 | G2a2b2a1a1c1a1a2a1a1a1c~ | A | 3344 |
| 1006 | II | 827 | 0 | 0 | 282 | 1 | 0 | M9- | 19 | | 7738468 | FGC12126 | G2a2b2a1a1c2 | | 1006 |
| 1199 | II | 827 | 0 | 0 | 282 | 1 | 0 | M9- | 19 | | 8190187 | FGC12129 | G2a2b2a1a1c2 | | 1199 |
| 1206 | II | 827 | 0 | 0 | 282 | 1 | 0 | M9- | 19 | | 8204404 | FGC12130 | G2a2b2a1a1c2 | | 1206 |
| 1418 | II | 827 | 0 | 0 | 282 | 1 | 0 | M9- | 19 | | 8509294 | FGC12135 | G2a2b2a1a1c2 | | 1418 |
| 17 | II | 827 | 0 | 0 | 281 | 2 | 0 | M9- | 20 | | 2659191 | PF3348 | G2a2b2b | | 17 |
| 475 | II | 827 | 0 | 0 | 281 | 2 | 0 | M9- | 20 | | 6668066 | PF3354 | G2a2b2b | | 475 |
| 1331 | III | 827 | 0 | 0 | 281 | 1 | 1 | M9- | 20 | MD | 8429419 | PF3362 | G2a2b2b1 | | 1331 |
| 5121 | II | 827 | 0 | 0 | 281 | 2 | 0 | M9- | 20 | | 22721202 | PF3430.2 | G2a2b2b1 | A | 5121 |
| 802 | II | 827 | 0 | 0 | 281 | 2 | 0 | M9- | 20 | | 7227975 | PF3356 | G2a2b2b1a1 | | 802 |
| 1121 | II | 827 | 0 | 0 | 281 | 2 | 0 | M9- | 20 | | 7987577 | F1079 | G2a2b2b1a1 | | 1121 |
| 1481 | II | 826 | 1 | 0 | 281 | 2 | 0 | BOTH | | PI | 8599422 | F1338 | G2a2b2b1a1 | | 1481 |
| 3752 | IV | 827 | 0 | 0 | 283 | 0 | 0 | Mono | Mono | | 18047373 | F2686.2 | G2a2b2b1a1b1~ | A | 3752 |
| 1630 | IV | 826 | 0 | 1 | 283 | 0 | 0 | Mono | Mono | | 9003725 | FGC52610 | G2a2b2b1b | | 1630 |
| 833 | IV | 827 | 0 | 0 | 283 | 0 | 0 | Mono | Mono | | 7322981 | M3115 | G2b | | 833 |
| 2336 | IV | 827 | 0 | 0 | 283 | 0 | 0 | Mono | Mono | | 14894401 | M3145 | G2b | | 2336 |
| 4531 | IV | 827 | 0 | 0 | 282 | 0 | 1 | Mono | Mono | | 21480778 | M3191 | G2b | | 4531 |
| 1372 | IV | 827 | 0 | 0 | 283 | 0 | 0 | Mono | Mono | | 8467136 | L183 | G2b1 | | 1372 |
| 2467 | IV | 827 | 0 | 0 | 283 | 0 | 0 | Mono | Mono | | 15027433 | M377 | G2b1 | | 2467 |
| 5381 | IV | 827 | 0 | 0 | 283 | 0 | 0 | Mono | Mono | | 23243887 | S315 | G2b1 | | 5381 |
| 2484 | IV | 827 | 0 | 0 | 283 | 0 | 0 | Mono | Mono | | 15031385 | M283 | G2b1a1 | | 2484 |
| 4758 | IV | 827 | 0 | 0 | 283 | 0 | 0 | Mono | Mono | | 21835021 | Y107627 | G2b2b | | 4758 |
| 2067 | III | 827 | 0 | 0 | 245 | 36 | 2 | M9- | 1 – 10 | MD | 14237670 | PF2657 | GHIJK | | 2067 |
| 569 | II | 827 | 0 | 0 | 282 | 1 | 0 | M9- | 21 | | 6855809 | Z4171 | H | | 569 |
| 1573 | IV | 819 | 0 | 8 | 280 | 0 | 3 | Mono | Mono | MD | 8781791 | Z13964 | H | | 1573 |
| 1722 | III | 820 | 0 | 7 | 281 | 1 | 1 | M9- | 21 | 1H | 9778243 | Z4205 | H | | 1722 |
| 3479 | II | 827 | 0 | 0 | 282 | 1 | 0 | M9- | 21 | | 17476211 | Z4265 | H | | 3479 |
| 3628 | II | 827 | 0 | 0 | 282 | 1 | 0 | M9- | 21 | | 17800761 | Z4273 | H | | 3628 |
| 3653 | III | 826 | 0 | 1 | 282 | 1 | 0 | M9- | 21 | | 17844304 | M2939 | H | | 3653 |
| 3676 | III | 825 | 0 | 2 | 282 | 1 | 0 | M9- | 21 | | 17887908 | Z4276 | H | | 3676 |
| 3708 | III | 825 | 0 | 2 | 281 | 1 | 1 | M9- | 21 | | 17941203 | Z4278 | H | | 3708 |
| 3820 | II | 827 | 0 | 0 | 282 | 1 | 0 | M9- | 21 | | 18182848 | Z4282 | H | | 3820 |
| 3981 | II | 827 | 0 | 0 | 282 | 1 | 0 | M9- | 21 | | 18788668 | Z4292 | H | | 3981 |
| 4318 | III | 826 | 0 | 1 | 282 | 1 | 0 | M9- | 21 | | 19535440 | Z4307 | H | | 4318 |
| 4325 | II | 827 | 0 | 0 | 282 | 1 | 0 | M9- | 21 | | 19563853 | Z4309 | H | | 4325 |
| 5111 | III | 824 | 0 | 3 | 282 | 1 | 0 | M9- | 21 | | 22701876 | Z4336 | H | | 5111 |
| 5262 | III | 826 | 0 | 1 | 281 | 1 | 1 | M9- | 21 | | 22951796 | Z4338 | H | | 5262 |
| 5296 | II | 827 | 0 | 0 | 282 | 1 | 0 | M9- | 21 | | 23033176 | Z4340 | H | A | 5296 |
| 2432 | II | 827 | 0 | 0 | 282 | 1 | 0 | M9- | 21 | | 15006597 | Z4221 | H~ | | 2432 |
| 5451 | III | 822 | 0 | 5 | 282 | 1 | 0 | M9- | 21 | | 23395750 | Z4354 | H~ | | 5451 |
| 4833 | II | 827 | 0 | 0 | 282 | 1 | 0 | M9- | 21 | | 21894058 | Page45 | H1a | | 4833 |
| 4697 | II | 827 | 0 | 0 | 282 | 1 | 0 | M9- | 21 | | 21753199 | M52 | H1a1 | | 4697 |
| 4074 | II | 827 | 0 | 0 | 282 | 1 | 0 | M9- | 21 | | 18979731 | Z4394 | H1a1a | | 4074 |
| 5497 | II | 827 | 0 | 0 | 282 | 1 | 0 | M9- | 21 | | 23501491 | M3079 | H1a1a | | 5497 |
| 4364 | IV | 827 | 0 | 0 | 282 | 0 | 1 | Mono | Mono | | 21071131 | Z14668 | H1a1a1 | | 4364 |
| 2461 | IV | 827 | 0 | 0 | 283 | 0 | 0 | Mono | Mono | | 15025598 | M197 | H1a1a1a | | 2461 |
| 4690 | IV | 827 | 0 | 0 | 283 | 0 | 0 | Mono | Mono | | 21751009 | M36 | H1a1a1a | | 4690 |
| 4732 | IV | 827 | 0 | 0 | 283 | 0 | 0 | Mono | Mono | | 21778713 | M97 | H1a1a2~ | | 4732 |
| 4644 | IV | 825 | 0 | 2 | 283 | 0 | 0 | Mono | Mono | | 21712855 | M138 | H1a1a3~ | | 4644 |
| 1318 | II | 827 | 0 | 0 | 282 | 1 | 0 | M9- | 21 | | 8416762 | Z4367 | H1a1a4 | | 1318 |
| 115 | III | 821 | 0 | 6 | 281 | 1 | 1 | M9- | 21 | | 2806920 | Z4361 | H1a1a4b | | 115 |
| 468 | II | 827 | 0 | 0 | 282 | 1 | 0 | M9- | 21 | | 6659608 | Z4362 | H1a1a4b | | 468 |
| 719 | III | 826 | 0 | 1 | 282 | 1 | 0 | M9- | 21 | | 7053185 | M2716 | H1a1a4b | | 719 |
| 905 | II | 827 | 0 | 0 | 282 | 1 | 0 | M9- | 21 | | 7585548 | Z4520 | H1a1a4b | | 905 |
| 1111 | II | 827 | 0 | 0 | 282 | 1 | 0 | M9- | 21 | | 7955599 | Z4531 | H1a1a4b | | 1111 |
| 1126 | IV | 826 | 0 | 1 | 282 | 0 | 1 | Mono | Mono | MD | 8001582 | Z4365 | H1a1a4b | | 1126 |
| 1174 | II | 827 | 0 | 0 | 282 | 1 | 0 | M9- | 21 | | 8123001 | M2737 | H1a1a4b | | 1174 |
| 1253 | II | 827 | 0 | 0 | 282 | 1 | 0 | M9- | 21 | | 8283064 | Z4366 | H1a1a4b | | 1253 |
| 1571 | II | 827 | 0 | 0 | 282 | 1 | 0 | M9- | 21 | | 8776074 | Z4369 | H1a1a4b | | 1571 |
| 1759 | IV | 821 | 0 | 6 | 276 | 0 | 7 | Mono | Mono | MD | 9853754 | M2785 | H1a1a4b | | 1759 |
| 1773 | II | 827 | 0 | 0 | 282 | 1 | 0 | M9- | 21 | | 9902767 | Z4371 | H1a1a4b | | 1773 |
| 1965 | III | 788 | 0 | 39 | 270 | 1 | 12 | M9- | 21 | | 14038430 | ZS5493 | H1a1a4b | A | 1965 |
| 2379 | II | 827 | 0 | 0 | 282 | 1 | 0 | M9- | 21 | | 14934778 | Z4374 | H1a1a4b | | 2379 |
| 2433 | III | 827 | 0 | 0 | 281 | 1 | 1 | M9- | 21 | | 15010005 | Z4375 | H1a1a4b | | 2433 |
| 2769 | II | 827 | 0 | 0 | 282 | 1 | 0 | M9- | 21 | | 15624558 | Z4376 | H1a1a4b | | 2769 |
| 2897 | II | 827 | 0 | 0 | 282 | 1 | 0 | M9- | 21 | | 15965479 | Z4378 | H1a1a4b | | 2897 |
| 3037 | III | 807 | 0 | 20 | 268 | 1 | 14 | M9- | 21 | | 16407903 | Z4593 | H1a1a4b | | 3037 |
| 3101 | IV | 819 | 0 | 8 | 278 | 0 | 5 | Mono | Mono | MD | 16556621 | M2880 | H1a1a4b | | 3101 |
| 3102 | II | 827 | 0 | 0 | 282 | 1 | 0 | M9- | 21 | | 16559404 | Z4598 | H1a1a4b | | 3102 |
| 3295 | III | 771 | 0 | 56 | 263 | 1 | 19 | M9- | 21 | | 16956661 | M2897 | H1a1a4b | | 3295 |
| 3333 | II | 827 | 0 | 0 | 282 | 1 | 0 | M9- | 21 | | 17100096 | Z4380 | H1a1a4b | | 3333 |
| 3385 | II | 827 | 0 | 0 | 282 | 1 | 0 | M9- | 21 | | 17242063 | Z4381 | H1a1a4b | | 3385 |
| 3467 | III | 826 | 0 | 1 | 281 | 1 | 1 | M9- | 21 | | 17447668 | M2918 | H1a1a4b | | 3467 |
| 3584 | II | 827 | 0 | 0 | 282 | 1 | 0 | M9- | 21 | | 17709958 | Z4383 | H1a1a4b | | 3584 |
| 3655 | III | 827 | 0 | 0 | 281 | 1 | 1 | M9- | 21 | | 17846704 | Z4384 | H1a1a4b | | 3655 |
| 3773 | III | 825 | 0 | 2 | 282 | 1 | 0 | M9- | 21 | | 18077490 | Z4386 | H1a1a4b | | 3773 |
| 3783 | II | 827 | 0 | 0 | 282 | 1 | 0 | M9- | 21 | | 18097518 | Z4387 | H1a1a4b | | 3783 |
| 3856 | II | 827 | 0 | 0 | 282 | 1 | 0 | M9- | 21 | | 18405356 | Z4388 | H1a1a4b | | 3856 |
| 3999 | III | 821 | 0 | 6 | 280 | 1 | 2 | M9- | 21 | | 18823705 | Z4392 | H1a1a4b | | 3999 |
| 4076 | III | 826 | 0 | 1 | 282 | 1 | 0 | M9- | 21 | | 18982181 | Z4635 | H1a1a4b | | 4076 |
| 4336 | II | 827 | 0 | 0 | 282 | 1 | 0 | M9- | 21 | | 20814117 | Z4396 | H1a1a4b | | 4336 |
| 4382 | II | 827 | 0 | 0 | 282 | 1 | 0 | M9- | 21 | | 21115642 | Z4397 | H1a1a4b | | 4382 |
| 4388 | II | 827 | 0 | 0 | 282 | 1 | 0 | M9- | 21 | | 21125953 | Z4398 | H1a1a4b | | 4388 |
| 4472 | II | 827 | 0 | 0 | 282 | 1 | 0 | M9- | 21 | | 21334230 | Z4399 | H1a1a4b | | 4472 |
| 4493 | II | 827 | 0 | 0 | 282 | 1 | 0 | M9- | 21 | | 21388368 | Z4400 | H1a1a4b | | 4493 |
| 4548 | II | 827 | 0 | 0 | 282 | 1 | 0 | M9- | 21 | | 21516489 | M3020 | H1a1a4b | | 4548 |
| 4757 | III | 816 | 0 | 11 | 278 | 1 | 4 | M9- | 21 | | 21832499 | Z4403 | H1a1a4b | | 4757 |
| 4763 | II | 827 | 0 | 0 | 282 | 1 | 0 | M9- | 21 | | 21844401 | Z4404 | H1a1a4b | | 4763 |
| 4873 | II | 827 | 0 | 0 | 282 | 1 | 0 | M9- | 21 | | 21912570 | Z4405 | H1a1a4b | | 4873 |
| 4982 | II | 827 | 0 | 0 | 282 | 1 | 0 | M9- | 21 | | 22167049 | Z4407 | H1a1a4b | | 4982 |
| 5049 | III | 825 | 0 | 2 | 282 | 1 | 0 | M9- | 21 | | 22557391 | Z4408 | H1a1a4b | | 5049 |
| 5070 | II | 827 | 0 | 0 | 282 | 1 | 0 | M9- | 21 | | 22588305 | Z4409 | H1a1a4b | | 5070 |
| 5422 | IV | 798 | 0 | 29 | 277 | 0 | 6 | Mono | Mono | MD | 23315541 | Z4411 | H1a1a4b | | 5422 |

| | | | M9+ | | | M9- | | | | | Build 37 | | ISOGG | | |
|---|---|---|---|---|---|---|---|---|---|---|---|---|---|---|---|
| List | Stage | 0 | 1 | . | 0 | 1 | . | M9? | Clade(s) | Prob. | Position | Name | Subgroup Name | Amb. | List |
| 2820 | IV | 813 | 0 | 14 | 280 | 0 | 3 | Mono | Mono | MD | 15748250 | M2853 | H1a1a4b2 | | 2820 |
| 4021 | II | 827 | 0 | 0 | 282 | 1 | 0 | M9- | 21 | | 18881280 | Z5875 | H1a1a4b2 | | 4021 |
| 1137 | IV | 827 | 0 | 0 | 282 | 0 | 1 | Mono | Mono | | 8030142 | Z14448 | H1a1a4b2a | | 1137 |
| 2918 | IV | 825 | 0 | 2 | 283 | 0 | 0 | Mono | Mono | | 16051872 | Z14449 | H1a1a4b2a | | 2918 |
| 3630 | IV | 827 | 0 | 0 | 283 | 0 | 0 | Mono | Mono | | 17802255 | Z5876 | H1a1a4b2a | | 3630 |
| 4380 | IV | 827 | 0 | 0 | 283 | 0 | 0 | Mono | Mono | | 21107320 | Z14450 | H1a1a4b2a | | 4380 |
| 5412 | IV | 827 | 0 | 0 | 283 | 0 | 0 | Mono | Mono | | 23284888 | Z14452 | H1a1a4b2a | | 5412 |
| 5694 | IV | 802 | 0 | 25 | 272 | 0 | 11 | Mono | Mono | | 24511005 | Z14453 | H1a1a4b2a | | 5694 |
| 1981 | IV | 826 | 0 | 1 | 283 | 0 | 0 | Mono | Mono | | 14064565 | Z14456 | H1a1a4b2b | | 1981 |
| 2154 | IV | 827 | 0 | 0 | 283 | 0 | 0 | Mono | Mono | | 14472817 | Z5878 | H1a1a4b2b | | 2154 |
| 4935 | IV | 827 | 0 | 0 | 283 | 0 | 0 | Mono | Mono | | 22046222 | M3038 | H1a1a4b2c | | 4935 |
| 1616 | IV | 826 | 0 | 1 | 283 | 0 | 0 | Mono | Mono | | 8894070 | Z14441 | H1a1a4b2c1 | | 1616 |
| 3505 | IV | 827 | 0 | 0 | 283 | 0 | 0 | Mono | Mono | | 17537444 | Z5882 | H1a1a4b2c1a | | 3505 |
| 3557 | IV | 827 | 0 | 0 | 283 | 0 | 0 | Mono | Mono | | 17654130 | Z14436 | H1a1a4b2c1a | | 3557 |
| 3732 | IV | 827 | 0 | 0 | 283 | 0 | 0 | Mono | Mono | | 18021446 | Z14437 | H1a1a4b2c1a | | 3732 |
| 3659 | IV | 827 | 0 | 0 | 283 | 0 | 0 | Mono | Mono | | 17855632 | Z5883 | H1a1a4b2c1a1 | | 3659 |
| 482 | IV | 827 | 0 | 0 | 283 | 0 | 0 | Mono | Mono | | 6680599 | Z4507 | H1a1a4b3 | | 482 |
| 4391 | IV | 827 | 0 | 0 | 283 | 0 | 0 | Mono | Mono | | 21143013 | Z4654 | H1a1a4b3 | | 4391 |
| 5407 | IV | 827 | 0 | 0 | 282 | 0 | 1 | Mono | Mono | | 23272722 | Z4695 | H1a1a4b3 | | 5407 |
| 753 | IV | 827 | 0 | 0 | 283 | 0 | 0 | Mono | Mono | | 7135731 | Z12552 | H1a1a4b3a | | 753 |
| 2609 | IV | 823 | 0 | 4 | 283 | 0 | 0 | Mono | Mono | | 15382783 | Z5888 | H1a1a4b3b | | 2609 |
| 5734 | IV | 824 | 0 | 3 | 283 | 0 | 0 | Mono | Mono | | 28789926 | Z12536 | H1a1a4b3b1 | | 5734 |
| 1693 | IV | 827 | 0 | 0 | 283 | 0 | 0 | Mono | Mono | | 9460224 | Z12534 | H1a1a4b3b1a | | 1693 |
| 4050 | IV | 827 | 0 | 0 | 283 | 0 | 0 | Mono | Mono | | 18927804 | Z5890 | H1a1a4b3b1a | | 4050 |
| 4204 | IV | 812 | 0 | 15 | 275 | 0 | 8 | Mono | Mono | | 19237845 | Z12533 | H1a1a4b3b1a | | 4204 |
| 2488 | IV | 827 | 0 | 0 | 283 | 0 | 0 | Mono | Mono | | 15039955 | V2778.2 | H1a1a4b3b1a7~ | A | 2488 |
| 158 | IV | 827 | 0 | 0 | 283 | 0 | 0 | Mono | Mono | | 2863725 | Z4489 | H1a1a4b4 | | 158 |
| 1000 | IV | 827 | 0 | 0 | 283 | 0 | 0 | Mono | Mono | | 7732481 | Z4525 | H1a1a4b4 | | 1000 |
| 1032 | IV | 827 | 0 | 0 | 283 | 0 | 0 | Mono | Mono | | 7792570 | Z4526 | H1a1a4b4 | | 1032 |
| 4932 | IV | 827 | 0 | 0 | 283 | 0 | 0 | Mono | Mono | | 22035997 | Z4469 | H1a1b | | 4932 |
| 1051 | IV | 826 | 0 | 1 | 283 | 0 | 0 | Mono | Mono | | 7831645 | Z4422 | H1a1b1 | | 1051 |
| 3461 | IV | 827 | 0 | 0 | 283 | 0 | 0 | Mono | Mono | | 17427783 | Z4451 | H1a1b1 | | 3461 |
| 5090 | IV | 827 | 0 | 0 | 282 | 0 | 1 | Mono | Mono | | 22646343 | Z4475 | H1a1b1 | | 5090 |
| 560 | IV | 827 | 0 | 0 | 283 | 0 | 0 | Mono | Mono | | 6849552 | Z4414 | H1a1b1a | | 560 |
| 657 | II | 826 | 1 | 0 | 283 | 0 | 0 | M9+ | NU | | 6938119 | Page75 | H1a1b1a | W | 657 |
| 834 | IV | 827 | 0 | 0 | 283 | 0 | 0 | Mono | Mono | | 7324263 | Z4417 | H1a1b1a | | 834 |
| 188 | IV | 826 | 0 | 1 | 283 | 0 | 0 | Mono | Mono | | 2893225 | Z13968 | H1a2a | | 188 |
| 717 | IV | 827 | 0 | 0 | 283 | 0 | 0 | Mono | Mono | | 7049177 | Z14001 | H1a2a | | 717 |
| 970 | IV | 827 | 0 | 0 | 283 | 0 | 0 | Mono | Mono | | 7683191 | Z14004 | H1a2a | | 970 |
| 1509 | IV | 826 | 0 | 1 | 283 | 0 | 0 | Mono | Mono | | 8635099 | Z14021 | H1a2a | | 1509 |
| 2055 | IV | 827 | 0 | 0 | 283 | 0 | 0 | Mono | Mono | | 14209170 | Z14055 | H1a2a | | 2055 |
| 2632 | IV | 811 | 0 | 16 | 279 | 0 | 4 | Mono | Mono | | 15436419 | Z14068 | H1a2a | | 2632 |
| 4038 | IV | 827 | 0 | 0 | 283 | 0 | 0 | Mono | Mono | | 18912198 | Z14120 | H1a2a | | 4038 |
| 5490 | IV | 827 | 0 | 0 | 283 | 0 | 0 | Mono | Mono | | 23496331 | Apt | H1a2a | | 5490 |
| 5618 | IV | 827 | 0 | 0 | 283 | 0 | 0 | Mono | Mono | | 23968298 | Z14191 | H1a2a | | 5618 |
| 37 | IV | 827 | 0 | 0 | 283 | 0 | 0 | Mono | Mono | | 2697625 | Z14258 | H1a2b | | 37 |
| 608 | IV | 825 | 0 | 2 | 282 | 0 | 1 | Mono | Mono | | 6901411 | Z14316 | H1a2b1a | | 608 |
| 4712 | IV | 823 | 0 | 4 | 283 | 0 | 0 | Mono | Mono | | 21764431 | M282 | H2 | | 4712 |
| 94 | IV | 814 | 0 | 13 | 277 | 0 | 6 | Mono | Mono | | 2759285 | Z5857 | H3 | | 94 |
| 142 | IV | 827 | 0 | 0 | 283 | 0 | 0 | Mono | Mono | | 2837192 | Z13449 | H3 | | 142 |
| 166 | IV | 827 | 0 | 0 | 283 | 0 | 0 | Mono | Mono | | 2873167 | Z13450 | H3 | | 166 |
| 682 | IV | 827 | 0 | 0 | 283 | 0 | 0 | Mono | Mono | | 6958386 | Z13474 | H3 | | 682 |
| 687 | IV | 827 | 0 | 0 | 283 | 0 | 0 | Mono | Mono | | 6964855 | Z13475 | H3 | | 687 |
| 695 | IV | 825 | 0 | 2 | 283 | 0 | 0 | Mono | Mono | | 6983363 | Z13476 | H3 | | 695 |
| 1225 | IV | 759 | 0 | 68 | 259 | 0 | 24 | Mono | Mono | | 8238824 | Z13487 | H3 | | 1225 |
| 1260 | IV | 827 | 0 | 0 | 283 | 0 | 0 | Mono | Mono | | 8291990 | S7655.2 | H3 | | 1260 |
| 1338 | IV | 827 | 0 | 0 | 283 | 0 | 0 | Mono | Mono | | 8435392 | Z13488 | H3 | | 1338 |
| 1443 | IV | 827 | 0 | 0 | 283 | 0 | 0 | Mono | Mono | | 8550637 | Z13489 | H3 | | 1443 |
| 1463 | IV | 826 | 0 | 1 | 283 | 0 | 0 | Mono | Mono | | 8576648 | Z13490 | H3 | | 1463 |
| 1471 | IV | 827 | 0 | 0 | 283 | 0 | 0 | Mono | Mono | | 8589767 | Z13491 | H3 | | 1471 |
| 1473 | IV | 827 | 0 | 0 | 283 | 0 | 0 | Mono | Mono | | 8590668 | Z13492 | H3 | | 1473 |
| 1479 | IV | 827 | 0 | 0 | 282 | 0 | 1 | Mono | Mono | | 8595552 | Z13493 | H3 | | 1479 |
| 1491 | IV | 827 | 0 | 0 | 283 | 0 | 0 | Mono | Mono | | 8610259 | Z13494 | H3 | | 1491 |
| 1585 | IV | 826 | 0 | 1 | 283 | 0 | 0 | Mono | Mono | | 8808439 | Z13498 | H3 | | 1585 |
| 1591 | IV | 827 | 0 | 0 | 283 | 0 | 0 | Mono | Mono | | 8828997 | Z13501 | H3 | | 1591 |
| 1593 | IV | 827 | 0 | 0 | 283 | 0 | 0 | Mono | Mono | | 8839116 | Z13502 | H3 | | 1593 |
| 1687 | IV | 826 | 0 | 1 | 283 | 0 | 0 | Mono | Mono | | 9418033 | Z13505 | H3 | | 1687 |
| 1721 | IV | 827 | 0 | 0 | 283 | 0 | 0 | Mono | Mono | | 9773377 | Z13507 | H3 | | 1721 |
| 1752 | IV | 827 | 0 | 0 | 283 | 0 | 0 | Mono | Mono | | 9847849 | Z13508 | H3 | | 1752 |
| 1771 | IV | 827 | 0 | 0 | 282 | 0 | 1 | Mono | Mono | | 9893934 | Z13509 | H3 | | 1771 |
| 1865 | IV | 811 | 0 | 16 | 276 | 0 | 7 | Mono | Mono | | 13668337 | Z13522 | H3 | | 1865 |
| 1921 | IV | 827 | 0 | 0 | 283 | 0 | 0 | Mono | Mono | | 13849996 | Z13526 | H3 | | 1921 |
| 2174 | IV | 799 | 0 | 28 | 270 | 0 | 13 | Mono | Mono | | 14503344 | Z13535 | H3 | | 2174 |
| 2264 | IV | 826 | 0 | 1 | 283 | 0 | 0 | Mono | Mono | | 14791866 | Z13536 | H3 | | 2264 |
| 2378 | IV | 827 | 0 | 0 | 283 | 0 | 0 | Mono | Mono | | 14934195 | Y11987 | H3 | | 2378 |
| 2415 | IV | 827 | 0 | 0 | 283 | 0 | 0 | Mono | Mono | | 14976485 | Z13538 | H3 | | 2415 |
| 2549 | IV | 827 | 0 | 0 | 283 | 0 | 0 | Mono | Mono | | 15207006 | Z13540 | H3 | | 2549 |
| 2793 | IV | 827 | 0 | 0 | 283 | 0 | 0 | Mono | Mono | | 15681064 | Z13543 | H3 | | 2793 |
| 3013 | IV | 827 | 0 | 0 | 283 | 0 | 0 | Mono | Mono | | 16363476 | Z13546 | H3 | | 3013 |
| 3138 | IV | 827 | 0 | 0 | 283 | 0 | 0 | Mono | Mono | | 16669102 | Z13549 | H3 | | 3138 |
| 3162 | IV | 827 | 0 | 0 | 283 | 0 | 0 | Mono | Mono | | 16749652 | Z13550 | H3 | | 3162 |
| 3401 | IV | 827 | 0 | 0 | 283 | 0 | 0 | Mono | Mono | | 17288291 | Z13553 | H3 | | 3401 |
| 3507 | IV | 827 | 0 | 0 | 283 | 0 | 0 | Mono | Mono | | 17544312 | Z13554 | H3 | | 3507 |
| 3544 | IV | 827 | 0 | 0 | 283 | 0 | 0 | Mono | Mono | | 17606413 | Z13555 | H3 | | 3544 |
| 3572 | IV | 827 | 0 | 0 | 283 | 0 | 0 | Mono | Mono | | 17677193 | Z13556 | H3 | | 3572 |
| 3575 | IV | 827 | 0 | 0 | 283 | 0 | 0 | Mono | Mono | | 17681150 | Z13557 | H3 | | 3575 |
| 3644 | IV | 827 | 0 | 0 | 283 | 0 | 0 | Mono | Mono | | 17821170 | Z13560 | H3 | | 3644 |
| 3787 | IV | 507 | 0 | 320 | 166 | 0 | 117 | Mono | Mono | | 18105101 | Z13563 | H3 | | 3787 |
| 3862 | IV | 827 | 0 | 0 | 283 | 0 | 0 | Mono | Mono | | 18422011 | Z13566 | H3 | | 3862 |
| 3978 | IV | 826 | 0 | 1 | 283 | 0 | 0 | Mono | Mono | | 18786572 | Z13568 | H3 | | 3978 |
| 3994 | IV | 827 | 0 | 0 | 283 | 0 | 0 | Mono | Mono | | 18817532 | Z13571 | H3 | | 3994 |
| 4114 | IV | 827 | 0 | 0 | 283 | 0 | 0 | Mono | Mono | | 19076816 | Z13575 | H3 | | 4114 |
| 4177 | IV | 827 | 0 | 0 | 283 | 0 | 0 | Mono | Mono | | 19200004 | Z13577 | H3 | | 4177 |
| 4199 | IV | 827 | 0 | 0 | 283 | 0 | 0 | Mono | Mono | | 19234590 | Z13578 | H3 | | 4199 |
| 4293 | IV | 827 | 0 | 0 | 283 | 0 | 0 | Mono | Mono | | 19475667 | Z13581 | H3 | | 4293 |
| 4408 | IV | 826 | 0 | 1 | 283 | 0 | 0 | Mono | Mono | | 21190678 | Z13586 | H3 | | 4408 |

| | | M9+ | | | M9- | | | | | | Build 37 | | ISOGG | | |
|---|---|---|---|---|---|---|---|---|---|---|---|---|---|---|---|
| List | Stage | 0 | 1 | . | 0 | 1 | . | M9? | Clade(s) | Prob. | Position | Name | Subgroup Name | Amb. | List |
| 4420 | IV | 827 | 0 | 0 | 283 | 0 | 0 | Mono | Mono | | 21226052 | Z13587 | H3 | | 4420 |
| 4563 | IV | 827 | 0 | 0 | 283 | 0 | 0 | Mono | Mono | | 21564841 | Z13591 | H3 | | 4563 |
| 4731 | IV | 827 | 0 | 0 | 283 | 0 | 0 | Mono | Mono | | 21778710 | Z13595 | H3 | | 4731 |
| 4802 | IV | 640 | 0 | 187 | 224 | 0 | 59 | Mono | Mono | | 21877554 | Z13598 | H3 | | 4802 |
| 4905 | IV | 825 | 0 | 2 | 283 | 0 | 0 | Mono | Mono | | 21963272 | Z13599 | H3 | | 4905 |
| 4934 | IV | 777 | 0 | 50 | 265 | 0 | 18 | Mono | Mono | | 22044605 | Z13600 | H3 | | 4934 |
| 4937 | IV | 826 | 0 | 1 | 283 | 0 | 0 | Mono | Mono | | 22053153 | Z13601 | H3 | | 4937 |
| 5047 | IV | 827 | 0 | 0 | 283 | 0 | 0 | Mono | Mono | | 22552528 | Z13609 | H3 | | 5047 |
| 5061 | IV | 827 | 0 | 0 | 283 | 0 | 0 | Mono | Mono | | 22573596 | Z13610 | H3 | | 5061 |
| 5087 | IV | 827 | 0 | 0 | 283 | 0 | 0 | Mono | Mono | | 22628301 | Z13612 | H3 | | 5087 |
| 5281 | IV | 826 | 0 | 1 | 283 | 0 | 0 | Mono | Mono | | 23003094 | Z13615 | H3 | | 5281 |
| 5322 | IV | 827 | 0 | 0 | 283 | 0 | 0 | Mono | Mono | | 23081504 | Z16816 | H3 | | 5322 |
| 5345 | IV | 827 | 0 | 0 | 283 | 0 | 0 | Mono | Mono | | 23134639 | Z13618 | H3 | | 5345 |
| 5443 | IV | 827 | 0 | 0 | 283 | 0 | 0 | Mono | Mono | | 23387399 | Z13621 | H3 | | 5443 |
| 5458 | IV | 827 | 0 | 0 | 283 | 0 | 0 | Mono | Mono | | 23408120 | Z13622 | H3 | | 5458 |
| 5471 | IV | 827 | 0 | 0 | 283 | 0 | 0 | Mono | Mono | | 23451560 | Z13623 | H3 | | 5471 |
| 5474 | IV | 827 | 0 | 0 | 283 | 0 | 0 | Mono | Mono | | 23456778 | Z13624 | H3 | | 5474 |
| 5480 | IV | 827 | 0 | 0 | 283 | 0 | 0 | Mono | Mono | | 23479007 | Z13625 | H3 | | 5480 |
| 5561 | IV | 827 | 0 | 0 | 282 | 0 | 1 | Mono | Mono | | 23772822 | Z13626 | H3 | | 5561 |
| 5667 | IV | 824 | 0 | 3 | 281 | 0 | 2 | Mono | Mono | | 24413499 | Z13627 | H3 | | 5667 |
| 1544 | IV | 827 | 0 | 0 | 283 | 0 | 0 | Mono | Mono | | 8692246 | Z13495 | H3~ | | 1544 |
| 1554 | IV | 826 | 0 | 1 | 282 | 0 | 1 | Mono | Mono | | 8712131 | Z13496 | H3~ | | 1554 |
| 1588 | IV | 827 | 0 | 0 | 283 | 0 | 0 | Mono | Mono | | 8816195 | Z13500 | H3~ | | 1588 |
| 1949 | IV | 827 | 0 | 0 | 283 | 0 | 0 | Mono | Mono | | 13999868 | Z13530 | H3~ | | 1949 |
| 1957 | IV | 827 | 0 | 0 | 282 | 0 | 1 | Mono | Mono | | 14017373 | Z13531 | H3~ | | 1957 |
| 2719 | IV | 827 | 0 | 0 | 283 | 0 | 0 | Mono | Mono | | 15559913 | Z13542 | H3~ | | 2719 |
| 4385 | IV | 827 | 0 | 0 | 283 | 0 | 0 | Mono | Mono | | 21121323 | Z13585 | H3~ | | 4385 |
| 4764 | IV | 827 | 0 | 0 | 283 | 0 | 0 | Mono | Mono | | 21850428 | Z13597 | H3~ | | 4764 |
| 644 | IV | 827 | 0 | 0 | 283 | 0 | 0 | Mono | Mono | | 6932688 | Z5866 | H3a | A | 644 |
| 3911 | IV | 827 | 0 | 0 | 283 | 0 | 0 | Mono | Mono | | 18622326 | Z9469.2 | H3a | | 3911 |
| 4352 | IV | 827 | 0 | 0 | 283 | 0 | 0 | Mono | Mono | | 21043320 | Z5864 | H3a1 | | 4352 |
| 26 | IV | 827 | 0 | 0 | 283 | 0 | 0 | Mono | Mono | | 2667783 | Z5865 | H3a2a | | 26 |
| 986 | IV | 827 | 0 | 0 | 283 | 0 | 0 | Mono | Mono | | 7711848 | Z12646 | H3a2a1 | | 986 |
| 2722 | IV | 827 | 0 | 0 | 283 | 0 | 0 | Mono | Mono | | 15567343 | Z12694 | H3a2a1 | | 2722 |
| 1276 | IV | 827 | 0 | 0 | 282 | 0 | 1 | Mono | Mono | | 8335702 | Z12656 | H3a2a2 | | 1276 |
| 174 | IV | 827 | 0 | 0 | 283 | 0 | 0 | Mono | Mono | | 2878605 | Z13871 | H3b | | 174 |
| 194 | IV | 827 | 0 | 0 | 283 | 0 | 0 | Mono | Mono | | 2913033 | Z13872 | H3b | | 194 |
| 1173 | IV | 827 | 0 | 0 | 283 | 0 | 0 | Mono | Mono | | 8121886 | Z13892 | H3b | | 1173 |
| 1919 | IV | 821 | 0 | 6 | 283 | 0 | 0 | Mono | Mono | | 13846844 | Z13907 | H3b | | 1919 |
| 3462 | IV | 827 | 0 | 0 | 283 | 0 | 0 | Mono | Mono | | 17445387 | Z13925 | H3b | | 3462 |
| 3871 | IV | 827 | 0 | 0 | 283 | 0 | 0 | Mono | Mono | | 18554090 | Z13931 | H3b | | 3871 |
| 4324 | IV | 825 | 0 | 2 | 283 | 0 | 0 | Mono | Mono | | 19555539 | ZS332.2 | H3b | A | 4324 |
| 4708 | IV | 827 | 0 | 0 | 283 | 0 | 0 | Mono | Mono | | 21763370 | Z13937 | H3b | | 4708 |
| 5054 | IV | 826 | 0 | 1 | 283 | 0 | 0 | Mono | Mono | | 22560423 | Z13945 | H3b | | 5054 |
| 5552 | IV | 825 | 0 | 2 | 283 | 0 | 0 | Mono | Mono | | 23754859 | Z13951 | H3b | | 5552 |
| 190 | IV | 822 | 0 | 5 | 282 | 0 | 1 | Mono | Mono | | 2894961 | Z5859 | H3b1 | | 190 |
| 2281 | IV | 827 | 0 | 0 | 283 | 0 | 0 | Mono | Mono | | 14818741 | M8081 | H3b1 | | 2281 |
| 792 | I | 827 | 0 | 0 | 225 | 58 | 0 | M9- | 1 – 20 | | 7202703 | S6378 | HIJK | | 792 |
| 34 | I | 827 | 0 | 0 | 82 | 201 | 0 | M9- | 22 – 59 | | 2688442 | PF3569 | I | | 34 |
| 63 | I | 827 | 0 | 0 | 82 | 201 | 0 | M9- | 22 – 59 | | 2723755 | V1031 | I | | 63 |
| 200 | I | 827 | 0 | 0 | 82 | 201 | 0 | M9- | 22 – 59 | | 2974782 | PF3574 | I | | 200 |
| 622 | III | 782 | 0 | 45 | 80 | 197 | 6 | M9- | 22 – 59 | MD | 6926038 | PF3629 | I | | 622 |
| 665 | I | 827 | 0 | 0 | 82 | 201 | 0 | M9- | 22 – 59 | | 6943522 | PF3630 | I | | 665 |
| 896 | I | 827 | 0 | 0 | 82 | 201 | 0 | M9- | 22 – 59 | | 7570370 | PF3639 | I | | 896 |
| 940 | I | 827 | 0 | 0 | 82 | 201 | 0 | M9- | 22 – 59 | | 7642823 | Y1865 | I | | 940 |
| 966 | I | 827 | 0 | 0 | 82 | 201 | 0 | M9- | 22 – 59 | | 7681156 | PF3640 | I | | 966 |
| 973 | I | 827 | 0 | 0 | 82 | 201 | 0 | M9- | 22 – 59 | | 7688470 | V1369 | I | | 973 |
| 1064 | III | 826 | 0 | 1 | 82 | 201 | 0 | M9- | 22 – 59 | | 7853028 | V1476 | I | | 1064 |
| 1068 | I | 827 | 0 | 0 | 82 | 201 | 0 | M9- | 22 – 59 | | 7856500 | YSC0000280 | I | | 1068 |
| 1144 | I | 827 | 0 | 0 | 82 | 201 | 0 | M9- | 22 – 59 | | 8046731 | V1582 | I | | 1144 |
| 1241 | III | 822 | 0 | 5 | 82 | 201 | 0 | M9- | 22 – 59 | | 8262092 | Y1869 | I | | 1241 |
| 1245 | I | 827 | 0 | 0 | 82 | 201 | 0 | M9- | 22 – 59 | | 8267857 | PF3653 | I | | 1245 |
| 1300 | I | 827 | 0 | 0 | 82 | 201 | 0 | M9- | 22 – 59 | | 8382265 | YSC0000281 | I | | 1300 |
| 1364 | I | 827 | 0 | 0 | 82 | 201 | 0 | M9- | 22 – 59 | | 8465165 | YSC0000283 | I | | 1364 |
| 1367 | I | 827 | 0 | 0 | 82 | 201 | 0 | M9- | 22 – 59 | | 8466652 | V2048 | I | | 1367 |
| 1397 | I | 827 | 0 | 0 | 82 | 201 | 0 | M9- | 22 – 59 | | 8484606 | V2111 | I | | 1397 |
| 1399 | I | 827 | 0 | 0 | 82 | 201 | 0 | M9- | 22 – 59 | | 8485677 | YSC0000284 | I | | 1399 |
| 1432 | I | 827 | 0 | 0 | 82 | 201 | 0 | M9- | 22 – 59 | | 8536868 | YSC0000285 | I | | 1432 |
| 1518 | I | 827 | 0 | 0 | 82 | 201 | 0 | M9- | 22 – 59 | | 8643763 | PF3665 | I | | 1518 |
| 1560 | I | 827 | 0 | 0 | 82 | 201 | 0 | M9- | 22 – 59 | | 8728974 | PF3666 | I | | 1560 |
| 1607 | I | 827 | 0 | 0 | 82 | 201 | 0 | M9- | 22 – 59 | | 8873160 | PF3668 | I | | 1607 |
| 1677 | III | 768 | 0 | 59 | 77 | 200 | 6 | M9- | 22 – 59 | MD | 9376351 | PF3672 | I | | 1677 |
| 1739 | III | 827 | 0 | 0 | 82 | 192 | 9 | M9- | 22 – 59 | MD | 9827411 | YSC0000256.2 | I | A | 1739 |
| 1988 | I | 827 | 0 | 0 | 82 | 201 | 0 | M9- | 22 – 59 | | 14073053 | PF3699 | I | | 1988 |
| 2058 | III | 826 | 0 | 1 | 82 | 201 | 0 | M9- | 22 – 59 | | 14214481 | PF3703 | I | | 2058 |
| 2112 | I | 827 | 0 | 0 | 82 | 201 | 0 | M9- | 22 – 59 | | 14337364 | PF3706 | I | | 2112 |
| 2119 | III | 827 | 0 | 0 | 81 | 201 | 1 | M9- | 22 – 59 | | 14352669 | PF3707 | I | | 2119 |
| 2297 | I | 827 | 0 | 0 | 82 | 201 | 0 | M9- | 22 – 59 | | 14847792 | PF3715 | I | | 2297 |
| 2413 | I | 827 | 0 | 0 | 82 | 201 | 0 | M9- | 22 – 59 | | 14974451 | YSC0000260 | I | | 2413 |
| 2424 | I | 827 | 0 | 0 | 82 | 201 | 0 | M9- | 22 – 59 | | 14986989 | V2740 | I | | 2424 |
| 2456 | I | 827 | 0 | 0 | 82 | 201 | 0 | M9- | 22 – 59 | | 15023364 | PF3721 | I | | 2456 |
| 2512 | I | 827 | 0 | 0 | 82 | 201 | 0 | M9- | 22 – 59 | | 15089989 | PF3722 | I | | 2512 |
| 2606 | I | 827 | 0 | 0 | 82 | 201 | 0 | M9- | 22 – 59 | | 15377802 | PF3725 | I | | 2606 |
| 2612 | I | 827 | 0 | 0 | 82 | 201 | 0 | M9- | 22 – 59 | | 15389836 | PF3868 | I | | 2612 |
| 2670 | I | 827 | 0 | 0 | 82 | 201 | 0 | M9- | 22 – 59 | | 15479899 | V2971 | I | | 2670 |
| 2683 | I | 827 | 0 | 0 | 82 | 201 | 0 | M9- | 22 – 59 | | 15506055 | V2992 | I | A | 2683 |
| 2703 | I | 827 | 0 | 0 | 82 | 201 | 0 | M9- | 22 – 59 | | 15536759 | V3027 | I | | 2703 |
| 2704 | I | 827 | 0 | 0 | 82 | 201 | 0 | M9- | 22 – 59 | | 15536870 | PF3729 | I | | 2704 |
| 2756 | I | 827 | 0 | 0 | 82 | 201 | 0 | M9- | 22 – 59 | | 15595624 | PF3730 | I | | 2756 |
| 2766 | I | 827 | 0 | 0 | 82 | 201 | 0 | M9- | 22 – 59 | | 15615533 | YSC0000263 | I | | 2766 |
| 2825 | I | 827 | 0 | 0 | 82 | 201 | 0 | M9- | 22 – 59 | | 15759200 | CTS4664 | I | | 2825 |
| 2839 | III | 799 | 0 | 28 | 81 | 158 | 44 | M9- | 22 – 59 | MD | 15793946 | PF3734 | I | | 2839 |
| 2840 | I | 827 | 0 | 0 | 82 | 201 | 0 | M9- | 22 – 59 | | 15799074 | PF3735 | I | | 2840 |
| 2843 | I | 826 | 1 | 0 | 82 | 201 | 0 | BOTH | | PI | 15801167 | ZS2268 | I | | 2843 |
| 2867 | I | 827 | 0 | 0 | 82 | 201 | 0 | M9- | 22 – 59 | | 15862842 | V3186 | I | | 2867 |
| 2890 | III | 827 | 0 | 0 | 82 | 200 | 1 | M9- | 22 – 59 | MD | 15937959 | V3251 | I | | 2890 |

| | | | M9+ | | | | M9- | | | | | Build 37 | | ISOGG | | |
|---|---|---|---|---|---|---|---|---|---|---|---|---|---|---|---|---|
| List | Stage | 0 | 1 | . | | 0 | 1 | . | M9? | Clade(s) | Prob. | Position | Name | Subgroup Name | Amb. | List |
| 2913 | I | 827 | 0 | 0 | | 82 | 201 | 0 | M9- | 22 – 59 | | 16039881 | PF3739 | I | | 2913 |
| 3039 | I | 827 | 0 | 0 | | 82 | 201 | 0 | M9- | 22 – 59 | | 16415916 | PF3744 | I | | 3039 |
| 3099 | I | 827 | 0 | 0 | | 82 | 201 | 0 | M9- | 22 – 59 | | 16548548 | PF3747 | I | | 3099 |
| 3105 | III | 824 | 0 | 3 | | 82 | 200 | 1 | M9- | 22 – 59 | MD | 16567253 | PF3748 | I | | 3105 |
| 3164 | I | 827 | 0 | 0 | | 82 | 201 | 0 | M9- | 22 – 59 | | 16751000 | PF3750 | I | | 3164 |
| 3179 | I | 827 | 0 | 0 | | 82 | 201 | 0 | M9- | 22 – 59 | | 16780748 | PF3871 | I | | 3179 |
| 3214 | I | 827 | 0 | 0 | | 82 | 201 | 0 | M9- | 22 – 59 | | 16836079 | PF3753 | I | | 3214 |
| 3279 | I | 827 | 0 | 0 | | 82 | 201 | 0 | M9- | 22 – 59 | | 16939794 | PF3756 | I | | 3279 |
| 3458 | I | 827 | 0 | 0 | | 82 | 201 | 0 | M9- | 22 – 59 | | 17424807 | CTS7329 | I | | 3458 |
| 3493 | I | 827 | 0 | 0 | | 82 | 201 | 0 | M9- | 22 – 59 | | 17511797 | PF3760 | I | | 3493 |
| 3502 | I | 827 | 0 | 0 | | 82 | 201 | 0 | M9- | 22 – 59 | | 17525137 | PF3761 | I | | 3502 |
| 3508 | I | 827 | 0 | 0 | | 82 | 201 | 0 | M9- | 22 – 59 | | 17548890 | PF3763 | I | | 3508 |
| 3578 | I | 827 | 0 | 0 | | 82 | 201 | 0 | M9- | 22 – 59 | | 17692855 | PF3766 | I | | 3578 |
| 3641 | I | 827 | 0 | 0 | | 82 | 201 | 0 | M9- | 22 – 59 | | 17818847 | PF3768 | I | | 3641 |
| 3698 | III | 826 | 0 | 1 | | 82 | 201 | 0 | M9- | 22 – 59 | | 17924382 | PF3770 | I | | 3698 |
| 3706 | I | 827 | 0 | 0 | | 82 | 201 | 0 | M9- | 22 – 59 | | 17940414 | PF3771 | I | | 3706 |
| 3714 | III | 804 | 0 | 23 | | 81 | 201 | 1 | M9- | 22 – 59 | | 17949402 | PF3772 | I | | 3714 |
| 3730 | I | 827 | 0 | 0 | | 82 | 201 | 0 | M9- | 22 – 59 | | 18018313 | PF3773 | I | | 3730 |
| 3775 | III | 826 | 0 | 1 | | 82 | 201 | 0 | M9- | 22 – 59 | | 18078759 | PF3775 | I | | 3775 |
| 3810 | I | 827 | 0 | 0 | | 82 | 201 | 0 | M9- | 22 – 59 | | 18172947 | PF3776 | I | | 3810 |
| 3837 | I | 827 | 0 | 0 | | 82 | 201 | 0 | M9- | 22 – 59 | | 18257568 | PF3778 | I | | 3837 |
| 3852 | I | 827 | 0 | 0 | | 82 | 201 | 0 | M9- | 22 – 59 | | 18394743 | YSC0000291 | I | | 3852 |
| 3855 | I | 827 | 0 | 0 | | 82 | 201 | 0 | M9- | 22 – 59 | | 18404486 | PF3780 | I | | 3855 |
| 3886 | I | 827 | 0 | 0 | | 82 | 201 | 0 | M9- | 22 – 59 | | 18582617 | CTS8963 | I | | 3886 |
| 3977 | I | 827 | 0 | 0 | | 82 | 201 | 0 | M9- | 22 – 59 | | 18786174 | V3496 | I | | 3977 |
| 4049 | I | 827 | 0 | 0 | | 82 | 201 | 0 | M9- | 22 – 59 | | 18927031 | PF3785 | I | | 4049 |
| 4100 | III | 820 | 0 | 7 | | 82 | 201 | 0 | M9- | 22 – 59 | | 19048602 | PF3787 | I | | 4100 |
| 4196 | I | 827 | 0 | 0 | | 82 | 201 | 0 | M9- | 22 – 59 | | 19233673 | CTS10058 | I | | 4196 |
| 4361 | I | 827 | 0 | 0 | | 82 | 201 | 0 | M9- | 22 – 59 | | 21067903 | PF3794 | I | | 4361 |
| 4479 | I | 827 | 0 | 0 | | 82 | 201 | 0 | M9- | 22 – 59 | | 21359407 | L503 | I | | 4479 |
| 4503 | I | 827 | 0 | 0 | | 82 | 201 | 0 | M9- | 22 – 59 | | 21402723 | PF3800 | I | | 4503 |
| 4520 | III | 827 | 0 | 0 | | 82 | 200 | 1 | M9- | 22 – 59 | MD | 21452125 | PF3803 | I | | 4520 |
| 4551 | III | 822 | 0 | 5 | | 82 | 199 | 2 | M9- | 22 – 59 | MD | 21525069 | PF3806 | I | | 4551 |
| 4553 | III | 827 | 0 | 0 | | 82 | 200 | 1 | M9- | 22 – 59 | MD | 21535086 | PF3807 | I | | 4553 |
| 4559 | I | 827 | 0 | 0 | | 82 | 201 | 0 | M9- | 22 – 59 | | 21556106 | PF3809 | I | | 4559 |
| 4598 | I | 827 | 0 | 0 | | 82 | 201 | 0 | M9- | 22 – 59 | | 21627180 | PF3811 | I | | 4598 |
| 4636 | I | 827 | 0 | 0 | | 82 | 201 | 0 | M9- | 22 – 59 | | 21689728 | Y1909 | I | | 4636 |
| 4759 | I | 827 | 0 | 0 | | 82 | 201 | 0 | M9- | 22 – 59 | | 21839183 | PF3814 | I | | 4759 |
| 4760 | I | 827 | 0 | 0 | | 82 | 201 | 0 | M9- | 22 – 59 | | 21841289 | PF3815 | I | | 4760 |
| 4951 | I | 827 | 0 | 0 | | 82 | 201 | 0 | M9- | 22 – 59 | | 22100087 | PF3819 | I | | 4951 |
| 4957 | I | 827 | 0 | 0 | | 82 | 201 | 0 | M9- | 22 – 59 | | 22115103 | YSC0000272 | I | | 4957 |
| 4998 | III | 827 | 0 | 0 | | 82 | 200 | 1 | M9- | 22 – 59 | MD | 22200336 | PF3822 | I | | 4998 |
| 5038 | I | 827 | 0 | 0 | | 82 | 201 | 0 | M9- | 22 – 59 | | 22525421 | PF3836 | I | | 5038 |
| 5062 | I | 827 | 0 | 0 | | 82 | 201 | 0 | M9- | 22 – 59 | | 22573702 | PF3837 | I | | 5062 |
| 5330 | III | 825 | 0 | 2 | | 82 | 201 | 0 | M9- | 22 – 59 | | 23113271 | CTS11441 | I | | 5330 |
| 5354 | I | 827 | 0 | 0 | | 82 | 201 | 0 | M9- | 22 – 59 | | 23154034 | YSC0000298 | I | | 5354 |
| 5355 | I | 827 | 0 | 0 | | 82 | 201 | 0 | M9- | 22 – 59 | | 23156725 | PF3842 | I | | 5355 |
| 5454 | I | 827 | 0 | 0 | | 82 | 201 | 0 | M9- | 22 – 59 | | 23401471 | PF3878 | I | | 5454 |
| 5482 | III | 824 | 0 | 3 | | 82 | 200 | 1 | M9- | 22 – 59 | MD | 23479970 | YSC0000300 | I | | 5482 |
| 659 | IV | 827 | 0 | 0 | | 283 | 0 | 0 | Mono | Mono | | 6938665 | Page76 | I (Notes) | | 659 |
| 2745 | IV | 827 | 0 | 0 | | 283 | 0 | 0 | Mono | Mono | | 15591474 | P259 | I (Notes) | | 2745 |
| 4666 | IV | 827 | 0 | 0 | | 283 | 0 | 0 | Mono | Mono | | 21732339 | M21 | I (Notes) | | 4666 |
| 18 | III | 825 | 0 | 2 | | 187 | 86 | 10 | M9- | 22 – 36 | MD | 2659347 | Z2676 | I1 | | 18 |
| 90 | I | 827 | 0 | 0 | | 187 | 96 | 0 | M9- | 22 – 36 | | 2753908 | Z2679 | I1 | | 90 |
| 121 | III | 826 | 0 | 1 | | 187 | 96 | 0 | M9- | 22 – 36 | | 2812119 | Z2680^ | I1 | | 121 |
| 479 | I | 827 | 0 | 0 | | 187 | 96 | 0 | M9- | 22 – 36 | | 6677619 | Y1863 | I1 | | 479 |
| 484 | I | 827 | 0 | 0 | | 187 | 96 | 0 | M9- | 22 – 36 | | 6681479 | Y1864 | I1 | | 484 |
| 559 | I | 827 | 0 | 0 | | 187 | 96 | 0 | M9- | 22 – 36 | | 6848560 | Z2700 | I1 | | 559 |
| 565 | III | 826 | 0 | 1 | | 187 | 96 | 0 | M9- | 22 – 36 | | 6854053 | Z2701 | I1 | | 565 |
| 620 | I | 827 | 0 | 0 | | 187 | 96 | 0 | M9- | 22 – 36 | | 6923525 | Z2702 | I1 | | 620 |
| 723 | I | 827 | 0 | 0 | | 187 | 96 | 0 | M9- | 22 – 36 | | 7066305 | Z2705 | I1 | | 723 |
| 844 | I | 827 | 0 | 0 | | 187 | 96 | 0 | M9- | 22 – 36 | | 7377905 | Z2708 | I1 | | 844 |
| 949 | I | 827 | 0 | 0 | | 187 | 96 | 0 | M9- | 22 – 36 | | 7652844 | YSC0000278 | I1 | | 949 |
| 1004 | I | 827 | 0 | 0 | | 187 | 96 | 0 | M9- | 22 – 36 | | 7737070 | Z2713^ | I1 | | 1004 |
| 1009 | III | 801 | 0 | 26 | | 180 | 96 | 7 | M9- | 22 – 36 | | 7742116 | Z2714 | I1 | | 1009 |
| 1027 | I | 827 | 0 | 0 | | 187 | 96 | 0 | M9- | 22 – 36 | | 7774013 | Z2717 | I1 | | 1027 |
| 1077 | I | 827 | 0 | 0 | | 187 | 96 | 0 | M9- | 22 – 36 | | 7888014 | Z2719 | I1 | | 1077 |
| 1166 | III | 826 | 0 | 1 | | 186 | 95 | 2 | M9- | 22 – 36 | MD | 8103323 | Z2723 | I1 | | 1166 |
| 1168 | I | 827 | 0 | 0 | | 187 | 96 | 0 | M9- | 22 – 36 | | 8108722 | Y1868 | I1 | | 1168 |
| 1248 | I | 827 | 0 | 0 | | 187 | 96 | 0 | M9- | 22 – 36 | | 8274659 | Z2724 | I1 | | 1248 |
| 1451 | III | 825 | 0 | 2 | | 187 | 94 | 2 | M9- | 22 – 36 | MD | 8563464 | Z2728^ | I1 | | 1451 |
| 1505 | I | 827 | 0 | 0 | | 187 | 96 | 0 | M9- | 22 – 36 | | 8629665 | Z2729 | I1 | A | 1505 |
| 1589 | I | 827 | 0 | 0 | | 187 | 96 | 0 | M9- | 22 – 36 | | 8820460 | Z2730^ | I1 | | 1589 |
| 1667 | III | 822 | 0 | 5 | | 187 | 95 | 1 | M9- | 22 – 36 | MD | 9148209 | Z2733 | I1 | | 1667 |
| 1760 | I | 827 | 0 | 0 | | 187 | 96 | 0 | M9- | 22 – 36 | | 9853969 | Z2737 | I1 | | 1760 |
| 1770 | I | 827 | 0 | 0 | | 187 | 96 | 0 | M9- | 22 – 36 | | 9885854 | Z2738 | I1 | | 1770 |
| 1833 | III | 827 | 0 | 0 | | 185 | 96 | 2 | M9- | 22 – 36 | | 13228969 | Z2742^ | I1 | | 1833 |
| 1836 | III | 827 | 0 | 0 | | 185 | 97 | 1 | M9- | | PI | 13237053 | Z2744 | I1 | | 1836 |
| 1908 | III | 826 | 0 | 1 | | 186 | 96 | 1 | M9- | 22 – 36 | | 13803650 | Y1879 | I1 | | 1908 |
| 1912 | III | 827 | 0 | 0 | | 187 | 95 | 1 | M9- | 22 – 36 | MD | 13822499 | Z2747 | I1 | | 1912 |
| 1970 | I | 827 | 0 | 0 | | 187 | 96 | 0 | M9- | 22 – 36 | | 14047137 | Z2749 | I1 | | 1970 |
| 1973 | I | 827 | 0 | 0 | | 187 | 96 | 0 | M9- | 22 – 36 | | 14049684 | Z2750^ | I1 | | 1973 |
| 1987 | I | 827 | 0 | 0 | | 187 | 96 | 0 | M9- | 22 – 36 | | 14071004 | Z2751 | I1 | | 1987 |
| 2085 | I | 827 | 0 | 0 | | 187 | 96 | 0 | M9- | 22 – 36 | | 14280509 | Z2754^ | I1 | | 2085 |
| 2094 | I | 827 | 0 | 0 | | 187 | 96 | 0 | M9- | 22 – 36 | | 14294544 | Z2755 | I1 | | 2094 |
| 2133 | I | 827 | 0 | 0 | | 187 | 96 | 0 | M9- | 22 – 36 | | 14402997 | Z2757 | I1 | | 2133 |
| 2147 | I | 827 | 0 | 0 | | 187 | 96 | 0 | M9- | 22 – 36 | | 14454653 | Z2759^ | I1 | | 2147 |
| 2170 | I | 827 | 0 | 0 | | 187 | 96 | 0 | M9- | 22 – 36 | | 14496753 | P30 | I1 | | 2170 |
| 2194 | III | 827 | 0 | 0 | | 187 | 95 | 1 | M9- | 22 – 36 | MD | 14596460 | Z2760 | I1 | | 2194 |
| 2210 | I | 827 | 0 | 0 | | 187 | 96 | 0 | M9- | 22 – 36 | | 14640715 | L80 | I1 | | 2210 |
| 2343 | I | 827 | 0 | 0 | | 187 | 96 | 0 | M9- | 22 – 36 | | 14898383 | Z2763 | I1 | A | 2343 |
| 2399 | I | 827 | 0 | 0 | | 187 | 96 | 0 | M9- | 22 – 36 | | 14955656 | Z2764 | I1 | | 2399 |
| 2418 | I | 827 | 0 | 0 | | 187 | 96 | 0 | M9- | 22 – 36 | | 14981302 | Z2765 | I1 | | 2418 |
| 2455 | I | 827 | 0 | 0 | | 187 | 96 | 0 | M9- | 22 – 36 | | 15022707 | M253 | I1 | | 2455 |
| 2623 | I | 827 | 0 | 0 | | 187 | 96 | 0 | M9- | 22 – 36 | | 15414257 | Z2769 | I1 | | 2623 |
| 2678 | III | 827 | 0 | 0 | | 186 | 95 | 2 | M9- | 22 – 36 | MD | 15500110 | L118 | I1 | | 2678 |

| | | M9+ | | | M9- | | | | | | Build 37 | | ISOGG | | |
|---|---|---|---|---|---|---|---|---|---|---|---|---|---|---|---|
| List | Stage | 0 | 1 | . | 0 | 1 | . | M9? | Clade(s) | Prob. | Position | Name | Subgroup Name | Amb. | List |
| 2714 | I | 827 | 0 | 0 | 187 | 96 | 0 | M9- | 22 – 36 | | 15553583 | Z2771 | I1 | | 2714 |
| 2823 | III | 826 | 0 | 1 | 187 | 96 | 0 | M9- | 22 – 36 | | 15754964 | Z2779 | I1 | | 2823 |
| 2922 | III | 825 | 0 | 2 | 187 | 95 | 1 | M9- | 22 – 36 | MD | 16084583 | Z2785 | I1 | | 2922 |
| 3000 | I | 827 | 0 | 0 | 187 | 96 | 0 | M9- | 22 – 36 | | 16333420 | Z2787^ | I1 | | 3000 |
| 3011 | I | 827 | 0 | 0 | 187 | 96 | 0 | M9- | 22 – 36 | | 16351523 | Z2788.2 | I1 | A | 3011 |
| 3023 | I | 827 | 0 | 0 | 187 | 96 | 0 | M9- | 22 – 36 | | 16377198 | S62 | I1 | | 3023 |
| 3028 | III | 826 | 0 | 1 | 186 | 96 | 1 | M9- | 22 – 36 | | 16392707 | Z2789 | I1 | | 3028 |
| 3048 | I | 827 | 0 | 0 | 187 | 96 | 0 | M9- | 22 – 36 | | 16443964 | Z2791 | I1 | | 3048 |
| 3091 | I | 827 | 0 | 0 | 187 | 96 | 0 | M9- | 22 – 36 | | 16537055 | Z2793 | I1 | | 3091 |
| 3117 | I | 827 | 0 | 0 | 187 | 96 | 0 | M9- | 22 – 36 | | 16599292 | Z2794 | I1 | | 3117 |
| 3146 | I | 827 | 0 | 0 | 187 | 96 | 0 | M9- | 22 – 36 | | 16693257 | L123 | I1 | | 3146 |
| 3149 | I | 827 | 0 | 0 | 187 | 96 | 0 | M9- | 22 – 36 | | 16695752 | Z2799 | I1 | | 3149 |
| 3160 | I | 827 | 0 | 0 | 187 | 96 | 0 | M9- | 22 – 36 | | 16737336 | Z2802^ | I1 | | 3160 |
| 3235 | I | 827 | 0 | 0 | 187 | 96 | 0 | M9- | 22 – 36 | | 16869992 | Z2803 | I1 | | 3235 |
| 3393 | III | 815 | 0 | 12 | 184 | 91 | 8 | M9- | 22 – 36 | MD, 1H | 17264749 | Z2807 | I1 | | 3393 |
| 3407 | III | 825 | 0 | 2 | 187 | 94 | 2 | M9- | 22 – 36 | MD | 17299483 | Z2808 | I1 | A | 3407 |
| 3439 | I | 827 | 0 | 0 | 187 | 96 | 0 | M9- | 22 – 36 | | 17390796 | Z2809^ | I1 | | 3439 |
| 3603 | III | 826 | 0 | 1 | 187 | 96 | 0 | M9- | 22 – 36 | | 17758299 | Z2814^ | I1 | | 3603 |
| 3610 | I | 827 | 0 | 0 | 187 | 96 | 0 | M9- | 22 – 36 | | 17766762 | S64 | I1 | | 3610 |
| 3723 | I | 827 | 0 | 0 | 187 | 96 | 0 | M9- | 22 – 36 | | 17976203 | Z2818 | I1 | | 3723 |
| 3772 | I | 827 | 0 | 0 | 187 | 96 | 0 | M9- | 22 – 36 | | 18077297 | L187 | I1 | | 3772 |
| 3844 | III | 826 | 0 | 1 | 187 | 96 | 0 | M9- | 22 – 36 | | 18387563 | Z2823 | I1 | | 3844 |
| 3904 | I | 827 | 0 | 0 | 187 | 96 | 0 | M9- | 22 – 36 | | 18614569 | Z2824 | I1 | | 3904 |
| 3959 | I | 827 | 0 | 0 | 187 | 96 | 0 | M9- | 22 – 36 | | 18759669 | S65 | I1 | | 3959 |
| 3984 | I | 827 | 0 | 0 | 187 | 96 | 0 | M9- | 22 – 36 | | 18802689 | Z2826 | I1 | | 3984 |
| 4013 | I | 827 | 0 | 0 | 187 | 96 | 0 | M9- | 22 – 36 | | 18855954 | Z2827^ | I1 | | 4013 |
| 4101 | I | 827 | 0 | 0 | 187 | 96 | 0 | M9- | 22 – 36 | | 19048605 | L75 | I1 | | 4101 |
| 4206 | I | 827 | 0 | 0 | 187 | 96 | 0 | M9- | 22 – 36 | | 19239993 | Z2835 | I1 | | 4206 |
| 4220 | I | 827 | 0 | 0 | 187 | 96 | 0 | M9- | 22 – 36 | | 19273317 | Z2836^ | I1 | | 4220 |
| 4228 | III | 826 | 0 | 1 | 187 | 96 | 0 | M9- | 22 – 36 | | 19284993 | Z2837 | I1 | | 4228 |
| 4260 | I | 826 | 1 | 0 | 187 | 96 | 0 | BOTH | | PI | 19372700 | Z2838.2 | I1 | A | 4260 |
| 4262 | III | 799 | 0 | 28 | 183 | 96 | 4 | M9- | 22 – 36 | | 19375292 | Z2839 | I1 | | 4262 |
| 4276 | I | 827 | 0 | 0 | 187 | 96 | 0 | M9- | 22 – 36 | | 19408727 | CTS10338 | I1 | | 4276 |
| 4326 | I | 827 | 0 | 0 | 187 | 96 | 0 | M9- | 22 – 36 | | 19563894 | Z2840 | I1 | | 4326 |
| 4374 | III | 826 | 0 | 1 | 185 | 93 | 5 | M9- | 22 – 36 | MD | 21097501 | Z2842 | I1 | | 4374 |
| 4377 | III | 827 | 0 | 0 | 187 | 72 | 24 | M9- | 22 – 36 | MD | 21100133 | Z2843 | I1 | | 4377 |
| 4449 | III | 826 | 0 | 1 | 187 | 96 | 0 | M9- | 22 – 36 | | 21288153 | Z2844 | I1 | | 4449 |
| 4458 | I | 827 | 0 | 0 | 187 | 96 | 0 | M9- | 22 – 36 | | 21309376 | Z2845 | I1 | | 4458 |
| 4504 | I | 827 | 0 | 0 | 187 | 96 | 0 | M9- | 22 – 36 | | 21402851 | Z2848 | I1 | | 4504 |
| 4561 | I | 827 | 0 | 0 | 187 | 96 | 0 | M9- | 22 – 36 | | 21561968 | Z2850 | I1 | | 4561 |
| 4925 | I | 827 | 0 | 0 | 187 | 96 | 0 | M9- | 22 – 36 | | 22011964 | Z2855^ | I1 | | 4925 |
| 4986 | III | 825 | 0 | 2 | 186 | 96 | 1 | M9- | 22 – 36 | 1H | 22174158 | Z2857^ | I1 | | 4986 |
| 5030 | I | 827 | 0 | 0 | 187 | 96 | 0 | M9- | 22 – 36 | | 22513726 | L81 | I1 | | 5030 |
| 5043 | III | 827 | 0 | 0 | 187 | 95 | 1 | M9- | 22 – 36 | MD | 22538925 | Z2861 | I1 | | 5043 |
| 5046 | I | 827 | 0 | 0 | 187 | 96 | 0 | M9- | 22 – 36 | | 22550691 | Z2862 | I1 | | 5046 |
| 5164 | I | 826 | 1 | 0 | 187 | 96 | 0 | BOTH | | PI | 22750951 | P203.2 | I1 | A | 5164 |
| 5231 | I | 827 | 0 | 0 | 187 | 96 | 0 | M9- | 22 – 36 | | 22909673 | Z2863 | I1 | | 5231 |
| 5232 | I | 827 | 0 | 0 | 187 | 96 | 0 | M9- | 22 – 36 | | 22914378 | S66 | I1 | | 5232 |
| 5248 | I | 827 | 0 | 0 | 187 | 96 | 0 | M9- | 22 – 36 | | 22928344 | Z2864 | I1 | | 5248 |
| 5261 | I | 827 | 0 | 0 | 187 | 96 | 0 | M9- | 22 – 36 | | 22948591 | Z2865 | I1 | | 5261 |
| 5351 | I | 827 | 0 | 0 | 187 | 96 | 0 | M9- | 22 – 36 | | 23150415 | Z2870 | I1 | A | 5351 |
| 5353 | I | 827 | 0 | 0 | 187 | 96 | 0 | M9- | 22 – 36 | | 23153786 | Z2871 | I1 | | 5353 |
| 5358 | I | 827 | 0 | 0 | 187 | 96 | 0 | M9- | 22 – 36 | | 23159643 | Z2872 | I1 | | 5358 |
| 5397 | I | 827 | 0 | 0 | 187 | 96 | 0 | M9- | 22 – 36 | | 23264430 | Z2875 | I1 | | 5397 |
| 5404 | I | 827 | 0 | 0 | 187 | 96 | 0 | M9- | 22 – 36 | | 23269094 | Z2876 | I1 | | 5404 |
| 5418 | III | 827 | 0 | 0 | 187 | 95 | 1 | M9- | 22 – 36 | MD | 23303290 | Z2878 | I1 | | 5418 |
| 5419 | I | 827 | 0 | 0 | 187 | 96 | 0 | M9- | 22 – 36 | | 23305273 | Z2879 | I1 | | 5419 |
| 5439 | I | 827 | 0 | 0 | 187 | 96 | 0 | M9- | 22 – 36 | | 23379549 | Z2880^ | I1 | | 5439 |
| 5613 | I | 827 | 0 | 0 | 187 | 96 | 0 | M9- | 22 – 36 | | 23957372 | Z2885 | I1 | | 5613 |
| 5691 | III | 825 | 0 | 2 | 187 | 96 | 0 | M9- | 22 – 36 | | 24499543 | Z2887 | I1 | | 5691 |
| 3090 | I | 827 | 0 | 0 | 187 | 96 | 0 | M9- | 22 – 36 | | 16536201 | Z2792 | I1~ | | 3090 |
| 5653 | III | 824 | 0 | 3 | 187 | 96 | 0 | M9- | 22 – 36 | | 24383715 | Z2886 | I1~ | | 5653 |
| 1164 | III | 825 | 0 | 2 | 187 | 95 | 1 | M9- | | PI | 8097201 | Z2891 | I1a~ | | 1164 |
| 4126 | III | 826 | 0 | 1 | 188 | 95 | 0 | M9- | | PI | 19103066 | V3772 | I1a~ | | 4126 |
| 3224 | III | 826 | 0 | 1 | 267 | 15 | 1 | M9- | 23 – 26 | | 16850779 | Z2336 | I1a1 | | 3224 |
| 2743 | IV | 826 | 0 | 1 | 281 | 0 | 2 | Mono | Mono | | 15591446 | M227 | I1a1a1a1a | | 2743 |
| 4836 | IV | 827 | 0 | 0 | 283 | 0 | 0 | Mono | Mono | | 21894407 | M72 | I1a1a1a1a2a1 | | 4836 |
| 637 | IV | 827 | 0 | 0 | 283 | 0 | 0 | Mono | Mono | | 6932171 | L817.2 | I1a1a1b5b1a1~ | A | 637 |
| 1461 | I | 827 | 0 | 0 | 269 | 14 | 0 | M9- | 24 – 26 | | 8576009 | S142 | I1a1b1 | | 1461 |
| 2630 | II | 827 | 0 | 0 | 281 | 2 | 0 | M9- | 25 | | 15426005 | P109 | I1a1b1a1 | | 2630 |
| 1157 | III | 823 | 0 | 4 | 281 | 2 | 0 | M9- | 26 | | 8073670 | Z74 | I1a1b1a4a | | 1157 |
| 4826 | IV | 827 | 0 | 0 | 283 | 0 | 0 | Mono | Mono | | 21892626 | S335 | I1a1b1a4a1a1 | | 4826 |
| 4868 | IV | 827 | 0 | 0 | 283 | 0 | 0 | Mono | Mono | | 21907394 | M120.2 | I1a1b1a4a1a1b1a3d~ | A | 4868 |
| 4784 | IV | 827 | 0 | 0 | 283 | 0 | 0 | Mono | Mono | | 21868140 | Y17701 | I1a1b1a4a1a1g3~ | | 4784 |
| 3832 | III | 826 | 0 | 1 | 261 | 22 | 0 | M9- | 27 – 30 | | 18234588 | Z60 | I1a2a1a1~ | | 3832 |
| 3662 | III | 827 | 0 | 0 | 266 | 13 | 4 | M9- | 28 – 30 | MD | 17863355 | Z140 | I1a2a1a1a | | 3662 |
| 4968 | I | 827 | 0 | 0 | 276 | 7 | 0 | M9- | 29 | | 22142382 | Z2535 | I1a2a1a1a1 | | 4968 |
| 3703 | I | 827 | 0 | 0 | 278 | 5 | 0 | M9- | 30 | | 17936279 | S2169.3 | I1a2a1a1a2 | A | 3703 |
| 1678 | IV | 826 | 0 | 1 | 282 | 0 | 1 | Mono | Mono | | 9381642 | Y6363 | I1a2a1a1a2b3b~ | | 1678 |
| 2636 | IV | 827 | 0 | 0 | 283 | 0 | 0 | Mono | Mono | | 15436915 | M219 | I1a2a1a1d2a1a~ | A | 2636 |
| 2805 | III | 826 | 0 | 1 | 265 | 18 | 0 | M9- | 31 – 32 | | 15711340 | Z138.2 | I1a2b | A | 2805 |
| 4450 | III | 827 | 0 | 0 | 265 | 1 | 17 | M9- | 31 – 32 | MD | 21288655 | Z139 | I1a2b | | 4450 |
| 5240 | IV | 827 | 0 | 0 | 283 | 0 | 0 | Mono | Mono | | 22920023 | F3371.2 | I1a2b1e1a~ | A | 5240 |
| 2131 | I | 827 | 0 | 0 | 254 | 29 | 0 | M9- | 33 – 36 | | 14401486 | Z63 | I1a3 | | 2131 |
| 2429 | I | 827 | 0 | 0 | 276 | 7 | 0 | M9- | 36 | | 14994820 | S2078 | I1a3a | | 2429 |
| 2335 | IV | 827 | 0 | 0 | 283 | 0 | 0 | Mono | Mono | | 14891535 | S16522 | I1a3a1a2b~ | | 2335 |
| 1396 | IV | 827 | 0 | 0 | 283 | 0 | 0 | Mono | Mono | | 8484412 | V2110.1 | I1a3b1a1 | A | 1396 |
| 1043 | III | 823 | 1 | 3 | 274 | 8 | 1 | BOTH | | PI | 7811520 | Z12228.3 | I1a3b2~ | A,W | 1043 |
| 3237 | IV | 827 | 0 | 0 | 283 | 0 | 0 | Mono | Mono | | 16870183 | CTS6397 | I1b1 | | 3237 |
| 994 | I | 827 | 0 | 0 | 178 | 105 | 0 | M9- | 37 – 59 | | 7728344 | Z2634 | I2 | | 994 |
| 1208 | I | 827 | 0 | 0 | 178 | 105 | 0 | M9- | 37 – 59 | | 8211182 | PF3651 | I2 | | 1208 |
| 1306 | I | 827 | 0 | 0 | 178 | 105 | 0 | M9- | 37 – 59 | | 8392226 | Z2636 | I2 | | 1306 |
| 1346 | III | 826 | 0 | 1 | 178 | 105 | 0 | M9- | 37 – 59 | | 8443332 | Z2637 | I2 | | 1346 |
| 1661 | III | 827 | 0 | 0 | 178 | 104 | 1 | M9- | 37 – 59 | MD | 9133937 | Z2642 | I2 | | 1661 |
| 3056 | I | 827 | 0 | 0 | 178 | 105 | 0 | M9- | 37 – 59 | | 16459388 | Z2653 | I2 | | 3056 |
| 3130 | III | 824 | 0 | 3 | 178 | 105 | 0 | M9- | 37 – 59 | | 16638804 | S31 | I2 | | 3130 |

| | | M9+ | | | M9- | | | | | | Build 37 | | ISOGG | | |
|---|---|---|---|---|---|---|---|---|---|---|---|---|---|---|---|
| List | Stage | 0 | 1 | . | 0 | 1 | . | M9? | Clade(s) | Prob. | Position | Name | Subgroup Name | Amb. | List |
| 3140 | I | 827 | 0 | 0 | 178 | 105 | 0 | M9- | 37 – 59 | | 16671358 | Z2654 | I2 | | 3140 |
| 3744 | III | 827 | 0 | 0 | 178 | 104 | 1 | M9- | 37 – 59 | MD | 18032844 | Z2658 | I2 | | 3744 |
| 3939 | I | 827 | 0 | 0 | 178 | 105 | 0 | M9- | 37 – 59 | | 18700150 | S329.2 | I2 | A | 3939 |
| 4250 | III | 826 | 0 | 1 | 178 | 105 | 0 | M9- | 37 – 59 | | 19347315 | Z2661 | I2 | | 4250 |
| 4955 | I | 827 | 0 | 0 | 178 | 105 | 0 | M9- | 37 – 59 | | 22109679 | Z2667 | I2 | | 4955 |
| 2068 | I | 827 | 0 | 0 | 178 | 105 | 0 | M9- | 37 – 59 | | 14238700 | Z2647 | I2a | | 2068 |
| 2596 | III | 826 | 0 | 1 | 178 | 105 | 0 | M9- | 37 – 59 | | 15349855 | Z2650 | I2a | | 2596 |
| 3885 | I | 827 | 0 | 0 | 178 | 105 | 0 | M9- | 37 – 59 | | 18581138 | Z2659 | I2a | | 3885 |
| 4511 | I | 827 | 0 | 0 | 178 | 105 | 0 | M9- | 37 – 59 | | 21414141 | Z2663 | I2a | | 4511 |
| 1073 | I | 827 | 0 | 0 | 182 | 101 | 0 | M9- | 37 – 58 | | 7879415 | S238 | I2a1 | | 1073 |
| 503 | I | 827 | 0 | 0 | 256 | 27 | 0 | M9- | 37 – 41 | | 6732120 | Z2609 | I2a1a | | 503 |
| 1097 | I | 827 | 0 | 0 | 256 | 27 | 0 | M9- | 37 – 41 | | 7919987 | Z2612 | I2a1a | | 1097 |
| 1979 | III | 813 | 0 | 14 | 250 | 27 | 6 | M9- | 37 – 41 | | 14063767 | Z2617 | I2a1a | | 1979 |
| 2124 | III | 820 | 0 | 7 | 255 | 27 | 1 | M9- | 37 – 41 | | 14371245 | S2648 | I2a1a | | 2124 |
| 2614 | I | 827 | 0 | 0 | 256 | 27 | 0 | M9- | 37 – 41 | | 15394811 | Z2651 | I2a1a | | 2614 |
| 2819 | I | 827 | 0 | 0 | 256 | 27 | 0 | M9- | 37 – 41 | | 15745246 | Z2619 | I2a1a | | 2819 |
| 2903 | I | 827 | 0 | 0 | 256 | 27 | 0 | M9- | 37 – 41 | | 15984708 | Z2620 | I2a1a | | 2903 |
| 3076 | III | 821 | 0 | 6 | 254 | 12 | 17 | M9- | 37 – 41 | MD | 16495009 | Z2671 | I2a1a | | 3076 |
| 3459 | III | 826 | 0 | 1 | 255 | 27 | 1 | M9- | 37 – 41 | | 17427444 | Z2622 | I2a1a | | 3459 |
| 3733 | I | 827 | 0 | 0 | 256 | 27 | 0 | M9- | 37 – 41 | | 18021838 | Z2624 | I2a1a | | 3733 |
| 4587 | I | 827 | 0 | 0 | 256 | 27 | 0 | M9- | 37 – 41 | | 21612591 | Z2625 | I2a1a | | 4587 |
| 4941 | I | 827 | 0 | 0 | 256 | 27 | 0 | M9- | 37 – 41 | | 22069173 | Z2626 | I2a1a | | 4941 |
| 583 | I | 827 | 0 | 0 | 265 | 18 | 0 | M9- | 37 – 39 | | 6874115 | CTS595.2 | I2a1a1 | A | 583 |
| 4772 | I | 827 | 0 | 0 | 272 | 11 | 0 | M9- | 37 – 38 | | 21865821 | PF4056 | I2a1a1a | | 4772 |
| 5491 | III | 827 | 0 | 0 | 272 | 6 | 5 | M9- | 37 – 38 | MD | 23496560 | S433 | I2a1a1a | | 5491 |
| 504 | III | 827 | 0 | 0 | 272 | 7 | 4 | M9- | 37 – 38 | MD | 6735361 | PF3950 | I2a1a1a~ | | 504 |
| 826 | I | 827 | 0 | 0 | 272 | 11 | 0 | M9- | 37 – 38 | | 7305474 | PF3958 | I2a1a1a~ | | 826 |
| 938 | III | 827 | 0 | 0 | 271 | 11 | 1 | M9- | 37 – 38 | | 7638008 | V1304 | I2a1a1a~ | | 938 |
| 1167 | I | 827 | 0 | 0 | 272 | 11 | 0 | M9- | 37 – 38 | | 8105907 | Y3997 | I2a1a1a~ | | 1167 |
| 1198 | I | 827 | 0 | 0 | 272 | 11 | 0 | M9- | 37 – 38 | | 8189057 | V1694 | I2a1a1a~ | | 1198 |
| 1336 | I | 827 | 0 | 0 | 272 | 11 | 0 | M9- | 37 – 38 | | 8432044 | V1959 | I2a1a1a~ | | 1336 |
| 1594 | IV | 827 | 0 | 0 | 272 | 0 | 11 | Mono | Mono | MD | 8839543 | Y3996 | I2a1a1a~ | | 1594 |
| 1743 | III | 824 | 0 | 3 | 271 | 11 | 1 | M9- | 37 – 38 | | 9832942 | PF3865 | I2a1a1a~ | | 1743 |
| 1764 | IV | 825 | 0 | 2 | 272 | 0 | 11 | Mono | Mono | MD | 9876142 | PF3986 | I2a1a1a~ | | 1764 |
| 1837 | III | 822 | 0 | 5 | 267 | 11 | 5 | M9- | 37 – 38 | | 13238818 | PF3988 | I2a1a1a~ | | 1837 |
| 2188 | I | 827 | 0 | 0 | 272 | 11 | 0 | M9- | 37 – 38 | | 14566306 | PF3710 | I2a1a1a~ | | 2188 |
| 2309 | I | 827 | 0 | 0 | 272 | 11 | 0 | M9- | 37 – 38 | | 14861707 | PF3866 | I2a1a1a~ | | 2309 |
| 2776 | I | 826 | 1 | 0 | 272 | 11 | 0 | BOTH | | PI | 15652249 | PF4009 | I2a1a1a~ | | 2776 |
| 3074 | I | 827 | 0 | 0 | 272 | 11 | 0 | M9- | 37 – 38 | | 16492382 | PF4015 | I2a1a1a~ | | 3074 |
| 3086 | III | 820 | 0 | 7 | 270 | 8 | 5 | M9- | 37 – 38 | MD | 16523608 | PF4017 | I2a1a1a~ | | 3086 |
| 3104 | III | 827 | 0 | 0 | 272 | 2 | 9 | M9- | 37 – 38 | MD | 16560849 | PF4018 | I2a1a1a~ | | 3104 |
| 3123 | I | 827 | 0 | 0 | 272 | 11 | 0 | M9- | 37 – 38 | | 16620529 | CTS6027 | I2a1a1a~ | A | 3123 |
| 3542 | III | 826 | 0 | 1 | 271 | 11 | 1 | M9- | 37 – 38 | | 17605878 | PF4026 | I2a1a1a~ | | 3542 |
| 3621 | I | 827 | 0 | 0 | 272 | 11 | 0 | M9- | 37 – 38 | | 17783792 | PF4029 | I2a1a1a~ | | 3621 |
| 3918 | III | 827 | 0 | 0 | 271 | 11 | 1 | M9- | 37 – 38 | | 18638039 | V3349 | I2a1a1a~ | | 3918 |
| 4187 | I | 827 | 0 | 0 | 272 | 11 | 0 | M9- | 37 – 38 | | 19218641 | V3915 | I2a1a1a~ | | 4187 |
| 4393 | III | 827 | 0 | 0 | 272 | 3 | 8 | M9- | 37 – 38 | MD | 21146700 | PF4046 | I2a1a1a~ | | 4393 |
| 4406 | I | 827 | 0 | 0 | 272 | 11 | 0 | M9- | 37 – 38 | | 21168729 | PF4047 | I2a1a1a~ | | 4406 |
| 4427 | III | 827 | 0 | 0 | 271 | 7 | 5 | M9- | 37 – 38 | MD | 21249535 | PF4048 | I2a1a1a~ | | 4427 |
| 4487 | I | 827 | 0 | 0 | 272 | 11 | 0 | M9- | 37 – 38 | | 21367607 | PF4049 | I2a1a1a~ | | 4487 |
| 4730 | III | 826 | 0 | 1 | 272 | 11 | 0 | M9- | 37 – 38 | | 21778131 | PF4054 | I2a1a1a~ | | 4730 |
| 4872 | I | 827 | 0 | 0 | 272 | 11 | 0 | M9- | 37 – 38 | | 21911727 | S6075 | I2a1a1a~ | A | 4872 |
| 4967 | I | 827 | 0 | 0 | 272 | 11 | 0 | M9- | 37 – 38 | | 22141167 | PF3875 | I2a1a1a~ | | 4967 |
| 5034 | III | 819 | 0 | 8 | 270 | 10 | 3 | M9- | 37 – 38 | MD | 22520043 | PF4068 | I2a1a1a~ | | 5034 |
| 5274 | III | 827 | 0 | 0 | 272 | 10 | 1 | M9- | 37 – 38 | MD | 22975528 | PF4070 | I2a1a1a~ | | 5274 |
| 5357 | III | 827 | 0 | 0 | 271 | 11 | 1 | M9- | 37 – 38 | | 23159136 | PF3877 | I2a1a1a~ | | 5357 |
| 5566 | III | 827 | 0 | 0 | 271 | 11 | 1 | M9- | 37 – 38 | | 23788111 | PF4075 | I2a1a1a~ | | 5566 |
| 5654 | III | 825 | 0 | 2 | 272 | 11 | 0 | M9- | 37 – 38 | | 24383802 | PF4076 | I2a1a1a~ | | 5654 |
| 5671 | IV | 820 | 0 | 7 | 271 | 0 | 12 | Mono | Mono | MD | 24433559 | PF4077 | I2a1a1a~ | | 5671 |
| 5687 | III | 824 | 0 | 3 | 272 | 11 | 0 | M9- | 37 – 38 | | 24495432 | PF4079 | I2a1a1a~ | | 5687 |
| 5693 | III | 825 | 0 | 2 | 271 | 11 | 1 | M9- | 37 – 38 | | 24510801 | PF4080 | I2a1a1a~ | | 5693 |
| 853 | I | 827 | 0 | 0 | 272 | 11 | 0 | M9- | 37 – 38 | | 7403261 | PF3961 | I2a1a1a1~ | | 853 |
| 1445 | I | 827 | 0 | 0 | 272 | 11 | 0 | M9- | 37 – 38 | | 8554849 | V2245 | I2a1a1a1~ | | 1445 |
| 1723 | I | 827 | 0 | 0 | 272 | 11 | 0 | M9- | 37 – 38 | | 9784940 | PF3984 | I2a1a1a1~ | | 1723 |
| 1856 | III | 827 | 0 | 0 | 270 | 6 | 7 | M9- | 37 – 38 | MD | 13653148 | PF3992 | I2a1a1a1~ | | 1856 |
| 2021 | I | 827 | 0 | 0 | 272 | 11 | 0 | M9- | 37 – 38 | | 14140435 | PF3996 | I2a1a1a1~ | | 2021 |
| 2063 | III | 827 | 0 | 0 | 272 | 10 | 1 | M9- | 37 – 38 | MD | 14224903 | PF3998 | I2a1a1a1~ | | 2063 |
| 3005 | I | 827 | 0 | 0 | 272 | 11 | 0 | M9- | 37 – 38 | | 16345907 | PF3741 | I2a1a1a1~ | | 3005 |
| 3888 | I | 827 | 0 | 0 | 272 | 11 | 0 | M9- | 37 – 38 | | 18584762 | V3296 | I2a1a1a1~ | | 3888 |
| 4317 | III | 827 | 0 | 0 | 272 | 9 | 2 | M9- | 37 – 38 | MD | 19530092 | PF3872 | I2a1a1a1~ | | 4317 |
| 4435 | I | 827 | 0 | 0 | 272 | 11 | 0 | M9- | 37 – 38 | | 21257341 | Y3992 | I2a1a1a1~ | | 4435 |
| 4522 | I | 827 | 0 | 0 | 272 | 11 | 0 | M9- | 37 – 38 | | 21453968 | PF4051 | I2a1a1a1~ | | 4522 |
| 5485 | IV | 827 | 0 | 0 | 272 | 0 | 11 | Mono | Mono | MD | 23485097 | PF3848 | I2a1a1a1~ | | 5485 |
| 801 | III | 827 | 0 | 0 | 271 | 11 | 1 | M9- | 37 – 38 | | 7227967 | PF3633 | I2a1a1a1a~ | | 801 |
| 2084 | III | 827 | 0 | 0 | 272 | 9 | 2 | M9- | 37 – 38 | MD | 14276262 | Z2050 | I2a1a1a1a~ | | 2084 |
| 5394 | I | 827 | 0 | 0 | 272 | 11 | 0 | M9- | 37 – 38 | | 23256670 | Z2053 | I2a1a1a1a~ | | 5394 |
| 708 | I | 827 | 0 | 0 | 279 | 4 | 0 | M9- | 38 | | 7030410 | PF4087 | I2a1a1a1a1 | | 708 |
| 1617 | I | 827 | 0 | 0 | 279 | 4 | 0 | M9- | 38 | | 8895456 | PF4089 | I2a1a1a1a1 | | 1617 |
| 2037 | I | 827 | 0 | 0 | 279 | 4 | 0 | M9- | 38 | | 14185731 | CTS2111 | I2a1a1a1a1 | | 2037 |
| 2946 | III | 826 | 0 | 1 | 279 | 4 | 0 | M9- | 38 | | 16202544 | S184 | I2a1a1a1a1 | | 2946 |
| 3240 | I | 827 | 0 | 0 | 279 | 4 | 0 | M9- | 38 | | 16876275 | PF4093 | I2a1a1a1a1 | | 3240 |
| 3426 | I | 827 | 0 | 0 | 279 | 4 | 0 | M9- | 38 | | 17344302 | PF4095 | I2a1a1a1a1 | | 3426 |
| 3919 | III | 826 | 0 | 1 | 280 | 3 | 0 | M9- | NU | | 18641477 | V3352 | I2a1a1a1a1 | | 3919 |
| 4588 | I | 827 | 0 | 0 | 279 | 4 | 0 | M9- | 38 | | 21614113 | PF4053 | I2a1a1a1a1 | | 4588 |
| 4956 | I | 827 | 0 | 0 | 279 | 4 | 0 | M9- | 38 | | 22111642 | PF4060 | I2a1a1a1a1 | | 4956 |
| 5228 | I | 827 | 0 | 0 | 279 | 4 | 0 | M9- | 38 | | 22908323 | PF4099 | I2a1a1a1a1 | | 5228 |
| 5291 | I | 827 | 0 | 0 | 279 | 4 | 0 | M9- | 38 | | 23023839 | PF4100 | I2a1a1a1a1 | | 5291 |
| 5450 | I | 827 | 0 | 0 | 279 | 4 | 0 | M9- | 38 | | 23392582 | PF4072 | I2a1a1a1a1 | | 5450 |
| 1534 | I | 827 | 0 | 0 | 279 | 4 | 0 | M9- | 38 | | 8681127 | V2447 | I2a1a1a1a1a | | 1534 |
| 4650 | IV | 827 | 0 | 0 | 283 | 0 | 0 | Mono | Mono | | 21717515 | PF4447 | I2a1a1a1a1a1e6a5~ | A | 4650 |
| 4575 | IV | 827 | 0 | 0 | 283 | 0 | 0 | Mono | Mono | | 21600514 | Z24547.2 | I2a1a1a1a2a1a~ | | 4575 |
| 2562 | IV | 827 | 0 | 0 | 283 | 0 | 0 | Mono | Mono | | 15241111 | Z27395.1 | I2a1a1a2~ | A | 2562 |
| 1468 | III | 825 | 0 | 2 | 271 | 10 | 2 | M9- | 37 – 38 | MD | 8586442 | S6233 | I2a1a1a3~ | W | 1468 |
| 1590 | I | 827 | 0 | 0 | 276 | 7 | 0 | M9- | 39 | | 8828242 | S11985 | I2a1a1b1a~ | A | 1590 |
| 829 | I | 827 | 0 | 0 | 274 | 9 | 0 | M9- | 40 – 41 | | 7317227 | Z2577 | I2a1a2 | | 829 |
| 2725 | III | 826 | 0 | 1 | 274 | 9 | 0 | M9- | 40 – 41 | | 15574052 | S328.1 | I2a1a2 | A | 2725 |

| | | M9+ | | | M9- | | | | | | Build 37 | | ISOGG | | |
|---|---|---|---|---|---|---|---|---|---|---|---|---|---|---|---|
| List | Stage | 0 | 1 | . | 0 | 1 | . | M9? | Clade(s) | Prob. | Position | Name | Subgroup Name | Amb. | List |
| 2962 | I | 827 | 0 | 0 | 274 | 9 | 0 | M9- | 40 – 41 | | 16233135 | Z2591 | I2a1a2 | | 2962 |
| 105 | I | 827 | 0 | 0 | 274 | 9 | 0 | M9- | 40 – 41 | | 2785672 | Z2575 | I2a1a2~ | | 105 |
| 3112 | I | 827 | 0 | 0 | 274 | 9 | 0 | M9- | 40 – 41 | | 16594452 | Z2592 | I2a1a2~ | | 3112 |
| 5029 | I | 827 | 0 | 0 | 277 | 6 | 0 | M9- | 40 | | 22513718 | S185.1 | I2a1a2a | | 5029 |
| 3962 | II | 827 | 0 | 0 | 280 | 3 | 0 | M9- | 41 | | 18760081 | S392 | I2a1a2b | | 3962 |
| 4246 | III | 827 | 0 | 0 | 279 | 3 | 1 | M9- | 41 | | 19334357 | V4023 | I2a1a2b1a1 | | 4246 |
| 2701 | II | 827 | 0 | 0 | 280 | 3 | 0 | M9- | 41 | | 15531354 | YP204 | I2a1a2b1a1a1 | | 2701 |
| 933 | I | 827 | 0 | 0 | 209 | 74 | 0 | M9- | 42 – 58 | | 7628484 | S23 | I2a1b | | 933 |
| 1947 | III | 827 | 0 | 0 | 209 | 73 | 1 | M9- | 42 – 58 | MD | 13992338 | S30 | I2a1b | | 1947 |
| 2723 | I | 827 | 0 | 0 | 209 | 74 | 0 | M9- | 42 – 58 | | 15567863 | S884.3 | I2a1b | A | 2723 |
| 3483 | III | 826 | 0 | 1 | 209 | 74 | 0 | M9- | 42 – 58 | | 17493630 | S32 | I2a1b | | 3483 |
| 3500 | I | 827 | 0 | 0 | 209 | 74 | 0 | M9- | 42 – 58 | | 17516123 | S153 | I2a1b | | 3500 |
| 3958 | III | 827 | 0 | 0 | 209 | 73 | 1 | M9- | 42 – 58 | MD | 18747493 | S33 | I2a1b | | 3958 |
| 4048 | I | 827 | 0 | 0 | 209 | 74 | 0 | M9- | 42 – 58 | | 18926388 | CTS9482 | I2a1b | | 4048 |
| 4118 | I | 827 | 0 | 0 | 209 | 74 | 0 | M9- | 42 – 58 | | 19077754 | L181 | I2a1b | | 4118 |
| 5124 | I | 827 | 0 | 0 | 209 | 74 | 0 | M9- | 42 – 58 | | 22725379 | S150 | I2a1b | | 5124 |
| 747 | I | 827 | 0 | 0 | 219 | 64 | 0 | M9- | 42 – 56 | | 7113556 | L59 | I2a1b1 | | 747 |
| 989 | I | 827 | 0 | 0 | 219 | 64 | 0 | M9- | 42 – 56 | | 7716262 | S151 | I2a1b1 | | 989 |
| 1284 | I | 827 | 0 | 0 | 219 | 64 | 0 | M9- | 42 – 56 | | 8353707 | S120 | I2a1b1 | | 1284 |
| 2693 | I | 827 | 0 | 0 | 219 | 64 | 0 | M9- | 42 – 56 | | 15517851 | S24 | I2a1b1 | | 2693 |
| 3150 | I | 827 | 0 | 0 | 219 | 64 | 0 | M9- | 42 – 56 | | 16699334 | S117 | I2a1b1 | | 3150 |
| 3520 | I | 827 | 0 | 0 | 219 | 64 | 0 | M9- | 42 – 56 | | 17570599 | S152 | I2a1b1 | | 3520 |
| 4025 | I | 827 | 0 | 0 | 219 | 64 | 0 | M9- | 42 – 56 | | 18888200 | U250 | I2a1b1 | | 4025 |
| 4648 | I | 827 | 0 | 0 | 219 | 64 | 0 | M9- | 42 – 56 | | 21717307 | M223 | I2a1b1 | | 4648 |
| 5685 | III | 825 | 0 | 2 | 219 | 64 | 0 | M9- | 42 – 56 | | 24475669 | S119 | I2a1b1 | | 5685 |
| 613 | I | 827 | 0 | 0 | 220 | 63 | 0 | M9- | 43 – 56 | | 6906332 | CTS616 | I2a1b1a | | 613 |
| 3949 | I | 827 | 0 | 0 | 220 | 63 | 0 | M9- | 43 – 56 | | 18732197 | CTS9183 | I2a1b1a | | 3949 |
| 1726 | I | 827 | 0 | 0 | 277 | 6 | 0 | M9- | 45 | | 9791250 | S166.1 | I2a1b1a1a1a | A | 1726 |
| 2780 | IV | 827 | 0 | 0 | 283 | 0 | 0 | Mono | Mono | | 15656524 | Y6003 | I2a1b1a1a1b1a2a~ | A | 2780 |
| 201 | III | 819 | 0 | 8 | 279 | 1 | 3 | M9- | 46 | | 2980725 | Z2054 | I2a1b1a1b1a1a | | 201 |
| 118 | IV | 824 | 0 | 3 | 283 | 0 | 0 | Mono | Mono | | 2809424 | BY138 | I2a1b1a1b1a1a4 | | 118 |
| 4194 | I | 827 | 0 | 0 | 232 | 51 | 0 | M9- | 47 – 56 | | 19232160 | CTS10057 | I2a1b1a2 | | 4194 |
| 4210 | I | 827 | 0 | 0 | 232 | 51 | 0 | M9- | 47 – 56 | | 19255890 | V3945 | I2a1b1a2 | | 4210 |
| 3259 | IV | 827 | 0 | 0 | 283 | 0 | 0 | Mono | Mono | | 16903025 | L1266.2 | I2a1b1a2a2a1b~ | A | 3259 |
| 4709 | I | 827 | 0 | 0 | 237 | 46 | 0 | M9- | 49 – 56 | | 21763755 | S390 | I2a1b1a2b1 | | 4709 |
| 3251 | I | 827 | 0 | 0 | 251 | 32 | 0 | M9- | 51 – 56 | | 16889964 | CTS6433 | I2a1b1a2b1a2 | | 3251 |
| 3857 | I | 827 | 0 | 0 | 278 | 5 | 0 | M9- | 53 – 54 | | 18405788 | L1198 | I2a1b1a2b1a2a1a1a1a | | 3857 |
| 3477 | II | 827 | 0 | 0 | 281 | 2 | 0 | M9- | 54 | | 17473966 | Z190 | I2a1b1a2b1a2a1a1a1a1a | A | 3477 |
| 5627 | III | 774 | 0 | 53 | 278 | 3 | 2 | M9- | 55 | | 23977120 | ZS20 | I2a1b1a2b1a2a2 | | 5627 |
| 629 | IV | 815 | 0 | 12 | 280 | 0 | 3 | Mono | Mono | | 6931960 | L1272 | I2a1b1a2b1a2a2a | | 629 |
| 509 | III | 822 | 3 | 2 | 281 | 0 | 2 | M9+ | | PI | 6740034 | M8963.2 | I2a1b1a2b1a2a3a1~ | W | 509 |
| 1762 | III | 827 | 0 | 0 | 270 | 5 | 8 | M9- | 56 | MD | 9869354 | M11760 | I2a1b1a2b1a2e2a3~ | A | 1762 |
| 4790 | IV | 827 | 0 | 0 | 280 | 0 | 3 | Mono | Mono | | 21869264 | M54.2 | I2a1b1a2b2b~ | A | 4790 |
| 2791 | I | 827 | 0 | 0 | 273 | 10 | 0 | M9- | 57 | | 15668070 | S154 | I2a1b2a | | 2791 |
| 2942 | I | 827 | 0 | 0 | 273 | 10 | 0 | M9- | 57 | | 16199051 | S155 | I2a1b2a | | 2942 |
| 2945 | I | 827 | 0 | 0 | 273 | 10 | 0 | M9- | 57 | | 16202267 | S156 | I2a1b2a | | 2945 |
| 3124 | I | 827 | 0 | 0 | 273 | 10 | 0 | M9- | 57 | | 16626617 | S159.2 | I2a1b2a | A | 3124 |
| 2042 | I | 827 | 0 | 0 | 279 | 4 | 0 | M9- | 59 | | 14197631 | S292 | I2a2 | A | 2042 |
| 4024 | III | 826 | 0 | 1 | 279 | 4 | 0 | M9- | 59 | | 18887888 | S333 | I2a2 | | 4024 |
| 5517 | III | 827 | 0 | 0 | 279 | 1 | 3 | M9- | 59 | MD | 23566624 | S6610 | I2a2 | | 5517 |
| 152 | IV | 827 | 0 | 0 | 283 | 0 | 0 | Mono | Mono | | 2846553 | F741 | I2a2a1a1a2 | A | 152 |
| 2565 | IV | 826 | 0 | 1 | 283 | 0 | 0 | Mono | Mono | | 15252409 | CTS3902.2 | I2a2b2~ | A | 2565 |
| 1326 | IV | 827 | 0 | 0 | 283 | 0 | 0 | Mono | Mono | | 8426321 | S332 | I2b | | 1326 |
| 1099 | III | 826 | 0 | 1 | 59 | 224 | 0 | M9- | 22 – 66 | | 7922133 | Y1866 | IJ | | 1099 |
| 1214 | III | 823 | 0 | 4 | 59 | 224 | 0 | M9- | 22 – 66 | | 8222299 | L403 | IJ | | 1214 |
| 1447 | I | 827 | 0 | 0 | 59 | 224 | 0 | M9- | 22 – 66 | | 8558969 | S22 | IJ | | 1447 |
| 1474 | I | 827 | 0 | 0 | 59 | 224 | 0 | M9- | 22 – 66 | | 8590752 | PF3526 | IJ | | 1474 |
| 1614 | I | 827 | 0 | 0 | 59 | 224 | 0 | M9- | 22 – 66 | | 8888300 | YSC0001300 | IJ | | 1614 |
| 1622 | I | 827 | 0 | 0 | 59 | 224 | 0 | M9- | 22 – 66 | | 8906357 | F1460 | IJ | | 1622 |
| 1872 | III | 827 | 0 | 0 | 59 | 222 | 2 | M9- | 22 – 66 | MD | 13676377 | FGC1574 | IJ | | 1872 |
| 1963 | I | 827 | 0 | 0 | 59 | 224 | 0 | M9- | 22 – 66 | | 14031334 | PF3535 | IJ | | 1963 |
| 2716 | I | 827 | 0 | 0 | 59 | 224 | 0 | M9- | 22 – 66 | | 15554989 | V3028 | IJ | | 2716 |
| 3217 | I | 827 | 0 | 0 | 59 | 224 | 0 | M9- | 22 – 66 | | 16839810 | FGC1575 | IJ | | 3217 |
| 3300 | I | 827 | 0 | 0 | 59 | 224 | 0 | M9- | 22 – 66 | | 16966024 | FGC1577 | IJ | | 3300 |
| 3309 | I | 827 | 0 | 0 | 59 | 224 | 0 | M9- | 22 – 66 | | 17012180 | YSC0000289 | IJ | | 3309 |
| 3320 | III | 825 | 0 | 2 | 59 | 224 | 0 | M9- | 22 – 66 | | 17040447 | FGC1584 | IJ | | 3320 |
| 3370 | III | 826 | 0 | 1 | 59 | 223 | 1 | M9- | 22 – 66 | MD | 17200869 | FGC1565 | IJ | | 3370 |
| 3496 | III | 825 | 0 | 2 | 59 | 221 | 3 | M9- | 22 – 66 | MD | 17513178 | FGC1576 | IJ | | 3496 |
| 3913 | III | 826 | 0 | 1 | 59 | 224 | 0 | M9- | 22 – 66 | | 18633084 | YSC0001311 | IJ | | 3913 |
| 3966 | III | 826 | 0 | 1 | 59 | 224 | 0 | M9- | 22 – 66 | | 18769674 | Y1894 | IJ | | 3966 |
| 4094 | I | 827 | 0 | 0 | 59 | 224 | 0 | M9- | 22 – 66 | | 19038302 | PF3553 | IJ | | 4094 |
| 4161 | I | 827 | 0 | 0 | 59 | 224 | 0 | M9- | 22 – 66 | | 19166861 | PF3554 | IJ | | 4161 |
| 4419 | I | 827 | 0 | 0 | 59 | 224 | 0 | M9- | 22 – 66 | | 21225770 | P126 | IJ | | 4419 |
| 4525 | I | 827 | 0 | 0 | 59 | 224 | 0 | M9- | 22 – 66 | | 21461202 | PF3562 | IJ | | 4525 |
| 5230 | III | 826 | 0 | 1 | 58 | 224 | 1 | M9- | 22 – 66 | | 22909101 | YSC0001317 | IJ | | 5230 |
| 5313 | I | 827 | 0 | 0 | 59 | 224 | 0 | M9- | 22 – 66 | | 23069523 | S2 | IJ | | 5313 |
| 5670 | III | 825 | 0 | 2 | 59 | 224 | 0 | M9- | 22 – 66 | | 24426007 | PF3568 | IJ | | 5670 |
| 521 | I | 827 | 0 | 0 | 224 | 59 | 0 | M9- | 1 – 21 | | 6753519 | S137 | IJK | | 521 |
| 780 | I | 827 | 0 | 0 | 224 | 59 | 0 | M9- | 1 – 21 | | 7173143 | S138 | IJK | | 780 |
| 936 | I | 827 | 0 | 0 | 224 | 59 | 0 | M9- | 1 – 21 | | 7629583 | Y2571 | IJK | | 936 |
| 981 | I | 827 | 0 | 0 | 224 | 59 | 0 | M9- | 1 – 21 | | 7702973 | YSC0001318 | IJK | | 981 |
| 1033 | III | 827 | 0 | 0 | 222 | 59 | 2 | M9- | 1 – 21 | | 7792789 | YSC0001319 | IJK | | 1033 |
| 4569 | I | 827 | 0 | 0 | 224 | 59 | 0 | M9- | 1 – 21 | | 21571895 | PF3500 | IJK | | 4569 |
| 716 | III | 814 | 0 | 13 | 254 | 23 | 6 | M9- | 60 – 66 | | 7048870 | PF4504.2 | J | A | 716 |
| 722 | I | 827 | 0 | 0 | 260 | 23 | 0 | M9- | 60 – 66 | | 7065239 | YSC0000064 | J | | 722 |
| 749 | I | 827 | 0 | 0 | 260 | 23 | 0 | M9- | 60 – 66 | | 7126936 | YSC0000066 | J | | 749 |
| 783 | I | 827 | 0 | 0 | 260 | 23 | 0 | M9- | 60 – 66 | | 7180976 | PF4508 | J | | 783 |
| 822 | I | 827 | 0 | 0 | 260 | 23 | 0 | M9- | 60 – 66 | | 7292720 | S34 | J | | 822 |
| 823 | I | 827 | 0 | 0 | 260 | 23 | 0 | M9- | 60 – 66 | | 7296343 | YSC0001255 | J | | 823 |
| 1018 | I | 827 | 0 | 0 | 260 | 23 | 0 | M9- | 60 – 66 | | 7759610 | PF4513 | J | | 1018 |
| 1308 | I | 827 | 0 | 0 | 260 | 23 | 0 | M9- | 60 – 66 | | 8393499 | YSC0001296 | J | | 1308 |
| 1527 | I | 827 | 0 | 0 | 260 | 23 | 0 | M9- | 60 – 66 | | 8669451 | PF4519 | J | | 1527 |
| 1540 | I | 827 | 0 | 0 | 260 | 23 | 0 | M9- | 60 – 66 | | 8687145 | PF4520 | J | | 1540 |
| 1732 | I | 825 | 2 | 0 | 260 | 23 | 0 | BOTH | | PI | 9815201 | PF4521 | J | | 1732 |
| 1820 | III | 824 | 0 | 3 | 259 | 23 | 1 | M9- | 60 – 66 | | 10038717 | Z7815 | J | | 1820 |
| 2079 | I | 827 | 0 | 0 | 260 | 23 | 0 | M9- | 60 – 66 | | 14264859 | YSC0001303 | J | | 2079 |

| | | M9+ | | | M9- | | | | | | Build 37 | | ISOGG | | |
|---|---|---|---|---|---|---|---|---|---|---|---|---|---|---|---|
| List | Stage | 0 | 1 | . | 0 | 1 | . | M9? | Clade(s) | Prob. | Position | Name | Subgroup Name | Amb. | List |
| 2156 | I | 827 | 0 | 0 | 260 | 23 | 0 | M9- | 60 – 66 | | 14476551 | PF4538 | J | | 2156 |
| 2303 | I | 827 | 0 | 0 | 260 | 23 | 0 | M9- | 60 – 66 | | 14850935 | PF4539 | J | | 2303 |
| 2558 | I | 827 | 0 | 0 | 260 | 23 | 0 | M9- | 60 – 66 | | 15237163 | PF4542 | J | A | 2558 |
| 2576 | I | 827 | 0 | 0 | 260 | 23 | 0 | M9- | 60 – 66 | | 15277466 | YSC0001275 | J | | 2576 |
| 2759 | III | 826 | 0 | 1 | 260 | 23 | 0 | M9- | 60 – 66 | | 15602183 | PF4547 | J | | 2759 |
| 2929 | III | 811 | 0 | 16 | 252 | 22 | 9 | M9- | 60 – 66 | MD | 16180103 | PF4550 | J | | 2929 |
| 2975 | III | 679 | 0 | 148 | 216 | 23 | 44 | M9- | 60 – 66 | OUT | 16262942 | PF4551 | J | | 2975 |
| 2978 | I | 827 | 0 | 0 | 260 | 23 | 0 | M9- | 60 – 66 | | 16268345 | YSC0000785 | J | | 2978 |
| 3030 | I | 827 | 0 | 0 | 260 | 23 | 0 | M9- | 60 – 66 | | 16401405 | PF4555 | J | | 3030 |
| 3064 | I | 827 | 0 | 0 | 260 | 23 | 0 | M9- | 60 – 66 | | 16475682 | YSC0001305 | J | | 3064 |
| 3347 | I | 827 | 0 | 0 | 260 | 23 | 0 | M9- | 60 – 66 | | 17136854 | PF4560 | J | | 3347 |
| 3388 | I | 827 | 0 | 0 | 260 | 23 | 0 | M9- | 60 – 66 | | 17246058 | PF4562 | J | | 3388 |
| 3434 | I | 827 | 0 | 0 | 260 | 23 | 0 | M9- | 60 – 66 | | 17367321 | YSC0001286 | J | | 3434 |
| 3484 | III | 825 | 0 | 2 | 260 | 23 | 0 | M9- | 60 – 66 | | 17495914 | PF4564 | J | | 3484 |
| 3543 | I | 827 | 0 | 0 | 260 | 23 | 0 | M9- | 60 – 66 | | 17605948 | YSC0000197 | J | | 3543 |
| 3552 | I | 827 | 0 | 0 | 260 | 23 | 0 | M9- | 60 – 66 | | 17637446 | PF4568 | J | | 3552 |
| 3580 | III | 702 | 0 | 125 | 223 | 23 | 37 | M9- | 60 – 66 | | 17693210 | PF4569 | J | | 3580 |
| 3728 | I | 827 | 0 | 0 | 260 | 23 | 0 | M9- | 60 – 66 | | 17996247 | PF4572 | J | | 3728 |
| 3779 | I | 827 | 0 | 0 | 260 | 23 | 0 | M9- | 60 – 66 | | 18084124 | PF4573 | J | | 3779 |
| 3836 | I | 827 | 0 | 0 | 260 | 23 | 0 | M9- | 60 – 66 | | 18257026 | YSC0001309 | J | | 3836 |
| 3869 | I | 827 | 0 | 0 | 260 | 23 | 0 | M9- | 60 – 66 | | 18552360 | PF4576 | J | | 3869 |
| 3876 | I | 827 | 0 | 0 | 260 | 23 | 0 | M9- | 60 – 66 | | 18567169 | PF4577 | J | | 3876 |
| 3937 | I | 827 | 0 | 0 | 260 | 23 | 0 | M9- | 60 – 66 | | 18695159 | PF4579 | J | | 3937 |
| 3970 | I | 827 | 0 | 0 | 260 | 23 | 0 | M9- | 60 – 66 | | 18773505 | PF4580 | J | | 3970 |
| 4167 | III | 827 | 0 | 0 | 260 | 22 | 1 | M9- | 60 – 66 | MD | 19179335 | PF4584 | J | | 4167 |
| 4175 | I | 827 | 0 | 0 | 260 | 23 | 0 | M9- | 60 – 66 | | 19194316 | YSC0001312 | J | | 4175 |
| 4311 | I | 827 | 0 | 0 | 260 | 23 | 0 | M9- | 60 – 66 | | 19508788 | YSC0001313 | J | | 4311 |
| 4375 | I | 827 | 0 | 0 | 260 | 23 | 0 | M9- | 60 – 66 | | 21097847 | PF4588 | J | | 4375 |
| 4444 | I | 827 | 0 | 0 | 260 | 23 | 0 | M9- | 60 – 66 | | 21281892 | PF4591 | J | | 4444 |
| 4766 | I | 827 | 0 | 0 | 260 | 23 | 0 | M9- | 60 – 66 | | 21858778 | PF4595 | J | | 4766 |
| 4881 | I | 827 | 0 | 0 | 260 | 23 | 0 | M9- | 60 – 66 | | 21923739 | Z7822 | J | | 4881 |
| 4959 | III | 826 | 0 | 1 | 259 | 23 | 1 | M9- | 60 – 66 | | 22118776 | PF4598 | J | | 4959 |
| 5114 | I | 827 | 0 | 0 | 260 | 23 | 0 | M9- | 60 – 66 | | 22708656 | YSC0001315 | J | | 5114 |
| 5161 | I | 827 | 0 | 0 | 260 | 23 | 0 | M9- | 60 – 66 | | 22749853 | PF4609 | J | | 5161 |
| 5184 | III | 814 | 0 | 13 | 257 | 23 | 3 | M9- | 60 – 66 | | 22785217 | YSC0001316 | J | | 5184 |
| 5191 | I | 827 | 0 | 0 | 260 | 23 | 0 | M9- | 60 – 66 | | 22808236 | PF4612 | J | | 5191 |
| 5326 | III | 816 | 0 | 11 | 256 | 23 | 4 | M9- | 60 – 66 | | 23088142 | YSC0000236 | J | | 5326 |
| 5362 | I | 827 | 0 | 0 | 260 | 23 | 0 | M9- | 60 – 66 | | 23163701 | PF4617.2 | J | A | 5362 |
| 5389 | III | 827 | 0 | 0 | 259 | 23 | 1 | M9- | 60 – 66 | | 23250894 | YSC0001250 | J | | 5389 |
| 5392 | I | 827 | 0 | 0 | 260 | 23 | 0 | M9- | 60 – 66 | | 23255729 | PF4620 | J | | 5392 |
| 5408 | I | 827 | 0 | 0 | 260 | 23 | 0 | M9- | 60 – 66 | | 23273016 | YSC0001251 | J | | 5408 |
| 5469 | I | 827 | 0 | 0 | 260 | 23 | 0 | M9- | 60 – 66 | | 23443976 | YSC0001253 | J | | 5469 |
| 182 | IV | 825 | 0 | 2 | 283 | 0 | 0 | Mono | Mono | | 2888477 | M369 | J (Notes) | | 182 |
| 903 | IV | 827 | 0 | 0 | 283 | 0 | 0 | Mono | Mono | | 7583254 | M62 | J (Notes) | | 903 |
| 4664 | IV | 827 | 0 | 0 | 283 | 0 | 0 | Mono | Mono | | 21730357 | M163 | J (Notes) | | 4664 |
| 4808 | IV | 827 | 0 | 0 | 283 | 0 | 0 | Mono | Mono | | 21878708 | M289 | J (Notes) | | 4808 |
| 4809 | IV | 827 | 0 | 0 | 283 | 0 | 0 | Mono | Mono | | 21878762 | M280 | J (Notes) | | 4809 |
| 4860 | IV | 827 | 0 | 0 | 283 | 0 | 0 | Mono | Mono | | 21904087 | M327 | J (Notes) | | 4860 |
| 2228 | III | 826 | 0 | 1 | 282 | 1 | 0 | M9- | 60 | | 14705158 | PF4701 | J1 | | 2228 |
| 4267 | III | 827 | 0 | 0 | 281 | 1 | 1 | M9- | 60 | | 19383622 | F4287 | J1 | | 4267 |
| 4300 | II | 827 | 0 | 0 | 282 | 1 | 0 | M9- | 60 | | 19489654 | YSC0001013 | J1 | | 4300 |
| 5144 | II | 827 | 0 | 0 | 282 | 1 | 0 | M9- | 60 | | 22741818 | PF4782 | J1 | | 5144 |
| 2959 | II | 827 | 0 | 0 | 282 | 1 | 0 | M9- | 60 | | 16227302 | Z2215 | J1a | | 2959 |
| 4141 | IV | 825 | 0 | 2 | 282 | 0 | 1 | Mono | Mono | | 19136821 | S4972 | J1a2a1a2c1a | | 4141 |
| 634 | IV | 827 | 0 | 0 | 283 | 0 | 0 | Mono | Mono | | 6932106 | L816 | J1a2a1a2c1a1 | | 634 |
| 4051 | III | 826 | 1 | 0 | 282 | 0 | 1 | M9+ | 1 | | 18937926 | CTS9501 | J1a2a1a2d2b2b2c4d2a2a5a1e4a5b1~ | W | 4051 |
| 29 | IV | 826 | 0 | 1 | 283 | 0 | 0 | Mono | Mono | | 2669716 | Z1828 | J1a2b | | 29 |
| 5179 | IV | 827 | 0 | 0 | 283 | 0 | 0 | Mono | Mono | | 22770232 | Y29697 | J1a3~ | A | 5179 |
| 3623 | IV | 826 | 0 | 1 | 254 | 0 | 29 | Mono | Mono | | 17792637 | F2594.2 | J1a3a~ | A | 3623 |
| 93 | IV | 827 | 0 | 0 | 283 | 0 | 0 | Mono | Mono | | 2757670 | V189 | J1a3b~ | A | 93 |
| 725 | I | 827 | 0 | 0 | 261 | 22 | 0 | M9- | 61 – 66 | | 7068942 | PF4893 | J2 | | 725 |
| 1026 | I | 827 | 0 | 0 | 261 | 22 | 0 | M9- | 61 – 66 | | 7771358 | S321 | J2 | | 1026 |
| 1917 | III | 810 | 0 | 17 | 257 | 22 | 4 | M9- | 61 – 66 | | 13832032 | PF4905 | J2 | | 1917 |
| 2410 | I | 827 | 0 | 0 | 261 | 22 | 0 | M9- | 61 – 66 | | 14969634 | PF4908 | J2 | | 2410 |
| 3057 | III | 826 | 0 | 1 | 261 | 22 | 0 | M9- | 61 – 66 | | 16459438 | PF4912 | J2 | | 3057 |
| 3095 | I | 827 | 0 | 0 | 261 | 22 | 0 | M9- | 61 – 66 | | 16542685 | PF4913 | J2 | | 3095 |
| 4073 | III | 827 | 0 | 0 | 260 | 22 | 1 | M9- | 61 – 66 | | 18978214 | PF4920 | J2 | | 4073 |
| 4172 | I | 827 | 0 | 0 | 261 | 22 | 0 | M9- | 61 – 66 | | 19181063 | PF4922 | J2 | | 4172 |
| 4454 | I | 827 | 0 | 0 | 261 | 22 | 0 | M9- | 61 – 66 | | 21292707 | PF4925 | J2 | | 4454 |
| 4930 | III | 826 | 0 | 1 | 261 | 22 | 0 | M9- | 61 – 66 | | 22024569 | PF4597 | J2 | | 4930 |
| 5107 | I | 827 | 0 | 0 | 261 | 22 | 0 | M9- | 61 – 66 | | 22687035 | PF4933 | J2 | | 5107 |
| 1282 | III | 820 | 0 | 7 | 273 | 9 | 1 | M9- | 61 – 64 | | 8351025 | PF4897 | J2a | | 1282 |
| 1319 | III | 809 | 0 | 18 | 269 | 9 | 5 | M9- | 61 – 64 | | 8417425 | PF4956 | J2a | | 1319 |
| 2802 | I | 827 | 0 | 0 | 274 | 9 | 0 | M9- | 61 – 64 | | 15704139 | PF4966 | J2a | | 2802 |
| 2941 | III | 825 | 0 | 2 | 274 | 9 | 0 | M9- | 61 – 64 | | 16196053 | PF4967 | J2a | | 2941 |
| 2988 | I | 827 | 0 | 0 | 274 | 9 | 0 | M9- | 61 – 64 | | 16310787 | PF4968 | J2a | | 2988 |
| 3315 | I | 827 | 0 | 0 | 274 | 9 | 0 | M9- | 61 – 64 | | 17032059 | F4221 | J2a | | 3315 |
| 3365 | III | 825 | 0 | 2 | 274 | 9 | 0 | M9- | 61 – 64 | | 17191877 | PF4972 | J2a | | 3365 |
| 3683 | I | 827 | 0 | 0 | 274 | 9 | 0 | M9- | 61 – 64 | | 17904970 | PF4974 | J2a | | 3683 |
| 3905 | I | 827 | 0 | 0 | 274 | 9 | 0 | M9- | 61 – 64 | | 18615144 | PF4976 | J2a | | 3905 |
| 3969 | I | 827 | 0 | 0 | 274 | 9 | 0 | M9- | 61 – 64 | | 18772238 | PF4918 | J2a | | 3969 |
| 4631 | I | 827 | 0 | 0 | 274 | 9 | 0 | M9- | 61 – 64 | | 21674327 | PF4986 | J2a | | 4631 |
| 4907 | I | 827 | 0 | 0 | 274 | 9 | 0 | M9- | 61 – 64 | | 21970721 | PF4987 | J2a | | 4907 |
| 5007 | IV | 0 | 827 | 0 | 0 | 282 | 1 | Mono | Mono | | 22243566 | L152 | J2a | W | 5007 |
| 5116 | I | 827 | 0 | 0 | 274 | 9 | 0 | M9- | 61 – 64 | | 22711465 | PF4988 | J2a | | 5116 |
| 5549 | I | 827 | 0 | 0 | 274 | 9 | 0 | M9- | 61 – 64 | | 23748578 | PF4990 | J2a | | 5549 |
| 1321 | I | 827 | 0 | 0 | 274 | 9 | 0 | M9- | 61 – 64 | | 8420738 | PF4957 | J2a~ | | 1321 |
| 2204 | III | 827 | 0 | 0 | 274 | 7 | 2 | M9- | 61 – 64 | MD | 14628068 | PF4962 | J2a~ | | 2204 |
| 1365 | III | 822 | 0 | 5 | 273 | 9 | 1 | M9- | 61 – 64 | | 8465758 | F1227 | J2a1 | | 1365 |
| 3548 | I | 827 | 0 | 0 | 274 | 9 | 0 | M9- | 61 – 64 | | 17614366 | PF5100 | J2a1 | A | 3548 |
| 4348 | III | 824 | 0 | 3 | 274 | 9 | 0 | M9- | 61 – 64 | | 20836109 | PF5105 | J2a1 | | 4348 |
| 5257 | I | 827 | 0 | 0 | 274 | 9 | 0 | M9- | 61 – 64 | | 22942897 | S57 | J2a1a | | 5257 |
| 5289 | III | 826 | 0 | 1 | 273 | 9 | 1 | M9- | 61 – 64 | | 23021978 | S396 | J2a1a | | 5289 |
| 2662 | IV | 827 | 0 | 0 | 283 | 0 | 0 | Mono | Mono | | 15469740 | M322 | J2a1a1a2a2b | | 2662 |
| 4691 | IV | 827 | 0 | 0 | 283 | 0 | 0 | Mono | Mono | | 21751330 | M47 | J2a1a1a2a2b | | 4691 |
| 4810 | I | 827 | 0 | 0 | 279 | 4 | 0 | M9- | 62 – 63 | | 21878809 | S51 | J2a1a1a2b2 | | 4810 |

| | | M9+ | | | M9- | | | | | | Build 37 | | ISOGG | | |
|---|---|---|---|---|---|---|---|---|---|---|---|---|---|---|---|
| List | Stage | 0 | 1 | . | 0 | 1 | . | M9? | Clade(s) | Prob. | Position | Name | Subgroup Name | Amb. | List |
| 2460 | III | 822 | 0 | 5 | 280 | 2 | 1 | M9- | 63 | | 15025506 | Page14 | J2a1a1a2b2a1a | | 2460 |
| 4859 | II | 827 | 0 | 0 | 281 | 2 | 0 | M9- | 63 | | 21904023 | M92 | J2a1a1a2b2a1a | | 4859 |
| 4718 | IV | 826 | 0 | 1 | 283 | 0 | 0 | Mono | Mono | | 21764694 | M166 | J2a1a1a2b2a2b1b2 | | 4718 |
| 3537 | IV | 827 | 0 | 0 | 283 | 0 | 0 | Mono | Mono | | 17595842 | L88.2 | J2a1a1b1a1a1a~ | | 3537 |
| 3771 | IV | 827 | 0 | 0 | 283 | 0 | 0 | Mono | Mono | | 18077016 | S287 | J2a1a1b2a1b1 | | 3771 |
| 5123 | IV | 827 | 0 | 0 | 282 | 0 | 1 | Mono | Mono | | 22725065 | PF5446 | J2a1a1b2a1b1 | | 5123 |
| 5382 | IV | 827 | 0 | 0 | 283 | 0 | 0 | Mono | Mono | | 23243897 | S320 | J2a1a1b2a1b1 | | 5382 |
| 5152 | IV | 825 | 0 | 2 | 283 | 0 | 0 | Mono | Mono | | 22744993 | M318 | J2a1a1b2a1b1b2b1 | | 5152 |
| 2087 | II | 827 | 0 | 0 | 282 | 1 | 0 | M9- | 64 | | 14286528 | S286 | J2a1a4 | | 2087 |
| 4645 | IV | 827 | 0 | 0 | 283 | 0 | 0 | Mono | Mono | | 21716366 | M158.2 | J2a1a4a | | 4645 |
| 4142 | II | 827 | 0 | 0 | 282 | 1 | 0 | M9- | 64 | | 19136822 | S399 | J2a1a4b | | 4142 |
| 4713 | IV | 827 | 0 | 0 | 283 | 0 | 0 | Mono | Mono | | 21764458 | M137 | J2a1a4b1a1a~ | | 4713 |
| 2315 | IV | 827 | 0 | 0 | 283 | 0 | 0 | Mono | Mono | | 14870114 | S398 | J2a2a | | 2315 |
| 175 | IV | 827 | 0 | 0 | 283 | 0 | 0 | Mono | Mono | | 2881377 | PF5065 | J2a2a1a1a2 | | 175 |
| 4807 | IV | 827 | 0 | 0 | 283 | 0 | 0 | Mono | Mono | | 21878699 | M340 | J2a2a1a1a2b1 | | 4807 |
| 1138 | III | 823 | 1 | 3 | 267 | 13 | 3 | BOTH | | PI | 8032311 | Z526.2 | J2b | A | 1138 |
| 2631 | I | 827 | 0 | 0 | 270 | 13 | 0 | M9- | 65 – 66 | | 15436316 | M221 | J2b | | 2631 |
| 4884 | III | 826 | 0 | 1 | 270 | 12 | 1 | M9- | 65 – 66 | MD | 21926112 | M102 | J2b | | 4884 |
| 5170 | I | 827 | 0 | 0 | 270 | 13 | 0 | M9- | 65 – 66 | | 22753079 | PF4939 | J2b | | 5170 |
| 2736 | IV | 827 | 0 | 0 | 283 | 0 | 0 | Mono | Mono | | 15587509 | M205 | J2b1 | | 2736 |
| 2440 | I | 827 | 0 | 0 | 270 | 13 | 0 | M9- | 65 – 66 | | 15018459 | M241 | J2b2a | | 2440 |
| 643 | III | 824 | 0 | 3 | 273 | 10 | 0 | M9- | 66 | | 6932663 | L283 | J2b2a1 | | 643 |
| 4535 | III | 821 | 0 | 6 | 272 | 6 | 5 | M9- | 66 | MD | 21490912 | Z1297 | J2b2a1a1a1a1a | | 4535 |
| 4663 | I | 827 | 0 | 0 | 0 | 283 | 0 | M9+/- | M9+/- | | 21730257 | PF5506 | K | | 4663 |
| 1532 | I | 827 | 0 | 0 | 0 | 283 | 0 | M9+/- | M9+/- | | 8679843 | PF5480 | K~ | | 1532 |
| 2668 | I | 827 | 0 | 0 | 0 | 283 | 0 | M9+/- | M9+/- | | 15472863 | PF5493 | K~ | | 2668 |
| 4349 | I | 827 | 0 | 0 | 0 | 283 | 0 | M9+/- | M9+/- | | 20837553 | PF5504 | K~ | | 4349 |
| 2955 | I | 824 | 3 | 0 | 0 | 283 | 0 | M9+ | 3 – 93 | | 16222561 | PF5911 | K2b | | 2955 |
| 1946 | IV | 827 | 0 | 0 | 283 | 0 | 0 | Mono | Mono | | 13992279 | P261 | K2c | | 1946 |
| 4404 | IV | 827 | 0 | 0 | 283 | 0 | 0 | Mono | Mono | | 21165969 | P263 | K2c | | 4404 |
| 5004 | IV | 827 | 0 | 0 | 283 | 0 | 0 | Mono | Mono | | 22214480 | P403 | K2d | | 5004 |
| 102 | IV | 826 | 0 | 1 | 283 | 0 | 0 | Mono | Mono | | 2774249 | PF5683 | L | | 102 |
| 580 | IV | 827 | 0 | 0 | 283 | 0 | 0 | Mono | Mono | | 6867589 | PF5693 | L | | 580 |
| 590 | IV | 827 | 0 | 0 | 283 | 0 | 0 | Mono | Mono | | 6881269 | PF5694 | L | | 590 |
| 623 | IV | 827 | 0 | 0 | 283 | 0 | 0 | Mono | Mono | | 6926083 | PF5695 | L | | 623 |
| 626 | IV | 794 | 0 | 33 | 267 | 0 | 16 | Mono | Mono | | 6931538 | L656 | L | | 626 |
| 838 | IV | 827 | 0 | 0 | 283 | 0 | 0 | Mono | Mono | | 7332400 | PF5699 | L | | 838 |
| 1057 | IV | 825 | 0 | 2 | 282 | 0 | 1 | Mono | Mono | | 7842405 | PF5705 | L | | 1057 |
| 1232 | IV | 826 | 0 | 1 | 282 | 0 | 1 | Mono | Mono | | 8246462 | PF5528 | L | | 1232 |
| 1233 | IV | 813 | 0 | 14 | 280 | 0 | 3 | Mono | Mono | | 8246586 | PF5715 | L | | 1233 |
| 1247 | IV | 826 | 0 | 1 | 283 | 0 | 0 | Mono | Mono | | 8270375 | PF5717 | L | | 1247 |
| 1511 | IV | 827 | 0 | 0 | 283 | 0 | 0 | Mono | Mono | | 8636585 | PF5532 | L | | 1511 |
| 1514 | IV | 827 | 0 | 0 | 283 | 0 | 0 | Mono | Mono | | 8640774 | PF5723 | L | | 1514 |
| 1691 | IV | 827 | 0 | 0 | 282 | 0 | 1 | Mono | Mono | | 9445206 | PF5729 | L | | 1691 |
| 1976 | IV | 826 | 0 | 1 | 283 | 0 | 0 | Mono | Mono | | 14060111 | PF5737 | L | | 1976 |
| 2069 | IV | 827 | 0 | 0 | 283 | 0 | 0 | Mono | Mono | | 14240717 | PF5743 | L | | 2069 |
| 2123 | IV | 827 | 0 | 0 | 283 | 0 | 0 | Mono | Mono | | 14371220 | PF5747 | L | | 2123 |
| 2353 | IV | 827 | 0 | 0 | 283 | 0 | 0 | Mono | Mono | | 14904859 | PF5755 | L | | 2353 |
| 2504 | IV | 769 | 0 | 58 | 260 | 0 | 23 | Mono | Mono | | 15070860 | PF5756 | L | | 2504 |
| 2537 | IV | 827 | 0 | 0 | 283 | 0 | 0 | Mono | Mono | | 15165158 | PF5757 | L | | 2537 |
| 2968 | IV | 827 | 0 | 0 | 283 | 0 | 0 | Mono | Mono | | 16252528 | PF5771 | L | | 2968 |
| 3541 | IV | 827 | 0 | 0 | 281 | 0 | 2 | Mono | Mono | | 17602759 | PF5791 | L | | 3541 |
| 3696 | IV | 827 | 0 | 0 | 283 | 0 | 0 | Mono | Mono | | 17921146 | PF5557 | L | | 3696 |
| 3701 | IV | 827 | 0 | 0 | 283 | 0 | 0 | Mono | Mono | | 17929784 | PF5802 | L | | 3701 |
| 3740 | IV | 825 | 0 | 2 | 283 | 0 | 0 | Mono | Mono | | 18029588 | PF5804 | L | | 3740 |
| 3758 | IV | 821 | 0 | 6 | 283 | 0 | 0 | Mono | Mono | | 18059295 | PF5806 | L | | 3758 |
| 3964 | IV | 827 | 0 | 0 | 283 | 0 | 0 | Mono | Mono | | 18765272 | PF5810 | L | | 3964 |
| 4084 | IV | 826 | 0 | 1 | 283 | 0 | 0 | Mono | Mono | | 19010882 | PF5813 | L | | 4084 |
| 4484 | IV | 827 | 0 | 0 | 283 | 0 | 0 | Mono | Mono | | 21366301 | PF5823 | L | | 4484 |
| 4665 | IV | 827 | 0 | 0 | 283 | 0 | 0 | Mono | Mono | | 21730647 | M11 | L | | 4665 |
| 4693 | IV | 827 | 0 | 0 | 283 | 0 | 0 | Mono | Mono | | 21751449 | Page43 | L | | 4693 |
| 5073 | IV | 827 | 0 | 0 | 283 | 0 | 0 | Mono | Mono | | 22595505 | PF5829 | L | | 5073 |
| 5127 | IV | 827 | 0 | 0 | 283 | 0 | 0 | Mono | Mono | | 22730070 | PF5830 | L | | 5127 |
| 5284 | IV | 826 | 0 | 1 | 281 | 0 | 2 | Mono | Mono | | 23010047 | L855 | L | | 5284 |
| 5304 | IV | 827 | 0 | 0 | 283 | 0 | 0 | Mono | Mono | | 23058840 | PF5832 | L | | 5304 |
| 5324 | IV | 827 | 0 | 0 | 283 | 0 | 0 | Mono | Mono | | 23081617 | PF5833 | L | A | 5324 |
| 5368 | IV | 827 | 0 | 0 | 283 | 0 | 0 | Mono | Mono | | 23184728 | PF5583 | L | | 5368 |
| 5426 | IV | 827 | 0 | 0 | 283 | 0 | 0 | Mono | Mono | | 23337218 | PF5837 | L | | 5426 |
| 5466 | IV | 827 | 0 | 0 | 283 | 0 | 0 | Mono | Mono | | 23437408 | PF5839 | L | | 5466 |
| 5579 | IV | 827 | 0 | 0 | 283 | 0 | 0 | Mono | Mono | | 23813544 | PF5841 | L | | 5579 |
| 5663 | IV | 825 | 0 | 2 | 283 | 0 | 0 | Mono | Mono | | 24406600 | PF5842 | L | | 5663 |
| 4670 | IV | 827 | 0 | 0 | 283 | 0 | 0 | Mono | Mono | | 21733454 | PF5570 | L or K1a | A | 4670 |
| 3486 | IV | 827 | 0 | 0 | 283 | 0 | 0 | Mono | Mono | | 17507398 | PF5790 | L~ | A | 3486 |
| 2262 | IV | 827 | 0 | 0 | 283 | 0 | 0 | Mono | Mono | | 14790163 | M22 | L1 | | 2262 |
| 3770 | IV | 827 | 0 | 0 | 283 | 0 | 0 | Mono | Mono | | 18076957 | L1304 | L1 | A | 3770 |
| 5153 | IV | 827 | 0 | 0 | 283 | 0 | 0 | Mono | Mono | | 22745051 | M295 | L1 | | 5153 |
| 3042 | IV | 827 | 0 | 0 | 283 | 0 | 0 | Mono | Mono | | 16418020 | Z20163 | L1a | | 3042 |
| 3517 | IV | 827 | 0 | 0 | 283 | 0 | 0 | Mono | Mono | | 17564121 | M2565 | L1a | | 3517 |
| 4303 | IV | 827 | 0 | 0 | 283 | 0 | 0 | Mono | Mono | | 19492036 | Z20168 | L1a | | 4303 |
| 4320 | IV | 827 | 0 | 0 | 283 | 0 | 0 | Mono | Mono | | 19542995 | M2605 | L1a | | 4320 |
| 5487 | IV | 827 | 0 | 0 | 283 | 0 | 0 | Mono | Mono | | 23489497 | M2664 | L1a | | 5487 |
| 1368 | II | 827 | 0 | 0 | 282 | 1 | 0 | M9- | NU | | 8466781 | P329 | L1a1 | W | 1368 |
| 1400 | IV | 827 | 0 | 0 | 283 | 0 | 0 | Mono | Mono | | 8485823 | L1319 | L1a1 | | 1400 |
| 4677 | IV | 827 | 0 | 0 | 283 | 0 | 0 | Mono | Mono | | 21739646 | M27 | L1a1 | | 4677 |
| 4700 | IV | 827 | 0 | 0 | 283 | 0 | 0 | Mono | Mono | | 21757316 | M76 | L1a1 | | 4700 |
| 179 | IV | 827 | 0 | 0 | 283 | 0 | 0 | Mono | Mono | | 2888252 | M357 | L1a2 | | 179 |
| 3257 | IV | 827 | 0 | 0 | 282 | 0 | 1 | Mono | Mono | | 16900247 | M5710 | L1a2 | | 3257 |
| 53 | IV | 827 | 0 | 0 | 283 | 0 | 0 | Mono | Mono | | 2713589 | Page113 | L1b1 | | 53 |
| 189 | IV | 826 | 0 | 1 | 283 | 0 | 0 | Mono | Mono | | 2893277 | M349 | L1b1 | | 189 |
| 4831 | IV | 827 | 0 | 0 | 283 | 0 | 0 | Mono | Mono | | 21893864 | Page116 | L1b1a | | 4831 |
| 5133 | IV | 827 | 0 | 0 | 283 | 0 | 0 | Mono | Mono | | 22737801 | M274 | L1b2a | | 5133 |
| 4200 | IV | 827 | 0 | 0 | 283 | 0 | 0 | Mono | Mono | | 19235022 | PH4366.1 | L1b2b | A | 4200 |
| 4678 | IV | 827 | 0 | 0 | 283 | 0 | 0 | Mono | Mono | | 21739651 | L595 | L2 | | 4678 |
| 701 | III | 824 | 1 | 2 | 283 | 0 | 0 | M9+ | 94 | | 6994764 | PF5525 | LT [K1] | | 701 |
| 1129 | III | 823 | 1 | 3 | 281 | 0 | 2 | M9+ | 94 | 1H | 8015672 | PF5526 | LT [K1] | | 1129 |
| 1373 | II | 826 | 1 | 0 | 283 | 0 | 0 | M9+ | 94 | | 8467290 | P326 | LT [K1] | A | 1373 |

| | | | M9+ | | | M9- | | | | | | Build 37 | | ISOGG | |
|---|---|---|---|---|---|---|---|---|---|---|---|---|---|---|---|
| List | Stage | 0 | 1 | . | 0 | 1 | . | M9? | Clade(s) | Prob. | Position | Name | Subgroup Name | Amb. | List |
| 1504 | III | 825 | 1 | 1 | 283 | 0 | 0 | M9+ | 94 | | 8628308 | PF5531 | LT [K1] | | 1504 |
| 2183 | II | 826 | 1 | 0 | 283 | 0 | 0 | M9+ | 94 | | 14542883 | PF5543 | LT [K1] | | 2183 |
| 2513 | II | 826 | 1 | 0 | 283 | 0 | 0 | M9+ | 94 | | 15094275 | CTS3648 | LT [K1] | | 2513 |
| 2847 | II | 826 | 1 | 0 | 283 | 0 | 0 | M9+ | 94 | | 15811928 | PF5548 | LT [K1] | | 2847 |
| 3358 | III | 805 | 1 | 21 | 278 | 0 | 5 | M9+ | 94 | | 17173391 | CTS6888 | LT [K1] | | 3358 |
| 4505 | II | 826 | 1 | 0 | 283 | 0 | 0 | M9+ | 94 | | 21406133 | PF5566 | LT [K1] | | 4505 |
| 5473 | II | 826 | 1 | 0 | 283 | 0 | 0 | M9+ | 94 | | 23454053 | PF5586 | LT [K1] | | 5473 |
| 2673 | IV | 0 | 0 | 827 | 0 | 0 | 283 | Mono | Mono | | 15481435 | S322 | M | | 2673 |
| 507 | IV | 827 | 0 | 0 | 283 | 0 | 0 | Mono | Mono | | 6736812 | Page1 | M1 | | 507 |
| 2388 | IV | 827 | 0 | 0 | 283 | 0 | 0 | Mono | Mono | | 14945705 | M189 | M1 | | 2388 |
| 4580 | IV | 827 | 0 | 0 | 283 | 0 | 0 | Mono | Mono | | 21609946 | P73 | M1 | | 4580 |
| 4773 | IV | 827 | 0 | 0 | 282 | 0 | 1 | Mono | Mono | | 21866424 | M106 | M1 | | 4773 |
| 2447 | IV | 827 | 0 | 0 | 283 | 0 | 0 | Mono | Mono | | 15019719 | P87 | M1a1a2a | | 2447 |
| 1499 | IV | 827 | 0 | 0 | 281 | 0 | 2 | Mono | Mono | | 8620324 | P22_2^^ | M1a1a2a1 | | 1499 |
| 2164 | IV | 827 | 0 | 0 | 283 | 0 | 0 | Mono | Mono | | 14492042 | P51 | M1a1a3a~ | | 2164 |
| 3574 | IV | 827 | 0 | 0 | 283 | 0 | 0 | Mono | Mono | | 17678803 | Page67 | M1a2a2a1~ | | 3574 |
| 16 | IV | 827 | 0 | 0 | 283 | 0 | 0 | Mono | Mono | | 2658869 | SRY9138^^ | M2a | | 16 |
| 991 | IV | 827 | 0 | 0 | 283 | 0 | 0 | Mono | Mono | | 7722315 | M2144 | N | | 991 |
| 999 | IV | 827 | 0 | 0 | 283 | 0 | 0 | Mono | Mono | | 7731706 | V1392 | N | | 999 |
| 1079 | IV | 827 | 0 | 0 | 283 | 0 | 0 | Mono | Mono | | 7891525 | V1495 | N | | 1079 |
| 1503 | IV | 827 | 0 | 0 | 283 | 0 | 0 | Mono | Mono | | 8623586 | V2374 | N | | 1503 |
| 1569 | IV | 813 | 0 | 14 | 272 | 0 | 11 | Mono | Mono | | 8771750 | M2161 | N | | 1569 |
| 2035 | IV | 827 | 0 | 0 | 283 | 0 | 0 | Mono | Mono | | 14174283 | S15571 | N | | 2035 |
| 2073 | IV | 814 | 0 | 13 | 282 | 0 | 1 | Mono | Mono | | 14244937 | M2169 | N | | 2073 |
| 2214 | IV | 824 | 0 | 3 | 281 | 0 | 2 | Mono | Mono | | 14647154 | V2528 | N | | 2214 |
| 2380 | IV | 827 | 0 | 0 | 283 | 0 | 0 | Mono | Mono | | 14935686 | V2700 | N | | 2380 |
| 2639 | IV | 827 | 0 | 0 | 283 | 0 | 0 | Mono | Mono | | 15437152 | V2948 | N | | 2639 |
| 2645 | IV | 827 | 0 | 0 | 283 | 0 | 0 | Mono | Mono | | 15439570 | V2951 | N | | 2645 |
| 2949 | IV | 827 | 0 | 0 | 283 | 0 | 0 | Mono | Mono | | 16203382 | M2200 | N | | 2949 |
| 3103 | IV | 827 | 0 | 0 | 283 | 0 | 0 | Mono | Mono | | 16560334 | S18745 | N | | 3103 |
| 3489 | IV | 827 | 0 | 0 | 283 | 0 | 0 | Mono | Mono | | 17509176 | S19819 | N | | 3489 |
| 3585 | IV | 788 | 0 | 39 | 275 | 0 | 8 | Mono | Mono | | 17710924 | S20099 | N | | 3585 |
| 3629 | IV | 827 | 0 | 0 | 283 | 0 | 0 | Mono | Mono | | 17800868 | S20231^ | N | | 3629 |
| 3695 | IV | 826 | 0 | 1 | 283 | 0 | 0 | Mono | Mono | | 17920486 | M2237 | N | | 3695 |
| 3760 | IV | 827 | 0 | 0 | 283 | 0 | 0 | Mono | Mono | | 18063802 | S20571^ | N | | 3760 |
| 4081 | IV | 827 | 0 | 0 | 283 | 0 | 0 | Mono | Mono | | 19004180 | V3683 | N | | 4081 |
| 4217 | IV | 827 | 0 | 0 | 283 | 0 | 0 | Mono | Mono | | 19269306 | V3957 | N | | 4217 |
| 4383 | IV | 827 | 0 | 0 | 283 | 0 | 0 | Mono | Mono | | 21116317 | S22586 | N | | 4383 |
| 4626 | IV | 827 | 0 | 0 | 283 | 0 | 0 | Mono | Mono | | 21666182 | S23418 | N | | 4626 |
| 4641 | IV | 824 | 0 | 3 | 282 | 0 | 1 | Mono | Mono | | 21707315 | M2270^ | N | | 4641 |
| 4987 | IV | 827 | 0 | 0 | 283 | 0 | 0 | Mono | Mono | | 22175388 | M2275 | N | | 4987 |
| 4999 | IV | 826 | 0 | 1 | 282 | 0 | 1 | Mono | Mono | | 22203556 | S24122 | N | | 4999 |
| 5396 | IV | 821 | 0 | 6 | 281 | 0 | 2 | Mono | Mono | | 23258160 | M2294 | N | | 5396 |
| 5425 | IV | 827 | 0 | 0 | 283 | 0 | 0 | Mono | Mono | | 23328431 | S25467 | N | | 5425 |
| 2661 | IV | 827 | 0 | 0 | 283 | 0 | 0 | Mono | Mono | | 15469724 | Page91 | N or K2a1a | A | 2661 |
| 1040 | IV | 827 | 0 | 0 | 283 | 0 | 0 | Mono | Mono | | 7806898 | M2146 | N~ | | 1040 |
| 1406 | IV | 827 | 0 | 0 | 283 | 0 | 0 | Mono | Mono | | 8496195 | V2137 | N~ | | 1406 |
| 2694 | IV | 827 | 0 | 0 | 283 | 0 | 0 | Mono | Mono | | 15517954 | V3006 | N~ | | 2694 |
| 5678 | IV | 827 | 0 | 0 | 283 | 0 | 0 | Mono | Mono | | 24451187 | M2302 | N~ | | 5678 |
| 471 | IV | 826 | 0 | 1 | 283 | 0 | 0 | Mono | Mono | | 6660143 | Z4835^ | N1 | | 471 |
| 856 | IV | 824 | 0 | 3 | 282 | 0 | 1 | Mono | Mono | | 7415220 | M2142 | N1 | | 856 |
| 2265 | IV | 827 | 0 | 0 | 283 | 0 | 0 | Mono | Mono | | 14792418 | V2603 | N1 | | 2265 |
| 2896 | IV | 827 | 0 | 0 | 283 | 0 | 0 | Mono | Mono | | 15955962 | V3270 | N1 | | 2896 |
| 3031 | IV | 824 | 0 | 3 | 280 | 0 | 3 | Mono | Mono | | 16402912 | M2202 | N1 | | 3031 |
| 3367 | IV | 827 | 0 | 0 | 282 | 0 | 1 | Mono | Mono | | 17192238 | S19333 | N1 | | 3367 |
| 3579 | IV | 827 | 0 | 0 | 283 | 0 | 0 | Mono | Mono | | 17692975 | S20081 | N1 | | 3579 |
| 3631 | IV | 827 | 0 | 0 | 283 | 0 | 0 | Mono | Mono | | 17803021 | M2234 | N1 | | 3631 |
| 4221 | IV | 827 | 0 | 0 | 283 | 0 | 0 | Mono | Mono | | 19274282 | V3963 | N1 | | 4221 |
| 4370 | IV | 827 | 0 | 0 | 283 | 0 | 0 | Mono | Mono | | 21087879 | S22547 | N1 | | 4370 |
| 5481 | IV | 827 | 0 | 0 | 283 | 0 | 0 | Mono | Mono | | 23479535 | S25619 | N1 | | 5481 |
| 5553 | IV | 827 | 0 | 0 | 283 | 0 | 0 | Mono | Mono | | 23755132 | M2299 | N1 | | 5553 |
| 2621 | IV | 827 | 0 | 0 | 283 | 0 | 0 | Mono | Mono | | 15409274 | V2928 | N1~ | | 2621 |
| 5242 | IV | 827 | 0 | 0 | 283 | 0 | 0 | Mono | Mono | | 22921768 | S323 | N1~ | A | 5242 |
| 5347 | IV | 826 | 0 | 1 | 283 | 0 | 0 | Mono | Mono | | 23139617 | M2291 | N1~ | | 5347 |
| 1343 | IV | 827 | 0 | 0 | 283 | 0 | 0 | Mono | Mono | | 8440417 | S11466 | N1a | | 1343 |
| 5214 | IV | 827 | 0 | 0 | 283 | 0 | 0 | Mono | Mono | | 22859747 | S24798 | N1a | | 5214 |
| 2365 | IV | 827 | 0 | 0 | 283 | 0 | 0 | Mono | Mono | | 14922583 | Tat | N1a1 | | 2365 |
| 4102 | IV | 827 | 0 | 0 | 282 | 0 | 1 | Mono | Mono | | 19048725 | M2080 | N1a1 | | 4102 |
| 4686 | IV | 827 | 0 | 0 | 283 | 0 | 0 | Mono | Mono | | 21741755 | M178 | N1a1a | | 4686 |
| 2206 | IV | 827 | 0 | 0 | 283 | 0 | 0 | Mono | Mono | | 14636086 | Z1953 | N1a1a~ | A | 2206 |
| 1582 | IV | 827 | 0 | 0 | 283 | 0 | 0 | Mono | Mono | | 8804414 | M2017 | N1a1a1~ | | 1582 |
| 935 | IV | 827 | 0 | 0 | 283 | 0 | 0 | Mono | Mono | | 7629512 | Z1951 | N1a1a1a | | 935 |
| 5273 | IV | 822 | 0 | 5 | 283 | 0 | 0 | Mono | Mono | | 22974783 | L839 | N1a1a1a | | 5273 |
| 4595 | IV | 827 | 0 | 0 | 283 | 0 | 0 | Mono | Mono | | 21621564 | P298 | N1a1a1a1 | | 4595 |
| 483 | IV | 827 | 0 | 0 | 283 | 0 | 0 | Mono | Mono | | 6681274 | S9317^ | N1a1a1a1a | | 483 |
| 1486 | IV | 827 | 0 | 0 | 282 | 0 | 1 | Mono | Mono | | 8602985 | S11709^ | N1a1a1a1a | | 1486 |
| 3280 | IV | 765 | 0 | 62 | 163 | 0 | 120 | Mono | Mono | | 16939816 | Z1961 | N1a1a1a1a | | 3280 |
| 3604 | IV | 827 | 0 | 0 | 283 | 0 | 0 | Mono | Mono | | 17761893 | S20163 | N1a1a1a1a | | 3604 |
| 4213 | IV | 827 | 0 | 0 | 283 | 0 | 0 | Mono | Mono | | 19260818 | M2083 | N1a1a1a1a | | 4213 |
| 5147 | IV | 827 | 0 | 0 | 283 | 0 | 0 | Mono | Mono | | 22743701 | S24635 | N1a1a1a1a | | 5147 |
| 5617 | IV | 825 | 0 | 2 | 283 | 0 | 0 | Mono | Mono | | 23964507 | M2124 | N1a1a1a1a | | 5617 |
| 2189 | IV | 827 | 0 | 0 | 283 | 0 | 0 | Mono | Mono | | 14570424 | VL29 | N1a1a1a1a1a | | 2189 |
| 4448 | IV | 827 | 0 | 0 | 283 | 0 | 0 | Mono | Mono | | 21286248 | S431 | N1a1a1a1a1a1a | | 4448 |
| 2016 | IV | 827 | 0 | 0 | 283 | 0 | 0 | Mono | Mono | | 14126006 | L1025 | N1a1a1a1a1a1a1 | | 2016 |
| 477 | III | 824 | 3 | 0 | 1 | 245 | 37 | BOTH | | PI | 6670461 | L58 | N1a1a1a1a1a1a2a1~ | w | 477 |
| 2643 | IV | 826 | 0 | 1 | 283 | 0 | 0 | Mono | Mono | | 15438219 | FGC14543 | N1a1a1a1a1a1a6a~ | | 2643 |
| 4526 | IV | 827 | 0 | 0 | 283 | 0 | 0 | Mono | Mono | | 21463326 | Z1936 | N1a1a1a1a2 | | 4526 |
| 4189 | IV | 827 | 0 | 0 | 283 | 0 | 0 | Mono | Mono | | 19219721 | Z1935 | N1a1a1a1a2a1a | | 4189 |
| 1966 | IV | 827 | 0 | 0 | 283 | 0 | 0 | Mono | Mono | | 14042164 | Z1927 | N1a1a1a1a2a1a1a | | 1966 |
| 173 | IV | 827 | 0 | 0 | 283 | 0 | 0 | Mono | Mono | | 2878213 | S27742 | N1a2a1b~ | | 173 |
| 1366 | IV | 827 | 0 | 0 | 283 | 0 | 0 | Mono | Mono | | 8466354 | F1228 | N1a3~ | | 1366 |
| 3677 | IV | 827 | 0 | 0 | 283 | 0 | 0 | Mono | Mono | | 17891210 | M1870 | N1b | | 3677 |
| 3699 | IV | 827 | 0 | 0 | 283 | 0 | 0 | Mono | Mono | | 17926690 | M1873 | N1b | | 3699 |
| 4121 | IV | 827 | 0 | 0 | 283 | 0 | 0 | Mono | Mono | | 19080602 | V3743 | N1b | | 4121 |
| 4148 | IV | 827 | 0 | 0 | 283 | 0 | 0 | Mono | Mono | | 19148481 | V3831 | N1b | | 4148 |
| 4253 | IV | 827 | 0 | 0 | 283 | 0 | 0 | Mono | Mono | | 19357794 | V4033 | N1b | | 4253 |

| | | M9+ | | | M9- | | | | | | Build 37 | | ISOGG | | |
|---|---|---|---|---|---|---|---|---|---|---|---|---|---|---|---|
| List | Stage | 0 | 1 | . | 0 | 1 | . | M9? | Clade(s) | Prob. | Position | Name | Subgroup Name | Amb. | List |
| 4275 | IV | 827 | 0 | 0 | 283 | 0 | 0 | Mono | Mono | | 19406997 | V4096 | N1b | | 4275 |
| 5558 | IV | 827 | 0 | 0 | 283 | 0 | 0 | Mono | Mono | | 23764923 | M1921 | N1b | | 5558 |
| 3075 | IV | 827 | 0 | 0 | 283 | 0 | 0 | Mono | Mono | | 16492454 | L731 | N1b1a2b1a~ | | 3075 |
| 3801 | IV | 827 | 0 | 0 | 282 | 0 | 1 | Mono | Mono | | 18146985 | L732 | N1b1a2b1a~ | A | 3801 |
| 535 | IV | 827 | 0 | 0 | 283 | 0 | 0 | Mono | Mono | | 6801568 | M1787 | N1b2 | | 535 |
| 579 | IV | 827 | 0 | 0 | 283 | 0 | 0 | Mono | Mono | | 6866201 | M1789 | N1b2 | | 579 |
| 684 | IV | 827 | 0 | 0 | 283 | 0 | 0 | Mono | Mono | | 6961447 | M1790 | N1b2 | | 684 |
| 741 | IV | 827 | 0 | 0 | 283 | 0 | 0 | Mono | Mono | | 7092819 | M1792 | N1b2 | | 741 |
| 827 | IV | 827 | 0 | 0 | 283 | 0 | 0 | Mono | Mono | | 7309148 | M1794 | N1b2 | | 827 |
| 828 | IV | 827 | 0 | 0 | 283 | 0 | 0 | Mono | Mono | | 7309210 | M1795^ | N1b2 | | 828 |
| 837 | IV | 451 | 0 | 376 | 149 | 0 | 134 | Mono | Mono | | 7330034 | M1796 | N1b2 | | 837 |
| 925 | IV | 827 | 0 | 0 | 283 | 0 | 0 | Mono | Mono | | 7605776 | M1797 | N1b2 | | 925 |
| 1165 | IV | 827 | 0 | 0 | 283 | 0 | 0 | Mono | Mono | | 8098341 | M1803 | N1b2 | | 1165 |
| 1181 | IV | 822 | 0 | 5 | 282 | 0 | 1 | Mono | Mono | | 8133554 | M1804 | N1b2 | | 1181 |
| 1310 | IV | 826 | 0 | 1 | 283 | 0 | 0 | Mono | Mono | | 8398614 | M1807^ | N1b2 | | 1310 |
| 1444 | IV | 827 | 0 | 0 | 283 | 0 | 0 | Mono | Mono | | 8550825 | M1809^ | N1b2 | A | 1444 |
| 1466 | IV | 821 | 0 | 6 | 280 | 0 | 3 | Mono | Mono | | 8579009 | M1810 | N1b2 | | 1466 |
| 1653 | IV | 827 | 0 | 0 | 283 | 0 | 0 | Mono | Mono | | 9108931 | M1814 | N1b2 | | 1653 |
| 2247 | IV | 827 | 0 | 0 | 283 | 0 | 0 | Mono | Mono | | 14770599 | M1832^ | N1b2 | | 2247 |
| 2268 | IV | 827 | 0 | 0 | 283 | 0 | 0 | Mono | Mono | | 14804668 | M1835 | N1b2 | | 2268 |
| 2427 | IV | 827 | 0 | 0 | 283 | 0 | 0 | Mono | Mono | | 14991805 | M1838 | N1b2 | | 2427 |
| 2603 | IV | 827 | 0 | 0 | 282 | 0 | 1 | Mono | Mono | | 15374340 | M1842 | N1b2 | | 2603 |
| 3021 | IV | 826 | 0 | 1 | 283 | 0 | 0 | Mono | Mono | | 16375764 | M1851 | N1b2 | | 3021 |
| 3067 | IV | 827 | 0 | 0 | 283 | 0 | 0 | Mono | Mono | | 16483478 | M1853 | N1b2 | | 3067 |
| 3220 | IV | 827 | 0 | 0 | 283 | 0 | 0 | Mono | Mono | | 16844417 | M1859 | N1b2 | | 3220 |
| 3357 | IV | 827 | 0 | 0 | 283 | 0 | 0 | Mono | Mono | | 17169008 | M1860^ | N1b2 | | 3357 |
| 3394 | IV | 827 | 0 | 0 | 282 | 0 | 1 | Mono | Mono | | 17271875 | M1862^ | N1b2 | | 3394 |
| 3592 | IV | 824 | 0 | 3 | 281 | 0 | 2 | Mono | Mono | | 17739376 | M1868 | N1b2 | | 3592 |
| 4119 | IV | 827 | 0 | 0 | 283 | 0 | 0 | Mono | Mono | | 19077866 | M1880 | N1b2 | | 4119 |
| 4265 | IV | 827 | 0 | 0 | 283 | 0 | 0 | Mono | Mono | | 19380397 | M1889^ | N1b2 | | 4265 |
| 4488 | IV | 827 | 0 | 0 | 283 | 0 | 0 | Mono | Mono | | 21369624 | M1896 | N1b2 | | 4488 |
| 4624 | IV | 827 | 0 | 0 | 283 | 0 | 0 | Mono | Mono | | 21663692 | M1898 | N1b2 | | 4624 |
| 5518 | IV | 827 | 0 | 0 | 283 | 0 | 0 | Mono | Mono | | 23566654 | M1919 | N1b2 | | 5518 |
| 5679 | IV | 823 | 0 | 4 | 283 | 0 | 0 | Mono | Mono | | 24456796 | M1923 | N1b2 | | 5679 |
| 1602 | IV | 827 | 0 | 0 | 283 | 0 | 0 | Mono | Mono | | 8861569 | M1812 | N1b2a | | 1602 |
| 5097 | IV | 826 | 0 | 1 | 283 | 0 | 0 | Mono | Mono | | 22669525 | PK3 | N1b2a | A | 5097 |
| 1363 | IV | 827 | 0 | 0 | 283 | 0 | 0 | Mono | Mono | | 8462601 | M1808 | N1b2a1b~ | | 1363 |
| 2044 | IV | 827 | 0 | 0 | 283 | 0 | 0 | Mono | Mono | | 14197977 | P189.2 | N2a~ | A | 2044 |
| 975 | II | 825 | 2 | 0 | 283 | 0 | 0 | M9+ | 1 – 2 | | 7690182 | Z4842 | NO [K2a] | | 975 |
| 1528 | II | 825 | 2 | 0 | 283 | 0 | 0 | M9+ | 1 – 2 | | 8674808 | Z4858 | NO [K2a] | | 1528 |
| 4313 | II | 825 | 2 | 0 | 283 | 0 | 0 | M9+ | 1 – 2 | | 19513070 | V4208 | NO [K2a] | | 4313 |
| 4743 | II | 825 | 2 | 0 | 283 | 0 | 0 | M9+ | 1 – 2 | | 21797754 | Z4952 | NO [K2a] | | 4743 |
| 5531 | II | 825 | 2 | 0 | 283 | 0 | 0 | M9+ | 1 – 2 | | 23617006 | M2346 | NO [K2a] | | 5531 |
| 686 | II | 825 | 2 | 0 | 283 | 0 | 0 | M9+ | 1 – 2 | | 6964531 | S9681 | NO1 [K1a1] | | 686 |
| 1013 | III | 825 | 2 | 0 | 282 | 0 | 1 | M9+ | 1 – 2 | | 7751812 | Z4843 | NO1 [K1a1] | | 1013 |
| 1133 | II | 825 | 2 | 0 | 283 | 0 | 0 | M9+ | 1 – 2 | | 8021340 | V1570 | NO1 [K1a1] | | 1133 |
| 2665 | III | 823 | 2 | 2 | 283 | 0 | 0 | M9+ | 1 – 2 | | 15471925 | Page39 | NO1 [K1a1] | | 2665 |
| 2882 | II | 825 | 2 | 0 | 283 | 0 | 0 | M9+ | 1 – 2 | | 15913078 | V3232 | NO1 [K1a1] | | 2882 |
| 3051 | II | 825 | 2 | 0 | 283 | 0 | 0 | M9+ | 1 – 2 | | 16450159 | M2324 | NO1 [K1a1] | | 3051 |
| 3081 | II | 825 | 2 | 0 | 283 | 0 | 0 | M9+ | 1 – 2 | | 16511867 | M2325 | NO1 [K1a1] | | 3081 |
| 3510 | II | 825 | 2 | 0 | 283 | 0 | 0 | M9+ | 1 – 2 | | 17551116 | S19886 | NO1 [K1a1] | | 3510 |
| 3688 | II | 825 | 2 | 0 | 283 | 0 | 0 | M9+ | 1 – 2 | | 17909881 | M2331 | NO1 [K1a1] | | 3688 |
| 3788 | II | 825 | 2 | 0 | 283 | 0 | 0 | M9+ | 1 – 2 | | 18106997 | S20648 | NO1 [K1a1] | | 3788 |
| 4224 | II | 825 | 2 | 0 | 283 | 0 | 0 | M9+ | 1 – 2 | | 19279765 | P192 | NO1 [K1a1] | | 4224 |
| 5003 | II | 825 | 2 | 0 | 283 | 0 | 0 | M9+ | 1 – 2 | | 22214221 | P193 | NO1 [K1a1] | | 5003 |
| 5094 | II | 825 | 2 | 0 | 283 | 0 | 0 | M9+ | 1 – 2 | | 22665262 | P195.2 | NO1 [K1a1] | A | 5094 |
| 5559 | II | 825 | 2 | 0 | 283 | 0 | 0 | M9+ | 1 – 2 | | 23765556 | S25824 | NO1 [K1a1] | | 5559 |
| 570 | III | 819 | 2 | 6 | 280 | 0 | 3 | M9+ | 1 – 2 | | 6858319 | M1739 | O | | 570 |
| 811 | II | 825 | 2 | 0 | 283 | 0 | 0 | M9+ | 1 – 2 | | 7257494 | M1740 | O | | 811 |
| 888 | II | 824 | 3 | 0 | 283 | 0 | 0 | M9+ | | PI | 7556207 | M1741 | O | | 888 |
| 894 | II | 825 | 2 | 0 | 283 | 0 | 0 | M9+ | 1 – 2 | | 7568568 | P186 | O | | 894 |
| 1350 | III | 820 | 2 | 5 | 282 | 0 | 1 | M9+ | 1 – 2 | | 8448515 | M1743 | O | | 1350 |
| 1586 | III | 823 | 2 | 2 | 283 | 0 | 0 | M9+ | 1 – 2 | | 8809390 | M1573 | O | | 1586 |
| 1624 | II | 825 | 2 | 0 | 283 | 0 | 0 | M9+ | 1 – 2 | | 8909837 | F175 | O | | 1624 |
| 1660 | III | 824 | 2 | 1 | 283 | 0 | 0 | M9+ | 1 – 2 | | 9128713 | M1744 | O | | 1660 |
| 1733 | III | 823 | 2 | 2 | 283 | 0 | 0 | M9+ | 1 – 2 | | 9818338 | M1745 | O | | 1733 |
| 1918 | III | 820 | 2 | 5 | 283 | 0 | 0 | M9+ | 1 – 2 | | 13840063 | FGC12434 | O | | 1918 |
| 2004 | II | 825 | 2 | 0 | 283 | 0 | 0 | M9+ | 1 – 2 | | 14096806 | M1747 | O | | 2004 |
| 2081 | II | 825 | 2 | 0 | 283 | 0 | 0 | M9+ | 1 – 2 | | 14266799 | CTS2340 | O | | 2081 |
| 2266 | III | 824 | 2 | 1 | 283 | 0 | 0 | M9+ | 1 – 2 | | 14796358 | M1749 | O | | 2266 |
| 2528 | II | 825 | 2 | 0 | 283 | 0 | 0 | M9+ | 1 – 2 | | 15122573 | M1359 | O | | 2528 |
| 2539 | II | 825 | 2 | 0 | 283 | 0 | 0 | M9+ | 1 – 2 | | 15167624 | M1750 | O | | 2539 |
| 2626 | II | 825 | 2 | 0 | 283 | 0 | 0 | M9+ | 1 – 2 | | 15415115 | P191 | O | | 2626 |
| 2822 | II | 825 | 2 | 0 | 283 | 0 | 0 | M9+ | 1 – 2 | | 15754313 | P196 | O | | 2822 |
| 2893 | III | 824 | 2 | 1 | 283 | 0 | 0 | M9+ | 1 – 2 | | 15948265 | M1751 | O | | 2893 |
| 3125 | III | 821 | 2 | 4 | 282 | 0 | 1 | M9+ | 1 – 2 | | 16628931 | M1754 | O | | 3125 |
| 3238 | II | 825 | 2 | 0 | 283 | 0 | 0 | M9+ | 1 – 2 | | 16870987 | M1756 | O | | 3238 |
| 3292 | II | 825 | 2 | 0 | 283 | 0 | 0 | M9+ | 1 – 2 | | 16954110 | S27659 | O | | 3292 |
| 3504 | II | 825 | 2 | 0 | 283 | 0 | 0 | M9+ | 1 – 2 | | 17535032 | M1759 | O | | 3504 |
| 3719 | III | 813 | 2 | 12 | 282 | 0 | 1 | M9+ | 1 – 2 | | 17968786 | M1762 | O | | 3719 |
| 3813 | III | 824 | 2 | 1 | 283 | 0 | 0 | M9+ | 1 – 2 | | 18175815 | M1763 | O | A | 3813 |
| 3831 | II | 825 | 2 | 0 | 283 | 0 | 0 | M9+ | 1 – 2 | | 18233782 | M1764 | O | | 3831 |
| 3841 | II | 825 | 2 | 0 | 283 | 0 | 0 | M9+ | 1 – 2 | | 18382410 | FGC12436 | O | | 3841 |
| 3910 | II | 825 | 2 | 0 | 283 | 0 | 0 | M9+ | 1 – 2 | | 18621300 | M1765 | O | | 3910 |
| 3912 | III | 824 | 2 | 1 | 283 | 0 | 0 | M9+ | 1 – 2 | | 18630185 | M1766 | O | | 3912 |
| 4243 | III | 820 | 2 | 5 | 282 | 0 | 1 | M9+ | 1 – 2 | | 19320942 | M1769 | O | | 4243 |
| 4269 | II | 825 | 2 | 0 | 283 | 0 | 0 | M9+ | 1 – 2 | | 19384549 | F540 | O | | 4269 |
| 4296 | II | 825 | 2 | 0 | 283 | 0 | 0 | M9+ | 1 – 2 | | 19480880 | M1770 | O | | 4296 |
| 4491 | II | 825 | 2 | 0 | 283 | 0 | 0 | M9+ | 1 – 2 | | 21385679 | M1771 | O | | 4491 |
| 4577 | II | 825 | 2 | 0 | 283 | 0 | 0 | M9+ | 1 – 2 | | 21604901 | M1773 | O | | 4577 |
| 4753 | II | 825 | 2 | 0 | 283 | 0 | 0 | M9+ | 1 – 2 | | 21815957 | M1775 | O | | 4753 |
| 4964 | II | 825 | 2 | 0 | 283 | 0 | 0 | M9+ | 1 – 2 | | 22133425 | M1776 | O | | 4964 |
| 4981 | III | 825 | 2 | 0 | 281 | 0 | 2 | M9+ | 1 – 2 | | 22166455 | M1777 | O | | 4981 |
| 5052 | III | 823 | 2 | 2 | 283 | 0 | 0 | M9+ | 1 – 2 | | 22559001 | M1778 | O | | 5052 |
| 5220 | II | 825 | 2 | 0 | 283 | 0 | 0 | M9+ | 1 – 2 | | 22880173 | M1780 | O | | 5220 |
| 5440 | II | 825 | 2 | 0 | 283 | 0 | 0 | M9+ | 1 – 2 | | 23382337 | M1781 | O | | 5440 |

| | | | M9+ | | | | M9- | | | | | Build 37 | | ISOGG | | |
|---|---|---|---|---|---|---|---|---|---|---|---|---|---|---|---|---|
| List | Stage | 0 | 1 | . | | 0 | 1 | . | M9? | Clade(s) | Prob. | Position | Name | Subgroup Name | Amb. | List |
| 5539 | II | 825 | 2 | 0 | | 283 | 0 | 0 | M9+ | 1 – 2 | | 23634362 | P188 | O | | 5539 |
| 5624 | III | 820 | 2 | 5 | | 282 | 0 | 1 | M9+ | 1 – 2 | | 23975319 | M1782 | O | | 5624 |
| 2746 | IV | 827 | 0 | 0 | | 283 | 0 | 0 | Mono | Mono | | 15591492 | N9 | O(Notes) | | 2746 |
| 4847 | IV | 827 | 0 | 0 | | 283 | 0 | 0 | Mono | Mono | | 21898279 | N11 | O(Notes) | | 4847 |
| 4850 | IV | 827 | 0 | 0 | | 283 | 0 | 0 | Mono | Mono | | 21900347 | N10 | O(Notes) | | 4850 |
| 4899 | IV | 827 | 0 | 0 | | 283 | 0 | 0 | Mono | Mono | | 21938532 | N4 | O(Notes) | | 4899 |
| 5156 | IV | 827 | 0 | 0 | | 283 | 0 | 0 | Mono | Mono | | 22746689 | M297 | O(Notes) | | 5156 |
| 855 | II | 825 | 2 | 0 | | 283 | 0 | 0 | M9+ | 1 – 2 | | 7410009 | M1297 | O1 | | 855 |
| 2231 | III | 821 | 2 | 4 | | 282 | 0 | 1 | M9+ | 1 – 2 | | 14727776 | M1354 | O1 | | 2231 |
| 3595 | III | 822 | 2 | 3 | | 283 | 0 | 0 | M9+ | 1 – 2 | | 17742266 | M1415 | O1 | | 3595 |
| 3769 | II | 825 | 2 | 0 | | 283 | 0 | 0 | M9+ | 1 – 2 | | 18071170 | M1422 | O1 | | 3769 |
| 4596 | III | 825 | 1 | 1 | | 283 | 0 | 0 | M9+ | 1 | | 21622006 | Page20 | O1a | | 4596 |
| 4629 | II | 826 | 1 | 0 | | 283 | 0 | 0 | M9+ | 1 | | 21674068 | L466 | O1a | | 4629 |
| 4706 | II | 826 | 1 | 0 | | 283 | 0 | 0 | M9+ | 1 | | 21762685 | M119 | O1a | | 4706 |
| 480 | III | 825 | 1 | 1 | | 283 | 0 | 0 | M9+ | 1 | | 6678315 | F31 | O1a1a | | 480 |
| 690 | II | 826 | 1 | 0 | | 283 | 0 | 0 | M9+ | 1 | | 6969303 | F54 | O1a1a | | 690 |
| 929 | II | 826 | 1 | 0 | | 283 | 0 | 0 | M9+ | 1 | | 7616941 | F89 | O1a1a | | 929 |
| 2357 | IV | 791 | 0 | 36 | | 274 | 0 | 9 | Mono | Mono | MD | 14911080 | CTS3422 | O1a1a | | 2357 |
| 2761 | II | 826 | 1 | 0 | | 283 | 0 | 0 | M9+ | 1 | | 15603353 | CTS4351 | O1a1a | | 2761 |
| 2906 | III | 825 | 1 | 1 | | 281 | 0 | 2 | M9+ | 1 | | 15994985 | CTS5059 | O1a1a | | 2906 |
| 3033 | II | 826 | 1 | 0 | | 283 | 0 | 0 | M9+ | 1 | | 16404175 | F333 | O1a1a | | 3033 |
| 3356 | II | 826 | 1 | 0 | | 283 | 0 | 0 | M9+ | 1 | | 17156422 | CTS6864 | O1a1a | | 3356 |
| 3383 | III | 825 | 1 | 1 | | 283 | 0 | 0 | M9+ | 1 | | 17237048 | CTS7015 | O1a1a | | 3383 |
| 3874 | II | 826 | 1 | 0 | | 283 | 0 | 0 | M9+ | 1 | | 18561615 | CTS8934 | O1a1a | | 3874 |
| 3997 | III | 818 | 2 | 7 | | 281 | 0 | 2 | M9+ | | PI | 18822568 | CTS9321 | O1a1a | W | 3997 |
| 5031 | III | 825 | 1 | 1 | | 283 | 0 | 0 | M9+ | 1 | | 22513920 | L83 | O1a1a | | 5031 |
| 2809 | III | 824 | 1 | 2 | | 279 | 0 | 4 | M9+ | 1 | | 15723358 | CTS4588 | O1a1a1 | | 2809 |
| 3685 | II | 826 | 1 | 0 | | 283 | 0 | 0 | M9+ | 1 | | 17909248 | F446 | O1a1a1 | | 3685 |
| 4355 | II | 826 | 1 | 0 | | 283 | 0 | 0 | M9+ | 1 | | 21052297 | F560 | O1a1a1 | | 4355 |
| 1501 | IV | 827 | 0 | 0 | | 283 | 0 | 0 | Mono | Mono | | 8621202 | F157 | O1a1a1a | | 1501 |
| 2256 | IV | 823 | 0 | 4 | | 283 | 0 | 0 | Mono | Mono | | 14780370 | CTS3265 | O1a1a1a | | 2256 |
| 4080 | IV | 827 | 0 | 0 | | 283 | 0 | 0 | Mono | Mono | | 18999749 | F518 | O1a1a1a | | 4080 |
| 4438 | IV | 827 | 0 | 0 | | 283 | 0 | 0 | Mono | Mono | | 21260029 | F571 | O1a1a1a | | 4438 |
| 863 | II | 826 | 1 | 0 | | 283 | 0 | 0 | M9+ | NU | | 7517784 | F78 | O1a1a1a1 | W | 863 |
| 1579 | IV | 798 | 0 | 29 | | 275 | 0 | 8 | Mono | Mono | | 8797778 | F168 | O1a1a1a1 | | 1579 |
| 3982 | IV | 797 | 0 | 30 | | 271 | 0 | 12 | Mono | Mono | | 18797957 | CTS9282 | O1a1a1a1 | | 3982 |
| 881 | IV | 827 | 0 | 0 | | 283 | 0 | 0 | Mono | Mono | | 7545605 | F81 | O1a1a1a1a | | 881 |
| 3050 | IV | 823 | 0 | 4 | | 283 | 0 | 0 | Mono | Mono | | 16445951 | CTS5709 | O1a1a1a1a | | 3050 |
| 2097 | IV | 827 | 0 | 0 | | 283 | 0 | 0 | Mono | Mono | | 14313964 | CTS2458 | O1a1a1a1a1 | | 2097 |
| 4214 | IV | 692 | 0 | 135 | | 231 | 0 | 52 | Mono | Mono | | 19261970 | F533 | O1a1a1a1a1a | | 4214 |
| 5093 | IV | 827 | 0 | 0 | | 283 | 0 | 0 | Mono | Mono | | 22657721 | F619 | O1a1a1a1a1a1 | | 5093 |
| 5405 | IV | 827 | 0 | 0 | | 283 | 0 | 0 | Mono | Mono | | 23269595 | CTS11784 | O1a1a1a1a1a1 | | 5405 |
| 5429 | IV | 824 | 0 | 3 | | 283 | 0 | 0 | Mono | Mono | | 23348259 | Z23478 | O1a1a1a1a1a1 | | 5429 |
| 2542 | IV | 827 | 0 | 0 | | 283 | 0 | 0 | Mono | Mono | | 15192338 | F285 | O1a1a1a1a1a1a3 | | 2542 |
| 3789 | IV | 827 | 0 | 0 | | 283 | 0 | 0 | Mono | Mono | | 18107379 | F469 | O1a1a1a1a1a1a3 | | 3789 |
| 5079 | IV | 827 | 0 | 0 | | 283 | 0 | 0 | Mono | Mono | | 22603043 | Y137197 | O1a1a1a1a1a1a3 | | 5079 |
| 4869 | IV | 825 | 0 | 2 | | 282 | 0 | 1 | Mono | Mono | | 21907648 | N8.2 | O1a1a1a3 | A | 4869 |
| 4649 | IV | 827 | 0 | 0 | | 283 | 0 | 0 | Mono | Mono | | 21717472 | M101 | O1a1a1b1a | | 4649 |
| 1751 | IV | 795 | 0 | 32 | | 271 | 0 | 12 | Mono | Mono | | 9845876 | F1620.2 | O1a1b | A | 1751 |
| 2964 | IV | 827 | 0 | 0 | | 283 | 0 | 0 | Mono | Mono | | 16247376 | F2104 | O1a1b | | 2964 |
| 3989 | IV | 827 | 0 | 0 | | 283 | 0 | 0 | Mono | Mono | | 18809368 | F2852^ | O1a1b | | 3989 |
| 4078 | IV | 824 | 0 | 3 | | 282 | 0 | 1 | Mono | Mono | | 18992733 | F2911 | O1a1b | | 4078 |
| 4291 | IV | 827 | 0 | 0 | | 283 | 0 | 0 | Mono | Mono | | 19474424 | F3063 | O1a1b | | 4291 |
| 5534 | IV | 827 | 0 | 0 | | 283 | 0 | 0 | Mono | Mono | | 23620488 | F3495 | O1a1b | | 5534 |
| 5677 | IV | 825 | 0 | 2 | | 283 | 0 | 0 | Mono | Mono | | 24448831 | F3640 | O1a1b | | 5677 |
| 1647 | IV | 827 | 0 | 0 | | 283 | 0 | 0 | Mono | Mono | | 9093029 | F1476 | O1a1b1a | | 1647 |
| 667 | IV | 827 | 0 | 0 | | 283 | 0 | 0 | Mono | Mono | | 6946901 | F881 | O1a2 | | 667 |
| 728 | IV | 827 | 0 | 0 | | 283 | 0 | 0 | Mono | Mono | | 7074018 | F908 | O1a2 | | 728 |
| 2416 | IV | 827 | 0 | 0 | | 282 | 0 | 1 | Mono | Mono | | 14979025 | F1880 | O1a2 | | 2416 |
| 3293 | IV | 827 | 0 | 0 | | 283 | 0 | 0 | Mono | Mono | | 16954537 | F2340 | O1a2 | | 3293 |
| 3337 | IV | 827 | 0 | 0 | | 282 | 0 | 1 | Mono | Mono | | 17110058 | F2380 | O1a2 | | 3337 |
| 3694 | IV | 827 | 0 | 0 | | 283 | 0 | 0 | Mono | Mono | | 17919250 | F2635 | O1a2 | | 3694 |
| 3731 | IV | 827 | 0 | 0 | | 282 | 0 | 1 | Mono | Mono | | 18018434 | F2671 | O1a2 | | 3731 |
| 4392 | IV | 827 | 0 | 0 | | 283 | 0 | 0 | Mono | Mono | | 21143756 | F3134 | O1a2 | | 4392 |
| 4414 | IV | 808 | 0 | 19 | | 280 | 0 | 3 | Mono | Mono | | 21216583 | F3152 | O1a2 | | 4414 |
| 4417 | IV | 826 | 0 | 1 | | 283 | 0 | 0 | Mono | Mono | | 21224636 | F3157 | O1a2 | | 4417 |
| 4786 | IV | 827 | 0 | 0 | | 283 | 0 | 0 | Mono | Mono | | 21868672 | M50 | O1a2 | | 4786 |
| 4867 | IV | 827 | 0 | 0 | | 283 | 0 | 0 | Mono | Mono | | 21907097 | M110 | O1a2 | | 4867 |
| 4895 | IV | 827 | 0 | 0 | | 283 | 0 | 0 | Mono | Mono | | 21936138 | M103 | O1a2 | | 4895 |
| 1720 | IV | 827 | 0 | 0 | | 283 | 0 | 0 | Mono | Mono | | 9769559 | F1600 | O1a2a2 | | 1720 |
| 4153 | IV | 827 | 0 | 0 | | 283 | 0 | 0 | Mono | Mono | | 19151833 | F2954 | O1a2a2 | | 4153 |
| 1353 | II | 826 | 1 | 0 | | 283 | 0 | 0 | M9+ | 2 | | 8452166 | F138 | O1b | | 1353 |
| 1578 | II | 826 | 1 | 0 | | 283 | 0 | 0 | M9+ | 2 | | 8796086 | L690 | O1b | | 1578 |
| 2105 | II | 826 | 1 | 0 | | 283 | 0 | 0 | M9+ | 2 | | 14332189 | M1341 | O1b | | 2105 |
| 2166 | II | 826 | 1 | 0 | | 283 | 0 | 0 | M9+ | 2 | | 14495243 | P31 | O1b | | 2166 |
| 2646 | III | 816 | 1 | 10 | | 278 | 0 | 5 | M9+ | 2 | | 15440730 | M1363 | O1b | | 2646 |
| 2983 | II | 826 | 1 | 0 | | 283 | 0 | 0 | M9+ | 2 | | 16301305 | L463 | O1b | | 2983 |
| 3634 | II | 826 | 1 | 0 | | 283 | 0 | 0 | M9+ | 2 | | 17809157 | M1417 | O1b | | 3634 |
| 4055 | II | 826 | 1 | 0 | | 283 | 0 | 0 | M9+ | 2 | | 18941356 | F516 | O1b | | 4055 |
| 5136 | II | 826 | 1 | 0 | | 283 | 0 | 0 | M9+ | 2 | | 22739301 | M268 | O1b | | 5136 |
| 3261 | III | 825 | 1 | 1 | | 283 | 0 | 0 | M9+ | 2 | | 16903082 | F2320 | O1b1 | | 3261 |
| 4233 | II | 826 | 1 | 0 | | 283 | 0 | 0 | M9+ | 2 | | 19292612 | M1441 | O1b1 | | 4233 |
| 4966 | II | 826 | 1 | 0 | | 283 | 0 | 0 | M9+ | 2 | | 22140302 | M1465 | O1b1 | | 4966 |
| 5065 | III | 825 | 1 | 1 | | 282 | 0 | 1 | M9+ | 2 | | 22582305 | M1470 | O1b1a | | 5065 |
| 79 | III | 826 | 1 | 0 | | 282 | 0 | 1 | M9+ | 2 | | 2737043 | M1279 | O1b1a1 | | 79 |
| 796 | III | 824 | 1 | 2 | | 283 | 0 | 0 | M9+ | 2 | | 7213725 | M1290 | O1b1a1 | | 796 |
| 1141 | III | 825 | 1 | 1 | | 283 | 0 | 0 | M9+ | 2 | | 8039638 | M1305 | O1b1a1 | | 1141 |
| 1226 | II | 826 | 1 | 0 | | 283 | 0 | 0 | M9+ | 2 | | 8239331 | M1306 | O1b1a1 | | 1226 |
| 1416 | II | 826 | 1 | 0 | | 283 | 0 | 0 | M9+ | 2 | | 8505317 | M1313.2 | O1b1a1 | A | 1416 |
| 1489 | II | 826 | 1 | 0 | | 283 | 0 | 0 | M9+ | 2 | | 8605895 | M1316 | O1b1a1 | | 1489 |
| 1498 | II | 826 | 1 | 0 | | 283 | 0 | 0 | M9+ | 2 | | 8619144 | M1317 | O1b1a1 | | 1498 |
| 2203 | III | 760 | 1 | 66 | | 262 | 0 | 21 | M9+ | 2 | | 14627760 | M1352 | O1b1a1 | | 2203 |
| 3254 | II | 826 | 1 | 0 | | 283 | 0 | 0 | M9+ | 2 | | 16898155 | F2317 | O1b1a1 | | 3254 |
| 3451 | II | 826 | 1 | 0 | | 283 | 0 | 0 | M9+ | 2 | | 17418386 | M1405 | O1b1a1 | | 3451 |
| 3944 | III | 826 | 1 | 0 | | 282 | 0 | 1 | M9+ | 2 | | 18717224 | CTS9150 | O1b1a1 | | 3944 |
| 4447 | II | 826 | 1 | 0 | | 283 | 0 | 0 | M9+ | 2 | | 21285962 | PK4 | O1b1a1 | | 4447 |

| | | M9+ | | | M9- | | | | | | Build 37 | | ISOGG | | |
|---|---|---|---|---|---|---|---|---|---|---|---|---|---|---|---|
| List | Stage | 0 | 1 | . | 0 | 1 | . | M9? | Clade(s) | Prob. | Position | Name | Subgroup Name | Amb. | List |
| 5403 | III | 826 | 1 | 0 | 282 | 0 | 1 | M9+ | 2 | | 23268915 | M1489 | O1b1a1 | | 5403 |
| 5495 | III | 825 | 1 | 1 | 283 | 0 | 0 | M9+ | 2 | | 23497646 | M1494 | O1b1a1 | | 5495 |
| 25 | III | 821 | 1 | 5 | 282 | 0 | 1 | M9+ | 2 | | 2667652 | F709 | O1b1a1a | | 25 |
| 880 | III | 822 | 1 | 4 | 281 | 0 | 2 | M9+ | 2 | 1H | 7543143 | M1300 | O1b1a1a | | 880 |
| 3065 | III | 825 | 1 | 1 | 283 | 0 | 0 | M9+ | 2 | | 16477410 | M1376 | O1b1a1a | | 3065 |
| 3169 | III | 821 | 1 | 5 | 281 | 0 | 2 | M9+ | 2 | | 16760911 | M1383 | O1b1a1a | | 3169 |
| 3446 | II | 826 | 1 | 0 | 283 | 0 | 0 | M9+ | 2 | | 17406441 | M1404 | O1b1a1a | | 3446 |
| 4898 | II | 826 | 1 | 0 | 283 | 0 | 0 | M9+ | 2 | | 21938444 | M95 | O1b1a1a | | 4898 |
| 4002 | IV | 826 | 0 | 1 | 283 | 0 | 0 | Mono | Mono | MD | 18835758 | F2861 | O1b1a1a1a1 | | 4002 |
| 4851 | II | 826 | 1 | 0 | 283 | 0 | 0 | M9+ | 2 | | 21900849 | M88 | O1b1a1a1a1 | | 4851 |
| 2495 | IV | 827 | 0 | 0 | 283 | 0 | 0 | Mono | Mono | | 15043403 | M1357 | O1b1a1a1b | | 2495 |
| 3512 | IV | 827 | 0 | 0 | 283 | 0 | 0 | Mono | Mono | | 17556777 | M1410 | O1b1a1a1b | | 3512 |
| 4342 | IV | 826 | 0 | 1 | 283 | 0 | 0 | Mono | Mono | | 20822799 | F3092 | O1b1a1a1b | | 4342 |
| 5050 | IV | 824 | 0 | 3 | 283 | 0 | 0 | Mono | Mono | | 22558351 | M1469 | O1b1a1a1b | | 5050 |
| 5441 | IV | 827 | 0 | 0 | 283 | 0 | 0 | Mono | Mono | | 23383729 | Z2953.2 | O1b1a1a2 | A | 5441 |
| 5154 | IV | 827 | 0 | 0 | 283 | 0 | 0 | Mono | Mono | | 22746267 | Page103 | O1b1a1a3~ | | 5154 |
| 4794 | IV | 824 | 0 | 3 | 281 | 0 | 2 | Mono | Mono | | 21869923 | Page59 | O1b1a2 | | 4794 |
| 3516 | IV | 827 | 0 | 0 | 283 | 0 | 0 | Mono | Mono | | 17562717 | M1654 | O1b1a2b | | 3516 |
| 2 | IV | 827 | 0 | 0 | 283 | 0 | 0 | Mono | Mono | | 2655180 | SRY465 | O1b2 | | 2 |
| 2169 | IV | 827 | 0 | 0 | 283 | 0 | 0 | Mono | Mono | | 14496407 | P49 | O1b2 | | 2169 |
| 2622 | IV | 827 | 0 | 0 | 283 | 0 | 0 | Mono | Mono | | 15409812 | Page92 | O1b2a | | 2622 |
| 5692 | IV | 825 | 0 | 2 | 282 | 0 | 1 | Mono | Mono | | 24510581 | IMS-JST022454 | O1b2a | | 5692 |
| 492 | IV | 827 | 0 | 0 | 283 | 0 | 0 | Mono | Mono | | 6702174 | CTS348 | O1b2a1 | | 492 |
| 1105 | IV | 827 | 0 | 0 | 283 | 0 | 0 | Mono | Mono | | 7940697 | Z24512 | O1b2a1 | | 1105 |
| 3975 | IV | 827 | 0 | 0 | 283 | 0 | 0 | Mono | Mono | | 18782782 | CTS9259 | O1b2a1 | | 3975 |
| 4106 | IV | 827 | 0 | 0 | 283 | 0 | 0 | Mono | Mono | | 19057095 | CTS9752 | O1b2a1 | | 4106 |
| 1424 | IV | 827 | 0 | 0 | 283 | 0 | 0 | Mono | Mono | | 8519034 | K10 | O1b2a1a | | 1424 |
| 1434 | IV | 827 | 0 | 0 | 283 | 0 | 0 | Mono | Mono | | 8538165 | F1290 | O1b2a1a | | 1434 |
| 1681 | IV | 826 | 0 | 1 | 283 | 0 | 0 | Mono | Mono | | 9387005 | F1530 | O1b2a1a | | 1681 |
| 2197 | IV | 827 | 0 | 0 | 283 | 0 | 0 | Mono | Mono | | 14604396 | F1805 | O1b2a1a | | 2197 |
| 3693 | IV | 825 | 0 | 2 | 283 | 0 | 0 | Mono | Mono | | 17917389 | F2634 | O1b2a1a | | 3693 |
| 4159 | IV | 826 | 0 | 1 | 283 | 0 | 0 | Mono | Mono | | 19162451 | F2960 | O1b2a1a | | 4159 |
| 4413 | IV | 826 | 0 | 1 | 282 | 0 | 1 | Mono | Mono | | 21205260 | F3147 | O1b2a1a | | 4413 |
| 5084 | IV | 812 | 0 | 15 | 276 | 0 | 7 | Mono | Mono | | 22619159 | F3327 | O1b2a1a | | 5084 |
| 5169 | IV | 826 | 0 | 1 | 283 | 0 | 0 | Mono | Mono | | 22752710 | F3356 | O1b2a1a | | 5169 |
| 4006 | IV | 827 | 0 | 0 | 283 | 0 | 0 | Mono | Mono | | 18842779 | F2868 | O1b2a1a2 | | 4006 |
| 4360 | IV | 826 | 0 | 1 | 283 | 0 | 0 | Mono | Mono | | 21066749 | F3110 | O1b2a1a2 | | 4360 |
| 4886 | IV | 827 | 0 | 0 | 283 | 0 | 0 | Mono | Mono | | 21930113 | L682 | O1b2a1a2a | | 4886 |
| 2406 | IV | 827 | 0 | 0 | 283 | 0 | 0 | Mono | Mono | | 14968449 | Page90 | O1b2a2~ | | 2406 |
| 3326 | IV | 827 | 0 | 0 | 283 | 0 | 0 | Mono | Mono | | 17053771 | P198 | O2 | | 3326 |
| 3616 | IV | 827 | 0 | 0 | 283 | 0 | 0 | Mono | Mono | | 17772380 | F433 | O2 | | 3616 |
| 3803 | IV | 827 | 0 | 0 | 283 | 0 | 0 | Mono | Mono | | 18148984 | F471 | O2 | | 3803 |
| 4717 | IV | 821 | 0 | 6 | 281 | 0 | 2 | Mono | Mono | | 21764674 | M122 | O2 | | 4717 |
| 5132 | IV | 827 | 0 | 0 | 283 | 0 | 0 | Mono | Mono | | 22735710 | CTS10736 | O2 | | 5132 |
| 5176 | IV | 827 | 0 | 0 | 283 | 0 | 0 | Mono | Mono | | 22758116 | CTS10753 | O2 | | 5176 |
| 5250 | IV | 827 | 0 | 0 | 283 | 0 | 0 | Mono | Mono | | 22931057 | F633 | O2 | | 5250 |
| 128 | IV | 827 | 0 | 0 | 283 | 0 | 0 | Mono | Mono | | 2821786 | M324 | O2a | | 128 |
| 443 | IV | 827 | 0 | 0 | 282 | 0 | 1 | Mono | Mono | | 6383533 | F27 | O2a | | 443 |
| 819 | IV | 821 | 0 | 6 | 281 | 0 | 2 | Mono | Mono | | 7280947 | F71 | O2a | | 819 |
| 1298 | IV | 827 | 0 | 0 | 282 | 0 | 1 | Mono | Mono | | 8381238 | F129 | O2a | | 1298 |
| 3060 | IV | 827 | 0 | 0 | 283 | 0 | 0 | Mono | Mono | | 16461916 | F341 | O2a | | 3060 |
| 3658 | IV | 827 | 0 | 0 | 283 | 0 | 0 | Mono | Mono | | 17854151 | CTS8153 | O2a | | 3658 |
| 3724 | IV | 827 | 0 | 0 | 283 | 0 | 0 | Mono | Mono | | 17977423 | CTS8399 | O2a | | 3724 |
| 3872 | IV | 827 | 0 | 0 | 283 | 0 | 0 | Mono | Mono | | 18560005 | P200 | O2a | | 3872 |
| 3921 | IV | 827 | 0 | 0 | 283 | 0 | 0 | Mono | Mono | | 18647042 | P199 | O2a | | 3921 |
| 4168 | IV | 827 | 0 | 0 | 283 | 0 | 0 | Mono | Mono | | 19179463 | P197 | O2a | | 4168 |
| 4439 | IV | 827 | 0 | 0 | 283 | 0 | 0 | Mono | Mono | | 21261489 | F572 | O2a | | 4439 |
| 2091 | IV | 827 | 0 | 0 | 283 | 0 | 0 | Mono | Mono | | 14288981 | L127.1 | O2a1 | | 2091 |
| 4007 | IV | 827 | 0 | 0 | 283 | 0 | 0 | Mono | Mono | | 18842841 | L465 | O2a1 | | 4007 |
| 5157 | IV | 827 | 0 | 0 | 283 | 0 | 0 | Mono | Mono | | 22746786 | L467 | O2a1 | | 5157 |
| 2408 | IV | 827 | 0 | 0 | 283 | 0 | 0 | Mono | Mono | | 14968529 | Page127 | O2a1a | | 2408 |
| 2373 | IV | 827 | 0 | 0 | 283 | 0 | 0 | Mono | Mono | | 14928001 | Page124 | O2a1a1a | | 2373 |
| 2624 | IV | 827 | 0 | 0 | 283 | 0 | 0 | Mono | Mono | | 15414633 | Page130 | O2a1a1b2a | | 2624 |
| 4978 | IV | 827 | 0 | 0 | 283 | 0 | 0 | Mono | Mono | | 22161564 | F4312 | O2a1a1b1 | | 4978 |
| 884 | IV | 827 | 0 | 0 | 283 | 0 | 0 | Mono | Mono | | 7546726 | IMS-JST002611 | O2a1b | | 884 |
| 123 | IV | 827 | 0 | 0 | 283 | 0 | 0 | Mono | Mono | | 2815303 | F11 | O2a1b1a1a1a | | 123 |
| 4165 | IV | 826 | 0 | 1 | 283 | 0 | 0 | Mono | Mono | | 19171953 | F526 | O2a1b1a1a1a1 | | 4165 |
| 1153 | II | 826 | 1 | 0 | 283 | 0 | 0 | M9+ | NU | | 8068096 | M11115 | O2a1b1a1a1a1a1 | W | 1153 |
| 640 | IV | 826 | 0 | 1 | 283 | 0 | 0 | Mono | Mono | | 6932199 | Page74.2 | O2a1b2 | A | 640 |
| 4699 | IV | 827 | 0 | 0 | 283 | 0 | 0 | Mono | Mono | | 21757306 | M164 | O2a1b2a1a1 | | 4699 |
| 134 | IV | 827 | 0 | 0 | 283 | 0 | 0 | Mono | Mono | | 2828196 | P201 | O2a2 | | 134 |
| 3679 | IV | 827 | 0 | 0 | 283 | 0 | 0 | Mono | Mono | | 17893116 | CTS8236 | O2a2 | | 3679 |
| 4147 | IV | 825 | 0 | 2 | 283 | 0 | 0 | Mono | Mono | | 19148128 | F525 | O2a2 | | 4147 |
| 2370 | IV | 827 | 0 | 0 | 283 | 0 | 0 | Mono | Mono | | 14924869 | M188 | O2a2a | | 2370 |
| 2220 | IV | 827 | 0 | 0 | 283 | 0 | 0 | Mono | Mono | | 14666926 | CTS3106 | O2a2a1 | | 2220 |
| 3607 | IV | 827 | 0 | 0 | 283 | 0 | 0 | Mono | Mono | | 17763772 | F2588 | O2a2a1 | | 3607 |
| 522 | IV | 827 | 0 | 0 | 283 | 0 | 0 | Mono | Mono | | 6756176 | CTS445 | O2a2a1a | | 522 |
| 3127 | IV | 745 | 0 | 82 | 262 | 0 | 21 | Mono | Mono | | 16631874 | F2221 | O2a2a1a | | 3127 |
| 3281 | IV | 826 | 0 | 1 | 279 | 0 | 4 | Mono | Mono | | 16940124 | F2329 | O2a2a1a | | 3281 |
| 3615 | IV | 827 | 0 | 0 | 283 | 0 | 0 | Mono | Mono | | 17771689 | F2592 | O2a2a1a | | 3615 |
| 5377 | IV | 827 | 0 | 0 | 282 | 0 | 1 | Mono | Mono | | 23240530 | CTS11727 | O2a2a1a | | 5377 |
| 5584 | IV | 824 | 0 | 3 | 283 | 0 | 0 | Mono | Mono | | 23828535 | F3559 | O2a2a1a | | 5584 |
| 4692 | IV | 827 | 0 | 0 | 283 | 0 | 0 | Mono | Mono | | 21751440 | Page96 | O2a2a1a1a | | 4692 |
| 272 | IV | 827 | 0 | 0 | 283 | 0 | 0 | Mono | Mono | | 4078217 | M7 | O2a2a1a2 | | 272 |
| 599 | IV | 823 | 0 | 4 | 283 | 0 | 0 | Mono | Mono | | 6892809 | F4066 | O2a2a1a2 | | 599 |
| 3614 | IV | 827 | 0 | 0 | 283 | 0 | 0 | Mono | Mono | | 17770871 | F2591 | O2a2a1a2 | | 3614 |
| 3743 | IV | 827 | 0 | 0 | 283 | 0 | 0 | Mono | Mono | | 18032765 | F2680 | O2a2a1a2 | | 3743 |
| 2728 | IV | 826 | 0 | 1 | 283 | 0 | 0 | Mono | Mono | | 15575790 | M209 | O2a2a1a2a | | 2728 |
| 4475 | IV | 827 | 0 | 0 | 283 | 0 | 0 | Mono | Mono | | 21354797 | F3177 | O2a2a1a2a | | 4475 |
| 5105 | IV | 826 | 0 | 1 | 283 | 0 | 0 | Mono | Mono | | 22683346 | F3342 | O2a2a1a2a | | 5105 |
| 534 | IV | 801 | 0 | 26 | 281 | 0 | 2 | Mono | Mono | | 6801231 | F4064 | O2a2a1a2a1 | A | 534 |
| 1427 | IV | 827 | 0 | 0 | 283 | 0 | 0 | Mono | Mono | | 8526023 | F1276 | O2a2a1a2a1 | | 1427 |
| 2190 | IV | 827 | 0 | 0 | 283 | 0 | 0 | Mono | Mono | | 14570631 | F4173 | O2a2a1a2a1 | | 2190 |
| 2494 | IV | 827 | 0 | 0 | 282 | 0 | 1 | Mono | Mono | | 15043284 | F1898 | O2a2a1a2a1 | | 2494 |
| 3382 | IV | 827 | 0 | 0 | 283 | 0 | 0 | Mono | Mono | | 17236531 | F2428 | O2a2a1a2a1 | | 3382 |
| 5336 | IV | 827 | 0 | 0 | 283 | 0 | 0 | Mono | Mono | | 23119516 | F3306 | O2a2a1a2a1 | | 5336 |

| | | M9+ | | | M9- | | | | | | Build 37 | | ISOGG | | |
|---|---|---|---|---|---|---|---|---|---|---|---|---|---|---|---|
| List | Stage | 0 | 1 | . | 0 | 1 | . | M9? | Clade(s) | Prob. | Position | Name | Subgroup Name | Amb. | List |
| 3245 | IV | 827 | 0 | 0 | 283 | 0 | 0 | Mono | Mono | | 16880955 | F2309 | O2a2a1a2a1a1 | | 3245 |
| 3637 | IV | 827 | 0 | 0 | 283 | 0 | 0 | Mono | Mono | | 17815008 | ZS3748.3 | O2a2a1a2a1a1 | A | 3637 |
| 4327 | IV | 827 | 0 | 0 | 283 | 0 | 0 | Mono | Mono | | 19566267 | F3085 | O2a2a1a2a1a1 | | 4327 |
| 4724 | IV | 824 | 0 | 3 | 283 | 0 | 0 | Mono | Mono | | 21768133 | M113 | O2a2a1a2a1a1a1a | | 4724 |
| 2304 | IV | 827 | 0 | 0 | 283 | 0 | 0 | Mono | Mono | | 14851538 | N5 | O2a2a1a2a1a1a2a1a1 | | 2304 |
| 1311 | IV | 784 | 0 | 43 | 265 | 0 | 18 | Mono | Mono | | 8398931 | F131 | O2a2b | | 1311 |
| 1951 | IV | 827 | 0 | 0 | 283 | 0 | 0 | Mono | Mono | | 14001232 | P164 | O2a2b | | 1951 |
| 2835 | IV | 827 | 0 | 0 | 283 | 0 | 0 | Mono | Mono | | 15783643 | CTS4723 | O2a2b | | 2835 |
| 2846 | IV | 827 | 0 | 0 | 283 | 0 | 0 | Mono | Mono | | 15810743 | F299 | O2a2b | | 2846 |
| 3534 | IV | 827 | 0 | 0 | 282 | 0 | 1 | Mono | Mono | | 17594625 | F422 | O2a2b | | 3534 |
| 3583 | IV | 827 | 0 | 0 | 283 | 0 | 0 | Mono | Mono | | 17703547 | F427 | O2a2b | | 3583 |
| 5265 | IV | 827 | 0 | 0 | 283 | 0 | 0 | Mono | Mono | | 22958012 | CTS11109 | O2a2b | | 5265 |
| 891 | IV | 827 | 0 | 0 | 283 | 0 | 0 | Mono | Mono | | 7562488 | F85 | O2a2b1 | | 891 |
| 1041 | IV | 827 | 0 | 0 | 283 | 0 | 0 | Mono | Mono | | 7809769 | M1546 | O2a2b1 | | 1041 |
| 1213 | IV | 827 | 0 | 0 | 282 | 0 | 1 | Mono | Mono | | 8221357 | M1556 | O2a2b1 | | 1213 |
| 1426 | IV | 827 | 0 | 0 | 283 | 0 | 0 | Mono | Mono | | 8524263 | F151 | O2a2b1 | | 1426 |
| 3700 | IV | 827 | 0 | 0 | 283 | 0 | 0 | Mono | Mono | | 17927014 | M1667 | O2a2b1a | A | 3700 |
| 734 | IV | 792 | 0 | 35 | 267 | 0 | 16 | Mono | Mono | | 7076840 | M1531 | O2a2b1a1 | | 734 |
| 1360 | IV | 827 | 0 | 0 | 283 | 0 | 0 | Mono | Mono | | 8461538 | M1564 | O2a2b1a1 | | 1360 |
| 1385 | IV | 827 | 0 | 0 | 283 | 0 | 0 | Mono | Mono | | 8480879 | F144 | O2a2b1a1 | | 1385 |
| 1993 | IV | 827 | 0 | 0 | 283 | 0 | 0 | Mono | Mono | | 14077529 | M1587 | O2a2b1a1 | | 1993 |
| 2908 | IV | 824 | 0 | 3 | 282 | 0 | 1 | Mono | Mono | | 15999244 | Page23 | O2a2b1a1 | | 2908 |
| 2911 | IV | 472 | 0 | 355 | 169 | 0 | 114 | Mono | Mono | | 16028396 | M1619 | O2a2b1a1 | | 2911 |
| 3242 | IV | 827 | 0 | 0 | 283 | 0 | 0 | Mono | Mono | | 16880085 | M1636 | O2a2b1a1 | | 3242 |
| 3312 | IV | 827 | 0 | 0 | 283 | 0 | 0 | Mono | Mono | | 17014495 | M1638 | O2a2b1a1 | | 3312 |
| 3826 | IV | 824 | 0 | 3 | 282 | 0 | 1 | Mono | Mono | | 18209413 | M1671 | O2a2b1a1 | | 3826 |
| 4473 | IV | 827 | 0 | 0 | 283 | 0 | 0 | Mono | Mono | | 21353681 | M1692 | O2a2b1a1 | | 4473 |
| 5051 | IV | 827 | 0 | 0 | 283 | 0 | 0 | Mono | Mono | | 22558796 | M1702 | O2a2b1a1 | | 5051 |
| 5386 | IV | 827 | 0 | 0 | 283 | 0 | 0 | Mono | Mono | | 23246261 | M1720 | O2a2b1a1 | A | 5386 |
| 498 | IV | 827 | 0 | 0 | 283 | 0 | 0 | Mono | Mono | | 6716850 | M6539 | O2a2b1a1a1a1a1a1 | | 498 |
| 763 | IV | 827 | 0 | 0 | 283 | 0 | 0 | Mono | Mono | | 7147695 | M1532 | O2a2b1a1a1c | | 763 |
| 947 | IV | 827 | 0 | 0 | 283 | 0 | 0 | Mono | Mono | | 7650862 | M1543 | O2a2b1a1a1c | | 947 |
| 5420 | IV | 827 | 0 | 0 | 283 | 0 | 0 | Mono | Mono | | 23311699 | M1722 | O2a2b1a1a1c1a1a | | 5420 |
| 2605 | IV | 827 | 0 | 0 | 283 | 0 | 0 | Mono | Mono | | 15377325 | M1604 | O2a2b1a1a1c1a1a2 | | 2605 |
| 1257 | IV | 827 | 0 | 0 | 282 | 0 | 1 | Mono | Mono | | 8287876 | F122 | O2a2b1a2 | | 1257 |
| 1273 | IV | 821 | 0 | 6 | 281 | 0 | 2 | Mono | Mono | | 8331546 | Y7101 | O2a2b1a2 | | 1273 |
| 1576 | IV | 827 | 0 | 0 | 283 | 0 | 0 | Mono | Mono | | 8790454 | Y7103^^ | O2a2b1a2 | | 1576 |
| 5619 | IV | 827 | 0 | 0 | 283 | 0 | 0 | Mono | Mono | | 23972702 | F667^^ | O2a2b1a2 | | 5619 |
| 1203 | IV | 827 | 0 | 0 | 283 | 0 | 0 | Mono | Mono | | 8198149 | F701 | O2a2b1a2a | | 1203 |
| 2392 | IV | 827 | 0 | 0 | 283 | 0 | 0 | Mono | Mono | | 14952697 | Page125 | O2a2b1a2a | | 2392 |
| 3423 | IV | 827 | 0 | 0 | 283 | 0 | 0 | Mono | Mono | | 17341468 | F403 | O2a2b1a2a | | 3423 |
| 3533 | IV | 809 | 0 | 18 | 275 | 0 | 8 | Mono | Mono | | 17591656 | CTS7655 | O2a2b1a2a | | 3533 |
| 3670 | IV | 827 | 0 | 0 | 283 | 0 | 0 | Mono | Mono | | 17880466 | F444 | O2a2b1a2a | | 3670 |
| 3925 | IV | 827 | 0 | 0 | 282 | 0 | 1 | Mono | Mono | | 18657388 | Y19 | O2a2b1a2a | | 3925 |
| 4701 | IV | 827 | 0 | 0 | 283 | 0 | 0 | Mono | Mono | | 21757881 | F593 | O2a2b1a2a | | 4701 |
| 1180 | IV | 815 | 0 | 12 | 280 | 0 | 3 | Mono | Mono | | 8133085 | F114 | O2a2b1a2a1a2 | | 1180 |
| 155 | IV | 827 | 0 | 0 | 282 | 0 | 1 | Mono | Mono | | 2851755 | F743 | O2a2b1a2a2 | | 155 |
| 2052 | IV | 826 | 0 | 1 | 282 | 0 | 1 | Mono | Mono | | 14202520 | F1725 | O2a2b1a2a2 | | 2052 |
| 2252 | IV | 826 | 0 | 1 | 283 | 0 | 0 | Mono | Mono | | 14776420 | F1835 | O2a2b1a2a2 | | 2252 |
| 3633 | IV | 825 | 0 | 2 | 283 | 0 | 0 | Mono | Mono | | 17806531 | F2600 | O2a2b1a2a2 | | 3633 |
| 889 | IV | 827 | 0 | 0 | 283 | 0 | 0 | Mono | Mono | | 7558650 | F996 | O2a2b2 | | 889 |
| 1716 | IV | 827 | 0 | 0 | 283 | 0 | 0 | Mono | Mono | | 9763046 | S3543 | O2a2b2 | | 1716 |
| 1959 | IV | 826 | 0 | 1 | 283 | 0 | 0 | Mono | Mono | | 14020486 | F1672 | O2a2b2 | | 1959 |
| 2001 | IV | 827 | 0 | 0 | 283 | 0 | 0 | Mono | Mono | | 14094513 | F1693 | O2a2b2 | | 2001 |
| 2875 | IV | 827 | 0 | 0 | 283 | 0 | 0 | Mono | Mono | | 15899637 | F2029 | O2a2b2 | | 2875 |
| 2943 | IV | 827 | 0 | 0 | 283 | 0 | 0 | Mono | Mono | | 16201548 | F2083 | O2a2b2 | | 2943 |
| 3008 | IV | 827 | 0 | 0 | 283 | 0 | 0 | Mono | Mono | | 16348471 | F2139 | O2a2b2 | | 3008 |
| 3745 | IV | 827 | 0 | 0 | 283 | 0 | 0 | Mono | Mono | | 18036943 | F2683 | O2a2b2 | | 3745 |
| 4610 | IV | 827 | 0 | 0 | 283 | 0 | 0 | Mono | Mono | | 21644600 | F3223 | O2a2b2 | | 4610 |
| 4643 | IV | 827 | 0 | 0 | 283 | 0 | 0 | Mono | Mono | | 21711624 | F3237 | O2a2b2 | | 4643 |
| 639 | IV | 827 | 0 | 0 | 283 | 0 | 0 | Mono | Mono | | 6932191 | F871 | O2a2b2a | | 639 |
| 4843 | IV | 824 | 0 | 3 | 283 | 0 | 0 | Mono | Mono | | 21897363 | N7 | O2a2b2a1 | | 4843 |
| 19 | IV | 827 | 0 | 0 | 283 | 0 | 0 | Mono | Mono | | 2659661 | F706 | O2a2b2a2 | | 19 |
| 1774 | IV | 826 | 0 | 1 | 283 | 0 | 0 | Mono | Mono | | 9904495 | F1645 | O2a2b2a2 | | 1774 |
| 3470 | IV | 827 | 0 | 0 | 283 | 0 | 0 | Mono | Mono | | 17460102 | F2486 | O2a2b2a2 | | 3470 |
| 4302 | IV | 827 | 0 | 0 | 283 | 0 | 0 | Mono | Mono | | 19490561 | F3069 | O2a2b2a2 | | 4302 |
| 57 | IV | 827 | 0 | 0 | 283 | 0 | 0 | Mono | Mono | | 2719706 | F717 | O2a2b2a2a | | 57 |
| 865 | IV | 826 | 0 | 1 | 283 | 0 | 0 | Mono | Mono | | 7521214 | F976 | O2a2b2a2a | | 865 |
| 906 | IV | 827 | 0 | 0 | 283 | 0 | 0 | Mono | Mono | | 7585706 | F1010 | O2a2b2a2a | | 906 |
| 2434 | IV | 827 | 0 | 0 | 283 | 0 | 0 | Mono | Mono | | 15013866 | F1884 | O2a2b2a2a | | 2434 |
| 3796 | IV | 827 | 0 | 0 | 283 | 0 | 0 | Mono | Mono | | 18128673 | F2724 | O2a2b2a2a | | 3796 |
| 4120 | IV | 827 | 0 | 0 | 283 | 0 | 0 | Mono | Mono | | 19078299 | F2929 | O2a2b2a2a | | 4120 |
| 4344 | IV | 827 | 0 | 0 | 283 | 0 | 0 | Mono | Mono | | 20828172 | F3095 | O2a2b2a2a | | 4344 |
| 4362 | IV | 827 | 0 | 0 | 283 | 0 | 0 | Mono | Mono | | 21068933 | F3820 | O2a2b2a2a | | 4362 |
| 4633 | IV | 827 | 0 | 0 | 283 | 0 | 0 | Mono | Mono | | 21683350 | F3230 | O2a2b2a2a | | 4633 |
| 5519 | IV | 827 | 0 | 0 | 283 | 0 | 0 | Mono | Mono | | 23568054 | F3473 | O2a2b2a2a | | 5519 |
| 3564 | IV | 796 | 0 | 31 | 279 | 0 | 4 | Mono | Mono | | 17669826 | F3819 | O2a2b2a2a1 | | 3564 |
| 5650 | IV | 827 | 0 | 0 | 283 | 0 | 0 | Mono | Mono | | 24367929 | F3612 | O2a2b2a2a1 | | 5650 |
| 1114 | I | 824 | 3 | 0 | 0 | 283 | 0 | M9+ | 3 – 93 | | 7963031 | S8 | P or K2b2 | | 1114 |
| 1184 | I | 824 | 3 | 0 | 0 | 283 | 0 | M9+ | 3 – 93 | | 8134704 | PF5870 | P or K2b2 | | 1184 |
| 33 | I | 824 | 3 | 0 | 0 | 283 | 0 | M9+ | 3 – 93 | | 2686727 | PF5991 | P1 or K2b2a | | 33 |
| 112 | I | 824 | 3 | 0 | 0 | 283 | 0 | M9+ | 3 – 93 | | 2803717 | PF5845 | P1 or K2b2a | | 112 |
| 464 | I | 824 | 3 | 0 | 0 | 283 | 0 | M9+ | 3 – 93 | | 6630931 | PF5857 | P1 or K2b2a | | 464 |
| 624 | I | 824 | 3 | 0 | 0 | 283 | 0 | M9+ | 3 – 93 | | 6931040 | PF5858 | P1 or K2b2a | | 624 |
| 886 | I | 824 | 3 | 0 | 0 | 283 | 0 | M9+ | 3 – 93 | | 7549445 | PF5861 | P1 or K2b2a | | 886 |
| 934 | I | 824 | 3 | 0 | 0 | 283 | 0 | M9+ | 3 – 93 | | 7628900 | V231 | P1 or K2b2a | | 934 |
| 1060 | III | 823 | 3 | 1 | 0 | 283 | 0 | M9+ | 3 – 93 | | 7848043 | PF5764 | P1 or K2b2a | | 1060 |
| 1096 | I | 824 | 3 | 0 | 0 | 283 | 0 | M9+ | 3 – 93 | | 7917959 | PF5865 | P1 or K2b2a | | 1096 |
| 1163 | I | 824 | 3 | 0 | 0 | 283 | 0 | M9+ | 3 – 93 | | 8096754 | PF5869 | P1 or K2b2a | | 1163 |
| 1237 | I | 824 | 3 | 0 | 0 | 283 | 0 | M9+ | 3 – 93 | | 8257661 | PF5871 | P1 or K2b2a | | 1237 |
| 1270 | I | 824 | 3 | 0 | 0 | 283 | 0 | M9+ | 3 – 93 | | 8326618 | PF5872 | P1 or K2b2a | | 1270 |
| 1649 | I | 824 | 3 | 0 | 0 | 283 | 0 | M9+ | 3 – 93 | | 9098922 | PF5881 | P1 or K2b2a | | 1649 |
| 1748 | I | 824 | 3 | 0 | 0 | 283 | 0 | M9+ | 3 – 93 | | 9841700 | PF5885 | P1 or K2b2a | | 1748 |
| 1769 | III | 824 | 3 | 0 | 0 | 281 | 2 | M9+ | 3 – 93 | | 9885733 | PF5886 | P1 or K2b2a | | 1769 |
| 1826 | I | 824 | 3 | 0 | 0 | 283 | 0 | M9+ | 3 – 93 | | 10077460 | PF5887 | P1 or K2b2a | | 1826 |
| 2223 | I | 3 | 824 | 0 | 283 | 0 | 0 | M9+ | 3 – 93 | | 14679362 | PF5898 | P1 or K2b2a | | 2223 |

| | | M9+ | | | M9- | | | | | | Build 37 | | ISOGG | | |
|---|---|---|---|---|---|---|---|---|---|---|---|---|---|---|---|
| List | Stage | 0 | 1 | . | 0 | 1 | . | M9? | Clade(s) | Prob. | Position | Name | Subgroup Name | Amb. | List |
| 2310 | I | 3 | 824 | 0 | 283 | 0 | 0 | M9+ | 3 – 93 | | 14864191 | PF5900 | P1 or K2b2a | | 2310 |
| 2524 | III | 3 | 823 | 1 | 281 | 0 | 2 | M9+ | 3 – 93 | | 15111148 | PF5904 | P1 or K2b2a | | 2524 |
| 2677 | I | 824 | 3 | 0 | 0 | 283 | 0 | M9+ | 3 – 93 | | 15497285 | PF5908 | P1 or K2b2a | | 2677 |
| 2773 | I | 824 | 3 | 0 | 0 | 283 | 0 | M9+ | 3 – 93 | | 15648415 | M1109 | P1 or K2b2a | | 2773 |
| 2883 | III | 815 | 3 | 9 | 0 | 282 | 1 | M9+ | 3 – 93 | 1H | 15918592 | PF5909 | P1 or K2b2a | | 2883 |
| 2891 | III | 823 | 3 | 1 | 0 | 283 | 0 | M9+ | 3 – 93 | | 15944828 | M1219 | P1 or K2b2a | A | 2891 |
| 2976 | I | 824 | 3 | 0 | 0 | 283 | 0 | M9+ | 3 – 93 | | 16263188 | PF5912 | P1 or K2b2a | | 2976 |
| 2998 | I | 824 | 3 | 0 | 0 | 283 | 0 | M9+ | 3 – 93 | | 16331558 | PF5914 | P1 or K2b2a | | 2998 |
| 3070 | I | 824 | 3 | 0 | 0 | 283 | 0 | M9+ | 3 – 93 | | 16489668 | PF5915 | P1 or K2b2a | | 3070 |
| 3077 | I | 824 | 3 | 0 | 0 | 283 | 0 | M9+ | 3 – 93 | | 16497020 | PF5916 | P1 or K2b2a | | 3077 |
| 3089 | I | 824 | 3 | 0 | 0 | 283 | 0 | M9+ | 3 – 93 | | 16535034 | PF5917 | P1 or K2b2a | | 3089 |
| 3151 | I | 824 | 3 | 0 | 0 | 283 | 0 | M9+ | 3 – 93 | | 16713287 | PF5920 | P1 or K2b2a | | 3151 |
| 3424 | I | 824 | 3 | 0 | 0 | 283 | 0 | M9+ | 3 – 93 | | 17341676 | PF5923 | P1 or K2b2a | | 3424 |
| 3514 | I | 824 | 3 | 0 | 0 | 283 | 0 | M9+ | 3 – 93 | | 17559652 | PF5928 | P1 or K2b2a | | 3514 |
| 3791 | I | 824 | 3 | 0 | 0 | 283 | 0 | M9+ | 3 – 93 | | 18116596 | PF5934 | P1 or K2b2a | | 3791 |
| 3922 | III | 824 | 3 | 0 | 0 | 282 | 1 | M9+ | 3 – 93 | | 18647146 | PF5936 | P1 or K2b2a | | 3922 |
| 3947 | I | 824 | 3 | 0 | 0 | 283 | 0 | M9+ | 3 – 93 | | 18720028 | PF5937 | P1 or K2b2a | | 3947 |
| 4000 | I | 824 | 3 | 0 | 0 | 283 | 0 | M9+ | 3 – 93 | | 18831084 | PF5940 | P1 or K2b2a | | 4000 |
| 4113 | III | 824 | 3 | 0 | 0 | 282 | 1 | M9+ | 3 – 93 | | 19067469 | PF5943 | P1 or K2b2a | | 4113 |
| 4146 | I | 824 | 3 | 0 | 0 | 283 | 0 | M9+ | 3 – 93 | | 19144375 | PF5945 | P1 or K2b2a | | 4146 |
| 4234 | I | 824 | 3 | 0 | 0 | 283 | 0 | M9+ | 3 – 93 | | 19303108 | M1149 | P1 or K2b2a | | 4234 |
| 4238 | I | 824 | 3 | 0 | 0 | 283 | 0 | M9+ | 3 – 93 | | 19315988 | PF5949 | P1 or K2b2a | | 4238 |
| 4332 | I | 824 | 3 | 0 | 0 | 283 | 0 | M9+ | 3 – 93 | | 20812228 | PF5951 | P1 or K2b2a | | 4332 |
| 4568 | I | 824 | 3 | 0 | 0 | 283 | 0 | M9+ | 3 – 93 | | 21570974 | PF5957 | P1 or K2b2a | | 4568 |
| 4576 | I | 824 | 3 | 0 | 0 | 283 | 0 | M9+ | 3 – 93 | | 21603164 | PF5958 | P1 or K2b2a | | 4576 |
| 4917 | I | 824 | 3 | 0 | 0 | 283 | 0 | M9+ | 3 – 93 | | 21990257 | PF5964 | P1 or K2b2a | | 4917 |
| 4921 | I | 824 | 3 | 0 | 0 | 283 | 0 | M9+ | 3 – 93 | | 22003770 | YSC0000227 | P1 or K2b2a | | 4921 |
| 5032 | I | 824 | 3 | 0 | 0 | 283 | 0 | M9+ | 3 – 93 | | 22514081 | PF5972 | P1 or K2b2a | | 5032 |
| 5395 | I | 824 | 3 | 0 | 0 | 283 | 0 | M9+ | 3 – 93 | | 23257819 | PF5975 | P1 or K2b2a | | 5395 |
| 5464 | I | 824 | 3 | 0 | 0 | 283 | 0 | M9+ | 3 – 93 | | 23431541 | PF5977 | P1 or K2b2a | | 5464 |
| 5486 | III | 811 | 3 | 13 | 0 | 283 | 0 | M9+ | 3 – 93 | | 23489203 | PF5978 | P1 or K2b2a | | 5486 |
| 5629 | III | 824 | 3 | 0 | 0 | 282 | 1 | M9+ | 3 – 93 | | 23979899 | PF5981 | P1 or K2b2a | | 5629 |
| 5637 | I | 824 | 3 | 0 | 0 | 283 | 0 | M9+ | 3 – 93 | | 23993156 | PF5982 | P1 or K2b2a | | 5637 |
| 5673 | III | 819 | 3 | 5 | 0 | 283 | 0 | M9+ | 3 – 93 | | 24443836 | PF5984 | P1 or K2b2a | | 5673 |
| 1274 | I | 824 | 3 | 0 | 0 | 283 | 0 | M9+ | 3 – 93 | | 8334875 | PF5873 | P1~ or K2b2a~ | | 1274 |
| 1538 | I | 824 | 3 | 0 | 0 | 283 | 0 | M9+ | 3 – 93 | | 8685083 | PF5874 | P1~ or K2b2a~ | | 1538 |
| 1597 | I | 824 | 3 | 0 | 0 | 283 | 0 | M9+ | 3 – 93 | | 8845380 | PF5879 | P1~ or K2b2a~ | | 1597 |
| 2141 | I | 3 | 824 | 0 | 283 | 0 | 0 | M9+ | 3 – 93 | | 14433100 | PF5896 | P1~ or K2b2a~ | | 2141 |
| 2338 | I | 3 | 824 | 0 | 283 | 0 | 0 | M9+ | 3 – 93 | | 14898094 | PF5901 | P1~ or K2b2a~ | | 2338 |
| 2671 | III | 792 | 3 | 32 | 0 | 283 | 0 | M9+ | 3 – 93 | | 15480547 | YSC0000251 | P1~ or K2b2a~ | | 2671 |
| 3475 | I | 824 | 3 | 0 | 0 | 283 | 0 | M9+ | 3 – 93 | | 17470112 | PF5925 | P1~ or K2b2a~ | | 3475 |
| 3487 | I | 824 | 3 | 0 | 0 | 283 | 0 | M9+ | 3 – 93 | | 17508337 | PF5927 | P1~ or K2b2a~ | | 3487 |
| 3671 | I | 824 | 3 | 0 | 0 | 283 | 0 | M9+ | 3 – 93 | | 17881230 | PF5930 | P1~ or K2b2a~ | | 3671 |
| 3737 | I | 824 | 3 | 0 | 0 | 283 | 0 | M9+ | 3 – 93 | | 18028661 | PF5932 | P1~ or K2b2a~ | | 3737 |
| 4070 | I | 824 | 3 | 0 | 0 | 283 | 0 | M9+ | 3 – 93 | | 18964263 | PF5941 | P1~ or K2b2a~ | | 4070 |
| 4176 | I | 824 | 3 | 0 | 0 | 283 | 0 | M9+ | 3 – 93 | | 19198212 | PF5946 | P1~ or K2b2a~ | | 4176 |
| 4747 | I | 824 | 3 | 0 | 0 | 283 | 0 | M9+ | 3 – 93 | | 21811108 | YSC0000297 | P1~ or K2b2a~ | | 4747 |
| 4779 | I | 824 | 3 | 0 | 0 | 283 | 0 | M9+ | 3 – 93 | | 21867787 | PF5962 | P1~ or K2b2a~ | | 4779 |
| 4822 | III | 821 | 3 | 3 | 0 | 280 | 3 | M9+ | 3 – 93 | | 21889767 | PF5963 | P1~ or K2b2a~ | | 4822 |
| 4943 | I | 824 | 3 | 0 | 0 | 283 | 0 | M9+ | 3 – 93 | | 22072340 | P284 | P1~ or K2b2a~ | | 4943 |
| 4949 | I | 824 | 3 | 0 | 0 | 283 | 0 | M9+ | 3 – 93 | | 22094491 | PF5966 | P1~ or K2b2a~ | | 4949 |
| 5427 | I | 824 | 3 | 0 | 0 | 283 | 0 | M9+ | 3 – 93 | | 23342709 | YSC0000274 | P1~ or K2b2a~ | | 5427 |
| 636 | IV | 827 | 0 | 0 | 283 | 0 | 0 | Mono | Mono | | 6932163 | M3873.2 | P2b1 | | 636 |
| 120 | II | 826 | 1 | 0 | 283 | 0 | 0 | M9+ | 3 | | 2810917 | V1090 | Q | | 120 |
| 525 | II | 826 | 1 | 0 | 283 | 0 | 0 | M9+ | 3 | | 6778043 | M1064 | Q | | 525 |
| 1172 | II | 826 | 1 | 0 | 283 | 0 | 0 | M9+ | 3 | | 8119406 | M1074 | Q | | 1172 |
| 1384 | II | 826 | 1 | 0 | 283 | 0 | 0 | M9+ | 3 | | 8479245 | V2083.2 | Q | A | 1384 |
| 2057 | II | 826 | 1 | 0 | 283 | 0 | 0 | M9+ | 3 | | 14210817 | M1092 | Q | | 2057 |
| 2235 | II | 826 | 1 | 0 | 283 | 0 | 0 | M9+ | 3 | | 14746296 | Z801 | Q | | 2235 |
| 2442 | II | 826 | 1 | 0 | 283 | 0 | 0 | M9+ | 3 | | 15018582 | M242 | Q | | 2442 |
| 2532 | II | 826 | 1 | 0 | 283 | 0 | 0 | M9+ | 3 | | 15134176 | M1105 | Q | | 2532 |
| 2834 | III | 823 | 1 | 3 | 282 | 0 | 1 | M9+ | 3 | | 15781852 | M997 | Q | | 2834 |
| 3499 | III | 820 | 1 | 6 | 282 | 0 | 1 | M9+ | 3 | | 17516095 | S432 | Q | | 3499 |
| 3636 | III | 826 | 1 | 0 | 281 | 0 | 2 | M9+ | 3 | | 17814673 | M1020 | Q | | 3636 |
| 4283 | II | 826 | 1 | 0 | 283 | 0 | 0 | M9+ | 3 | | 19431928 | V4116 | Q | | 4283 |
| 4410 | III | 825 | 1 | 1 | 283 | 0 | 0 | M9+ | 3 | | 21193181 | M1154 | Q | | 4410 |
| 4919 | III | 824 | 1 | 2 | 283 | 0 | 0 | M9+ | 3 | | 22001258 | M1166 | Q | | 4919 |
| 67 | IV | 826 | 0 | 1 | 283 | 0 | 0 | Mono | Mono | | 2727311 | V1032 | Q1 | | 67 |
| 541 | IV | 827 | 0 | 0 | 283 | 0 | 0 | Mono | Mono | | 6812699 | Z796 | Q1 | | 541 |
| 706 | IV | 827 | 0 | 0 | 283 | 0 | 0 | Mono | Mono | | 7014317 | L472 | Q1 | | 706 |
| 1595 | IV | 827 | 0 | 0 | 283 | 0 | 0 | Mono | Mono | | 8840134 | M1083 | Q1 | | 1595 |
| 1999 | IV | 827 | 0 | 0 | 283 | 0 | 0 | Mono | Mono | | 14092227 | M1087 | Q1 | | 1999 |
| 2028 | IV | 826 | 0 | 1 | 280 | 0 | 3 | Mono | Mono | | 14151622 | M1088 | Q1 | | 2028 |
| 2529 | IV | 815 | 0 | 12 | 282 | 0 | 1 | Mono | Mono | | 15125752 | M1104 | Q1 | | 2529 |
| 2940 | IV | 827 | 0 | 0 | 282 | 0 | 1 | Mono | Mono | | 16195023 | Z804 | Q1 | | 2940 |
| 3069 | IV | 752 | 0 | 75 | 257 | 0 | 26 | Mono | Mono | | 16487674 | M1122 | Q1 | | 3069 |
| 3314 | IV | 827 | 0 | 0 | 283 | 0 | 0 | Mono | Mono | | 17030116 | M1128 | Q1 | | 3314 |
| 3549 | IV | 827 | 0 | 0 | 283 | 0 | 0 | Mono | Mono | | 17616914 | M1136 | Q1 | | 3549 |
| 3720 | IV | 825 | 0 | 2 | 282 | 0 | 1 | Mono | Mono | | 17970530 | M1140 | Q1 | | 3720 |
| 3739 | IV | 826 | 0 | 1 | 283 | 0 | 0 | Mono | Mono | | 18029008 | L528 | Q1 | | 3739 |
| 3971 | IV | 827 | 0 | 0 | 283 | 0 | 0 | Mono | Mono | | 18775212 | V3486 | Q1 | | 3971 |
| 4432 | IV | 827 | 0 | 0 | 283 | 0 | 0 | Mono | Mono | | 21254696 | M1155 | Q1 | | 4432 |
| 4524 | IV | 827 | 0 | 0 | 283 | 0 | 0 | Mono | Mono | | 21455411 | M1158 | Q1 | | 4524 |
| 4974 | IV | 827 | 0 | 0 | 283 | 0 | 0 | Mono | Mono | | 22155597 | M1168 | Q1 | | 4974 |
| 5177 | IV | 826 | 0 | 1 | 283 | 0 | 0 | Mono | Mono | | 22768373 | M1170 | Q1 | | 5177 |
| 5269 | IV | 827 | 0 | 0 | 283 | 0 | 0 | Mono | Mono | | 22970360 | M1173 | Q1 | | 5269 |
| 1354 | IV | 827 | 0 | 0 | 283 | 0 | 0 | Mono | Mono | | 8454150 | F1215 | Q1a | | 1354 |
| 1404 | IV | 827 | 0 | 0 | 283 | 0 | 0 | Mono | Mono | | 8491335 | F1251 | Q1a | | 1404 |
| 4656 | IV | 827 | 0 | 0 | 283 | 0 | 0 | Mono | Mono | | 21723218 | F3243 | Q1a | | 4656 |
| 178 | IV | 827 | 0 | 0 | 283 | 0 | 0 | Mono | Mono | | 2888083 | NWT01 | Q1a1 | | 178 |
| 2478 | IV | 826 | 0 | 1 | 283 | 0 | 0 | Mono | Mono | | 15030650 | N14 | Q1a1a | | 2478 |
| 4776 | IV | 827 | 0 | 0 | 283 | 0 | 0 | Mono | Mono | | 21866664 | M25 | Q1a2 | | 4776 |
| 4823 | IV | 824 | 0 | 3 | 283 | 0 | 0 | Mono | Mono | | 21889818 | M143 | Q1a2 | | 4823 |
| 562 | IV | 826 | 0 | 1 | 283 | 0 | 0 | Mono | Mono | | 6851494 | L712 | Q1a2a | | 562 |
| 47 | II | 826 | 1 | 0 | 283 | 0 | 0 | M9+ | NU | | 2710308 | L697.2 | Q1a2a1~ | W | 47 |
| 176 | IV | 825 | 0 | 2 | 283 | 0 | 0 | Mono | Mono | | 2887156 | M346 | Q1b | | 176 |

| | | M9+ | | | M9- | | | | | | Build 37 | | ISOGG | | |
|---|---|---|---|---|---|---|---|---|---|---|---|---|---|---|---|
| List | Stage | 0 | 1 | . | 0 | 1 | . | M9? | Clade(s) | Prob. | Position | Name | Subgroup Name | Amb. | List |
| 1189 | IV | 827 | 0 | 0 | 283 | 0 | 0 | Mono | Mono | | 8148869 | S324 | Q1b | | 1189 |
| 1405 | IV | 827 | 0 | 0 | 283 | 0 | 0 | Mono | Mono | | 8493022 | V2131 | Q1b | | 1405 |
| 2485 | IV | 827 | 0 | 0 | 283 | 0 | 0 | Mono | Mono | | 15033712 | L474 | Q1b | | 2485 |
| 2726 | IV | 826 | 0 | 1 | 283 | 0 | 0 | Mono | Mono | | 15574102 | L57 | Q1b | | 2726 |
| 2832 | IV | 827 | 0 | 0 | 283 | 0 | 0 | Mono | Mono | | 15781062 | Z802 | Q1b | | 2832 |
| 2984 | IV | 827 | 0 | 0 | 283 | 0 | 0 | Mono | Mono | | 16305155 | M1118 | Q1b | | 2984 |
| 3432 | IV | 827 | 0 | 0 | 283 | 0 | 0 | Mono | Mono | | 17359287 | M1132 | Q1b | | 3432 |
| 1223 | IV | 827 | 0 | 0 | 283 | 0 | 0 | Mono | Mono | | 8235033 | SA23 | Q1b1 | | 1223 |
| 3800 | IV | 826 | 0 | 1 | 283 | 0 | 0 | Mono | Mono | | 18146921 | L475 | Q1b1 | | 3800 |
| 4277 | IV | 827 | 0 | 0 | 283 | 0 | 0 | Mono | Mono | | 19413335 | L55 | Q1b1 | | 4277 |
| 4607 | IV | 827 | 0 | 0 | 283 | 0 | 0 | Mono | Mono | | 21642296 | S326 | Q1b1 | | 4607 |
| 5414 | IV | 827 | 0 | 0 | 283 | 0 | 0 | Mono | Mono | | 23292782 | L54 | Q1b1a | | 5414 |
| 5448 | IV | 827 | 0 | 0 | 282 | 0 | 1 | Mono | Mono | | 23391298 | M930 | Q1b1a1 | | 5448 |
| 3416 | IV | 827 | 0 | 0 | 283 | 0 | 0 | Mono | Mono | | 17317841 | Z788 | Q1b1a1a | | 3416 |
| 4125 | IV | 827 | 0 | 0 | 283 | 0 | 0 | Mono | Mono | | 19096363 | M3 | Q1b1a1a | | 4125 |
| 5449 | IV | 827 | 0 | 0 | 283 | 0 | 0 | Mono | Mono | | 23391307 | M931 | Q1b1a1a | A | 5449 |
| 4669 | IV | 827 | 0 | 0 | 283 | 0 | 0 | Mono | Mono | | 21733231 | M19 | Q1b1a1a1a~ | | 4669 |
| 2435 | IV | 827 | 0 | 0 | 283 | 0 | 0 | Mono | Mono | | 15014550 | M194 | Q1b1a1a1b~ | | 2435 |
| 2481 | IV | 827 | 0 | 0 | 283 | 0 | 0 | Mono | Mono | | 15030767 | P292 | Q1b1a1a1c~ | A | 2481 |
| 1010 | IV | 827 | 0 | 0 | 283 | 0 | 0 | Mono | Mono | | 7742989 | Z777 | Q1b1a2 | | 1010 |
| 1982 | IV | 827 | 0 | 0 | 283 | 0 | 0 | Mono | Mono | | 14064827 | M981 | Q1b1a2 | | 1982 |
| 2146 | IV | 827 | 0 | 0 | 283 | 0 | 0 | Mono | Mono | | 14449243 | M988 | Q1b1a2 | | 2146 |
| 4570 | IV | 827 | 0 | 0 | 283 | 0 | 0 | Mono | Mono | | 21573011 | Z791 | Q1b1a2 | | 4570 |
| 5307 | IV | 827 | 0 | 0 | 283 | 0 | 0 | Mono | Mono | | 23060094 | Z795 | Q1b1a2 | | 5307 |
| 3421 | IV | 827 | 0 | 0 | 283 | 0 | 0 | Mono | Mono | | 17335310 | Z789 | Q1b1a2a1b | A | 3421 |
| 2352 | IV | 827 | 0 | 0 | 283 | 0 | 0 | Mono | Mono | | 14903417 | CTS3407 | Q1b1a2a1b1a~ | | 2352 |
| 2159 | IV | 826 | 0 | 1 | 282 | 0 | 1 | Mono | Mono | | 14482079 | L334 | Q1b1a3 | | 2159 |
| 3611 | IV | 827 | 0 | 0 | 283 | 0 | 0 | Mono | Mono | | 17766807 | L330 | Q1b1a3 | | 3611 |
| 1442 | IV | 827 | 0 | 0 | 283 | 0 | 0 | Mono | Mono | | 8550111 | Z35941 | Q1b1a3b | | 1442 |
| 555 | IV | 827 | 0 | 0 | 283 | 0 | 0 | Mono | Mono | | 6846280 | L940 | Q1b2b~ | | 555 |
| 2992 | IV | 827 | 0 | 0 | 283 | 0 | 0 | Mono | Mono | | 16316263 | F2125 | Q1b2b~ | | 2992 |
| 593 | IV | 827 | 0 | 0 | 283 | 0 | 0 | Mono | Mono | | 6886008 | L941 | Q1b2b1b2~ | | 593 |
| 5489 | IV | 827 | 0 | 0 | 283 | 0 | 0 | Mono | Mono | | 23496062 | L938 | Q1b2b1b2~ | | 5489 |
| 35 | III | 826 | 1 | 0 | 282 | 0 | 1 | M9+ | 3 | | 2692142 | L612 | Q2 | | 35 |
| 481 | II | 826 | 1 | 0 | 283 | 0 | 0 | M9+ | 3 | | 6679787 | F803 | Q2 | | 481 |
| 1140 | II | 826 | 1 | 0 | 283 | 0 | 0 | M9+ | 3 | | 8039511 | F108 | Q2 | A | 1140 |
| 4123 | II | 826 | 1 | 0 | 283 | 0 | 0 | M9+ | 3 | | 19089909 | F2934 | Q2 | | 4123 |
| 4143 | II | 826 | 1 | 0 | 283 | 0 | 0 | M9+ | 3 | | 19136888 | L275 | Q2 | | 4143 |
| 1352 | III | 825 | 1 | 1 | 283 | 0 | 0 | M9+ | 3 | | 8451013 | F1213 | Q2a | | 1352 |
| 2065 | II | 826 | 1 | 0 | 283 | 0 | 0 | M9+ | 3 | | 14235022 | F1734 | Q2a | | 2065 |
| 2331 | II | 826 | 1 | 0 | 283 | 0 | 0 | M9+ | 3 | | 14889974 | S325 | Q2a | | 2331 |
| 2468 | II | 826 | 1 | 0 | 283 | 0 | 0 | M9+ | 3 | | 15027507 | Page100 | Q2a1 | | 2468 |
| 4866 | II | 826 | 1 | 0 | 283 | 0 | 0 | M9+ | 3 | | 21906607 | S289 | Q2a1 | | 4866 |
| 797 | II | 826 | 1 | 0 | 283 | 0 | 0 | M9+ | 3 | | 7219594 | Y2121 | Q2a1a | | 797 |
| 3756 | II | 826 | 1 | 0 | 283 | 0 | 0 | M9+ | 3 | | 18051798 | Y2209 | Q2a1a1a | | 3756 |
| 1525 | III | 826 | 1 | 0 | 281 | 0 | 2 | M9+ | 3 | | 8662585 | Y2224 | Q2a1a1a1 | | 1525 |
| 2634 | IV | 827 | 0 | 0 | 283 | 0 | 0 | Mono | Mono | | 15436590 | YP734 | Q2a1a1a2b | | 2634 |
| 907 | IV | 826 | 0 | 1 | 282 | 0 | 1 | Mono | Mono | | 7586919 | V220.2 | Q2a1a2a1a1a2~ | A | 907 |
| 2403 | IV | 827 | 0 | 0 | 283 | 0 | 0 | Mono | Mono | | 14959017 | BZ5065 | Q2a1a4b~ | | 2403 |
| 96 | IV | 826 | 0 | 1 | 283 | 0 | 0 | Mono | Mono | | 2765038 | Y2247 | Q2a1c | | 96 |
| 491 | I | 823 | 4 | 0 | 0 | 283 | 0 | M9+ | 4 – 93 | | 6701239 | PF6013 | R | | 491 |
| 581 | I | 823 | 4 | 0 | 0 | 283 | 0 | M9+ | 4 – 93 | | 6868118 | Y472 | R | | 581 |
| 781 | I | 823 | 4 | 0 | 0 | 283 | 0 | M9+ | 4 – 93 | | 7177189 | PF6016 | R | | 781 |
| 885 | III | 821 | 4 | 2 | 0 | 281 | 2 | M9+ | 4 – 93 | | 7548900 | M620 | R | | 885 |
| 1148 | I | 823 | 4 | 0 | 0 | 283 | 0 | M9+ | 4 – 93 | | 8050994 | PF6019 | R | | 1148 |
| 2237 | III | 822 | 4 | 1 | 0 | 283 | 0 | M9+ | 4 – 93 | | 14750668 | YSC0001265 | R | | 2237 |
| 2506 | I | 4 | 823 | 0 | 283 | 0 | 0 | M9+ | 4 – 93 | | 15078469 | PF6037 | R | | 2506 |
| 2731 | I | 823 | 4 | 0 | 0 | 283 | 0 | M9+ | 4 – 93 | | 15581983 | UTY2 | R | | 2731 |
| 2755 | I | 823 | 4 | 0 | 0 | 283 | 0 | M9+ | 4 – 93 | | 15594523 | M685 | R | | 2755 |
| 2789 | I | 823 | 4 | 0 | 0 | 283 | 0 | M9+ | 4 – 93 | | 15667208 | FGC1168 | R | | 2789 |
| 3071 | I | 823 | 4 | 0 | 0 | 283 | 0 | M9+ | 4 – 93 | | 16491135 | M696 | R | | 3071 |
| 3122 | I | 823 | 4 | 0 | 0 | 283 | 0 | M9+ | 4 – 93 | | 16615413 | YSC0000287 | R | | 3122 |
| 3126 | I | 823 | 4 | 0 | 0 | 283 | 0 | M9+ | 4 – 93 | | 16629782 | PF5919 | R | | 3126 |
| 3229 | I | 823 | 4 | 0 | 0 | 283 | 0 | M9+ | 4 – 93 | | 16856357 | Y479 | R | | 3229 |
| 3397 | I | 823 | 4 | 0 | 0 | 283 | 0 | M9+ | 4 – 93 | | 17285993 | PF6050 | R | | 3397 |
| 3729 | I | 823 | 4 | 0 | 0 | 283 | 0 | M9+ | 4 – 93 | | 18017528 | Y482 | R | | 3729 |
| 3762 | I | 823 | 4 | 0 | 0 | 283 | 0 | M9+ | 4 – 93 | | 18066156 | YSC0000201 | R | | 3762 |
| 4216 | III | 821 | 4 | 2 | 0 | 283 | 0 | M9+ | 4 – 93 | | 19267344 | P285 | R | | 4216 |
| 4415 | I | 823 | 4 | 0 | 0 | 283 | 0 | M9+ | 4 – 93 | | 21219443 | Y506 | R | | 4415 |
| 4440 | I | 823 | 4 | 0 | 0 | 283 | 0 | M9+ | 4 – 93 | | 21263029 | PF5953 | R | | 4440 |
| 4761 | I | 823 | 4 | 0 | 0 | 283 | 0 | M9+ | 4 – 93 | | 21843090 | P280 | R | | 4761 |
| 5195 | I | 823 | 4 | 0 | 0 | 283 | 0 | M9+ | 4 – 93 | | 22818334 | YSC0000233 | R | | 5195 |
| 5298 | I | 823 | 4 | 0 | 0 | 283 | 0 | M9+ | 4 – 93 | | 23035132 | P232 | R | | 5298 |
| 5346 | I | 823 | 4 | 0 | 0 | 283 | 0 | M9+ | 4 – 93 | | 23134896 | M799 | R | | 5346 |
| 5645 | III | 821 | 4 | 2 | 0 | 283 | 0 | M9+ | 4 – 93 | | 24360964 | F675 | R | | 5645 |
| 2741 | IV | 823 | 0 | 4 | 282 | 0 | 1 | Mono | Mono | | 15591201 | M204 | R (Notes) | | 2741 |
| 4780 | IV | 827 | 0 | 0 | 283 | 0 | 0 | Mono | Mono | | 21867854 | M157.1 | R (Notes) | | 4780 |
| 4801 | IV | 827 | 0 | 0 | 283 | 0 | 0 | Mono | Mono | | 21872886 | M56 | R (Notes) | | 4801 |
| 4857 | IV | 826 | 0 | 1 | 283 | 0 | 0 | Mono | Mono | | 21903383 | Page44.2 | R (Notes) | A | 4857 |
| 4863 | IV | 827 | 0 | 0 | 283 | 0 | 0 | Mono | Mono | | 21906109 | M87 | R (Notes) | | 4863 |
| 945 | I | 823 | 4 | 0 | 0 | 283 | 0 | M9+ | 4 – 93 | | 7647357 | PF6113 | R1 | | 945 |
| 962 | I | 823 | 4 | 0 | 0 | 283 | 0 | M9+ | 4 – 93 | | 7671535 | PF6114 | R1 | | 962 |
| 1025 | I | 823 | 4 | 0 | 0 | 283 | 0 | M9+ | 4 – 93 | | 7771131 | PF6115 | R1 | | 1025 |
| 1065 | I | 823 | 4 | 0 | 0 | 283 | 0 | M9+ | 4 – 93 | | 7854412 | PF6116 | R1 | | 1065 |
| 1317 | I | 823 | 4 | 0 | 0 | 283 | 0 | M9+ | 4 – 93 | | 8416058 | M632 | R1 | | 1317 |
| 1507 | I | 823 | 4 | 0 | 0 | 283 | 0 | M9+ | 4 – 93 | | 8633545 | PF6117 | R1 | | 1507 |
| 1910 | III | 823 | 4 | 0 | 0 | 281 | 2 | M9+ | 4 – 93 | | 13807475 | Y305 | R1 | | 1910 |
| 2186 | I | 4 | 823 | 0 | 283 | 0 | 0 | M9+ | 4 – 93 | | 14556851 | PF6123 | R1 | | 2186 |
| 2463 | I | 4 | 823 | 0 | 283 | 0 | 0 | M9+ | 4 – 93 | | 15026424 | Page29 | R1 | | 2463 |
| 2739 | I | 823 | 4 | 0 | 0 | 283 | 0 | M9+ | 4 – 93 | | 15590342 | P225 | R1 | | 2739 |
| 2856 | I | 823 | 4 | 0 | 0 | 283 | 0 | M9+ | 4 – 93 | | 15818409 | M689 | R1 | | 2856 |
| 3161 | I | 823 | 4 | 0 | 0 | 283 | 0 | M9+ | 4 – 93 | | 16742224 | YSC0000288 | R1 | | 3161 |
| 3587 | III | 822 | 4 | 1 | 0 | 281 | 2 | M9+ | 4 – 93 | | 17716251 | PF6136 | R1 | | 3587 |
| 3620 | I | 823 | 4 | 0 | 0 | 283 | 0 | M9+ | 4 – 93 | | 17782178 | P236 | R1 | | 3620 |
| 4097 | I | 823 | 4 | 0 | 0 | 283 | 0 | M9+ | 4 – 93 | | 19045552 | YSC0000207 | R1 | | 4097 |
| 4384 | I | 823 | 4 | 0 | 0 | 283 | 0 | M9+ | 4 – 93 | | 21117888 | P234 | R1 | | 4384 |

| | | | M9+ | | | M9- | | | | | Build 37 | | ISOGG | | |
|---|---|---|---|---|---|---|---|---|---|---|---|---|---|---|---|
| List | Stage | 0 | 1 | . | 0 | 1 | . | M9? | Clade(s) | Prob. | Position | Name | Subgroup Name | Amb. | List |
| 4405 | I | 823 | 4 | 0 | 0 | 283 | 0 | M9+ | 4 – 93 | | 21166358 | P233 | R1 | | 4405 |
| 4877 | I | 823 | 4 | 0 | 0 | 283 | 0 | M9+ | 4 – 93 | | 21917832 | M781 | R1 | | 4877 |
| 5077 | I | 823 | 4 | 0 | 0 | 283 | 0 | M9+ | 4 – 93 | | 22601068 | YSC0000230 | R1 | | 5077 |
| 5163 | I | 823 | 4 | 0 | 0 | 283 | 0 | M9+ | 4 – 93 | | 22750583 | S1 | R1 | | 5163 |
| 2019 | III | 741 | 31 | 55 | 263 | 0 | 20 | M9+ | 4 – 16 | | 14138745 | PF6175 | R1a | | 2019 |
| 3678 | I | 796 | 31 | 0 | 283 | 0 | 0 | M9+ | 4 – 16 | | 17891241 | PF6200 | R1a | | 3678 |
| 3807 | I | 796 | 31 | 0 | 283 | 0 | 0 | M9+ | 4 – 16 | | 18162834 | PF6203 | R1a | | 3807 |
| 5477 | III | 796 | 30 | 1 | 283 | 0 | 0 | M9+ | 4 – 16 | MD | 23473201 | PF6229 | R1a | | 5477 |
| 680 | I | 796 | 31 | 0 | 283 | 0 | 0 | M9+ | 4 – 16 | | 6955868 | PF6153 | R1a~ | | 680 |
| 791 | I | 796 | 31 | 0 | 283 | 0 | 0 | M9+ | 4 – 16 | | 7201215 | M616 | R1a~ | | 791 |
| 1150 | I | 796 | 31 | 0 | 283 | 0 | 0 | M9+ | 4 – 16 | | 8053594 | PF6160 | R1a~ | | 1150 |
| 2111 | III | 793 | 31 | 3 | 283 | 0 | 0 | M9+ | 4 – 16 | | 14336840 | PF6179 | R1a~ | | 2111 |
| 3045 | I | 796 | 31 | 0 | 283 | 0 | 0 | M9+ | 4 – 16 | | 16436872 | PF6191 | R1a~ | | 3045 |
| 4145 | III | 765 | 31 | 31 | 276 | 0 | 7 | M9+ | 4 – 16 | | 19141755 | V3820 | R1a~ | | 4145 |
| 5224 | I | 796 | 31 | 0 | 283 | 0 | 0 | M9+ | 4 – 16 | | 22897863 | PF6222 | R1a~ | | 5224 |
| 5515 | III | 795 | 31 | 1 | 282 | 0 | 1 | M9+ | 4 – 16 | | 23559768 | PF7534 | R1a~ | | 5515 |
| 5615 | III | 794 | 31 | 2 | 282 | 0 | 1 | M9+ | 4 – 16 | | 23962104 | PF6233 | R1a~ | | 5615 |
| 5682 | III | 796 | 29 | 2 | 282 | 0 | 1 | M9+ | 4 – 16 | MD | 24462117 | F3650 | R1a~ | | 5682 |
| 13 | I | 796 | 31 | 0 | 283 | 0 | 0 | M9+ | 4 – 16 | | 2657176 | SRY1532.2 | R1a1 | A | 13 |
| 612 | I | 796 | 31 | 0 | 283 | 0 | 0 | M9+ | 4 – 16 | | 6906074 | PF6235 | R1a1 | | 612 |
| 3085 | I | 796 | 31 | 0 | 283 | 0 | 0 | M9+ | 4 – 16 | | 16520444 | PF6237 | R1a1 | | 3085 |
| 812 | I | 796 | 31 | 0 | 283 | 0 | 0 | M9+ | 4 – 16 | | 7260177 | F937 | R1a1~ | | 812 |
| 1301 | III | 795 | 31 | 1 | 283 | 0 | 0 | M9+ | 4 – 16 | | 8384434 | V1855 | R1a1~ | | 1301 |
| 1685 | I | 796 | 31 | 0 | 283 | 0 | 0 | M9+ | 4 – 16 | | 9405443 | PF6168 | R1a1~ | | 1685 |
| 2872 | I | 796 | 31 | 0 | 283 | 0 | 0 | M9+ | 4 – 16 | | 15879017 | PF6236 | R1a1~ | | 2872 |
| 3118 | I | 796 | 31 | 0 | 283 | 0 | 0 | M9+ | 4 – 16 | | 16601264 | PF6194 | R1a1~ | | 3118 |
| 3274 | III | 796 | 17 | 14 | 283 | 0 | 0 | M9+ | 4 – 16 | MD | 16933931 | PF7522 | R1a1~ | | 3274 |
| 4046 | I | 796 | 31 | 0 | 283 | 0 | 0 | M9+ | 4 – 16 | | 18920886 | V3610 | R1a1~ | | 4046 |
| 4499 | III | 793 | 31 | 3 | 281 | 0 | 2 | M9+ | 4 – 16 | | 21400746 | M774 | R1a1~ | | 4499 |
| 1975 | I | 796 | 31 | 0 | 283 | 0 | 0 | M9+ | 4 – 16 | | 14054623 | M515 | R1a1a | | 1975 |
| 2480 | III | 758 | 31 | 38 | 277 | 0 | 6 | M9+ | 4 – 16 | | 15030752 | PF6238 | R1a1a | | 2480 |
| 2991 | I | 796 | 31 | 0 | 283 | 0 | 0 | M9+ | 4 – 16 | | 16315153 | PF6239 | R1a1a | | 2991 |
| 4263 | III | 791 | 31 | 5 | 283 | 0 | 0 | M9+ | 4 – 16 | | 19375294 | PF6240 | R1a1a | | 4263 |
| 883 | III | 786 | 31 | 10 | 279 | 0 | 4 | M9+ | 4 – 16 | | 7545969 | V1190 | R1a1a~ | | 883 |
| 988 | III | 793 | 31 | 3 | 282 | 0 | 1 | M9+ | 4 – 16 | | 7715717 | V1380 | R1a1a~ | | 988 |
| 1680 | IV | 0 | 823 | 4 | 0 | 283 | 0 | Mono | Mono | | 9384631 | F4138 | R1a1a~ | A,W | 1680 |
| 3750 | III | 795 | 31 | 1 | 283 | 0 | 0 | M9+ | 4 – 16 | | 18044223 | PF6201 | R1a1a~ | | 3750 |
| 4485 | I | 796 | 31 | 0 | 283 | 0 | 0 | M9+ | 4 – 16 | | 21366991 | M771 | R1a1a~ | | 4485 |
| 4496 | I | 796 | 31 | 0 | 283 | 0 | 0 | M9+ | 4 – 16 | | 21392030 | M773 | R1a1a~ | | 4496 |
| 5102 | III | 795 | 31 | 1 | 283 | 0 | 0 | M9+ | 4 – 16 | | 22676818 | F3337 | R1a1a~ | | 5102 |
| 1431 | I | 796 | 31 | 0 | 283 | 0 | 0 | M9+ | 4 – 16 | | 8533735 | M417 | R1a1a1 | | 1431 |
| 2173 | I | 796 | 31 | 0 | 283 | 0 | 0 | M9+ | 4 – 16 | | 14498990 | Page7 | R1a1a1 | | 2173 |
| 4157 | III | 795 | 31 | 1 | 278 | 5 | 0 | BOTH | | PI | 19160342 | V3842 | R1a1a1~ | W | 4157 |
| 4418 | I | 796 | 31 | 0 | 283 | 0 | 0 | M9+ | 4 – 16 | | 21225537 | M761 | R1a1a1~ | | 4418 |
| 5570 | I | 796 | 31 | 0 | 283 | 0 | 0 | M9+ | 4 – 16 | | 23800360 | PF6231 | R1a1a1~ | | 5570 |
| 969 | I | 807 | 20 | 0 | 283 | 0 | 0 | M9+ | 6 – 16 | | 7683058 | Z647 | R1a1a1b | | 969 |
| 1231 | I | 807 | 20 | 0 | 283 | 0 | 0 | M9+ | 6 – 16 | | 8245045 | Z645 | R1a1a1b | | 1231 |
| 4279 | I | 807 | 20 | 0 | 283 | 0 | 0 | M9+ | 6 – 16 | | 19419865 | Z651 | R1a1a1b~ | | 4279 |
| 4909 | III | 809 | 17 | 1 | 283 | 0 | 0 | M9+ | 6 – 14 | | 21976303 | Z283 | R1a1a1b1 | | 4909 |
| 2737 | I | 810 | 17 | 0 | 283 | 0 | 0 | M9+ | 6 – 14 | | 15588401 | Z282 | R1a1a1b1a | | 2737 |
| 5647 | II | 826 | 1 | 0 | 283 | 0 | 0 | M9+ | 6 | | 24366464 | PF6241 | R1a1a1b1a1a | | 5647 |
| 3704 | IV | 827 | 0 | 0 | 283 | 0 | 0 | Mono | Mono | | 17937260 | S4554 | R1a1a1b1a1a1c1 | | 3704 |
| 2869 | IV | 827 | 0 | 0 | 283 | 0 | 0 | Mono | Mono | | 15868478 | YP417 | R1a1a1b1a1a1c1c | | 2869 |
| 65 | IV | 819 | 0 | 8 | 280 | 0 | 3 | Mono | Mono | | 2724506 | Z660 | R1a1a1b1a2a | | 65 |
| 3528 | IV | 827 | 0 | 0 | 283 | 0 | 0 | Mono | Mono | | 17585001 | Z92 | R1a1a1b1a2a | | 3528 |
| 3777 | IV | 827 | 0 | 0 | 283 | 0 | 0 | Mono | Mono | | 18082508 | Z661 | R1a1a1b1a2a | | 3777 |
| 816 | II | 825 | 2 | 0 | 283 | 0 | 0 | M9+ | 7 | | 7271722 | S3357 | R1a1a1b1a2b | | 816 |
| 1370 | IV | 827 | 0 | 0 | 283 | 0 | 0 | Mono | Mono | | 8467053 | P278.2 | R1a1a1b1a2b1 | A | 1370 |
| 2337 | III | 825 | 2 | 0 | 273 | 0 | 10 | M9+ | 7 | | 14896020 | V2670 | R1a1a1b1a2b3a | | 2337 |
| 4602 | IV | 827 | 0 | 0 | 283 | 0 | 0 | Mono | Mono | | 21628667 | S468 | R1a1a1b1a2b3a1a1b1 | | 4602 |
| 1556 | I | 813 | 14 | 0 | 283 | 0 | 0 | M9+ | 8 – 14 | | 8717196 | Z284 | R1a1a1b1a3a | | 1556 |
| 627 | III | 811 | 9 | 7 | 281 | 0 | 2 | M9+ | 9 – 11 | | 6931809 | S200 | R1a1a1b1a3a1 | | 627 |
| 2651 | I | 821 | 6 | 0 | 283 | 0 | 0 | M9+ | 10 – 11 | | 15453490 | CTS4179 | R1a1a1b1a3a1a | | 2651 |
| 3402 | II | 825 | 2 | 0 | 283 | 0 | 0 | M9+ | 12 – 13 | | 17293093 | Z287 | R1a1a1b1a3a2 | | 3402 |
| 3812 | II | 825 | 2 | 0 | 283 | 0 | 0 | M9+ | 12 – 13 | | 18175765 | Z288 | R1a1a1b1a3a2 | | 3812 |
| 3294 | II | 826 | 1 | 0 | 283 | 0 | 0 | M9+ | 13 | | 16954812 | Z88 | R1a1a1b1a3a2a1 | | 3294 |
| 2592 | I | 822 | 5 | 0 | 283 | 0 | 0 | M9+ | | PI | 15342222 | V2895.1 | R1a1a1b1a3a3a | A,W | 2592 |
| 887 | II | 824 | 3 | 0 | 283 | 0 | 0 | M9+ | 15 – 16 | | 7552356 | Z93 | R1a1a1b2 | | 887 |
| 5612 | II | 826 | 1 | 0 | 283 | 0 | 0 | M9+ | 16 | | 23956870 | Z95 | R1a1a1b2a~ | | 5612 |
| 5104 | IV | 827 | 0 | 0 | 283 | 0 | 0 | Mono | Mono | | 22679436 | Y6 | R1a1a1b2a1a1a | | 5104 |
| 4294 | IV | 827 | 0 | 0 | 283 | 0 | 0 | Mono | Mono | | 19475863 | Y7 | R1a1a1b2a1a2c | | 4294 |
| 2157 | IV | 827 | 0 | 0 | 283 | 0 | 0 | Mono | Mono | | 14477397 | Z2122 | R1a1a1b2a2b | | 2157 |
| 1488 | IV | 827 | 0 | 0 | 283 | 0 | 0 | Mono | Mono | | 8605676 | Z2472 | R1a1a1b2a2b1 | | 1488 |
| 4215 | IV | 826 | 0 | 1 | 283 | 0 | 0 | Mono | Mono | | 19262425 | F2997 | R1a1a1b2a2b1~ | | 4215 |
| 14 | IV | 827 | 0 | 0 | 283 | 0 | 0 | Mono | Mono | | 2657349 | Z2469 | R1a1a1b2a2b1a | | 14 |
| 2888 | IV | 827 | 0 | 0 | 283 | 0 | 0 | Mono | Mono | | 15934901 | Z874.2 | R1a2a~ | A | 2888 |
| 4056 | III | 825 | 0 | 2 | 282 | 1 | 0 | M9- | NU | | 18945773 | Z16200.2 | R1a2a~ | A,W | 4056 |
| 177 | I | 792 | 35 | 0 | 0 | 283 | 0 | M9+ | 17 – 93 | | 2887824 | PF6242 | R1b | | 177 |
| 1108 | I | 792 | 35 | 0 | 0 | 283 | 0 | M9+ | 17 – 93 | | 7948701 | PF6246 | R1b | | 1108 |
| 1113 | I | 792 | 35 | 0 | 0 | 283 | 0 | M9+ | 17 – 93 | | 7960019 | YSC0000075 | R1b | | 1113 |
| 1342 | III | 786 | 35 | 6 | 0 | 283 | 0 | M9+ | 17 – 93 | | 8439542 | PF6250 | R1b | | 1342 |
| 1672 | I | 792 | 35 | 0 | 0 | 283 | 0 | M9+ | 17 – 93 | | 9170545 | PF6251 | R1b | | 1672 |
| 2041 | I | 792 | 35 | 0 | 0 | 283 | 0 | M9+ | 17 – 93 | | 14193384 | CTS2134 | R1b | | 2041 |
| 4040 | I | 792 | 35 | 0 | 0 | 283 | 0 | M9+ | 17 – 93 | | 18914441 | L278 | R1b | | 4040 |
| 4918 | I | 792 | 35 | 0 | 0 | 283 | 0 | M9+ | 17 – 93 | | 21995972 | PF6267 | R1b | | 4918 |
| 5588 | III | 791 | 35 | 1 | 0 | 283 | 0 | M9+ | 17 – 93 | | 23845409 | PF6270 | R1b | | 5588 |
| 5636 | I | 792 | 35 | 0 | 0 | 283 | 0 | M9+ | 17 – 93 | | 23992762 | PF6272 | R1b | | 5636 |
| 1082 | I | 792 | 35 | 0 | 0 | 283 | 0 | M9+ | 17 – 93 | | 7900883 | Y108 | R1b1 | | 1082 |
| 1209 | III | 792 | 35 | 0 | 0 | 276 | 7 | M9+ | 17 – 93 | | 8214827 | PF6249 | R1b1 | | 1209 |
| 1819 | I | 792 | 35 | 0 | 0 | 283 | 0 | M9+ | 17 – 93 | | 10038192 | A702 | R1b1 | | 1819 |
| 1913 | III | 780 | 35 | 12 | 0 | 283 | 0 | M9+ | 17 – 93 | | 13822833 | Y409 | R1b1 | | 1913 |
| 2209 | III | 35 | 791 | 1 | 283 | 0 | 0 | M9+ | 17 – 93 | | 14637352 | V2515 | R1b1 | | 2209 |
| 2686 | III | 784 | 35 | 8 | 0 | 282 | 1 | M9+ | 17 – 93 | | 15510064 | YSC0001279 | R1b1 | | 2686 |
| 3176 | I | 792 | 35 | 0 | 0 | 283 | 0 | M9+ | 17 – 93 | | 16773870 | YSC0000266 | R1b1 | | 3176 |
| 3736 | I | 792 | 35 | 0 | 0 | 283 | 0 | M9+ | 17 – 93 | | 18026855 | PF6259 | R1b1 | | 3736 |
| 3790 | III | 792 | 35 | 0 | 0 | 282 | 1 | M9+ | 17 – 93 | | 18109555 | PF6260 | R1b1 | | 3790 |

| | | M9+ | | | M9- | | | | | | Build 37 | | ISOGG | | |
|---|---|---|---|---|---|---|---|---|---|---|---|---|---|---|---|
| List | Stage | 0 | 1 | . | 0 | 1 | . | M9? | Clade(s) | Prob. | Position | Name | Subgroup Name | Amb. | List |
| 4170 | I | 792 | 35 | 0 | 0 | 283 | 0 | M9+ | 17 – 93 | | 19179606 | PF6261 | R1b1 | | 4170 |
| 4401 | I | 792 | 35 | 0 | 0 | 283 | 0 | M9+ | 17 – 93 | | 21159055 | PF6263 | R1b1 | | 4401 |
| 4560 | I | 792 | 35 | 0 | 0 | 283 | 0 | M9+ | 17 – 93 | | 21558298 | YSC0000224 | R1b1 | | 4560 |
| 5223 | III | 792 | 35 | 0 | 0 | 282 | 1 | M9+ | 17 – 93 | | 22889018 | YSC0000022 | R1b1 | | 5223 |
| 5630 | I | 792 | 35 | 0 | 0 | 283 | 0 | M9+ | 17 – 93 | | 23984056 | PF6271 | R1b1 | | 5630 |
| 5723 | I | 792 | 35 | 0 | 0 | 283 | 0 | M9+ | 17 – 93 | | 28733101 | PF6531 | R1b1a | | 5723 |
| 1019 | III | 792 | 35 | 0 | 0 | 282 | 1 | M9+ | 17 – 93 | | 7759944 | PF6424 | R1b1a1 | | 1019 |
| 2012 | I | 792 | 35 | 0 | 0 | 283 | 0 | M9+ | 17 – 93 | | 14116584 | PF6451 | R1b1a1 | | 2012 |
| 2559 | III | 35 | 791 | 1 | 283 | 0 | 0 | M9+ | 17 – 93 | | 15239181 | PF6458 | R1b1a1 | | 2559 |
| 2933 | I | 792 | 35 | 0 | 0 | 283 | 0 | M9+ | 17 – 93 | | 16183412 | PF6463 | R1b1a1 | | 2933 |
| 3022 | I | 792 | 35 | 0 | 0 | 283 | 0 | M9+ | 17 – 93 | | 16376495 | PF6464 | R1b1a1 | | 3022 |
| 3590 | I | 792 | 35 | 0 | 0 | 283 | 0 | M9+ | 17 – 93 | | 17732408 | PF6471 | R1b1a1 | | 3590 |
| 3600 | I | 792 | 35 | 0 | 0 | 283 | 0 | M9+ | 17 – 93 | | 17755905 | PF6472 | R1b1a1 | | 3600 |
| 3727 | I | 792 | 35 | 0 | 0 | 283 | 0 | M9+ | 17 – 93 | | 17986687 | YSC0000269 | R1b1a1 | | 3727 |
| 3851 | I | 792 | 35 | 0 | 0 | 283 | 0 | M9+ | 17 – 93 | | 18394634 | PF6483 | R1b1a1 | | 3851 |
| 3909 | I | 792 | 35 | 0 | 0 | 283 | 0 | M9+ | 17 – 93 | | 18617596 | PF6484 | R1b1a1 | | 3909 |
| 3924 | I | 792 | 35 | 0 | 0 | 283 | 0 | M9+ | 17 – 93 | | 18656508 | PF6398 | R1b1a1 | | 3924 |
| 4085 | I | 792 | 35 | 0 | 0 | 283 | 0 | M9+ | 17 – 93 | | 19020340 | PF6487 | R1b1a1 | | 4085 |
| 4460 | III | 788 | 35 | 4 | 0 | 283 | 0 | M9+ | 17 – 93 | | 21312064 | PF6498 | R1b1a1 | | 4460 |
| 4477 | I | 792 | 35 | 0 | 0 | 283 | 0 | M9+ | 17 – 93 | | 21358553 | PF6499 | R1b1a1 | | 4477 |
| 4744 | I | 792 | 35 | 0 | 0 | 283 | 0 | M9+ | 17 – 93 | | 21801722 | PF6506 | R1b1a1 | | 4744 |
| 5456 | I | 792 | 35 | 0 | 0 | 283 | 0 | M9+ | 17 – 93 | | 23403749 | PF6523 | R1b1a1 | | 5456 |
| 5472 | I | 792 | 35 | 0 | 0 | 283 | 0 | M9+ | 17 – 93 | | 23452965 | PF6524 | R1b1a1 | | 5472 |
| 5470 | IV | 827 | 0 | 0 | 283 | 0 | 0 | Mono | Mono | | 23444054 | M478 | R1b1a1a1 | | 5470 |
| 27 | I | 792 | 35 | 0 | 0 | 283 | 0 | M9+ | 17 – 93 | | 2668456 | S10 | R1b1a1b | | 27 |
| 285 | I | 792 | 35 | 0 | 0 | 283 | 0 | M9+ | 17 – 93 | | 4352151 | PF6409 | R1b1a1b | | 285 |
| 618 | I | 792 | 35 | 0 | 0 | 283 | 0 | M9+ | 17 – 93 | | 6912992 | PF6419 | R1b1a1b | | 618 |
| 727 | I | 792 | 35 | 0 | 0 | 283 | 0 | M9+ | 17 – 93 | | 7073423 | YSC0001293 | R1b1a1b | | 727 |
| 798 | I | 792 | 35 | 0 | 0 | 283 | 0 | M9+ | 17 – 93 | | 7220727 | YSC0000276 | R1b1a1b | | 798 |
| 1021 | I | 792 | 35 | 0 | 0 | 283 | 0 | M9+ | 17 – 93 | | 7762947 | PF6425 | R1b1a1b | | 1021 |
| 1024 | I | 792 | 35 | 0 | 0 | 283 | 0 | M9+ | 17 – 93 | | 7766712 | PF6426 | R1b1a1b | | 1024 |
| 1071 | III | 792 | 35 | 0 | 0 | 282 | 1 | M9+ | 17 – 93 | | 7863189 | PF6427 | R1b1a1b | | 1071 |
| 1118 | I | 792 | 35 | 0 | 0 | 283 | 0 | M9+ | 17 – 93 | | 7971505 | PF6429 | R1b1a1b | | 1118 |
| 1154 | I | 792 | 35 | 0 | 0 | 283 | 0 | M9+ | 17 – 93 | | 8070532 | PF6430 | R1b1a1b | | 1154 |
| 1191 | I | 792 | 35 | 0 | 0 | 283 | 0 | M9+ | 17 – 93 | | 8149348 | PF6431 | R1b1a1b | | 1191 |
| 1202 | I | 792 | 35 | 0 | 0 | 283 | 0 | M9+ | 17 – 93 | | 8194310 | PF6432 | R1b1a1b | | 1202 |
| 1222 | I | 792 | 35 | 0 | 0 | 283 | 0 | M9+ | 17 – 93 | | 8233186 | L483 | R1b1a1b | | 1222 |
| 1314 | I | 792 | 35 | 0 | 0 | 283 | 0 | M9+ | 17 – 93 | | 8411202 | PF6434 | R1b1a1b | | 1314 |
| 1526 | I | 792 | 35 | 0 | 0 | 283 | 0 | M9+ | 17 – 93 | | 8667179 | PF6435 | R1b1a1b | | 1526 |
| 1848 | III | 792 | 35 | 0 | 0 | 281 | 2 | M9+ | 17 – 93 | | 13511147 | PF6443 | R1b1a1b | | 1848 |
| 1933 | I | 792 | 35 | 0 | 0 | 283 | 0 | M9+ | 17 – 93 | | 13887941 | PF6252 | R1b1a1b | | 1933 |
| 1967 | I | 792 | 35 | 0 | 0 | 283 | 0 | M9+ | 17 – 93 | | 14042701 | PF6449 | R1b1a1b | | 1967 |
| 1994 | III | 790 | 35 | 2 | 0 | 282 | 1 | M9+ | 17 – 93 | | 14079811 | YSC0000258 | R1b1a1b | | 1994 |
| 2138 | I | 35 | 792 | 0 | 283 | 0 | 0 | M9+ | 17 – 93 | | 14416216 | PF6454 | R1b1a1b | | 2138 |
| 2400 | I | 35 | 792 | 0 | 283 | 0 | 0 | M9+ | 17 – 93 | | 14956117 | V2713 | R1b1a1b | | 2400 |
| 2486 | III | 35 | 791 | 1 | 281 | 0 | 2 | M9+ | 17 – 93 | | 15037433 | PF6457 | R1b1a1b | | 2486 |
| 3311 | I | 792 | 35 | 0 | 0 | 283 | 0 | M9+ | 17 – 93 | | 17013730 | YSC0000194 | R1b1a1b | | 3311 |
| 3395 | I | 792 | 35 | 0 | 1 | 282 | 0 | BOTH | | PI | 17281258 | Y1271 | R1b1a1b | W | 3395 |
| 3471 | I | 792 | 35 | 0 | 0 | 283 | 0 | M9+ | 17 – 93 | | 17461478 | PF6469 | R1b1a1b | | 3471 |
| 3535 | I | 792 | 35 | 0 | 0 | 283 | 0 | M9+ | 17 – 93 | | 17594966 | PF6470 | R1b1a1b | | 3535 |
| 3780 | I | 792 | 35 | 0 | 0 | 283 | 0 | M9+ | 17 – 93 | | 18095336 | PF6477 | R1b1a1b | | 3780 |
| 3792 | I | 792 | 35 | 0 | 0 | 283 | 0 | M9+ | 17 – 93 | | 18117193 | PF6478 | R1b1a1b | | 3792 |
| 3799 | I | 792 | 35 | 0 | 0 | 283 | 0 | M9+ | 17 – 93 | | 18137831 | PF6479 | R1b1a1b | | 3799 |
| 3808 | I | 792 | 35 | 0 | 0 | 283 | 0 | M9+ | 17 – 93 | | 18167403 | S13 | R1b1a1b | | 3808 |
| 3818 | I | 792 | 35 | 0 | 0 | 283 | 0 | M9+ | 17 – 93 | | 18180446 | PF6481 | R1b1a1b | | 3818 |
| 3840 | I | 792 | 35 | 0 | 0 | 283 | 0 | M9+ | 17 – 93 | | 18381735 | YSC0000203 | R1b1a1b | | 3840 |
| 3946 | I | 792 | 35 | 0 | 0 | 283 | 0 | M9+ | 17 – 93 | | 18719565 | S3 | R1b1a1b | | 3946 |
| 4015 | III | 791 | 35 | 1 | 0 | 282 | 1 | M9+ | 17 – 93 | | 18865298 | YSC0000018 | R1b1a1b | | 4015 |
| 4105 | I | 792 | 35 | 0 | 0 | 283 | 0 | M9+ | 17 – 93 | | 19054889 | YSC0000293 | R1b1a1b | | 4105 |
| 4169 | I | 792 | 35 | 0 | 0 | 283 | 0 | M9+ | 17 – 93 | | 19179540 | YSC0000294 | R1b1a1b | | 4169 |
| 4231 | I | 792 | 35 | 0 | 0 | 283 | 0 | M9+ | 17 – 93 | | 19291359 | V3993 | R1b1a1b | | 4231 |
| 4331 | I | 792 | 35 | 0 | 0 | 283 | 0 | M9+ | 17 – 93 | | 20811307 | PF6494 | R1b1a1b | | 4331 |
| 4345 | I | 792 | 35 | 0 | 0 | 283 | 0 | M9+ | 17 – 93 | | 20828795 | PF6495 | R1b1a1b | | 4345 |
| 4350 | I | 792 | 35 | 0 | 0 | 283 | 0 | M9+ | 17 – 93 | | 20838224 | YSC0000213 | R1b1a1b | | 4350 |
| 4416 | I | 792 | 35 | 0 | 0 | 283 | 0 | M9+ | 17 – 93 | | 21222868 | YSC0000219 | R1b1a1b | | 4416 |
| 4506 | I | 792 | 35 | 0 | 0 | 283 | 0 | M9+ | 17 – 93 | | 21410840 | PF6500 | R1b1a1b | | 4506 |
| 4558 | I | 792 | 35 | 0 | 0 | 283 | 0 | M9+ | 17 – 93 | | 21554468 | PF6265 | R1b1a1b | | 4558 |
| 4737 | I | 792 | 35 | 0 | 0 | 283 | 0 | M9+ | 17 – 93 | | 21784286 | YSC0000225 | R1b1a1b | | 4737 |
| 4914 | I | 792 | 35 | 0 | 0 | 283 | 0 | M9+ | 17 – 93 | | 21983827 | PF6507 | R1b1a1b | | 4914 |
| 4995 | I | 792 | 35 | 0 | 0 | 283 | 0 | M9+ | 17 – 93 | | 22190371 | PF6509 | R1b1a1b | | 4995 |
| 5137 | I | 792 | 35 | 0 | 0 | 283 | 0 | M9+ | 17 – 93 | | 22739367 | PF6517 | R1b1a1b | | 5137 |
| 5188 | I | 792 | 35 | 0 | 0 | 283 | 0 | M9+ | 17 – 93 | | 22796697 | CTS10834 | R1b1a1b | | 5188 |
| 5342 | I | 792 | 35 | 0 | 0 | 283 | 0 | M9+ | 17 – 93 | | 23124367 | PF6520 | R1b1a1b | | 5342 |
| 5380 | I | 792 | 35 | 0 | 0 | 283 | 0 | M9+ | 17 – 93 | | 23242935 | YSC0000238 | R1b1a1b | | 5380 |
| 5438 | I | 792 | 35 | 0 | 0 | 283 | 0 | M9+ | 17 – 93 | | 23379254 | PF6522 | R1b1a1b | | 5438 |
| 5658 | III | 792 | 33 | 2 | 0 | 283 | 0 | M9+ | 17 – 93 | MD | 24394612 | PF6527 | R1b1a1b | | 5658 |
| 5675 | III | 792 | 33 | 2 | 0 | 283 | 0 | M9+ | 17 – 93 | MD, 2H | 24444622 | YSC0000240 | R1b1a1b | | 5675 |
| 5728 | III | 775 | 35 | 17 | 0 | 280 | 3 | M9+ | 17 – 93 | | 28771116 | PF6532 | R1b1a1b | | 5728 |
| 520 | I | 791 | 36 | 0 | 0 | 283 | 0 | M9+ | 17 – 92 | | 6753511 | S141 | R1b1a1b1 | | 520 |
| 1413 | I | 782 | 45 | 0 | 0 | 283 | 0 | M9+ | 18 – 92 | | 8502236 | S167 | R1b1a1b1a | | 1413 |
| 1645 | I | 782 | 45 | 0 | 0 | 283 | 0 | M9+ | 18 – 92 | | 9084870 | YSC0000082 | R1b1a1b1a1 | | 1645 |
| 3835 | I | 782 | 45 | 0 | 0 | 283 | 0 | M9+ | 18 – 92 | | 18248698 | S128 | R1b1a1b1a1 | | 3835 |
| 4035 | I | 782 | 45 | 0 | 0 | 283 | 0 | M9+ | 18 – 92 | | 18907236 | S129 | R1b1a1b1a1 | | 4035 |
| 3167 | I | 782 | 45 | 0 | 0 | 283 | 0 | M9+ | 18 – 92 | | 16751825 | YSC0000191 | R1b1a1b1a1a | | 3167 |
| 3652 | I | 782 | 45 | 0 | 0 | 283 | 0 | M9+ | 18 – 92 | | 17844018 | L11 | R1b1a1b1a1a | | 3652 |
| 1577 | I | 604 | 223 | 0 | 283 | 0 | 0 | M9+ | 23 – 46 | | 8796078 | U106 | R1b1a1b1a1a1 | | 1577 |
| 2830 | I | 821 | 6 | 0 | 283 | 0 | 0 | M9+ | 25 | | 15779994 | Z8056 | R1b1a1b1a1a1a | | 2830 |
| 2426 | I | 811 | 16 | 0 | 283 | 0 | 0 | M9+ | 26 - 27 | | 14991735 | Z18 | R1b1a1b1a1a1b | | 2426 |
| 4933 | III | 817 | 7 | 3 | 283 | 0 | 0 | M9+ | 27 | | 22038215 | Z372 | R1b1a1b1a1a1b1a | | 4933 |
| 2831 | III | 770 | 56 | 1 | 283 | 0 | 0 | M9+ | 28 – 29 | | 15780341 | Z156 | R1b1a1b1a1a1c1 | | 2831 |
| 4098 | III | 788 | 25 | 14 | 283 | 0 | 0 | M9+ | 29 | MD | 19047132 | Z307 | R1b1a1b1a1a1c1a | | 4098 |
| 5080 | III | 768 | 9 | 50 | 272 | 0 | 11 | M9+ | 29 | MD | 22605163 | Z305 | R1b1a1b1a1a1c1a | | 5080 |
| 2301 | IV | 826 | 0 | 1 | 283 | 0 | 0 | Mono | Mono | | 14850465 | P89.2 | R1b1a1b1a1a1c1a2b1a1a | A | 2301 |
| 5333 | I | 809 | 18 | 0 | 283 | 0 | 0 | M9+ | 30 | | 23115473 | S1688 | R1b1a1b1a1a1c2a | | 5333 |
| 3215 | I | 809 | 18 | 0 | 283 | 0 | 0 | M9+ | 30 | | 16839499 | U198 | R1b1a1b1a1a1c2a1 | | 3215 |
| 5529 | I | 713 | 114 | 0 | 283 | 0 | 0 | M9+ | 31 – 45 | | 23612197 | S162 | R1b1a1b1a1a1c2b | | 5529 |

| List | Stage | M9+ | | | M9- | | | | | | Build 37 | | ISOGG | | |
|---|---|---|---|---|---|---|---|---|---|---|---|---|---|---|---|
| List | Stage | 0 | 1 | . | 0 | 1 | . | M9? | Clade(s) | Prob. | Position | Name | Subgroup Name | Amb. | List |
| 4988 | I | 813 | 14 | 0 | 283 | 0 | 0 | M9+ | 32 – 34 | | 22178569 | S170 | R1b1a1b1a1a1c2b1 | | 4988 |
| 187 | III | 825 | 1 | 1 | 283 | 0 | 0 | M9+ | 32 | | 2892150 | S171 | R1b1a1b1a1a1c2b1a | | 187 |
| 849 | IV | 726 | 0 | 101 | 243 | 0 | 40 | Mono | Mono | MD | 7400278 | S352 | R1b1a1b1a1a1c2b1a1 | | 849 |
| 1974 | II | 826 | 1 | 0 | 283 | 0 | 0 | M9+ | 32 | | 14053205 | L525 | R1b1a1b1a1a1c2b1a1a | | 1974 |
| 4116 | II | 826 | 1 | 0 | 283 | 0 | 0 | M9+ | 32 | | 19077409 | S172 | R1b1a1b1a1a1c2b1a1a | | 4116 |
| 2527 | II | 826 | 1 | 0 | 283 | 0 | 0 | M9+ | 32 | | 15116138 | S353 | R1b1a1b1a1a1c2b1a1a1 | | 2527 |
| 2799 | II | 826 | 1 | 0 | 283 | 0 | 0 | M9+ | 32 | | 15701859 | S502 | R1b1a1b1a1a1c2b1a1a1 | | 2799 |
| 5457 | IV | 825 | 0 | 2 | 283 | 0 | 0 | Mono | Mono | MD | 23405659 | CTS11988 | R1b1a1b1a1a1c2b1a1a1 | | 5457 |
| 159 | IV | 827 | 0 | 0 | 283 | 0 | 0 | Mono | Mono | | 2867288 | L477 | R1b1a1b1a1a1c2b1a1a1a | A | 159 |
| 2842 | IV | 827 | 0 | 0 | 283 | 0 | 0 | Mono | Mono | | 15800326 | L493 | R1b1a1b1a1a1c2b1a1a1a | | 2842 |
| 5229 | I | 814 | 13 | 0 | 283 | 0 | 0 | M9+ | 33 – 34 | | 22908970 | Z350 | R1b1a1b1a1a1c2b1b | | 5229 |
| 1339 | III | 825 | 1 | 1 | 283 | 0 | 0 | M9+ | 34 | | 8436030 | V1970.2 | R1b1a1b1a1a1c2b1b2a1 | A | 1339 |
| 531 | I | 733 | 94 | 0 | 283 | 0 | 0 | M9+ | 35 – 45 | | 6788390 | Z9 | R1b1a1b1a1a1c2b2 | | 531 |
| 3609 | I | 733 | 94 | 0 | 283 | 0 | 0 | M9+ | 35 – 45 | | 17765734 | Z348 | R1b1a1b1a1a1c2b2 | | 3609 |
| 4639 | I | 733 | 94 | 0 | 283 | 0 | 0 | M9+ | 35 – 45 | | 21700810 | Z28 | R1b1a1b1a1a1c2b2 | | 4639 |
| 5270 | I | 749 | 78 | 0 | 283 | 0 | 0 | M9+ | 35 – 43 | | 22971392 | Z30 | R1b1a1b1a1a1c2b2a | | 5270 |
| 1074 | I | 754 | 73 | 0 | 283 | 0 | 0 | M9+ | 37 – 43 | | 7883339 | Z2 | R1b1a1b1a1a1c2b2a1b | | 1074 |
| 3440 | I | 767 | 60 | 0 | 283 | 0 | 0 | M9+ | 38 – 43 | | 17391560 | Z7 | R1b1a1b1a1a1c2b2a1b1 | | 3440 |
| 5574 | III | 766 | 60 | 1 | 282 | 0 | 1 | M9+ | 38 – 43 | | 23805264 | Z31 | R1b1a1b1a1a1c2b2a1b1 | | 5574 |
| 1457 | I | 775 | 52 | 0 | 283 | 0 | 0 | M9+ | 39 – 42 | | 8568930 | Z22 | R1b1a1b1a1a1c2b2a1b1a | | 1457 |
| 2236 | I | 775 | 52 | 0 | 283 | 0 | 0 | M9+ | 39 – 42 | | 14746493 | Z24.2 | R1b1a1b1a1a1c2b2a1b1a | A | 2236 |
| 4222 | I | 775 | 52 | 0 | 283 | 0 | 0 | M9+ | 39 – 42 | | 19276174 | Z26 | R1b1a1b1a1a1c2b2a1b1a | | 4222 |
| 4996 | I | 775 | 52 | 0 | 283 | 0 | 0 | M9+ | 39 – 42 | | 22191144 | Z8 | R1b1a1b1a1a1c2b2a1b1a | | 4996 |
| 2937 | I | 812 | 15 | 0 | 283 | 0 | 0 | M9+ | 41 – 42 | | 16190880 | Z11 | R1b1a1b1a1a1c2b2a1b1a1 | | 2937 |
| 4532 | III | 815 | 11 | 1 | 283 | 0 | 0 | M9+ | 42 | | 21481454 | Z12 | R1b1a1b1a1a1c2b2a1b1a1a | | 4532 |
| 5201 | I | 821 | 6 | 0 | 283 | 0 | 0 | M9+ | 43 | | 22823352 | S3595 | R1b1a1b1a1a1c2b2a1b1b1 | | 5201 |
| 1122 | IV | 827 | 0 | 0 | 283 | 0 | 0 | Mono | Mono | | 7989319 | S27458 | R1b1a1b1a1a1c2b2a1b1b1a4b | | 1122 |
| 2030 | IV | 827 | 0 | 0 | 283 | 0 | 0 | Mono | Mono | | 14159864 | S347.2 | R1b1a1b1a1a1c2b2a1b3 | A | 2030 |
| 3563 | I | 811 | 16 | 0 | 283 | 0 | 0 | M9+ | 44 – 45 | | 17663668 | Z347 | R1b1a1b1a1a1c2b2b | | 3563 |
| 4083 | III | 809 | 16 | 2 | 283 | 0 | 0 | M9+ | 44 – 45 | | 19007975 | Z334 | R1b1a1b1a1a1c2b2b | | 4083 |
| 3110 | I | 815 | 12 | 0 | 283 | 0 | 0 | M9+ | 45 | | 16585994 | Z330 | R1b1a1b1a1a1c2b2b1 | | 3110 |
| 1034 | I | 815 | 12 | 0 | 283 | 0 | 0 | M9+ | 45 | | 7792995 | Z329 | R1b1a1b1a1a1c2b2b1a | | 1034 |
| 4929 | I | 815 | 12 | 0 | 283 | 0 | 0 | M9+ | 45 | | 22023565 | Z337 | R1b1a1b1a1a1c2b2b1a | | 4929 |
| 5621 | III | 815 | 12 | 0 | 282 | 0 | 1 | M9+ | 45 | | 23973367 | Z326 | R1b1a1b1a1a1c2b2b1a | | 5621 |
| 4252 | III | 808 | 12 | 7 | 282 | 0 | 1 | M9+ | 45 | | 19349905 | Z335 | R1b1a1b1a1a1c2b2b1a~ | | 4252 |
| 4778 | II | 826 | 1 | 0 | 283 | 0 | 0 | M9+ | 46 | | 21867718 | M323.2 | R1b1a1b1a1a1c3 | A | 4778 |
| 705 | IV | 826 | 0 | 1 | 283 | 0 | 0 | Mono | Mono | | 6997451 | Y3444 | R1b1a1b1a1a1d | | 705 |
| 3115 | IV | 827 | 0 | 0 | 283 | 0 | 0 | Mono | Mono | | 16596916 | L86.2 | R1b1a1b1a1a2~ | A,W | 3115 |
| 3697 | I | 759 | 68 | 0 | 283 | 0 | 0 | M9+ | 47 – 58 | | 17922066 | Z195 | R1b1a1b1a1a2a1 | | 3697 |
| 2685 | III | 788 | 22 | 17 | 282 | 0 | 1 | M9+ | 50 – 57 | | 15508140 | Z209 | R1b1a1b1a1a2a1a1 | | 2685 |
| 2987 | I | 805 | 22 | 0 | 283 | 0 | 0 | M9+ | 50 – 57 | | 16310705 | Z220 | R1b1a1b1a1a2a1a1 | | 2987 |
| 899 | I | 805 | 22 | 0 | 283 | 0 | 0 | M9+ | 50 – 57 | | 7571621 | Z268 | R1b1a1b1a1a2a1a1~ | | 899 |
| 3809 | I | 821 | 6 | 0 | 283 | 0 | 0 | M9+ | 54 – 56 | | 18167479 | Z278 | R1b1a1b1a1a2a1a1a1 | | 3809 |
| 1356 | III | 812 | 5 | 10 | 281 | 0 | 2 | M9+ | 53 – 56 | MD | 8456571 | Z211 | R1b1a1b1a1a2a1a1a1~ | | 1356 |
| 3931 | I | 821 | 6 | 0 | 283 | 0 | 0 | M9+ | 54 – 56 | | 18670504 | Z212 | R1b1a1b1a1a2a1a1a1~ | | 3931 |
| 4756 | II | 826 | 1 | 0 | 283 | 0 | 0 | M9+ | 56 | | 21827794 | Z214 | R1b1a1b1a1a2a1a1a1a | | 4756 |
| 4640 | IV | 827 | 0 | 0 | 283 | 0 | 0 | Mono | Mono | | 21706360 | M153 | R1b1a1b1a1a2a1a1a1a1 | | 4640 |
| 2602 | II | 826 | 1 | 0 | 283 | 0 | 0 | M9+ | 57 | | 15372323 | Z2355 | R1b1a1b1a1a2a1a1a2 | | 2602 |
| 15 | I | 800 | 27 | 0 | 283 | 0 | 0 | M9+ | 58 | | 2658271 | SRY2627 | R1b1a1b1a1a2a1b1a1 | | 15 |
| 2833 | III | 800 | 25 | 2 | 283 | 0 | 0 | M9+ | 58 | MD | 15781305 | CTS4716 | R1b1a1b1a1a2a1b1a1~ | | 2833 |
| 2814 | IV | 827 | 0 | 0 | 279 | 0 | 4 | Mono | Mono | | 15726862 | PF742.2 | R1b1a1b1a1a2a1b1a1a~ | | 2814 |
| 1487 | II | 824 | 3 | 0 | 283 | 0 | 0 | M9+ | 59 | | 8603396 | Z225 | R1b1a1b1a1a2a5 | | 1487 |
| 1377 | II | 824 | 3 | 0 | 283 | 0 | 0 | M9+ | 59 | | 8469661 | Z229 | R1b1a1b1a1a2a5~ | | 1377 |
| 2588 | I | 780 | 47 | 0 | 283 | 0 | 0 | M9+ | 60 – 63 | | 15333149 | U152 | R1b1a1b1a1a2b | A | 2588 |
| 412 | I | 37 | 790 | 0 | 0 | 283 | 0 | M9+ | 60 – 63 | | 5755550 | S139 | R1b1a1b1a1a2b1 | | 412 |
| 1731 | I | 17 | 810 | 0 | 0 | 283 | 0 | M9+ | 62 – 63 | | 9810794 | Z367 | R1b1a1b1a1a2b1a | | 1731 |
| 3155 | III | 17 | 809 | 1 | 0 | 283 | 0 | M9+ | 62 – 63 | | 16720013 | Z384 | R1b1a1b1a1a2b1a~ | | 3155 |
| 2064 | I | 10 | 817 | 0 | 0 | 283 | 0 | M9+ | 63 | | 14231292 | S144 | R1b1a1b1a1a2b1a1 | | 2064 |
| 5521 | IV | 827 | 0 | 0 | 283 | 0 | 0 | Mono | Mono | | 23574525 | Z291 | R1b1a1b1a1a2b1a1~ | W | 5521 |
| 5662 | III | 10 | 815 | 2 | 0 | 283 | 0 | M9+ | 63 | | 24405862 | Z383 | R1b1a1b1a1a2b1a1~ | | 5662 |
| 5666 | III | 474 | 225 | 128 | 282 | 0 | 1 | M9+ | 64 – 89 | MD | 24411932 | Z260 | R1b1a1b1a1a2c | | 5666 |
| 2777 | I | 476 | 351 | 0 | 283 | 0 | 0 | M9+ | 64 – 89 | | 15654428 | S145 | R1b1a1b1a1a2c1 | | 2777 |
| 3674 | I | 481 | 346 | 0 | 283 | 0 | 0 | M9+ | 66 – 89 | | 17885577 | Z2542 | R1b1a1b1a1a2c1a | | 3674 |
| 5131 | III | 768 | 54 | 5 | 279 | 0 | 4 | M9+ | 70 – 74 | | 22735599 | S474 | R1b1a1b1a1a2c1a1a | | 5131 |
| 1066 | I | 789 | 38 | 0 | 283 | 0 | 0 | M9+ | 73 – 74 | | 7855252 | Z2961 | R1b1a1b1a1a2c1a1a1a1a | | 1066 |
| 2349 | I | 793 | 34 | 0 | 283 | 0 | 0 | M9+ | 74 | | 14902414 | USP9Y+3636 | R1b1a1b1a1a2c1a1a1a1a1 | | 2349 |
| 1971 | III | 809 | 4 | 14 | 282 | 0 | 1 | M9+ | 75 | | 14048611 | S3666 | R1b1a1b1a1a2c1a1b | | 1971 |
| 3305 | I | 820 | 7 | 0 | 283 | 0 | 0 | M9+ | 76 | | 16992602 | S524 | R1b1a1b1a1a2c1a1d | | 3305 |
| 4134 | IV | 820 | 0 | 7 | 283 | 0 | 0 | Mono | Mono | | 19118561 | Z17470.2 | R1b1a1b1a1a2c1a1d3c | A | 4134 |
| 915 | IV | 827 | 0 | 0 | 283 | 0 | 0 | Mono | Mono | | 7596723 | A874 | R1b1a1b1a1a2c1a1d4 | | 915 |
| 3349 | I | 820 | 7 | 0 | 283 | 0 | 0 | M9+ | 77 | | 17138487 | CTS6838 | R1b1a1b1a1a2c1a1f1 | | 3349 |
| 3389 | III | 820 | 7 | 0 | 282 | 0 | 1 | M9+ | 77 | | 17246645 | S735 | R1b1a1b1a1a2c1a1f1 | | 3389 |
| 4014 | I | 819 | 8 | 0 | 270 | 13 | 0 | BOTH | | PI | 18861458 | S738^^ | R1b1a1b1a1a2c1a1f1 | W | 4014 |
| 5375 | I | 820 | 7 | 0 | 283 | 0 | 0 | M9+ | 77 | | 23237322 | S749 | R1b1a1b1a1a2c1a1f1 | | 5375 |
| 45 | IV | 827 | 0 | 0 | 281 | 0 | 2 | Mono | Mono | | 2710015 | Z17611 | R1b1a1b1a1a2c1a1f1a1b | | 45 |
| 2351 | IV | 827 | 0 | 0 | 283 | 0 | 0 | Mono | Mono | | 14902456 | L96 | R1b1a1b1a1a2c1a3a1 | | 2351 |
| 2781 | III | 791 | 34 | 2 | 283 | 0 | 0 | M9+ | | PI | 15660495 | S1136 | R1b1a1b1a1a2c1a3a2 | | 2781 |
| 2801 | I | 795 | 32 | 0 | 283 | 0 | 0 | M9+ | 79 | | 15702775 | Z255 | R1b1a1b1a1a2c1a4a | | 2801 |
| 788 | I | 757 | 70 | 0 | 283 | 0 | 0 | M9+ | 80 – 84 | | 7193034 | Z253 | R1b1a1b1a1a2c1a4b | | 788 |
| 4091 | I | 776 | 51 | 0 | 283 | 0 | 0 | M9+ | 82 – 84 | | 19033817 | Z2534 | R1b1a1b1a1a2c1a4b2 | | 4091 |
| 2591 | I | 801 | 26 | 0 | 283 | 0 | 0 | M9+ | 83 – 84 | | 15339857 | Z2184 | R1b1a1b1a1a2c1a4b2c | | 2591 |
| 4171 | III | 800 | 26 | 1 | 283 | 0 | 0 | M9+ | 83 – 84 | | 19180465 | Z2185 | R1b1a1b1a1a2c1a4b2c | | 4171 |
| 814 | I | 803 | 24 | 0 | 283 | 0 | 0 | M9+ | 84 | | 7265766 | L1066.2 | R1b1a1b1a1a2c1a4b2c1a | A | 814 |
| 1441 | IV | 826 | 0 | 1 | 283 | 0 | 0 | Mono | Mono | | 8548391 | Z2189 | R1b1a1b1a1a2c1a4b2d | | 1441 |
| 3327 | IV | 796 | 0 | 31 | 268 | 0 | 15 | Mono | Mono | | 17054132 | PF825.2 | R1b1a1b1a1a2c1a4b3a1 | A | 3327 |
| 5626 | III | 826 | 1 | 0 | 280 | 2 | 1 | BOTH | | PI | 23977118 | F4036.2 | R1b1a1b1a1a2c1a4b8~ | W | 5626 |
| 1765 | IV | 795 | 0 | 32 | 283 | 0 | 0 | Mono | Mono | MD | 9878915 | S192 | R1b1a1b1a1a2c1a5 | | 1765 |
| 2794 | II | 824 | 3 | 0 | 283 | 0 | 0 | M9+ | 85 | | 15683936 | S3058 | R1b1a1b1a1a2c1a5a2a | | 2794 |
| 130 | III | 811 | 16 | 0 | 282 | 0 | 1 | M9+ | 86 – 88 | | 2826066 | S253 | R1b1a1b1a1a2c1a5b1a1 | | 130 |
| 2137 | III | 814 | 12 | 1 | 283 | 0 | 0 | M9+ | 87 – 88 | | 14407732 | Z248 | R1b1a1b1a1a2c1a5b1a1a | | 2137 |
| 3850 | III | 815 | 11 | 1 | 283 | 0 | 0 | M9+ | 87 – 88 | MD | 18393814 | S191 | R1b1a1b1a1a2c1a5b1a1a | | 3850 |
| 2517 | I | 822 | 5 | 0 | 283 | 0 | 0 | M9+ | 88 | | 15095379 | S3787 | R1b1a1b1a1a2c1a5b1a1a2 | | 2517 |
| 3455 | IV | 827 | 0 | 0 | 283 | 0 | 0 | Mono | Mono | | 17421349 | V75.2 | R1b1a1b1a1a2c1a5b1a1a2b1a1 | A | 3455 |
| 4540 | IV | 810 | 0 | 17 | 280 | 0 | 3 | Mono | Mono | MD | 21497008 | Z3017 | R1b1a1b1a1a2c1a5d | | 4540 |
| 1536 | II | 826 | 1 | 0 | 283 | 0 | 0 | M9+ | 89 | | 8684816 | Z3001 | R1b1a1b1a1a2c1a5d3 | | 1536 |
| 4379 | IV | 827 | 0 | 0 | 283 | 0 | 0 | Mono | Mono | | 21101380 | Z13583.2 | R1b1a1b1a1a2c1a6b~ | A | 4379 |

| | | M9+ | | | M9- | | | | | | Build 37 | | ISOGG | | |
|---|---|---|---|---|---|---|---|---|---|---|---|---|---|---|---|
| List | Stage | 0 | 1 | . | 0 | 1 | . | M9? | Clade(s) | Prob. | Position | Name | Subgroup Name | Amb. | List |
| 4430 | IV | 827 | 0 | 0 | 283 | 0 | 0 | Mono | Mono | | 21253443 | S182 | R1b1a1b1a1a2d | | 4430 |
| 2095 | III | 816 | 10 | 1 | 282 | 0 | 1 | M9+ | 90 – 92 | | 14301499 | S232 | R1b1a1b1a1a2e | | 2095 |
| 3488 | I | 820 | 7 | 0 | 283 | 0 | 0 | M9+ | 91 | | 17508792 | S4298 | R1b1a1b1a1a2e1 | | 3488 |
| 3422 | IV | 823 | 0 | 4 | 283 | 0 | 0 | Mono | Mono | | 17340220 | FGC37100 | R1b1a1b1a1a4 | | 3422 |
| 1490 | II | 826 | 1 | 0 | 283 | 0 | 0 | M9+ | 93 | | 8610246 | V2347 | R1b1a1b2a | | 1490 |
| 4667 | IV | 827 | 0 | 0 | 283 | 0 | 0 | Mono | Mono | | 21733162 | M18 | R1b1b1 | | 4667 |
| 539 | IV | 826 | 0 | 1 | 282 | 0 | 1 | Mono | Mono | | 6812012 | V35 | R1b1b2a1 | | 539 |
| 2462 | IV | 821 | 0 | 6 | 283 | 0 | 0 | Mono | Mono | | 15026395 | M335 | R1b2a | | 2462 |
| 1327 | IV | 826 | 0 | 1 | 283 | 0 | 0 | Mono | Mono | | 8426362 | PF6108 | R2 | | 1327 |
| 4347 | IV | 827 | 0 | 0 | 283 | 0 | 0 | Mono | Mono | | 20834667 | PF6107 | R2 | | 4347 |
| 514 | IV | 827 | 0 | 0 | 283 | 0 | 0 | Mono | Mono | | 6740704 | Page4 | R2a | | 514 |
| 1275 | IV | 826 | 0 | 1 | 283 | 0 | 0 | Mono | Mono | | 8335203 | P249 | R2a | | 1275 |
| 4715 | IV | 826 | 0 | 1 | 283 | 0 | 0 | Mono | Mono | | 21764501 | M124 | R2a | | 4715 |
| 3738 | IV | 826 | 0 | 1 | 283 | 0 | 0 | Mono | Mono | | 18028669 | PF6109 | R2a2 | | 3738 |
| 4733 | IV | 827 | 0 | 0 | 283 | 0 | 0 | Mono | Mono | | 21778964 | L723 | R2a2b1b2b1 | | 4733 |
| 60 | IV | 827 | 0 | 0 | 283 | 0 | 0 | Mono | Mono | | 2722677 | FGC50198 | R2b | | 60 |
| 1305 | IV | 826 | 0 | 1 | 283 | 0 | 0 | Mono | Mono | | 8391108 | P405 | S1a~ | A | 1305 |
| 1950 | IV | 826 | 0 | 1 | 283 | 0 | 0 | Mono | Mono | | 14001024 | P202 | S1a1b | | 1950 |
| 2672 | IV | 781 | 0 | 46 | 264 | 0 | 19 | Mono | Mono | | 15481372 | M230 | S1a1b | | 2672 |
| 2161 | IV | 827 | 0 | 0 | 283 | 0 | 0 | Mono | Mono | | 14487066 | P57 | S1a1b1a | | 2161 |
| 22 | IV | 827 | 0 | 0 | 283 | 0 | 0 | Mono | Mono | | 2661836 | P307 | S1a2a | | 22 |
| 511 | IV | 827 | 0 | 0 | 283 | 0 | 0 | Mono | Mono | | 6740377 | P79 | S1a2a | A | 511 |
| 1420 | IV | 827 | 0 | 0 | 283 | 0 | 0 | Mono | Mono | | 8510540 | Z780.2 | S1a2a | A | 1420 |
| 97 | II | 826 | 1 | 0 | 283 | 0 | 0 | M9+ | 94 | | 2765271 | CTS150 | T | | 97 |
| 528 | II | 826 | 1 | 0 | 283 | 0 | 0 | M9+ | 94 | | 6781398 | PF5596 | T | | 528 |
| 532 | II | 826 | 1 | 0 | 283 | 0 | 0 | M9+ | 94 | | 6794129 | PF5597 | T | | 532 |
| 566 | III | 825 | 1 | 1 | 283 | 0 | 0 | M9+ | 94 | | 6854139 | CTS573 | T | | 566 |
| 2340 | II | 826 | 1 | 0 | 283 | 0 | 0 | M9+ | 94 | | 14898163 | USP9Y+3178 | T | A | 2340 |
| 2666 | III | 826 | 1 | 0 | 282 | 0 | 1 | M9+ | 94 | | 15472563 | Page129 | T | | 2666 |
| 5135 | II | 826 | 1 | 0 | 283 | 0 | 0 | M9+ | 94 | | 22738775 | PF5667 | T | | 5135 |
| 2179 | II | 826 | 1 | 0 | 283 | 0 | 0 | M9+ | 94 | | 14529630 | L490 | T1 | | 2179 |
| 3113 | II | 826 | 1 | 0 | 283 | 0 | 0 | M9+ | 94 | | 16595426 | CTS5987 | T1 | | 3113 |
| 2299 | III | 821 | 1 | 5 | 282 | 0 | 1 | M9+ | 94 | | 14848650 | Page78 | T1a | | 2299 |
| 2953 | II | 826 | 1 | 0 | 283 | 0 | 0 | M9+ | 94 | | 16212441 | PF7472.2 | T1a | A | 2953 |
| 2958 | II | 826 | 1 | 0 | 283 | 0 | 0 | M9+ | 94 | | 16224726 | CTS5364 | T1a | | 2958 |
| 4832 | II | 826 | 1 | 0 | 283 | 0 | 0 | M9+ | 94 | | 21893881 | PF5662 | T1a | | 4832 |
| 529 | II | 826 | 1 | 0 | 283 | 0 | 0 | M9+ | 94 | | 6782868 | CTS484 | T1a1 | | 529 |
| 549 | II | 826 | 1 | 0 | 283 | 0 | 0 | M9+ | 94 | | 6837868 | CTS550 | T1a1 | | 549 |
| 2257 | III | 825 | 1 | 1 | 283 | 0 | 0 | M9+ | 94 | | 14784140 | CTS3271 | T1a1 | | 2257 |
| 3010 | II | 826 | 1 | 0 | 283 | 0 | 0 | M9+ | 94 | | 16350661 | PF5627 | T1a1 | | 3010 |
| 2128 | II | 826 | 1 | 0 | 283 | 0 | 0 | M9+ | 94 | | 14389760 | CTS2611 | T1a1a | | 2128 |
| 2611 | II | 826 | 1 | 0 | 283 | 0 | 0 | M9+ | 94 | | 15385073 | PF5620 | T1a1a | | 2611 |
| 597 | III | 823 | 1 | 3 | 283 | 0 | 0 | M9+ | 94 | | 6890722 | Z709 | T1a1a1b2b | | 597 |
| 1263 | II | 826 | 1 | 0 | 283 | 0 | 0 | M9+ | 94 | | 8304456 | Z710 | T1a1a1b2b2 | | 1263 |
| 2115 | II | 826 | 1 | 0 | 283 | 0 | 0 | M9+ | 94 | | 14343435 | Z713 | T1a1a1b2b2 | | 2115 |
| 3511 | II | 826 | 1 | 0 | 283 | 0 | 0 | M9+ | 94 | | 17553410 | Z714 | T1a1a1b2b2 | | 3511 |
| 3759 | II | 826 | 1 | 0 | 283 | 0 | 0 | M9+ | 94 | | 18061004 | CTS8512 | T1a1a1b2b2 | | 3759 |
| 628 | III | 819 | 1 | 7 | 280 | 0 | 3 | M9+ | 94 | 1H | 6931957 | L906 | T1a1a1b2b2b | | 628 |
| 2177 | II | 826 | 1 | 0 | 283 | 0 | 0 | M9+ | 94 | | 14529342 | CTS2860 | T1a1a1b2b2b1a | | 2177 |
| 3433 | II | 826 | 1 | 0 | 283 | 0 | 0 | M9+ | 94 | | 17364892 | L907 | T1a1a1b2b2b1a | | 3433 |
| 5447 | II | 826 | 1 | 0 | 283 | 0 | 0 | M9+ | 94 | | 23390048 | CTS11968 | T1a1a1b2b2b1a | | 5447 |
| 3289 | II | 826 | 1 | 0 | 283 | 0 | 0 | M9+ | 94 | | 16949553 | CTS6507 | T1a1a1b2b2b1a1a | | 3289 |
| 2051 | IV | 827 | 0 | 0 | 283 | 0 | 0 | Mono | Mono | | 14201429 | CTS2157 | T1a2 | | 2051 |
| 4261 | IV | 827 | 0 | 0 | 283 | 0 | 0 | Mono | Mono | | 19372808 | L131 | T1a2 | | 4261 |
| 3136 | IV | 827 | 0 | 0 | 283 | 0 | 0 | Mono | Mono | | 16660785 | L446 | T1a2b | | 3136 |
| 4885 | IV | 825 | 0 | 2 | 282 | 0 | 1 | Mono | Mono | | 21926143 | L1255 | T1a3b1 | | 4885 |
| 1 | IV | 826 | 0 | 1 | 282 | 0 | 1 | Mono | Mono | | 2654333 | | | | 1 |
| 3 | IV | 827 | 0 | 0 | 283 | 0 | 0 | Mono | Mono | | 2655265 | | | | 3 |
| 4 | IV | 825 | 0 | 2 | 283 | 0 | 0 | Mono | Mono | | 2655328 | | | | 4 |
| 5 | IV | 826 | 0 | 1 | 283 | 0 | 0 | Mono | Mono | | 2655362 | | | | 5 |
| 6 | IV | 825 | 0 | 2 | 283 | 0 | 0 | Mono | Mono | | 2655371 | | | | 6 |
| 7 | IV | 827 | 0 | 0 | 283 | 0 | 0 | Mono | Mono | | 2655375 | | | | 7 |
| 8 | IV | 0 | 0 | 827 | 0 | 0 | 283 | Mono | Mono | | 2655436 | | | | 8 |
| 9 | IV | 827 | 0 | 0 | 283 | 0 | 0 | Mono | Mono | | 2655467 | | | | 9 |
| 10 | IV | 827 | 0 | 0 | 283 | 0 | 0 | Mono | Mono | | 2655471 | | | | 10 |
| 11 | IV | 827 | 0 | 0 | 283 | 0 | 0 | Mono | Mono | | 2655592 | | | | 11 |
| 12 | IV | 827 | 0 | 0 | 283 | 0 | 0 | Mono | Mono | | 2655633 | | | | 12 |
| 20 | IV | 827 | 0 | 0 | 283 | 0 | 0 | Mono | Mono | | 2661306 | | | | 20 |
| 21 | IV | 0 | 827 | 0 | 0 | 283 | 0 | Mono | Mono | | 2661694 | | | | 21 |
| 24 | IV | 827 | 0 | 0 | 283 | 0 | 0 | Mono | Mono | | 2663685 | | | | 24 |
| 32 | II | 826 | 1 | 0 | 283 | 0 | 0 | M9+ | NU | | 2683014 | | | | 32 |
| 36 | IV | 827 | 0 | 0 | 283 | 0 | 0 | Mono | Mono | | 2693993 | | | | 36 |
| 38 | IV | 826 | 0 | 1 | 282 | 0 | 1 | Mono | Mono | | 2703038 | | | | 38 |
| 39 | IV | 818 | 0 | 9 | 278 | 0 | 5 | Mono | Mono | | 2703239 | | | | 39 |
| 40 | IV | 827 | 0 | 0 | 283 | 0 | 0 | Mono | Mono | | 2705854 | | | | 40 |
| 42 | IV | 827 | 0 | 0 | 283 | 0 | 0 | Mono | Mono | | 2709555 | | | | 42 |
| 43 | IV | 827 | 0 | 0 | 283 | 0 | 0 | Mono | Mono | | 2710000 | | | | 43 |
| 51 | IV | 827 | 0 | 0 | 283 | 0 | 0 | Mono | Mono | | 2712237 | | | | 51 |
| 54 | IV | 827 | 0 | 0 | 283 | 0 | 0 | Mono | Mono | | 2713753 | | | | 54 |
| 58 | IV | 827 | 0 | 0 | 283 | 0 | 0 | Mono | Mono | | 2722413 | | | | 58 |
| 61 | IV | 827 | 0 | 0 | 283 | 0 | 0 | Mono | Mono | | 2722814 | | | | 61 |
| 64 | IV | 818 | 0 | 9 | 281 | 0 | 2 | Mono | Mono | | 2724189 | | | | 64 |
| 69 | IV | 827 | 0 | 0 | 283 | 0 | 0 | Mono | Mono | | 2727858 | | | | 69 |
| 73 | IV | 827 | 0 | 0 | 283 | 0 | 0 | Mono | Mono | | 2733235 | | | | 73 |
| 75 | II | 826 | 1 | 0 | 283 | 0 | 0 | M9+ | 1 | | 2734250 | | | | 75 |
| 76 | IV | 827 | 0 | 0 | 283 | 0 | 0 | Mono | Mono | | 2734776 | | | | 76 |
| 78 | IV | 827 | 0 | 0 | 283 | 0 | 0 | Mono | Mono | | 2734919 | | | | 78 |
| 82 | I | 827 | 0 | 0 | 246 | 37 | 0 | M9- | 1 – 10 | | 2743754 | | | | 82 |
| 84 | IV | 826 | 0 | 1 | 283 | 0 | 0 | Mono | Mono | | 2749045 | | | | 84 |
| 85 | IV | 827 | 0 | 0 | 283 | 0 | 0 | Mono | Mono | | 2749046 | | | | 85 |
| 92 | IV | 827 | 0 | 0 | 283 | 0 | 0 | Mono | Mono | | 2756980 | | | | 92 |
| 95 | IV | 825 | 0 | 2 | 283 | 0 | 0 | Mono | Mono | | 2762492 | | | | 95 |
| 98 | IV | 826 | 0 | 1 | 282 | 0 | 1 | Mono | Mono | | 2766172 | | | | 98 |
| 99 | IV | 827 | 0 | 0 | 283 | 0 | 0 | Mono | Mono | | 2768216 | | | | 99 |
| 109 | IV | 825 | 0 | 2 | 281 | 0 | 2 | Mono | Mono | | 2802847 | | | | 109 |
| 110 | IV | 814 | 0 | 13 | 278 | 0 | 5 | Mono | Mono | | 2803135 | | | | 110 |

| | | | M9+ | | | M9- | | | | | Build 37 | | ISOGG | | |
|---|---|---|---|---|---|---|---|---|---|---|---|---|---|---|---|
| List | Stage | 0 | 1 | . | 0 | 1 | . | M9? | Clade(s) | Prob. | Position | Name | Subgroup Name | Amb. | List |
| 117 | IV | 826 | 0 | 1 | 282 | 0 | 1 | Mono | Mono | | 2808518 | | | | 117 |
| 119 | IV | 827 | 0 | 0 | 283 | 0 | 0 | Mono | Mono | | 2810628 | | | | 119 |
| 124 | IV | 827 | 0 | 0 | 283 | 0 | 0 | Mono | Mono | | 2815425 | | | | 124 |
| 127 | IV | 827 | 0 | 0 | 283 | 0 | 0 | Mono | Mono | | 2818883 | | | | 127 |
| 129 | IV | 827 | 0 | 0 | 283 | 0 | 0 | Mono | Mono | | 2825845 | | | | 129 |
| 136 | III | 817 | 3 | 7 | 280 | 0 | 3 | M9+ | 67 | | 2829379 | | | | 136 |
| 137 | IV | 827 | 0 | 0 | 283 | 0 | 0 | Mono | Mono | | 2829550 | | | | 137 |
| 138 | IV | 771 | 0 | 56 | 266 | 0 | 17 | Mono | Mono | | 2829674 | | | | 138 |
| 139 | IV | 827 | 0 | 0 | 283 | 0 | 0 | Mono | Mono | | 2830361 | | | | 139 |
| 146 | I | 791 | 36 | 0 | 0 | 283 | 0 | M9+ | 17 – 92 | | 2842212 | | | | 146 |
| 149 | IV | 827 | 0 | 0 | 283 | 0 | 0 | Mono | Mono | | 2844882 | | | | 149 |
| 150 | IV | 823 | 0 | 4 | 281 | 0 | 2 | Mono | Mono | | 2845849 | | | | 150 |
| 153 | IV | 827 | 0 | 0 | 283 | 0 | 0 | Mono | Mono | | 2846959 | | | | 153 |
| 154 | IV | 827 | 0 | 0 | 283 | 0 | 0 | Mono | Mono | | 2847810 | | | | 154 |
| 156 | IV | 827 | 0 | 0 | 283 | 0 | 0 | Mono | Mono | | 2854520 | | | | 156 |
| 167 | I | 827 | 0 | 0 | 247 | 36 | 0 | M9- | 2 – 10 | | 2874542 | | | | 167 |
| 168 | III | 826 | 0 | 1 | 247 | 36 | 0 | M9- | 2 – 10 | | 2874544 | | | | 168 |
| 171 | IV | 827 | 0 | 0 | 283 | 0 | 0 | Mono | Mono | | 2877478 | | | | 171 |
| 191 | I | 827 | 0 | 0 | 0 | 283 | 0 | M9+/- | M9+/- | | 2902839 | | | | 191 |
| 195 | IV | 827 | 0 | 0 | 283 | 0 | 0 | Mono | Mono | | 2943636 | | | | 195 |
| 196 | IV | 824 | 0 | 3 | 283 | 0 | 0 | Mono | Mono | | 2948252 | | | | 196 |
| 197 | IV | 827 | 0 | 0 | 282 | 0 | 1 | Mono | Mono | | 2951972 | | | | 197 |
| 198 | IV | 807 | 0 | 20 | 277 | 0 | 6 | Mono | Mono | | 2957180 | | | | 198 |
| 199 | IV | 826 | 0 | 1 | 283 | 0 | 0 | Mono | Mono | | 2967867 | | | | 199 |
| 202 | IV | 820 | 0 | 7 | 276 | 0 | 7 | Mono | Mono | | 2994033 | | | | 202 |
| 203 | II | 826 | 1 | 0 | 283 | 0 | 0 | M9+ | NU | | 3017407 | | | | 203 |
| 204 | IV | 824 | 0 | 3 | 283 | 0 | 0 | Mono | Mono | | 3026733 | | | | 204 |
| 205 | IV | 827 | 0 | 0 | 283 | 0 | 0 | Mono | Mono | | 3029098 | | | | 205 |
| 206 | IV | 820 | 0 | 7 | 281 | 0 | 2 | Mono | Mono | | 3040556 | | | | 206 |
| 207 | IV | 827 | 0 | 0 | 282 | 0 | 1 | Mono | Mono | | 3059025 | | | | 207 |
| 208 | I | 827 | 0 | 0 | 59 | 224 | 0 | M9- | 22 – 66 | | 3074753 | | | | 208 |
| 209 | I | 827 | 0 | 0 | 187 | 96 | 0 | M9- | 22 – 36 | | 3084401 | | | | 209 |
| 210 | IV | 822 | 0 | 5 | 282 | 0 | 1 | Mono | Mono | | 3098262 | | | | 210 |
| 211 | III | 789 | 35 | 3 | 0 | 283 | 0 | M9+ | 17 – 93 | | 3118152 | | | | 211 |
| 212 | IV | 827 | 0 | 0 | 282 | 0 | 1 | Mono | Mono | | 3126520 | | | | 212 |
| 213 | IV | 812 | 0 | 15 | 278 | 0 | 5 | Mono | Mono | | 3127531 | | | | 213 |
| 214 | IV | 827 | 0 | 0 | 283 | 0 | 0 | Mono | Mono | | 3179117 | | | | 214 |
| 215 | IV | 827 | 0 | 0 | 283 | 0 | 0 | Mono | Mono | | 3199276 | | | | 215 |
| 216 | IV | 823 | 0 | 4 | 281 | 0 | 2 | Mono | Mono | | 3245636 | | | | 216 |
| 217 | IV | 826 | 0 | 1 | 281 | 0 | 2 | Mono | Mono | | 3269280 | | | | 217 |
| 218 | IV | 827 | 0 | 0 | 283 | 0 | 0 | Mono | Mono | | 3285148 | | | | 218 |
| 219 | IV | 806 | 0 | 21 | 274 | 0 | 9 | Mono | Mono | | 3310270 | | | | 219 |
| 220 | IV | 827 | 0 | 0 | 282 | 0 | 1 | Mono | Mono | | 3327371 | | | | 220 |
| 221 | IV | 827 | 0 | 0 | 283 | 0 | 0 | Mono | Mono | | 3342922 | | | | 221 |
| 222 | IV | 827 | 0 | 0 | 282 | 0 | 1 | Mono | Mono | | 3364187 | | | | 222 |
| 223 | I | 827 | 0 | 0 | 82 | 201 | 0 | M9- | 22 – 59 | | 3366638 | | | | 223 |
| 224 | IV | 821 | 0 | 6 | 280 | 0 | 3 | Mono | Mono | | 3370708 | | | | 224 |
| 225 | IV | 826 | 0 | 1 | 282 | 0 | 1 | Mono | Mono | | 3370714 | | | | 225 |
| 226 | IV | 827 | 0 | 0 | 283 | 0 | 0 | Mono | Mono | | 3380902 | | | | 226 |
| 227 | III | 823 | 3 | 1 | 0 | 224 | 59 | M9+ | 4 – 93 | MD | 3385259 | | | | 227 |
| 228 | IV | 825 | 0 | 2 | 282 | 0 | 1 | Mono | Mono | | 3409772 | | | | 228 |
| 229 | IV | 818 | 0 | 9 | 279 | 0 | 4 | Mono | Mono | | 3439085 | | | | 229 |
| 230 | IV | 827 | 0 | 0 | 282 | 0 | 1 | Mono | Mono | | 3443398 | | | | 230 |
| 231 | IV | 827 | 0 | 0 | 283 | 0 | 0 | Mono | Mono | | 3446321 | | | | 231 |
| 232 | IV | 827 | 0 | 0 | 282 | 0 | 1 | Mono | Mono | | 3453914 | | | | 232 |
| 233 | IV | 827 | 0 | 0 | 283 | 0 | 0 | Mono | Mono | | 3456712 | | | | 233 |
| 234 | IV | 815 | 0 | 12 | 280 | 0 | 3 | Mono | Mono | | 3471480 | | | | 234 |
| 235 | III | 826 | 0 | 1 | 273 | 10 | 0 | M9- | 15 – 19 | | 3501139 | | | | 235 |
| 236 | IV | 825 | 0 | 2 | 281 | 0 | 2 | Mono | Mono | | 3506956 | | | | 236 |
| 237 | IV | 826 | 0 | 1 | 283 | 0 | 0 | Mono | Mono | | 3509300 | | | | 237 |
| 238 | IV | 821 | 0 | 6 | 281 | 0 | 2 | Mono | Mono | | 3518178 | | | | 238 |
| 239 | IV | 826 | 0 | 1 | 282 | 0 | 1 | Mono | Mono | | 3521762 | | | | 239 |
| 240 | IV | 826 | 0 | 1 | 283 | 0 | 0 | Mono | Mono | | 3531404 | | | | 240 |
| 241 | III | 823 | 1 | 3 | 283 | 0 | 0 | M9+ | 1 | | 3545167 | | | | 241 |
| 242 | IV | 810 | 0 | 17 | 272 | 0 | 11 | Mono | Mono | | 3553080 | | | | 242 |
| 243 | IV | 824 | 0 | 3 | 282 | 0 | 1 | Mono | Mono | | 3597108 | | | | 243 |
| 244 | III | 826 | 0 | 1 | 247 | 36 | 0 | M9- | 2 – 10 | | 3600143 | | | | 244 |
| 245 | IV | 820 | 0 | 7 | 282 | 0 | 1 | Mono | Mono | | 3624238 | | | | 245 |
| 246 | IV | 827 | 0 | 0 | 283 | 0 | 0 | Mono | Mono | | 3624644 | | | | 246 |
| 247 | IV | 653 | 0 | 174 | 231 | 0 | 52 | Mono | Mono | | 3662845 | | | | 247 |
| 248 | IV | 827 | 0 | 0 | 283 | 0 | 0 | Mono | Mono | | 3667776 | | | | 248 |
| 249 | III | 663 | 0 | 164 | 226 | 1 | 56 | M9- | 21 | | 3683471 | | | | 249 |
| 250 | IV | 825 | 0 | 2 | 280 | 0 | 3 | Mono | Mono | | 3721328 | | | | 250 |
| 251 | IV | 827 | 0 | 0 | 283 | 0 | 0 | Mono | Mono | | 3757978 | | | | 251 |
| 252 | IV | 827 | 0 | 0 | 283 | 0 | 0 | Mono | Mono | | 3777548 | | | | 252 |
| 253 | IV | 827 | 0 | 0 | 283 | 0 | 0 | Mono | Mono | | 3780899 | | | | 253 |
| 254 | IV | 825 | 0 | 2 | 282 | 0 | 1 | Mono | Mono | | 3781791 | | | | 254 |
| 255 | IV | 827 | 0 | 0 | 282 | 0 | 1 | Mono | Mono | | 3798995 | | | | 255 |
| 256 | IV | 827 | 0 | 0 | 283 | 0 | 0 | Mono | Mono | | 3818738 | | | | 256 |
| 257 | IV | 827 | 0 | 0 | 283 | 0 | 0 | Mono | Mono | | 3829722 | | | | 257 |
| 258 | IV | 826 | 0 | 1 | 283 | 0 | 0 | Mono | Mono | | 3830130 | | | | 258 |
| 259 | IV | 815 | 0 | 12 | 280 | 0 | 3 | Mono | Mono | | 3833961 | | | | 259 |
| 260 | IV | 825 | 0 | 2 | 282 | 0 | 1 | Mono | Mono | | 3875488 | | | | 260 |
| 261 | IV | 825 | 0 | 2 | 283 | 0 | 0 | Mono | Mono | | 3894810 | | | | 261 |
| 262 | IV | 812 | 0 | 15 | 281 | 0 | 2 | Mono | Mono | | 3931440 | | | | 262 |
| 263 | IV | 827 | 0 | 0 | 283 | 0 | 0 | Mono | Mono | | 3966392 | | | | 263 |
| 264 | III | 792 | 35 | 0 | 0 | 282 | 1 | M9+ | 17 – 93 | | 3966708 | | | | 264 |
| 265 | IV | 794 | 0 | 33 | 276 | 0 | 7 | Mono | Mono | | 3977240 | | | | 265 |
| 266 | IV | 806 | 0 | 21 | 279 | 0 | 4 | Mono | Mono | | 3982762 | | | | 266 |
| 267 | IV | 822 | 0 | 5 | 281 | 0 | 2 | Mono | Mono | | 3988042 | | | | 267 |
| 268 | III | 824 | 2 | 1 | 247 | 36 | 0 | BOTH | | PI | 4009014 | | | | 268 |
| 269 | IV | 826 | 0 | 1 | 282 | 0 | 1 | Mono | Mono | | 4009690 | | | | 269 |
| 270 | IV | 823 | 0 | 4 | 281 | 0 | 2 | Mono | Mono | | 4037971 | | | | 270 |
| 271 | IV | 827 | 0 | 0 | 282 | 0 | 1 | Mono | Mono | | 4065381 | | | | 271 |
| 273 | IV | 827 | 0 | 0 | 283 | 0 | 0 | Mono | Mono | | 4087052 | | | | 273 |
| 274 | III | 827 | 0 | 0 | 278 | 4 | 1 | M9- | 9 – 10 | | 4113969 | | | | 274 |

| | | | M9+ | | | M9- | | | | | Build 37 | | ISOGG | | |
|---|---|---|---|---|---|---|---|---|---|---|---|---|---|---|---|
| List | Stage | 0 | 1 | . | 0 | 1 | . | M9? | Clade(s) | Prob. | Position | Name | Subgroup Name | Amb. | List |
| 275 | II | 827 | 0 | 0 | 282 | 1 | 0 | M9- | 21 | | 4155345 | | | | 275 |
| 276 | IV | 827 | 0 | 0 | 262 | 0 | 21 | Mono | Mono | | 4179056 | | | | 276 |
| 277 | IV | 811 | 0 | 16 | 278 | 0 | 5 | Mono | Mono | | 4183123 | | | | 277 |
| 278 | IV | 827 | 0 | 0 | 282 | 0 | 1 | Mono | Mono | | 4206439 | | | | 278 |
| 279 | IV | 827 | 0 | 0 | 283 | 0 | 0 | Mono | Mono | | 4221644 | | | | 279 |
| 280 | I | 792 | 35 | 0 | 0 | 283 | 0 | M9+ | 17 – 93 | | 4223442 | | | | 280 |
| 281 | IV | 827 | 0 | 0 | 283 | 0 | 0 | Mono | Mono | | 4249478 | | | | 281 |
| 282 | IV | 825 | 0 | 2 | 283 | 0 | 0 | Mono | Mono | | 4253380 | | | | 282 |
| 283 | IV | 827 | 0 | 0 | 283 | 0 | 0 | Mono | Mono | | 4280570 | | | | 283 |
| 284 | IV | 808 | 0 | 19 | 278 | 0 | 5 | Mono | Mono | | 4328236 | | | | 284 |
| 286 | IV | 827 | 0 | 0 | 282 | 0 | 1 | Mono | Mono | | 4353235 | | | | 286 |
| 287 | IV | 825 | 0 | 2 | 283 | 0 | 0 | Mono | Mono | | 4369067 | | | | 287 |
| 288 | IV | 827 | 0 | 0 | 283 | 0 | 0 | Mono | Mono | | 4374214 | | | | 288 |
| 289 | IV | 826 | 0 | 1 | 283 | 0 | 0 | Mono | Mono | | 4374216 | | | | 289 |
| 290 | IV | 827 | 0 | 0 | 283 | 0 | 0 | Mono | Mono | | 4388629 | | | | 290 |
| 291 | IV | 827 | 0 | 0 | 283 | 0 | 0 | Mono | Mono | | 4389694 | | | | 291 |
| 292 | IV | 827 | 0 | 0 | 283 | 0 | 0 | Mono | Mono | | 4399143 | | | | 292 |
| 293 | IV | 826 | 0 | 1 | 282 | 0 | 1 | Mono | Mono | | 4436752 | | | | 293 |
| 294 | IV | 825 | 0 | 2 | 283 | 0 | 0 | Mono | Mono | | 4451126 | | | | 294 |
| 295 | III | 822 | 4 | 1 | 0 | 283 | 0 | M9+ | 4 – 93 | | 4461159 | | | | 295 |
| 296 | IV | 826 | 0 | 1 | 283 | 0 | 0 | Mono | Mono | | 4465673 | | | | 296 |
| 297 | IV | 827 | 0 | 0 | 283 | 0 | 0 | Mono | Mono | | 4465675 | | | | 297 |
| 298 | IV | 827 | 0 | 0 | 283 | 0 | 0 | Mono | Mono | | 4482533 | | | | 298 |
| 299 | IV | 825 | 0 | 2 | 281 | 0 | 2 | Mono | Mono | | 4498690 | | | | 299 |
| 300 | IV | 823 | 0 | 4 | 282 | 0 | 1 | Mono | Mono | | 4518978 | | | | 300 |
| 301 | IV | 826 | 0 | 1 | 283 | 0 | 0 | Mono | Mono | | 4527646 | | | | 301 |
| 302 | IV | 824 | 0 | 3 | 282 | 0 | 1 | Mono | Mono | | 4531473 | | | | 302 |
| 303 | IV | 823 | 0 | 4 | 280 | 0 | 3 | Mono | Mono | | 4573475 | | | | 303 |
| 304 | IV | 827 | 0 | 0 | 283 | 0 | 0 | Mono | Mono | | 4593374 | | | | 304 |
| 305 | IV | 823 | 0 | 4 | 281 | 0 | 2 | Mono | Mono | | 4593378 | | | | 305 |
| 306 | IV | 812 | 0 | 15 | 279 | 0 | 4 | Mono | Mono | | 4599754 | | | | 306 |
| 307 | IV | 814 | 0 | 13 | 279 | 0 | 4 | Mono | Mono | | 4642548 | | | | 307 |
| 308 | IV | 826 | 0 | 1 | 283 | 0 | 0 | Mono | Mono | | 4656348 | | | | 308 |
| 309 | IV | 824 | 0 | 3 | 282 | 0 | 1 | Mono | Mono | | 4662029 | | | | 309 |
| 310 | IV | 827 | 0 | 0 | 282 | 0 | 1 | Mono | Mono | | 4670423 | | | | 310 |
| 311 | IV | 827 | 0 | 0 | 283 | 0 | 0 | Mono | Mono | | 4672316 | | | | 311 |
| 312 | IV | 827 | 0 | 0 | 283 | 0 | 0 | Mono | Mono | | 4685801 | | | | 312 |
| 313 | IV | 827 | 0 | 0 | 283 | 0 | 0 | Mono | Mono | | 4696701 | | | | 313 |
| 314 | IV | 827 | 0 | 0 | 282 | 0 | 1 | Mono | Mono | | 4716925 | | | | 314 |
| 315 | IV | 827 | 0 | 0 | 282 | 0 | 1 | Mono | Mono | | 4724101 | | | | 315 |
| 316 | IV | 824 | 0 | 3 | 277 | 0 | 6 | Mono | Mono | | 4743576 | | | | 316 |
| 317 | IV | 825 | 0 | 2 | 282 | 0 | 1 | Mono | Mono | | 4771928 | | | | 317 |
| 318 | IV | 793 | 0 | 34 | 271 | 0 | 12 | Mono | Mono | | 4780764 | | | | 318 |
| 319 | IV | 827 | 0 | 0 | 282 | 0 | 1 | Mono | Mono | | 4813975 | | | | 319 |
| 320 | IV | 827 | 0 | 0 | 282 | 0 | 1 | Mono | Mono | | 4815709 | | | | 320 |
| 321 | IV | 816 | 0 | 11 | 278 | 0 | 5 | Mono | Mono | | 4816154 | | | | 321 |
| 322 | IV | 801 | 0 | 26 | 273 | 0 | 10 | Mono | Mono | | 4824672 | | | | 322 |
| 323 | IV | 822 | 0 | 5 | 281 | 0 | 2 | Mono | Mono | | 4826650 | | | | 323 |
| 324 | IV | 823 | 0 | 4 | 281 | 0 | 2 | Mono | Mono | | 4829860 | | | | 324 |
| 325 | IV | 827 | 0 | 0 | 283 | 0 | 0 | Mono | Mono | | 4849444 | | | | 325 |
| 326 | IV | 818 | 0 | 9 | 281 | 0 | 2 | Mono | Mono | | 4860430 | | | | 326 |
| 327 | IV | 827 | 0 | 0 | 283 | 0 | 0 | Mono | Mono | | 4861963 | | | | 327 |
| 328 | IV | 827 | 0 | 0 | 283 | 0 | 0 | Mono | Mono | | 4862439 | | | | 328 |
| 329 | IV | 814 | 0 | 13 | 280 | 0 | 3 | Mono | Mono | | 4869632 | | | | 329 |
| 330 | III | 826 | 0 | 1 | 0 | 283 | 0 | M9+/- | M9+/- | | 4874629 | | | | 330 |
| 331 | IV | 820 | 0 | 7 | 281 | 0 | 2 | Mono | Mono | | 4885328 | | | | 331 |
| 332 | IV | 827 | 0 | 0 | 283 | 0 | 0 | Mono | Mono | | 4886615 | | | | 332 |
| 333 | IV | 819 | 0 | 8 | 277 | 0 | 6 | Mono | Mono | | 4890139 | | | | 333 |
| 334 | IV | 0 | 827 | 0 | 0 | 283 | 0 | Mono | Mono | | 4890141 | | | | 334 |
| 335 | IV | 827 | 0 | 0 | 283 | 0 | 0 | Mono | Mono | | 4918862 | | | | 335 |
| 336 | IV | 827 | 0 | 0 | 282 | 0 | 1 | Mono | Mono | | 4920361 | | | | 336 |
| 337 | IV | 819 | 0 | 8 | 280 | 0 | 3 | Mono | Mono | | 4923729 | | | | 337 |
| 338 | IV | 827 | 0 | 0 | 283 | 0 | 0 | Mono | Mono | | 4924937 | | | | 338 |
| 339 | IV | 820 | 0 | 7 | 280 | 0 | 3 | Mono | Mono | | 4925004 | | | | 339 |
| 340 | IV | 825 | 0 | 2 | 282 | 0 | 1 | Mono | Mono | | 4925084 | | | | 340 |
| 341 | IV | 823 | 0 | 4 | 278 | 0 | 5 | Mono | Mono | | 4935539 | | | | 341 |
| 342 | IV | 827 | 0 | 0 | 283 | 0 | 0 | Mono | Mono | | 4936805 | | | | 342 |
| 343 | IV | 826 | 0 | 1 | 283 | 0 | 0 | Mono | Mono | | 4943232 | | | | 343 |
| 344 | IV | 827 | 0 | 0 | 283 | 0 | 0 | Mono | Mono | | 4959569 | | | | 344 |
| 345 | IV | 827 | 0 | 0 | 282 | 0 | 1 | Mono | Mono | | 4971950 | | | | 345 |
| 346 | III | 826 | 0 | 1 | 0 | 283 | 0 | M9+/- | M9+/- | | 4975602 | | | | 346 |
| 347 | IV | 827 | 0 | 0 | 283 | 0 | 0 | Mono | Mono | | 4988954 | | | | 347 |
| 348 | IV | 821 | 0 | 6 | 282 | 0 | 1 | Mono | Mono | | 4997130 | | | | 348 |
| 349 | IV | 819 | 0 | 8 | 280 | 0 | 3 | Mono | Mono | | 5013778 | | | | 349 |
| 350 | IV | 827 | 0 | 0 | 283 | 0 | 0 | Mono | Mono | | 5017056 | | | | 350 |
| 351 | IV | 827 | 0 | 0 | 283 | 0 | 0 | Mono | Mono | | 5021851 | | | | 351 |
| 352 | IV | 827 | 0 | 0 | 283 | 0 | 0 | Mono | Mono | | 5029265 | | | | 352 |
| 353 | III | 824 | 0 | 3 | 262 | 21 | 0 | M9- | 11 – 20 | | 5030803 | | | | 353 |
| 354 | IV | 816 | 0 | 11 | 279 | 0 | 4 | Mono | Mono | | 5034233 | | | | 354 |
| 355 | IV | 826 | 0 | 1 | 283 | 0 | 0 | Mono | Mono | | 5036720 | | | | 355 |
| 356 | IV | 825 | 0 | 2 | 282 | 0 | 1 | Mono | Mono | | 5038186 | | | | 356 |
| 357 | IV | 827 | 0 | 0 | 283 | 0 | 0 | Mono | Mono | | 5039828 | | | | 357 |
| 358 | IV | 827 | 0 | 0 | 283 | 0 | 0 | Mono | Mono | | 5049404 | | | | 358 |
| 359 | II | 826 | 1 | 0 | 283 | 0 | 0 | M9+ | 1 | | 5063793 | | | | 359 |
| 360 | III | 826 | 0 | 1 | 247 | 36 | 0 | M9- | 2 – 10 | | 5081588 | | | | 360 |
| 361 | IV | 824 | 0 | 3 | 283 | 0 | 0 | Mono | Mono | | 5086412 | | | | 361 |
| 362 | IV | 826 | 0 | 1 | 282 | 0 | 1 | Mono | Mono | | 5092017 | | | | 362 |
| 363 | IV | 822 | 0 | 5 | 282 | 0 | 1 | Mono | Mono | | 5092035 | | | | 363 |
| 364 | IV | 827 | 0 | 0 | 283 | 0 | 0 | Mono | Mono | | 5104133 | | | | 364 |
| 365 | IV | 820 | 0 | 7 | 282 | 0 | 1 | Mono | Mono | | 5122557 | | | | 365 |
| 366 | IV | 827 | 0 | 0 | 283 | 0 | 0 | Mono | Mono | | 5127976 | | | | 366 |
| 367 | IV | 826 | 0 | 1 | 283 | 0 | 0 | Mono | Mono | | 5166488 | | | | 367 |
| 368 | IV | 815 | 0 | 12 | 282 | 0 | 1 | Mono | Mono | | 5190930 | | | | 368 |
| 369 | IV | 827 | 0 | 0 | 283 | 0 | 0 | Mono | Mono | | 5205541 | | | | 369 |
| 370 | IV | 827 | 0 | 0 | 283 | 0 | 0 | Mono | Mono | | 5228166 | | | | 370 |
| 371 | IV | 826 | 0 | 1 | 282 | 0 | 1 | Mono | Mono | | 5233227 | | | | 371 |

| | | M9+ | | | M9- | | | | | | Build 37 | | ISOGG | | |
|---|---|---|---|---|---|---|---|---|---|---|---|---|---|---|---|
| List | Stage | 0 | 1 | . | 0 | 1 | . | M9? | Clade(s) | Prob. | Position | Name | Subgroup Name | Amb. | List |
| 372 | IV | 827 | 0 | 0 | 283 | 0 | 0 | Mono | Mono | | 5243188 | | | | 372 |
| 373 | IV | 827 | 0 | 0 | 283 | 0 | 0 | Mono | Mono | | 5247381 | | | | 373 |
| 374 | IV | 827 | 0 | 0 | 282 | 0 | 1 | Mono | Mono | | 5249934 | | | | 374 |
| 375 | IV | 825 | 0 | 2 | 283 | 0 | 0 | Mono | Mono | | 5252099 | | | | 375 |
| 376 | IV | 827 | 0 | 0 | 283 | 0 | 0 | Mono | Mono | | 5270504 | | | | 376 |
| 377 | IV | 827 | 0 | 0 | 282 | 0 | 1 | Mono | Mono | | 5287832 | | | | 377 |
| 378 | IV | 820 | 0 | 7 | 281 | 0 | 2 | Mono | Mono | | 5298470 | | | | 378 |
| 379 | IV | 821 | 0 | 6 | 280 | 0 | 3 | Mono | Mono | | 5340671 | | | | 379 |
| 380 | IV | 827 | 0 | 0 | 283 | 0 | 0 | Mono | Mono | | 5354507 | | | | 380 |
| 381 | IV | 826 | 0 | 1 | 283 | 0 | 0 | Mono | Mono | | 5358346 | | | | 381 |
| 382 | IV | 822 | 0 | 5 | 279 | 0 | 4 | Mono | Mono | | 5360092 | | | | 382 |
| 383 | IV | 826 | 0 | 1 | 281 | 0 | 2 | Mono | Mono | | 5372983 | | | | 383 |
| 384 | IV | 827 | 0 | 0 | 283 | 0 | 0 | Mono | Mono | | 5377315 | | | | 384 |
| 385 | IV | 824 | 0 | 3 | 282 | 0 | 1 | Mono | Mono | | 5384673 | | | | 385 |
| 386 | IV | 827 | 0 | 0 | 282 | 0 | 1 | Mono | Mono | | 5407917 | | | | 386 |
| 387 | IV | 826 | 0 | 1 | 282 | 0 | 1 | Mono | Mono | | 5413643 | | | | 387 |
| 388 | IV | 827 | 0 | 0 | 283 | 0 | 0 | Mono | Mono | | 5426528 | | | | 388 |
| 389 | IV | 820 | 0 | 7 | 279 | 0 | 4 | Mono | Mono | | 5426745 | | | | 389 |
| 390 | IV | 826 | 0 | 1 | 282 | 0 | 1 | Mono | Mono | | 5427360 | | | | 390 |
| 391 | IV | 827 | 0 | 0 | 283 | 0 | 0 | Mono | Mono | | 5449667 | | | | 391 |
| 392 | IV | 826 | 0 | 1 | 283 | 0 | 0 | Mono | Mono | | 5470887 | | | | 392 |
| 393 | IV | 827 | 0 | 0 | 283 | 0 | 0 | Mono | Mono | | 5504106 | | | | 393 |
| 394 | IV | 826 | 0 | 1 | 281 | 0 | 2 | Mono | Mono | | 5521648 | | | | 394 |
| 395 | IV | 827 | 0 | 0 | 283 | 0 | 0 | Mono | Mono | | 5532299 | | | | 395 |
| 396 | IV | 826 | 0 | 1 | 283 | 0 | 0 | Mono | Mono | | 5548849 | | | | 396 |
| 397 | IV | 827 | 0 | 0 | 283 | 0 | 0 | Mono | Mono | | 5557491 | | | | 397 |
| 398 | III | 814 | 1 | 12 | 280 | 0 | 3 | M9+ | 1 | | 5557632 | | | | 398 |
| 399 | IV | 827 | 0 | 0 | 283 | 0 | 0 | Mono | Mono | | 5566038 | | | | 399 |
| 400 | IV | 827 | 0 | 0 | 283 | 0 | 0 | Mono | Mono | | 5573860 | | | | 400 |
| 401 | IV | 826 | 0 | 1 | 282 | 0 | 1 | Mono | Mono | | 5586370 | | | | 401 |
| 402 | IV | 827 | 0 | 0 | 283 | 0 | 0 | Mono | Mono | | 5603640 | | | | 402 |
| 403 | IV | 827 | 0 | 0 | 283 | 0 | 0 | Mono | Mono | | 5626072 | | | | 403 |
| 404 | IV | 827 | 0 | 0 | 282 | 0 | 1 | Mono | Mono | | 5645719 | | | | 404 |
| 405 | IV | 817 | 0 | 10 | 280 | 0 | 3 | Mono | Mono | | 5646254 | | | | 405 |
| 406 | IV | 827 | 0 | 0 | 283 | 0 | 0 | Mono | Mono | | 5650504 | | | | 406 |
| 407 | IV | 826 | 0 | 1 | 283 | 0 | 0 | Mono | Mono | | 5680894 | | | | 407 |
| 408 | IV | 802 | 0 | 25 | 273 | 0 | 10 | Mono | Mono | | 5697256 | | | | 408 |
| 409 | IV | 818 | 0 | 9 | 280 | 0 | 3 | Mono | Mono | | 5708471 | | | | 409 |
| 410 | IV | 821 | 0 | 6 | 281 | 0 | 2 | Mono | Mono | | 5729144 | | | | 410 |
| 411 | IV | 827 | 0 | 0 | 283 | 0 | 0 | Mono | Mono | | 5747681 | | | | 411 |
| 413 | III | 817 | 0 | 10 | 183 | 96 | 4 | M9- | 22 – 36 | | 5818860 | | | | 413 |
| 414 | IV | 820 | 0 | 7 | 280 | 0 | 3 | Mono | Mono | | 5832456 | | | | 414 |
| 415 | I | 824 | 3 | 0 | 274 | 9 | 0 | BOTH | | PI | 5836035 | | | | 415 |
| 416 | IV | 827 | 0 | 0 | 282 | 0 | 1 | Mono | Mono | | 5853004 | | | | 416 |
| 417 | IV | 827 | 0 | 0 | 283 | 0 | 0 | Mono | Mono | | 5857806 | | | | 417 |
| 418 | IV | 820 | 0 | 7 | 281 | 0 | 2 | Mono | Mono | | 5881606 | | | | 418 |
| 419 | IV | 822 | 0 | 5 | 281 | 0 | 2 | Mono | Mono | | 5897758 | | | | 419 |
| 420 | IV | 827 | 0 | 0 | 283 | 0 | 0 | Mono | Mono | | 5905252 | | | | 420 |
| 421 | IV | 827 | 0 | 0 | 282 | 0 | 1 | Mono | Mono | | 5917202 | | | | 421 |
| 422 | IV | 826 | 0 | 1 | 283 | 0 | 0 | Mono | Mono | | 5928113 | | | | 422 |
| 423 | IV | 825 | 0 | 2 | 282 | 0 | 1 | Mono | Mono | | 5928852 | | | | 423 |
| 424 | IV | 824 | 0 | 3 | 283 | 0 | 0 | Mono | Mono | | 5965169 | | | | 424 |
| 425 | IV | 826 | 0 | 1 | 283 | 0 | 0 | Mono | Mono | | 5969507 | | | | 425 |
| 426 | IV | 817 | 0 | 10 | 282 | 0 | 1 | Mono | Mono | | 5977725 | | | | 426 |
| 427 | IV | 827 | 0 | 0 | 283 | 0 | 0 | Mono | Mono | | 5984066 | | | | 427 |
| 428 | II | 826 | 1 | 0 | 283 | 0 | 0 | M9+ | 1 | | 5985179 | | | | 428 |
| 429 | IV | 827 | 0 | 0 | 283 | 0 | 0 | Mono | Mono | | 5990878 | | | | 429 |
| 430 | I | 827 | 0 | 0 | 246 | 37 | 0 | M9- | 1 – 10 | | 6000464 | | | | 430 |
| 431 | IV | 826 | 0 | 1 | 283 | 0 | 0 | Mono | Mono | | 6019479 | | | | 431 |
| 432 | IV | 827 | 0 | 0 | 282 | 0 | 1 | Mono | Mono | | 6027715 | | | | 432 |
| 433 | IV | 823 | 0 | 4 | 282 | 0 | 1 | Mono | Mono | | 6039351 | | | | 433 |
| 434 | IV | 822 | 0 | 5 | 283 | 0 | 0 | Mono | Mono | | 6051738 | | | | 434 |
| 435 | IV | 827 | 0 | 0 | 282 | 0 | 1 | Mono | Mono | | 6073852 | | | | 435 |
| 436 | IV | 827 | 0 | 0 | 283 | 0 | 0 | Mono | Mono | | 6093865 | | | | 436 |
| 437 | IV | 820 | 0 | 7 | 279 | 0 | 4 | Mono | Mono | | 6190214 | | | | 437 |
| 438 | III | 825 | 0 | 2 | 277 | 6 | 0 | M9- | 12 | | 6345565 | | | | 438 |
| 439 | IV | 827 | 0 | 0 | 282 | 0 | 1 | Mono | Mono | | 6356478 | | | | 439 |
| 440 | IV | 827 | 0 | 0 | 283 | 0 | 0 | Mono | Mono | | 6364661 | | | | 440 |
| 441 | IV | 827 | 0 | 0 | 283 | 0 | 0 | Mono | Mono | | 6372196 | | | | 441 |
| 442 | IV | 827 | 0 | 0 | 283 | 0 | 0 | Mono | Mono | | 6373152 | | | | 442 |
| 444 | IV | 827 | 0 | 0 | 283 | 0 | 0 | Mono | Mono | | 6396326 | | | | 444 |
| 445 | IV | 827 | 0 | 0 | 282 | 0 | 1 | Mono | Mono | | 6399711 | | | | 445 |
| 446 | IV | 808 | 0 | 19 | 277 | 0 | 6 | Mono | Mono | | 6401978 | | | | 446 |
| 447 | IV | 819 | 0 | 8 | 278 | 0 | 5 | Mono | Mono | | 6404526 | | | | 447 |
| 448 | I | 827 | 0 | 0 | 178 | 105 | 0 | M9- | 37 – 59 | | 6424387 | | | | 448 |
| 450 | IV | 827 | 0 | 0 | 283 | 0 | 0 | Mono | Mono | | 6425538 | | | | 450 |
| 451 | IV | 827 | 0 | 0 | 283 | 0 | 0 | Mono | Mono | | 6425539 | | | | 451 |
| 452 | IV | 819 | 0 | 8 | 282 | 0 | 1 | Mono | Mono | | 6460016 | | | | 452 |
| 453 | III | 792 | 33 | 2 | 0 | 275 | 8 | M9+ | 17 – 93 | MD | 6482140 | | | | 453 |
| 454 | IV | 0 | 808 | 19 | 0 | 279 | 4 | Mono | Mono | | 6523383 | | | | 454 |
| 455 | IV | 827 | 0 | 0 | 283 | 0 | 0 | Mono | Mono | | 6530338 | | | | 455 |
| 456 | IV | 814 | 0 | 13 | 275 | 0 | 8 | Mono | Mono | | 6541648 | | | | 456 |
| 457 | IV | 824 | 0 | 3 | 282 | 0 | 1 | Mono | Mono | | 6575897 | | | | 457 |
| 458 | IV | 782 | 0 | 45 | 271 | 0 | 12 | Mono | Mono | | 6586098 | | | | 458 |
| 459 | IV | 827 | 0 | 0 | 283 | 0 | 0 | Mono | Mono | | 6601364 | | | | 459 |
| 460 | IV | 827 | 0 | 0 | 283 | 0 | 0 | Mono | Mono | | 6615362 | | | | 460 |
| 461 | IV | 820 | 0 | 7 | 282 | 0 | 1 | Mono | Mono | | 6624215 | | | | 461 |
| 465 | IV | 827 | 0 | 0 | 283 | 0 | 0 | Mono | Mono | | 6635162 | | | | 465 |
| 469 | IV | 827 | 0 | 0 | 283 | 0 | 0 | Mono | Mono | | 6659729 | | | | 469 |
| 472 | IV | 827 | 0 | 0 | 283 | 0 | 0 | Mono | Mono | | 6661837 | | | | 472 |
| 473 | IV | 827 | 0 | 0 | 283 | 0 | 0 | Mono | Mono | | 6661973 | | | | 473 |
| 478 | IV | 827 | 0 | 0 | 283 | 0 | 0 | Mono | Mono | | 6670598 | | | | 478 |
| 487 | IV | 826 | 0 | 1 | 283 | 0 | 0 | Mono | Mono | | 6692993 | | | | 487 |
| 489 | II | 826 | 1 | 0 | 283 | 0 | 0 | M9+ | 1 | | 6698445 | | | | 489 |
| 493 | IV | 827 | 0 | 0 | 283 | 0 | 0 | Mono | Mono | | 6707367 | | | | 493 |
| 494 | IV | 0 | 827 | 0 | 0 | 283 | 0 | Mono | Mono | | 6707890 | | | | 494 |

| | | M9+ | | | M9- | | | | | | Build 37 | | ISOGG | | |
|---|---|---|---|---|---|---|---|---|---|---|---|---|---|---|---|
| List | Stage | 0 | 1 | . | 0 | 1 | . | M9? | Clade(s) | Prob. | Position | Name | Subgroup Name | Amb. | List |
| 495 | II | 826 | 1 | 0 | 283 | 0 | 0 | M9+ | 1 | | 6709045 | | | | 495 |
| 499 | IV | 827 | 0 | 0 | 283 | 0 | 0 | Mono | Mono | | 6717693 | | | | 499 |
| 501 | IV | 827 | 0 | 0 | 282 | 0 | 1 | Mono | Mono | | 6728092 | | | | 501 |
| 502 | IV | 827 | 0 | 0 | 282 | 0 | 1 | Mono | Mono | | 6728094 | | | | 502 |
| 505 | IV | 827 | 0 | 0 | 283 | 0 | 0 | Mono | Mono | | 6736127 | | | | 505 |
| 506 | IV | 827 | 0 | 0 | 283 | 0 | 0 | Mono | Mono | | 6736371 | | | | 506 |
| 512 | IV | 827 | 0 | 0 | 283 | 0 | 0 | Mono | Mono | | 6740510 | | | | 512 |
| 513 | IV | 827 | 0 | 0 | 283 | 0 | 0 | Mono | Mono | | 6740599 | | | | 513 |
| 516 | IV | 827 | 0 | 0 | 283 | 0 | 0 | Mono | Mono | | 6743405 | | | | 516 |
| 526 | III | 825 | 1 | 1 | 282 | 0 | 1 | M9+ | 1 | | 6779609 | | | | 526 |
| 533 | IV | 827 | 0 | 0 | 282 | 0 | 1 | Mono | Mono | | 6796065 | | | | 533 |
| 538 | IV | 827 | 0 | 0 | 283 | 0 | 0 | Mono | Mono | | 6811475 | | | | 538 |
| 540 | IV | 827 | 0 | 0 | 283 | 0 | 0 | Mono | Mono | | 6812031 | | | | 540 |
| 548 | II | 826 | 1 | 0 | 283 | 0 | 0 | M9+ | 1 | | 6835196 | | | | 548 |
| 552 | IV | 825 | 0 | 2 | 283 | 0 | 0 | Mono | Mono | | 6845657 | | | | 552 |
| 558 | IV | 827 | 0 | 0 | 283 | 0 | 0 | Mono | Mono | | 6848390 | | | | 558 |
| 561 | IV | 827 | 0 | 0 | 283 | 0 | 0 | Mono | Mono | | 6849811 | | | | 561 |
| 563 | IV | 826 | 0 | 1 | 283 | 0 | 0 | Mono | Mono | | 6851541 | | | | 563 |
| 567 | IV | 826 | 0 | 1 | 283 | 0 | 0 | Mono | Mono | | 6854765 | | | | 567 |
| 575 | IV | 826 | 0 | 1 | 283 | 0 | 0 | Mono | Mono | | 6863631 | | | | 575 |
| 606 | IV | 827 | 0 | 0 | 283 | 0 | 0 | Mono | Mono | | 6896546 | | | | 606 |
| 607 | IV | 827 | 0 | 0 | 283 | 0 | 0 | Mono | Mono | | 6899215 | | | | 607 |
| 619 | IV | 827 | 0 | 0 | 283 | 0 | 0 | Mono | Mono | | 6919614 | | | | 619 |
| 621 | IV | 827 | 0 | 0 | 282 | 0 | 1 | Mono | Mono | | 6924694 | | | | 621 |
| 631 | IV | 827 | 0 | 0 | 283 | 0 | 0 | Mono | Mono | | 6932074 | | | | 631 |
| 632 | IV | 827 | 0 | 0 | 283 | 0 | 0 | Mono | Mono | | 6932080 | | | | 632 |
| 633 | IV | 827 | 0 | 0 | 283 | 0 | 0 | Mono | Mono | | 6932104 | | | | 633 |
| 638 | IV | 827 | 0 | 0 | 283 | 0 | 0 | Mono | Mono | | 6932182 | | | | 638 |
| 645 | IV | 0 | 827 | 0 | 0 | 283 | 0 | Mono | Mono | | 6932790 | | | | 645 |
| 652 | II | 826 | 1 | 0 | 283 | 0 | 0 | M9+ | 2 | | 6933016 | | | | 652 |
| 656 | II | 826 | 1 | 0 | 283 | 0 | 0 | M9+ | 1 | | 6936636 | | | | 656 |
| 658 | IV | 827 | 0 | 0 | 283 | 0 | 0 | Mono | Mono | | 6938314 | | | | 658 |
| 660 | IV | 826 | 0 | 1 | 283 | 0 | 0 | Mono | Mono | | 6938864 | | | | 660 |
| 661 | IV | 827 | 0 | 0 | 283 | 0 | 0 | Mono | Mono | | 6939706 | | | | 661 |
| 662 | IV | 827 | 0 | 0 | 283 | 0 | 0 | Mono | Mono | | 6939851 | | | | 662 |
| 666 | IV | 827 | 0 | 0 | 283 | 0 | 0 | Mono | Mono | | 6943622 | | | | 666 |
| 669 | IV | 825 | 0 | 2 | 283 | 0 | 0 | Mono | Mono | | 6952003 | | | | 669 |
| 670 | IV | 827 | 0 | 0 | 283 | 0 | 0 | Mono | Mono | | 6952023 | | | | 670 |
| 671 | IV | 827 | 0 | 0 | 283 | 0 | 0 | Mono | Mono | | 6952036 | | | | 671 |
| 672 | IV | 827 | 0 | 0 | 283 | 0 | 0 | Mono | Mono | | 6952061 | | | | 672 |
| 675 | IV | 827 | 0 | 0 | 283 | 0 | 0 | Mono | Mono | | 6954340 | | | | 675 |
| 677 | IV | 826 | 0 | 1 | 283 | 0 | 0 | Mono | Mono | | 6955423 | | | | 677 |
| 678 | IV | 827 | 0 | 0 | 283 | 0 | 0 | Mono | Mono | | 6955451 | | | | 678 |
| 683 | IV | 827 | 0 | 0 | 282 | 0 | 1 | Mono | Mono | | 6961386 | | | | 683 |
| 685 | IV | 827 | 0 | 0 | 283 | 0 | 0 | Mono | Mono | | 6961471 | | | | 685 |
| 689 | IV | 827 | 0 | 0 | 283 | 0 | 0 | Mono | Mono | | 6968550 | | | | 689 |
| 691 | IV | 827 | 0 | 0 | 283 | 0 | 0 | Mono | Mono | | 6970565 | | | | 691 |
| 692 | IV | 827 | 0 | 0 | 283 | 0 | 0 | Mono | Mono | | 6973766 | | | | 692 |
| 693 | IV | 826 | 0 | 1 | 283 | 0 | 0 | Mono | Mono | | 6976734 | | | | 693 |
| 694 | IV | 827 | 0 | 0 | 283 | 0 | 0 | Mono | Mono | | 6983355 | | | | 694 |
| 696 | IV | 827 | 0 | 0 | 283 | 0 | 0 | Mono | Mono | | 6986981 | | | | 696 |
| 697 | IV | 827 | 0 | 0 | 283 | 0 | 0 | Mono | Mono | | 6988555 | | | | 697 |
| 699 | II | 826 | 1 | 0 | 283 | 0 | 0 | M9+ | 1 | | 6992150 | | | | 699 |
| 703 | IV | 827 | 0 | 0 | 283 | 0 | 0 | Mono | Mono | | 6995947 | | | | 703 |
| 704 | IV | 826 | 0 | 1 | 283 | 0 | 0 | Mono | Mono | | 6995952 | | | | 704 |
| 709 | IV | 827 | 0 | 0 | 283 | 0 | 0 | Mono | Mono | | 7035173 | | | | 709 |
| 710 | IV | 827 | 0 | 0 | 283 | 0 | 0 | Mono | Mono | | 7035520 | | | | 710 |
| 713 | IV | 827 | 0 | 0 | 283 | 0 | 0 | Mono | Mono | | 7045854 | | | | 713 |
| 715 | IV | 827 | 0 | 0 | 283 | 0 | 0 | Mono | Mono | | 7046114 | | | | 715 |
| 718 | IV | 827 | 0 | 0 | 283 | 0 | 0 | Mono | Mono | | 7050339 | | | | 718 |
| 720 | IV | 824 | 0 | 3 | 282 | 0 | 1 | Mono | Mono | | 7057319 | | | | 720 |
| 721 | IV | 827 | 0 | 0 | 283 | 0 | 0 | Mono | Mono | | 7057795 | | | | 721 |
| 724 | IV | 827 | 0 | 0 | 283 | 0 | 0 | Mono | Mono | | 7067665 | | | | 724 |
| 732 | IV | 827 | 0 | 0 | 283 | 0 | 0 | Mono | Mono | | 7075557 | | | | 732 |
| 738 | IV | 826 | 0 | 1 | 283 | 0 | 0 | Mono | Mono | | 7087279 | | | | 738 |
| 739 | IV | 826 | 0 | 1 | 283 | 0 | 0 | Mono | Mono | | 7087315 | | | | 739 |
| 744 | III | 826 | 1 | 0 | 281 | 0 | 2 | M9+ | 1 | | 7103167 | | | | 744 |
| 746 | IV | 819 | 0 | 8 | 280 | 0 | 3 | Mono | Mono | | 7109122 | | | | 746 |
| 748 | IV | 826 | 0 | 1 | 283 | 0 | 0 | Mono | Mono | | 7126454 | | | | 748 |
| 751 | IV | 815 | 0 | 12 | 279 | 0 | 4 | Mono | Mono | | 7134120 | | | | 751 |
| 752 | IV | 824 | 0 | 3 | 283 | 0 | 0 | Mono | Mono | | 7134400 | | | | 752 |
| 754 | IV | 827 | 0 | 0 | 283 | 0 | 0 | Mono | Mono | | 7136012 | | | | 754 |
| 756 | IV | 826 | 0 | 1 | 283 | 0 | 0 | Mono | Mono | | 7136294 | | | | 756 |
| 757 | IV | 826 | 0 | 1 | 283 | 0 | 0 | Mono | Mono | | 7137632 | | | | 757 |
| 758 | IV | 826 | 0 | 1 | 283 | 0 | 0 | Mono | Mono | | 7141536 | | | | 758 |
| 759 | IV | 814 | 0 | 13 | 279 | 0 | 4 | Mono | Mono | | 7142860 | | | | 759 |
| 760 | IV | 827 | 0 | 0 | 283 | 0 | 0 | Mono | Mono | | 7143700 | | | | 760 |
| 762 | IV | 827 | 0 | 0 | 283 | 0 | 0 | Mono | Mono | | 7145620 | | | | 762 |
| 764 | IV | 827 | 0 | 0 | 283 | 0 | 0 | Mono | Mono | | 7149287 | | | | 764 |
| 765 | IV | 827 | 0 | 0 | 283 | 0 | 0 | Mono | Mono | | 7149682 | | | | 765 |
| 766 | IV | 827 | 0 | 0 | 283 | 0 | 0 | Mono | Mono | | 7149924 | | | | 766 |
| 767 | IV | 827 | 0 | 0 | 283 | 0 | 0 | Mono | Mono | | 7151887 | | | | 767 |
| 769 | IV | 819 | 0 | 8 | 280 | 0 | 3 | Mono | Mono | | 7154345 | | | | 769 |
| 770 | IV | 826 | 0 | 1 | 280 | 0 | 3 | Mono | Mono | | 7155343 | | | | 770 |
| 772 | IV | 827 | 0 | 0 | 283 | 0 | 0 | Mono | Mono | | 7158945 | | | | 772 |
| 774 | IV | 827 | 0 | 0 | 283 | 0 | 0 | Mono | Mono | | 7160115 | | | | 774 |
| 775 | IV | 827 | 0 | 0 | 283 | 0 | 0 | Mono | Mono | | 7162736 | | | | 775 |
| 777 | III | 810 | 0 | 17 | 269 | 11 | 3 | M9- | 37 – 38 | | 7170563 | | | | 777 |
| 778 | IV | 827 | 0 | 0 | 283 | 0 | 0 | Mono | Mono | | 7171069 | | | | 778 |
| 782 | IV | 820 | 0 | 7 | 274 | 0 | 9 | Mono | Mono | | 7180774 | | | | 782 |
| 784 | III | 816 | 0 | 11 | 244 | 37 | 2 | M9- | 1 – 10 | | 7184141 | | | | 784 |
| 785 | IV | 827 | 0 | 0 | 283 | 0 | 0 | Mono | Mono | | 7188046 | | | | 785 |
| 786 | IV | 827 | 0 | 0 | 283 | 0 | 0 | Mono | Mono | | 7188055 | | | | 786 |
| 787 | IV | 827 | 0 | 0 | 283 | 0 | 0 | Mono | Mono | | 7192109 | | | | 787 |
| 789 | IV | 825 | 0 | 2 | 282 | 0 | 1 | Mono | Mono | | 7193928 | | | | 789 |
| 790 | IV | 826 | 0 | 1 | 283 | 0 | 0 | Mono | Mono | | 7196932 | | | | 790 |

| | | | M9+ | | | M9- | | | | | Build 37 | | ISOGG | | |
|---|---|---|---|---|---|---|---|---|---|---|---|---|---|---|---|
| List | Stage | 0 | 1 | . | 0 | 1 | . | M9? | Clade(s) | Prob. | Position | Name | Subgroup Name | Amb. | List |
| 793 | IV | 827 | 0 | 0 | 283 | 0 | 0 | Mono | Mono | | 7209616 | | | | 793 |
| 794 | IV | 827 | 0 | 0 | 283 | 0 | 0 | Mono | Mono | | 7209642 | | | | 794 |
| 795 | IV | 827 | 0 | 0 | 283 | 0 | 0 | Mono | Mono | | 7211776 | | | | 795 |
| 799 | IV | 746 | 0 | 81 | 254 | 0 | 29 | Mono | Mono | | 7224075 | | | | 799 |
| 800 | IV | 827 | 0 | 0 | 283 | 0 | 0 | Mono | Mono | | 7225848 | | | | 800 |
| 803 | IV | 827 | 0 | 0 | 283 | 0 | 0 | Mono | Mono | | 7233512 | | | | 803 |
| 804 | II | 827 | 0 | 0 | 281 | 2 | 0 | M9- | 8 | | 7234609 | | | | 804 |
| 805 | IV | 827 | 0 | 0 | 283 | 0 | 0 | Mono | Mono | | 7241447 | | | | 805 |
| 806 | IV | 827 | 0 | 0 | 283 | 0 | 0 | Mono | Mono | | 7241448 | | | | 806 |
| 807 | IV | 825 | 0 | 2 | 283 | 0 | 0 | Mono | Mono | | 7241566 | | | | 807 |
| 808 | IV | 827 | 0 | 0 | 283 | 0 | 0 | Mono | Mono | | 7245239 | | | | 808 |
| 809 | IV | 827 | 0 | 0 | 283 | 0 | 0 | Mono | Mono | | 7249562 | | | | 809 |
| 818 | IV | 827 | 0 | 0 | 283 | 0 | 0 | Mono | Mono | | 7279986 | | | | 818 |
| 836 | IV | 827 | 0 | 0 | 283 | 0 | 0 | Mono | Mono | | 7324971 | | | | 836 |
| 839 | IV | 827 | 0 | 0 | 283 | 0 | 0 | Mono | Mono | | 7355673 | | | | 839 |
| 841 | I | 827 | 0 | 0 | 260 | 23 | 0 | M9- | 60 – 66 | | 7367950 | | | | 841 |
| 843 | IV | 827 | 0 | 0 | 283 | 0 | 0 | Mono | Mono | | 7377160 | | | | 843 |
| 846 | IV | 827 | 0 | 0 | 281 | 0 | 2 | Mono | Mono | | 7390996 | | | | 846 |
| 847 | IV | 827 | 0 | 0 | 282 | 0 | 1 | Mono | Mono | | 7395839 | | | | 847 |
| 852 | IV | 827 | 0 | 0 | 283 | 0 | 0 | Mono | Mono | | 7403244 | | | | 852 |
| 854 | II | 826 | 1 | 0 | 283 | 0 | 0 | M9+ | 1 | | 7403484 | | | | 854 |
| 859 | IV | 827 | 0 | 0 | 283 | 0 | 0 | Mono | Mono | | 7433394 | | | | 859 |
| 860 | IV | 827 | 0 | 0 | 283 | 0 | 0 | Mono | Mono | | 7508686 | | | | 860 |
| 861 | IV | 821 | 0 | 6 | 282 | 0 | 1 | Mono | Mono | | 7512329 | | | | 861 |
| 868 | IV | 810 | 0 | 17 | 276 | 0 | 7 | Mono | Mono | | 7527958 | | | | 868 |
| 869 | IV | 827 | 0 | 0 | 283 | 0 | 0 | Mono | Mono | | 7528917 | | | | 869 |
| 872 | IV | 827 | 0 | 0 | 283 | 0 | 0 | Mono | Mono | | 7533750 | | | | 872 |
| 873 | III | 824 | 0 | 3 | 245 | 37 | 1 | M9- | 1 – 10 | 1H | 7534418 | | | | 873 |
| 877 | IV | 819 | 0 | 8 | 279 | 0 | 4 | Mono | Mono | | 7542628 | | | | 877 |
| 878 | IV | 826 | 0 | 1 | 283 | 0 | 0 | Mono | Mono | | 7543038 | | | | 878 |
| 890 | IV | 827 | 0 | 0 | 283 | 0 | 0 | Mono | Mono | | 7558795 | | | | 890 |
| 895 | II | 826 | 1 | 0 | 283 | 0 | 0 | M9+ | 1 | | 7569265 | | | | 895 |
| 897 | IV | 827 | 0 | 0 | 283 | 0 | 0 | Mono | Mono | | 7570678 | | | | 897 |
| 898 | IV | 827 | 0 | 0 | 283 | 0 | 0 | Mono | Mono | | 7571122 | | | | 898 |
| 902 | IV | 827 | 0 | 0 | 283 | 0 | 0 | Mono | Mono | | 7583210 | | | | 902 |
| 919 | IV | 827 | 0 | 0 | 283 | 0 | 0 | Mono | Mono | | 7599930 | | | | 919 |
| 920 | IV | 827 | 0 | 0 | 283 | 0 | 0 | Mono | Mono | | 7602368 | | | | 920 |
| 922 | III | 825 | 1 | 1 | 283 | 0 | 0 | M9+ | 1 | | 7602738 | | | | 922 |
| 927 | IV | 827 | 0 | 0 | 283 | 0 | 0 | Mono | Mono | | 7611416 | | | | 927 |
| 932 | IV | 827 | 0 | 0 | 283 | 0 | 0 | Mono | Mono | | 7626710 | | | | 932 |
| 937 | IV | 823 | 0 | 4 | 282 | 0 | 1 | Mono | Mono | | 7635648 | | | | 937 |
| 951 | IV | 826 | 0 | 1 | 283 | 0 | 0 | Mono | Mono | | 7654523 | | | | 951 |
| 957 | IV | 826 | 0 | 1 | 282 | 0 | 1 | Mono | Mono | | 7660612 | | | | 957 |
| 958 | IV | 826 | 0 | 1 | 283 | 0 | 0 | Mono | Mono | | 7660613 | | | | 958 |
| 960 | IV | 825 | 0 | 2 | 283 | 0 | 0 | Mono | Mono | | 7665048 | | | | 960 |
| 967 | IV | 827 | 0 | 0 | 283 | 0 | 0 | Mono | Mono | | 7681463 | | | | 967 |
| 971 | IV | 825 | 0 | 2 | 283 | 0 | 0 | Mono | Mono | | 7684952 | | | | 971 |
| 976 | IV | 827 | 0 | 0 | 283 | 0 | 0 | Mono | Mono | | 7690447 | | | | 976 |
| 977 | IV | 827 | 0 | 0 | 283 | 0 | 0 | Mono | Mono | | 7690471 | | | | 977 |
| 980 | IV | 826 | 0 | 1 | 283 | 0 | 0 | Mono | Mono | | 7702181 | | | | 980 |
| 983 | IV | 827 | 0 | 0 | 283 | 0 | 0 | Mono | Mono | | 7707483 | | | | 983 |
| 985 | IV | 827 | 0 | 0 | 283 | 0 | 0 | Mono | Mono | | 7709218 | | | | 985 |
| 987 | I | 824 | 3 | 0 | 0 | 283 | 0 | M9+ | 3 – 93 | | 7714986 | | | | 987 |
| 998 | III | 826 | 1 | 0 | 282 | 0 | 1 | M9+ | NU | | 7730515 | | | | 998 |
| 1002 | IV | 827 | 0 | 0 | 283 | 0 | 0 | Mono | Mono | | 7736134 | | | | 1002 |
| 1016 | IV | 826 | 0 | 1 | 283 | 0 | 0 | Mono | Mono | | 7756702 | | | | 1016 |
| 1017 | I | 827 | 0 | 0 | 260 | 23 | 0 | M9- | 60 – 66 | | 7759160 | | | | 1017 |
| 1029 | IV | 827 | 0 | 0 | 283 | 0 | 0 | Mono | Mono | | 7782651 | | | | 1029 |
| 1030 | IV | 827 | 0 | 0 | 283 | 0 | 0 | Mono | Mono | | 7790703 | | | | 1030 |
| 1035 | IV | 827 | 0 | 0 | 283 | 0 | 0 | Mono | Mono | | 7794147 | | | | 1035 |
| 1037 | IV | 827 | 0 | 0 | 283 | 0 | 0 | Mono | Mono | | 7796708 | | | | 1037 |
| 1046 | IV | 827 | 0 | 0 | 283 | 0 | 0 | Mono | Mono | | 7813450 | | | | 1046 |
| 1047 | I | 827 | 0 | 0 | 274 | 9 | 0 | M9- | 61 – 64 | | 7813674 | | | | 1047 |
| 1050 | II | 826 | 1 | 0 | 283 | 0 | 0 | M9+ | 2 | | 7830165 | | | | 1050 |
| 1052 | IV | 827 | 0 | 0 | 283 | 0 | 0 | Mono | Mono | | 7833418 | | | | 1052 |
| 1053 | IV | 826 | 0 | 1 | 282 | 0 | 1 | Mono | Mono | | 7835840 | | | | 1053 |
| 1054 | IV | 827 | 0 | 0 | 283 | 0 | 0 | Mono | Mono | | 7835847 | | | | 1054 |
| 1059 | IV | 0 | 827 | 0 | 0 | 283 | 0 | Mono | Mono | | 7844490 | | | | 1059 |
| 1062 | IV | 827 | 0 | 0 | 283 | 0 | 0 | Mono | Mono | | 7850855 | | | | 1062 |
| 1069 | IV | 827 | 0 | 0 | 283 | 0 | 0 | Mono | Mono | | 7857743 | | | | 1069 |
| 1070 | IV | 822 | 0 | 5 | 281 | 0 | 2 | Mono | Mono | | 7858871 | | | | 1070 |
| 1072 | IV | 827 | 0 | 0 | 283 | 0 | 0 | Mono | Mono | | 7872203 | | | | 1072 |
| 1075 | IV | 827 | 0 | 0 | 283 | 0 | 0 | Mono | Mono | | 7887185 | | | | 1075 |
| 1081 | IV | 827 | 0 | 0 | 283 | 0 | 0 | Mono | Mono | | 7900462 | | | | 1081 |
| 1087 | IV | 827 | 0 | 0 | 283 | 0 | 0 | Mono | Mono | | 7907021 | | | | 1087 |
| 1089 | II | 826 | 1 | 0 | 283 | 0 | 0 | M9+ | 1 | | 7908027 | | | | 1089 |
| 1090 | IV | 827 | 0 | 0 | 283 | 0 | 0 | Mono | Mono | | 7908950 | | | | 1090 |
| 1091 | IV | 827 | 0 | 0 | 283 | 0 | 0 | Mono | Mono | | 7912180 | | | | 1091 |
| 1092 | IV | 827 | 0 | 0 | 283 | 0 | 0 | Mono | Mono | | 7912385 | | | | 1092 |
| 1093 | IV | 827 | 0 | 0 | 283 | 0 | 0 | Mono | Mono | | 7912877 | | | | 1093 |
| 1095 | IV | 827 | 0 | 0 | 283 | 0 | 0 | Mono | Mono | | 7913691 | | | | 1095 |
| 1102 | III | 825 | 0 | 2 | 187 | 96 | 0 | M9- | 22 – 36 | | 7935992 | | | | 1102 |
| 1104 | IV | 827 | 0 | 0 | 283 | 0 | 0 | Mono | Mono | | 7940644 | | | | 1104 |
| 1107 | IV | 782 | 0 | 45 | 263 | 0 | 20 | Mono | Mono | | 7948480 | | | | 1107 |
| 1112 | IV | 827 | 0 | 0 | 283 | 0 | 0 | Mono | Mono | | 7956783 | | | | 1112 |
| 1115 | IV | 827 | 0 | 0 | 283 | 0 | 0 | Mono | Mono | | 7964326 | | | | 1115 |
| 1119 | IV | 826 | 0 | 1 | 283 | 0 | 0 | Mono | Mono | | 7971707 | | | | 1119 |
| 1124 | IV | 822 | 0 | 5 | 283 | 0 | 0 | Mono | Mono | | 7992061 | | | | 1124 |
| 1132 | IV | 827 | 0 | 0 | 283 | 0 | 0 | Mono | Mono | | 8020092 | | | | 1132 |
| 1134 | IV | 827 | 0 | 0 | 283 | 0 | 0 | Mono | Mono | | 8024880 | | | | 1134 |
| 1135 | IV | 827 | 0 | 0 | 283 | 0 | 0 | Mono | Mono | | 8025051 | | | | 1135 |
| 1139 | IV | 827 | 0 | 0 | 283 | 0 | 0 | Mono | Mono | | 8033198 | | | | 1139 |
| 1142 | IV | 827 | 0 | 0 | 283 | 0 | 0 | Mono | Mono | | 8039719 | | | | 1142 |
| 1146 | IV | 827 | 0 | 0 | 283 | 0 | 0 | Mono | Mono | | 8048991 | | | | 1146 |
| 1147 | IV | 827 | 0 | 0 | 283 | 0 | 0 | Mono | Mono | | 8050318 | | | | 1147 |
| 1149 | IV | 826 | 0 | 1 | 283 | 0 | 0 | Mono | Mono | | 8052975 | | | | 1149 |

| | | M9+ | | | M9- | | | | | | Build 37 | | ISOGG | | |
|---|---|---|---|---|---|---|---|---|---|---|---|---|---|---|---|
| List | Stage | 0 | 1 | . | 0 | 1 | . | M9? | Clade(s) | Prob. | Position | Name | Subgroup Name | Amb. | List |
| 1151 | I | 827 | 0 | 0 | 246 | 37 | 0 | M9- | 1 – 10 | | 8059894 | | | | 1151 |
| 1152 | III | 826 | 1 | 0 | 282 | 0 | 1 | M9+ | 1 | | 8060903 | | | | 1152 |
| 1155 | IV | 825 | 0 | 2 | 283 | 0 | 0 | Mono | Mono | | 8070774 | | | | 1155 |
| 1159 | III | 824 | 1 | 2 | 281 | 0 | 2 | M9+ | 1 | | 8082806 | | | | 1159 |
| 1161 | IV | 0 | 827 | 0 | 0 | 283 | 0 | Mono | Mono | | 8086815 | | | | 1161 |
| 1169 | IV | 826 | 0 | 1 | 283 | 0 | 0 | Mono | Mono | | 8115897 | | | | 1169 |
| 1171 | IV | 827 | 0 | 0 | 283 | 0 | 0 | Mono | Mono | | 8117450 | | | | 1171 |
| 1182 | IV | 827 | 0 | 0 | 283 | 0 | 0 | Mono | Mono | | 8133670 | | | | 1182 |
| 1186 | IV | 827 | 0 | 0 | 283 | 0 | 0 | Mono | Mono | | 8140098 | | | | 1186 |
| 1193 | IV | 827 | 0 | 0 | 283 | 0 | 0 | Mono | Mono | | 8162893 | | | | 1193 |
| 1200 | IV | 827 | 0 | 0 | 283 | 0 | 0 | Mono | Mono | | 8191818 | | | | 1200 |
| 1204 | IV | 827 | 0 | 0 | 283 | 0 | 0 | Mono | Mono | | 8202356 | | | | 1204 |
| 1207 | IV | 827 | 0 | 0 | 283 | 0 | 0 | Mono | Mono | | 8207142 | | | | 1207 |
| 1210 | IV | 826 | 0 | 1 | 283 | 0 | 0 | Mono | Mono | | 8217320 | | | | 1210 |
| 1224 | IV | 827 | 0 | 0 | 283 | 0 | 0 | Mono | Mono | | 8235187 | | | | 1224 |
| 1234 | IV | 827 | 0 | 0 | 283 | 0 | 0 | Mono | Mono | | 8247581 | | | | 1234 |
| 1240 | I | 822 | 5 | 0 | 279 | 4 | 0 | BOTH | | PI | 8261193 | | | | 1240 |
| 1244 | III | 826 | 0 | 1 | 178 | 105 | 0 | M9- | 37 – 59 | | 8267123 | | | | 1244 |
| 1250 | I | 827 | 0 | 0 | 82 | 201 | 0 | M9- | 22 – 59 | | 8278628 | | | | 1250 |
| 1254 | IV | 827 | 0 | 0 | 283 | 0 | 0 | Mono | Mono | | 8285596 | | | | 1254 |
| 1258 | IV | 827 | 0 | 0 | 283 | 0 | 0 | Mono | Mono | | 8290465 | | | | 1258 |
| 1259 | IV | 826 | 0 | 1 | 283 | 0 | 0 | Mono | Mono | | 8291446 | | | | 1259 |
| 1264 | IV | 827 | 0 | 0 | 283 | 0 | 0 | Mono | Mono | | 8305710 | | | | 1264 |
| 1265 | IV | 827 | 0 | 0 | 283 | 0 | 0 | Mono | Mono | | 8306262 | | | | 1265 |
| 1267 | IV | 827 | 0 | 0 | 283 | 0 | 0 | Mono | Mono | | 8311795 | | | | 1267 |
| 1269 | IV | 827 | 0 | 0 | 283 | 0 | 0 | Mono | Mono | | 8315255 | | | | 1269 |
| 1278 | II | 826 | 1 | 0 | 283 | 0 | 0 | M9+ | 1 | | 8347488 | | | | 1278 |
| 1279 | IV | 827 | 0 | 0 | 283 | 0 | 0 | Mono | Mono | | 8348932 | | | | 1279 |
| 1283 | IV | 827 | 0 | 0 | 283 | 0 | 0 | Mono | Mono | | 8353524 | | | | 1283 |
| 1285 | IV | 827 | 0 | 0 | 283 | 0 | 0 | Mono | Mono | | 8360818 | | | | 1285 |
| 1289 | IV | 826 | 0 | 1 | 283 | 0 | 0 | Mono | Mono | | 8364460 | | | | 1289 |
| 1291 | IV | 827 | 0 | 0 | 283 | 0 | 0 | Mono | Mono | | 8373972 | | | | 1291 |
| 1293 | II | 826 | 1 | 0 | 283 | 0 | 0 | M9+ | 1 | | 8375595 | | | | 1293 |
| 1294 | IV | 827 | 0 | 0 | 283 | 0 | 0 | Mono | Mono | | 8380220 | | | | 1294 |
| 1295 | IV | 817 | 0 | 10 | 279 | 0 | 4 | Mono | Mono | | 8380497 | | | | 1295 |
| 1296 | IV | 827 | 0 | 0 | 283 | 0 | 0 | Mono | Mono | | 8380670 | | | | 1296 |
| 1297 | II | 826 | 1 | 0 | 283 | 0 | 0 | M9+ | 1 | | 8381017 | | | | 1297 |
| 1303 | I | 827 | 0 | 0 | 246 | 37 | 0 | M9- | 1 – 10 | | 8389216 | | | | 1303 |
| 1307 | IV | 827 | 0 | 0 | 283 | 0 | 0 | Mono | Mono | | 8392350 | | | | 1307 |
| 1312 | II | 826 | 1 | 0 | 283 | 0 | 0 | M9+ | 2 | | 8401239 | | | | 1312 |
| 1315 | II | 826 | 1 | 0 | 283 | 0 | 0 | M9+ | 1 | | 8411430 | | | | 1315 |
| 1322 | IV | 827 | 0 | 0 | 283 | 0 | 0 | Mono | Mono | | 8422356 | | | | 1322 |
| 1330 | IV | 827 | 0 | 0 | 283 | 0 | 0 | Mono | Mono | | 8428729 | | | | 1330 |
| 1332 | IV | 827 | 0 | 0 | 283 | 0 | 0 | Mono | Mono | | 8429649 | | | | 1332 |
| 1335 | IV | 827 | 0 | 0 | 283 | 0 | 0 | Mono | Mono | | 8431604 | | | | 1335 |
| 1337 | IV | 827 | 0 | 0 | 283 | 0 | 0 | Mono | Mono | | 8432095 | | | | 1337 |
| 1344 | IV | 827 | 0 | 0 | 283 | 0 | 0 | Mono | Mono | | 8442334 | | | | 1344 |
| 1357 | IV | 826 | 0 | 1 | 283 | 0 | 0 | Mono | Mono | | 8457361 | | | | 1357 |
| 1361 | IV | 827 | 0 | 0 | 283 | 0 | 0 | Mono | Mono | | 8461895 | | | | 1361 |
| 1380 | III | 820 | 1 | 6 | 281 | 0 | 2 | M9+ | 1 | | 8475654 | | | | 1380 |
| 1381 | IV | 817 | 0 | 10 | 279 | 0 | 4 | Mono | Mono | | 8475689 | | | | 1381 |
| 1386 | III | 782 | 4 | 41 | 249 | 22 | 12 | BOTH | | PI | 8481036 | | | | 1386 |
| 1387 | IV | 827 | 0 | 0 | 283 | 0 | 0 | Mono | Mono | | 8481920 | | | | 1387 |
| 1388 | IV | 818 | 0 | 9 | 281 | 0 | 2 | Mono | Mono | | 8482010 | | | | 1388 |
| 1403 | IV | 827 | 0 | 0 | 283 | 0 | 0 | Mono | Mono | | 8490706 | | | | 1403 |
| 1407 | IV | 827 | 0 | 0 | 283 | 0 | 0 | Mono | Mono | | 8497599 | | | | 1407 |
| 1411 | IV | 826 | 0 | 1 | 283 | 0 | 0 | Mono | Mono | | 8501357 | | | | 1411 |
| 1412 | IV | 827 | 0 | 0 | 283 | 0 | 0 | Mono | Mono | | 8501449 | | | | 1412 |
| 1415 | IV | 827 | 0 | 0 | 283 | 0 | 0 | Mono | Mono | | 8504927 | | | | 1415 |
| 1417 | IV | 827 | 0 | 0 | 282 | 0 | 1 | Mono | Mono | | 8506663 | | | | 1417 |
| 1419 | IV | 827 | 0 | 0 | 283 | 0 | 0 | Mono | Mono | | 8509737 | | | | 1419 |
| 1422 | IV | 827 | 0 | 0 | 283 | 0 | 0 | Mono | Mono | | 8513390 | | | | 1422 |
| 1429 | IV | 827 | 0 | 0 | 282 | 0 | 1 | Mono | Mono | | 8530679 | | | | 1429 |
| 1438 | IV | 827 | 0 | 0 | 283 | 0 | 0 | Mono | Mono | | 8542774 | | | | 1438 |
| 1446 | III | 821 | 1 | 5 | 281 | 0 | 2 | M9+ | 1 | | 8556580 | | | | 1446 |
| 1456 | IV | 827 | 0 | 0 | 283 | 0 | 0 | Mono | Mono | | 8568831 | | | | 1456 |
| 1460 | IV | 827 | 0 | 0 | 283 | 0 | 0 | Mono | Mono | | 8574958 | | | | 1460 |
| 1467 | IV | 824 | 0 | 3 | 283 | 0 | 0 | Mono | Mono | | 8581659 | | | | 1467 |
| 1484 | IV | 827 | 0 | 0 | 283 | 0 | 0 | Mono | Mono | | 8602518 | | | | 1484 |
| 1500 | IV | 827 | 0 | 0 | 283 | 0 | 0 | Mono | Mono | | 8620877 | | | | 1500 |
| 1502 | IV | 827 | 0 | 0 | 283 | 0 | 0 | Mono | Mono | | 8622207 | | | | 1502 |
| 1513 | IV | 827 | 0 | 0 | 283 | 0 | 0 | Mono | Mono | | 8640367 | | | | 1513 |
| 1516 | IV | 827 | 0 | 0 | 283 | 0 | 0 | Mono | Mono | | 8642055 | | | | 1516 |
| 1522 | IV | 827 | 0 | 0 | 283 | 0 | 0 | Mono | Mono | | 8649167 | | | | 1522 |
| 1523 | I | 827 | 0 | 0 | 246 | 37 | 0 | M9- | 1 – 10 | | 8660432 | | | | 1523 |
| 1529 | IV | 827 | 0 | 0 | 282 | 0 | 1 | Mono | Mono | | 8675410 | | | | 1529 |
| 1530 | IV | 827 | 0 | 0 | 283 | 0 | 0 | Mono | Mono | | 8675467 | | | | 1530 |
| 1539 | IV | 826 | 0 | 1 | 283 | 0 | 0 | Mono | Mono | | 8685230 | | | | 1539 |
| 1545 | IV | 0 | 827 | 0 | 0 | 283 | 0 | Mono | Mono | | 8694916 | | | | 1545 |
| 1547 | IV | 826 | 0 | 1 | 283 | 0 | 0 | Mono | Mono | | 8698064 | | | | 1547 |
| 1552 | IV | 827 | 0 | 0 | 283 | 0 | 0 | Mono | Mono | | 8709342 | | | | 1552 |
| 1555 | IV | 827 | 0 | 0 | 283 | 0 | 0 | Mono | Mono | | 8714022 | | | | 1555 |
| 1558 | IV | 827 | 0 | 0 | 283 | 0 | 0 | Mono | Mono | | 8722850 | | | | 1558 |
| 1562 | IV | 827 | 0 | 0 | 283 | 0 | 0 | Mono | Mono | | 8738037 | | | | 1562 |
| 1563 | II | 826 | 1 | 0 | 283 | 0 | 0 | M9+ | NU | | 8738392 | | | | 1563 |
| 1564 | IV | 827 | 0 | 0 | 283 | 0 | 0 | Mono | Mono | | 8745230 | | | | 1564 |
| 1566 | IV | 827 | 0 | 0 | 283 | 0 | 0 | Mono | Mono | | 8758629 | | | | 1566 |
| 1567 | IV | 827 | 0 | 0 | 283 | 0 | 0 | Mono | Mono | | 8758632 | | | | 1567 |
| 1572 | IV | 827 | 0 | 0 | 283 | 0 | 0 | Mono | Mono | | 8781088 | | | | 1572 |
| 1574 | III | 823 | 1 | 3 | 282 | 0 | 1 | M9+ | 1 | | 8783363 | | | | 1574 |
| 1580 | III | 826 | 0 | 1 | 261 | 21 | 1 | M9- | 11 – 20 | | 8801406 | | | | 1580 |
| 1583 | IV | 826 | 0 | 1 | 283 | 0 | 0 | Mono | Mono | | 8805000 | | | | 1583 |
| 1584 | IV | 0 | 827 | 0 | 0 | 282 | 1 | Mono | Mono | | 8805525 | | | | 1584 |
| 1592 | IV | 826 | 0 | 1 | 282 | 0 | 1 | Mono | Mono | | 8838729 | | | | 1592 |
| 1598 | IV | 827 | 0 | 0 | 283 | 0 | 0 | Mono | Mono | | 8845759 | | | | 1598 |
| 1599 | IV | 827 | 0 | 0 | 283 | 0 | 0 | Mono | Mono | | 8846370 | | | | 1599 |

| | | M9+ | | | M9- | | | | | | Build 37 | | ISOGG | | |
|---|---|---|---|---|---|---|---|---|---|---|---|---|---|---|---|
| List | Stage | 0 | 1 | . | 0 | 1 | . | M9? | Clade(s) | Prob. | Position | Name | Subgroup Name | Amb. | List |
| 1605 | IV | 827 | 0 | 0 | 283 | 0 | 0 | Mono | Mono | | 8869608 | | | | 1605 |
| 1608 | IV | 827 | 0 | 0 | 283 | 0 | 0 | Mono | Mono | | 8873548 | | | | 1608 |
| 1612 | IV | 827 | 0 | 0 | 283 | 0 | 0 | Mono | Mono | | 8882497 | | | | 1612 |
| 1613 | I | 827 | 0 | 0 | 209 | 74 | 0 | M9- | 42 – 58 | | 8885865 | | | | 1613 |
| 1615 | IV | 827 | 0 | 0 | 283 | 0 | 0 | Mono | Mono | | 8890588 | | | | 1615 |
| 1618 | IV | 826 | 0 | 1 | 282 | 0 | 1 | Mono | Mono | | 8898284 | | | | 1618 |
| 1619 | IV | 827 | 0 | 0 | 283 | 0 | 0 | Mono | Mono | | 8899540 | | | | 1619 |
| 1621 | IV | 827 | 0 | 0 | 282 | 0 | 1 | Mono | Mono | | 8906097 | | | | 1621 |
| 1625 | II | 826 | 1 | 0 | 283 | 0 | 0 | M9+ | 2 | | 8910408 | | | | 1625 |
| 1626 | IV | 827 | 0 | 0 | 283 | 0 | 0 | Mono | Mono | | 8967006 | | | | 1626 |
| 1627 | IV | 827 | 0 | 0 | 283 | 0 | 0 | Mono | Mono | | 8986136 | | | | 1627 |
| 1628 | I | 827 | 0 | 0 | 279 | 4 | 0 | M9- | 9 – 10 | | 8990496 | | | | 1628 |
| 1629 | IV | 827 | 0 | 0 | 283 | 0 | 0 | Mono | Mono | | 8996352 | | | | 1629 |
| 1631 | II | 825 | 2 | 0 | 283 | 0 | 0 | M9+ | 1 – 2 | | 9004556 | | | | 1631 |
| 1632 | IV | 826 | 0 | 1 | 283 | 0 | 0 | Mono | Mono | | 9011473 | | | | 1632 |
| 1633 | IV | 827 | 0 | 0 | 283 | 0 | 0 | Mono | Mono | | 9021351 | | | | 1633 |
| 1634 | IV | 826 | 0 | 1 | 283 | 0 | 0 | Mono | Mono | | 9030715 | | | | 1634 |
| 1635 | IV | 825 | 0 | 2 | 283 | 0 | 0 | Mono | Mono | | 9030752 | | | | 1635 |
| 1636 | III | 815 | 1 | 11 | 278 | 0 | 5 | M9+ | 2 | | 9030938 | | | | 1636 |
| 1637 | IV | 827 | 0 | 0 | 283 | 0 | 0 | Mono | Mono | | 9044535 | | | | 1637 |
| 1638 | II | 827 | 0 | 0 | 282 | 1 | 0 | M9- | 21 | | 9055812 | | | | 1638 |
| 1639 | IV | 826 | 0 | 1 | 283 | 0 | 0 | Mono | Mono | | 9055962 | | | | 1639 |
| 1640 | IV | 827 | 0 | 0 | 283 | 0 | 0 | Mono | Mono | | 9060781 | | | | 1640 |
| 1641 | III | 826 | 0 | 1 | 246 | 37 | 0 | M9- | 1 – 10 | | 9064682 | | | | 1641 |
| 1643 | I | 827 | 0 | 0 | 247 | 36 | 0 | M9- | 2 – 10 | | 9083940 | | | | 1643 |
| 1644 | IV | 827 | 0 | 0 | 283 | 0 | 0 | Mono | Mono | | 9084849 | | | | 1644 |
| 1646 | I | 827 | 0 | 0 | 247 | 36 | 0 | M9- | 2 – 10 | | 9091871 | | | | 1646 |
| 1648 | IV | 827 | 0 | 0 | 283 | 0 | 0 | Mono | Mono | | 9098400 | | | | 1648 |
| 1652 | IV | 827 | 0 | 0 | 283 | 0 | 0 | Mono | Mono | | 9107281 | | | | 1652 |
| 1654 | III | 826 | 0 | 1 | 262 | 18 | 3 | M9- | 11 – 20 | MD | 9113774 | | | | 1654 |
| 1655 | I | 827 | 0 | 0 | 274 | 9 | 0 | M9- | 61 – 64 | | 9118058 | | | | 1655 |
| 1657 | IV | 818 | 0 | 9 | 277 | 0 | 6 | Mono | Mono | | 9119893 | | | | 1657 |
| 1659 | IV | 827 | 0 | 0 | 283 | 0 | 0 | Mono | Mono | | 9127755 | | | | 1659 |
| 1663 | III | 824 | 0 | 3 | 261 | 18 | 4 | M9- | 11 – 20 | MD | 9136546 | | | | 1663 |
| 1664 | IV | 826 | 0 | 1 | 283 | 0 | 0 | Mono | Mono | | 9142205 | | | | 1664 |
| 1665 | III | 791 | 32 | 4 | 187 | 96 | 0 | BOTH | | PI | 9142914 | | | | 1665 |
| 1668 | IV | 821 | 0 | 6 | 281 | 0 | 2 | Mono | Mono | | 9155266 | | | | 1668 |
| 1670 | IV | 827 | 0 | 0 | 283 | 0 | 0 | Mono | Mono | | 9161225 | | | | 1670 |
| 1671 | II | 826 | 1 | 0 | 283 | 0 | 0 | M9+ | 1 | | 9161969 | | | | 1671 |
| 1673 | IV | 827 | 0 | 0 | 283 | 0 | 0 | Mono | Mono | | 9171639 | | | | 1673 |
| 1674 | IV | 0 | 827 | 0 | 0 | 283 | 0 | Mono | Mono | | 9172332 | | | | 1674 |
| 1675 | IV | 827 | 0 | 0 | 283 | 0 | 0 | Mono | Mono | | 9193040 | | | | 1675 |
| 1676 | IV | 827 | 0 | 0 | 283 | 0 | 0 | Mono | Mono | | 9354054 | | | | 1676 |
| 1679 | III | 825 | 1 | 1 | 283 | 0 | 0 | M9+ | NU | | 9384122 | | | | 1679 |
| 1686 | IV | 827 | 0 | 0 | 283 | 0 | 0 | Mono | Mono | | 9417393 | | | | 1686 |
| 1688 | IV | 827 | 0 | 0 | 283 | 0 | 0 | Mono | Mono | | 9424766 | | | | 1688 |
| 1694 | II | 826 | 1 | 0 | 283 | 0 | 0 | M9+ | 1 | | 9460359 | | | | 1694 |
| 1695 | IV | 827 | 0 | 0 | 283 | 0 | 0 | Mono | Mono | | 9463712 | | | | 1695 |
| 1696 | IV | 827 | 0 | 0 | 283 | 0 | 0 | Mono | Mono | | 9466878 | | | | 1696 |
| 1697 | I | 827 | 0 | 0 | 187 | 96 | 0 | M9- | 22 – 36 | | 9477753 | | | | 1697 |
| 1698 | IV | 827 | 0 | 0 | 283 | 0 | 0 | Mono | Mono | | 9480167 | | | | 1698 |
| 1699 | IV | 827 | 0 | 0 | 282 | 0 | 1 | Mono | Mono | | 9496791 | | | | 1699 |
| 1700 | IV | 826 | 0 | 1 | 283 | 0 | 0 | Mono | Mono | | 9499034 | | | | 1700 |
| 1701 | III | 826 | 0 | 1 | 282 | 1 | 0 | M9- | NU | | 9500347 | | | | 1701 |
| 1702 | IV | 824 | 0 | 3 | 281 | 0 | 2 | Mono | Mono | | 9507122 | | | | 1702 |
| 1703 | IV | 827 | 0 | 0 | 282 | 0 | 1 | Mono | Mono | | 9507451 | | | | 1703 |
| 1704 | I | 827 | 0 | 0 | 83 | 200 | 0 | M9- | | PI | 9516653 | | | | 1704 |
| 1705 | I | 827 | 0 | 0 | 270 | 13 | 0 | M9- | 65 – 66 | | 9520802 | | | | 1705 |
| 1706 | IV | 827 | 0 | 0 | 282 | 0 | 1 | Mono | Mono | | 9523217 | | | | 1706 |
| 1708 | IV | 825 | 0 | 2 | 283 | 0 | 0 | Mono | Mono | | 9524762 | | | | 1708 |
| 1709 | IV | 826 | 0 | 1 | 282 | 0 | 1 | Mono | Mono | | 9642917 | | | | 1709 |
| 1710 | I | 790 | 37 | 0 | 0 | 283 | 0 | M9+ | | PI | 9646980 | | | | 1710 |
| 1711 | IV | 827 | 0 | 0 | 283 | 0 | 0 | Mono | Mono | | 9740811 | | | | 1711 |
| 1712 | IV | 827 | 0 | 0 | 282 | 0 | 1 | Mono | Mono | | 9741848 | | | | 1712 |
| 1714 | III | 824 | 1 | 2 | 282 | 0 | 1 | M9+ | 1 | | 9760575 | | | | 1714 |
| 1715 | II | 826 | 1 | 0 | 283 | 0 | 0 | M9+ | 1 | | 9760879 | | | | 1715 |
| 1717 | IV | 827 | 0 | 0 | 283 | 0 | 0 | Mono | Mono | | 9764409 | | | | 1717 |
| 1718 | IV | 827 | 0 | 0 | 283 | 0 | 0 | Mono | Mono | | 9764623 | | | | 1718 |
| 1727 | IV | 827 | 0 | 0 | 283 | 0 | 0 | Mono | Mono | | 9794806 | | | | 1727 |
| 1728 | IV | 825 | 0 | 2 | 283 | 0 | 0 | Mono | Mono | | 9795076 | | | | 1728 |
| 1730 | IV | 827 | 0 | 0 | 283 | 0 | 0 | Mono | Mono | | 9807098 | | | | 1730 |
| 1734 | IV | 827 | 0 | 0 | 283 | 0 | 0 | Mono | Mono | | 9819120 | | | | 1734 |
| 1735 | IV | 827 | 0 | 0 | 283 | 0 | 0 | Mono | Mono | | 9819205 | | | | 1735 |
| 1736 | IV | 827 | 0 | 0 | 283 | 0 | 0 | Mono | Mono | | 9819254 | | | | 1736 |
| 1737 | IV | 825 | 0 | 2 | 282 | 0 | 1 | Mono | Mono | | 9819301 | | | | 1737 |
| 1742 | IV | 806 | 0 | 21 | 277 | 0 | 6 | Mono | Mono | | 9831559 | | | | 1742 |
| 1745 | IV | 827 | 0 | 0 | 283 | 0 | 0 | Mono | Mono | | 9833726 | | | | 1745 |
| 1746 | IV | 827 | 0 | 0 | 283 | 0 | 0 | Mono | Mono | | 9836911 | | | | 1746 |
| 1750 | IV | 827 | 0 | 0 | 283 | 0 | 0 | Mono | Mono | | 9842606 | | | | 1750 |
| 1754 | III | 818 | 7 | 2 | 257 | 24 | 2 | BOTH | | PI | 9850653 | | | | 1754 |
| 1755 | IV | 824 | 0 | 3 | 283 | 0 | 0 | Mono | Mono | | 9850700 | | | | 1755 |
| 1761 | IV | 826 | 0 | 1 | 283 | 0 | 0 | Mono | Mono | | 9854154 | | | | 1761 |
| 1763 | IV | 825 | 0 | 2 | 282 | 0 | 1 | Mono | Mono | | 9869534 | | | | 1763 |
| 1768 | IV | 827 | 0 | 0 | 283 | 0 | 0 | Mono | Mono | | 9883072 | | | | 1768 |
| 1776 | IV | 0 | 825 | 2 | 0 | 283 | 0 | Mono | Mono | | 9908071 | | | | 1776 |
| 1777 | IV | 827 | 0 | 0 | 283 | 0 | 0 | Mono | Mono | | 9932720 | | | | 1777 |
| 1778 | IV | 827 | 0 | 0 | 283 | 0 | 0 | Mono | Mono | | 9936979 | | | | 1778 |
| 1779 | IV | 827 | 0 | 0 | 283 | 0 | 0 | Mono | Mono | | 9937130 | | | | 1779 |
| 1780 | IV | 730 | 0 | 97 | 249 | 0 | 34 | Mono | Mono | | 9937387 | | | | 1780 |
| 1781 | I | 827 | 0 | 0 | 246 | 37 | 0 | M9- | 1 – 10 | | 9940615 | | | | 1781 |
| 1782 | IV | 827 | 0 | 0 | 283 | 0 | 0 | Mono | Mono | | 9942979 | | | | 1782 |
| 1783 | IV | 827 | 0 | 0 | 283 | 0 | 0 | Mono | Mono | | 9953885 | | | | 1783 |
| 1784 | IV | 827 | 0 | 0 | 283 | 0 | 0 | Mono | Mono | | 9956566 | | | | 1784 |
| 1785 | II | 826 | 1 | 0 | 283 | 0 | 0 | M9+ | 1 | | 9960679 | | | | 1785 |
| 1786 | IV | 827 | 0 | 0 | 283 | 0 | 0 | Mono | Mono | | 9963448 | | | | 1786 |
| 1787 | IV | 827 | 0 | 0 | 283 | 0 | 0 | Mono | Mono | | 9963477 | | | | 1787 |

| | | | M9+ | | | | M9- | | | | | Build 37 | | ISOGG | | |
|---|---|---|---|---|---|---|---|---|---|---|---|---|---|---|---|---|
| List | Stage | 0 | 1 | . | | 0 | 1 | . | M9? | Clade(s) | Prob. | Position | Name | Subgroup Name | Amb. | List |
| 1788 | IV | 827 | 0 | 0 | | 283 | 0 | 0 | Mono | Mono | | 9968357 | | | | 1788 |
| 1789 | IV | 827 | 0 | 0 | | 283 | 0 | 0 | Mono | Mono | | 9972115 | | | | 1789 |
| 1790 | IV | 825 | 0 | 2 | | 283 | 0 | 0 | Mono | Mono | | 9978271 | | | | 1790 |
| 1791 | IV | 827 | 0 | 0 | | 283 | 0 | 0 | Mono | Mono | | 9980841 | | | | 1791 |
| 1792 | IV | 826 | 0 | 1 | | 283 | 0 | 0 | Mono | Mono | | 9982280 | | | | 1792 |
| 1793 | IV | 827 | 0 | 0 | | 283 | 0 | 0 | Mono | Mono | | 10003838 | | | | 1793 |
| 1794 | IV | 827 | 0 | 0 | | 283 | 0 | 0 | Mono | Mono | | 10004228 | | | | 1794 |
| 1795 | IV | 826 | 0 | 1 | | 283 | 0 | 0 | Mono | Mono | | 10007881 | | | | 1795 |
| 1796 | IV | 827 | 0 | 0 | | 283 | 0 | 0 | Mono | Mono | | 10007932 | | | | 1796 |
| 1797 | IV | 826 | 0 | 1 | | 283 | 0 | 0 | Mono | Mono | | 10009621 | | | | 1797 |
| 1798 | IV | 827 | 0 | 0 | | 283 | 0 | 0 | Mono | Mono | | 10011231 | | | | 1798 |
| 1799 | IV | 826 | 0 | 1 | | 283 | 0 | 0 | Mono | Mono | | 10012273 | | | | 1799 |
| 1800 | IV | 796 | 0 | 31 | | 269 | 0 | 14 | Mono | Mono | | 10019072 | | | | 1800 |
| 1801 | III | 826 | 0 | 1 | | 247 | 36 | 0 | M9- | 2 – 10 | | 10019293 | | | | 1801 |
| 1802 | III | 818 | 1 | 8 | | 269 | 11 | 3 | BOTH | | PI | 10020053 | | | | 1802 |
| 1803 | IV | 825 | 0 | 2 | | 282 | 0 | 1 | Mono | Mono | | 10021948 | | | | 1803 |
| 1804 | IV | 827 | 0 | 0 | | 283 | 0 | 0 | Mono | Mono | | 10022636 | | | | 1804 |
| 1805 | IV | 827 | 0 | 0 | | 283 | 0 | 0 | Mono | Mono | | 10022705 | | | | 1805 |
| 1806 | IV | 826 | 0 | 1 | | 283 | 0 | 0 | Mono | Mono | | 10023120 | | | | 1806 |
| 1807 | IV | 817 | 0 | 10 | | 280 | 0 | 3 | Mono | Mono | | 10023410 | | | | 1807 |
| 1808 | IV | 826 | 0 | 1 | | 283 | 0 | 0 | Mono | Mono | | 10024996 | | | | 1808 |
| 1809 | IV | 827 | 0 | 0 | | 283 | 0 | 0 | Mono | Mono | | 10026900 | | | | 1809 |
| 1810 | IV | 827 | 0 | 0 | | 282 | 0 | 1 | Mono | Mono | | 10027873 | | | | 1810 |
| 1811 | II | 826 | 1 | 0 | | 283 | 0 | 0 | M9+ | 1 | | 10028676 | | | | 1811 |
| 1812 | IV | 827 | 0 | 0 | | 283 | 0 | 0 | Mono | Mono | | 10028786 | | | | 1812 |
| 1813 | IV | 827 | 0 | 0 | | 283 | 0 | 0 | Mono | Mono | | 10029396 | | | | 1813 |
| 1814 | IV | 827 | 0 | 0 | | 283 | 0 | 0 | Mono | Mono | | 10029690 | | | | 1814 |
| 1815 | IV | 827 | 0 | 0 | | 283 | 0 | 0 | Mono | Mono | | 10029711 | | | | 1815 |
| 1816 | IV | 827 | 0 | 0 | | 283 | 0 | 0 | Mono | Mono | | 10029759 | | | | 1816 |
| 1817 | IV | 826 | 0 | 1 | | 283 | 0 | 0 | Mono | Mono | | 10034855 | | | | 1817 |
| 1818 | IV | 827 | 0 | 0 | | 283 | 0 | 0 | Mono | Mono | | 10035382 | | | | 1818 |
| 1822 | IV | 827 | 0 | 0 | | 283 | 0 | 0 | Mono | Mono | | 10047932 | | | | 1822 |
| 1824 | IV | 811 | 0 | 16 | | 278 | 0 | 5 | Mono | Mono | | 10059544 | | | | 1824 |
| 1827 | IV | 827 | 0 | 0 | | 283 | 0 | 0 | Mono | Mono | | 13133499 | | | | 1827 |
| 1828 | III | 827 | 0 | 0 | | 245 | 36 | 2 | M9- | 2 – 10 | | 13195605 | | | | 1828 |
| 1830 | IV | 827 | 0 | 0 | | 281 | 0 | 2 | Mono | Mono | | 13212571 | | | | 1830 |
| 1831 | IV | 826 | 0 | 1 | | 281 | 0 | 2 | Mono | Mono | | 13220827 | | | | 1831 |
| 1834 | IV | 827 | 0 | 0 | | 281 | 0 | 2 | Mono | Mono | | 13234104 | | | | 1834 |
| 1835 | III | 3 | 824 | 0 | | 281 | 0 | 2 | M9+ | 3 – 93 | | 13234865 | | | | 1835 |
| 1838 | IV | 826 | 0 | 1 | | 281 | 0 | 2 | Mono | Mono | | 13259989 | | | | 1838 |
| 1839 | IV | 827 | 0 | 0 | | 281 | 0 | 2 | Mono | Mono | | 13420650 | | | | 1839 |
| 1840 | III | 821 | 0 | 6 | | 241 | 37 | 5 | M9- | 1 – 10 | | 13424256 | | | | 1840 |
| 1841 | IV | 827 | 0 | 0 | | 281 | 0 | 2 | Mono | Mono | | 13434514 | | | | 1841 |
| 1842 | IV | 827 | 0 | 0 | | 280 | 0 | 3 | Mono | Mono | | 13457272 | | | | 1842 |
| 1843 | IV | 825 | 0 | 2 | | 281 | 0 | 2 | Mono | Mono | | 13457275 | | | | 1843 |
| 1844 | IV | 827 | 0 | 0 | | 281 | 0 | 2 | Mono | Mono | | 13457347 | | | | 1844 |
| 1845 | IV | 827 | 0 | 0 | | 281 | 0 | 2 | Mono | Mono | | 13464820 | | | | 1845 |
| 1846 | IV | 826 | 0 | 1 | | 280 | 0 | 3 | Mono | Mono | | 13472852 | | | | 1846 |
| 1847 | IV | 825 | 0 | 2 | | 281 | 0 | 2 | Mono | Mono | | 13481779 | | | | 1847 |
| 1849 | IV | 823 | 0 | 4 | | 280 | 0 | 3 | Mono | Mono | | 13527667 | | | | 1849 |
| 1850 | IV | 825 | 0 | 2 | | 281 | 0 | 2 | Mono | Mono | | 13531656 | | | | 1850 |
| 1851 | IV | 820 | 0 | 7 | | 279 | 0 | 4 | Mono | Mono | | 13540419 | | | | 1851 |
| 1852 | III | 826 | 0 | 1 | | 243 | 37 | 3 | M9- | 1 – 10 | | 13559017 | | | | 1852 |
| 1853 | IV | 827 | 0 | 0 | | 281 | 0 | 2 | Mono | Mono | | 13650039 | | | | 1853 |
| 1854 | IV | 827 | 0 | 0 | | 281 | 0 | 2 | Mono | Mono | | 13650841 | | | | 1854 |
| 1855 | IV | 827 | 0 | 0 | | 281 | 0 | 2 | Mono | Mono | | 13652289 | | | | 1855 |
| 1857 | IV | 827 | 0 | 0 | | 281 | 0 | 2 | Mono | Mono | | 13656512 | | | | 1857 |
| 1858 | IV | 826 | 0 | 1 | | 281 | 0 | 2 | Mono | Mono | | 13659237 | | | | 1858 |
| 1859 | IV | 823 | 0 | 4 | | 281 | 0 | 2 | Mono | Mono | | 13659442 | | | | 1859 |
| 1860 | IV | 825 | 0 | 2 | | 279 | 0 | 4 | Mono | Mono | | 13659588 | | | | 1860 |
| 1861 | IV | 819 | 0 | 8 | | 279 | 0 | 4 | Mono | Mono | | 13660871 | | | | 1861 |
| 1862 | IV | 827 | 0 | 0 | | 281 | 0 | 2 | Mono | Mono | | 13662451 | | | | 1862 |
| 1863 | IV | 826 | 0 | 1 | | 281 | 0 | 2 | Mono | Mono | | 13666052 | | | | 1863 |
| 1866 | IV | 827 | 0 | 0 | | 281 | 0 | 2 | Mono | Mono | | 13668684 | | | | 1866 |
| 1867 | IV | 826 | 0 | 1 | | 281 | 0 | 2 | Mono | Mono | | 13668686 | | | | 1867 |
| 1868 | IV | 826 | 0 | 1 | | 281 | 0 | 2 | Mono | Mono | | 13668714 | | | | 1868 |
| 1870 | IV | 827 | 0 | 0 | | 281 | 0 | 2 | Mono | Mono | | 13669387 | | | | 1870 |
| 1871 | IV | 827 | 0 | 0 | | 281 | 0 | 2 | Mono | Mono | | 13669390 | | | | 1871 |
| 1873 | IV | 827 | 0 | 0 | | 281 | 0 | 2 | Mono | Mono | | 13676540 | | | | 1873 |
| 1874 | IV | 822 | 0 | 5 | | 281 | 0 | 2 | Mono | Mono | | 13678931 | | | | 1874 |
| 1877 | IV | 826 | 0 | 1 | | 280 | 0 | 3 | Mono | Mono | | 13684735 | | | | 1877 |
| 1878 | IV | 807 | 0 | 20 | | 244 | 0 | 39 | Mono | Mono | | 13686953 | | | | 1878 |
| 1879 | IV | 0 | 827 | 0 | | 0 | 283 | 0 | Mono | Mono | | 13686963 | | | | 1879 |
| 1880 | IV | 0 | 827 | 0 | | 0 | 283 | 0 | Mono | Mono | | 13687112 | | | | 1880 |
| 1881 | IV | 0 | 827 | 0 | | 0 | 283 | 0 | Mono | Mono | | 13687378 | | | | 1881 |
| 1882 | IV | 0 | 827 | 0 | | 0 | 283 | 0 | Mono | Mono | | 13687415 | | | | 1882 |
| 1883 | IV | 0 | 827 | 0 | | 0 | 283 | 0 | Mono | Mono | | 13687697 | | | | 1883 |
| 1884 | IV | 0 | 826 | 1 | | 0 | 283 | 0 | Mono | Mono | | 13687740 | | | | 1884 |
| 1885 | IV | 0 | 827 | 0 | | 0 | 283 | 0 | Mono | Mono | | 13687747 | | | | 1885 |
| 1886 | IV | 0 | 820 | 7 | | 0 | 279 | 4 | Mono | Mono | | 13689634 | | | | 1886 |
| 1887 | IV | 0 | 827 | 0 | | 0 | 282 | 1 | Mono | Mono | | 13692231 | | | | 1887 |
| 1888 | IV | 0 | 431 | 396 | | 0 | 176 | 107 | Mono | Mono | | 13692590 | | | | 1888 |
| 1889 | IV | 0 | 827 | 0 | | 0 | 283 | 0 | Mono | Mono | | 13693205 | | | | 1889 |
| 1890 | IV | 827 | 0 | 0 | | 281 | 0 | 2 | Mono | Mono | | 13693534 | | | | 1890 |
| 1891 | IV | 827 | 0 | 0 | | 281 | 0 | 2 | Mono | Mono | | 13693698 | | | | 1891 |
| 1892 | IV | 0 | 766 | 61 | | 0 | 274 | 9 | Mono | Mono | | 13693802 | | | | 1892 |
| 1893 | IV | 0 | 827 | 0 | | 0 | 281 | 2 | Mono | Mono | | 13694929 | | | | 1893 |
| 1894 | IV | 0 | 809 | 18 | | 0 | 274 | 9 | Mono | Mono | | 13695208 | | | | 1894 |
| 1895 | IV | 825 | 0 | 2 | | 278 | 0 | 5 | Mono | Mono | | 13695249 | | | | 1895 |
| 1896 | IV | 0 | 826 | 1 | | 0 | 281 | 2 | Mono | Mono | | 13695298 | | | | 1896 |
| 1897 | IV | 0 | 827 | 0 | | 0 | 281 | 2 | Mono | Mono | | 13707656 | | | | 1897 |
| 1898 | III | 827 | 0 | 0 | | 281 | 1 | 1 | M9- | NU | | 13711929 | | | | 1898 |
| 1899 | IV | 827 | 0 | 0 | | 281 | 0 | 2 | Mono | Mono | | 13713684 | | | | 1899 |
| 1900 | IV | 0 | 827 | 0 | | 0 | 281 | 2 | Mono | Mono | | 13716757 | | | | 1900 |
| 1901 | IV | 0 | 827 | 0 | | 0 | 280 | 3 | Mono | Mono | | 13717500 | | | | 1901 |
| 1902 | IV | 827 | 0 | 0 | | 283 | 0 | 0 | Mono | Mono | | 13799151 | | | | 1902 |

| | | M9+ | | | M9- | | | | | | Build 37 | | ISOGG | | |
|---|---|---|---|---|---|---|---|---|---|---|---|---|---|---|---|
| List | Stage | 0 | 1 | . | 0 | 1 | . | M9? | Clade(s) | Prob. | Position | Name | Subgroup Name | Amb. | List |
| 1903 | IV | 827 | 0 | 0 | 283 | 0 | 0 | Mono | Mono | | 13799872 | | | | 1903 |
| 1904 | IV | 827 | 0 | 0 | 283 | 0 | 0 | Mono | Mono | | 13801064 | | | | 1904 |
| 1906 | IV | 823 | 0 | 4 | 282 | 0 | 1 | Mono | Mono | | 13802287 | | | | 1906 |
| 1907 | IV | 826 | 0 | 1 | 283 | 0 | 0 | Mono | Mono | | 13802302 | | | | 1907 |
| 1909 | IV | 827 | 0 | 0 | 283 | 0 | 0 | Mono | Mono | | 13805274 | | | | 1909 |
| 1911 | IV | 827 | 0 | 0 | 283 | 0 | 0 | Mono | Mono | | 13808780 | | | | 1911 |
| 1915 | IV | 827 | 0 | 0 | 283 | 0 | 0 | Mono | Mono | | 13824097 | | | | 1915 |
| 1922 | IV | 826 | 0 | 1 | 283 | 0 | 0 | Mono | Mono | | 13855597 | | | | 1922 |
| 1923 | I | 823 | 4 | 0 | 0 | 283 | 0 | M9+ | 4 – 93 | | 13857331 | | | | 1923 |
| 1924 | IV | 827 | 0 | 0 | 283 | 0 | 0 | Mono | Mono | | 13858215 | | | | 1924 |
| 1925 | IV | 825 | 0 | 2 | 281 | 0 | 2 | Mono | Mono | | 13862569 | | | | 1925 |
| 1927 | IV | 827 | 0 | 0 | 283 | 0 | 0 | Mono | Mono | | 13864547 | | | | 1927 |
| 1928 | IV | 789 | 0 | 38 | 271 | 0 | 12 | Mono | Mono | | 13865385 | | | | 1928 |
| 1930 | IV | 774 | 0 | 53 | 268 | 0 | 15 | Mono | Mono | | 13871209 | | | | 1930 |
| 1931 | IV | 827 | 0 | 0 | 281 | 0 | 2 | Mono | Mono | | 13874161 | | | | 1931 |
| 1932 | IV | 827 | 0 | 0 | 283 | 0 | 0 | Mono | Mono | | 13880411 | | | | 1932 |
| 1935 | IV | 824 | 0 | 3 | 283 | 0 | 0 | Mono | Mono | | 13894235 | | | | 1935 |
| 1936 | III | 825 | 0 | 2 | 0 | 282 | 1 | M9+/- | M9+/- | | 13907799 | | | | 1936 |
| 1937 | III | 716 | 97 | 14 | 248 | 33 | 2 | BOTH | | PI | 13921953 | | | | 1937 |
| 1938 | IV | 827 | 0 | 0 | 283 | 0 | 0 | Mono | Mono | | 13924059 | | | | 1938 |
| 1939 | IV | 827 | 0 | 0 | 283 | 0 | 0 | Mono | Mono | | 13945848 | | | | 1939 |
| 1940 | IV | 827 | 0 | 0 | 283 | 0 | 0 | Mono | Mono | | 13955033 | | | | 1940 |
| 1941 | I | 827 | 0 | 0 | 82 | 201 | 0 | M9- | 22 – 59 | | 13961890 | | | | 1941 |
| 1942 | IV | 825 | 0 | 2 | 283 | 0 | 0 | Mono | Mono | | 13966891 | | | | 1942 |
| 1943 | III | 826 | 0 | 1 | 272 | 11 | 0 | M9- | 37 – 38 | | 13975598 | | | | 1943 |
| 1944 | IV | 827 | 0 | 0 | 283 | 0 | 0 | Mono | Mono | | 13987489 | | | | 1944 |
| 1948 | III | 824 | 1 | 2 | 283 | 0 | 0 | M9+ | 1 | | 13992420 | | | | 1948 |
| 1953 | IV | 827 | 0 | 0 | 283 | 0 | 0 | Mono | Mono | | 14003440 | | | | 1953 |
| 1954 | IV | 827 | 0 | 0 | 283 | 0 | 0 | Mono | Mono | | 14003581 | | | | 1954 |
| 1955 | IV | 827 | 0 | 0 | 283 | 0 | 0 | Mono | Mono | | 14005951 | | | | 1955 |
| 1960 | IV | 827 | 0 | 0 | 283 | 0 | 0 | Mono | Mono | | 14024615 | | | | 1960 |
| 1961 | III | 788 | 1 | 38 | 247 | 21 | 15 | BOTH | | PI, 1H | 14027381 | | | | 1961 |
| 1964 | I | 827 | 0 | 0 | 262 | 21 | 0 | M9- | 11 – 20 | | 14034508 | | | | 1964 |
| 1968 | IV | 821 | 0 | 6 | 280 | 0 | 3 | Mono | Mono | | 14043151 | | | | 1968 |
| 1969 | IV | 827 | 0 | 0 | 283 | 0 | 0 | Mono | Mono | | 14046864 | | | | 1969 |
| 1977 | IV | 827 | 0 | 0 | 283 | 0 | 0 | Mono | Mono | | 14060159 | | | | 1977 |
| 1983 | IV | 827 | 0 | 0 | 283 | 0 | 0 | Mono | Mono | | 14064856 | | | | 1983 |
| 1989 | IV | 824 | 0 | 3 | 283 | 0 | 0 | Mono | Mono | | 14074877 | | | | 1989 |
| 1995 | II | 826 | 1 | 0 | 283 | 0 | 0 | M9+ | 1 | | 14082442 | | | | 1995 |
| 1997 | II | 826 | 1 | 0 | 283 | 0 | 0 | M9+ | 2 | | 14085009 | | | | 1997 |
| 1998 | III | 800 | 0 | 27 | 268 | 11 | 4 | M9- | 37 – 38 | | 14089323 | | | | 1998 |
| 2005 | III | 810 | 1 | 16 | 279 | 0 | 4 | M9+ | 72 | | 14102798 | | | | 2005 |
| 2031 | IV | 827 | 0 | 0 | 283 | 0 | 0 | Mono | Mono | | 14170424 | | | | 2031 |
| 2036 | IV | 826 | 0 | 1 | 283 | 0 | 0 | Mono | Mono | | 14177422 | | | | 2036 |
| 2048 | IV | 824 | 0 | 3 | 282 | 0 | 1 | Mono | Mono | | 14199835 | | | | 2048 |
| 2050 | IV | 827 | 0 | 0 | 283 | 0 | 0 | Mono | Mono | | 14201316 | | | | 2050 |
| 2059 | IV | 826 | 0 | 1 | 283 | 0 | 0 | Mono | Mono | | 14219307 | | | | 2059 |
| 2060 | IV | 827 | 0 | 0 | 283 | 0 | 0 | Mono | Mono | | 14219849 | | | | 2060 |
| 2072 | IV | 827 | 0 | 0 | 283 | 0 | 0 | Mono | Mono | | 14243581 | | | | 2072 |
| 2074 | IV | 827 | 0 | 0 | 283 | 0 | 0 | Mono | Mono | | 14245444 | | | | 2074 |
| 2076 | III | 788 | 0 | 39 | 271 | 2 | 10 | M9- | 8 | | 14253637 | | | | 2076 |
| 2080 | I | 827 | 0 | 0 | 276 | 7 | 0 | M9- | 39 | | 14266305 | | | | 2080 |
| 2086 | IV | 827 | 0 | 0 | 283 | 0 | 0 | Mono | Mono | | 14280847 | | | | 2086 |
| 2089 | IV | 827 | 0 | 0 | 283 | 0 | 0 | Mono | Mono | | 14287097 | | | | 2089 |
| 2100 | I | 827 | 0 | 0 | 246 | 37 | 0 | M9- | 1 – 10 | | 14324714 | | | | 2100 |
| 2102 | I | 827 | 0 | 0 | 0 | 283 | 0 | M9+/- | M9+/- | | 14330570 | | | | 2102 |
| 2103 | IV | 824 | 0 | 3 | 281 | 0 | 2 | Mono | Mono | | 14331142 | | | | 2103 |
| 2104 | IV | 0 | 827 | 0 | 0 | 283 | 0 | Mono | Mono | | 14331272 | | | | 2104 |
| 2107 | IV | 824 | 0 | 3 | 281 | 0 | 2 | Mono | Mono | | 14334030 | | | | 2107 |
| 2110 | IV | 814 | 0 | 13 | 276 | 0 | 7 | Mono | Mono | | 14336165 | | | | 2110 |
| 2114 | IV | 827 | 0 | 0 | 283 | 0 | 0 | Mono | Mono | | 14340428 | | | | 2114 |
| 2117 | I | 827 | 0 | 0 | 187 | 96 | 0 | M9- | 22 – 36 | | 14345705 | | | | 2117 |
| 2118 | II | 826 | 1 | 0 | 283 | 0 | 0 | M9+ | 1 | | 14346901 | | | | 2118 |
| 2121 | IV | 827 | 0 | 0 | 283 | 0 | 0 | Mono | Mono | | 14365984 | | | | 2121 |
| 2122 | I | 827 | 0 | 0 | 246 | 37 | 0 | M9- | 1 – 10 | | 14367181 | | | | 2122 |
| 2125 | IV | 826 | 0 | 1 | 283 | 0 | 0 | Mono | Mono | | 14375731 | | | | 2125 |
| 2132 | IV | 827 | 0 | 0 | 283 | 0 | 0 | Mono | Mono | | 14402596 | | | | 2132 |
| 2136 | IV | 792 | 0 | 35 | 269 | 0 | 14 | Mono | Mono | | 14407089 | | | | 2136 |
| 2142 | III | 35 | 787 | 5 | 283 | 0 | 0 | M9+ | 17 – 93 | | 14435792 | | | | 2142 |
| 2144 | IV | 826 | 0 | 1 | 283 | 0 | 0 | Mono | Mono | | 14442048 | | | | 2144 |
| 2145 | IV | 827 | 0 | 0 | 283 | 0 | 0 | Mono | Mono | | 14443391 | | | | 2145 |
| 2149 | IV | 826 | 0 | 1 | 282 | 0 | 1 | Mono | Mono | | 14466371 | | | | 2149 |
| 2152 | IV | 823 | 0 | 4 | 282 | 0 | 1 | Mono | Mono | | 14470825 | | | | 2152 |
| 2153 | II | 826 | 1 | 0 | 283 | 0 | 0 | M9+ | 1 | | 14471212 | | | | 2153 |
| 2160 | IV | 827 | 0 | 0 | 283 | 0 | 0 | Mono | Mono | | 14484899 | | | | 2160 |
| 2163 | IV | 825 | 0 | 2 | 283 | 0 | 0 | Mono | Mono | | 14491707 | | | | 2163 |
| 2175 | IV | 827 | 0 | 0 | 283 | 0 | 0 | Mono | Mono | | 14515265 | | | | 2175 |
| 2176 | IV | 826 | 0 | 1 | 283 | 0 | 0 | Mono | Mono | | 14517684 | | | | 2176 |
| 2178 | IV | 827 | 0 | 0 | 283 | 0 | 0 | Mono | Mono | | 14529518 | | | | 2178 |
| 2180 | IV | 827 | 0 | 0 | 283 | 0 | 0 | Mono | Mono | | 14531350 | | | | 2180 |
| 2181 | III | 821 | 1 | 5 | 282 | 0 | 1 | M9+ | 1 | | 14534569 | | | | 2181 |
| 2184 | IV | 827 | 0 | 0 | 283 | 0 | 0 | Mono | Mono | | 14548082 | | | | 2184 |
| 2185 | IV | 827 | 0 | 0 | 283 | 0 | 0 | Mono | Mono | | 14552147 | | | | 2185 |
| 2187 | IV | 823 | 0 | 4 | 283 | 0 | 0 | Mono | Mono | | 14564336 | | | | 2187 |
| 2192 | II | 826 | 1 | 0 | 283 | 0 | 0 | M9+ | 1 | | 14579178 | | | | 2192 |
| 2193 | IV | 820 | 0 | 7 | 282 | 0 | 1 | Mono | Mono | | 14585108 | | | | 2193 |
| 2195 | III | 3 | 822 | 2 | 283 | 0 | 0 | M9+ | 3 – 93 | | 14598808 | | | | 2195 |
| 2198 | IV | 827 | 0 | 0 | 283 | 0 | 0 | Mono | Mono | | 14604583 | | | | 2198 |
| 2199 | IV | 820 | 0 | 7 | 281 | 0 | 2 | Mono | Mono | | 14605172 | | | | 2199 |
| 2200 | IV | 827 | 0 | 0 | 283 | 0 | 0 | Mono | Mono | | 14605758 | | | | 2200 |
| 2208 | I | 827 | 0 | 0 | 246 | 37 | 0 | M9- | 1 – 10 | | 14636530 | | | | 2208 |
| 2211 | IV | 827 | 0 | 0 | 283 | 0 | 0 | Mono | Mono | | 14641599 | | | | 2211 |
| 2212 | IV | 827 | 0 | 0 | 283 | 0 | 0 | Mono | Mono | | 14643410 | | | | 2212 |
| 2224 | IV | 825 | 0 | 2 | 282 | 0 | 1 | Mono | Mono | | 14680165 | | | | 2224 |
| 2229 | IV | 0 | 827 | 0 | 0 | 283 | 0 | Mono | Mono | | 14709276 | | | | 2229 |
| 2232 | IV | 827 | 0 | 0 | 283 | 0 | 0 | Mono | Mono | | 14734176 | | | | 2232 |

| | | | M9+ | | | M9- | | | | | Build 37 | | ISOGG | | |
|---|---|---|---|---|---|---|---|---|---|---|---|---|---|---|---|
| List | Stage | 0 | 1 | . | 0 | 1 | . | M9? | Clade(s) | Prob. | Position | Name | Subgroup Name | Amb. | List |
| 2233 | IV | 827 | 0 | 0 | 283 | 0 | 0 | Mono | Mono | | 14734195 | | | | 2233 |
| 2234 | IV | 827 | 0 | 0 | 283 | 0 | 0 | Mono | Mono | | 14734216 | | | | 2234 |
| 2238 | IV | 0 | 827 | 0 | 0 | 283 | 0 | Mono | Mono | | 14753989 | | | | 2238 |
| 2241 | IV | 826 | 0 | 1 | 283 | 0 | 0 | Mono | Mono | | 14763172 | | | | 2241 |
| 2242 | IV | 827 | 0 | 0 | 283 | 0 | 0 | Mono | Mono | | 14763240 | | | | 2242 |
| 2244 | IV | 827 | 0 | 0 | 282 | 0 | 1 | Mono | Mono | | 14769786 | | | | 2244 |
| 2245 | IV | 827 | 0 | 0 | 283 | 0 | 0 | Mono | Mono | | 14769935 | | | | 2245 |
| 2246 | IV | 827 | 0 | 0 | 283 | 0 | 0 | Mono | Mono | | 14770351 | | | | 2246 |
| 2249 | IV | 827 | 0 | 0 | 283 | 0 | 0 | Mono | Mono | | 14773984 | | | | 2249 |
| 2254 | IV | 0 | 827 | 0 | 0 | 283 | 0 | Mono | Mono | | 14777842 | | | | 2254 |
| 2259 | IV | 827 | 0 | 0 | 283 | 0 | 0 | Mono | Mono | | 14786868 | | | | 2259 |
| 2261 | IV | 826 | 0 | 1 | 282 | 0 | 1 | Mono | Mono | | 14788344 | | | | 2261 |
| 2263 | III | 823 | 1 | 3 | 281 | 0 | 2 | M9+ | 1 | | 14790357 | | | | 2263 |
| 2269 | I | 827 | 0 | 0 | 178 | 105 | 0 | M9- | 37 – 59 | | 14810672 | | | | 2269 |
| 2271 | IV | 827 | 0 | 0 | 283 | 0 | 0 | Mono | Mono | | 14812876 | | | | 2271 |
| 2275 | IV | 827 | 0 | 0 | 283 | 0 | 0 | Mono | Mono | | 14814539 | | | | 2275 |
| 2276 | IV | 827 | 0 | 0 | 283 | 0 | 0 | Mono | Mono | | 14814540 | | | | 2276 |
| 2277 | IV | 827 | 0 | 0 | 283 | 0 | 0 | Mono | Mono | | 14814550 | | | | 2277 |
| 2278 | IV | 827 | 0 | 0 | 283 | 0 | 0 | Mono | Mono | | 14814602 | | | | 2278 |
| 2279 | IV | 827 | 0 | 0 | 283 | 0 | 0 | Mono | Mono | | 14817309 | | | | 2279 |
| 2284 | I | 827 | 0 | 0 | 246 | 37 | 0 | M9- | 1 – 10 | | 14825719 | | | | 2284 |
| 2285 | III | 5 | 821 | 1 | 283 | 0 | 0 | M9+ | | PI | 14825721 | | | | 2285 |
| 2286 | IV | 827 | 0 | 0 | 283 | 0 | 0 | Mono | Mono | | 14827703 | | | | 2286 |
| 2287 | II | 826 | 1 | 0 | 283 | 0 | 0 | M9+ | 1 | | 14828362 | | | | 2287 |
| 2288 | IV | 827 | 0 | 0 | 283 | 0 | 0 | Mono | Mono | | 14832645 | | | | 2288 |
| 2290 | IV | 827 | 0 | 0 | 283 | 0 | 0 | Mono | Mono | | 14838533 | | | | 2290 |
| 2291 | IV | 827 | 0 | 0 | 283 | 0 | 0 | Mono | Mono | | 14838688 | | | | 2291 |
| 2294 | IV | 827 | 0 | 0 | 283 | 0 | 0 | Mono | Mono | | 14839786 | | | | 2294 |
| 2298 | IV | 827 | 0 | 0 | 283 | 0 | 0 | Mono | Mono | | 14848117 | | | | 2298 |
| 2305 | IV | 815 | 0 | 12 | 278 | 0 | 5 | Mono | Mono | | 14851961 | | | | 2305 |
| 2306 | IV | 827 | 0 | 0 | 283 | 0 | 0 | Mono | Mono | | 14854941 | | | | 2306 |
| 2313 | IV | 827 | 0 | 0 | 283 | 0 | 0 | Mono | Mono | | 14869180 | | | | 2313 |
| 2314 | IV | 827 | 0 | 0 | 283 | 0 | 0 | Mono | Mono | | 14869183 | | | | 2314 |
| 2319 | IV | 827 | 0 | 0 | 283 | 0 | 0 | Mono | Mono | | 14875232 | | | | 2319 |
| 2322 | IV | 827 | 0 | 0 | 283 | 0 | 0 | Mono | Mono | | 14883004 | | | | 2322 |
| 2323 | IV | 827 | 0 | 0 | 283 | 0 | 0 | Mono | Mono | | 14883033 | | | | 2323 |
| 2325 | IV | 827 | 0 | 0 | 282 | 0 | 1 | Mono | Mono | | 14885546 | | | | 2325 |
| 2326 | IV | 827 | 0 | 0 | 283 | 0 | 0 | Mono | Mono | | 14885779 | | | | 2326 |
| 2327 | IV | 827 | 0 | 0 | 283 | 0 | 0 | Mono | Mono | | 14885815 | | | | 2327 |
| 2332 | IV | 823 | 0 | 4 | 281 | 0 | 2 | Mono | Mono | | 14890152 | | | | 2332 |
| 2333 | IV | 827 | 0 | 0 | 283 | 0 | 0 | Mono | Mono | | 14890604 | | | | 2333 |
| 2334 | IV | 827 | 0 | 0 | 283 | 0 | 0 | Mono | Mono | | 14890987 | | | | 2334 |
| 2341 | IV | 827 | 0 | 0 | 283 | 0 | 0 | Mono | Mono | | 14898180 | | | | 2341 |
| 2342 | IV | 827 | 0 | 0 | 283 | 0 | 0 | Mono | Mono | | 14898241 | | | | 2342 |
| 2344 | II | 827 | 0 | 0 | 282 | 1 | 0 | M9- | NU | | 14898465 | | | | 2344 |
| 2345 | IV | 827 | 0 | 0 | 283 | 0 | 0 | Mono | Mono | | 14898546 | | | | 2345 |
| 2346 | IV | 827 | 0 | 0 | 283 | 0 | 0 | Mono | Mono | | 14898570 | | | | 2346 |
| 2347 | IV | 827 | 0 | 0 | 283 | 0 | 0 | Mono | Mono | | 14898678 | | | | 2347 |
| 2348 | IV | 793 | 0 | 34 | 283 | 0 | 0 | Mono | Mono | | 14902412 | | | | 2348 |
| 2356 | IV | 827 | 0 | 0 | 283 | 0 | 0 | Mono | Mono | | 14906886 | | | | 2356 |
| 2360 | IV | 827 | 0 | 0 | 283 | 0 | 0 | Mono | Mono | | 14917573 | | | | 2360 |
| 2362 | IV | 827 | 0 | 0 | 283 | 0 | 0 | Mono | Mono | | 14917922 | | | | 2362 |
| 2363 | IV | 827 | 0 | 0 | 283 | 0 | 0 | Mono | Mono | | 14920900 | | | | 2363 |
| 2364 | IV | 823 | 0 | 4 | 282 | 0 | 1 | Mono | Mono | | 14922451 | | | | 2364 |
| 2366 | IV | 827 | 0 | 0 | 283 | 0 | 0 | Mono | Mono | | 14922817 | | | | 2366 |
| 2368 | IV | 827 | 0 | 0 | 283 | 0 | 0 | Mono | Mono | | 14923457 | | | | 2368 |
| 2371 | IV | 827 | 0 | 0 | 283 | 0 | 0 | Mono | Mono | | 14927957 | | | | 2371 |
| 2372 | IV | 827 | 0 | 0 | 283 | 0 | 0 | Mono | Mono | | 14927960 | | | | 2372 |
| 2375 | IV | 826 | 0 | 1 | 282 | 0 | 1 | Mono | Mono | | 14928564 | | | | 2375 |
| 2377 | IV | 827 | 0 | 0 | 283 | 0 | 0 | Mono | Mono | | 14930502 | | | | 2377 |
| 2383 | IV | 827 | 0 | 0 | 283 | 0 | 0 | Mono | Mono | | 14944433 | | | | 2383 |
| 2384 | IV | 827 | 0 | 0 | 283 | 0 | 0 | Mono | Mono | | 14945191 | | | | 2384 |
| 2385 | IV | 827 | 0 | 0 | 283 | 0 | 0 | Mono | Mono | | 14945628 | | | | 2385 |
| 2386 | IV | 826 | 0 | 1 | 283 | 0 | 0 | Mono | Mono | | 14945646 | | | | 2386 |
| 2387 | IV | 827 | 0 | 0 | 283 | 0 | 0 | Mono | Mono | | 14945678 | | | | 2387 |
| 2390 | IV | 827 | 0 | 0 | 283 | 0 | 0 | Mono | Mono | | 14948644 | | | | 2390 |
| 2391 | IV | 827 | 0 | 0 | 283 | 0 | 0 | Mono | Mono | | 14952324 | | | | 2391 |
| 2393 | IV | 827 | 0 | 0 | 283 | 0 | 0 | Mono | Mono | | 14953040 | | | | 2393 |
| 2395 | IV | 827 | 0 | 0 | 283 | 0 | 0 | Mono | Mono | | 14954119 | | | | 2395 |
| 2396 | IV | 827 | 0 | 0 | 283 | 0 | 0 | Mono | Mono | | 14954378 | | | | 2396 |
| 2397 | IV | 827 | 0 | 0 | 283 | 0 | 0 | Mono | Mono | | 14954404 | | | | 2397 |
| 2398 | IV | 827 | 0 | 0 | 283 | 0 | 0 | Mono | Mono | | 14954978 | | | | 2398 |
| 2402 | IV | 827 | 0 | 0 | 283 | 0 | 0 | Mono | Mono | | 14958901 | | | | 2402 |
| 2404 | IV | 827 | 0 | 0 | 283 | 0 | 0 | Mono | Mono | | 14959244 | | | | 2404 |
| 2409 | IV | 827 | 0 | 0 | 283 | 0 | 0 | Mono | Mono | | 14968722 | | | | 2409 |
| 2411 | IV | 827 | 0 | 0 | 283 | 0 | 0 | Mono | Mono | | 14971316 | | | | 2411 |
| 2414 | IV | 827 | 0 | 0 | 283 | 0 | 0 | Mono | Mono | | 14976252 | | | | 2414 |
| 2420 | IV | 827 | 0 | 0 | 283 | 0 | 0 | Mono | Mono | | 14982984 | | | | 2420 |
| 2422 | IV | 827 | 0 | 0 | 283 | 0 | 0 | Mono | Mono | | 14984713 | | | | 2422 |
| 2423 | IV | 827 | 0 | 0 | 283 | 0 | 0 | Mono | Mono | | 14985806 | | | | 2423 |
| 2428 | IV | 825 | 0 | 2 | 283 | 0 | 0 | Mono | Mono | | 14994696 | | | | 2428 |
| 2436 | IV | 827 | 0 | 0 | 283 | 0 | 0 | Mono | Mono | | 15015396 | | | | 2436 |
| 2444 | IV | 827 | 0 | 0 | 283 | 0 | 0 | Mono | Mono | | 15019124 | | | | 2444 |
| 2448 | IV | 809 | 0 | 18 | 277 | 0 | 6 | Mono | Mono | | 15020304 | | | | 2448 |
| 2450 | IV | 827 | 0 | 0 | 283 | 0 | 0 | Mono | Mono | | 15021104 | | | | 2450 |
| 2451 | II | 826 | 1 | 0 | 283 | 0 | 0 | M9+ | 2 | | 15021516 | | | | 2451 |
| 2453 | IV | 820 | 0 | 7 | 279 | 0 | 4 | Mono | Mono | | 15022184 | | | | 2453 |
| 2458 | IV | 827 | 0 | 0 | 283 | 0 | 0 | Mono | Mono | | 15024151 | | | | 2458 |
| 2459 | IV | 827 | 0 | 0 | 283 | 0 | 0 | Mono | Mono | | 15024924 | | | | 2459 |
| 2465 | IV | 827 | 0 | 0 | 283 | 0 | 0 | Mono | Mono | | 15026818 | | | | 2465 |
| 2466 | IV | 827 | 0 | 0 | 283 | 0 | 0 | Mono | Mono | | 15027132 | | | | 2466 |
| 2469 | IV | 827 | 0 | 0 | 283 | 0 | 0 | Mono | Mono | | 15027510 | | | | 2469 |
| 2471 | I | 821 | 6 | 0 | 283 | 0 | 0 | M9+ | 24 | | 15027638 | | | | 2471 |
| 2472 | IV | 826 | 0 | 1 | 283 | 0 | 0 | Mono | Mono | | 15028176 | | | | 2472 |
| 2473 | IV | 827 | 0 | 0 | 283 | 0 | 0 | Mono | Mono | | 15028771 | | | | 2473 |
| 2474 | IV | 826 | 0 | 1 | 283 | 0 | 0 | Mono | Mono | | 15028846 | | | | 2474 |

| List | Stage | M9+ 0 | M9+ 1 | M9+ . | M9- 0 | M9- 1 | M9- . | M9? | Clade(s) | Prob. | Build 37 Position | Name | ISOGG Subgroup Name | Amb. | List |
|---|---|---|---|---|---|---|---|---|---|---|---|---|---|---|---|
| 2475 | III | 820 | 1 | 6 | 282 | 0 | 1 | M9+ | 13 | | 15029413 | | | | 2475 |
| 2477 | IV | 827 | 0 | 0 | 283 | 0 | 0 | Mono | Mono | | 15029968 | | | | 2477 |
| 2482 | IV | 827 | 0 | 0 | 283 | 0 | 0 | Mono | Mono | | 15030915 | | | | 2482 |
| 2483 | IV | 826 | 0 | 1 | 283 | 0 | 0 | Mono | Mono | | 15031109 | | | | 2483 |
| 2487 | I | 0 | 827 | 0 | 21 | 262 | 0 | M9- | 11 – 20 | | 15037754 | | | | 2487 |
| 2491 | II | 826 | 1 | 0 | 283 | 0 | 0 | M9+ | 1 | | 15041346 | | | | 2491 |
| 2493 | IV | 827 | 0 | 0 | 283 | 0 | 0 | Mono | Mono | | 15042327 | | | | 2493 |
| 2496 | IV | 827 | 0 | 0 | 283 | 0 | 0 | Mono | Mono | | 15043778 | | | | 2496 |
| 2498 | IV | 827 | 0 | 0 | 283 | 0 | 0 | Mono | Mono | | 15053296 | | | | 2498 |
| 2499 | IV | 827 | 0 | 0 | 283 | 0 | 0 | Mono | Mono | | 15055439 | | | | 2499 |
| 2507 | IV | 827 | 0 | 0 | 283 | 0 | 0 | Mono | Mono | | 15082677 | | | | 2507 |
| 2508 | IV | 827 | 0 | 0 | 283 | 0 | 0 | Mono | Mono | | 15083396 | | | | 2508 |
| 2514 | IV | 0 | 825 | 2 | 0 | 283 | 0 | Mono | Mono | | 15094400 | | | | 2514 |
| 2523 | I | 0 | 827 | 0 | 21 | 262 | 0 | M9- | 11 – 20 | | 15110364 | | | | 2523 |
| 2530 | IV | 827 | 0 | 0 | 283 | 0 | 0 | Mono | Mono | | 15126199 | | | | 2530 |
| 2531 | IV | 0 | 827 | 0 | 0 | 283 | 0 | Mono | Mono | | 15129415 | | | | 2531 |
| 2533 | IV | 827 | 0 | 0 | 283 | 0 | 0 | Mono | Mono | | 15140670 | | | | 2533 |
| 2534 | I | 0 | 827 | 0 | 21 | 262 | 0 | M9- | 11 – 20 | | 15152317 | | | | 2534 |
| 2535 | IV | 827 | 0 | 0 | 283 | 0 | 0 | Mono | Mono | | 15156388 | | | | 2535 |
| 2536 | II | 826 | 1 | 0 | 283 | 0 | 0 | M9+ | 1 | | 15160809 | | | | 2536 |
| 2538 | IV | 824 | 0 | 3 | 283 | 0 | 0 | Mono | Mono | | 15165471 | | | | 2538 |
| 2540 | IV | 0 | 827 | 0 | 0 | 283 | 0 | Mono | Mono | | 15169439 | | | | 2540 |
| 2541 | IV | 827 | 0 | 0 | 283 | 0 | 0 | Mono | Mono | | 15182178 | | | | 2541 |
| 2547 | IV | 827 | 0 | 0 | 283 | 0 | 0 | Mono | Mono | | 15206476 | | | | 2547 |
| 2548 | IV | 826 | 0 | 1 | 283 | 0 | 0 | Mono | Mono | | 15206662 | | | | 2548 |
| 2552 | IV | 827 | 0 | 0 | 283 | 0 | 0 | Mono | Mono | | 15222399 | | | | 2552 |
| 2554 | IV | 827 | 0 | 0 | 283 | 0 | 0 | Mono | Mono | | 15222764 | | | | 2554 |
| 2555 | IV | 810 | 0 | 17 | 268 | 0 | 15 | Mono | Mono | | 15227283 | | | | 2555 |
| 2560 | IV | 827 | 0 | 0 | 283 | 0 | 0 | Mono | Mono | | 15239377 | | | | 2560 |
| 2563 | IV | 827 | 0 | 0 | 283 | 0 | 0 | Mono | Mono | | 15242077 | | | | 2563 |
| 2564 | IV | 827 | 0 | 0 | 283 | 0 | 0 | Mono | Mono | | 15242249 | | | | 2564 |
| 2567 | IV | 827 | 0 | 0 | 283 | 0 | 0 | Mono | Mono | | 15255494 | | | | 2567 |
| 2570 | IV | 827 | 0 | 0 | 283 | 0 | 0 | Mono | Mono | | 15262035 | | | | 2570 |
| 2571 | IV | 827 | 0 | 0 | 283 | 0 | 0 | Mono | Mono | | 15262755 | | | | 2571 |
| 2572 | IV | 827 | 0 | 0 | 283 | 0 | 0 | Mono | Mono | | 15262779 | | | | 2572 |
| 2573 | IV | 826 | 0 | 1 | 283 | 0 | 0 | Mono | Mono | | 15264052 | | | | 2573 |
| 2574 | IV | 827 | 0 | 0 | 283 | 0 | 0 | Mono | Mono | | 15266547 | | | | 2574 |
| 2577 | III | 827 | 0 | 0 | 260 | 22 | 1 | M9- | 60 – 66 | MD | 15278569 | | | | 2577 |
| 2578 | III | 824 | 0 | 3 | 246 | 37 | 0 | M9- | 1 – 10 | | 15281135 | | | | 2578 |
| 2579 | IV | 827 | 0 | 0 | 283 | 0 | 0 | Mono | Mono | | 15302894 | | | | 2579 |
| 2580 | IV | 826 | 0 | 1 | 283 | 0 | 0 | Mono | Mono | | 15302901 | | | | 2580 |
| 2581 | IV | 827 | 0 | 0 | 283 | 0 | 0 | Mono | Mono | | 15309625 | | | | 2581 |
| 2586 | IV | 827 | 0 | 0 | 283 | 0 | 0 | Mono | Mono | | 15321839 | | | | 2586 |
| 2593 | IV | 827 | 0 | 0 | 283 | 0 | 0 | Mono | Mono | | 15343127 | | | | 2593 |
| 2595 | IV | 827 | 0 | 0 | 283 | 0 | 0 | Mono | Mono | | 15346546 | | | | 2595 |
| 2597 | IV | 827 | 0 | 0 | 283 | 0 | 0 | Mono | Mono | | 15350615 | | | | 2597 |
| 2598 | IV | 827 | 0 | 0 | 283 | 0 | 0 | Mono | Mono | | 15350616 | | | | 2598 |
| 2599 | IV | 827 | 0 | 0 | 283 | 0 | 0 | Mono | Mono | | 15353217 | | | | 2599 |
| 2601 | IV | 827 | 0 | 0 | 283 | 0 | 0 | Mono | Mono | | 15372130 | | | | 2601 |
| 2610 | III | 825 | 1 | 1 | 283 | 0 | 0 | M9+ | 1 | | 15383004 | | | | 2610 |
| 2613 | IV | 827 | 0 | 0 | 283 | 0 | 0 | Mono | Mono | | 15393044 | | | | 2613 |
| 2615 | IV | 823 | 0 | 4 | 282 | 0 | 1 | Mono | Mono | | 15398613 | | | | 2615 |
| 2616 | IV | 827 | 0 | 0 | 283 | 0 | 0 | Mono | Mono | | 15399195 | | | | 2616 |
| 2618 | IV | 827 | 0 | 0 | 283 | 0 | 0 | Mono | Mono | | 15401876 | | | | 2618 |
| 2619 | IV | 827 | 0 | 0 | 283 | 0 | 0 | Mono | Mono | | 15401877 | | | | 2619 |
| 2620 | IV | 827 | 0 | 0 | 283 | 0 | 0 | Mono | Mono | | 15404832 | | | | 2620 |
| 2625 | IV | 827 | 0 | 0 | 283 | 0 | 0 | Mono | Mono | | 15414820 | | | | 2625 |
| 2627 | IV | 827 | 0 | 0 | 283 | 0 | 0 | Mono | Mono | | 15420555 | | | | 2627 |
| 2633 | IV | 827 | 0 | 0 | 283 | 0 | 0 | Mono | Mono | | 15436444 | | | | 2633 |
| 2641 | IV | 825 | 0 | 2 | 283 | 0 | 0 | Mono | Mono | | 15437410 | | | | 2641 |
| 2644 | IV | 827 | 0 | 0 | 283 | 0 | 0 | Mono | Mono | | 15438777 | | | | 2644 |
| 2648 | IV | 825 | 0 | 2 | 283 | 0 | 0 | Mono | Mono | | 15442970 | | | | 2648 |
| 2649 | IV | 827 | 0 | 0 | 283 | 0 | 0 | Mono | Mono | | 15445765 | | | | 2649 |
| 2653 | IV | 826 | 0 | 1 | 283 | 0 | 0 | Mono | Mono | | 15464526 | | | | 2653 |
| 2654 | IV | 827 | 0 | 0 | 283 | 0 | 0 | Mono | Mono | | 15466915 | | | | 2654 |
| 2655 | IV | 825 | 0 | 2 | 283 | 0 | 0 | Mono | Mono | | 15466948 | | | | 2655 |
| 2656 | IV | 827 | 0 | 0 | 283 | 0 | 0 | Mono | Mono | | 15466966 | | | | 2656 |
| 2657 | IV | 827 | 0 | 0 | 283 | 0 | 0 | Mono | Mono | | 15467064 | | | | 2657 |
| 2658 | IV | 827 | 0 | 0 | 283 | 0 | 0 | Mono | Mono | | 15467106 | | | | 2658 |
| 2659 | II | 827 | 0 | 0 | 282 | 1 | 0 | M9- | NU | | 15467230 | | | | 2659 |
| 2663 | IV | 827 | 0 | 0 | 283 | 0 | 0 | Mono | Mono | | 15469768 | | | | 2663 |
| 2664 | IV | 827 | 0 | 0 | 283 | 0 | 0 | Mono | Mono | | 15470057 | | | | 2664 |
| 2676 | IV | 826 | 0 | 1 | 283 | 0 | 0 | Mono | Mono | | 15496600 | | | | 2676 |
| 2679 | IV | 827 | 0 | 0 | 282 | 0 | 1 | Mono | Mono | | 15502402 | | | | 2679 |
| 2684 | IV | 827 | 0 | 0 | 283 | 0 | 0 | Mono | Mono | | 15507686 | | | | 2684 |
| 2699 | IV | 827 | 0 | 0 | 283 | 0 | 0 | Mono | Mono | | 15528107 | | | | 2699 |
| 2702 | IV | 827 | 0 | 0 | 283 | 0 | 0 | Mono | Mono | | 15533845 | | | | 2702 |
| 2705 | IV | 827 | 0 | 0 | 283 | 0 | 0 | Mono | Mono | | 15538487 | | | | 2705 |
| 2708 | IV | 827 | 0 | 0 | 283 | 0 | 0 | Mono | Mono | | 15542883 | | | | 2708 |
| 2711 | IV | 827 | 0 | 0 | 283 | 0 | 0 | Mono | Mono | | 15549050 | | | | 2711 |
| 2712 | IV | 827 | 0 | 0 | 283 | 0 | 0 | Mono | Mono | | 15551636 | | | | 2712 |
| 2713 | I | 827 | 0 | 0 | 246 | 37 | 0 | M9- | 1 – 10 | | 15553123 | | | | 2713 |
| 2715 | IV | 826 | 0 | 1 | 283 | 0 | 0 | Mono | Mono | | 15554755 | | | | 2715 |
| 2720 | IV | 827 | 0 | 0 | 283 | 0 | 0 | Mono | Mono | | 15564625 | | | | 2720 |
| 2721 | IV | 827 | 0 | 0 | 283 | 0 | 0 | Mono | Mono | | 15566388 | | | | 2721 |
| 2733 | IV | 827 | 0 | 0 | 283 | 0 | 0 | Mono | Mono | | 15585558 | | | | 2733 |
| 2747 | II | 826 | 1 | 0 | 283 | 0 | 0 | M9+ | 8 | | 15591497 | | | | 2747 |
| 2749 | IV | 827 | 0 | 0 | 283 | 0 | 0 | Mono | Mono | | 15592033 | | | | 2749 |
| 2750 | IV | 827 | 0 | 0 | 283 | 0 | 0 | Mono | Mono | | 15592635 | | | | 2750 |
| 2751 | IV | 827 | 0 | 0 | 283 | 0 | 0 | Mono | Mono | | 15592640 | | | | 2751 |
| 2752 | IV | 827 | 0 | 0 | 283 | 0 | 0 | Mono | Mono | | 15592788 | | | | 2752 |
| 2753 | IV | 827 | 0 | 0 | 283 | 0 | 0 | Mono | Mono | | 15592813 | | | | 2753 |
| 2757 | IV | 827 | 0 | 0 | 283 | 0 | 0 | Mono | Mono | | 15600671 | | | | 2757 |
| 2758 | IV | 827 | 0 | 0 | 283 | 0 | 0 | Mono | Mono | | 15601452 | | | | 2758 |
| 2779 | IV | 827 | 0 | 0 | 283 | 0 | 0 | Mono | Mono | | 15655759 | | | | 2779 |
| 2784 | III | 825 | 1 | 1 | 283 | 0 | 0 | M9+ | 1 | | 15660684 | | | | 2784 |

| List | Stage | M9+ 0 | M9+ 1 | M9+ . | M9- 0 | M9- 1 | M9- . | M9? | Clade(s) | Prob. | Build 37 Position | Name | ISOGG Subgroup Name | Amb. | List |
|---|---|---|---|---|---|---|---|---|---|---|---|---|---|---|---|
| 2785 | IV | 824 | 0 | 3 | 283 | 0 | 0 | Mono | Mono | | 15660852 | | | | 2785 |
| 2788 | IV | 826 | 0 | 1 | 282 | 0 | 1 | Mono | Mono | | 15665459 | | | | 2788 |
| 2792 | IV | 827 | 0 | 0 | 283 | 0 | 0 | Mono | Mono | | 15679655 | | | | 2792 |
| 2796 | IV | 827 | 0 | 0 | 283 | 0 | 0 | Mono | Mono | | 15693300 | | | | 2796 |
| 2798 | IV | 827 | 0 | 0 | 283 | 0 | 0 | Mono | Mono | | 15698201 | | | | 2798 |
| 2803 | IV | 827 | 0 | 0 | 283 | 0 | 0 | Mono | Mono | | 15704602 | | | | 2803 |
| 2806 | III | 826 | 0 | 1 | 245 | 37 | 1 | M9- | 1 – 10 | | 15712801 | | | | 2806 |
| 2807 | IV | 827 | 0 | 0 | 283 | 0 | 0 | Mono | Mono | | 15717300 | | | | 2807 |
| 2810 | IV | 827 | 0 | 0 | 279 | 0 | 4 | Mono | Mono | | 15724290 | | | | 2810 |
| 2813 | IV | 827 | 0 | 0 | 279 | 0 | 4 | Mono | Mono | | 15726484 | | | | 2813 |
| 2816 | IV | 827 | 0 | 0 | 283 | 0 | 0 | Mono | Mono | | 15736328 | | | | 2816 |
| 2824 | IV | 827 | 0 | 0 | 283 | 0 | 0 | Mono | Mono | | 15755918 | | | | 2824 |
| 2828 | IV | 701 | 0 | 126 | 240 | 0 | 43 | Mono | Mono | | 15772103 | | | | 2828 |
| 2836 | IV | 827 | 0 | 0 | 283 | 0 | 0 | Mono | Mono | | 15784412 | | | | 2836 |
| 2837 | IV | 827 | 0 | 0 | 283 | 0 | 0 | Mono | Mono | | 15784413 | | | | 2837 |
| 2848 | IV | 827 | 0 | 0 | 283 | 0 | 0 | Mono | Mono | | 15814487 | | | | 2848 |
| 2849 | IV | 827 | 0 | 0 | 283 | 0 | 0 | Mono | Mono | | 15815443 | | | | 2849 |
| 2850 | IV | 825 | 0 | 2 | 283 | 0 | 0 | Mono | Mono | | 15816093 | | | | 2850 |
| 2851 | IV | 826 | 0 | 1 | 283 | 0 | 0 | Mono | Mono | | 15816112 | | | | 2851 |
| 2852 | IV | 827 | 0 | 0 | 283 | 0 | 0 | Mono | Mono | | 15816113 | | | | 2852 |
| 2853 | IV | 827 | 0 | 0 | 283 | 0 | 0 | Mono | Mono | | 15816140 | | | | 2853 |
| 2854 | IV | 827 | 0 | 0 | 283 | 0 | 0 | Mono | Mono | | 15816142 | | | | 2854 |
| 2855 | IV | 827 | 0 | 0 | 283 | 0 | 0 | Mono | Mono | | 15816157 | | | | 2855 |
| 2858 | IV | 826 | 0 | 1 | 283 | 0 | 0 | Mono | Mono | | 15822352 | | | | 2858 |
| 2862 | III | 826 | 0 | 1 | 0 | 283 | 0 | M9+/- | M9+/- | | 15842844 | | | | 2862 |
| 2863 | I | 827 | 0 | 0 | 247 | 36 | 0 | M9- | 2 – 10 | | 15843782 | | | | 2863 |
| 2868 | IV | 827 | 0 | 0 | 283 | 0 | 0 | Mono | Mono | | 15865875 | | | | 2868 |
| 2871 | IV | 827 | 0 | 0 | 283 | 0 | 0 | Mono | Mono | | 15877995 | | | | 2871 |
| 2878 | IV | 812 | 0 | 15 | 279 | 0 | 4 | Mono | Mono | | 15900954 | | | | 2878 |
| 2884 | IV | 827 | 0 | 0 | 283 | 0 | 0 | Mono | Mono | | 15922523 | | | | 2884 |
| 2885 | II | 826 | 1 | 0 | 283 | 0 | 0 | M9+ | 1 | | 15925244 | | | | 2885 |
| 2886 | IV | 825 | 0 | 2 | 283 | 0 | 0 | Mono | Mono | | 15927028 | | | | 2886 |
| 2894 | IV | 827 | 0 | 0 | 283 | 0 | 0 | Mono | Mono | | 15950750 | | | | 2894 |
| 2898 | IV | 827 | 0 | 0 | 283 | 0 | 0 | Mono | Mono | | 15966502 | | | | 2898 |
| 2899 | IV | 822 | 0 | 5 | 280 | 0 | 3 | Mono | Mono | | 15969185 | | | | 2899 |
| 2901 | IV | 827 | 0 | 0 | 283 | 0 | 0 | Mono | Mono | | 15980942 | | | | 2901 |
| 2902 | III | 824 | 2 | 1 | 282 | 1 | 0 | BOTH | | PI | 15984461 | | | | 2902 |
| 2905 | IV | 827 | 0 | 0 | 283 | 0 | 0 | Mono | Mono | | 15992237 | | | | 2905 |
| 2909 | IV | 827 | 0 | 0 | 282 | 0 | 1 | Mono | Mono | | 16004863 | | | | 2909 |
| 2910 | IV | 827 | 0 | 0 | 283 | 0 | 0 | Mono | Mono | | 16010606 | | | | 2910 |
| 2914 | IV | 821 | 0 | 6 | 282 | 0 | 1 | Mono | Mono | | 16040682 | | | | 2914 |
| 2915 | IV | 827 | 0 | 0 | 283 | 0 | 0 | Mono | Mono | | 16041344 | | | | 2915 |
| 2916 | I | 827 | 0 | 0 | 247 | 36 | 0 | M9- | 2 – 10 | | 16045476 | | | | 2916 |
| 2917 | IV | 821 | 0 | 6 | 281 | 0 | 2 | Mono | Mono | | 16046168 | | | | 2917 |
| 2919 | IV | 826 | 0 | 1 | 231 | 0 | 52 | Mono | Mono | | 16052881 | | | | 2919 |
| 2920 | IV | 827 | 0 | 0 | 283 | 0 | 0 | Mono | Mono | | 16071026 | | | | 2920 |
| 2921 | IV | 827 | 0 | 0 | 283 | 0 | 0 | Mono | Mono | | 16074836 | | | | 2921 |
| 2923 | III | 826 | 1 | 0 | 281 | 0 | 2 | M9+ | 1 | | 16086306 | | | | 2923 |
| 2924 | IV | 827 | 0 | 0 | 283 | 0 | 0 | Mono | Mono | | 16087575 | | | | 2924 |
| 2925 | IV | 826 | 0 | 1 | 283 | 0 | 0 | Mono | Mono | | 16096330 | | | | 2925 |
| 2927 | IV | 827 | 0 | 0 | 283 | 0 | 0 | Mono | Mono | | 16172915 | | | | 2927 |
| 2935 | III | 824 | 1 | 2 | 282 | 0 | 1 | M9+ | 2 | | 16188674 | | | | 2935 |
| 2936 | IV | 827 | 0 | 0 | 283 | 0 | 0 | Mono | Mono | | 16190764 | | | | 2936 |
| 2951 | IV | 827 | 0 | 0 | 283 | 0 | 0 | Mono | Mono | | 16205566 | | | | 2951 |
| 2952 | II | 827 | 0 | 0 | 282 | 1 | 0 | M9- | 60 | | 16211449 | | | | 2952 |
| 2954 | IV | 827 | 0 | 0 | 283 | 0 | 0 | Mono | Mono | | 16213100 | | | | 2954 |
| 2960 | III | 575 | 0 | 252 | 62 | 113 | 108 | M9- | 22 – 59 | MD | 16227992 | | | | 2960 |
| 2965 | IV | 827 | 0 | 0 | 283 | 0 | 0 | Mono | Mono | | 16249652 | | | | 2965 |
| 2966 | IV | 827 | 0 | 0 | 283 | 0 | 0 | Mono | Mono | | 16250659 | | | | 2966 |
| 2971 | IV | 827 | 0 | 0 | 283 | 0 | 0 | Mono | Mono | | 16255201 | | | | 2971 |
| 2979 | IV | 827 | 0 | 0 | 283 | 0 | 0 | Mono | Mono | | 16271151 | | | | 2979 |
| 2980 | IV | 826 | 0 | 1 | 283 | 0 | 0 | Mono | Mono | | 16271262 | | | | 2980 |
| 2981 | IV | 825 | 0 | 2 | 283 | 0 | 0 | Mono | Mono | | 16283803 | | | | 2981 |
| 2982 | IV | 827 | 0 | 0 | 283 | 0 | 0 | Mono | Mono | | 16299885 | | | | 2982 |
| 2985 | II | 826 | 1 | 0 | 283 | 0 | 0 | M9+ | 1 | | 16306319 | | | | 2985 |
| 2986 | IV | 827 | 0 | 0 | 283 | 0 | 0 | Mono | Mono | | 16306764 | | | | 2986 |
| 2993 | III | 823 | 1 | 3 | 283 | 0 | 0 | M9+ | 2 | | 16316281 | | | | 2993 |
| 3002 | IV | 827 | 0 | 0 | 283 | 0 | 0 | Mono | Mono | | 16335488 | | | | 3002 |
| 3004 | IV | 827 | 0 | 0 | 283 | 0 | 0 | Mono | Mono | | 16339916 | | | | 3004 |
| 3014 | IV | 827 | 0 | 0 | 282 | 0 | 1 | Mono | Mono | | 16363518 | | | | 3014 |
| 3020 | III | 825 | 1 | 1 | 283 | 0 | 0 | M9+ | 2 | | 16373480 | | | | 3020 |
| 3024 | IV | 827 | 0 | 0 | 283 | 0 | 0 | Mono | Mono | | 16388815 | | | | 3024 |
| 3025 | IV | 827 | 0 | 0 | 282 | 0 | 1 | Mono | Mono | | 16389872 | | | | 3025 |
| 3027 | IV | 827 | 0 | 0 | 283 | 0 | 0 | Mono | Mono | | 16391135 | | | | 3027 |
| 3034 | IV | 819 | 0 | 8 | 280 | 0 | 3 | Mono | Mono | | 16404579 | | | | 3034 |
| 3035 | IV | 827 | 0 | 0 | 283 | 0 | 0 | Mono | Mono | | 16404607 | | | | 3035 |
| 3041 | IV | 827 | 0 | 0 | 283 | 0 | 0 | Mono | Mono | | 16417704 | | | | 3041 |
| 3044 | IV | 827 | 0 | 0 | 283 | 0 | 0 | Mono | Mono | | 16424452 | | | | 3044 |
| 3046 | IV | 827 | 0 | 0 | 283 | 0 | 0 | Mono | Mono | | 16439230 | | | | 3046 |
| 3049 | II | 826 | 1 | 0 | 283 | 0 | 0 | M9+ | 1 | | 16445319 | | | | 3049 |
| 3053 | II | 826 | 1 | 0 | 283 | 0 | 0 | M9+ | 1 | | 16452321 | | | | 3053 |
| 3054 | IV | 827 | 0 | 0 | 283 | 0 | 0 | Mono | Mono | | 16454368 | | | | 3054 |
| 3055 | IV | 827 | 0 | 0 | 283 | 0 | 0 | Mono | Mono | | 16457357 | | | | 3055 |
| 3058 | IV | 827 | 0 | 0 | 283 | 0 | 0 | Mono | Mono | | 16459892 | | | | 3058 |
| 3059 | IV | 827 | 0 | 0 | 283 | 0 | 0 | Mono | Mono | | 16460301 | | | | 3059 |
| 3062 | II | 826 | 1 | 0 | 283 | 0 | 0 | M9+ | 1 | | 16467374 | | | | 3062 |
| 3066 | II | 826 | 1 | 0 | 283 | 0 | 0 | M9+ | 2 | | 16480738 | | | | 3066 |
| 3083 | IV | 827 | 0 | 0 | 282 | 0 | 1 | Mono | Mono | | 16519029 | | | | 3083 |
| 3084 | IV | 827 | 0 | 0 | 282 | 0 | 1 | Mono | Mono | | 16519057 | | | | 3084 |
| 3087 | IV | 821 | 0 | 6 | 282 | 0 | 1 | Mono | Mono | | 16527693 | | | | 3087 |
| 3092 | IV | 827 | 0 | 0 | 283 | 0 | 0 | Mono | Mono | | 16537380 | | | | 3092 |
| 3093 | IV | 827 | 0 | 0 | 283 | 0 | 0 | Mono | Mono | | 16539688 | | | | 3093 |
| 3096 | IV | 827 | 0 | 0 | 283 | 0 | 0 | Mono | Mono | | 16543462 | | | | 3096 |
| 3097 | IV | 827 | 0 | 0 | 283 | 0 | 0 | Mono | Mono | | 16545698 | | | | 3097 |
| 3098 | III | 823 | 2 | 2 | 282 | 0 | 1 | M9+ | | PI | 16548244 | | | | 3098 |
| 3100 | III | 826 | 1 | 0 | 282 | 0 | 1 | M9+ | 1 | | 16549178 | | | | 3100 |

| | | M9+ | | | M9- | | | | | | Build 37 | | ISOGG | | |
|---|---|---|---|---|---|---|---|---|---|---|---|---|---|---|---|
| List | Stage | 0 | 1 | . | 0 | 1 | . | M9? | Clade(s) | Prob. | Position | Name | Subgroup Name | Amb. | List |
| 3106 | I | 827 | 0 | 0 | 246 | 37 | 0 | M9- | 1 – 10 | | 16567551 | | | | 3106 |
| 3107 | IV | 826 | 0 | 1 | 283 | 0 | 0 | Mono | Mono | | 16571915 | | | | 3107 |
| 3131 | IV | 811 | 0 | 16 | 277 | 0 | 6 | Mono | Mono | | 16643296 | | | | 3131 |
| 3132 | IV | 827 | 0 | 0 | 283 | 0 | 0 | Mono | Mono | | 16647978 | | | | 3132 |
| 3133 | IV | 823 | 0 | 4 | 282 | 0 | 1 | Mono | Mono | | 16648116 | | | | 3133 |
| 3137 | I | 827 | 0 | 0 | 246 | 37 | 0 | M9- | 1 – 10 | | 16661107 | | | | 3137 |
| 3139 | IV | 827 | 0 | 0 | 283 | 0 | 0 | Mono | Mono | | 16670327 | | | | 3139 |
| 3141 | IV | 826 | 0 | 1 | 283 | 0 | 0 | Mono | Mono | | 16676564 | | | | 3141 |
| 3145 | IV | 827 | 0 | 0 | 283 | 0 | 0 | Mono | Mono | | 16692495 | | | | 3145 |
| 3153 | IV | 827 | 0 | 0 | 283 | 0 | 0 | Mono | Mono | | 16716818 | | | | 3153 |
| 3154 | III | 826 | 0 | 1 | 187 | 93 | 3 | M9- | 22 – 36 | MD | 16718694 | | | | 3154 |
| 3158 | IV | 824 | 0 | 3 | 282 | 0 | 1 | Mono | Mono | | 16735462 | | | | 3158 |
| 3163 | IV | 827 | 0 | 0 | 283 | 0 | 0 | Mono | Mono | | 16750812 | | | | 3163 |
| 3166 | IV | 827 | 0 | 0 | 283 | 0 | 0 | Mono | Mono | | 16751727 | | | | 3166 |
| 3168 | IV | 827 | 0 | 0 | 283 | 0 | 0 | Mono | Mono | | 16758450 | | | | 3168 |
| 3171 | IV | 827 | 0 | 0 | 283 | 0 | 0 | Mono | Mono | | 16767517 | | | | 3171 |
| 3174 | IV | 819 | 0 | 8 | 279 | 0 | 4 | Mono | Mono | | 16772143 | | | | 3174 |
| 3177 | IV | 827 | 0 | 0 | 283 | 0 | 0 | Mono | Mono | | 16774007 | | | | 3177 |
| 3178 | II | 826 | 1 | 0 | 283 | 0 | 0 | M9+ | 1 | | 16778194 | | | | 3178 |
| 3180 | IV | 827 | 0 | 0 | 283 | 0 | 0 | Mono | Mono | | 16786382 | | | | 3180 |
| 3181 | IV | 827 | 0 | 0 | 283 | 0 | 0 | Mono | Mono | | 16788383 | | | | 3181 |
| 3182 | IV | 827 | 0 | 0 | 283 | 0 | 0 | Mono | Mono | | 16788427 | | | | 3182 |
| 3183 | IV | 827 | 0 | 0 | 283 | 0 | 0 | Mono | Mono | | 16788431 | | | | 3183 |
| 3184 | IV | 827 | 0 | 0 | 283 | 0 | 0 | Mono | Mono | | 16788539 | | | | 3184 |
| 3185 | IV | 821 | 0 | 6 | 281 | 0 | 2 | Mono | Mono | | 16788581 | | | | 3185 |
| 3186 | IV | 827 | 0 | 0 | 283 | 0 | 0 | Mono | Mono | | 16788665 | | | | 3186 |
| 3187 | IV | 827 | 0 | 0 | 283 | 0 | 0 | Mono | Mono | | 16801352 | | | | 3187 |
| 3189 | I | 827 | 0 | 0 | 260 | 23 | 0 | M9- | 60 – 66 | | 16803249 | | | | 3189 |
| 3190 | IV | 827 | 0 | 0 | 283 | 0 | 0 | Mono | Mono | | 16804852 | | | | 3190 |
| 3191 | IV | 827 | 0 | 0 | 283 | 0 | 0 | Mono | Mono | | 16806198 | | | | 3191 |
| 3194 | IV | 827 | 0 | 0 | 283 | 0 | 0 | Mono | Mono | | 16809337 | | | | 3194 |
| 3195 | II | 825 | 2 | 0 | 283 | 0 | 0 | M9+ | 15 | | 16809478 | | | | 3195 |
| 3197 | IV | 827 | 0 | 0 | 283 | 0 | 0 | Mono | Mono | | 16817026 | | | | 3197 |
| 3198 | IV | 827 | 0 | 0 | 283 | 0 | 0 | Mono | Mono | | 16817030 | | | | 3198 |
| 3201 | IV | 827 | 0 | 0 | 283 | 0 | 0 | Mono | Mono | | 16819308 | | | | 3201 |
| 3202 | IV | 827 | 0 | 0 | 283 | 0 | 0 | Mono | Mono | | 16819471 | | | | 3202 |
| 3203 | IV | 827 | 0 | 0 | 283 | 0 | 0 | Mono | Mono | | 16820469 | | | | 3203 |
| 3206 | IV | 827 | 0 | 0 | 283 | 0 | 0 | Mono | Mono | | 16826015 | | | | 3206 |
| 3207 | IV | 827 | 0 | 0 | 283 | 0 | 0 | Mono | Mono | | 16831338 | | | | 3207 |
| 3210 | IV | 827 | 0 | 0 | 283 | 0 | 0 | Mono | Mono | | 16834840 | | | | 3210 |
| 3211 | IV | 827 | 0 | 0 | 283 | 0 | 0 | Mono | Mono | | 16834855 | | | | 3211 |
| 3212 | IV | 827 | 0 | 0 | 283 | 0 | 0 | Mono | Mono | | 16835022 | | | | 3212 |
| 3213 | IV | 820 | 0 | 7 | 283 | 0 | 0 | Mono | Mono | | 16835087 | | | | 3213 |
| 3218 | IV | 825 | 0 | 2 | 282 | 0 | 1 | Mono | Mono | | 16840817 | | | | 3218 |
| 3222 | IV | 827 | 0 | 0 | 283 | 0 | 0 | Mono | Mono | | 16849860 | | | | 3222 |
| 3223 | IV | 827 | 0 | 0 | 283 | 0 | 0 | Mono | Mono | | 16849994 | | | | 3223 |
| 3226 | IV | 825 | 0 | 2 | 283 | 0 | 0 | Mono | Mono | | 16854794 | | | | 3226 |
| 3227 | IV | 826 | 0 | 1 | 283 | 0 | 0 | Mono | Mono | | 16855080 | | | | 3227 |
| 3228 | II | 825 | 2 | 0 | 283 | 0 | 0 | M9+ | 1 – 2 | | 16855781 | | | | 3228 |
| 3230 | II | 826 | 1 | 0 | 283 | 0 | 0 | M9+ | 2 | | 16857131 | | | | 3230 |
| 3231 | IV | 827 | 0 | 0 | 283 | 0 | 0 | Mono | Mono | | 16859582 | | | | 3231 |
| 3236 | IV | 827 | 0 | 0 | 283 | 0 | 0 | Mono | Mono | | 16870008 | | | | 3236 |
| 3243 | I | 827 | 0 | 0 | 247 | 36 | 0 | M9- | 2 – 10 | | 16880603 | | | | 3243 |
| 3244 | IV | 827 | 0 | 0 | 283 | 0 | 0 | Mono | Mono | | 16880730 | | | | 3244 |
| 3246 | I | 827 | 0 | 0 | 247 | 36 | 0 | M9- | 2 – 10 | | 16881142 | | | | 3246 |
| 3247 | IV | 827 | 0 | 0 | 283 | 0 | 0 | Mono | Mono | | 16883075 | | | | 3247 |
| 3248 | IV | 827 | 0 | 0 | 283 | 0 | 0 | Mono | Mono | | 16883773 | | | | 3248 |
| 3249 | IV | 827 | 0 | 0 | 283 | 0 | 0 | Mono | Mono | | 16886911 | | | | 3249 |
| 3252 | II | 826 | 1 | 0 | 283 | 0 | 0 | M9+ | 2 | | 16894620 | | | | 3252 |
| 3255 | IV | 827 | 0 | 0 | 283 | 0 | 0 | Mono | Mono | | 16898428 | | | | 3255 |
| 3262 | IV | 827 | 0 | 0 | 283 | 0 | 0 | Mono | Mono | | 16903692 | | | | 3262 |
| 3263 | IV | 827 | 0 | 0 | 283 | 0 | 0 | Mono | Mono | | 16904772 | | | | 3263 |
| 3264 | IV | 827 | 0 | 0 | 283 | 0 | 0 | Mono | Mono | | 16904849 | | | | 3264 |
| 3266 | IV | 827 | 0 | 0 | 283 | 0 | 0 | Mono | Mono | | 16907962 | | | | 3266 |
| 3270 | IV | 827 | 0 | 0 | 283 | 0 | 0 | Mono | Mono | | 16923335 | | | | 3270 |
| 3276 | IV | 826 | 0 | 1 | 283 | 0 | 0 | Mono | Mono | | 16937297 | | | | 3276 |
| 3277 | IV | 826 | 0 | 1 | 283 | 0 | 0 | Mono | Mono | | 16937384 | | | | 3277 |
| 3278 | IV | 827 | 0 | 0 | 283 | 0 | 0 | Mono | Mono | | 16937432 | | | | 3278 |
| 3282 | I | 824 | 3 | 0 | 252 | 31 | 0 | BOTH | | PI | 16940903 | | | | 3282 |
| 3284 | I | 823 | 4 | 0 | 0 | 283 | 0 | M9+ | 4 – 93 | | 16944053 | | | | 3284 |
| 3286 | IV | 827 | 0 | 0 | 283 | 0 | 0 | Mono | Mono | | 16947820 | | | | 3286 |
| 3291 | IV | 825 | 0 | 2 | 282 | 0 | 1 | Mono | Mono | | 16951875 | | | | 3291 |
| 3299 | IV | 826 | 0 | 1 | 283 | 0 | 0 | Mono | Mono | | 16962383 | | | | 3299 |
| 3301 | IV | 819 | 0 | 8 | 280 | 0 | 3 | Mono | Mono | | 16981140 | | | | 3301 |
| 3302 | IV | 827 | 0 | 0 | 283 | 0 | 0 | Mono | Mono | | 16981954 | | | | 3302 |
| 3303 | IV | 827 | 0 | 0 | 283 | 0 | 0 | Mono | Mono | | 16984055 | | | | 3303 |
| 3304 | IV | 827 | 0 | 0 | 283 | 0 | 0 | Mono | Mono | | 16992232 | | | | 3304 |
| 3306 | IV | 778 | 0 | 49 | 266 | 0 | 17 | Mono | Mono | | 17001055 | | | | 3306 |
| 3307 | IV | 827 | 0 | 0 | 283 | 0 | 0 | Mono | Mono | | 17004094 | | | | 3307 |
| 3310 | I | 827 | 0 | 0 | 209 | 74 | 0 | M9- | 42 – 58 | | 17012932 | | | | 3310 |
| 3313 | II | 826 | 1 | 0 | 283 | 0 | 0 | M9+ | 1 | | 17022695 | | | | 3313 |
| 3316 | IV | 827 | 0 | 0 | 283 | 0 | 0 | Mono | Mono | | 17032174 | | | | 3316 |
| 3319 | IV | 824 | 0 | 3 | 281 | 0 | 2 | Mono | Mono | | 17038316 | | | | 3319 |
| 3323 | IV | 827 | 0 | 0 | 283 | 0 | 0 | Mono | Mono | | 17048384 | | | | 3323 |
| 3324 | III | 824 | 1 | 2 | 283 | 0 | 0 | M9+ | 2 | | 17048682 | | | | 3324 |
| 3328 | I | 827 | 0 | 0 | 246 | 37 | 0 | M9- | 1 – 10 | | 17054936 | | | | 3328 |
| 3336 | I | 824 | 3 | 0 | 272 | 11 | 0 | BOTH | | PI | 17106445 | | | | 3336 |
| 3341 | I | 827 | 0 | 0 | 246 | 37 | 0 | M9- | 1 – 10 | | 17115298 | | | | 3341 |
| 3346 | IV | 827 | 0 | 0 | 283 | 0 | 0 | Mono | Mono | | 17135154 | | | | 3346 |
| 3352 | IV | 827 | 0 | 0 | 283 | 0 | 0 | Mono | Mono | | 17141732 | | | | 3352 |
| 3354 | II | 826 | 1 | 0 | 283 | 0 | 0 | M9+ | 1 | | 17151126 | | | | 3354 |
| 3362 | IV | 826 | 0 | 1 | 282 | 0 | 1 | Mono | Mono | | 17182858 | | | | 3362 |
| 3364 | IV | 825 | 0 | 2 | 282 | 0 | 1 | Mono | Mono | | 17190810 | | | | 3364 |
| 3366 | IV | 827 | 0 | 0 | 283 | 0 | 0 | Mono | Mono | | 17191930 | | | | 3366 |
| 3369 | IV | 827 | 0 | 0 | 283 | 0 | 0 | Mono | Mono | | 17197796 | | | | 3369 |
| 3375 | IV | 827 | 0 | 0 | 283 | 0 | 0 | Mono | Mono | | 17209692 | | | | 3375 |

| | | | M9+ | | | M9- | | | | | Build 37 | | ISOGG | | |
|---|---|---|---|---|---|---|---|---|---|---|---|---|---|---|---|
| List | Stage | 0 | 1 | . | 0 | 1 | . | M9? | Clade(s) | Prob. | Position | Name | Subgroup Name | Amb. | List |
| 3376 | IV | 827 | 0 | 0 | 283 | 0 | 0 | Mono | Mono | | 17209790 | | | | 3376 |
| 3380 | IV | 827 | 0 | 0 | 283 | 0 | 0 | Mono | Mono | | 17231905 | | | | 3380 |
| 3384 | I | 825 | 2 | 0 | 281 | 2 | 0 | BOTH | | PI | 17239322 | | | | 3384 |
| 3386 | IV | 827 | 0 | 0 | 283 | 0 | 0 | Mono | Mono | | 17242145 | | | | 3386 |
| 3387 | IV | 826 | 0 | 1 | 283 | 0 | 0 | Mono | Mono | | 17242819 | | | | 3387 |
| 3390 | IV | 827 | 0 | 0 | 283 | 0 | 0 | Mono | Mono | | 17252369 | | | | 3390 |
| 3391 | IV | 827 | 0 | 0 | 283 | 0 | 0 | Mono | Mono | | 17252385 | | | | 3391 |
| 3400 | IV | 827 | 0 | 0 | 283 | 0 | 0 | Mono | Mono | | 17287672 | | | | 3400 |
| 3406 | IV | 821 | 0 | 6 | 282 | 0 | 1 | Mono | Mono | | 17298476 | | | | 3406 |
| 3408 | III | 822 | 4 | 1 | 0 | 282 | 1 | M9+ | 4 – 93 | | 17301401 | | | | 3408 |
| 3410 | IV | 827 | 0 | 0 | 283 | 0 | 0 | Mono | Mono | | 17304846 | | | | 3410 |
| 3411 | IV | 827 | 0 | 0 | 283 | 0 | 0 | Mono | Mono | | 17304901 | | | | 3411 |
| 3412 | II | 827 | 0 | 0 | 281 | 2 | 0 | M9- | 48 | | 17307191 | | | | 3412 |
| 3413 | IV | 827 | 0 | 0 | 283 | 0 | 0 | Mono | Mono | | 17310145 | | | | 3413 |
| 3415 | IV | 827 | 0 | 0 | 283 | 0 | 0 | Mono | Mono | | 17312195 | | | | 3415 |
| 3417 | IV | 827 | 0 | 0 | 283 | 0 | 0 | Mono | Mono | | 17321074 | | | | 3417 |
| 3420 | I | 827 | 0 | 0 | 262 | 21 | 0 | M9- | 11 – 20 | | 17329861 | | | | 3420 |
| 3425 | IV | 826 | 0 | 1 | 283 | 0 | 0 | Mono | Mono | | 17343510 | | | | 3425 |
| 3430 | IV | 827 | 0 | 0 | 281 | 0 | 2 | Mono | Mono | | 17358315 | | | | 3430 |
| 3431 | IV | 826 | 0 | 1 | 283 | 0 | 0 | Mono | Mono | | 17358410 | | | | 3431 |
| 3435 | IV | 827 | 0 | 0 | 283 | 0 | 0 | Mono | Mono | | 17380269 | | | | 3435 |
| 3436 | IV | 827 | 0 | 0 | 283 | 0 | 0 | Mono | Mono | | 17384648 | | | | 3436 |
| 3438 | IV | 827 | 0 | 0 | 283 | 0 | 0 | Mono | Mono | | 17389379 | | | | 3438 |
| 3444 | IV | 0 | 825 | 2 | 0 | 283 | 0 | Mono | Mono | | 17397645 | | | | 3444 |
| 3447 | IV | 827 | 0 | 0 | 283 | 0 | 0 | Mono | Mono | | 17409375 | | | | 3447 |
| 3448 | I | 827 | 0 | 0 | 246 | 37 | 0 | M9- | 1 – 10 | | 17412198 | | | | 3448 |
| 3452 | IV | 827 | 0 | 0 | 283 | 0 | 0 | Mono | Mono | | 17418389 | | | | 3452 |
| 3460 | IV | 827 | 0 | 0 | 283 | 0 | 0 | Mono | Mono | | 17427710 | | | | 3460 |
| 3463 | IV | 827 | 0 | 0 | 283 | 0 | 0 | Mono | Mono | | 17446588 | | | | 3463 |
| 3464 | IV | 827 | 0 | 0 | 283 | 0 | 0 | Mono | Mono | | 17446611 | | | | 3464 |
| 3465 | IV | 827 | 0 | 0 | 283 | 0 | 0 | Mono | Mono | | 17446768 | | | | 3465 |
| 3466 | IV | 827 | 0 | 0 | 283 | 0 | 0 | Mono | Mono | | 17447638 | | | | 3466 |
| 3469 | IV | 827 | 0 | 0 | 283 | 0 | 0 | Mono | Mono | | 17454580 | | | | 3469 |
| 3474 | II | 826 | 1 | 0 | 283 | 0 | 0 | M9+ | 1 | | 17468036 | | | | 3474 |
| 3476 | II | 826 | 1 | 0 | 283 | 0 | 0 | M9+ | 1 | | 17472151 | | | | 3476 |
| 3490 | IV | 822 | 0 | 5 | 283 | 0 | 0 | Mono | Mono | | 17509926 | | | | 3490 |
| 3491 | IV | 826 | 0 | 1 | 283 | 0 | 0 | Mono | Mono | | 17509929 | | | | 3491 |
| 3495 | IV | 827 | 0 | 0 | 283 | 0 | 0 | Mono | Mono | | 17512003 | | | | 3495 |
| 3506 | IV | 827 | 0 | 0 | 283 | 0 | 0 | Mono | Mono | | 17537643 | | | | 3506 |
| 3509 | IV | 827 | 0 | 0 | 283 | 0 | 0 | Mono | Mono | | 17550661 | | | | 3509 |
| 3513 | IV | 827 | 0 | 0 | 283 | 0 | 0 | Mono | Mono | | 17558118 | | | | 3513 |
| 3515 | IV | 826 | 0 | 1 | 283 | 0 | 0 | Mono | Mono | | 17560292 | | | | 3515 |
| 3518 | III | 819 | 1 | 7 | 281 | 0 | 2 | M9+ | 1 | | 17566936 | | | | 3518 |
| 3519 | IV | 827 | 0 | 0 | 283 | 0 | 0 | Mono | Mono | | 17569251 | | | | 3519 |
| 3521 | IV | 827 | 0 | 0 | 283 | 0 | 0 | Mono | Mono | | 17571473 | | | | 3521 |
| 3523 | IV | 827 | 0 | 0 | 283 | 0 | 0 | Mono | Mono | | 17571623 | | | | 3523 |
| 3525 | II | 826 | 1 | 0 | 283 | 0 | 0 | M9+ | 1 | | 17572846 | | | | 3525 |
| 3526 | IV | 827 | 0 | 0 | 283 | 0 | 0 | Mono | Mono | | 17576942 | | | | 3526 |
| 3529 | IV | 827 | 0 | 0 | 283 | 0 | 0 | Mono | Mono | | 17588338 | | | | 3529 |
| 3530 | IV | 826 | 0 | 1 | 283 | 0 | 0 | Mono | Mono | | 17588597 | | | | 3530 |
| 3531 | IV | 826 | 0 | 1 | 283 | 0 | 0 | Mono | Mono | | 17589340 | | | | 3531 |
| 3540 | IV | 827 | 0 | 0 | 283 | 0 | 0 | Mono | Mono | | 17601992 | | | | 3540 |
| 3545 | IV | 827 | 0 | 0 | 283 | 0 | 0 | Mono | Mono | | 17609833 | | | | 3545 |
| 3555 | IV | 827 | 0 | 0 | 283 | 0 | 0 | Mono | Mono | | 17650073 | | | | 3555 |
| 3556 | II | 825 | 2 | 0 | 283 | 0 | 0 | M9+ | 68 | | 17651573 | | | | 3556 |
| 3562 | I | 827 | 0 | 0 | 0 | 283 | 0 | M9+/- | M9+/- | | 17660074 | | | | 3562 |
| 3570 | IV | 827 | 0 | 0 | 283 | 0 | 0 | Mono | Mono | | 17672931 | | | | 3570 |
| 3577 | I | 827 | 0 | 0 | 246 | 37 | 0 | M9- | 1 – 10 | | 17689245 | | | | 3577 |
| 3581 | IV | 827 | 0 | 0 | 283 | 0 | 0 | Mono | Mono | | 17693443 | | | | 3581 |
| 3582 | IV | 827 | 0 | 0 | 283 | 0 | 0 | Mono | Mono | | 17695251 | | | | 3582 |
| 3586 | IV | 820 | 0 | 7 | 282 | 0 | 1 | Mono | Mono | | 17714426 | | | | 3586 |
| 3588 | IV | 827 | 0 | 0 | 283 | 0 | 0 | Mono | Mono | | 17717953 | | | | 3588 |
| 3589 | IV | 827 | 0 | 0 | 283 | 0 | 0 | Mono | Mono | | 17730754 | | | | 3589 |
| 3594 | IV | 827 | 0 | 0 | 283 | 0 | 0 | Mono | Mono | | 17742219 | | | | 3594 |
| 3601 | II | 825 | 2 | 0 | 283 | 0 | 0 | M9+ | 1 – 2 | | 17756426 | | | | 3601 |
| 3605 | IV | 827 | 0 | 0 | 283 | 0 | 0 | Mono | Mono | | 17761948 | | | | 3605 |
| 3608 | IV | 827 | 0 | 0 | 283 | 0 | 0 | Mono | Mono | | 17765230 | | | | 3608 |
| 3612 | IV | 826 | 0 | 1 | 283 | 0 | 0 | Mono | Mono | | 17768046 | | | | 3612 |
| 3613 | IV | 827 | 0 | 0 | 283 | 0 | 0 | Mono | Mono | | 17768441 | | | | 3613 |
| 3617 | II | 826 | 1 | 0 | 283 | 0 | 0 | M9+ | 1 | | 17779649 | | | | 3617 |
| 3622 | IV | 827 | 0 | 0 | 283 | 0 | 0 | Mono | Mono | | 17789323 | | | | 3622 |
| 3632 | IV | 827 | 0 | 0 | 283 | 0 | 0 | Mono | Mono | | 17805633 | | | | 3632 |
| 3638 | IV | 825 | 0 | 2 | 283 | 0 | 0 | Mono | Mono | | 17815397 | | | | 3638 |
| 3639 | IV | 827 | 0 | 0 | 283 | 0 | 0 | Mono | Mono | | 17816507 | | | | 3639 |
| 3640 | IV | 827 | 0 | 0 | 283 | 0 | 0 | Mono | Mono | | 17816694 | | | | 3640 |
| 3643 | IV | 827 | 0 | 0 | 283 | 0 | 0 | Mono | Mono | | 17820684 | | | | 3643 |
| 3654 | IV | 0 | 826 | 1 | 0 | 282 | 1 | Mono | Mono | | 17844781 | | | | 3654 |
| 3660 | IV | 0 | 827 | 0 | 0 | 283 | 0 | Mono | Mono | | 17860729 | | | | 3660 |
| 3661 | I | 823 | 4 | 0 | 281 | 2 | 0 | BOTH | | PI | 17861975 | | | | 3661 |
| 3667 | III | 822 | 1 | 4 | 283 | 0 | 0 | M9+ | 1 | | 17871401 | | | | 3667 |
| 3668 | IV | 827 | 0 | 0 | 283 | 0 | 0 | Mono | Mono | | 17872179 | | | | 3668 |
| 3669 | II | 826 | 1 | 0 | 283 | 0 | 0 | M9+ | 2 | | 17879035 | | | | 3669 |
| 3673 | IV | 826 | 0 | 1 | 283 | 0 | 0 | Mono | Mono | | 17881712 | | | | 3673 |
| 3689 | IV | 827 | 0 | 0 | 283 | 0 | 0 | Mono | Mono | | 17913958 | | | | 3689 |
| 3691 | IV | 827 | 0 | 0 | 283 | 0 | 0 | Mono | Mono | | 17916397 | | | | 3691 |
| 3692 | IV | 827 | 0 | 0 | 283 | 0 | 0 | Mono | Mono | | 17916624 | | | | 3692 |
| 3709 | IV | 827 | 0 | 0 | 283 | 0 | 0 | Mono | Mono | | 17944100 | | | | 3709 |
| 3710 | IV | 827 | 0 | 0 | 283 | 0 | 0 | Mono | Mono | | 17944430 | | | | 3710 |
| 3715 | IV | 0 | 827 | 0 | 0 | 283 | 0 | Mono | Mono | | 17951267 | | | | 3715 |
| 3721 | IV | 827 | 0 | 0 | 283 | 0 | 0 | Mono | Mono | | 17974575 | | | | 3721 |
| 3726 | IV | 827 | 0 | 0 | 283 | 0 | 0 | Mono | Mono | | 17985841 | | | | 3726 |
| 3746 | IV | 827 | 0 | 0 | 283 | 0 | 0 | Mono | Mono | | 18037104 | | | | 3746 |
| 3747 | IV | 827 | 0 | 0 | 283 | 0 | 0 | Mono | Mono | | 18038953 | | | | 3747 |
| 3748 | IV | 827 | 0 | 0 | 283 | 0 | 0 | Mono | Mono | | 18038955 | | | | 3748 |
| 3753 | IV | 827 | 0 | 0 | 283 | 0 | 0 | Mono | Mono | | 18050087 | | | | 3753 |
| 3754 | IV | 827 | 0 | 0 | 283 | 0 | 0 | Mono | Mono | | 18051212 | | | | 3754 |

| | | M9+ | | | M9- | | | | | | Build 37 | | ISOGG | | |
|---|---|---|---|---|---|---|---|---|---|---|---|---|---|---|---|
| List | Stage | 0 | 1 | . | 0 | 1 | . | M9? | Clade(s) | Prob. | Position | Name | Subgroup Name | Amb. | List |
| 3761 | IV | 827 | 0 | 0 | 283 | 0 | 0 | Mono | Mono | | 18064300 | | | | 3761 |
| 3766 | IV | 827 | 0 | 0 | 283 | 0 | 0 | Mono | Mono | | 18068361 | | | | 3766 |
| 3784 | IV | 827 | 0 | 0 | 283 | 0 | 0 | Mono | Mono | | 18099341 | | | | 3784 |
| 3785 | I | 827 | 0 | 0 | 246 | 37 | 0 | M9- | 1 – 10 | | 18103101 | | | | 3785 |
| 3786 | IV | 827 | 0 | 0 | 283 | 0 | 0 | Mono | Mono | | 18104102 | | | | 3786 |
| 3794 | I | 827 | 0 | 0 | 246 | 37 | 0 | M9- | 1 – 10 | | 18121600 | | | | 3794 |
| 3795 | IV | 822 | 0 | 5 | 282 | 0 | 1 | Mono | Mono | | 18123819 | | | | 3795 |
| 3798 | IV | 827 | 0 | 0 | 283 | 0 | 0 | Mono | Mono | | 18129736 | | | | 3798 |
| 3802 | IV | 827 | 0 | 0 | 283 | 0 | 0 | Mono | Mono | | 18147185 | | | | 3802 |
| 3811 | II | 826 | 1 | 0 | 283 | 0 | 0 | M9+ | 1 | | 18175761 | | | | 3811 |
| 3824 | III | 823 | 0 | 4 | 282 | 1 | 0 | M9- | 21 | | 18207718 | | | | 3824 |
| 3825 | IV | 818 | 0 | 9 | 281 | 0 | 2 | Mono | Mono | | 18207719 | | | | 3825 |
| 3827 | II | 826 | 1 | 0 | 283 | 0 | 0 | M9+ | 2 | | 18213440 | | | | 3827 |
| 3828 | IV | 825 | 0 | 2 | 283 | 0 | 0 | Mono | Mono | | 18223950 | | | | 3828 |
| 3829 | IV | 827 | 0 | 0 | 283 | 0 | 0 | Mono | Mono | | 18224142 | | | | 3829 |
| 3830 | III | 816 | 1 | 10 | 279 | 0 | 4 | M9+ | 1 | | 18233001 | | | | 3830 |
| 3833 | IV | 827 | 0 | 0 | 282 | 0 | 1 | Mono | Mono | | 18238370 | | | | 3833 |
| 3838 | IV | 827 | 0 | 0 | 283 | 0 | 0 | Mono | Mono | | 18259415 | | | | 3838 |
| 3839 | IV | 827 | 0 | 0 | 283 | 0 | 0 | Mono | Mono | | 18264657 | | | | 3839 |
| 3843 | II | 826 | 1 | 0 | 283 | 0 | 0 | M9+ | 2 | | 18387196 | | | | 3843 |
| 3846 | III | 826 | 0 | 1 | 260 | 23 | 0 | M9- | 60 – 66 | | 18391106 | | | | 3846 |
| 3854 | IV | 826 | 0 | 1 | 283 | 0 | 0 | Mono | Mono | | 18401414 | | | | 3854 |
| 3859 | IV | 827 | 0 | 0 | 283 | 0 | 0 | Mono | Mono | | 18417772 | | | | 3859 |
| 3866 | IV | 827 | 0 | 0 | 283 | 0 | 0 | Mono | Mono | | 18546164 | | | | 3866 |
| 3867 | III | 826 | 1 | 0 | 282 | 0 | 1 | M9+ | 1 | | 18548828 | | | | 3867 |
| 3868 | IV | 827 | 0 | 0 | 272 | 0 | 11 | Mono | Mono | | 18551135 | | | | 3868 |
| 3870 | IV | 827 | 0 | 0 | 283 | 0 | 0 | Mono | Mono | | 18553075 | | | | 3870 |
| 3877 | IV | 827 | 0 | 0 | 283 | 0 | 0 | Mono | Mono | | 18567437 | | | | 3877 |
| 3887 | IV | 826 | 0 | 1 | 283 | 0 | 0 | Mono | Mono | | 18584283 | | | | 3887 |
| 3894 | III | 827 | 0 | 0 | 245 | 37 | 1 | M9- | 1 – 10 | | 18596847 | | | | 3894 |
| 3897 | IV | 809 | 0 | 18 | 277 | 0 | 6 | Mono | Mono | | 18600862 | | | | 3897 |
| 3899 | IV | 827 | 0 | 0 | 283 | 0 | 0 | Mono | Mono | | 18608096 | | | | 3899 |
| 3900 | II | 826 | 1 | 0 | 283 | 0 | 0 | M9+ | 2 | | 18612761 | | | | 3900 |
| 3901 | III | 820 | 4 | 3 | 282 | 1 | 0 | BOTH | | PI | 18612789 | | | | 3901 |
| 3917 | IV | 827 | 0 | 0 | 283 | 0 | 0 | Mono | Mono | | 18637416 | | | | 3917 |
| 3927 | IV | 827 | 0 | 0 | 283 | 0 | 0 | Mono | Mono | | 18663485 | | | | 3927 |
| 3928 | IV | 826 | 0 | 1 | 283 | 0 | 0 | Mono | Mono | | 18666823 | | | | 3928 |
| 3933 | IV | 827 | 0 | 0 | 283 | 0 | 0 | Mono | Mono | | 18675236 | | | | 3933 |
| 3938 | II | 826 | 1 | 0 | 283 | 0 | 0 | M9+ | 2 | | 18696215 | | | | 3938 |
| 3940 | IV | 827 | 0 | 0 | 283 | 0 | 0 | Mono | Mono | | 18706036 | | | | 3940 |
| 3942 | IV | 827 | 0 | 0 | 283 | 0 | 0 | Mono | Mono | | 18710853 | | | | 3942 |
| 3948 | IV | 827 | 0 | 0 | 282 | 0 | 1 | Mono | Mono | | 18725936 | | | | 3948 |
| 3953 | IV | 827 | 0 | 0 | 283 | 0 | 0 | Mono | Mono | | 18742602 | | | | 3953 |
| 3956 | I | 827 | 0 | 0 | 262 | 21 | 0 | M9- | 11 – 20 | | 18744995 | | | | 3956 |
| 3957 | III | 822 | 0 | 5 | 261 | 21 | 1 | M9- | 11 – 20 | | 18744996 | | | | 3957 |
| 3968 | IV | 827 | 0 | 0 | 283 | 0 | 0 | Mono | Mono | | 18771855 | | | | 3968 |
| 3972 | I | 827 | 0 | 0 | 246 | 37 | 0 | M9- | 1 – 10 | | 18776629 | | | | 3972 |
| 3974 | IV | 827 | 0 | 0 | 283 | 0 | 0 | Mono | Mono | | 18782266 | | | | 3974 |
| 3976 | IV | 827 | 0 | 0 | 283 | 0 | 0 | Mono | Mono | | 18785737 | | | | 3976 |
| 3983 | I | 824 | 3 | 0 | 0 | 283 | 0 | M9+ | 3 – 93 | | 18801906 | | | | 3983 |
| 3985 | IV | 827 | 0 | 0 | 283 | 0 | 0 | Mono | Mono | | 18803875 | | | | 3985 |
| 3986 | IV | 826 | 0 | 1 | 283 | 0 | 0 | Mono | Mono | | 18803914 | | | | 3986 |
| 3987 | IV | 827 | 0 | 0 | 283 | 0 | 0 | Mono | Mono | | 18803990 | | | | 3987 |
| 3991 | IV | 827 | 0 | 0 | 283 | 0 | 0 | Mono | Mono | | 18817127 | | | | 3991 |
| 3992 | IV | 827 | 0 | 0 | 283 | 0 | 0 | Mono | Mono | | 18817521 | | | | 3992 |
| 3993 | IV | 827 | 0 | 0 | 283 | 0 | 0 | Mono | Mono | | 18817522 | | | | 3993 |
| 3995 | IV | 824 | 0 | 3 | 283 | 0 | 0 | Mono | Mono | | 18820048 | | | | 3995 |
| 3996 | IV | 827 | 0 | 0 | 283 | 0 | 0 | Mono | Mono | | 18821912 | | | | 3996 |
| 4008 | IV | 827 | 0 | 0 | 283 | 0 | 0 | Mono | Mono | | 18843088 | | | | 4008 |
| 4010 | IV | 826 | 0 | 1 | 283 | 0 | 0 | Mono | Mono | | 18847802 | | | | 4010 |
| 4011 | IV | 827 | 0 | 0 | 283 | 0 | 0 | Mono | Mono | | 18847810 | | | | 4011 |
| 4016 | IV | 820 | 0 | 7 | 282 | 0 | 1 | Mono | Mono | | 18867706 | | | | 4016 |
| 4018 | IV | 827 | 0 | 0 | 282 | 0 | 1 | Mono | Mono | | 18868481 | | | | 4018 |
| 4019 | IV | 827 | 0 | 0 | 283 | 0 | 0 | Mono | Mono | | 18871216 | | | | 4019 |
| 4020 | I | 827 | 0 | 0 | 246 | 37 | 0 | M9- | 1 – 10 | | 18880574 | | | | 4020 |
| 4027 | IV | 827 | 0 | 0 | 283 | 0 | 0 | Mono | Mono | | 18893048 | | | | 4027 |
| 4028 | IV | 827 | 0 | 0 | 283 | 0 | 0 | Mono | Mono | | 18893538 | | | | 4028 |
| 4029 | IV | 826 | 0 | 1 | 283 | 0 | 0 | Mono | Mono | | 18893800 | | | | 4029 |
| 4031 | IV | 827 | 0 | 0 | 283 | 0 | 0 | Mono | Mono | | 18898795 | | | | 4031 |
| 4034 | II | 826 | 1 | 0 | 283 | 0 | 0 | M9+ | 2 | | 18903362 | | | | 4034 |
| 4041 | IV | 825 | 0 | 2 | 283 | 0 | 0 | Mono | Mono | | 18916849 | | | | 4041 |
| 4052 | IV | 827 | 0 | 0 | 283 | 0 | 0 | Mono | Mono | | 18940129 | | | | 4052 |
| 4054 | IV | 827 | 0 | 0 | 283 | 0 | 0 | Mono | Mono | | 18940964 | | | | 4054 |
| 4059 | IV | 827 | 0 | 0 | 283 | 0 | 0 | Mono | Mono | | 18954148 | | | | 4059 |
| 4063 | IV | 827 | 0 | 0 | 283 | 0 | 0 | Mono | Mono | | 18958586 | | | | 4063 |
| 4068 | IV | 827 | 0 | 0 | 283 | 0 | 0 | Mono | Mono | | 18963337 | | | | 4068 |
| 4077 | IV | 827 | 0 | 0 | 283 | 0 | 0 | Mono | Mono | | 18992566 | | | | 4077 |
| 4093 | IV | 827 | 0 | 0 | 283 | 0 | 0 | Mono | Mono | | 19037067 | | | | 4093 |
| 4096 | III | 824 | 0 | 3 | 231 | 51 | 1 | M9- | | PI | 19044813 | | | | 4096 |
| 4104 | IV | 827 | 0 | 0 | 283 | 0 | 0 | Mono | Mono | | 19053889 | | | | 4104 |
| 4110 | IV | 827 | 0 | 0 | 283 | 0 | 0 | Mono | Mono | | 19060742 | | | | 4110 |
| 4112 | IV | 827 | 0 | 0 | 283 | 0 | 0 | Mono | Mono | | 19065314 | | | | 4112 |
| 4115 | I | 827 | 0 | 0 | 246 | 37 | 0 | M9- | 1 – 10 | | 19077394 | | | | 4115 |
| 4127 | IV | 826 | 0 | 1 | 283 | 0 | 0 | Mono | Mono | | 19103154 | | | | 4127 |
| 4131 | IV | 827 | 0 | 0 | 283 | 0 | 0 | Mono | Mono | | 19117451 | | | | 4131 |
| 4137 | IV | 825 | 0 | 2 | 283 | 0 | 0 | Mono | Mono | | 19126462 | | | | 4137 |
| 4138 | IV | 827 | 0 | 0 | 283 | 0 | 0 | Mono | Mono | | 19128168 | | | | 4138 |
| 4150 | IV | 827 | 0 | 0 | 283 | 0 | 0 | Mono | Mono | | 19150014 | | | | 4150 |
| 4151 | IV | 827 | 0 | 0 | 282 | 0 | 1 | Mono | Mono | | 19150774 | | | | 4151 |
| 4154 | II | 826 | 1 | 0 | 283 | 0 | 0 | M9+ | 2 | | 19151936 | | | | 4154 |
| 4156 | II | 826 | 1 | 0 | 283 | 0 | 0 | M9+ | 1 | | 19158022 | | | | 4156 |
| 4160 | IV | 827 | 0 | 0 | 283 | 0 | 0 | Mono | Mono | | 19165461 | | | | 4160 |
| 4178 | IV | 827 | 0 | 0 | 283 | 0 | 0 | Mono | Mono | | 19200208 | | | | 4178 |
| 4181 | IV | 827 | 0 | 0 | 283 | 0 | 0 | Mono | Mono | | 19206845 | | | | 4181 |
| 4191 | IV | 827 | 0 | 0 | 283 | 0 | 0 | Mono | Mono | | 19224007 | | | | 4191 |
| 4193 | IV | 827 | 0 | 0 | 283 | 0 | 0 | Mono | Mono | | 19230498 | | | | 4193 |

| | | M9+ | | | M9- | | | | | | Build 37 | | ISOGG | | |
|---|---|---|---|---|---|---|---|---|---|---|---|---|---|---|---|
| List | Stage | 0 | 1 | . | 0 | 1 | . | M9? | Clade(s) | Prob. | Position | Name | Subgroup Name | Amb. | List |
| 4208 | IV | 826 | 0 | 1 | 283 | 0 | 0 | Mono | Mono | | 19244931 | | | | 4208 |
| 4211 | II | 826 | 1 | 0 | 283 | 0 | 0 | M9+ | 1 | | 19257852 | | | | 4211 |
| 4212 | IV | 827 | 0 | 0 | 283 | 0 | 0 | Mono | Mono | | 19259569 | | | | 4212 |
| 4218 | IV | 827 | 0 | 0 | 283 | 0 | 0 | Mono | Mono | | 19272427 | | | | 4218 |
| 4223 | IV | 826 | 0 | 1 | 283 | 0 | 0 | Mono | Mono | | 19276877 | | | | 4223 |
| 4226 | IV | 827 | 0 | 0 | 283 | 0 | 0 | Mono | Mono | | 19281922 | | | | 4226 |
| 4227 | IV | 827 | 0 | 0 | 283 | 0 | 0 | Mono | Mono | | 19281932 | | | | 4227 |
| 4229 | IV | 827 | 0 | 0 | 283 | 0 | 0 | Mono | Mono | | 19285054 | | | | 4229 |
| 4232 | IV | 826 | 0 | 1 | 283 | 0 | 0 | Mono | Mono | | 19292547 | | | | 4232 |
| 4237 | IV | 827 | 0 | 0 | 283 | 0 | 0 | Mono | Mono | | 19306942 | | | | 4237 |
| 4242 | IV | 827 | 0 | 0 | 283 | 0 | 0 | Mono | Mono | | 19320885 | | | | 4242 |
| 4245 | IV | 827 | 0 | 0 | 283 | 0 | 0 | Mono | Mono | | 19332760 | | | | 4245 |
| 4254 | I | 827 | 0 | 0 | 0 | 283 | 0 | M9+/- | M9+/- | | 19360192 | | | | 4254 |
| 4255 | IV | 827 | 0 | 0 | 283 | 0 | 0 | Mono | Mono | | 19360642 | | | | 4255 |
| 4256 | IV | 827 | 0 | 0 | 283 | 0 | 0 | Mono | Mono | | 19370523 | | | | 4256 |
| 4258 | II | 826 | 1 | 0 | 283 | 0 | 0 | M9+ | 2 | | 19371568 | | | | 4258 |
| 4259 | I | 790 | 37 | 0 | 283 | 0 | 0 | M9+ | | PI | 19372386 | | | | 4259 |
| 4266 | IV | 813 | 0 | 14 | 280 | 0 | 3 | Mono | Mono | | 19383158 | | | | 4266 |
| 4274 | IV | 827 | 0 | 0 | 282 | 0 | 1 | Mono | Mono | | 19399252 | | | | 4274 |
| 4282 | IV | 825 | 0 | 2 | 282 | 0 | 1 | Mono | Mono | | 19422325 | | | | 4282 |
| 4286 | IV | 825 | 0 | 2 | 283 | 0 | 0 | Mono | Mono | | 19443641 | | | | 4286 |
| 4288 | III | 825 | 1 | 1 | 283 | 0 | 0 | M9+ | 1 | | 19445302 | | | | 4288 |
| 4289 | IV | 827 | 0 | 0 | 283 | 0 | 0 | Mono | Mono | | 19446121 | | | | 4289 |
| 4295 | IV | 825 | 0 | 2 | 282 | 0 | 1 | Mono | Mono | | 19478026 | | | | 4295 |
| 4298 | III | 825 | 1 | 1 | 283 | 0 | 0 | M9+ | 1 | | 19484499 | | | | 4298 |
| 4305 | IV | 827 | 0 | 0 | 283 | 0 | 0 | Mono | Mono | | 19496663 | | | | 4305 |
| 4307 | IV | 827 | 0 | 0 | 283 | 0 | 0 | Mono | Mono | | 19500150 | | | | 4307 |
| 4308 | IV | 827 | 0 | 0 | 273 | 0 | 10 | Mono | Mono | | 19501311 | | | | 4308 |
| 4312 | IV | 827 | 0 | 0 | 283 | 0 | 0 | Mono | Mono | | 19510566 | | | | 4312 |
| 4315 | IV | 827 | 0 | 0 | 282 | 0 | 1 | Mono | Mono | | 19518344 | | | | 4315 |
| 4322 | IV | 827 | 0 | 0 | 282 | 0 | 1 | Mono | Mono | | 19550408 | | | | 4322 |
| 4323 | IV | 827 | 0 | 0 | 283 | 0 | 0 | Mono | Mono | | 19550740 | | | | 4323 |
| 4328 | IV | 827 | 0 | 0 | 283 | 0 | 0 | Mono | Mono | | 19567693 | | | | 4328 |
| 4329 | IV | 826 | 0 | 1 | 283 | 0 | 0 | Mono | Mono | | 19567718 | | | | 4329 |
| 4330 | IV | 825 | 0 | 2 | 283 | 0 | 0 | Mono | Mono | | 20804835 | | | | 4330 |
| 4334 | IV | 818 | 0 | 9 | 277 | 0 | 6 | Mono | Mono | | 20812349 | | | | 4334 |
| 4335 | IV | 827 | 0 | 0 | 283 | 0 | 0 | Mono | Mono | | 20812848 | | | | 4335 |
| 4337 | IV | 827 | 0 | 0 | 282 | 0 | 1 | Mono | Mono | | 20815225 | | | | 4337 |
| 4338 | IV | 827 | 0 | 0 | 283 | 0 | 0 | Mono | Mono | | 20815739 | | | | 4338 |
| 4339 | IV | 827 | 0 | 0 | 283 | 0 | 0 | Mono | Mono | | 20820125 | | | | 4339 |
| 4341 | IV | 827 | 0 | 0 | 283 | 0 | 0 | Mono | Mono | | 20821862 | | | | 4341 |
| 4346 | IV | 827 | 0 | 0 | 283 | 0 | 0 | Mono | Mono | | 20832009 | | | | 4346 |
| 4353 | IV | 827 | 0 | 0 | 283 | 0 | 0 | Mono | Mono | | 21045242 | | | | 4353 |
| 4354 | IV | 827 | 0 | 0 | 283 | 0 | 0 | Mono | Mono | | 21051193 | | | | 4354 |
| 4357 | IV | 827 | 0 | 0 | 283 | 0 | 0 | Mono | Mono | | 21062598 | | | | 4357 |
| 4367 | IV | 825 | 0 | 2 | 283 | 0 | 0 | Mono | Mono | | 21081179 | | | | 4367 |
| 4372 | IV | 827 | 0 | 0 | 283 | 0 | 0 | Mono | Mono | | 21094173 | | | | 4372 |
| 4378 | IV | 826 | 0 | 1 | 283 | 0 | 0 | Mono | Mono | | 21100759 | | | | 4378 |
| 4381 | II | 826 | 1 | 0 | 283 | 0 | 0 | M9+ | 1 | | 21111776 | | | | 4381 |
| 4389 | IV | 827 | 0 | 0 | 274 | 0 | 9 | Mono | Mono | | 21132000 | | | | 4389 |
| 4390 | IV | 827 | 0 | 0 | 283 | 0 | 0 | Mono | Mono | | 21138078 | | | | 4390 |
| 4397 | IV | 823 | 0 | 4 | 281 | 0 | 2 | Mono | Mono | | 21150727 | | | | 4397 |
| 4399 | IV | 827 | 0 | 0 | 282 | 0 | 1 | Mono | Mono | | 21152524 | | | | 4399 |
| 4400 | IV | 827 | 0 | 0 | 283 | 0 | 0 | Mono | Mono | | 21157531 | | | | 4400 |
| 4403 | II | 826 | 1 | 0 | 283 | 0 | 0 | M9+ | 1 | | 21163707 | | | | 4403 |
| 4422 | IV | 827 | 0 | 0 | 283 | 0 | 0 | Mono | Mono | | 21229652 | | | | 4422 |
| 4423 | IV | 827 | 0 | 0 | 283 | 0 | 0 | Mono | Mono | | 21235895 | | | | 4423 |
| 4429 | II | 826 | 1 | 0 | 283 | 0 | 0 | M9+ | 1 | | 21250852 | | | | 4429 |
| 4433 | IV | 827 | 0 | 0 | 283 | 0 | 0 | Mono | Mono | | 21255649 | | | | 4433 |
| 4441 | IV | 827 | 0 | 0 | 283 | 0 | 0 | Mono | Mono | | 21264006 | | | | 4441 |
| 4446 | IV | 803 | 0 | 24 | 273 | 0 | 10 | Mono | Mono | | 21285194 | | | | 4446 |
| 4456 | IV | 827 | 0 | 0 | 283 | 0 | 0 | Mono | Mono | | 21302682 | | | | 4456 |
| 4459 | IV | 827 | 0 | 0 | 283 | 0 | 0 | Mono | Mono | | 21309993 | | | | 4459 |
| 4461 | III | 821 | 1 | 5 | 281 | 0 | 2 | M9+ | 1 | | 21312531 | | | | 4461 |
| 4465 | IV | 827 | 0 | 0 | 283 | 0 | 0 | Mono | Mono | | 21327167 | | | | 4465 |
| 4466 | IV | 826 | 0 | 1 | 283 | 0 | 0 | Mono | Mono | | 21327172 | | | | 4466 |
| 4471 | IV | 827 | 0 | 0 | 283 | 0 | 0 | Mono | Mono | | 21332123 | | | | 4471 |
| 4478 | IV | 827 | 0 | 0 | 283 | 0 | 0 | Mono | Mono | | 21358927 | | | | 4478 |
| 4482 | IV | 827 | 0 | 0 | 283 | 0 | 0 | Mono | Mono | | 21364755 | | | | 4482 |
| 4483 | IV | 826 | 0 | 1 | 283 | 0 | 0 | Mono | Mono | | 21365772 | | | | 4483 |
| 4486 | IV | 827 | 0 | 0 | 283 | 0 | 0 | Mono | Mono | | 21367143 | | | | 4486 |
| 4498 | I | 827 | 0 | 0 | 246 | 37 | 0 | M9- | 1 – 10 | | 21398584 | | | | 4498 |
| 4502 | IV | 827 | 0 | 0 | 283 | 0 | 0 | Mono | Mono | | 21402296 | | | | 4502 |
| 4508 | IV | 827 | 0 | 0 | 283 | 0 | 0 | Mono | Mono | | 21412205 | | | | 4508 |
| 4512 | IV | 827 | 0 | 0 | 283 | 0 | 0 | Mono | Mono | | 21425325 | | | | 4512 |
| 4513 | IV | 827 | 0 | 0 | 283 | 0 | 0 | Mono | Mono | | 21429125 | | | | 4513 |
| 4516 | IV | 814 | 0 | 13 | 275 | 0 | 8 | Mono | Mono | | 21438830 | | | | 4516 |
| 4517 | IV | 827 | 0 | 0 | 283 | 0 | 0 | Mono | Mono | | 21439873 | | | | 4517 |
| 4519 | IV | 827 | 0 | 0 | 283 | 0 | 0 | Mono | Mono | | 21447557 | | | | 4519 |
| 4527 | I | 827 | 0 | 0 | 82 | 201 | 0 | M9- | 22 – 59 | | 21465033 | | | | 4527 |
| 4529 | IV | 827 | 0 | 0 | 283 | 0 | 0 | Mono | Mono | | 21470958 | | | | 4529 |
| 4530 | I | 823 | 4 | 0 | 0 | 283 | 0 | M9+ | 4 – 93 | | 21476910 | | | | 4530 |
| 4541 | III | 822 | 4 | 1 | 1 | 282 | 0 | BOTH | | PI | 21499869 | | | | 4541 |
| 4542 | IV | 827 | 0 | 0 | 283 | 0 | 0 | Mono | Mono | | 21502776 | | | | 4542 |
| 4543 | IV | 826 | 0 | 1 | 283 | 0 | 0 | Mono | Mono | | 21503271 | | | | 4543 |
| 4544 | I | 827 | 0 | 0 | 0 | 283 | 0 | M9+/- | M9+/- | | 21507294 | | | | 4544 |
| 4546 | IV | 827 | 0 | 0 | 283 | 0 | 0 | Mono | Mono | | 21510491 | | | | 4546 |
| 4552 | IV | 827 | 0 | 0 | 282 | 0 | 1 | Mono | Mono | | 21533003 | | | | 4552 |
| 4555 | IV | 826 | 0 | 1 | 283 | 0 | 0 | Mono | Mono | | 21544238 | | | | 4555 |
| 4557 | II | 826 | 1 | 0 | 283 | 0 | 0 | M9+ | 1 | | 21550754 | | | | 4557 |
| 4562 | IV | 827 | 0 | 0 | 282 | 0 | 1 | Mono | Mono | | 21562247 | | | | 4562 |
| 4566 | I | 827 | 0 | 0 | 246 | 37 | 0 | M9- | 1 – 10 | | 21567932 | | | | 4566 |
| 4567 | IV | 827 | 0 | 0 | 283 | 0 | 0 | Mono | Mono | | 21569908 | | | | 4567 |
| 4572 | IV | 826 | 0 | 1 | 283 | 0 | 0 | Mono | Mono | | 21590100 | | | | 4572 |
| 4573 | IV | 827 | 0 | 0 | 283 | 0 | 0 | Mono | Mono | | 21590322 | | | | 4573 |
| 4586 | IV | 827 | 0 | 0 | 283 | 0 | 0 | Mono | Mono | | 21612268 | | | | 4586 |

| List | Stage | M9+ 0 | M9+ 1 | M9+ . | M9- 0 | M9- 1 | M9- . | M9? | Clade(s) | Prob. | Build 37 Position | Name | ISOGG Subgroup Name | Amb. | List |
|---|---|---|---|---|---|---|---|---|---|---|---|---|---|---|---|
| 4589 | II | 826 | 1 | 0 | 283 | 0 | 0 | M9+ | 2 | | 21614288 | | | | 4589 |
| 4603 | IV | 827 | 0 | 0 | 283 | 0 | 0 | Mono | Mono | | 21629209 | | | | 4603 |
| 4606 | IV | 827 | 0 | 0 | 283 | 0 | 0 | Mono | Mono | | 21641094 | | | | 4606 |
| 4609 | IV | 827 | 0 | 0 | 283 | 0 | 0 | Mono | Mono | | 21644268 | | | | 4609 |
| 4621 | IV | 827 | 0 | 0 | 283 | 0 | 0 | Mono | Mono | | 21653414 | | | | 4621 |
| 4637 | IV | 827 | 0 | 0 | 282 | 0 | 1 | Mono | Mono | | 21691104 | | | | 4637 |
| 4642 | IV | 827 | 0 | 0 | 283 | 0 | 0 | Mono | Mono | | 21708134 | | | | 4642 |
| 4652 | IV | 824 | 0 | 3 | 282 | 0 | 1 | Mono | Mono | | 21721419 | | | | 4652 |
| 4657 | IV | 827 | 0 | 0 | 283 | 0 | 0 | Mono | Mono | | 21724699 | | | | 4657 |
| 4659 | IV | 827 | 0 | 0 | 283 | 0 | 0 | Mono | Mono | | 21729056 | | | | 4659 |
| 4660 | IV | 827 | 0 | 0 | 283 | 0 | 0 | Mono | Mono | | 21729491 | | | | 4660 |
| 4668 | I | 796 | 31 | 0 | 283 | 0 | 0 | M9+ | 4 – 16 | | 21733164 | | | | 4668 |
| 4671 | IV | 827 | 0 | 0 | 283 | 0 | 0 | Mono | Mono | | 21735456 | | | | 4671 |
| 4675 | IV | 826 | 0 | 1 | 283 | 0 | 0 | Mono | Mono | | 21738257 | | | | 4675 |
| 4676 | IV | 827 | 0 | 0 | 283 | 0 | 0 | Mono | Mono | | 21739006 | | | | 4676 |
| 4684 | IV | 827 | 0 | 0 | 283 | 0 | 0 | Mono | Mono | | 21740760 | | | | 4684 |
| 4685 | IV | 822 | 0 | 5 | 282 | 0 | 1 | Mono | Mono | | 21741485 | | | | 4685 |
| 4694 | IV | 825 | 0 | 2 | 283 | 0 | 0 | Mono | Mono | | 21751551 | | | | 4694 |
| 4695 | IV | 827 | 0 | 0 | 283 | 0 | 0 | Mono | Mono | | 21752037 | | | | 4695 |
| 4698 | IV | 827 | 0 | 0 | 283 | 0 | 0 | Mono | Mono | | 21754097 | | | | 4698 |
| 4721 | IV | 827 | 0 | 0 | 283 | 0 | 0 | Mono | Mono | | 21766113 | | | | 4721 |
| 4725 | I | 827 | 0 | 0 | 246 | 37 | 0 | M9- | 1 – 10 | | 21768404 | | | | 4725 |
| 4726 | IV | 827 | 0 | 0 | 283 | 0 | 0 | Mono | Mono | | 21769037 | | | | 4726 |
| 4727 | IV | 827 | 0 | 0 | 283 | 0 | 0 | Mono | Mono | | 21772208 | | | | 4727 |
| 4729 | IV | 827 | 0 | 0 | 283 | 0 | 0 | Mono | Mono | | 21777403 | | | | 4729 |
| 4749 | IV | 827 | 0 | 0 | 283 | 0 | 0 | Mono | Mono | | 21811903 | | | | 4749 |
| 4755 | IV | 827 | 0 | 0 | 283 | 0 | 0 | Mono | Mono | | 21817303 | | | | 4755 |
| 4765 | IV | 827 | 0 | 0 | 283 | 0 | 0 | Mono | Mono | | 21853357 | | | | 4765 |
| 4768 | I | 826 | 1 | 0 | 82 | 201 | 0 | BOTH | | PI | 21862684 | | | | 4768 |
| 4769 | IV | 827 | 0 | 0 | 283 | 0 | 0 | Mono | Mono | | 21864291 | | | | 4769 |
| 4770 | III | 789 | 0 | 38 | 254 | 21 | 8 | M9- | 11 – 20 | | 21864828 | | | | 4770 |
| 4775 | IV | 827 | 0 | 0 | 283 | 0 | 0 | Mono | Mono | | 21866659 | | | | 4775 |
| 4781 | IV | 827 | 0 | 0 | 283 | 0 | 0 | Mono | Mono | | 21867979 | | | | 4781 |
| 4782 | IV | 827 | 0 | 0 | 283 | 0 | 0 | Mono | Mono | | 21868035 | | | | 4782 |
| 4785 | IV | 827 | 0 | 0 | 283 | 0 | 0 | Mono | Mono | | 21868519 | | | | 4785 |
| 4791 | IV | 824 | 0 | 3 | 282 | 0 | 1 | Mono | Mono | | 21869266 | | | | 4791 |
| 4792 | IV | 827 | 0 | 0 | 283 | 0 | 0 | Mono | Mono | | 21869383 | | | | 4792 |
| 4795 | IV | 824 | 0 | 3 | 283 | 0 | 0 | Mono | Mono | | 21870325 | | | | 4795 |
| 4797 | IV | 827 | 0 | 0 | 283 | 0 | 0 | Mono | Mono | | 21871502 | | | | 4797 |
| 4798 | IV | 826 | 0 | 1 | 283 | 0 | 0 | Mono | Mono | | 21871549 | | | | 4798 |
| 4799 | IV | 827 | 0 | 0 | 282 | 0 | 1 | Mono | Mono | | 21871614 | | | | 4799 |
| 4803 | IV | 827 | 0 | 0 | 283 | 0 | 0 | Mono | Mono | | 21878072 | | | | 4803 |
| 4804 | IV | 827 | 0 | 0 | 283 | 0 | 0 | Mono | Mono | | 21878191 | | | | 4804 |
| 4805 | IV | 827 | 0 | 0 | 283 | 0 | 0 | Mono | Mono | | 21878322 | | | | 4805 |
| 4806 | IV | 827 | 0 | 0 | 283 | 0 | 0 | Mono | Mono | | 21878677 | | | | 4806 |
| 4815 | IV | 827 | 0 | 0 | 283 | 0 | 0 | Mono | Mono | | 21882724 | | | | 4815 |
| 4816 | IV | 827 | 0 | 0 | 283 | 0 | 0 | Mono | Mono | | 21882889 | | | | 4816 |
| 4817 | IV | 827 | 0 | 0 | 283 | 0 | 0 | Mono | Mono | | 21883162 | | | | 4817 |
| 4818 | III | 820 | 1 | 6 | 283 | 0 | 0 | M9+ | 1 | | 21885026 | | | | 4818 |
| 4819 | IV | 827 | 0 | 0 | 283 | 0 | 0 | Mono | Mono | | 21888865 | | | | 4819 |
| 4820 | IV | 826 | 0 | 1 | 283 | 0 | 0 | Mono | Mono | | 21888873 | | | | 4820 |
| 4834 | IV | 827 | 0 | 0 | 283 | 0 | 0 | Mono | Mono | | 21894072 | | | | 4834 |
| 4835 | II | 827 | 0 | 0 | 282 | 1 | 0 | M9- | 21 | | 21894171 | | | | 4835 |
| 4844 | IV | 827 | 0 | 0 | 283 | 0 | 0 | Mono | Mono | | 21897548 | | | | 4844 |
| 4845 | IV | 827 | 0 | 0 | 283 | 0 | 0 | Mono | Mono | | 21897640 | | | | 4845 |
| 4846 | IV | 827 | 0 | 0 | 283 | 0 | 0 | Mono | Mono | | 21897796 | | | | 4846 |
| 4848 | III | 826 | 0 | 1 | 280 | 3 | 0 | M9- | 9 | | 21898354 | | | | 4848 |
| 4852 | IV | 827 | 0 | 0 | 283 | 0 | 0 | Mono | Mono | | 21901436 | | | | 4852 |
| 4853 | IV | 827 | 0 | 0 | 283 | 0 | 0 | Mono | Mono | | 21901440 | | | | 4853 |
| 4854 | IV | 827 | 0 | 0 | 283 | 0 | 0 | Mono | Mono | | 21901463 | | | | 4854 |
| 4856 | IV | 827 | 0 | 0 | 283 | 0 | 0 | Mono | Mono | | 21902969 | | | | 4856 |
| 4861 | IV | 827 | 0 | 0 | 283 | 0 | 0 | Mono | Mono | | 21905043 | | | | 4861 |
| 4871 | IV | 827 | 0 | 0 | 283 | 0 | 0 | Mono | Mono | | 21909842 | | | | 4871 |
| 4875 | IV | 827 | 0 | 0 | 283 | 0 | 0 | Mono | Mono | | 21916548 | | | | 4875 |
| 4878 | IV | 827 | 0 | 0 | 283 | 0 | 0 | Mono | Mono | | 21917939 | | | | 4878 |
| 4879 | IV | 827 | 0 | 0 | 283 | 0 | 0 | Mono | Mono | | 21919016 | | | | 4879 |
| 4880 | IV | 827 | 0 | 0 | 283 | 0 | 0 | Mono | Mono | | 21921675 | | | | 4880 |
| 4887 | IV | 827 | 0 | 0 | 283 | 0 | 0 | Mono | Mono | | 21930258 | | | | 4887 |
| 4894 | IV | 827 | 0 | 0 | 283 | 0 | 0 | Mono | Mono | | 21935866 | | | | 4894 |
| 4901 | IV | 824 | 0 | 3 | 283 | 0 | 0 | Mono | Mono | | 21949062 | | | | 4901 |
| 4902 | I | 827 | 0 | 0 | 269 | 14 | 0 | M9- | 13 – 20 | | 21950308 | | | | 4902 |
| 4903 | IV | 0 | 827 | 0 | 0 | 283 | 0 | Mono | Mono | | 21952388 | | | | 4903 |
| 4906 | IV | 827 | 0 | 0 | 283 | 0 | 0 | Mono | Mono | | 21969615 | | | | 4906 |
| 4908 | IV | 827 | 0 | 0 | 283 | 0 | 0 | Mono | Mono | | 21973836 | | | | 4908 |
| 4911 | IV | 827 | 0 | 0 | 283 | 0 | 0 | Mono | Mono | | 21978583 | | | | 4911 |
| 4912 | IV | 827 | 0 | 0 | 283 | 0 | 0 | Mono | Mono | | 21981844 | | | | 4912 |
| 4915 | IV | 827 | 0 | 0 | 283 | 0 | 0 | Mono | Mono | | 21984942 | | | | 4915 |
| 4920 | IV | 827 | 0 | 0 | 283 | 0 | 0 | Mono | Mono | | 22002616 | | | | 4920 |
| 4922 | IV | 817 | 0 | 10 | 279 | 0 | 4 | Mono | Mono | | 22004336 | | | | 4922 |
| 4924 | IV | 826 | 0 | 1 | 283 | 0 | 0 | Mono | Mono | | 22011578 | | | | 4924 |
| 4926 | II | 826 | 1 | 0 | 283 | 0 | 0 | M9+ | 2 | | 22019711 | | | | 4926 |
| 4927 | IV | 827 | 0 | 0 | 283 | 0 | 0 | Mono | Mono | | 22019768 | | | | 4927 |
| 4928 | IV | 827 | 0 | 0 | 283 | 0 | 0 | Mono | Mono | | 22019769 | | | | 4928 |
| 4936 | IV | 827 | 0 | 0 | 283 | 0 | 0 | Mono | Mono | | 22047290 | | | | 4936 |
| 4939 | IV | 827 | 0 | 0 | 283 | 0 | 0 | Mono | Mono | | 22063297 | | | | 4939 |
| 4946 | IV | 826 | 0 | 1 | 283 | 0 | 0 | Mono | Mono | | 22086350 | | | | 4946 |
| 4947 | IV | 826 | 0 | 1 | 283 | 0 | 0 | Mono | Mono | | 22091374 | | | | 4947 |
| 4948 | IV | 825 | 0 | 2 | 283 | 0 | 0 | Mono | Mono | | 22092956 | | | | 4948 |
| 4952 | IV | 827 | 0 | 0 | 283 | 0 | 0 | Mono | Mono | | 22108050 | | | | 4952 |
| 4960 | IV | 824 | 0 | 3 | 283 | 0 | 0 | Mono | Mono | | 22127976 | | | | 4960 |
| 4961 | IV | 826 | 0 | 1 | 283 | 0 | 0 | Mono | Mono | | 22127984 | | | | 4961 |
| 4962 | IV | 827 | 0 | 0 | 283 | 0 | 0 | Mono | Mono | | 22127998 | | | | 4962 |
| 4963 | III | 817 | 0 | 10 | 258 | 20 | 5 | M9- | 11 – 20 | MD | 22130140 | | | | 4963 |
| 4971 | II | 826 | 1 | 0 | 283 | 0 | 0 | M9+ | 1 | | 22146007 | | | | 4971 |
| 4973 | IV | 827 | 0 | 0 | 283 | 0 | 0 | Mono | Mono | | 22149593 | | | | 4973 |
| 4980 | IV | 827 | 0 | 0 | 283 | 0 | 0 | Mono | Mono | | 22165294 | | | | 4980 |

| List | Stage | M9+ | | | M9- | | | | | | Build 37 | | ISOGG | | |
|---|---|---|---|---|---|---|---|---|---|---|---|---|---|---|---|
| List | Stage | 0 | 1 | . | 0 | 1 | . | M9? | Clade(s) | Prob. | Position | Name | Subgroup Name | Amb. | List |
| 4985 | III | 824 | 1 | 2 | 282 | 0 | 1 | M9+ | 2 | | 22173931 | | | | 4985 |
| 4990 | IV | 827 | 0 | 0 | 283 | 0 | 0 | Mono | Mono | | 22179053 | | | | 4990 |
| 4997 | IV | 827 | 0 | 0 | 283 | 0 | 0 | Mono | Mono | | 22192044 | | | | 4997 |
| 5000 | III | 824 | 0 | 3 | 282 | 1 | 0 | M9- | 1 | | 22205325 | | | | 5000 |
| 5005 | IV | 0 | 798 | 29 | 0 | 232 | 51 | Mono | Mono | | 22222070 | | | | 5005 |
| 5006 | IV | 0 | 827 | 0 | 0 | 280 | 3 | Mono | Mono | | 22239128 | | | | 5006 |
| 5008 | IV | 0 | 827 | 0 | 0 | 274 | 9 | Mono | Mono | | 22246045 | | | | 5008 |
| 5009 | IV | 0 | 817 | 10 | 0 | 281 | 2 | Mono | Mono | | 22262889 | | | | 5009 |
| 5010 | IV | 0 | 827 | 0 | 0 | 282 | 1 | Mono | Mono | | 22268487 | | | | 5010 |
| 5011 | IV | 821 | 0 | 6 | 281 | 0 | 2 | Mono | Mono | | 22272087 | | | | 5011 |
| 5012 | IV | 0 | 827 | 0 | 0 | 279 | 4 | Mono | Mono | | 22273443 | | | | 5012 |
| 5013 | IV | 827 | 0 | 0 | 278 | 0 | 5 | Mono | Mono | | 22279481 | | | | 5013 |
| 5014 | IV | 0 | 824 | 3 | 0 | 278 | 5 | Mono | Mono | | 22279707 | | | | 5014 |
| 5015 | IV | 0 | 827 | 0 | 0 | 281 | 2 | Mono | Mono | | 22299844 | | | | 5015 |
| 5016 | IV | 0 | 827 | 0 | 0 | 281 | 2 | Mono | Mono | | 22302537 | | | | 5016 |
| 5017 | IV | 827 | 0 | 0 | 281 | 0 | 2 | Mono | Mono | | 22304246 | | | | 5017 |
| 5018 | IV | 0 | 707 | 120 | 0 | 160 | 123 | Mono | Mono | | 22318999 | | | | 5018 |
| 5019 | IV | 0 | 827 | 0 | 0 | 283 | 0 | Mono | Mono | | 22319261 | | | | 5019 |
| 5020 | IV | 0 | 827 | 0 | 0 | 283 | 0 | Mono | Mono | | 22437744 | | | | 5020 |
| 5021 | IV | 0 | 827 | 0 | 0 | 274 | 9 | Mono | Mono | | 22442215 | | | | 5021 |
| 5022 | IV | 816 | 0 | 11 | 265 | 0 | 18 | Mono | Mono | | 22446575 | | | | 5022 |
| 5023 | IV | 827 | 0 | 0 | 283 | 0 | 0 | Mono | Mono | | 22467833 | | | | 5023 |
| 5024 | IV | 0 | 826 | 1 | 0 | 280 | 3 | Mono | Mono | | 22475403 | | | | 5024 |
| 5025 | IV | 0 | 746 | 81 | 0 | 248 | 35 | Mono | Mono | | 22477053 | | | | 5025 |
| 5026 | IV | 826 | 0 | 1 | 274 | 0 | 9 | Mono | Mono | | 22508286 | | | | 5026 |
| 5027 | IV | 717 | 0 | 110 | 188 | 0 | 95 | Mono | Mono | | 22512891 | | | | 5027 |
| 5028 | IV | 821 | 0 | 6 | 280 | 0 | 3 | Mono | Mono | | 22513646 | | | | 5028 |
| 5036 | IV | 826 | 0 | 1 | 282 | 0 | 1 | Mono | Mono | | 22521822 | | | | 5036 |
| 5037 | IV | 827 | 0 | 0 | 283 | 0 | 0 | Mono | Mono | | 22521995 | | | | 5037 |
| 5039 | IV | 827 | 0 | 0 | 283 | 0 | 0 | Mono | Mono | | 22528821 | | | | 5039 |
| 5040 | IV | 827 | 0 | 0 | 283 | 0 | 0 | Mono | Mono | | 22528868 | | | | 5040 |
| 5044 | IV | 827 | 0 | 0 | 283 | 0 | 0 | Mono | Mono | | 22539004 | | | | 5044 |
| 5053 | IV | 825 | 0 | 2 | 283 | 0 | 0 | Mono | Mono | | 22560254 | | | | 5053 |
| 5055 | IV | 827 | 0 | 0 | 283 | 0 | 0 | Mono | Mono | | 22560797 | | | | 5055 |
| 5057 | IV | 827 | 0 | 0 | 283 | 0 | 0 | Mono | Mono | | 22568525 | | | | 5057 |
| 5058 | IV | 827 | 0 | 0 | 283 | 0 | 0 | Mono | Mono | | 22569417 | | | | 5058 |
| 5060 | IV | 826 | 0 | 1 | 281 | 0 | 2 | Mono | Mono | | 22572966 | | | | 5060 |
| 5064 | IV | 827 | 0 | 0 | 283 | 0 | 0 | Mono | Mono | | 22582201 | | | | 5064 |
| 5066 | IV | 827 | 0 | 0 | 283 | 0 | 0 | Mono | Mono | | 22583507 | | | | 5066 |
| 5071 | IV | 826 | 0 | 1 | 283 | 0 | 0 | Mono | Mono | | 22590025 | | | | 5071 |
| 5072 | III | 784 | 1 | 42 | 271 | 0 | 12 | M9+ | 1 | | 22592640 | | | | 5072 |
| 5075 | IV | 827 | 0 | 0 | 283 | 0 | 0 | Mono | Mono | | 22597106 | | | | 5075 |
| 5076 | I | 827 | 0 | 0 | 272 | 11 | 0 | M9- | 37 – 38 | | 22597961 | | | | 5076 |
| 5078 | IV | 826 | 0 | 1 | 282 | 0 | 1 | Mono | Mono | | 22602532 | | | | 5078 |
| 5081 | III | 822 | 1 | 4 | 282 | 0 | 1 | M9+ | 1 | | 22613624 | | | | 5081 |
| 5083 | IV | 826 | 0 | 1 | 283 | 0 | 0 | Mono | Mono | | 22618475 | | | | 5083 |
| 5092 | IV | 826 | 0 | 1 | 283 | 0 | 0 | Mono | Mono | | 22653755 | | | | 5092 |
| 5101 | IV | 827 | 0 | 0 | 283 | 0 | 0 | Mono | Mono | | 22675702 | | | | 5101 |
| 5103 | IV | 827 | 0 | 0 | 283 | 0 | 0 | Mono | Mono | | 22677409 | | | | 5103 |
| 5112 | IV | 827 | 0 | 0 | 283 | 0 | 0 | Mono | Mono | | 22707203 | | | | 5112 |
| 5113 | IV | 826 | 0 | 1 | 283 | 0 | 0 | Mono | Mono | | 22708133 | | | | 5113 |
| 5115 | IV | 827 | 0 | 0 | 283 | 0 | 0 | Mono | Mono | | 22710656 | | | | 5115 |
| 5118 | IV | 827 | 0 | 0 | 283 | 0 | 0 | Mono | Mono | | 22715814 | | | | 5118 |
| 5134 | IV | 827 | 0 | 0 | 283 | 0 | 0 | Mono | Mono | | 22738079 | | | | 5134 |
| 5138 | IV | 823 | 0 | 4 | 281 | 0 | 2 | Mono | Mono | | 22740356 | | | | 5138 |
| 5139 | IV | 827 | 0 | 0 | 283 | 0 | 0 | Mono | Mono | | 22741578 | | | | 5139 |
| 5142 | IV | 827 | 0 | 0 | 283 | 0 | 0 | Mono | Mono | | 22741770 | | | | 5142 |
| 5155 | IV | 827 | 0 | 0 | 283 | 0 | 0 | Mono | Mono | | 22746366 | | | | 5155 |
| 5158 | IV | 827 | 0 | 0 | 283 | 0 | 0 | Mono | Mono | | 22747810 | | | | 5158 |
| 5160 | IV | 827 | 0 | 0 | 283 | 0 | 0 | Mono | Mono | | 22749784 | | | | 5160 |
| 5165 | IV | 827 | 0 | 0 | 283 | 0 | 0 | Mono | Mono | | 22751581 | | | | 5165 |
| 5166 | IV | 826 | 0 | 1 | 283 | 0 | 0 | Mono | Mono | | 22751711 | | | | 5166 |
| 5168 | IV | 827 | 0 | 0 | 283 | 0 | 0 | Mono | Mono | | 22752620 | | | | 5168 |
| 5171 | IV | 827 | 0 | 0 | 283 | 0 | 0 | Mono | Mono | | 22753445 | | | | 5171 |
| 5172 | IV | 827 | 0 | 0 | 283 | 0 | 0 | Mono | Mono | | 22754570 | | | | 5172 |
| 5173 | IV | 827 | 0 | 0 | 283 | 0 | 0 | Mono | Mono | | 22754670 | | | | 5173 |
| 5174 | I | 827 | 0 | 0 | 246 | 37 | 0 | M9- | 1 – 10 | | 22755855 | | | | 5174 |
| 5178 | IV | 827 | 0 | 0 | 283 | 0 | 0 | Mono | Mono | | 22769564 | | | | 5178 |
| 5180 | IV | 827 | 0 | 0 | 283 | 0 | 0 | Mono | Mono | | 22771704 | | | | 5180 |
| 5181 | IV | 827 | 0 | 0 | 283 | 0 | 0 | Mono | Mono | | 22774608 | | | | 5181 |
| 5185 | IV | 823 | 0 | 4 | 281 | 0 | 2 | Mono | Mono | | 22786871 | | | | 5185 |
| 5186 | IV | 827 | 0 | 0 | 282 | 0 | 1 | Mono | Mono | | 22792221 | | | | 5186 |
| 5187 | IV | 827 | 0 | 0 | 282 | 0 | 1 | Mono | Mono | | 22796548 | | | | 5187 |
| 5190 | IV | 827 | 0 | 0 | 283 | 0 | 0 | Mono | Mono | | 22802939 | | | | 5190 |
| 5192 | III | 817 | 1 | 9 | 232 | 50 | 1 | BOTH | | PI | 22815525 | | | | 5192 |
| 5196 | IV | 827 | 0 | 0 | 283 | 0 | 0 | Mono | Mono | | 22822081 | | | | 5196 |
| 5197 | IV | 827 | 0 | 0 | 283 | 0 | 0 | Mono | Mono | | 22822229 | | | | 5197 |
| 5198 | IV | 827 | 0 | 0 | 283 | 0 | 0 | Mono | Mono | | 22822230 | | | | 5198 |
| 5199 | IV | 826 | 0 | 1 | 283 | 0 | 0 | Mono | Mono | | 22822346 | | | | 5199 |
| 5215 | III | 824 | 0 | 3 | 0 | 283 | 0 | M9+/- | M9+/- | | 22866703 | | | | 5215 |
| 5217 | I | 827 | 0 | 0 | 246 | 37 | 0 | M9- | 1 – 10 | | 22872115 | | | | 5217 |
| 5218 | IV | 827 | 0 | 0 | 283 | 0 | 0 | Mono | Mono | | 22876678 | | | | 5218 |
| 5222 | III | 790 | 36 | 1 | 184 | 99 | 0 | BOTH | | PI | 22884954 | | | | 5222 |
| 5233 | II | 826 | 1 | 0 | 283 | 0 | 0 | M9+ | 1 | | 22914760 | | | | 5233 |
| 5237 | III | 795 | 0 | 32 | 80 | 201 | 2 | M9- | 22 – 59 | | 22919332 | | | | 5237 |
| 5238 | IV | 827 | 0 | 0 | 283 | 0 | 0 | Mono | Mono | | 22919840 | | | | 5238 |
| 5244 | IV | 826 | 0 | 1 | 282 | 0 | 1 | Mono | Mono | | 22923226 | | | | 5244 |
| 5245 | IV | 827 | 0 | 0 | 283 | 0 | 0 | Mono | Mono | | 22924348 | | | | 5245 |
| 5246 | IV | 827 | 0 | 0 | 283 | 0 | 0 | Mono | Mono | | 22927023 | | | | 5246 |
| 5247 | IV | 827 | 0 | 0 | 283 | 0 | 0 | Mono | Mono | | 22928067 | | | | 5247 |
| 5249 | IV | 827 | 0 | 0 | 283 | 0 | 0 | Mono | Mono | | 22928699 | | | | 5249 |
| 5256 | IV | 824 | 0 | 3 | 282 | 0 | 1 | Mono | Mono | | 22942886 | | | | 5256 |
| 5258 | IV | 827 | 0 | 0 | 283 | 0 | 0 | Mono | Mono | | 22944601 | | | | 5258 |
| 5260 | IV | 827 | 0 | 0 | 283 | 0 | 0 | Mono | Mono | | 22947915 | | | | 5260 |
| 5263 | IV | 827 | 0 | 0 | 283 | 0 | 0 | Mono | Mono | | 22953408 | | | | 5263 |
| 5266 | IV | 827 | 0 | 0 | 283 | 0 | 0 | Mono | Mono | | 22959424 | | | | 5266 |

| | | | M9+ | | | M9- | | | | | Build 37 | | ISOGG | | |
|---|---|---|---|---|---|---|---|---|---|---|---|---|---|---|---|
| List | Stage | 0 | 1 | . | 0 | 1 | . | M9? | Clade(s) | Prob. | Position | Name | Subgroup Name | Amb. | List |
| 5267 | IV | 827 | 0 | 0 | 283 | 0 | 0 | Mono | Mono | | 22961946 | | | | 5267 |
| 5271 | I | 827 | 0 | 0 | 260 | 23 | 0 | M9- | 60 – 66 | | 22972939 | | | | 5271 |
| 5272 | IV | 827 | 0 | 0 | 283 | 0 | 0 | Mono | Mono | | 22973793 | | | | 5272 |
| 5276 | I | 827 | 0 | 0 | 246 | 37 | 0 | M9- | 1 – 10 | | 22978713 | | | | 5276 |
| 5277 | IV | 827 | 0 | 0 | 283 | 0 | 0 | Mono | Mono | | 22993780 | | | | 5277 |
| 5279 | IV | 823 | 0 | 4 | 283 | 0 | 0 | Mono | Mono | | 22995399 | | | | 5279 |
| 5285 | IV | 826 | 0 | 1 | 283 | 0 | 0 | Mono | Mono | | 23010681 | | | | 5285 |
| 5293 | IV | 827 | 0 | 0 | 283 | 0 | 0 | Mono | Mono | | 23025118 | | | | 5293 |
| 5294 | IV | 827 | 0 | 0 | 283 | 0 | 0 | Mono | Mono | | 23031377 | | | | 5294 |
| 5303 | IV | 827 | 0 | 0 | 281 | 0 | 2 | Mono | Mono | | 23048693 | | | | 5303 |
| 5310 | IV | 827 | 0 | 0 | 283 | 0 | 0 | Mono | Mono | | 23065366 | | | | 5310 |
| 5311 | IV | 827 | 0 | 0 | 283 | 0 | 0 | Mono | Mono | | 23066368 | | | | 5311 |
| 5316 | IV | 827 | 0 | 0 | 283 | 0 | 0 | Mono | Mono | | 23072627 | | | | 5316 |
| 5332 | IV | 827 | 0 | 0 | 283 | 0 | 0 | Mono | Mono | | 23113764 | | | | 5332 |
| 5339 | II | 826 | 1 | 0 | 283 | 0 | 0 | M9+ | 2 | | 23122114 | | | | 5339 |
| 5343 | IV | 827 | 0 | 0 | 283 | 0 | 0 | Mono | Mono | | 23126356 | | | | 5343 |
| 5344 | IV | 824 | 0 | 3 | 283 | 0 | 0 | Mono | Mono | | 23128801 | | | | 5344 |
| 5348 | IV | 827 | 0 | 0 | 283 | 0 | 0 | Mono | Mono | | 23144801 | | | | 5348 |
| 5349 | IV | 827 | 0 | 0 | 283 | 0 | 0 | Mono | Mono | | 23148103 | | | | 5349 |
| 5350 | IV | 827 | 0 | 0 | 283 | 0 | 0 | Mono | Mono | | 23148323 | | | | 5350 |
| 5356 | IV | 827 | 0 | 0 | 281 | 0 | 2 | Mono | Mono | | 23156865 | | | | 5356 |
| 5364 | IV | 827 | 0 | 0 | 283 | 0 | 0 | Mono | Mono | | 23164144 | | | | 5364 |
| 5367 | IV | 827 | 0 | 0 | 283 | 0 | 0 | Mono | Mono | | 23165992 | | | | 5367 |
| 5370 | IV | 827 | 0 | 0 | 282 | 0 | 1 | Mono | Mono | | 23195236 | | | | 5370 |
| 5373 | IV | 827 | 0 | 0 | 283 | 0 | 0 | Mono | Mono | | 23234163 | | | | 5373 |
| 5376 | I | 827 | 0 | 0 | 246 | 37 | 0 | M9- | 1 – 10 | | 23238985 | | | | 5376 |
| 5390 | I | 827 | 0 | 0 | 260 | 23 | 0 | M9- | 60 – 66 | | 23251880 | | | | 5390 |
| 5399 | IV | 827 | 0 | 0 | 283 | 0 | 0 | Mono | Mono | | 23265885 | | | | 5399 |
| 5400 | IV | 827 | 0 | 0 | 283 | 0 | 0 | Mono | Mono | | 23265887 | | | | 5400 |
| 5401 | IV | 827 | 0 | 0 | 283 | 0 | 0 | Mono | Mono | | 23265910 | | | | 5401 |
| 5406 | IV | 827 | 0 | 0 | 283 | 0 | 0 | Mono | Mono | | 23270897 | | | | 5406 |
| 5409 | IV | 827 | 0 | 0 | 283 | 0 | 0 | Mono | Mono | | 23273446 | | | | 5409 |
| 5411 | IV | 827 | 0 | 0 | 283 | 0 | 0 | Mono | Mono | | 23282689 | | | | 5411 |
| 5413 | IV | 827 | 0 | 0 | 283 | 0 | 0 | Mono | Mono | | 23290087 | | | | 5413 |
| 5423 | IV | 827 | 0 | 0 | 283 | 0 | 0 | Mono | Mono | | 23317006 | | | | 5423 |
| 5431 | IV | 827 | 0 | 0 | 283 | 0 | 0 | Mono | Mono | | 23353025 | | | | 5431 |
| 5433 | IV | 0 | 827 | 0 | 0 | 283 | 0 | Mono | Mono | | 23354909 | | | | 5433 |
| 5434 | IV | 827 | 0 | 0 | 283 | 0 | 0 | Mono | Mono | | 23363475 | | | | 5434 |
| 5436 | IV | 827 | 0 | 0 | 283 | 0 | 0 | Mono | Mono | | 23374941 | | | | 5436 |
| 5437 | IV | 827 | 0 | 0 | 283 | 0 | 0 | Mono | Mono | | 23376017 | | | | 5437 |
| 5444 | IV | 827 | 0 | 0 | 283 | 0 | 0 | Mono | Mono | | 23387696 | | | | 5444 |
| 5445 | II | 826 | 1 | 0 | 283 | 0 | 0 | M9+ | 1 | | 23388869 | | | | 5445 |
| 5446 | IV | 827 | 0 | 0 | 283 | 0 | 0 | Mono | Mono | | 23389470 | | | | 5446 |
| 5452 | IV | 827 | 0 | 0 | 283 | 0 | 0 | Mono | Mono | | 23398687 | | | | 5452 |
| 5453 | IV | 827 | 0 | 0 | 283 | 0 | 0 | Mono | Mono | | 23399871 | | | | 5453 |
| 5459 | IV | 827 | 0 | 0 | 283 | 0 | 0 | Mono | Mono | | 23411643 | | | | 5459 |
| 5462 | IV | 827 | 0 | 0 | 283 | 0 | 0 | Mono | Mono | | 23424266 | | | | 5462 |
| 5463 | III | 827 | 0 | 0 | 246 | 36 | 1 | M9- | 1 – 10 | MD | 23430285 | | | | 5463 |
| 5465 | IV | 827 | 0 | 0 | 283 | 0 | 0 | Mono | Mono | | 23435325 | | | | 5465 |
| 5479 | IV | 827 | 0 | 0 | 283 | 0 | 0 | Mono | Mono | | 23476831 | | | | 5479 |
| 5483 | IV | 827 | 0 | 0 | 283 | 0 | 0 | Mono | Mono | | 23482521 | | | | 5483 |
| 5484 | IV | 827 | 0 | 0 | 283 | 0 | 0 | Mono | Mono | | 23482828 | | | | 5484 |
| 5488 | IV | 827 | 0 | 0 | 283 | 0 | 0 | Mono | Mono | | 23494959 | | | | 5488 |
| 5492 | III | 825 | 1 | 1 | 283 | 0 | 0 | M9+ | 1 | | 23496809 | | | | 5492 |
| 5498 | IV | 827 | 0 | 0 | 283 | 0 | 0 | Mono | Mono | | 23502275 | | | | 5498 |
| 5499 | IV | 827 | 0 | 0 | 283 | 0 | 0 | Mono | Mono | | 23531992 | | | | 5499 |
| 5502 | I | 827 | 0 | 0 | 260 | 23 | 0 | M9- | 60 – 66 | | 23538307 | | | | 5502 |
| 5504 | IV | 827 | 0 | 0 | 283 | 0 | 0 | Mono | Mono | | 23544836 | | | | 5504 |
| 5505 | IV | 827 | 0 | 0 | 283 | 0 | 0 | Mono | Mono | | 23545377 | | | | 5505 |
| 5506 | IV | 827 | 0 | 0 | 283 | 0 | 0 | Mono | Mono | | 23545578 | | | | 5506 |
| 5507 | IV | 821 | 0 | 6 | 282 | 0 | 1 | Mono | Mono | | 23547264 | | | | 5507 |
| 5513 | IV | 825 | 0 | 2 | 283 | 0 | 0 | Mono | Mono | | 23555515 | | | | 5513 |
| 5520 | IV | 827 | 0 | 0 | 283 | 0 | 0 | Mono | Mono | | 23571646 | | | | 5520 |
| 5523 | IV | 827 | 0 | 0 | 283 | 0 | 0 | Mono | Mono | | 23580343 | | | | 5523 |
| 5525 | IV | 824 | 0 | 3 | 282 | 0 | 1 | Mono | Mono | | 23584117 | | | | 5525 |
| 5528 | IV | 827 | 0 | 0 | 283 | 0 | 0 | Mono | Mono | | 23608077 | | | | 5528 |
| 5532 | IV | 827 | 0 | 0 | 283 | 0 | 0 | Mono | Mono | | 23618148 | | | | 5532 |
| 5537 | IV | 818 | 0 | 9 | 283 | 0 | 0 | Mono | Mono | | 23627949 | | | | 5537 |
| 5542 | IV | 827 | 0 | 0 | 283 | 0 | 0 | Mono | Mono | | 23652416 | | | | 5542 |
| 5543 | I | 824 | 3 | 0 | 0 | 283 | 0 | M9+ | 3 – 93 | | 23655112 | | | | 5543 |
| 5544 | IV | 818 | 0 | 9 | 281 | 0 | 2 | Mono | Mono | | 23655662 | | | | 5544 |
| 5545 | I | 826 | 1 | 0 | 246 | 37 | 0 | BOTH | | PI | 23729951 | | | | 5545 |
| 5546 | III | 824 | 1 | 2 | 283 | 0 | 0 | M9+ | 1 | | 23732809 | | | | 5546 |
| 5548 | IV | 827 | 0 | 0 | 283 | 0 | 0 | Mono | Mono | | 23742476 | | | | 5548 |
| 5551 | IV | 827 | 0 | 0 | 283 | 0 | 0 | Mono | Mono | | 23749157 | | | | 5551 |
| 5555 | IV | 827 | 0 | 0 | 283 | 0 | 0 | Mono | Mono | | 23758186 | | | | 5555 |
| 5556 | III | 826 | 0 | 1 | 178 | 105 | 0 | M9- | 37 – 59 | | 23759054 | | | | 5556 |
| 5557 | IV | 827 | 0 | 0 | 283 | 0 | 0 | Mono | Mono | | 23759180 | | | | 5557 |
| 5564 | IV | 827 | 0 | 0 | 283 | 0 | 0 | Mono | Mono | | 23781770 | | | | 5564 |
| 5565 | I | 827 | 0 | 0 | 247 | 36 | 0 | M9- | 2 – 10 | | 23785250 | | | | 5565 |
| 5567 | IV | 804 | 0 | 23 | 279 | 0 | 4 | Mono | Mono | | 23791923 | | | | 5567 |
| 5571 | IV | 825 | 0 | 2 | 282 | 0 | 1 | Mono | Mono | | 23800407 | | | | 5571 |
| 5573 | IV | 826 | 0 | 1 | 283 | 0 | 0 | Mono | Mono | | 23801764 | | | | 5573 |
| 5575 | IV | 827 | 0 | 0 | 283 | 0 | 0 | Mono | Mono | | 23807304 | | | | 5575 |
| 5577 | IV | 826 | 0 | 1 | 283 | 0 | 0 | Mono | Mono | | 23810724 | | | | 5577 |
| 5578 | I | 792 | 35 | 0 | 0 | 283 | 0 | M9+ | 17 – 93 | | 23811761 | | | | 5578 |
| 5580 | IV | 826 | 0 | 1 | 283 | 0 | 0 | Mono | Mono | | 23817649 | | | | 5580 |
| 5581 | IV | 826 | 0 | 1 | 283 | 0 | 0 | Mono | Mono | | 23818841 | | | | 5581 |
| 5582 | III | 824 | 0 | 3 | 246 | 37 | 0 | M9- | 1 – 10 | | 23821790 | | | | 5582 |
| 5583 | IV | 827 | 0 | 0 | 283 | 0 | 0 | Mono | Mono | | 23826836 | | | | 5583 |
| 5585 | IV | 827 | 0 | 0 | 283 | 0 | 0 | Mono | Mono | | 23832020 | | | | 5585 |
| 5586 | IV | 806 | 0 | 21 | 277 | 0 | 6 | Mono | Mono | | 23841255 | | | | 5586 |
| 5587 | IV | 826 | 0 | 1 | 283 | 0 | 0 | Mono | Mono | | 23842147 | | | | 5587 |
| 5589 | IV | 826 | 0 | 1 | 283 | 0 | 0 | Mono | Mono | | 23851447 | | | | 5589 |
| 5590 | IV | 827 | 0 | 0 | 283 | 0 | 0 | Mono | Mono | | 23852284 | | | | 5590 |
| 5591 | IV | 827 | 0 | 0 | 283 | 0 | 0 | Mono | Mono | | 23856058 | | | | 5591 |

| | | | M9+ | | | M9- | | | | | Build 37 | | ISOGG | | |
|---|---|---|---|---|---|---|---|---|---|---|---|---|---|---|---|
| List | Stage | 0 | 1 | . | 0 | 1 | . | M9? | Clade(s) | Prob. | Position | Name | Subgroup Name | Amb. | List |
| 5592 | IV | 827 | 0 | 0 | 283 | 0 | 0 | Mono | Mono | | 23863301 | | | | 5592 |
| 5593 | I | 827 | 0 | 0 | 246 | 37 | 0 | M9- | 1 – 10 | | 23870139 | | | | 5593 |
| 5595 | IV | 827 | 0 | 0 | 283 | 0 | 0 | Mono | Mono | | 23878211 | | | | 5595 |
| 5596 | III | 823 | 2 | 2 | 280 | 2 | 1 | BOTH | | PI | 23881461 | | | | 5596 |
| 5597 | IV | 827 | 0 | 0 | 283 | 0 | 0 | Mono | Mono | | 23883619 | | | | 5597 |
| 5598 | IV | 826 | 0 | 1 | 283 | 0 | 0 | Mono | Mono | | 23897175 | | | | 5598 |
| 5599 | IV | 827 | 0 | 0 | 283 | 0 | 0 | Mono | Mono | | 23897221 | | | | 5599 |
| 5600 | IV | 821 | 0 | 6 | 283 | 0 | 0 | Mono | Mono | | 23897924 | | | | 5600 |
| 5601 | IV | 826 | 0 | 1 | 282 | 0 | 1 | Mono | Mono | | 23898257 | | | | 5601 |
| 5602 | IV | 827 | 0 | 0 | 283 | 0 | 0 | Mono | Mono | | 23898319 | | | | 5602 |
| 5603 | IV | 827 | 0 | 0 | 283 | 0 | 0 | Mono | Mono | | 23898535 | | | | 5603 |
| 5604 | IV | 827 | 0 | 0 | 283 | 0 | 0 | Mono | Mono | | 23898898 | | | | 5604 |
| 5605 | IV | 0 | 819 | 8 | 0 | 283 | 0 | Mono | Mono | | 23899170 | | | | 5605 |
| 5606 | IV | 827 | 0 | 0 | 283 | 0 | 0 | Mono | Mono | | 23899533 | | | | 5606 |
| 5607 | IV | 827 | 0 | 0 | 283 | 0 | 0 | Mono | Mono | | 23899893 | | | | 5607 |
| 5608 | IV | 827 | 0 | 0 | 283 | 0 | 0 | Mono | Mono | | 23899906 | | | | 5608 |
| 5609 | IV | 826 | 0 | 1 | 282 | 0 | 1 | Mono | Mono | | 23900003 | | | | 5609 |
| 5610 | IV | 827 | 0 | 0 | 283 | 0 | 0 | Mono | Mono | | 23900125 | | | | 5610 |
| 5614 | IV | 826 | 0 | 1 | 283 | 0 | 0 | Mono | Mono | | 23959349 | | | | 5614 |
| 5616 | IV | 827 | 0 | 0 | 283 | 0 | 0 | Mono | Mono | | 23963405 | | | | 5616 |
| 5620 | IV | 826 | 0 | 1 | 281 | 0 | 2 | Mono | Mono | | 23972908 | | | | 5620 |
| 5623 | IV | 827 | 0 | 0 | 283 | 0 | 0 | Mono | Mono | | 23974513 | | | | 5623 |
| 5628 | II | 826 | 1 | 0 | 283 | 0 | 0 | M9+ | 1 | | 23977583 | | | | 5628 |
| 5631 | IV | 817 | 0 | 10 | 281 | 0 | 2 | Mono | Mono | | 23988956 | | | | 5631 |
| 5632 | IV | 827 | 0 | 0 | 283 | 0 | 0 | Mono | Mono | | 23989547 | | | | 5632 |
| 5638 | IV | 825 | 0 | 2 | 281 | 0 | 2 | Mono | Mono | | 24007524 | | | | 5638 |
| 5639 | IV | 827 | 0 | 0 | 283 | 0 | 0 | Mono | Mono | | 24007623 | | | | 5639 |
| 5640 | IV | 822 | 0 | 5 | 281 | 0 | 2 | Mono | Mono | | 24007925 | | | | 5640 |
| 5646 | IV | 825 | 0 | 2 | 283 | 0 | 0 | Mono | Mono | | 24366204 | | | | 5646 |
| 5648 | II | 825 | 2 | 0 | 283 | 0 | 0 | M9+ | 5 | | 24366649 | | | | 5648 |
| 5649 | IV | 825 | 0 | 2 | 283 | 0 | 0 | Mono | Mono | | 24367067 | | | | 5649 |
| 5657 | IV | 827 | 0 | 0 | 283 | 0 | 0 | Mono | Mono | | 24392881 | | | | 5657 |
| 5665 | IV | 825 | 0 | 2 | 283 | 0 | 0 | Mono | Mono | | 24410769 | | | | 5665 |
| 5668 | IV | 820 | 0 | 7 | 281 | 0 | 2 | Mono | Mono | | 24415101 | | | | 5668 |
| 5669 | IV | 825 | 0 | 2 | 270 | 0 | 13 | Mono | Mono | | 24415720 | | | | 5669 |
| 5676 | IV | 0 | 825 | 2 | 0 | 283 | 0 | Mono | Mono | | 24447989 | | | | 5676 |
| 5680 | IV | 827 | 0 | 0 | 283 | 0 | 0 | Mono | Mono | | 24457245 | | | | 5680 |
| 5689 | IV | 827 | 0 | 0 | 283 | 0 | 0 | Mono | Mono | | 24497922 | | | | 5689 |
| 5695 | IV | 824 | 0 | 3 | 282 | 0 | 1 | Mono | Mono | | 24511620 | | | | 5695 |
| 5697 | IV | 825 | 0 | 2 | 282 | 0 | 1 | Mono | Mono | | 24522209 | | | | 5697 |
| 5698 | IV | 827 | 0 | 0 | 283 | 0 | 0 | Mono | Mono | | 24522333 | | | | 5698 |
| 5699 | IV | 824 | 0 | 3 | 283 | 0 | 0 | Mono | Mono | | 27035220 | | | | 5699 |
| 5700 | III | 826 | 0 | 1 | 262 | 19 | 2 | M9- | | PI | 27390536 | | | | 5700 |
| 5701 | III | 827 | 0 | 0 | 246 | 14 | 23 | M9- | 1 – 10 | MD | 28459738 | | | | 5701 |
| 5702 | III | 827 | 0 | 0 | 254 | 27 | 2 | M9- | 37 – 41 | | 28475000 | | | | 5702 |
| 5703 | IV | 827 | 0 | 0 | 283 | 0 | 0 | Mono | Mono | | 28504055 | | | | 5703 |
| 5704 | IV | 826 | 0 | 1 | 282 | 0 | 1 | Mono | Mono | | 28509482 | | | | 5704 |
| 5705 | IV | 826 | 0 | 1 | 283 | 0 | 0 | Mono | Mono | | 28547314 | | | | 5705 |
| 5706 | I | 827 | 0 | 0 | 270 | 13 | 0 | M9- | 14 – 20 | | 28556572 | | | | 5706 |
| 5707 | IV | 827 | 0 | 0 | 283 | 0 | 0 | Mono | Mono | | 28568089 | | | | 5707 |
| 5708 | IV | 814 | 0 | 13 | 276 | 0 | 7 | Mono | Mono | | 28583394 | | | | 5708 |
| 5710 | III | 825 | 0 | 2 | 246 | 36 | 1 | M9- | 2 – 10 | | 28612905 | | | | 5710 |
| 5711 | IV | 827 | 0 | 0 | 283 | 0 | 0 | Mono | Mono | | 28617025 | | | | 5711 |
| 5712 | I | 827 | 0 | 0 | 262 | 21 | 0 | M9- | 11 – 20 | | 28632813 | | | | 5712 |
| 5714 | IV | 827 | 0 | 0 | 283 | 0 | 0 | Mono | Mono | | 28659645 | | | | 5714 |
| 5715 | IV | 825 | 0 | 2 | 283 | 0 | 0 | Mono | Mono | | 28688867 | | | | 5715 |
| 5716 | IV | 827 | 0 | 0 | 283 | 0 | 0 | Mono | Mono | | 28688884 | | | | 5716 |
| 5717 | IV | 827 | 0 | 0 | 283 | 0 | 0 | Mono | Mono | | 28693558 | | | | 5717 |
| 5718 | I | 827 | 0 | 0 | 224 | 59 | 0 | M9- | 1 – 21 | | 28699018 | | | | 5718 |
| 5719 | IV | 781 | 0 | 46 | 264 | 0 | 19 | Mono | Mono | | 28701583 | | | | 5719 |
| 5720 | IV | 816 | 0 | 11 | 283 | 0 | 0 | Mono | Mono | | 28710492 | | | | 5720 |
| 5721 | IV | 827 | 0 | 0 | 283 | 0 | 0 | Mono | Mono | | 28727063 | | | | 5721 |
| 5722 | IV | 827 | 0 | 0 | 283 | 0 | 0 | Mono | Mono | | 28733089 | | | | 5722 |
| 5725 | IV | 827 | 0 | 0 | 283 | 0 | 0 | Mono | Mono | | 28755784 | | | | 5725 |
| 5726 | IV | 827 | 0 | 0 | 283 | 0 | 0 | Mono | Mono | | 28760588 | | | | 5726 |
| 5729 | IV | 827 | 0 | 0 | 283 | 0 | 0 | Mono | Mono | | 28777064 | | | | 5729 |
| 5730 | IV | 827 | 0 | 0 | 283 | 0 | 0 | Mono | Mono | | 28784903 | | | | 5730 |
| 5731 | IV | 827 | 0 | 0 | 283 | 0 | 0 | Mono | Mono | | 28788467 | | | | 5731 |
| 5732 | IV | 825 | 0 | 2 | 281 | 0 | 2 | Mono | Mono | | 28788472 | | | | 5732 |
| 5733 | IV | 827 | 0 | 0 | 283 | 0 | 0 | Mono | Mono | | 28789331 | | | | 5733 |
| 5735 | IV | 824 | 0 | 3 | 283 | 0 | 0 | Mono | Mono | | 28791555 | | | | 5735 |
| 5736 | IV | 0 | 824 | 3 | 0 | 280 | 3 | Mono | Mono | | 28791769 | | | | 5736 |
| 5737 | IV | 827 | 0 | 0 | 283 | 0 | 0 | Mono | Mono | | 28797325 | | | | 5737 |
| 5738 | IV | 827 | 0 | 0 | 283 | 0 | 0 | Mono | Mono | | 28802564 | | | | 5738 |
| 5739 | IV | 825 | 0 | 2 | 283 | 0 | 0 | Mono | Mono | | 28804190 | | | | 5739 |
| 5740 | IV | 827 | 0 | 0 | 283 | 0 | 0 | Mono | Mono | | 28804197 | | | | 5740 |
| 5741 | IV | 0 | 826 | 1 | 0 | 282 | 1 | Mono | Mono | | 28804383 | | | | 5741 |
| 5742 | IV | 825 | 0 | 2 | 283 | 0 | 0 | Mono | Mono | | 28804848 | | | | 5742 |
| 5743 | IV | 0 | 826 | 1 | 0 | 283 | 0 | Mono | Mono | | 28805547 | | | | 5743 |
| 5744 | IV | 0 | 827 | 0 | 0 | 283 | 0 | Mono | Mono | | 28807236 | | | | 5744 |
| 5745 | IV | 0 | 827 | 0 | 0 | 283 | 0 | Mono | Mono | | 28808002 | | | | 5745 |
| 5746 | IV | 827 | 0 | 0 | 283 | 0 | 0 | Mono | Mono | | 28808049 | | | | 5746 |
| 5747 | IV | 824 | 0 | 3 | 280 | 0 | 3 | Mono | Mono | | 28809587 | | | | 5747 |
| 5748 | IV | 0 | 827 | 0 | 0 | 283 | 0 | Mono | Mono | | 28810859 | | | | 5748 |
| 5749 | IV | 827 | 0 | 0 | 283 | 0 | 0 | Mono | Mono | | 28813634 | | | | 5749 |
| 5750 | IV | 827 | 0 | 0 | 283 | 0 | 0 | Mono | Mono | | 28815253 | | | | 5750 |
| 5751 | IV | 827 | 0 | 0 | 283 | 0 | 0 | Mono | Mono | | 28816607 | | | | 5751 |
| 5752 | IV | 827 | 0 | 0 | 283 | 0 | 0 | Mono | Mono | | 28816703 | | | | 5752 |
| 5753 | IV | 827 | 0 | 0 | 283 | 0 | 0 | Mono | Mono | | 28816755 | | | | 5753 |
| 5754 | IV | 827 | 0 | 0 | 283 | 0 | 0 | Mono | Mono | | 28817114 | | | | 5754 |
| 5755 | IV | 827 | 0 | 0 | 283 | 0 | 0 | Mono | Mono | | 28817191 | | | | 5755 |
| 5756 | IV | 827 | 0 | 0 | 283 | 0 | 0 | Mono | Mono | | 28817442 | | | | 5756 |
| 5757 | IV | 827 | 0 | 0 | 283 | 0 | 0 | Mono | Mono | | 28817458 | | | | 5757 |
| 5758 | IV | 827 | 0 | 0 | 283 | 0 | 0 | Mono | Mono | | 28817494 | | | | 5758 |
| 5759 | IV | 827 | 0 | 0 | 283 | 0 | 0 | Mono | Mono | | 28817579 | | | | 5759 |
| 5760 | IV | 0 | 827 | 0 | 0 | 281 | 2 | Mono | Mono | | 28817636 | | | | 5760 |

| | | M9+ | | | | M9- | | | | | | Build 37 | | ISOGG | | |
|---|---|---|---|---|---|---|---|---|---|---|---|---|---|---|---|---|
| List | Stage | 0 | 1 | . | 0 | 1 | . | M9? | Clade(s) | Prob. | | Position | Name | Subgroup Name | Amb. | List |
| 5761 | IV | 0 | 826 | 1 | 0 | 282 | 1 | Mono | Mono | | | 58843097 | | | | 5761 |

**Columns:**

List -- Numerical, ordered by Build 37 Position.

Stage -- I through IV

M9+/M9- (0, 1, .) -- these seven columns are the summary genotype data (counts) for each SNP

M9? -- M9- are clades splitting off prior to M9; M9+ are those after M9; M9-/M9+ are the M9 split itself; Mono are monomorphic SNPs; phylogenetically inconsistent (PI) SNPs with M9- and M9+ variability are labeled BOTH.

Clades -- Simple numerical designation of terminal branches or clades, essentially following alphabetical order given by inferred ISOGG Subgroup name. 1 -- 66 for M9- Clades, 1 -- 94 for M9+ Clades. NU: not used, for SNPs in which a single genotypi

Prob. -- Problematic SNPs. PI -- phylogenetically inconsistent (56 SNPs); S -- singleton SNPs without additional support, inferred to be genotyping errors (xx); 1H = one heterozygous genotype (13 SNPs), or 2H = heterozygous genotypes (1 SNP), conv

Build 37 Position -- basepair position of SNP according to HG Build 37

Name -- SNP's identifier

ISOGG Subgroup Name -- Subgroup Name assigned each SNP, when available

Amb. -- ISOGG label ambiguity: A = multiple distinct ISOGG labels assigned; W = wrong for our data (see Supplementary Table ?3?)

List -- Numerical, ordered by Build 37 Position. Should be identical to first column.
