## Supplementary material for "Characterization of the Y Chromosome in Newfoundland and Labrador: Evidence of a Founder Effect": Table S2

| M9 | Clade | ISOGG | Count (n = 1110) | Count (n = 831) |
|---|---|---|---|---|
| M9- | 1 | E1a2a1 | 1 | 0 |
| M9- | 2 | E1b1b1a1a1 | 2 | 1 |
| M9- | 3 | E1b1b1a1b1 | 1 | 1 |
| M9- | 4 | E1b1b1a1b1a | 20 | 14 |
| M9- | 5 | E1b1b1a1b1a^ | 2 | 0 |
| M9- | 6 | E1b1b1a1b2 | 3 | 1 |
| M9- | 7 | E1b1b1a1b2^ | 2 | 2 |
| M9- | 8 | E1b1b1b1b | 2 | 1 |
| M9- | 9 | E1b1b1b2a1a1~ | 3 | 1 |
| M9- | 10 | E1b1b1b2a1a4 | 1 | 1 |
| M9- | 11 | G2a1a1a1a1 | 1 | 0 |
| M9- | 12 | G2a2a1a2a1a | 6 | 4 |
| M9- | 13 | G2a2b1 | 1 | 1 |
| M9- | 14 | G2a2b2a1 | 1 | 1 |
| M9- | 15 | G2a2b2a1a1 | 1 | 0 |
| M9- | 16 | G2a2b2a1a1a1a1 | 1 | 1 |
| M9- | 17 | G2a2b2a1a1b1a | 3 | 2 |
| M9- | 18 | G2a2b2a1a1b1a1a2a1 | 4 | 2 |
| M9- | 19 | G2a2b2a1a1c2 | 1 | 0 |
| M9- | 20 | G2a2b2b1a1 | 2 | 2 |
| M9- | 21 | H1a1a4b2 | 1 | 0 |
| M9- | 22 | I1a~ | 12 | 10 |
| M9- | 23 | I1a1 | 1 | 0 |
| M9- | 24 | I1a1b1 | 10 | 7 |
| M9- | 25 | I1a1b1a1 | 2 | 2 |
| M9- | 26 | I1a1b1a4a | 2 | 1 |
| M9- | 27 | I1a2a1a1~ | 7 | 5 |
| M9- | 28 | I1a2a1a1a | 3 | 3 |
| M9- | 29 | I1a2a1a1a1 | 7 | 5 |
| M9- | 30 | I1a2a1a1a2 | 5 | 3 |
| M9- | 31 | I1a2b | 13 | 7 |
| M9- | 32 | I1a2b^ | 5 | 4 |
| M9- | 33 | I1a3~ | 18 | 13 |
| M9- | 34 | I1a3~^ | 2 | 2 |
| M9- | 35 | I1a3~^^ | 2 | 1 |
| M9- | 36 | I1a3a | 7 | 6 |
| M9- | 37 | I2a1a1a1a~ | 7 | 7 |
| M9- | 38 | I2a1a1a1a1a | 4 | 1 |
| M9- | 39 | I2a1a1b1a~ | 7 | 5 |
| M9- | 40 | I2a1a2a | 6 | 5 |
| M9- | 41 | I2a1a2b1a1a1 | 3 | 0 |
| M9- | 42 | I2a1b1 | 1 | 0 |
| M9- | 43 | I2a1b1a | 1 | 1 |
| M9- | 44 | I2a1b1a~ | 4 | 3 |

| M9 | Clade | ISOGG | Count (n = 1110) | Count (n = 831) |
|---|---|---|---|---|
| M9- | 45 | I2a1b1a1a1a | 6 | 5 |
| M9- | 46 | I2a1b1a1b1a1a | 1 | 1 |
| M9- | 47 | I2a1b1a2 | 3 | 3 |
| M9- | 48 | I2a1b1a2^ | 2 | 1 |
| M9- | 49 | I2a1b1a2b1 | 12 | 9 |
| M9- | 50 | I2a1b1a2b1^ | 2 | 1 |
| M9- | 51 | I2a1b1a2b1a2 | 8 | 5 |
| M9- | 52 | I2a1b1a2b1a2^ | 3 | 3 |
| M9- | 53 | I2a1b1a2b1a2a1a1a1a | 3 | 2 |
| M9- | 54 | I2a1b1a2b1a2a1a1a1a1a | 2 | 2 |
| M9- | 55 | I2a1b1a2b1a2a2 | 3 | 2 |
| M9- | 56 | I2a1b1a2b1a2e2a3~ | 13 | 10 |
| M9- | 57 | I2a1b2a | 9 | 9 |
| M9- | 58 | I2a1b2a^ | 1 | 0 |
| M9- | 59 | I2a2 | 4 | 4 |
| M9- | 60 | J1a | 1 | 0 |
| M9- | 61 | J2a1a | 4 | 3 |
| M9- | 62 | J2a1a1a2b2 | 2 | 0 |
| M9- | 63 | J2a1a1a2b2a1a | 2 | 2 |
| M9- | 64 | J2a1a4b | 1 | 0 |
| M9- | 65 | J2b2a | 3 | 2 |
| M9- | 66 | J2b2a1a1a1a1a | 10 | 6 |
| M9+ | 1 | O1a1a1 | 1 | 0 |
| M9+ | 2 | O1b1a1a1a1a1 | 1 | 0 |
| M9+ | 3 | Q2a1a1a1 | 1 | 0 |
| M9+ | 4 | R1a1a1~ | 9 | 7 |
| M9+ | 5 | R1a1a1~^ | 2 | 1 |
| M9+ | 6 | R1a1a1b1a1a | 1 | 0 |
| M9+ | 7 | R1a1a1b1a2b3a | 2 | 2 |
| M9+ | 8 | R1a1a1b1a3a^ | 1 | 0 |
| M9+ | 9 | R1a1a1b1a3a1 | 3 | 2 |
| M9+ | 10 | R1a1a1b1a3a1a | 4 | 3 |
| M9+ | 11 | R1a1a1b1a3a1a^ | 2 | 1 |
| M9+ | 12 | R1a1a1b1a3a2 | 1 | 1 |
| M9+ | 13 | R1a1a1b1a3a2a1 | 1 | 0 |
| M9+ | 14 | R1a1a1b1a3a3a | 2 | 2 |
| M9+ | 15 | R1a1a1b2^ | 2 | 1 |
| M9+ | 16 | R1a1a1b2a~ | 1 | 1 |
| M9+ | 17 | R1b1a1b1 | 9 | 8 |
| M9+ | 18 | R1b1a1b1a1a | 65 | 51 |
| M9+ | 19 | R1b1a1b1a1a^ | 3 | 3 |
| M9+ | 20 | R1b1a1b1a1a^^ | 2 | 2 |
| M9+ | 21 | R1b1a1b1a1a^^^ | 4 | 4 |
| M9+ | 22 | R1b1a1b1a1a^^^^ | 6 | 5 |
| M9+ | 23 | R1b1a1b1a1a1 | 6 | 5 |

| M9 | Clade | ISOGG | Count (n = 1110) | Count (n = 831) |
|---|---|---|---|---|
| M9+ | 24 | R1b1a1b1a1a1^ | 6 | 5 |
| M9+ | 25 | R1b1a1b1a1a1a | 6 | 2 |
| M9+ | 26 | R1b1a1b1a1a1b | 9 | 7 |
| M9+ | 27 | R1b1a1b1a1a1b1a | 7 | 5 |
| M9+ | 28 | R1b1a1b1a1a1c1 | 30 | 21 |
| M9+ | 29 | R1b1a1b1a1a1c1a | 26 | 24 |
| M9+ | 30 | R1b1a1b1a1a1c2a1 | 18 | 17 |
| M9+ | 31 | R1b1a1b1a1a1c2b | 6 | 6 |
| M9+ | 32 | R1b1a1b1a1a1c2b1a1a1 | 1 | 0 |
| M9+ | 33 | R1b1a1b1a1a1c2b1b | 12 | 12 |
| M9+ | 34 | R1b1a1b1a1a1c2b1b2a1 | 1 | 1 |
| M9+ | 35 | R1b1a1b1a1a1c2b2a | 4 | 2 |
| M9+ | 36 | R1b1a1b1a1a1c2b2a^ | 1 | 1 |
| M9+ | 37 | R1b1a1b1a1a1c2b2a1b | 13 | 13 |
| M9+ | 38 | R1b1a1b1a1a1c2b2a1b1 | 2 | 0 |
| M9+ | 39 | R1b1a1b1a1a1c2b2a1b1a | 35 | 26 |
| M9+ | 40 | R1b1a1b1a1a1c2b2a1b1a^ | 2 | 2 |
| M9+ | 41 | R1b1a1b1a1a1c2b2a1b1a1 | 4 | 2 |
| M9+ | 42 | R1b1a1b1a1a1c2b2a1b1a1a | 11 | 9 |
| M9+ | 43 | R1b1a1b1a1a1c2b2a1b1b1 | 6 | 5 |
| M9+ | 44 | R1b1a1b1a1a1c2b2b | 4 | 0 |
| M9+ | 45 | R1b1a1b1a1a1c2b2b1a~ | 12 | 8 |
| M9+ | 46 | R1b1a1b1a1a1c3 | 1 | 1 |
| M9+ | 47 | R1b1a1b1a1a2a1 | 13 | 9 |
| M9+ | 48 | R1b1a1b1a1a2a1^ | 5 | 3 |
| M9+ | 49 | R1b1a1b1a1a2a1^^ | 1 | 0 |
| M9+ | 50 | R1b1a1b1a1a2a1a1~ | 10 | 9 |
| M9+ | 51 | R1b1a1b1a1a2a1a1~^ | 2 | 2 |
| M9+ | 52 | R1b1a1b1a1a2a1a1~^^ | 2 | 2 |
| M9+ | 53 | R1b1a1b1a1a2a1a1a1~ | 1 | 1 |
| M9+ | 54 | R1b1a1b1a1a2a1a1a1~^ | 3 | 3 |
| M9+ | 55 | R1b1a1b1a1a2a1a1a1~^^ | 2 | 2 |
| M9+ | 56 | R1b1a1b1a1a2a1a1a1a | 1 | 1 |
| M9+ | 57 | R1b1a1b1a1a2a1a1a2 | 1 | 0 |
| M9+ | 58 | R1b1a1b1a1a2a1b1a1~ | 27 | 20 |
| M9+ | 59 | R1b1a1b1a1a2a5~ | 3 | 1 |
| M9+ | 60 | R1b1a1b1a1a2b | 10 | 7 |
| M9+ | 61 | R1b1a1b1a1a2b1 | 20 | 14 |
| M9+ | 62 | R1b1a1b1a1a2b1a~ | 7 | 4 |
| M9+ | 63 | R1b1a1b1a1a2b1a1~ | 10 | 7 |
| M9+ | 64 | R1b1a1b1a1a2c1 | 4 | 3 |
| M9+ | 65 | R1b1a1b1a1a2c1^ | 1 | 0 |
| M9+ | 66 | R1b1a1b1a1a2c1a | 112 | 86 |
| M9+ | 67 | R1b1a1b1a1a2c1a^ | 3 | 1 |
| M9+ | 68 | R1b1a1b1a1a2c1a^^ | 2 | 2 |

| M9 | Clade | ISOGG | Count (n = 1110) | Count (n = 831) |
|---|---|---|---|---|
| M9+ | 69 | R1b1a1b1a1a2c1a^^^ | 2 | 2 |
| M9+ | 70 | R1b1a1b1a1a2c1a1a | 12 | 12 |
| M9+ | 71 | R1b1a1b1a1a2c1a1a^ | 3 | 2 |
| M9+ | 72 | R1b1a1b1a1a2c1a1a^^ | 1 | 0 |
| M9+ | 73 | R1b1a1b1a1a2c1a1a1a | 4 | 3 |
| M9+ | 74 | R1b1a1b1a1a2c1a1a1a1 | 34 | 22 |
| M9+ | 75 | R1b1a1b1a1a2c1a1b | 4 | 3 |
| M9+ | 76 | R1b1a1b1a1a2c1a1d | 7 | 7 |
| M9+ | 77 | R1b1a1b1a1a2c1a1f1 | 7 | 5 |
| M9+ | 78 | R1b1a1b1a1a2c1a3a2 | 33 | 27 |
| M9+ | 79 | R1b1a1b1a1a2c1a4a | 32 | 21 |
| M9+ | 80 | R1b1a1b1a1a2c1a4b | 17 | 12 |
| M9+ | 81 | R1b1a1b1a1a2c1a4b^ | 2 | 0 |
| M9+ | 82 | R1b1a1b1a1a2c1a4b2 | 25 | 23 |
| M9+ | 83 | R1b1a1b1a1a2c1a4b2c | 2 | 2 |
| M9+ | 84 | R1b1a1b1a1a2c1a4b2c1a | 24 | 20 |
| M9+ | 85 | R1b1a1b1a1a2c1a5a2a | 3 | 0 |
| M9+ | 86 | R1b1a1b1a1a2c1a5b1a1 | 4 | 4 |
| M9+ | 87 | R1b1a1b1a1a2c1a5b1a1a | 7 | 4 |
| M9+ | 88 | R1b1a1b1a1a2c1a5b1a1a2 | 5 | 3 |
| M9+ | 89 | R1b1a1b1a1a2c1a5d3 | 1 | 1 |
| M9+ | 90 | R1b1a1b1a1a2e | 3 | 3 |
| M9+ | 91 | R1b1a1b1a1a2e1 | 6 | 5 |
| M9+ | 92 | R1b1a1b1a1a2e1^ | 1 | 0 |
| M9+ | 93 | R1b1a1b2a | 1 | 1 |
| M9+ | 94 | T1a1a1b2b2b1a1a | 1 | 1 |
