## Supplementary material for "Characterization of the Y Chromosome in Newfoundland and Labrador: Evidence of a Founder Effect": Table S3

| | | M9+ | | | M9- | | | | | | Build 37 | | ISOGG | | |
|---|---|---|---|---|---|---|---|---|---|---|---|---|---|---|---|
| List | Stage | 0 | 1 | . | 0 | 1 | . | M9? | Clade(s) | Res. | Position | Name | Subgroup Name | Amb. | List |
| 415 | I | 824 | 3 | 0 | 274 | 9 | 0 | BOTH | | PI | 5836035 | | | | 415 |
| 415.1 | V | - | - | - | - | 9 | - | M9- | 61 – 64 | H | | | | | 415.1 |
| 415.2 | V | - | 2 | - | - | - | - | M9+ | 81 | H | | | | | 415.2 |
| 415.3 | V | - | 1 | - | - | - | - | M9+ | NU | H,S | | | | | 415.3 |
| 1240 | I | 822 | 5 | 0 | 279 | 4 | 0 | BOTH | | PI | 8261193 | | | | 1240 |
| 1240.1 | V | - | - | - | - | 2 | - | M9- | 48 | H | | | | | 1240.1 |
| 1240.2 | V | - | - | - | - | 1 | - | M9- | 42 | H,S | | | | | 1240.2 |
| 1240.3 | V | - | - | - | - | 1 | - | M9- | NU | H,S | | | | | 1240.3 |
| 1240.4 | V | - | 4 | - | - | - | - | M9+ | 21 | H | | | | | 1240.4 |
| 1240.5 | V | - | 1 | - | - | - | - | M9+ | 56 | H,S | | | | | 1240.5 |
| 1548 | I | 824 | 3 | 0 | 262 | 21 | 0 | BOTH | | PI | 8700380 | PF2892 | G | | 1548 |
| 1548.1 | V | - | - | - | - | 21 | - | M9- | 11 – 20 | H | | | G | | 1548.1 |
| 1548.2 | V | - | 3 | - | - | - | - | M9+ | 71 | H | | | G | W | 1548.2 |
| 1704 | I | 827 | 0 | 0 | 83 | 200 | 0 | M9- | | PI | 9516653 | | | | 1704 |
| 1704.1 | V | 827 | 0 | 0 | 83 | 200 | 0 | M9- | 22 – 59 | 1E | | | | | 1704.1 |
| 1710 | I | 790 | 37 | 0 | 0 | 283 | 0 | M9+ | | PI | 9646980 | | | | 1710 |
| 1710.1 | V | 790 | 37 | 0 | 0 | 283 | 0 | M9+ | 17 – 93 | 2E | | | | | 1710.1 |
| 1732 | I | 825 | 2 | 0 | 260 | 23 | 0 | BOTH | | PI | 9815201 | PF4521 | J | | 1732 |
| 1732.1 | V | - | - | - | - | 23 | - | M9- | 60 – 66 | H | | | J | | 1732.1 |
| 1732.2 | V | - | 2 | - | - | - | - | M9+ | 20 | H | | | J | W | 1732.2 |
| 1738 | I | 826 | 1 | 0 | 270 | 13 | 0 | BOTH | | PI | 9823657 | PF3985 | E1b1b1b1b | A | 1738 |
| 1738.1 | V | - | - | - | - | 2 | - | M9- | 8 | H | | | E1b1b1b1b | A | 1738.1 |
| 1738.2 | V | - | - | - | - | 11 | - | M9- | 37 – 38 | H | | | I2a1a1a1~ | A | 1738.2 |
| 1738.3 | V | - | 1 | - | - | - | - | M9+ | NU | H,S | | | | A,W | 1738.3 |
| 2592 | I | 822 | 5 | 0 | 283 | 0 | 0 | M9+ | | PI | 15342222 | V2895.1 | R1a1a1b1a3a3a | A,W | 2592 |
| 2592.1 | V | - | 2 | 0 | - | - | - | M9+ | 55 | H | | | R1a1a1b1a3a3a | A,W | 2592.1 |
| 2592.2 | V | - | 2 | 0 | - | - | - | M9+ | 14 | H | | | R1a1a1b1a3a3a | A | 2592.2 |
| 2592.3 | V | - | 1 | 0 | - | - | - | M9+ | NU | H,S | | | R1a1a1b1a3a3a | A,W | 2592.3 |
| 2675 | I | 796 | 31 | 0 | 272 | 11 | 0 | BOTH | | PI | 15491978 | FGC22105 | C2a1a1a1a1a1~ | W | 2675 |
| 2675.1 | V | - | - | - | - | 10 | - | M9- | 44 – 45 | H | | | C2a1a1a1a1a1~ | W | 2675.1 |
| 2675.2 | V | - | - | - | - | 1 | - | M9- | NU | H,S | | | C2a1a1a1a1a1~ | W | 2675.2 |
| 2675.3 | V | - | 29 | - | - | - | - | M9+ | 4 – 16 | H,2E | | | C2a1a1a1a1a1~ | W | 2675.3 |
| 2675.4 | V | - | 1 | - | - | - | - | M9+ | NU | H,S | | | C2a1a1a1a1a1~ | W | 2675.4 |
| 2675.5 | V | - | 1 | - | - | - | - | M9+ | 49 | H,S | | | C2a1a1a1a1a1~ | W | 2675.5 |
| 2776 | I | 826 | 1 | 0 | 272 | 11 | 0 | BOTH | | PI | 15652249 | PF4009 | I2a1a1a~ | | 2776 |
| 2776.1 | V | - | - | - | - | 11 | - | M9- | 37 – 38 | H | | | I2a1a1a~ | | 2776.1 |
| 2776.2 | V | - | 1 | - | - | - | - | M9+ | NU | H,S | | | I2a1a1a~ | W | 2776.2 |
| 2843 | I | 826 | 1 | 0 | 82 | 201 | 0 | BOTH | | PI | 15801167 | ZS2268 | I | | 2843 |
| 2843.1 | V | - | - | - | - | 201 | - | M9- | 22 – 59 | H | | | I | | 2843.1 |
| 2843.2 | V | - | 1 | - | - | - | - | M9+ | 8 | H,S | | | I | W | 2843.2 |
| 3282 | I | 824 | 3 | 0 | 252 | 31 | 0 | BOTH | | PI | 16940903 | | | | 3282 |
| 3282.1 | V | - | - | - | - | 30 | - | M9- | 2 – 7 | H | | | | | 3282.1 |
| 3282.2 | V | - | - | - | - | 1 | - | M9- | NU | H,S | | | | | 3282.2 |
| 3282.3 | V | - | 2 | - | - | - | - | M9+ | 69 | H | | | | | 3282.3 |
| 3282.4 | V | - | 1 | - | - | - | - | M9+ | 92 | H,S | | | | | 3282.4 |
| 3336 | I | 824 | 3 | 0 | 272 | 11 | 0 | BOTH | | PI | 17106445 | | | | 3336 |
| 3336.1 | V | - | - | - | - | 11 | - | M9- | 37 – 38 | H | | | | | 3336.1 |
| 3336.2 | V | - | 1 | - | - | - | - | M9+ | NU | H,S | | | | | 3336.2 |
| 3336.3 | V | - | 1 | - | - | - | - | M9+ | 92 | H,S | | | | | 3336.3 |
| 3336.4 | V | - | 1 | - | - | - | - | M9+ | NU | H,S | | | | | 3336.4 |
| 3384 | I | 825 | 2 | 0 | 281 | 2 | 0 | BOTH | | PI | 17239322 | | | | 3384 |
| 3384.1 | V | - | - | - | - | 2 | - | M9- | 50 | H | | | | | 3384.1 |
| 3384.2 | V | - | 1 | - | - | - | - | M9+ | 94 | H,S | | | | | 3384.2 |
| 3384.3 | V | - | 1 | - | - | - | - | M9+ | NU | H,S | | | | | 3384.3 |
| 3395 | I | 792 | 35 | 0 | 1 | 282 | 0 | BOTH | | PI | 17281258 | Y1271 | R1b1a1b | W | 3395 |
| 3395.1 | V | - | - | - | 1 | - | - | M9- | NU | H,S | | | R1b1a1b | W | 3395.1 |
| 3395.2 | V | 792 | - | - | - | - | - | M9+ | 17 – 93 | H | | | R1b1a1b | | 3395.2 |
| 3661 | I | 823 | 4 | 0 | 281 | 2 | 0 | BOTH | | PI | 17861975 | | | | 3661 |
| 3661.1 | V | - | - | - | - | 2 | - | M9- | 5 | H | | | | | 3661.1 |
| 3661.2 | V | - | 2 | - | - | - | - | M9+ | 52 | H | | | | | 3661.2 |
| 3661.3 | V | - | 1 | - | - | - | - | M9+ | 3 | H,S | | | | | 3661.3 |
| 3661.4 | V | - | 1 | - | - | - | - | M9+ | 1 | H,S | | | | | 3661.4 |
| 4014 | I | 819 | 8 | 0 | 270 | 13 | 0 | BOTH | | PI | 18861458 | S738^^ | R1b1a1b1a1a2c1a1f1 | W | 4014 |
| 4014.1 | V | - | - | - | - | 8 | - | M9- | 61 – 64 | H,1E | | | R1b1a1b1a1a2c1a1f1 | W | 4014.1 |
| 4014.2 | V | - | - | - | - | 4 | - | M9- | 59 | H | | | R1b1a1b1a1a2c1a1f1 | W | 4014.2 |
| 4014.3 | V | - | - | - | - | 1 | - | M9- | NU | H,S | | | R1b1a1b1a1a2c1a1f1 | W | 4014.3 |
| 4014.4 | V | - | 7 | - | - | - | - | M9+ | 77 | H | | | R1b1a1b1a1a2c1a1f1 | | 4014.4 |
| 4014.5 | V | - | 1 | - | - | - | - | M9+ | 2 | H,S | | | R1b1a1b1a1a2c1a1f1 | W | 4014.5 |

| | | M9+ | | | M9- | | | | | | Build 37 | | ISOGG | | |
|---|---|---|---|---|---|---|---|---|---|---|---|---|---|---|---|
| List | Stage | 0 | 1 | . | 0 | 1 | . | M9? | Clade(s) | Res. | Position | Name | Subgroup Name | Amb. | List |
| 4259 | I | 790 | 37 | 0 | 283 | 0 | 0 | M9+ | | PI | 19372386 | | | | 4259 |
| 4259.1 | V | - | 6 | - | - | - | - | M9+ | 22 | H | | | | | 4259.1 |
| 4259.2 | V | - | 31 | - | - | - | - | M9+ | 4 – 16 | H | | | | | 4259.2 |
| 4260 | I | 826 | 1 | 0 | 187 | 96 | 0 | BOTH | | PI | 19372700 | Z2838.2 | I1 | A | 4260 |
| 4260.1 | V | - | - | - | - | 96 | - | M9- | 22 – 36 | H | | | I1 | A | 4260.1 |
| 4260.2 | V | - | 1 | - | - | - | - | M9+ | 2 | H,S | | | I1 | A,W | 4260.2 |
| 4768 | I | 826 | 1 | 0 | 82 | 201 | 0 | BOTH | | PI | 21862684 | | | | 4768 |
| 4768.1 | V | - | - | - | - | 201 | - | M9- | 22 – 59 | H | | | | | 4768.1 |
| 4768.2 | V | - | 1 | - | - | - | - | M9+ | NU | H,S | | | | | 4768.2 |
| 5164 | I | 826 | 1 | 0 | 187 | 96 | 0 | BOTH | | PI | 22750951 | P203.2 | I1 | A | 5164 |
| 5164.1 | V | - | - | - | - | 96 | - | M9- | 22 – 36 | H | | | I1 | A | 5164.1 |
| 5164.2 | V | - | 1 | - | - | - | - | M9+ | 1 | H,S | | | I1 | A,W | 5164.2 |
| 5545 | I | 826 | 1 | 0 | 246 | 37 | 0 | BOTH | | PI | 23729951 | | | | 5545 |
| 5545.1 | V | - | - | - | - | 37 | - | M9- | 1 – 10 | H | | | | | 5545.1 |
| 5545.2 | V | - | 1 | - | - | - | - | M9+ | NU | H,S | | | | | 5545.2 |
| 888 | II | 824 | 3 | 0 | 283 | 0 | 0 | M9+ | | PI | 7556207 | M1741 | O | | 888 |
| 888.1 | V | - | 2 | - | - | - | - | M9+ | 1 – 2 | H | | | O | | 888.1 |
| 888.2 | V | - | 1 | - | - | - | - | M9+ | NU | H,S | | | O | W | 888.2 |
| 1481 | II | 826 | 1 | 0 | 281 | 2 | 0 | BOTH | | PI | 8599422 | F1338 | G2a2b2b1a1 | | 1481 |
| 1481.1 | V | - | - | - | - | 2 | - | M9- | 20 | H | | | G2a2b2b1a1 | | 1481.1 |
| 1481.2 | V | - | 1 | - | - | - | - | M9+ | 72 | H,S | | | G2a2b2b1a1 | W | 1481.2 |
| 3363 | II | 826 | 1 | 0 | 281 | 2 | 0 | BOTH | | PI | 17190078 | PF2372 | E1b1b1b1 | | 3363 |
| 3363.1 | V | - | - | - | - | 2 | - | M9- | 8 | H | | | E1b1b1b1 | | 3363.1 |
| 3363.2 | V | - | 1 | - | - | - | - | M9+ | NU | H,S | | | E1b1b1b1 | W | 3363.2 |
| 268 | III | 824 | 2 | 1 | 247 | 36 | 0 | BOTH | | PI | 4009014 | | | | 268 |
| 268.1 | V | - | - | - | - | 36 | - | M9- | 2 – 10 | H | | | | | 268.1 |
| 268.2 | V | - | 2 | - | - | - | - | M9+ | 51 | H | | | | | 268.2 |
| 477 | III | 824 | 3 | 0 | 1 | 245 | 37 | BOTH | | PI | 6670461 | L58 | N1a1a1a1a1a2a1~ | W | 477 |
| 477.1 | V | - | - | - | 1 | - | - | M9- | NU | H,S | | | N1a1a1a1a1a2a1~ | W | 477.1 |
| 477.2 | V | 824 | - | - | - | - | - | M9+ | 3 – 93 | H | | | N1a1a1a1a1a2a1~ | W | 477.2 |
| 509 | III | 822 | 3 | 2 | 281 | 0 | 2 | M9+ | | PI | 6740034 | M8963.2 | I2a1b1a2b1a2a3a1~ | W | 509 |
| 509.1 | V | - | 1 | - | - | - | - | M9+ | NU | H,S | | | I2a1b1a2b1a2a3a1~ | W | 509.1 |
| 509.2 | V | - | 1 | - | - | - | - | M9+ | 49 | H,S | | | I2a1b1a2b1a2a3a1~ | W | 509.2 |
| 509.3 | V | - | 1 | - | - | - | - | M9+ | NU | H,S | | | I2a1b1a2b1a2a3a1~ | W | 509.3 |
| 1043 | III | 823 | 1 | 3 | 274 | 8 | 1 | BOTH | | PI | 7811520 | Z12228.3 | I1a3b2~ | A,W | 1043 |
| 1043.1 | V | - | - | - | - | 6 | - | M9- | 12 | H | | | I1a3b2~ | A,W | 1043.1 |
| 1043.2 | V | - | - | - | - | 1 | - | M9- | NU | H,S | | | I1a3b2~ | A,W | 1043.2 |
| 1043.3 | V | - | - | - | - | 1 | - | M9- | NU | H,S | | | I1a3b2~ | A,W | 1043.3 |
| 1043.4 | V | - | 1 | - | - | - | - | M9+ | NU | H,S | | | I1a3b2~ | A,W | 1043.4 |
| 1138 | III | 823 | 1 | 3 | 267 | 13 | 3 | BOTH | | PI | 8032311 | Z526.2 | J2b | A | 1138 |
| 1138.1 | V | - | - | - | - | 13 | - | M9- | 65 – 66 | H | | | J2b | A | 1138.1 |
| 1138.2 | V | - | 1 | - | - | - | - | M9+ | 94 | H,S | | | T1a1 | A | 1138.2 |
| 1164 | III | 825 | 0 | 2 | 187 | 95 | 1 | M9- | | PI | 8097201 | Z2891 | I1a~ | | 1164 |
| 1164.1 | V | - | - | - | - | 95 | - | M9- | 22 – 36 | 1E | | | I1a~ | | 1164.1 |
| 1386 | III | 782 | 4 | 41 | 249 | 22 | 12 | BOTH | | PI | 8481036 | | | | 1386 |
| 1386.1 | V | - | - | - | - | 21 | - | M9- | 11 – 20 | H | | | | | 1386.1 |
| 1386.2 | V | - | - | - | - | 1 | - | M9- | 58 | H,S | | | | | 1386.2 |
| 1386.3 | V | - | 4 | - | - | - | - | M9+ | BAD | BAD | | | | | 1386.3 |
| 1665 | III | 791 | 32 | 4 | 187 | 96 | 0 | BOTH | | PI | 9142914 | | | | 1665 |
| 1665.1 | V | - | - | - | - | 96 | - | M9- | 22 – 36 | H | | | | | 1665.1 |
| 1665.2 | V | - | 5 | - | - | - | - | M9+ | 48 | H | | | | | 1665.2 |
| 1665.3 | V | - | 27 | - | - | - | - | M9+ | 58 | H | | | | | 1665.3 |
| 1754 | III | 818 | 7 | 2 | 257 | 24 | 2 | BOTH | | PI | 9850653 | | | | 1754 |
| 1754.1 | V | - | - | - | - | 1 | - | M9- | 21 | H,S | | | | | 1754.1 |
| 1754.2 | V | - | - | - | - | 23 | - | M9- | 60 – 66 | H | | | | | 1754.2 |
| 1754.3 | V | - | 7 | - | - | - | - | M9+ | BAD | BAD | | | | | 1754.3 |
| 1802 | III | 818 | 1 | 8 | 269 | 11 | 3 | BOTH | | PI | 10020053 | | | | 1802 |
| 1802.1 | V | - | - | - | - | 10 | - | M9- | 66 | H | | | | | 1802.1 |
| 1802.2 | V | - | - | - | - | 1 | - | M9- | 1 | H,S | | | | | 1802.2 |
| 1802.3 | V | - | 1 | - | - | - | - | M9+ | NU | H,S | | | | | 1802.3 |
| 1836 | III | 827 | 0 | 0 | 185 | 97 | 1 | M9- | | PI | 13237053 | Z2744 | I1 | | 1836 |
| 1836.1 | V | - | - | - | - | 96 | - | M9- | 22 – 36 | H | | | I1 | | 1836.1 |
| 1836.2 | V | - | - | - | - | 1 | - | M9- | NU | H,S | | | I1 | W | 1836.2 |
| 1937 | III | 716 | 97 | 14 | 248 | 33 | 2 | BOTH | | PI | 13921953 | | | | 1937 |
| 1937.1 | V | - | - | - | - | 33 | - | M9- | BAD | BAD | | | | | 1937.1 |
| 1937.2 | V | - | 97 | - | - | - | - | M9+ | BAD | BAD | | | | | 1937.2 |

| List | Stage | M9+ 0 | M9+ 1 | M9+ . | M9- 0 | M9- 1 | M9- . | M9? | Clade(s) | Res. | Build 37 Position | Name | ISOGG Subgroup Name | Amb. | List |
|---|---|---|---|---|---|---|---|---|---|---|---|---|---|---|---|
| 1961 | III | 788 | 1 | 38 | 247 | 21 | 15 | BOTH | | PI, 1H | 14027381 | | | | 1961 |
| 1961.1 | V | - | - | - | - | 21 | - | M9- | 11 – 20 | H | | | | | 1961.1 |
| 1961.2 | V | - | 1 | - | - | - | - | M9+ | NU | H,S | | | | | 1961.2 |
| 2202 | III | 1 | 820 | 6 | 21 | 260 | 2 | BOTH | | PI | 14624254 | PF3499 | Freq. Mut. SNP in R | W | 2202 |
| 2202.1 | V | - | - | - | 21 | - | - | M9- | 11 – 20 | H | | | Freq. Mut. SNP in R | W | 2202.1 |
| 2202.2 | V | 1 | - | - | - | - | - | M9+ | NU | H,S | | | Freq. Mut. SNP in R | | 2202.2 |
| 2285 | III | 5 | 821 | 1 | 283 | 0 | 0 | M9+ | | PI | 14825721 | | | | 2285 |
| 2285.1 | V | - | 821 | 1 | - | - | - | M9+ | 3 – 93 | H, MD | | | | | 2285.1 |
| 2285.2 | V | 1 | - | - | - | - | - | M9+ | NU | H,S | | | | | 2285.2 |
| 2285.3 | V | 1 | - | - | - | - | - | M9+ | NU | H,S | | | | | 2285.3 |
| 2781 | III | 791 | 34 | 2 | 283 | 0 | 0 | M9+ | | PI | 15660495 | S1136 | R1b1a1b1a1a2c1a3a2 | | 2781 |
| 2781.1 | V | - | 33 | - | - | - | - | M9+ | 78 | H | | | R1b1a1b1a1a2c1a3a2 | | 2781.1 |
| 2781.2 | V | - | 1 | - | - | - | - | M9+ | NU | H,S | | | R1b1a1b1a1a2c1a3a2 | W | 2781.2 |
| 2902 | III | 824 | 2 | 1 | 282 | 1 | 0 | BOTH | | PI | 15984461 | | | | 2902 |
| 2902.1 | V | - | - | - | - | 1 | - | M9- | 1 | H,S | | | | | 2902.1 |
| 2902.2 | V | - | 1 | - | - | - | - | M9+ | 93 | H,S | | | | | 2902.2 |
| 2902.3 | V | - | 1 | - | - | - | - | M9+ | NU | H,S | | | | | 2902.3 |
| 3098 | III | 823 | 2 | 2 | 282 | 0 | 1 | M9+ | | PI | 16548244 | | | | 3098 |
| 3098.1 | V | - | 1 | - | - | - | - | M9+ | 65 | H,S | | | | | 3098.1 |
| 3098.2 | V | - | 1 | - | - | - | - | M9+ | 1 | H,S | | | | | 3098.2 |
| 3901 | III | 820 | 4 | 3 | 282 | 1 | 0 | BOTH | | PI | 18612789 | | | | 3901 |
| 3901.1 | V | - | - | - | - | 1 | - | M9- | NU | H,S | | | | | 3901.1 |
| 3901.2 | V | - | 1 | - | - | - | - | M9+ | NU | H,S | | | | | 3901.2 |
| 3901.3 | V | - | 2 | - | - | - | - | M9+ | 40 | H | | | | | 3901.3 |
| 3901.4 | V | - | 1 | - | - | - | - | M9+ | NU | H,S | | | | | 3901.4 |
| 3997 | III | 818 | 2 | 7 | 281 | 0 | 2 | M9+ | | PI | 18822568 | CTS9321 | O1a1a | W | 3997 |
| 3997.1 | V | - | 1 | - | - | - | - | M9+ | NU | H,S | | | O1a1a | W | 3997.1 |
| 3997.2 | V | - | 1 | - | - | - | - | M9+ | 1 | H,S | | | O1a1a | | 3997.2 |
| 4096 | III | 824 | 0 | 3 | 231 | 51 | 1 | M9- | | PI | 19044813 | | | | 4096 |
| 4096.1 | V | - | - | - | - | 21 | - | M9- | 11 – 20 | H | | | | | 4096.1 |
| 4096.2 | V | - | - | - | - | 30 | - | M9- | 2 – 7 | H | | | | | 4096.2 |
| 4126 | III | 826 | 0 | 1 | 188 | 95 | 0 | M9- | | PI | 19103066 | V3772 | I1a~ | | 4126 |
| 4126.1 | V | - | - | - | - | 95 | - | M9- | 22 – 36 | 1E | | | I1a~ | | 4126.1 |
| 4157 | III | 795 | 31 | 1 | 278 | 5 | 0 | BOTH | | PI | 19160342 | V3842 | R1a1a1~ | W | 4157 |
| 4157.1 | V | - | - | - | - | 5 | - | M9- | 32 | H | | | R1a1a1~ | W | 4157.1 |
| 4157.2 | V | - | 31 | - | - | - | - | M9+ | 4 – 16 | H | | | R1a1a1~ | | 4157.2 |
| 4541 | III | 822 | 4 | 1 | 1 | 282 | 0 | BOTH | | PI | 21499869 | | | | 4541 |
| 4541.1 | V | - | - | - | 1 | - | - | M9- | 21 | H,S | | | | | 4541.1 |
| 4541.2 | V | 822 | - | - | - | - | - | M9+ | 4 – 93 | H | | | | | 4541.2 |
| 5192 | III | 817 | 1 | 9 | 232 | 50 | 1 | BOTH | | PI | 22815525 | | | | 5192 |
| 5192.1 | V | - | - | - | - | 36 | - | M9- | 1 – 10 | H,1E | | | | | 5192.1 |
| 5192.2 | V | - | - | - | - | 13 | - | M9- | 56 | H | | | | | 5192.2 |
| 5192.3 | V | - | - | - | - | 1 | - | M9- | 58 | H,S | | | | | 5192.3 |
| 5192.4 | V | - | 1 | - | - | - | - | M9+ | NU | H,S | | | | | 5192.4 |
| 5222 | III | 790 | 36 | 1 | 184 | 99 | 0 | BOTH | | PI | 22884954 | | | | 5222 |
| 5222.1 | V | - | - | - | - | 96 | - | M9- | 22 – 36 | H | | | | | 5222.1 |
| 5222.2 | V | - | - | - | - | 3 | - | M9- | 52 | H | | | | | 5222.2 |
| 5222.3 | V | - | 3 | - | - | - | - | M9+ | 19 | H | | | | | 5222.3 |
| 5222.4 | V | - | 33 | 1 | - | - | - | M9+ | 74 | H, MD | | | | | 5222.4 |
| 5321 | III | 827 | 0 | 0 | 277 | 4 | 2 | M9- | | PI | 23080197 | Z759 | G2a2b2a1a1b1a | | 5321 |
| 5321.1 | V | - | - | - | - | 4 | 2 | M9- | 17 – 18 | MD, 1E | | | G2a2b2a1a1b1a | | 5321.1 |
| 5596 | III | 823 | 2 | 2 | 280 | 2 | 1 | BOTH | | PI | 23881461 | | | | 5596 |
| 5596.1 | V | - | - | - | - | 2 | - | M9- | 8 | H | | | | | 5596.1 |
| 5596.2 | V | - | 1 | - | - | - | - | M9+ | NU | H,S | | | | | 5596.2 |
| 5596.3 | V | - | 1 | - | - | - | - | M9+ | 3 | H,S | | | | | 5596.3 |
| 5626 | III | 826 | 1 | 0 | 280 | 2 | 1 | BOTH | | PI | 23977118 | F4036.2 | R1b1a1b1a1a2c1a4b8~ | W | 5626 |
| 5626.1 | V | - | - | - | - | 2 | - | M9- | 35 | H | | | R1b1a1b1a1a2c1a4b8~ | W | 5626.1 |
| 5626.2 | V | - | 1 | - | - | - | - | M9+ | 36 | H,S | | | R1b1a1b1a1a2c1a4b8~ | W | 5626.2 |
| 5672 | III | 825 | 0 | 2 | 248 | 35 | 0 | M9- | | PI | 24437979 | S7474 | E | | 5672 |
| 5672.1 | V | - | - | - | - | 35 | - | M9- | 1 – 10 | 2E | | | E | | 5672.1 |
| 5700 | III | 826 | 0 | 1 | 262 | 19 | 2 | M9- | | PI | 27390536 | | | | 5700 |
| 5700.1 | V | - | - | - | - | 19 | 2 | M9- | 11 – 20 | MD, 1E | | | | | 5700.1 |

**Columns:**

List -- Numerical, ordered by Build 37 Position.

Stage -- I through V

M9+/M9- (0, 1, .) -- these seven columns are the genotype data for each SNP

Note: For each SNP, the summary SNP data is given, identical to that in Supplementary Table 1.

| | | M9+ | | | | M9- | | | | | | Build 37 | | ISOGG | | |
|---|---|---|---|---|---|---|---|---|---|---|---|---|---|---|---|---|
| List | Stage | 0 | 1 | . | | 0 | 1 | . | M9? | Clade(s) | Res. | Position | Name | Subgroup Name | Amb. | List |

Then, multiple extensions, .1, .2, .3, .4 are given as necessary with the specific genotype numbers pertinent to each recognized clade (see next 3 columns JKL).

M9? -- as in Supplementary Table 1, M9- are clades splitting off prior to M9; M9+ are those after M9. BOTH, in the first line summary for each SNP, indicates that variation on both sides of M9 was evaluated for potential clades.

Clades -- as in Supplementary Table 1, simple numerical designation of terminal branches or clades, essentially following alphabetical order given by inferred ISOGG Subgroup name. 1 -- 66 for M9- Clades, 1 -- 94 for M9+ Clades.

Note: to identify those singleton calls that were NOT supported by additional data (i.e., criteria C1 and/or C2 in Methods), NU is given instead of a clade number. For 3 SNPs, at least a portion of the data were deemed BAD and these SNPs excluded.

Res. -- PI SNP Resolution: PI (phylogenetically inconsistent) in each SNP's summary line, followed by one or more of: H (homoplasy), 1E or 2E (1 or 2 genotyping errors predicted), S (flag for singleton calls),

MD = missing relevent data, BAD (as above in Clades column).

Build 37 Position -- basepair position of SNP according to HG Build 37 (suppressed in extensions).

Name -- SNP's identifier (suppressed in extensions).

ISOGG Subgroup Name -- Subgroup Name assigned each SNP, when available

Amb. -- ISOGG label ambiguity: A for multiple distinct ISOGG labels available, W for wrong for our data (see Supplementary Table 3).

List -- Numerical, ordered by Build 37 Position.  Should be identical to first column.
