## Supplementary material for "Characterization of the Y Chromosome in Newfoundland and Labrador: Evidence of a Founder Effect": Table S4

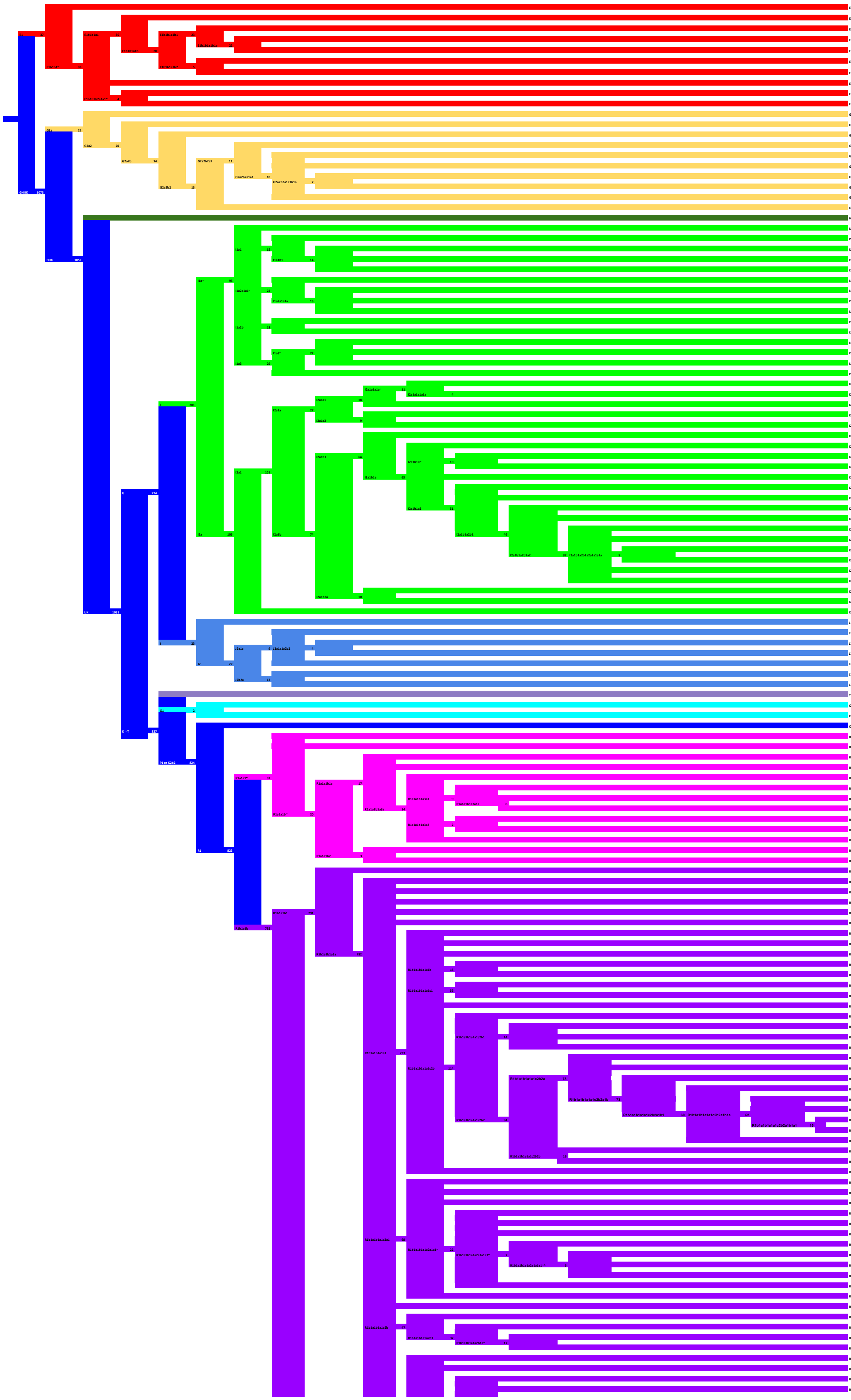

| ISOGG | Count | I9+/M9- | Clade |
|---|---|---|---|
| E1a2a1 | 1 | M9- | 1 |
| E1b1b1a1a1 | 2 | M9- | 2 |
| E1b1b1a1b1 | 1 | M9- | 3 |
| E1b1b1a1b1a | 20 | M9- | 4 |
| E1b1b1a1b1a^ | 2 | M9- | 5 |
| E1b1b1a1b2 | 3 | M9- | 6 |
| E1b1b1a1b2^ | 2 | M9- | 7 |
| E1b1b1b1b | 2 | M9- | 8 |
| E1b1b1b2a1a1~ | 3 | M9- | 9 |
| E1b1b1b2a1a4 | 1 | M9- | 10 |
| G2a1a1a1a1 | 1 | M9- | 11 |
| G2a2a1a2a1a | 6 | M9- | 12 |
| G2a2b1 | 1 | M9- | 13 |
| G2a2b2a1 | 1 | M9- | 14 |
| G2a2b2a1a1 | 1 | M9- | 15 |
| G2a2b2a1a1a1a1 | 1 | M9- | 16 |
| G2a2b2a1a1b1a | 3 | M9- | 17 |
| G2a2b2a1a1b1a1a2a1 | 4 | M9- | 18 |
| G2a2b2a1a1c2 | 1 | M9- | 19 |
| G2a2b2b1a1 | 2 | M9- | 20 |
| H1a1a4b2 | 1 | M9- | 21 |
| I1a~ | 12 | M9- | 22 |
| I1a1 | 1 | M9- | 23 |
| I1a1b1 | 10 | M9- | 24 |
| I1a1b1a1 | 2 | M9- | 25 |
| I1a1b1a4a | 2 | M9- | 26 |
| I1a2a1a1~ | 7 | M9- | 27 |
| I1a2a1a1a | 3 | M9- | 28 |
| I1a2a1a1a1 | 7 | M9- | 29 |
| I1a2a1a1a2 | 5 | M9- | 30 |
| I1a2b | 13 | M9- | 31 |
| I1a2b^ | 5 | M9- | 32 |
| I1a3~ | 18 | M9- | 33 |
| I1a3~^ | 2 | M9- | 34 |
| I1a3~^^ | 2 | M9- | 35 |
| I1a3a | 7 | M9- | 36 |
| I2a1a1a1a~ | 7 | M9- | 37 |
| I2a1a1a1a1a | 4 | M9- | 38 |
| I2a1a1b1a~ | 7 | M9- | 39 |
| I2a1a2a | 6 | M9- | 40 |
| I2a1a2b1a1a1 | 3 | M9- | 41 |
| I2a1b1 | 1 | M9- | 42 |
| I2a1b1a | 1 | M9- | 43 |
| I2a1b1a~ | 4 | M9- | 44 |
| I2a1b1a1a1a | 6 | M9- | 45 |
| I2a1b1a1b1a1a | 1 | M9- | 46 |
| I2a1b1a2 | 3 | M9- | 47 |
| I2a1b1a2^ | 2 | M9- | 48 |
| I2a1b1a2b1 | 12 | M9- | 49 |
| I2a1b1a2b1^ | 2 | M9- | 50 |
| I2a1b1a2b1a2 | 8 | M9- | 51 |
| I2a1b1a2b1a2^ | 3 | M9- | 52 |
| I2a1b1a2b1a2a1a1a1a | 3 | M9- | 53 |
| I2a1b1a2b1a2a1a1a1a1a | 2 | M9- | 54 |
| I2a1b1a2b1a2a2 | 3 | M9- | 55 |
| I2a1b1a2b1a2e2a3~ | 13 | M9- | 56 |
| I2a1b2a | 9 | M9- | 57 |
| I2a1b2a^ | 1 | M9- | 58 |
| I2a2 | 4 | M9- | 59 |
| J1a | 1 | M9- | 60 |
| J2a1a | 4 | M9- | 61 |
| J2a1a1a2b2 | 2 | M9- | 62 |
| J2a1a1a2b2a1a | 2 | M9- | 63 |
| J2a1a4b | 1 | M9- | 64 |
| J2b2a | 3 | M9- | 65 |
| J2b2a1a1a1a1a | 10 | M9- | 66 |
| T1a1a1b2b2b1a1a | 1 | M9+ | 94 |
| O1a1a1 | 1 | M9+ | 1 |
| O1b1a1a1a1a1 | 1 | M9+ | 2 |
| Q2a1a1a1 | 1 | M9+ | 3 |
| R1a1a1~ | 9 | M9+ | 4 |
| R1a1a1~^ | 2 | M9+ | 5 |
| R1a1a1b1a1a | 1 | M9+ | 6 |
| R1a1a1b1a2b3a | 2 | M9+ | 7 |
| R1a1a1b1a3a^ | 1 | M9+ | 8 |
| R1a1a1b1a3a1 | 3 | M9+ | 9 |
| R1a1a1b1a3a1a | 4 | M9+ | 10 |
| R1a1a1b1a3a1a^ | 2 | M9+ | 11 |
| R1a1a1b1a3a2 | 1 | M9+ | 12 |
| R1a1a1b1a3a2a1 | 1 | M9+ | 13 |
| R1a1a1b1a3a3a | 2 | M9+ | 14 |
| R1a1a1b2^ | 2 | M9+ | 15 |
| R1a1a1b2a~ | 1 | M9+ | 16 |
| R1b1a1b1 | 9 | M9+ | 17 |
| R1b1a1b1a1a | 65 | M9+ | 18 |
| R1b1a1b1a1a^ | 3 | M9+ | 19 |
| R1b1a1b1a1a^^ | 2 | M9+ | 20 |
| R1b1a1b1a1a^^^ | 4 | M9+ | 21 |
| R1b1a1b1a1a^^^^ | 6 | M9+ | 22 |
| R1b1a1b1a1a1 | 6 | M9+ | 23 |
| R1b1a1b1a1a1^ | 6 | M9+ | 24 |
| R1b1a1b1a1a1a | 6 | M9+ | 25 |
| R1b1a1b1a1a1b | 9 | M9+ | 26 |
| R1b1a1b1a1a1b1a | 7 | M9+ | 27 |
| R1b1a1b1a1a1c1 | 30 | M9+ | 28 |
| R1b1a1b1a1a1c1a | 26 | M9+ | 29 |
| R1b1a1b1a1a1c2a1 | 18 | M9+ | 30 |
| R1b1a1b1a1a1c2b | 6 | M9+ | 31 |
| R1b1a1b1a1a1c2b1a1a1 | 1 | M9+ | 32 |
| R1b1a1b1a1a1c2b1b | 12 | M9+ | 33 |
| R1b1a1b1a1a1c2b1b2a1 | 1 | M9+ | 34 |
| R1b1a1b1a1a1c2b2a | 4 | M9+ | 35 |
| R1b1a1b1a1a1c2b2a^ | 1 | M9+ | 36 |
| R1b1a1b1a1a1c2b2a1b | 13 | M9+ | 37 |
| R1b1a1b1a1a1c2b2a1b1 | 2 | M9+ | 38 |
| R1b1a1b1a1a1c2b2a1b1a | 35 | M9+ | 39 |
| R1b1a1b1a1a1c2b2a1b1a^ | 2 | M9+ | 40 |
| R1b1a1b1a1a1c2b2a1b1a1 | 4 | M9+ | 41 |
| R1b1a1b1a1a1c2b2a1b1a1a | 11 | M9+ | 42 |
| R1b1a1b1a1a1c2b2a1b1b1 | 6 | M9+ | 43 |
| R1b1a1b1a1a1c2b2b | 4 | M9+ | 44 |
| R1b1a1b1a1a1c2b2b1a~ | 12 | M9+ | 45 |
| R1b1a1b1a1a1c3 | 1 | M9+ | 46 |
| R1b1a1b1a1a2a1 | 13 | M9+ | 47 |
| R1b1a1b1a1a2a1^ | 5 | M9+ | 48 |
| R1b1a1b1a1a2a1^^ | 1 | M9+ | 49 |
| R1b1a1b1a1a2a1a1~ | 10 | M9+ | 50 |
| R1b1a1b1a1a2a1a1~^ | 2 | M9+ | 51 |
| R1b1a1b1a1a2a1a1~^^ | 2 | M9+ | 52 |
| R1b1a1b1a1a2a1a1a1~ | 1 | M9+ | 53 |
| R1b1a1b1a1a2a1a1a1~^ | 3 | M9+ | 54 |
| R1b1a1b1a1a2a1a1a1~^^ | 2 | M9+ | 55 |
| R1b1a1b1a1a2a1a1a1a | 1 | M9+ | 56 |
| R1b1a1b1a1a2a1a1a2 | 1 | M9+ | 57 |
| R1b1a1b1a1a2a1b1a1~ | 27 | M9+ | 58 |
| R1b1a1b1a1a2a5~ | 3 | M9+ | 59 |
| R1b1a1b1a1a2b | 10 | M9+ | 60 |
| R1b1a1b1a1a2b1 | 20 | M9+ | 61 |
| R1b1a1b1a1a2b1a~ | 7 | M9+ | 62 |
| R1b1a1b1a1a2b1a1~ | 10 | M9+ | 63 |
| R1b1a1b1a1a2c1 | 4 | M9+ | 64 |
| R1b1a1b1a1a2c1^ | 1 | M9+ | 65 |
| R1b1a1b1a1a2c1a | 112 | M9+ | 66 |
| R1b1a1b1a1a2c1a^ | 3 | M9+ | 67 |

105

31

479

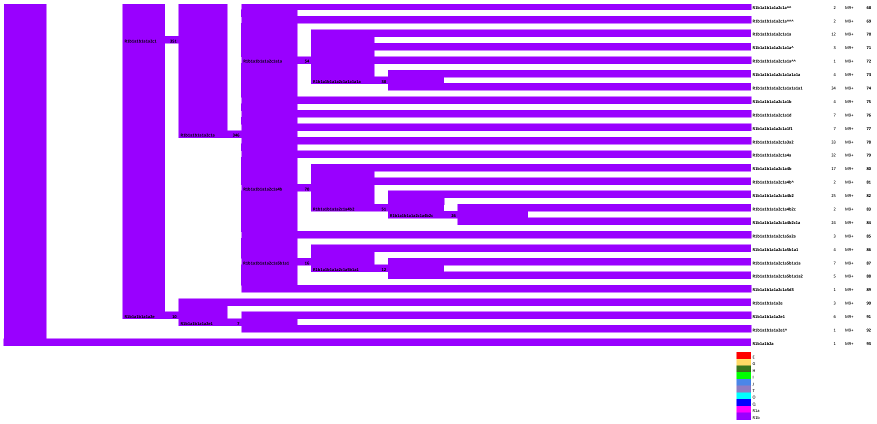
R1b1a1b1a1a2c1a^^ 2 M9+ 68
R1b1a1b1a1a2c1a^^^ 2 M9+ 69
R1b1a1b1a1a2c1a1a 12 M9+ 70
R1b1a1b1a1a2c1a1a^ 3 M9+ 71
R1b1a1b1a1a2c1a1a^^ 1 M9+ 72
R1b1a1b1a1a2c1a1a1a1a 4 M9+ 73
R1b1a1b1a1a2c1a1a1a1a1 34 M9+ 74
R1b1a1b1a1a2c1a1b 4 M9+ 75
R1b1a1b1a1a2c1a1d 7 M9+ 76
R1b1a1b1a1a2c1a1f1 7 M9+ 77
R1b1a1b1a1a2c1a3a2 33 M9+ 78
R1b1a1b1a1a2c1a4a 32 M9+ 79
R1b1a1b1a1a2c1a4b 17 M9+ 80
R1b1a1b1a1a2c1a4b^ 2 M9+ 81
R1b1a1b1a1a2c1a4b2 25 M9+ 82
R1b1a1b1a1a2c1a4b2c 2 M9+ 83
R1b1a1b1a1a2c1a4b2c1a 24 M9+ 84
R1b1a1b1a1a2c1a5a2a 3 M9+ 85
R1b1a1b1a1a2c1a5b1a1 4 M9+ 86
R1b1a1b1a1a2c1a5b1a1a 7 M9+ 87
R1b1a1b1a1a2c1a5b1a1a2 5 M9+ 88
R1b1a1b1a1a2c1a5d3 1 M9+ 89
R1b1a1b1a1a2e 3 M9+ 90
R1b1a1b1a1a2e1 6 M9+ 91
R1b1a1b1a1a2e1^ 1 M9+ 92
R1b1a1b2a 1 M9+ 93
E
G
H
I
J
T
O
Q
R1a
R1b
